# Zinc finger protein SALL4 functions through an AT-rich motif to regulate gene expression

**DOI:** 10.1101/2020.07.03.186783

**Authors:** Nikki R. Kong, Mahmoud A. Bassal, Hong Kee Tan, Jesse V. Kurland, Kol Jia Yong, John J. Young, Yang Yang, Fudong Li, Jonathan Lee, Yue Liu, Chan-Shuo Wu, Alicia Stein, Hongbo Luo, Leslie E. Silberstein, Martha L. Bulyk, Daniel G. Tenen, Li Chai

## Abstract

The zinc finger transcription factor SALL4 is highly expressed in embryonic stem cells, down-regulated in most adult tissues, but reactivated in many aggressive cancers. This unique expression pattern makes SALL4 an attractive target for designing therapeutic strategies. However, whether SALL4 binds DNA directly to regulate gene expression is unclear and many of its targets in cancer cells remain elusive. Here, through an unbiased screen of protein binding microarray (PBM) and Cleavage Under Targets and Release Using Nuclease (CUT&RUN) experiments, we identified and validated the DNA binding domain of SALL4 and its consensus binding sequence. Combined with RNA-seq analyses after SALL4 knockdown, we discovered hundreds of new SALL4 target genes that it directly regulates in aggressive liver cancer cells, including genes encoding a family of Histone 3 Lysine 9-specific Demethylases (KDMs). Taken together, these results elucidated the mechanism of SALL4 DNA binding and revealed novel pathways and molecules to target in SALL4-dependent tumors.

## Introduction

SALL4 is a nuclear factor which plays an important role in murine and human embryonic development (Elling et al., 2006; Sakaki-Yumoto et al., 2006; Zhang et al., 2006). Normally, SALL4 is down-regulated in most adult tissues except germ cells (Chan et al., 2017; Hobbs et al., 2012; Yamaguchi et al., 2015) and hematopoietic stem cells (Gao et al., 2013b). However, it is dysregulated in hematopoietic pre-leukemias and leukemias, including high-grade myelodysplastic syndrome (MDS) and acute myeloid leukemia (AML) (Gao et al., 2013a; Ma et al., 2006). SALL4 is also reactivated in a significant fraction of almost all solid tumors, including lung cancer, endometrial cancer, germ cell tumors, and hepatocellular carcinomas (Li et al., 2013a; Miettinen et al., 2014; Yong et al., 2013; 2016). This unique expression pattern demonstrates that SALL4 can be a potential link between pluripotency and cancer, and thus targeted therapeutically with limited side effects in adult normal tissues. Accordingly, SALL4-positive liver cancers share a similar gene expression signature to that of fetal liver tissues, and are associated with a more aggressive cancer phenotype, drug resistance, and worse patient survival (Oikawa et al., 2013; Yong et al., 2013).

Despite its important roles in pluripotency and association with certain types of cancers, it is still unclear how SALL4 functions as a transcription factor. SALL4 has two main isoforms, the full length SALL4A and a spliced variant, SALL4B (Tatetsu et al., 2016). SALL4A has four zinc finger clusters (ZFC), three of which contain either a pair or a trio of C2H2-type zinc fingers, which are though to confer nucleic acid-binding activity (Al-Baradie et al., 2002). However, these clusters are scattered throughout the linear polypeptide sequence and it is not known which ZFC of SALL4 is responsible for DNA binding. Demonstrating their functional importance, SALL4 ZFC with either missense or frameshift mutations are frequently found in patients with Okihiro Syndrome (Borozdin et al., 2004; Kohlhase et al., 2003; Miertus et al., 2006; Terhal et al., 2006), which is proposed to be a result of impaired zinc finger function. Furthermore, thalidomide-mediated SALL4 degradation depends on its zinc finger amino acid sequences that show species-specific selectivity (Donovan et al., 2018; Matyskiela et al., 2018). Despite evidence of their functional importance, it is not known whether SALL4 binds DNA through its ZFCs directly or if so, which ZFC is responsible for binding. It is also unclear what consensus sequence SALL4 prefers. Finally, while SALL4 has been shown to function as a transcriptional repressor by recruiting the Nucleosome remodeling and histone deacetylase complex (NuRD) (Gao et al., 2013a; Lu et al., 2009; Yong et al., 2013), many of its target genes and downstream pathways have yet to be elucidated. Its association with the NuRD complex has led to the hypothesis that SALL4 may play a role in global chromatin regulation. However, its direct involvement with heterochromatin or euchromatin has yet to be determined (Böhm et al., 2007; Kim et al., 2017; Sathyan et al., 2011).

Here, we have used an unbiased screen to discover that SALL4 binds AT-rich motif through its C-terminal ZFC. These result were further confirmed using a recently developed method of targeted in-situ genome wide profiling (CUT&RUN) (Skene et al., 2018) to identify true SALL4 binding sites in liver cancer cells. These experiments, coupled with RNA-seq after SALL4 KD, allowed us to unveil SALL4’s transcriptional regulation of a family of Hisone 3 lysine 9-specific demethylases (KDM3/4), through which it can regulate the chromatin landscape in cancer cells. Understanding its mechanism as a transcription factor can thus provide new insight of how SALL4-dependent pathways can be targeted in therapeutic approaches.

## Results

### SALL4 binds DNA through an AT-rich motif

In order to identify the SALL4 consensus binding sequence(s), we used the universal protein binding microarray (PBM) technology (Berger et al., 2006) to conduct an unbiased analysis of all possible DNA sequences to which purified SALL4 binds. Using this method, we discovered that SALL4 prefers to bind an AT-rich sequence with little degeneracy: AA[A/T]TAT[T/G][A/G][T/A] (Figure 1A), in which the WTATB in the center of the motif represents the core sequence. In addition, this sequence is highly specific compared to other AT-rich sequences on the array (Supplementary Figure 1A), and the control FLAG peptide alone does not bind this sequence (Supplementary Figure 1B). Along its linear polypeptide sequence, full length SALL4 protein (SALL4A) has three C2H2-type zinc finger clusters (ZFC 2-4) either in pairs or in a triplet, as well as one C2HC-type zinc finger. Zinc finger motifs are frequently associated with nucleic acid binding (Struhl, 1989), however, it is not known which zinc fingers of SALL4 are responsible for its DNA binding activity. Therefore, we deleted two of the clusters individually and generated SALL4A mutants that lack either their ZFC2 or ZFC4 domains, hereafter referred to as AΔZFC2 and AΔZFC4, respectively, and repeated the PBM experiments. Interestingly, the AΔZFC2 mutant was unaffected compared to wild type (WT) SALL4A protein in DNA binding specificity, but the AΔZFC4 mutant was unable to bind the AT-rich consensus motif (Figure 1B, C), suggesting that ZFC4 is responsible for sequence recognition by the protein. SALL4 also has a shorter isoform resulting from alternative splicing, SALL4B, which shares only the ZFC4 domain with SALL4A. Supporting our finding that ZFC4 is the DNA sequence recognition domain of SALL4, we confirmed that WT SALL4B also binds the specific AT-rich motif, but SALL4BΔZFC4 mutant lacking this domain does not (Supplementary Figure 1C, D). To validate our PBM results, we performed electrophoretic mobility shift assays (EMSA) with two randomly picked oligonucleotides on the PBM chip, both containing the AT-rich consensus binding site. These assays demonstrated that SALL4 could shift biotinylated oligonucleotides containing the WT AT-rich sequence but not those with the probe sequence randomly scrambled (Figure 1D, Supplementary Figure 2A, and Supplementary Table 1). Furthermore, anti-FLAG or anti-SALL4 antibodies were able to diminish or super-shift the signal, the latter through binding to and slowing down the electrophoretic mobility of the SALL4-DNA complex, while mouse IgG isotype control could not (Figure 1E, and Figure 2A lanes 4-7 compared to lane 2). This demonstrated that the binding event was highly specific to SALL4, and could not be attributed to any other proteins co-purified with SALL4.

**Figure 1.**
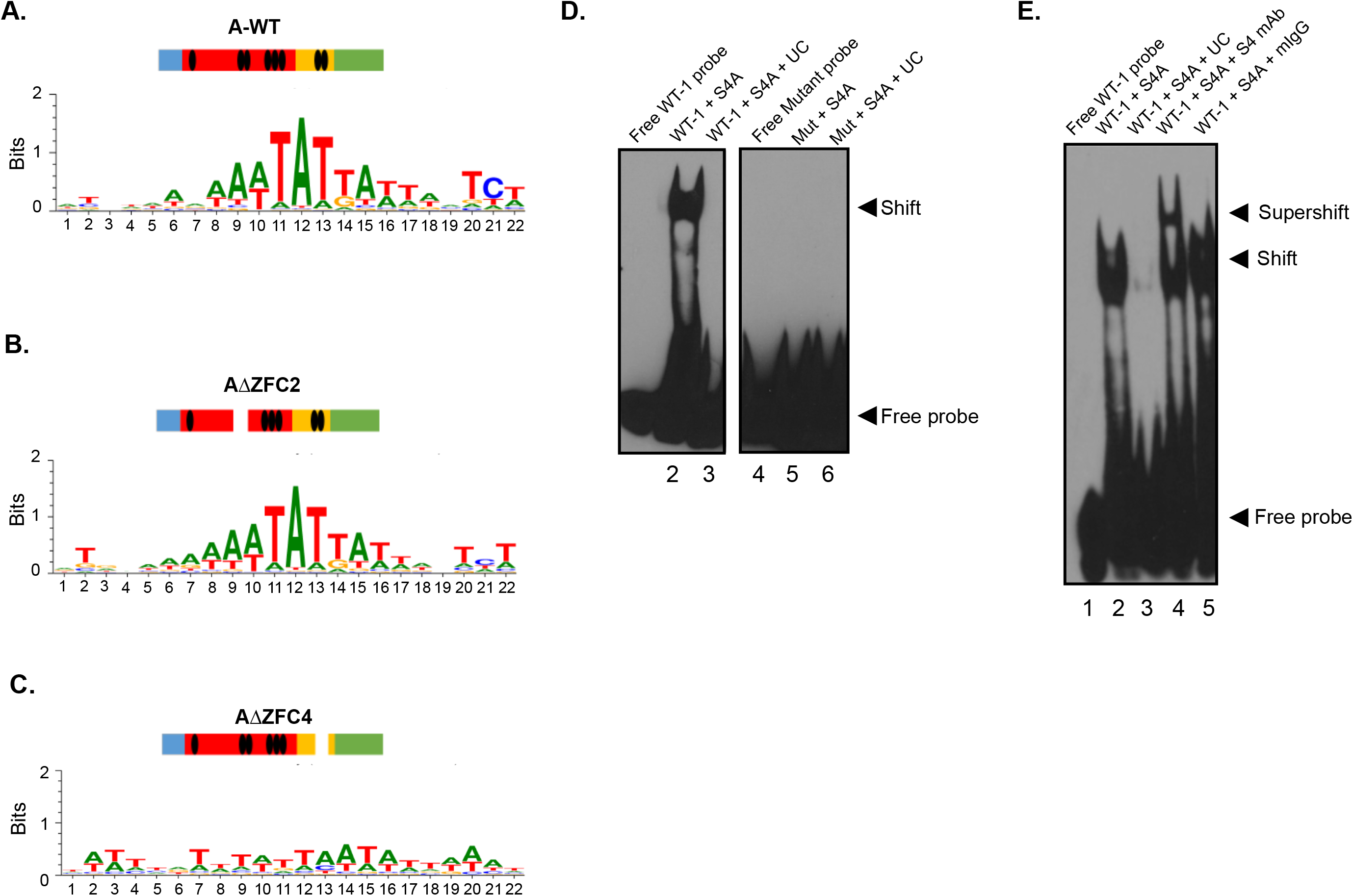
Discovery of a novel SALL4 DNA binding motif. **A)** Specific DNA sequence motif bound by SALL4A in universal PBM assays (Berger et al., 2006).The color bar above the position weighted matrix (PWM) indicates the linear structure of SALL4 and the 4 zinc finger clusters are denoted by black ovals. **B)** DNA sequences bound by SALL4A mutant lacking zinc finger cluster 2 (AΔZFC2). **C)** DNA sequences bound by SALL4A mutant lacking zinc finger cluster 4 (AΔZFC4). **D)** EMSA showing SALL4A shifts the AT-rich motifcontaining oligos (lanes 1-3) but not when the motif is scrambled (lanes 4-6, gel cut for clarity); S4A=SALL4A, UC=unlabeled WT-1 probe **E)** SALL4-DNA complex is super-shifted by a SALL4-specific monoclonal antibody (lane 4) but not by mouse IgG isotype control (lane 5); S4 mAb=mouse SALL4 antibdy (Santa Cruz EE-30), mIgG=mouse isotype control IgG. All EMSA reactions contain poly dI:dC competitor to reduce background binding.

**Figure 2.**
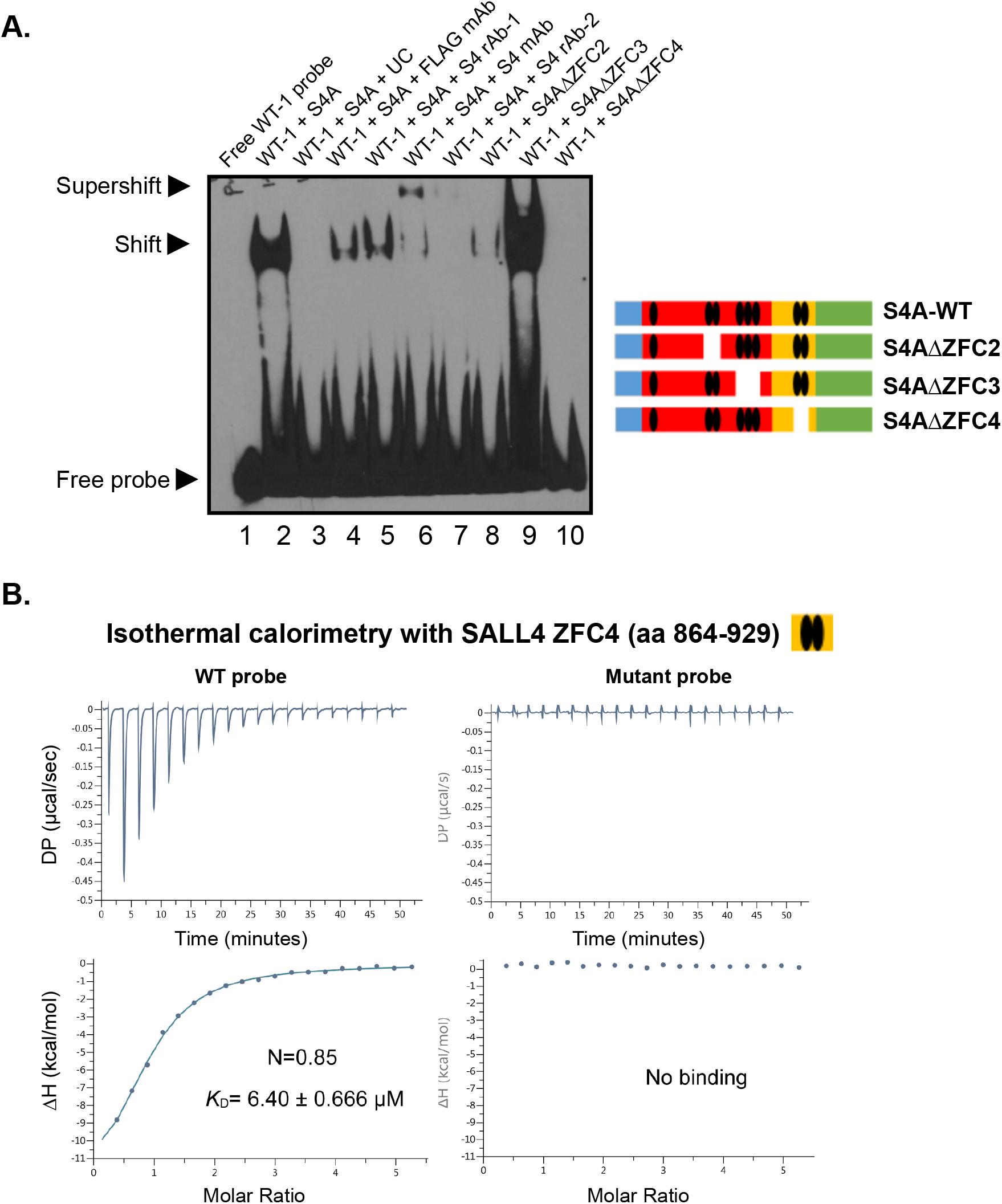
SALL4 binds DNA through its fourth zinc finger cluster. **A)** EMSA showing that SALL4-DNA complex was reduced or super-shifted in the presence of FLAG or SALL4 antibodies (1ug pre-incubated with proteins before the addition of DNA probes); S4A=SALL4A, UC=unlabeled WT-1 probe, S4 rAb-1=SALL4 rabbit antibody (Cell Signaling D16H12), S4 rAb-2=SALL4 rabbit antibody (Abcam ab57577). Lanes 8-10 in the right panel show that the AΔZFC3 mutant binds to the same WT sequence, whereas binding by AΔZFC2 and AΔZFC4 is abrogated. Sequences for the oligonucleotides used for EMSA can be found in Supplementary Table 1, all EMSA reactions contain poly dI:dC competitor. **B)** Isothermal titration calorimetry (ITC) experiments showing purified SALL4 ZFC4 (amino acids 864-929) binds DNA oligos containing the WTATB motif (left, WT sequence: GAGTTATTAATG) and not when the motif was mutated (right, mutant sequence: GAGTCGCTAATG).

Next, we performed EMSA experiments with AΔZFC2 and AΔZFC4 mutants described above along with a SALL4A mutant lacking ZFC3 (refers to as AΔZFC3). While AΔZFC3 can still bind WT oligonucleotides (Figure 2A, compare lane 2 to lane 9), suggesting ZFC3 is not involved in DNA binding, the AΔZFC4 mutant was unable to bind the oligos (Figure 2A, compare lane 2 to lane 10), suggesting that deleting ZFC4 completely abrogated SALL4 DNA binding ability. In addition, deletion of AΔZFC2 had impaired DNA binding, suggesting that ZFC2 contributes to the ability of ZFC4 to bind to DNA in the context of full length SALL4 (Figure 2A, compare lane 2 to lane 8). These results were consistent with our observation that the smaller SALL4B isoform lacking ZFC2 and ZFC3 appears to bind DNA less strongly than the longer A isoform (Supplementary Figure 2B, compare lanes 6-9 for SALL4B to lanes 2-5 for SALL4A).

To further validate the SALL4 DNA binding domain, we performed isothermal titration calorimetry (ITC) experiments with purified ZFC4 domain of SALL4 with either WT probes containing the AT-rich motif or mutated probes with only the core motif changed (Supplementary Table 1). ITC experiments demonstrated that while SALL4’s C-terminal ZFC4 domain can bind WT probes with Kd of 6.4μM for probe 1 and 6.88μM for probe2, it cannot bind mutated probes (Figure 2B, Supplementary Figure 3). Results from these experiments supported our PBM and EMSA findings of a specific SALL4 motif that is AT-rich, as well as the importance of SALL ZFC4 domain in DNA sequence recognition and binding.

### The SALL4 motif is enriched in CUT&RUN binding experiments

Given the *in vitro* results demonstrating that SALL4 binds to an AT-rich DNA motif, we sought to determine if it binds a similar motif in cells. Previously, we have performed SALL4 ChIP-seq experiments in human cells and found they were challenging because SALL4 is located in the chromatin fractions which can be difficult to sonicate (Supplementary Figure 4A). Further, our previous method of interrogating the chromatin fraction still required cross-linking, while it did not generate many SALL4 peaks or yield a SALL4 motif (Liu et al., 2018a). Here, we took advantage of the availability of a highly specific antibody against human SALL4 (Cell Signaling Technology, clone D16H12, lot 2) and performed the Cleavage Under Targets and Release Using Nuclease (CUT&RUN) assay (Skene et al., 2018), which is an in-situ profiling of protein/DNA binding that eliminates the cross-linking step while generates reads with low background and more precise localization.

We chose SNU398 liver cancer cells to identify endogenous SALL4 binding sites genome-wide because i) these cells have high SALL4 expression compared to other cancer cells (Supplementary Figure 4B) and SALL4 ChIP data are available in these cells (Liu et al., 2018a); ii) SALL4 is required for the viability of a large fraction of hepatocellular carcinoma cells (Yong et al., 2013), and iii) SALL4 serves as a biomarker for worse prognosis in liver cancer (Oikawa et al., 2013; Yong et al., 2013).

Three separate CUT&RUN experiments using SNU398 liver cancer cell nuclei revealed SALL4 binds over 11,200 common peaks genome-wide at least two-fold above isotype control IgG peak enrichment (representative track shown in Figure 3A, Supplementary Table 2). Furthermore, SALL4 peaks were distributed 33.8% intergenic, 30% intronic, and 21% within the proximal promoter (Figure 3B) and can be annotated near 4,364 genes. On average, the reads are about 80-100bp long, allowing for better identification of SALL4 binding motif. Consequently, when *de novo* motif search was performed on SALL4 peaks, we observed a significant enrichment of the AT-rich motif with the core WTATB motif that was independently identified from PBM experiments (Figure 3C and Supplementary Figure 4C). This motif was present in similar percentages of peaks in all three SALL4 CUT&RUN replicates through both *de no* and direct searches using Motif 2, which is a shortened Motif 1 (Figure 3D).

**Figure 3.**
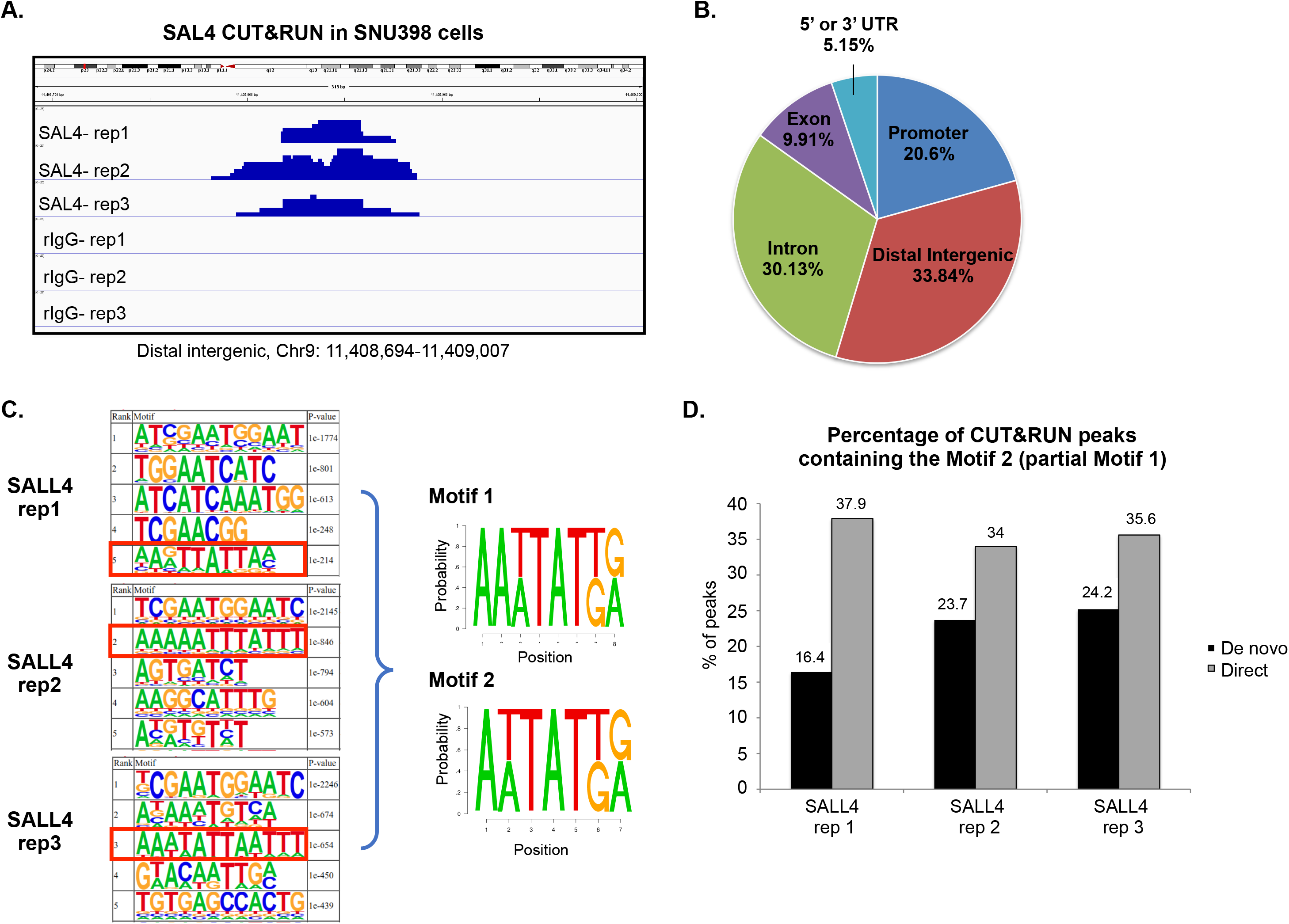
SALL4 CUT&RUN showing its binding genome-wide in liver cancer cells. **A)** Representative genomic tracks of three SALL4 CUT&RUN replicates and their isotype rabbit IgG control experiments. Scale is 0-25 **B)** The genomic distribution of SALL4 CUT&RUN peaks from three biological replicates (rep). **C)** The top five HOMER motifs from *de novo* analysis of three CUT&RUN replicates with their respective p-values, resulting in the two composite motifs shown on right. **D)** Bar graph showing the percentage of peaks containing Motif 2 in *de novo* (black bars) and direct (gray bars) from three SALL4 CUT&RUN replicates.

### Discovery of new SALL4 gene targets in liver cancer cells

The SALL4 target genes in liver cancer that are associated with transcriptional and chromatin regulation are not well defined. Furthermore, it has been shown that SALL4 can act like a transcriptional activator and/or repressor dependent on the cellular context (Gao et al., 2013a; Li et al., 2013b; Lu et al., 2009; Ma et al., 2006; Young et al., 2014; Zhang et al., 2006). To understand how SALL4 DNA binding affects the expression of its downstream target genes, we performed RNA-seq at 72 hours after SALL4 knock-down (KD) (Supplementary Figure 5A and Supplementary Table 3) in biological triplicates. We then compared the RNA-seq results to our CUT&RUN peaks and found that among 2,696 significantly differentially expressed genes (red circles in Figure 4A, Supplementary Table 4), 430 genes had SALL4 peaks (totaling 1,192 peaks, Figure 4B, Supplementary Table 5), suggesting they were directly regulated by SALL4. When *de novo* motif search was performed on the peaks, the core WTATB motif was among the top hits (Supplementary Figure 5B).

**Figure 4.**
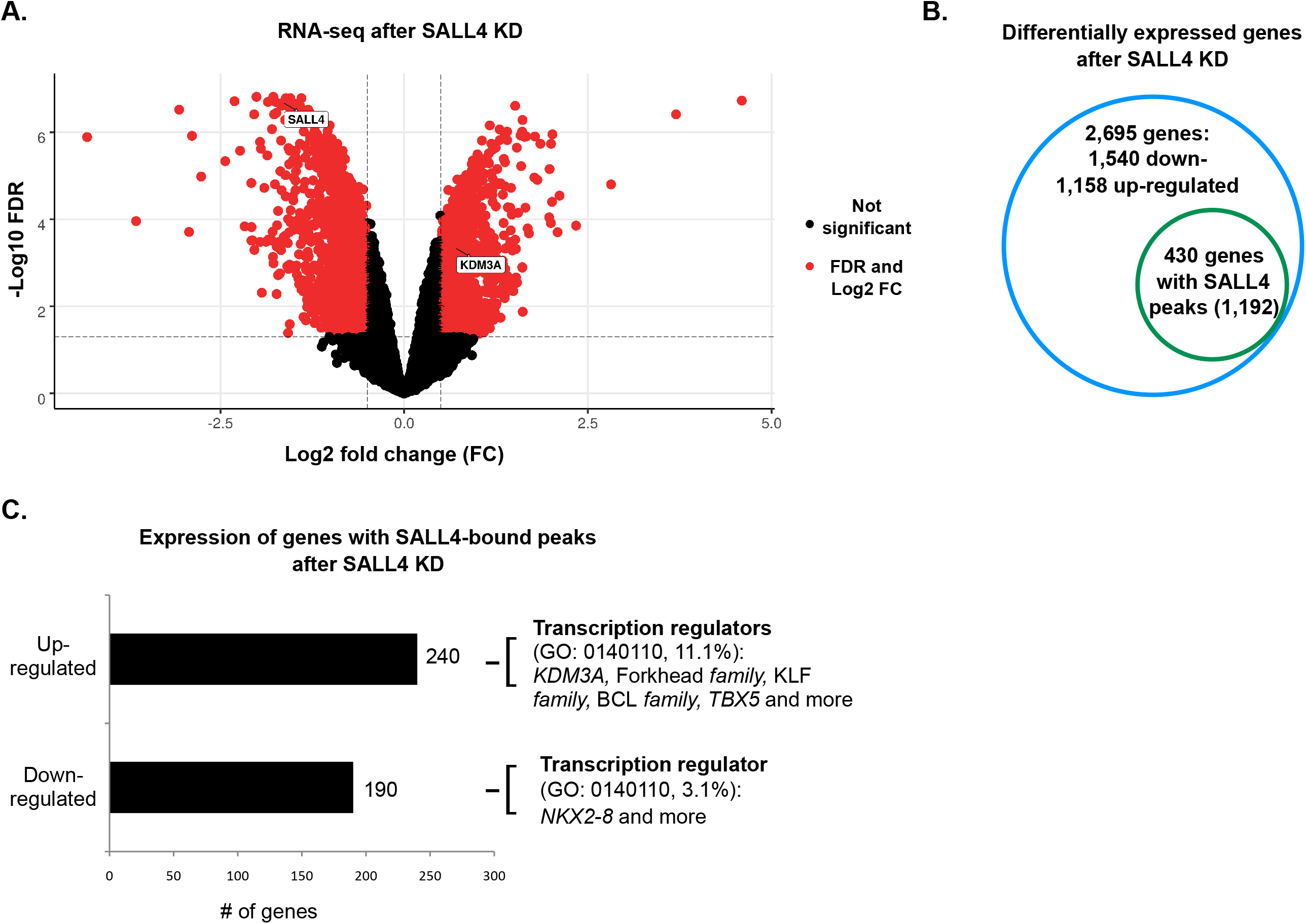
RNA-seq data revealed direct SALL4 target genes. **A)** Volcano plot showing genes that are down- or up-regulated significantly after SALL4 KD with log2 fold change (FC) represented on the x-axis; red circles denote differentially expressed genes with FDR <0.05; experiment performed in biological triplicates with shSALL4-2 compared to scrambled control shRNA (SCR) 72 hours after transduction. **B)** Number of differentially expressed genes after SALL4 KD as well as those containing SALL4 peaks (Supplementary Table 4 and 5). **C)** Bar graph representing number of up- and down-regulated genes after SALL4 KD (Supplementary Table 5), and their gene ontology (GO) molecular pathway analysis (www.pantherdb.org) focusing on GO 0140110.

For genes with SALL4 CUT&RUN peaks and are differentially expressed after KD, 240 were up-regulated (repressed by SALL4) and 190 were down-regulated (activated by SALL4). When gene ontology analysis was performed, we found that the most differentially represented molecular pathway that was repressed by SALL4 was transcription regulation, accounting for 11.1 % of the up-regulated genes after SALL4 KD, compared to 3.1 % of down-regulated genes after SALL4 KD (Figure 4C). These categories of SALL4-repressed genes included those encoding chromatin modifiers such as *KDM3A* and several family of transcription factors such as Forkhead (*FOXA1, FOXO1*), BCL (*BCL11A, BCL11B*, and *BCL6*), and KLF (*KLF10, KLF12*), as well as *TBX5*. Many of these targets were not previously reported in liver cancer cells(Liu et al., 2018a), which demonstrates the sensitivity of the CUT&RUN technique. Taken together, these genome-wide binding assays have confirmed the bona fide SALL4 motif we found *in vitro*.

### SALL4 regulates the expression of histone demethylases

SALL4 can interact with the HDAC-containing nucleosome remodeling and deacetylase (NuRD) complex (Lu et al., 2009) (Supplementary Figure 5C), and we have previously shown that blocking SALL4’s transcriptional repressive function by interrupting its interaction with NuRD was an effective therapeutic approach in liver cancer cells (Gao et al., 2013a; Liu et al., 2018a). Furthermore, SALL4 has been shown to localize in and regulate chromatin in cells (Böhm et al., 2007; Chan et al., 2017; Hobbs et al., 2012; Kim et al., 2017; Xiong et al., 2016). Therefore, we focused on chromatin-associated genes that were up-regulated after SALL4 KD, as well as potential direct targets identified by CUT&RUN. One of the up-regulated genes *KDM3A*, encoding a Histone 3 Lysine 9 (H3K9)-specific Demethlyase (Gray et al., 2005), as well as several other members of the KDM demethylase family (Figure 5A, Supplementary Figure 6A, B). We validated the RNA-seq data by performing qPCR after SALL4 KD (Figure 5B), and we validated the binding by ChIP-qPCR with primers targeting the SALL4 binding site at the *KDM3A* promoter (Supplementary Figure 6C, and Supplementary Table 6). Then we used the *KDM3A* peak to design probes for EMSA experiments. We found that SALL4A binds double-stranded oligonucleotides containing the SALL4 motif, but not when this motif was mutated (Figure 5C, Supplementary Table 1).

**Figure 5.**
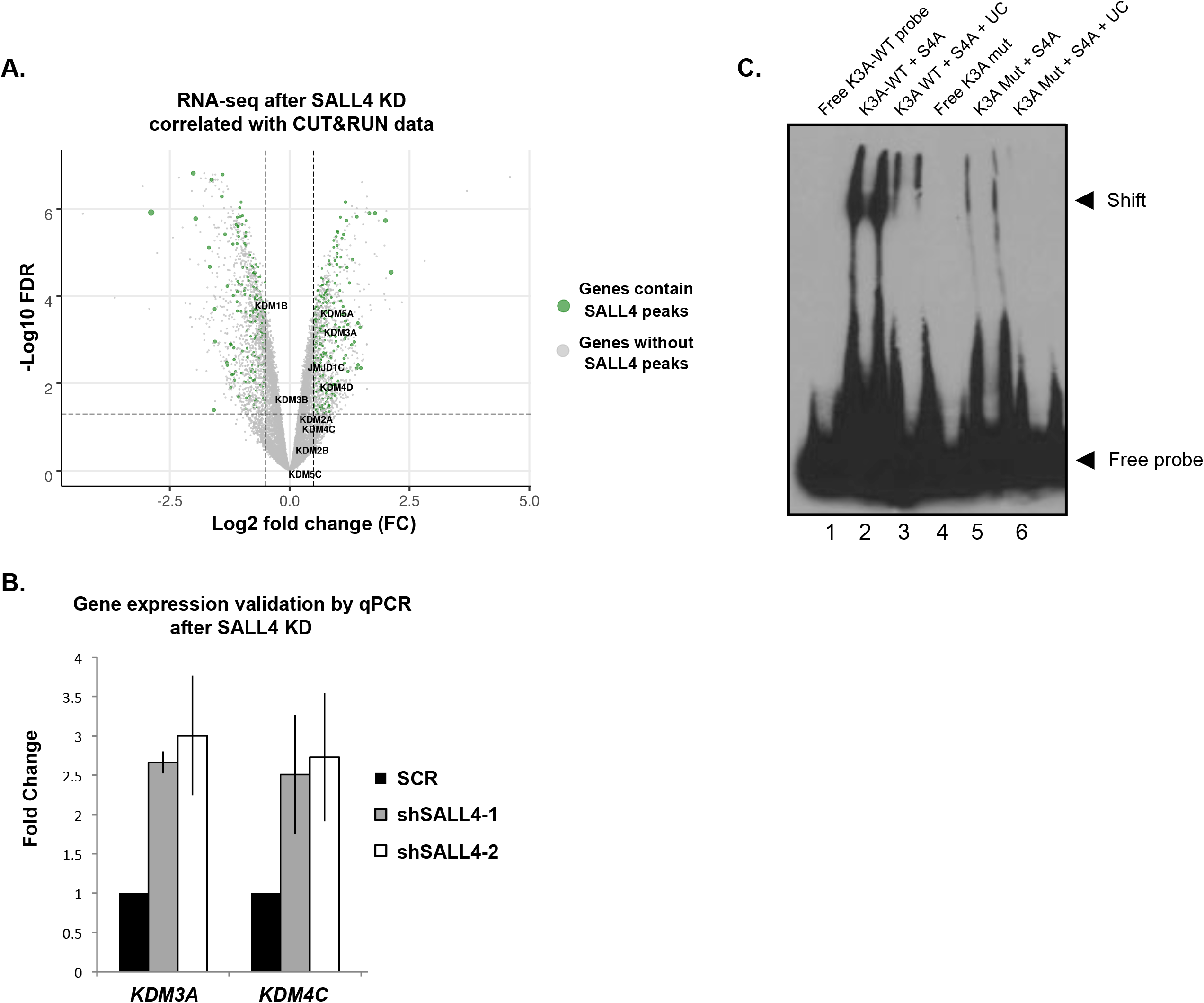
SALL4 regulates histone lysine demethylases. **A)** Volcano plot from Figure 4A with genes encoding KDM proteins labeled. Green circles denote genes containing SALL4 CUT&RUN peaks, the size of the circles corresponds to log_2_FC in expression **B)** Quantitative real-time PCR (qPCR) analysis of three SALL4 direct targets 40 hours after SALL4 KD, results are summarized from either three or four independent experiments (primer sequences found in Supplementary Table 3). **C)** EMSA showing that SALL4 binds *KDM3A* promoter region containing the AT-rich motif (WT sequence: TCTTCATTTATCCTTCAAAA), but not the mutated sequence (TCTTCTTTTAACCTTCAAAA).

In order to confirm the importance of SALL4 binding on *KDM3A* gene expression, we further used CRISPR/Cas9 to disrupt one of the SALL4 motifs in its binding site on the *KDM3A* promoter (targeted region can be found in Supplementary Table 3). After sorting cells that had successful CRISPR targeting through GFP expression on the lentiviral vector, we collected mRNA and performed qPCR. We observed a two-fold increase in expression of *KDM3A* in cells with deleted SALL4 binding site compared to mock-transfected control cells (Supplementary Figure 6D).

Our combined analyses of RNA-seq and CUT&RUN data, in addition to EMSA and gene editing experiments, demonstrated that SALL4 directly regulates a subset of chromatin modifying genes in cancer cells, raising the possibility that SALL4 could regulate the global chromatin landscape of cells.

### SALL4 and heterochromatin

It has been shown previously that SALL4 KD in liver cancer cells led to cell death (Yong et al., 2013), and we confirmed that cells could not survive after prolonged loss of SALL4 expression via KD by two different shRNAs (Supplementary Figure 7). However, at an earlier time point (40 hours post-viral transduction), SALL4 KD was highly efficient (Figure 7A, and detected by GFP expression from the shRNA lentiviral plasmid), yet no significant cell death was observed, as GFP and DAPI double positive cells did not increase until 72 hours post-transduction (Figure 7B). This observation allows a window to assess of true SALL4-dependent cellular functions with fewer potential secondary effects.

Because of SALL4’s known association with the NuRD complex and its regulation of *KDM3/4* pathway (Figure 5), we sought to ascertain whether there were any changes in the global chromatin landscape. We first confirmed our RNA-seq data by showing that KDM3A protein was up-regulated upon SALL44 KD at the early time point of 40hr after transduction (Figure 6A). Given that KDM3A is known to demethylate H3K9me2/3 (Gray et al., 2005), we found that while total histone 3 levels were unchanged, there was a marked reduction of global H3K9me2/3 levels after SALL4 KD (Figure 6A).

**Figure 6.**
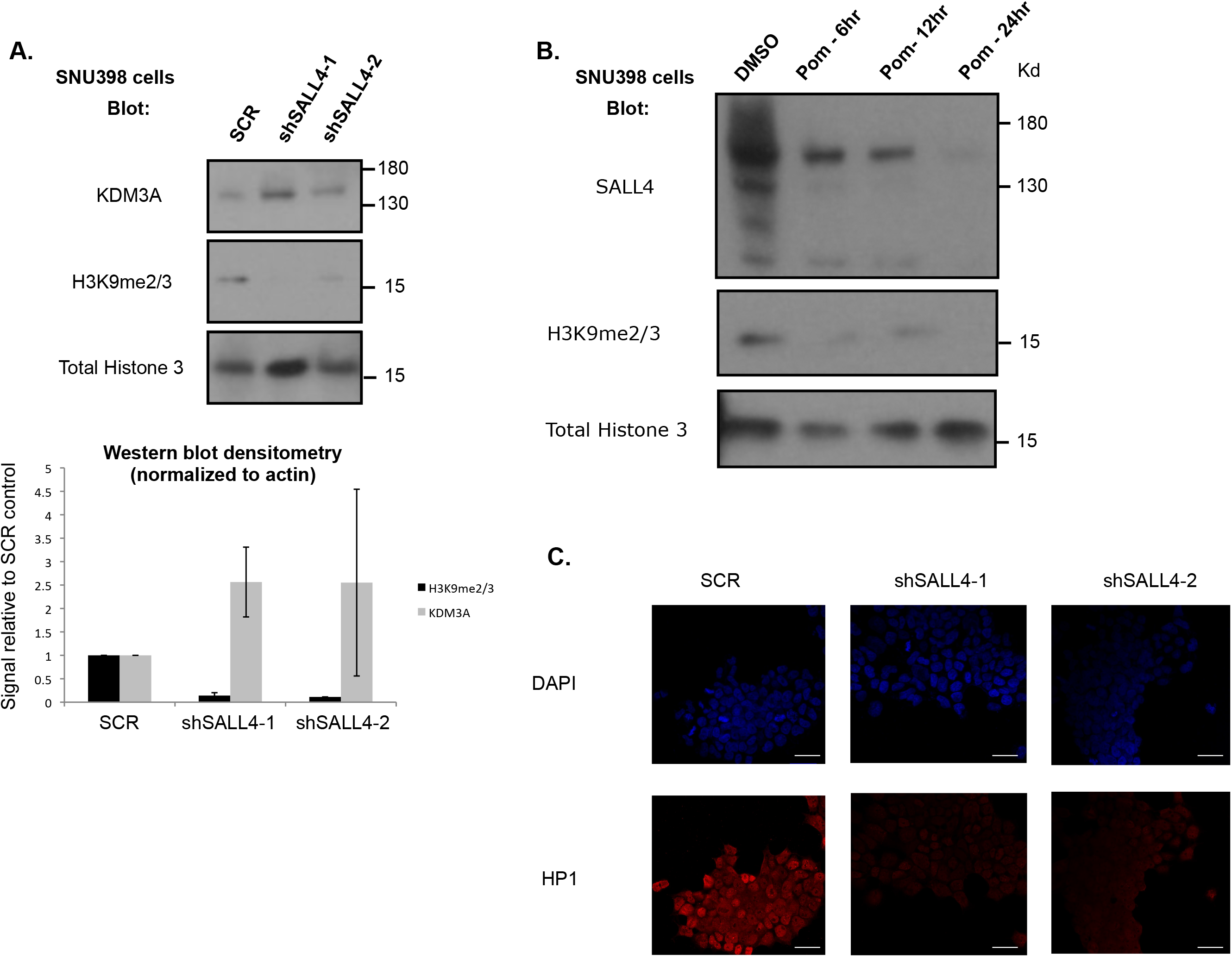
SALL4 and heterochromatin. **A)** Western blotting of SNU398 liver cancer cell lysates at 40hr after SALL4 KD, gel cut for clarity, molecular weight marker is indicated on the right. Bottom: summary of western blot densitometry from two separate SALL4 KD experiments of H3K9me2/3 protein expression (black bars) and KDM3A expression (gray bars). **B)** Western blotting of SNU398 cell lysates collected after either control DMSO or pomalidomide (Pom, 10μM) treatments at the indicated time points. **C)** Immunofluorescence staining of HP1 of SNU398 liver cancer cells transduced with SCR control, or two shRNAs against SALL4; white scale bar denotes 33μm.

In addition to shRNA-mediated kD of SALL4 expression, we utilized a second approach to pharmacologically delete SALL4. It has been shown that as a neo-substrate of the E3 ligase CRBN, SALL4 can be induced to degrade via treatment with immunomodulatory drugs such as thalidomide (Donovan et al., 2018; Matyskiela et al., 2018). Therefore, we treated SNU398 liver cancer cells with a thalidomide analog, pomalidomide, for 6, 12, or 24 hours and collected protein lysates. We observed robust SALL4 degradation as soon as 6-12 hours after treatment, at which points H3K9me2/3 marks were already diminished significantly, while total histone 3 levels remained unchanged (Figure 6B). We further confirmed SALL4’s importance in heterochromatin regulation by performing immunofluoresence for HP1α protein after SALL4 KD. HP1 binds H3K9me2/3 and is a hallmark of heterochromatin (Zeng et al., 2010). Again, using two shRNAs targeting SALL4 (Supplementary Figure 8), we observed significant reduction of HP1 staining in the nuclei of KD cells compared to SCR control shRNA-targeted cells (Figure 6C).

## Discussion

Although SALL4 has been referred to as a transcriptional regulator, biochemical evidence of direct DNA binding has been scant. Its zinc finger clusters have been individually tested in EMSA assays and shown to preferentially bind 5-hydroxymethylcytosine, and thus SALL4 was proposed to stabilize Tet2-DNA interaction during DNA demethylation in murine ESCs (Xiong et al., 2016). However, the role of SALL4 DNA binding to regulate specific gene expression is still unclear. Here, we discovered an AT-rich SALL4 binding motif via PBM assays and confirmed this finding via EMSA and ITC assays. In addition, through biochemical approaches we demonstrated that the fourth ZFC of SALL4 is responsible for DNA recognition and binding. Our DNA motif was further supported by CUT&RUN assays, which resulted in much smaller DNA fragments than previous ChIP-seq experiments, allowing for better *de novo* motif discovery. In all, we utilized three separate biochemical and two cellular assays to demonstrate SALL4 binds a unique AT-rich motif through its fourth zinc finger cluster.

Our novel CUT&RUN results in cancer cells, coupled with RNA-seq data from SALL4 KD in the same cells, identified hundreds of genes that SALL4 directly regulates. Of note, many of these targets are involved in transcriptional regulation or chromatin modification. We found that SALL4 binds and represses members of the *KDM* family of genes, resulting in changes in the methylation status of H3K9 and chromatin as assessed by staining with HP1. Our identification of the link between SALL4 KD and decreased heterchromatin marks presents a previously undescribed mechanism by which SALL4 acts as a regulator of global chromatin landscape. One of the few adult tissues where SALL4 is expressed is in the spermatogonial progenitor cells, where it antagonizes PLZF transcription factor function to drive differentiation (Hobbs et al., 2012). Interestingly, KDM3A has been shown to be highly expressed in post-meiotic male germ cells to control transcription of protamine in spermatids (Okada et al., 2007). It remains to be seen whether SALL4 directly represses KDM3A in the testes, which would suggest that the SALL4-KDM3A pathway discovered here is applicable to other progenitor cell tissues. KDM3A has also been shown to be important in maintaining self-renewal property of ESCs (Kuroki et al., 2018). Therefore, the SALL4/KDM pathway in embryonic development, spermatogenesis, and adult cancer should be examined more closely, wherein SALL4 expression is potentially important to prevent premature over-expression of KDM3A. An unbiased shRNA screen found that KDM3A promotes a form of epithelial cell apoptosis, anoikis, through activating pro-apoptotic *BNIP3* genes (Pedanou et al., 2016).Therefore, SALL4 regulation of the KDM pathway may contribute to its regulation of both heterochromatin and cell death.

Altogether, these results suggest that SALL4 regulates gene expression and chromatin in two ways: i) by directly recruiting HDAC-containing NuRD complex to deacetylate histones; and ii) by indirectly affecting heterochromatin levels through transcriptional repression of KDM demethylases of H3K9me2/3. Therefore, SALL4 is part of a transcriptional regulatory network in liver cancer cells, where it is required for both cell survival and maintenance of the heterochromatin status at certain important target genes. These findings contribute to further understanding of SALL4 function in cancer cells and the underlying molecular mechanisms, thereby uncovering novel therapeutic targets in SALL4-positive cancers.

## STAR Methods

### Lead contact and materials availability

Further information and requests for resources such as recombinant DNA plasmids generated in this study should be directed to and will be fulfilled by the Lead Contact, Daniel G. Tenen

### Experimental model and subject details

SNU398 and SNU387 hepatocellular carcinoma cells (ATCC) were cultured in RPMI media with 10% fetal bovine serum (FBS, Gibco). HEK293T and HeLa cells were cultured in DMEM media with 10% FBS.

### Method Details

#### Protein binding microarray (PBM)

SALL4 proteins were purified using M2 FLAG agarose beads (Sigma) from nuclear extracts of HEK293T cells that were transfected with FLAG-tagged SALL4. “Universal” all 10-mer double-stranded oligonucleotide arrays in 8 x 60K, GSE format (Agilent Technologies; AMADID #030236) were used to perform PBM experiments following previously described experimental protocols (Berger and Bulyk, 2009; Berger et al., 2006). Each SALL4A WT or mutant protein was assayed in PBM at 600nM. The PBM scan images were obtained using a GenePix 4000A Microarray Scanner (Molecular Devices).

#### Electrophoretic mobility shift assay

Biotinylated probes were designed based on PBM data and obtained from Integrated DNA Technology. Oligonucleotide annealing was performed by heating mixed oligonucleotides to 95 degrees for 5 minutes, and slowly cooled in a water bath (initially 70 degrees) overnight. SALL4 proteins (purified as described for PBM) were premixed with unlabeled probes or appropriate antibodies for 20 minutes at 4 degrees, then mixed with labeled probes and incubated for 20 minutes at room temperature. Binding buffers B1 and B2 containing poly dIdC blocking DNA, binding buffer C1, and stabilization buffer D (Active Motif) were used. Free DNA and protein-DNA complexes were run for 2 hours in the cold room in 6% polyacrylamide gels in tris-borate-EDTA, then transferred onto nylon membranes (0.45um pore), and visualized via streptavidin-HRP according to manufacturer’s instructions (Thermo Fisher). 1ug of antibody against SALL4 (Santa Cruz EE-30) or isotype control IgG (Santa Cruz) was pre-incubated with the proteins, before incubation with the DNA probes. All EMSA probe sequences can be found in Supplementary Table 1.

#### Isothermal titration calorimetry (ITC)

The DNA fragments encoding SALL4 ZFC4 (residues 864-929) cloned into modified pGEX-4T1 vector (GE Healthcare) with a tobacco etch virus (TEV) cleavage site after the GST tag. All the proteins were expressed in *Escherichia coli* BL21 (DE3) cells (Novagen), purified using glutathione sepharose (GE healthcare), and cleaved by TEV protease overnight at 4 degrees to remove the GST tag. The cleaved protein was further purified by size-exclusion chromatography on a HiLoad 16/60 Superdex 75 column (GE healthcare), dialyzed with buffer C (20 mM Tris-HCl (pH 7.5), 150 mM NaCl), and concentrated for subsequent experiments. ITC assays were carried out on a Microcal PEAQ-ITC instrument (Malvern) at 25 degrees. The titration protocol consisted of a single initial injection of 1 μl, followed by 19 injections of 2 μl of the protein (concentration: 0.5-1mM) into the sample cell containing double stranded DNA oligos (concentration: 20 μM).

#### Cleavage under targets and release using nucleases (CUT&RUN)

A detailed protocol can be found on protocol.io from the Henikoff lab (<u>dx.doi.org/10.17504/protocols.io.mgjc3un</u>) (Skene et al., 2018). Briefly, 2 million cell nuclei were immobilized on Concanavalin A beads after washing. SALL4 (CST D16H12) or H3K9me2/3 (Cell Signaling D4W1U) antibodies, or normal rabbit IgG (Cell Signaling DA1E) were incubated with the nuclei overnight in the presence of 0.02% digitonin at 4 degrees. The next day, 700ng/mL of proteinA-micrococcal nuclease (pA-Mnase purified in house with vector from Addgene 86973, protocol from Schmid et al.) (Schmid et al., 2004) were incubated with the nuclei at 4 degrees for an hour. After washing, the tubes were placed in heat blocks on ice set to 0 degrees, CaCl2 (1mM) was added and incubated for 30 minutes before 2x Stop buffer containing EDTA was added. DNA was eluted by heat and high-speed spin, then phenol-chloroform extracted. Qubit was used to quantify purified DNA and Bioanalyzer (2100) traces were run to determine the size of the cleaved products. NEBNext Ultra II DNA library prep kit (NEB E7645) was used to make the libraries according to Liu et al.’s protocol, outlined on protocols.io (<u>dx.doi.org/10.17504/protocols.io.wvgfe3w</u>) (Liu et al., 2018b). Pair-end (42bp) Illumina sequencing was performed on the bar-coded and amplified libraries.

#### Lentivirus-mediated gene expression knockdown and western blotting

Two shRNAs targeting SALL4 were previously described (Gao et al., 2013b): E5 and 507 (Supplementary Table 3); both target exon 2 of SALL4 mRNA. A pLL.3 vector containing the shRNA and GFP were transfected into HEK293T cells using TransIT-Lenti (Mirus). Viruses were collected at 48 hours and 72 hours post-transfection. After cell debris was filtered out with 45micron syringe filters, viral supernatants were spun at 20,000 RPM at 4 degrees for 2 hours, and re-suspended in RPMI media (Gibco). The viral titer was calculated by serial dilution and transduction of HeLa cells. MOI of 2 were used for these cells. Transduction was performed with polybrene (8ug/mL) and spinning at 70g at room temperature for an hour. SNU398 cells were >90% GFP positive starting at 40 hours posttransduction. At either 40 or 72 hours after transduction, cells were either counted with trypan blue exclusion method or collected and stained with DAPI nuclear stain. GFP and DAPI+ dead cells were counted using a Canto II flow cytometer (BD Biosciences). Western blotting was performed by running the collected cell lysates in a 4-20% gradient tris-glycine SDS-PAGE gel, transferred onto methylcellulose, and blotted with antibodies raised against SALL4 (Santa Cruz, clone EE30), Actin (Sigma, clone AC-74), total histone 3 (Cell Signaling Technology, clone D1H2), di/tri-methyl histone 3 lysine 9 (CST, D4W1U), tri-methyl histone 3 lysine 9 (EMD/Millipore, catalog 07-442), or KDM3A (Abcam, catalog ab80598).

#### RNA-seq

RNA was extracted by Trizol in three biological replicates 72 hours after SALL4 KD and libraries were made following manufacturer’s instructions (Illumina). Pair-end Illumina sequencing was performed on the bar-coded and amplified libraries.

#### Co-immunoprecipitation

Cells were lysed with RIPA lysis buffer and sonicated with a microtip sonicator at 90% duty, 15 bursts. The lysates were incubated with SALL4 antibody (same as ChIP) over night at 4 degrees, followed by 6hr incubation with protein A/G beads at 4 degrees. After washing, the beads were boiled in 2x SDS sample buffer containing beta-mercaptoethanol and the supernatant was separated in Tris-glycine gels. Western blotting was performed with SALL4 antibody (EE30, Santa Cruz Biotechnology) and HDAC1/2 antibodies (Cell Signaling 8349).

#### Immunofluorescence staining

40 hours after SALL4 KD with shRNA 1 and 2, SNU398 liver cancer cells were fixed with 4% paraformaldehyde in PBS for 15 minutes. Fixed cells were permeabilized with 0.1% Triton-x in PBS, washed with PBS containing 0.1% Tween-20, and blocked with 3% bovine serum albumin (BSA). Primary antibodies against HP1 (Abcam ab77256) or SALL4 (Abcam ab57577) were incubated with cells overnight at 4 degrees, in PBS containing 0.3% BSA and 0.1% Tween-20. The next day, cells were washed with Tween-20/PBS, incubated with secondary anti-goat or anti-rabbit antibodies conjugated to Alexa Fluorophore 594 at room temperature for an hour. Cells were washed and stained with DAPI DNA stain for 5 minutes and mounted with Vectashield mounting medium. Images were taken with a confocal microscope (Zeiss LSM710) with the same settings for all samples.

### Quantification and statistical analysis

#### PBM analysis

PBM image data were processed using GenePix Pro v7.2 to obtain signal intensity data for each spot. The data were then further processed by using Masliner software (v1.02) (Berger et al., 2006; Dudley et al., 2002) to combine scans from different intensity settings, increasing the effective dynamic range of the signal intensity values. If a dataset had any negative background-subtracted intensity (BSI) values (which can occur if the region surrounding a spot is brighter than the spot itself), consistent pseudocounts were added to all BSI values such that they all became nonnegative. All BSI values were normalized using the software for spatial de-trending providing in the Universal PBM Analysis Suite(Berger and Bulyk, 2009). Motifs were derived using the Seed-and-Wobble algorithm, and Enologos was used to generate logos from PWMs, as previously described (Berger and Bulyk, 2009; Berger et al., 2006).

#### ITC analysis

Data obtained from ITC assays were fitted to one-site binding model via the MicroCal PEAQ-ITC analysis software provided by the manufacturer and the oligonucleotide sequences can be found in Supplementary Table 1.

#### CUT&RUN analysis

Detailed data analysis combining Henikoff (Skene et al., 2018) and Orkin (Liu et al., 2018b) labs’ pipelines can be found on github (https://github.com/mbassalbioinformatics/CnRAP). Briefly, raw fastq files were trimmed with Trimmomatic v0.36 (Bolger et al., 2014) in pair-end mode. Next, the kseq trimmer developed by the Orkin lab was run on each fastq file. BWA (v0.7.17-r1188) (Li and Durbin, 2009) was first run in “aln” mode on a masked hg38 genome downloaded form UCSC to create *.sai files; then BWA was run in “sampe” mode with the flag “-n 20” on the *.sai files. Afterwards, Stampy (v1.0.32) (Lunter and Goodson, 2011) was in “—sensitive” mode. Next, using SAMtools (v1.5)(Li et al., 2009), bam files were sorted (“sort - | 0 –O bam”), had read pair mates fixed (“fixmate”), and indexed (“index”). Bam coverage maps were generated using bamCoverage from the deepTools suite (v2.5.7) (Ramírez et al., 2014). The same procedure was run to align fastq files to a masked *Saccharomyces Cerevisiae* v3 (sacCer3) genome for spike-in control DNA, also downloaded form UCSC. A normalization factor was determined for each hg38 aligned replicate based on the corresponding number of proper-pairs aligned to the sacCer3 genome, as recommended in the Henikoff pipeline, this was calculated as follow: normalization factor = 10,000,000#”proper_pairs’2. Next, from the hg38 aligned bam files, “proper-paired” reads were extracted using SAMtools with the output piped into Bedtools (Quinlan and Hall, 2010), producing BED files of reads that have been normalized to the number of reds aligned to the sacCer3 genome. BedGraphs of these files were generated as intermediary fiels to facilitate generation of BigWig coverage maps using the bedGraphToBigWig tool from UCSC (v4) (Kent et al., 2010). For peak calling, the recently developed SEACR (v.1.1) (Meers et al., 2019)was utilized and run in “relaxed’ mode to produce peak files as the BED files used were already normalized to the number of yeast spike-in reads. Subsequent peak file columns were re-arranged to facilitate motif discovery using HOMER (v4.10) (Heinz et al., 2010). Peaks were annotate using the R package ChIPSeeker (v.1.20.0)(Yu et al., 2015). Overlapping peak subsets within 3kb of each other were generated using mergePeaks.py from the HOMER suite (Heinz et al., 2010). Peak positions for those that are common to all three replicates and at least 2-fold above IgG control can be found in Supplementary Table 2. Heatmap of SALL4-bound genes encoding lysine demethylases was generated using the R package pheatmap (https://CRAN.R-project.org/package=pheatmap, Raivo Kolde. Pheatmap under R 3.6.2.

#### RNA-seq analysis

Raw fastq files had optical duplicates removed using clumpify form BBMap (https://sourceforge.net/projects/bbmap/). Next, adapt trimming was performed using BBDuk (from BBMap) and reads were trimmed using trimmomatic (Bolger et al., 2014). After read cleanup, reads were aligned to hg38 genome using STAR (Dobin et al., 2013). BamCoverage (Ramírez et al., 2014) maps were generated using default parameters and read distributions were calculated using read_distribution.py from the RseQC suite of tools (Wang et al., 2012). Counts tables were generated using htseq-count (Anders et al., 2015). The fold change was plotted as a volcano plot using EnhancedVolcano (https://github.com/kevinblighe/EnhancedVolcano) with FDR cut-off of 0.05. Quantitative real-time PCR for selected targets were performed with primer sequences found in Supplementary Table 6.

#### Motif search

*De novo* motif search was performed using both HOMER (Heinz et al., 2010) with the flags “-size given –mask –S 50” and MEME-ChIP (Bailey et al., 2009) with the flags “-drene-m 50 –meme-nmotifs 50”. For directed motif search wherein we searched for the abundance of our motif of interest (Motif 2 in Figure 3C) in called CUT&RUN peaks, HOMER was utilized with a calculated motif position weight matrix and the flags “-find pos_weight_matrix.motif”.

#### Data and code availability

The datasets and code utilized in this study are available at GEO (accession GSE136332) and on GitHub at https://github.com/mbassalbioinformatics/CnRAP

## Declarations

All authors have consented to publication and declare that they have no conflict of interest.

## Author contributions are

N.R.K., L.C., D.G.T., M.L.B., F.L., and S.P. conceived and designed the experiments; H.L and L.E.S. helped with the design of the experiments; N.R.K., H.K.T., J.V.K., K.J.Y., Y.Y., J.L., Y.L., and A.S. performed the experiments; N.R.K., M.A.B., J.J.Y., and C.S.W. performed the analyses; N.R.K., MA.B., L.C., and D.G.T. prepared the manuscript.

