## Supplementary figures and legends for "Zinc finger protein SALL4 functions through an AT-rich motif to regulate gene expression"

**Supplementary Figure 1.** SALL4 binds an AT-rich motif in PBM experiments. **A)** The mean PBM enrichment (E)-scores of FLAG-tagged SALL4A WT, AΔZFC2 and AΔZFC4 mutants for most AT-rich motifs represented in PBM, with FLAG alone as the negative control. The IUPAC motif representing the PBM-derived SALL4 motif shown in Figure 1A is indicated by the red bracket. **B)** FLAG peptide control alone does not bind the SALL4 consensus sequence. **C)** SALL4B also binds the AT-rich SALL4A binding motif but SALL4B mutant **(D)** lacking zinc finger cluster 4 (BΔZFC4) does not bind (right). The colored bars above the PWM indicate the linear structure of WT SALL4B or BΔZFC4.

**Supplementary Figure 2.** EMSA experiments demonstrated that SALL4 binds DNA. **A)** SALL4A shifts a PBM probe (WT-2) different from the probe used in Figure 1D, also containing the AT-rich motif (Supplementary Table 1); S4A=SALL4A. **B)** SALL4A binds DNA (lanes 2-5) more strongly than SALL4B (lanes 6-9); S4B=SALL4B, different concentrations of the purified proteins used in the EMSA reactions are noted. All EMSA reactions contain poly dI:dC competitor to reduce background signals.

**Supplementary Figure 3.** ITC experiments showing purified SALL4 ZFC4 binds DNA oligos containing the WTATB motif that is different from oligos in Figure 2B. ZFC4 binds WT oligos (left, WT sequence: GATAAATATTTG) and not when the motif was mutated (right, mutant sequence: GATAAACGCTTG).

**Supplementary Figure 4.** SALL4 binds AT-rich motif in cells. **A)** SALL4 is localized exclusively in the chromatin fraction in SNU398 liver cancer cells. **B)** SALL4 expression is high in SNU398 liver cancer cells compared to other similar liver cancer cell lines such as SNU387. **C)** Top MEME-ChIP motifs from *de novo* analysis of three CUT&RUN replicates with their respective E-values.

**Supplementary Figure 5.** Related to Figure 4. **A)** Representative western blot showing SALL4 knockdown (KD) efficiency in liver cancer cells using two different shRNAs and scrambled (SCR) as control; beta-Actin levels served as loading control; shSALL-2 used in RNA-seq experiments. **B)** *De novo* motif search focusing on 1,192 SALL4 CUT&RUN peaks near the differentially expressed genes. **C)** SALL4 binds components of Nucleosome remodeling and histone deacetlyase (NuRD) complex. Co-IP of SALL4 and HDAC2 in K562 cells over-expressing either WT (lanes 2 and 3) or mutant SALL4 (lane 4), with EV-K562 cells served as a negative control (lane 1). Rep1 and rep2 denote experiments performed with two separate cell clones with stable SALL4A over-expression. 3A5A mutant has two amino acid substitutions at positions 3 and 5, previously shown to not bind the NuRD complex.

**Supplementary Figure 6.** SALL4 regulates histone demethylases. Representative CUT&RUN genomic tracks at *KDM3A* (**A**) and *KDM4C* (**B**) in SNU398 liver cancer cells. scaless shown are 0-15 for both. Regions highlighted show the called peaks from the analysis. **C)** ChIP-qPCR validation of SALL4 target gene, *KDM3A*, percentage of input was used to calculate relative fold enrichment between positive and negative regions, primer sequences can be found in Supplementary Table 5; results shown from three biological replicates. **D)** *KDM3A* gene expression in SNU398 liver cancer cells after one of its SALL4 motifs in peak 2 shown in (A) was mutated via transient expression of Cas9-GFP and single-guide RNA. Successfully transfected cells were sorted by GFP expression, experiment was performed in biological duplicates.

**Supplementary Figure 7.** Prolonged SALL4 KD led to cell death. **A)** Western blotting of SNU398 liver cancer cell lysates at 40, 72, and 120hr after SALL4 KD with two different shRNAs, molecular weight marker is indicated on the right. **B)** Percentages of DAPI+ dead cells that were also successfully transduced by shRNAs (GFP+) at 40, 72, or 120hr after SALL4 KD. Experiments were performed in biological duplicates.

**Supplementary Figure 8.** Immunofluorescence staining of SALL4 (Abcam ab57577) in SNU398 liver cancer cells after successful transduction with SCR control or two different shSALL4s (GFP+). White scale bars denote 33μm.

**Supplementary Table 1.** Probe sequences used in EMSA assays.

**Supplementary Table 2.** SALL4 CUT&RUN peaks in SNU398 liver cancer cells common among three biological triplicates. Peaks within 5kb of each other were collapsed into a single peak.

**Supplementary Table 3.** shSALL4 target sequences and CRISPR/Cas9 target sequence

**Supplementary Table 4.** List of differentially expressed genes after SALL4 KD in SNU398 liver cancer cells.

**Supplementary Table 5**. List of differentially expressed genes bound by SALL4 in CUT&RUN.

**Supplementary Table 7.** List of ChIP-qPCR and qPCR primers.

#### Supplementary Figure 1

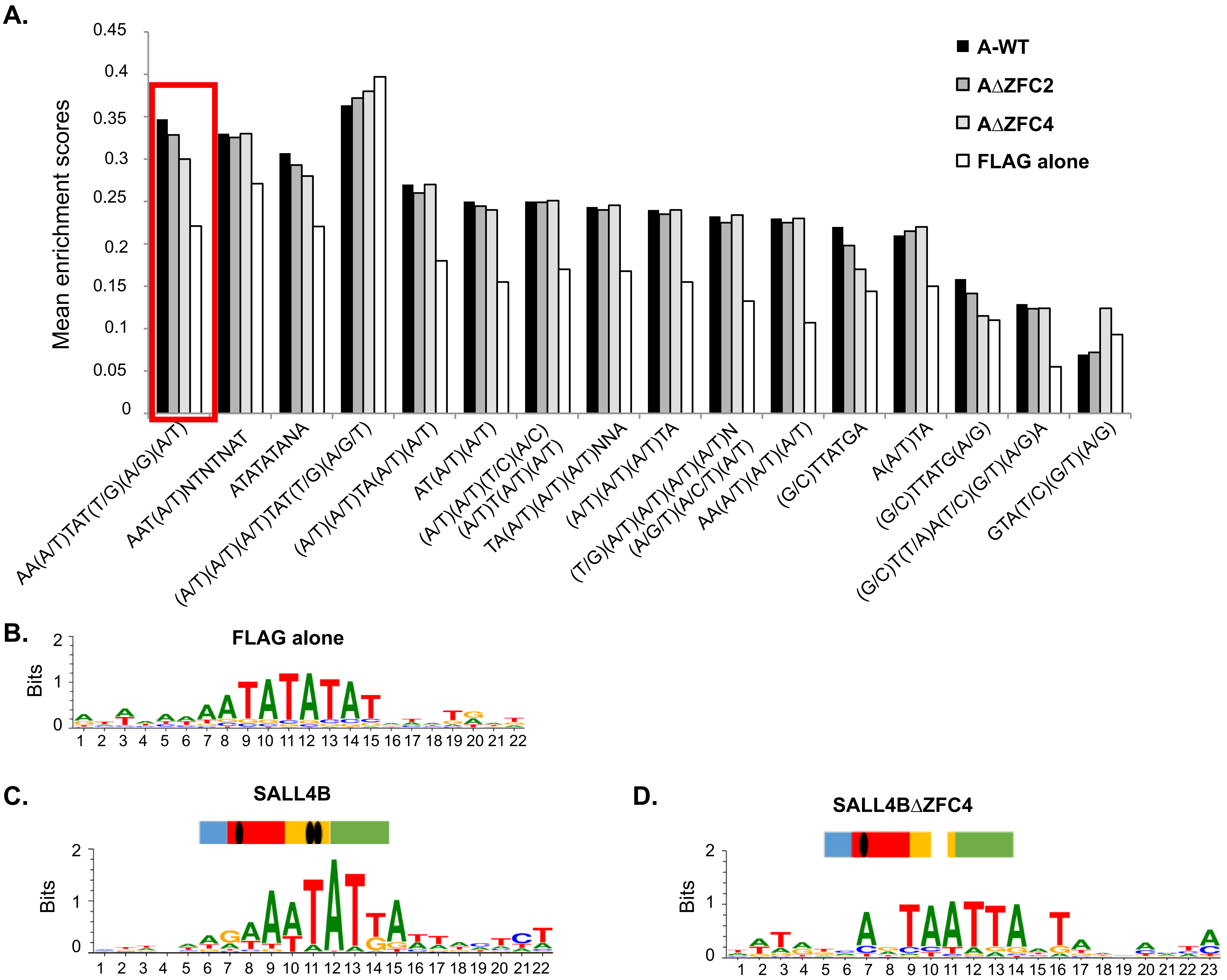

**Supplementary Figure 2**

**A.**

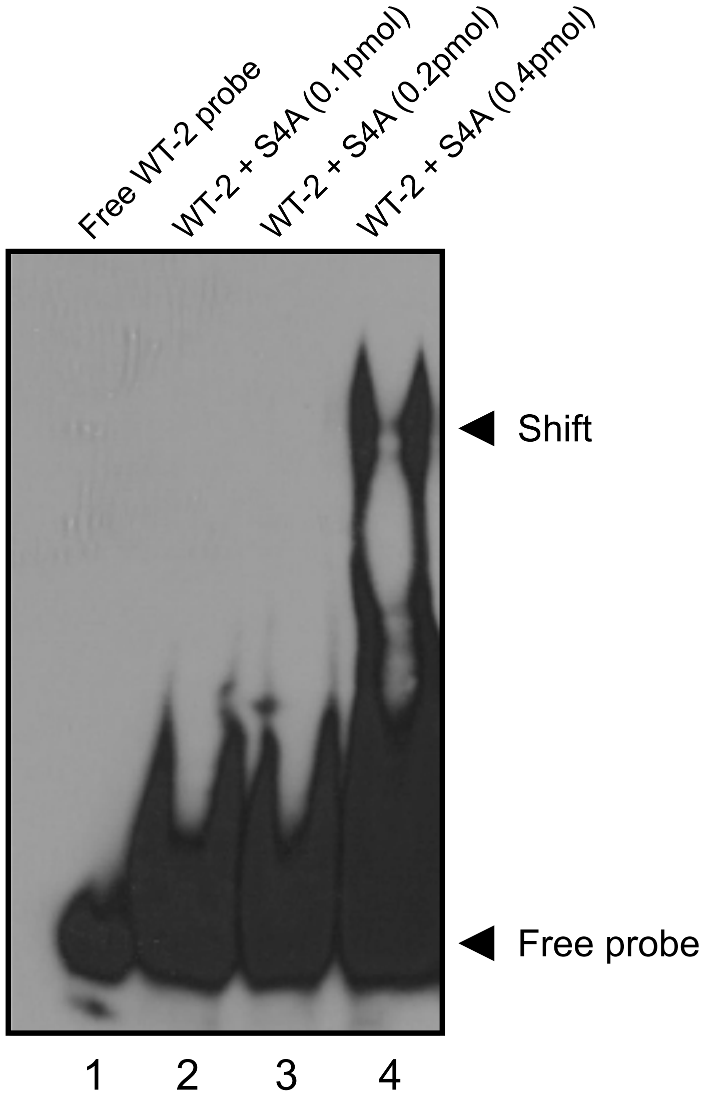


**B.**

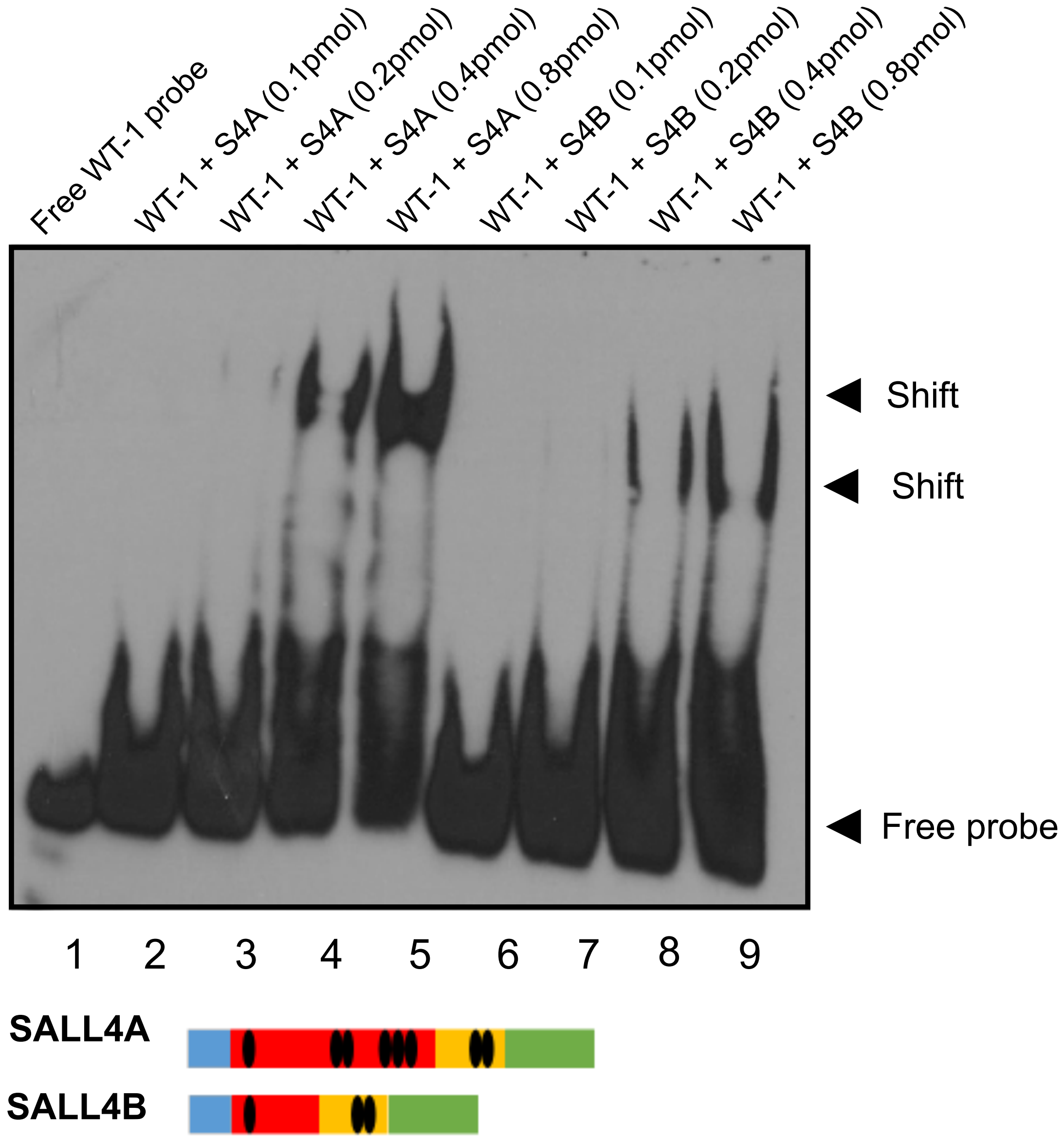

#### Supplementary Figure 3

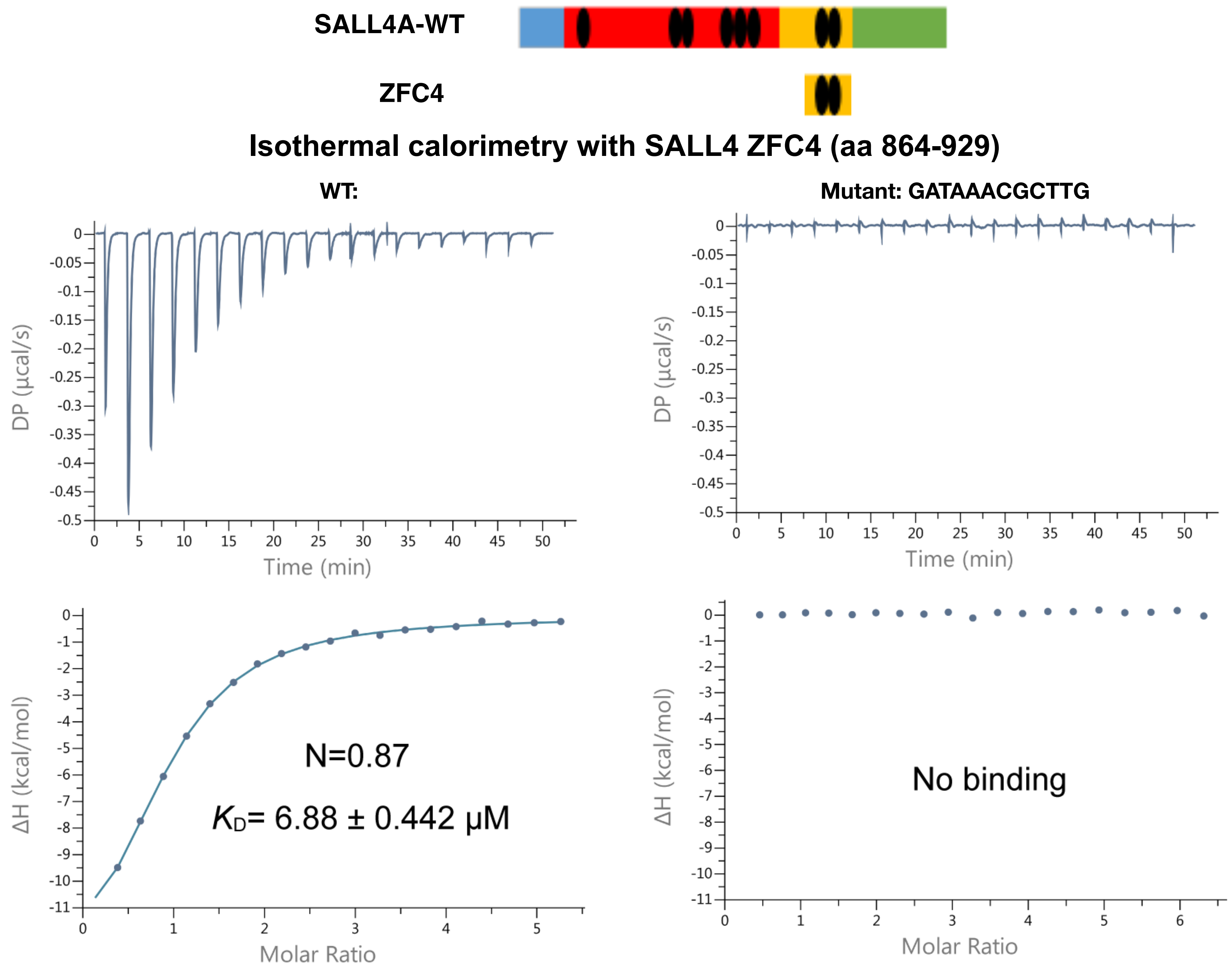

### Supplementary Figure 4

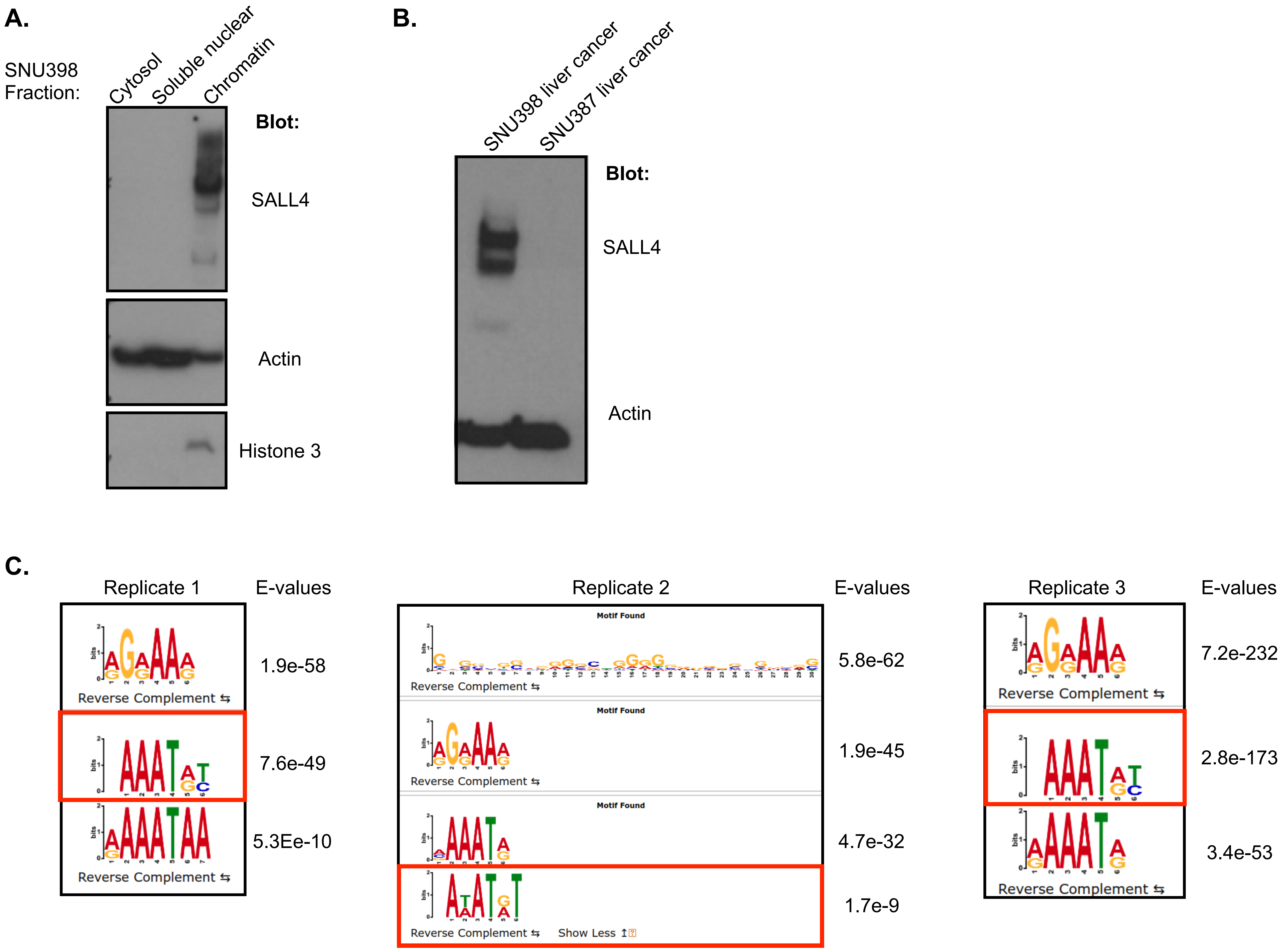

#### Supplementary Figure 5

**A.**

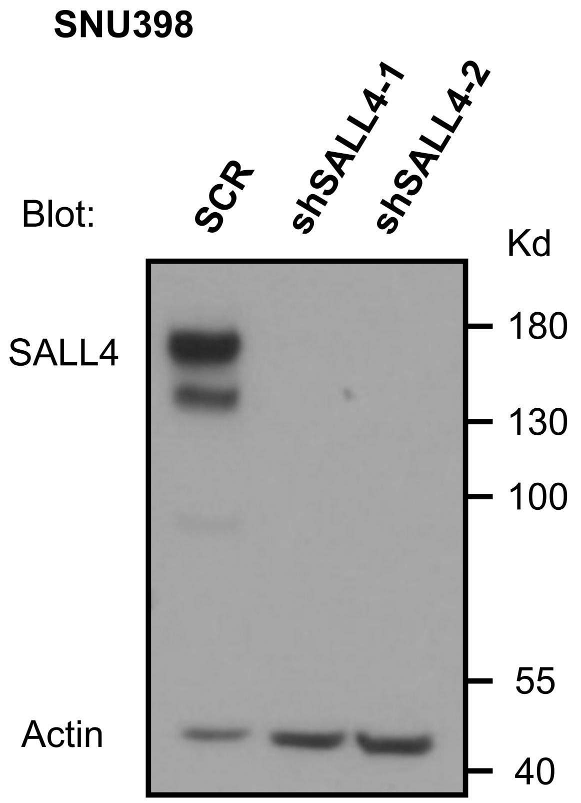


**B.**

**De novo motif search**
**SALL4-bound differentially expressed genes**

| Rank | Motif | P-value | log P-pvalue |
|---|---|---|---|
| 1 | GAAATTTTCATG | 1e-26 | -6.097e+01 |
| 2 | ACAATTTGTGGC | 1e-22 | -5.079e+01 |
| 3 | TAATATTGCAAT | 1e-21 | -4.968e+01 |
| 4 | GCAATTATTGCT | 1e-20 | -4.827e+01 |
| 5 | GTTTAACTGA | 1e-19 | -4.484e+01 |

**C.**

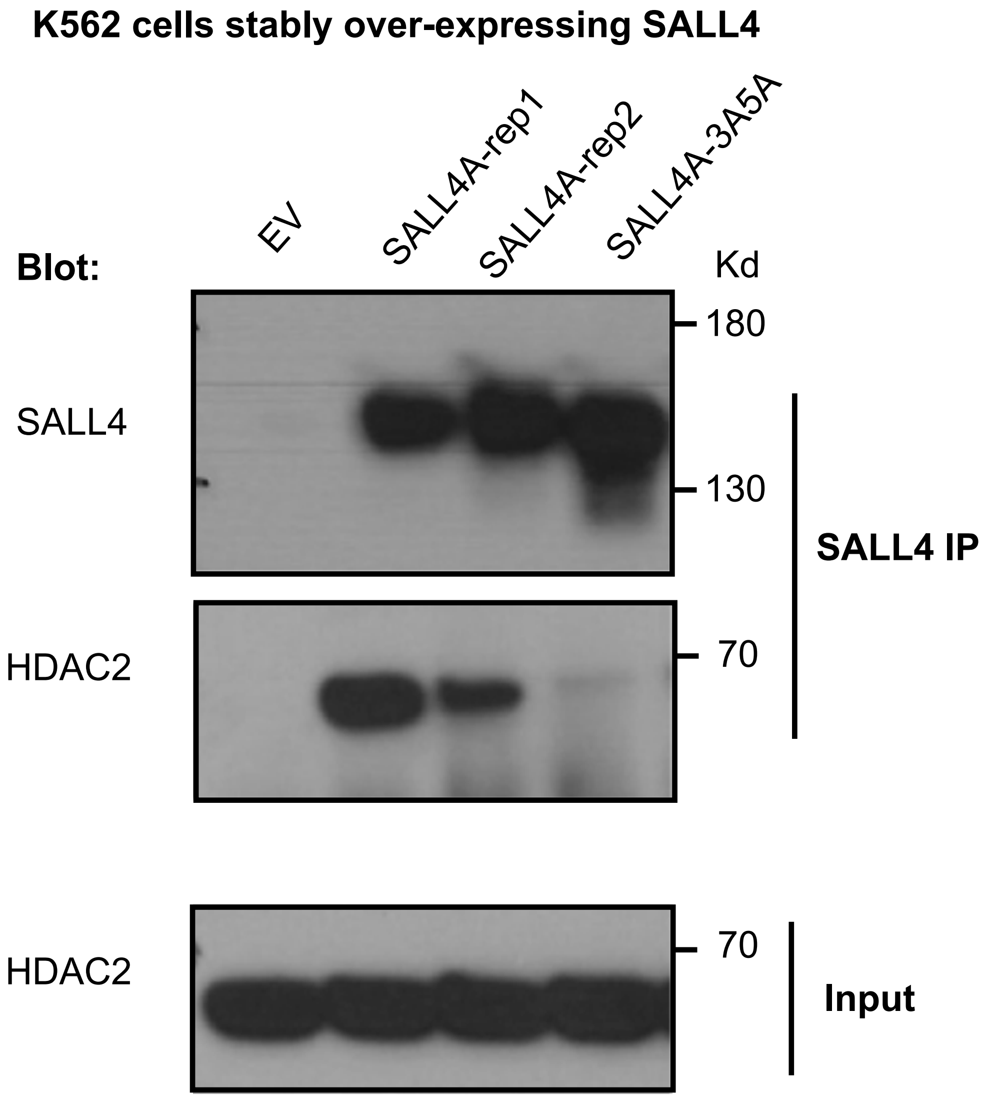

#### Supplementary Figure 6

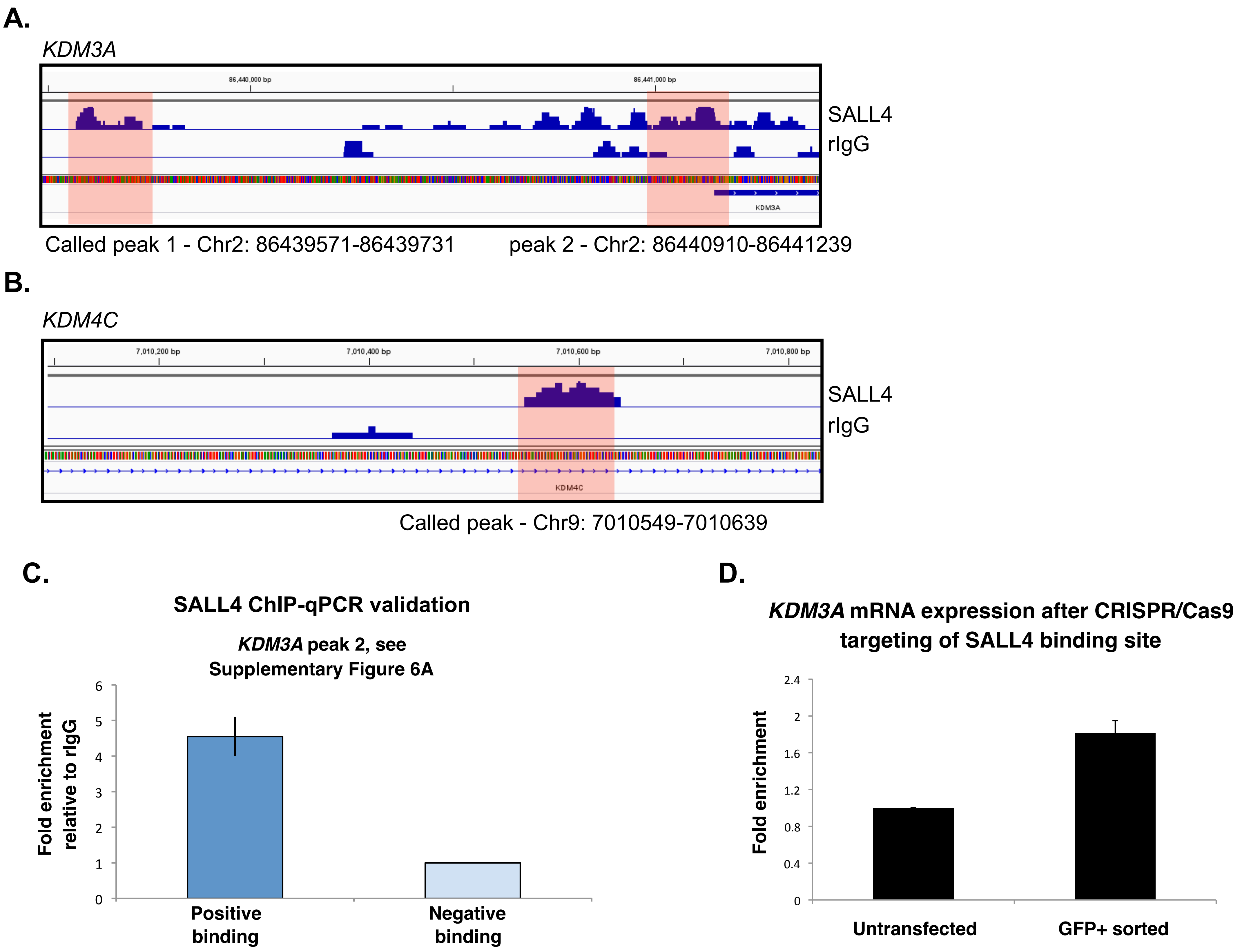

### Supplementary Figure 7

**A.**

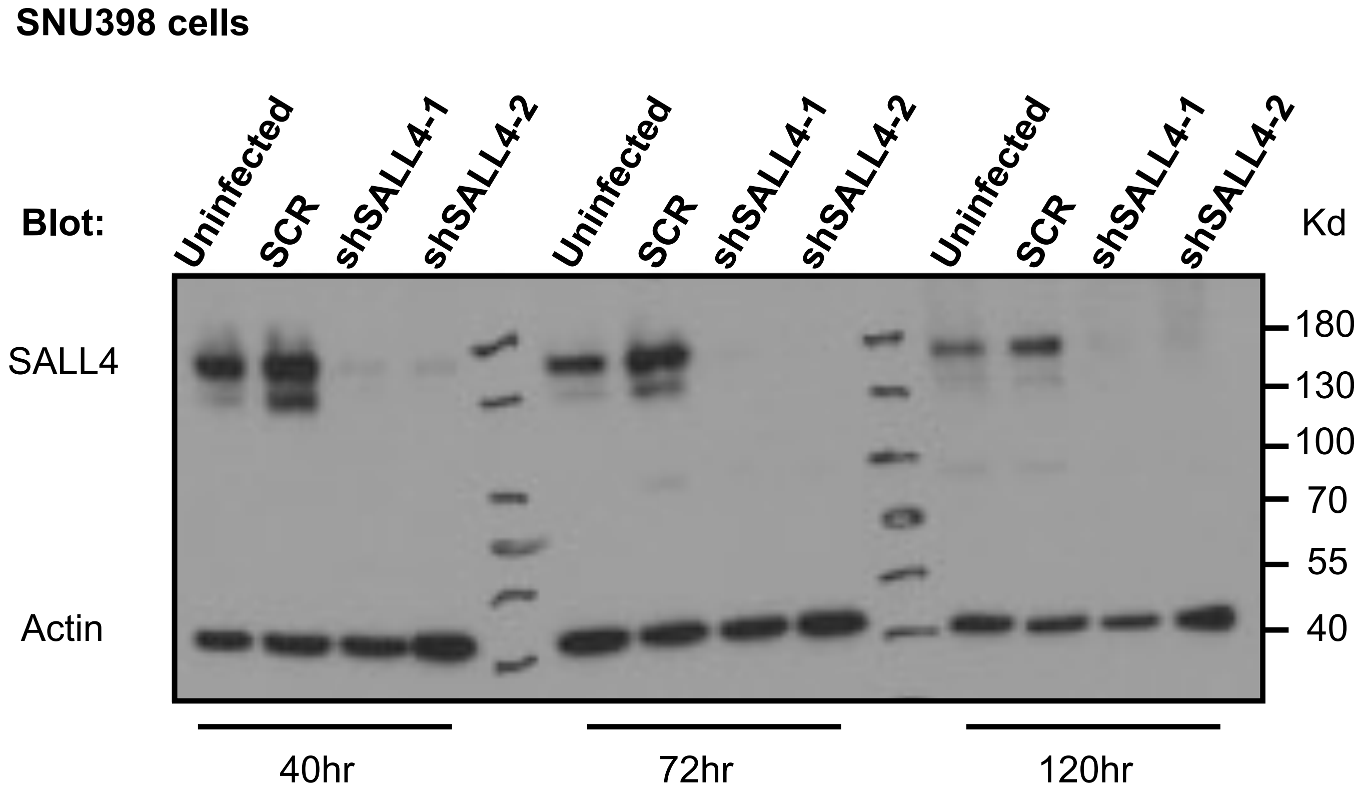


**B.**

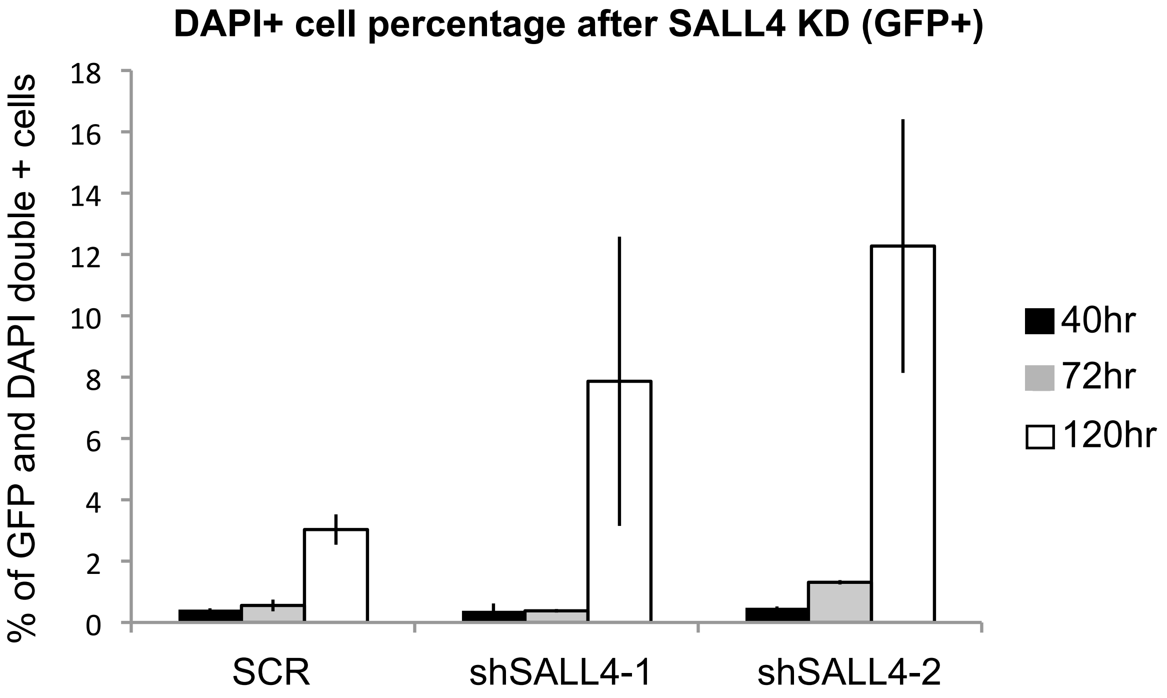

#### Supplementary Figure 8

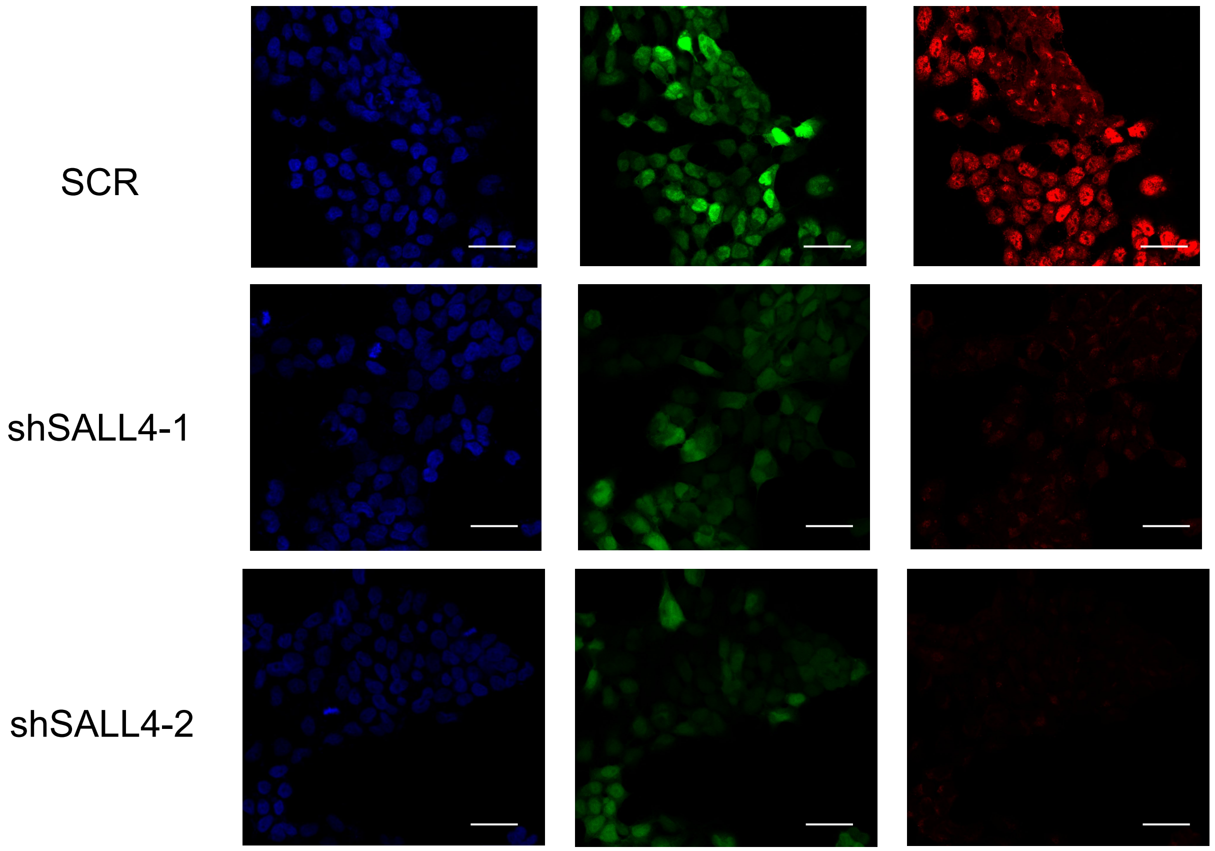


Blue: DAPI
Green: shRNA - GFP
Red: SALL4-Abcam antibody; Alexa 546
