## Supplementary Tables for "Zinc finger protein SALL4 functions through an AT-rich motif to regulate gene expression": Suppl_Table4 DGEs.pdf

Supplementary Table 4. List of differentially expressed genes after SALL4 KD in SNU398 liver cancer cells.

| gene_id | logFC | AveExpr | t | P.Value | adj.P.Val |
|---|---|---|---|---|---|
| RRBP1 | -2.8401893 | 6.95688834 | -29.668744 | 5.9146E-13 | 8.9554E-09 |
| GNB4 | -2.0096768 | 5.40672299 | -21.913718 | 2.4296E-11 | 1.5462E-07 |
| HDGFL3 | -1.7798554 | 6.28037176 | -21.500693 | 3.0635E-11 | 1.5462E-07 |
| KTN1 | -1.3968519 | 8.37472703 | -20.445608 | 5.6498E-11 | 1.6555E-07 |
| TTL | -1.6086999 | 6.80196757 | -20.413263 | 5.7595E-11 | 1.6555E-07 |
| ARCN1 | -1.5501681 | 7.18905864 | -20.195571 | 6.5604E-11 | 1.6555E-07 |
| C19orf73 | 4.59498371 | 2.65828904 | 19.7304172 | 8.7037E-11 | 1.8826E-07 |
| SH3BGRL3 | -2.3079998 | 4.25332212 | -19.460918 | 1.0282E-10 | 1.946E-07 |
| ITPRIPL2 | -1.7505574 | 5.61318356 | -19.144786 | 1.2538E-10 | 2.0197E-07 |
| SUCLG2 | -1.853352 | 5.0740771 | -19.04694 | 1.3339E-10 | 2.0197E-07 |
| HMGB3 | -1.585145 | 6.07777549 | -18.590631 | 1.7883E-10 | 2.1802E-07 |
| PLEKHB2 | -1.5018616 | 6.52390379 | -18.577355 | 1.8038E-10 | 2.1802E-07 |
| LMBR1 | -1.5912793 | 6.20796686 | -18.403058 | 2.0213E-10 | 2.1802E-07 |
| CMTM4 | -1.6680181 | 6.1265086 | -18.259878 | 2.2211E-10 | 2.1802E-07 |
| BUD23 | -1.5462202 | 6.43387587 | -18.240789 | 2.2493E-10 | 2.1802E-07 |
| SALL4 | -1.6295135 | 5.62655801 | -18.20459 | 2.3039E-10 | 2.1802E-07 |
| COX7C | 1.5109857 | 6.4292686 | 17.7743106 | 3.0738E-10 | 2.4852E-07 |
| IDH1 | -1.6723163 | 7.51925691 | -17.740441 | 3.1452E-10 | 2.4852E-07 |
| PPID | -1.5936512 | 5.55460127 | -17.719554 | 3.1901E-10 | 2.4852E-07 |
| OSTM1 | -1.4164391 | 6.69379047 | -17.677468 | 3.2828E-10 | 2.4852E-07 |
| CDIPT | -1.5481019 | 5.76528199 | -17.342826 | 4.1314E-10 | 2.9787E-07 |
| HES2 | -3.0615166 | 2.48397623 | -17.190725 | 4.5927E-10 | 3.0423E-07 |
| MAPK6 | -1.3059075 | 7.13594892 | -17.181803 | 4.6214E-10 | 3.0423E-07 |
| GSPT1 | -1.2834972 | 7.2907667 | -16.808864 | 6.0141E-10 | 3.5981E-07 |
| RBBP7 | -1.4695388 | 6.57934003 | -16.789198 | 6.0992E-10 | 3.5981E-07 |
| DENND6A | -1.7386685 | 4.975521 | -16.771077 | 6.1787E-10 | 3.5981E-07 |
| C1orf61 | 3.70022623 | 1.87993435 | 16.5898181 | 7.0382E-10 | 3.8995E-07 |
| ZNF468 | -1.7692541 | 4.46650198 | -16.549654 | 7.2455E-10 | 3.8995E-07 |
| AHNAK2 | -2.0412526 | 4.0525795 | -16.507752 | 7.4689E-10 | 3.8995E-07 |
| PLBD1 | -1.37937 | 6.23830436 | -16.402631 | 8.0626E-10 | 4.0692E-07 |
| DIXDC1 | -1.4123488 | 6.13750561 | -15.994601 | 1.0896E-09 | 5.2546E-07 |
| HDAC5 | 1.61104217 | 5.22922908 | 15.9323682 | 1.1416E-09 | 5.2546E-07 |
| ABCB10 | -1.6196808 | 4.82867598 | -15.928074 | 1.1453E-09 | 5.2546E-07 |
| ITGAV | -1.0135644 | 8.93550924 | -15.501621 | 1.5829E-09 | 6.979E-07 |
| CACUL1 | 1.16564962 | 7.39983835 | 15.4769181 | 1.6133E-09 | 6.979E-07 |
| PIK3CD | -1.7989119 | 4.21028847 | -15.17905 | 2.0329E-09 | 8.5502E-07 |
| ARPC3 | 1.29897475 | 6.58643172 | 15.1300021 | 2.1127E-09 | 8.6454E-07 |

| | | | | | |
|---|---|---|---|---|---|
| CYCS | -1.2153129 | 7.07727081 | -15.04275 | 2.2629E-09 | 9.0166E-07 |
| TXNRD1 | -1.0410416 | 8.47228229 | -14.953438 | 2.4287E-09 | 9.4291E-07 |
| STK24 | -1.3658196 | 5.40866686 | -14.890282 | 2.5538E-09 | 9.6669E-07 |
| ANKRD40 | 1.4015178 | 6.3607788 | 14.8501046 | 2.637E-09 | 9.7268E-07 |
| TULP3 | 1.6059441 | 4.43834853 | 14.8214531 | 2.6981E-09 | 9.7268E-07 |
| SOD2 | -1.18716 | 6.67706937 | -14.639437 | 3.1237E-09 | 1.0999E-06 |
| ZNF853 | 2.01801858 | 3.36744204 | 14.587924 | 3.2569E-09 | 1.1207E-06 |
| TXNIP | 1.58411403 | 5.39282062 | 14.4889495 | 3.5302E-09 | 1.1878E-06 |
| BNIP3L | -2.8868756 | 4.09147179 | -14.434789 | 3.6901E-09 | 1.2146E-06 |
| LRRC75A | -1.2822872 | 6.67390639 | -14.37251 | 3.8837E-09 | 1.2511E-06 |
| BCL11B | 1.77976061 | 5.37960507 | 14.3286728 | 4.0265E-09 | 1.2587E-06 |
| OCA2 | 1.66096816 | 4.84491764 | 14.2949636 | 4.1401E-09 | 1.2587E-06 |
| SKA2 | 1.36490605 | 5.91597893 | 14.2901991 | 4.1564E-09 | 1.2587E-06 |
| HTRA2 | 1.63595744 | 6.62630257 | 14.2384815 | 4.3383E-09 | 1.2745E-06 |
| RRN3 | -1.1766507 | 6.21616236 | -14.227774 | 4.377E-09 | 1.2745E-06 |
| TAS1R1 | -4.3140764 | 1.39921885 | -14.185913 | 4.532E-09 | 1.2947E-06 |
| RPL14 | -1.0640891 | 7.39668322 | -14.12344 | 4.7744E-09 | 1.3387E-06 |
| ADAR | -0.9834041 | 8.46985341 | -14.018595 | 5.213E-09 | 1.4351E-06 |
| PHF5A | -1.2359498 | 5.68361461 | -13.917313 | 5.678E-09 | 1.5191E-06 |
| SAMD8 | -1.030572 | 7.79544348 | -13.873628 | 5.8922E-09 | 1.5191E-06 |
| CASC1 | 1.40183266 | 4.79848259 | 13.862841 | 5.9465E-09 | 1.5191E-06 |
| RAB15 | 1.30284778 | 5.89298443 | 13.8529327 | 5.9967E-09 | 1.5191E-06 |
| BYSL | -1.2501336 | 5.76530526 | -13.848442 | 6.0197E-09 | 1.5191E-06 |
| ATXN7L3B | 1.05847432 | 7.04940712 | 13.7988148 | 6.2795E-09 | 1.5578E-06 |
| PAQR5 | -1.6751444 | 4.36000006 | -13.774086 | 6.4135E-09 | 1.5578E-06 |
| ABCD3 | -1.0909903 | 6.73349843 | -13.755949 | 6.5137E-09 | 1.5578E-06 |
| ANKRD46 | -1.2812074 | 5.38234036 | -13.730705 | 6.656E-09 | 1.5578E-06 |
| TYRO3 | -1.2167494 | 5.72076939 | -13.725137 | 6.6878E-09 | 1.5578E-06 |
| CEND1 | -1.961073 | 3.0666186 | -13.626231 | 7.2814E-09 | 1.6704E-06 |
| GATM | -1.1017185 | 6.80101027 | -13.50454 | 8.0905E-09 | 1.8283E-06 |
| DUSP22 | 1.18709585 | 5.73990306 | 13.4792359 | 8.2706E-09 | 1.8415E-06 |
| SLC39A6 | -1.1373629 | 6.12772278 | -13.438227 | 8.5717E-09 | 1.8492E-06 |
| RFK | -1.4198482 | 4.27205268 | -13.430229 | 8.6318E-09 | 1.8492E-06 |
| TNFAIP3 | 1.85468352 | 3.52750106 | 13.4249922 | 8.6714E-09 | 1.8492E-06 |
| DHRS2 | 1.99946634 | 3.86639432 | 13.4046622 | 8.8269E-09 | 1.8505E-06 |
| PSAP | -0.9099007 | 9.23057814 | -13.392439 | 8.9219E-09 | 1.8505E-06 |
| NMD3 | -1.1632538 | 5.68410929 | -13.36709 | 9.1223E-09 | 1.8665E-06 |
| LGALS1 | -1.1108951 | 6.49326882 | -13.275788 | 9.8853E-09 | 1.9957E-06 |
| DDHD1 | -1.0751132 | 6.75673319 | -13.209743 | 1.048E-08 | 2.0879E-06 |
| ELOVL6 | -1.3500163 | 4.63950879 | -13.192576 | 1.0641E-08 | 2.0924E-06 |
| CAPZA2 | -1.0546588 | 6.55591741 | -13.17264 | 1.0831E-08 | 2.1024E-06 |
| MTMR2 | -0.9925992 | 7.06076327 | -13.058718 | 1.1989E-08 | 2.267E-06 |
| CMTM3 | 1.24664496 | 5.491062 | 13.0499909 | 1.2083E-08 | 2.267E-06 |
| ILF3-DT | 1.53344991 | 4.74136292 | 13.0226993 | 1.2383E-08 | 2.267E-06 |
| TAPBP | -1.1353162 | 6.54118379 | -13.015589 | 1.2462E-08 | 2.267E-06 |

| | | | | | |
|---|---|---|---|---|---|
| IARS2 | -1.0105287 | 7.86223051 | -13.007617 | 1.2551E-08 | 2.267E-06 |
| LAP3 | -1.0466505 | 6.60283264 | -13.005336 | 1.2577E-08 | 2.267E-06 |
| UBE2Q2 | -1.1677544 | 5.46728108 | -12.955332 | 1.3156E-08 | 2.3038E-06 |
| SLC7A1 | -1.1144666 | 6.9692252 | -12.948504 | 1.3237E-08 | 2.3038E-06 |
| MMP14 | -1.0456883 | 6.6786012 | -12.948494 | 1.3238E-08 | 2.3038E-06 |
| MARS | -0.9298354 | 8.85455312 | -12.922929 | 1.3547E-08 | 2.3145E-06 |
| ALG12 | -1.5740473 | 3.92917422 | -12.918204 | 1.3605E-08 | 2.3145E-06 |
| PIK3AP1 | -1.9497842 | 3.19823426 | -12.864328 | 1.4285E-08 | 2.3942E-06 |
| RAB6A | -1.22423 | 6.38696333 | -12.845712 | 1.4528E-08 | 2.3942E-06 |
| PROCR | -1.0676034 | 6.36413547 | -12.844211 | 1.4548E-08 | 2.3942E-06 |
| QSOX1 | -1.0685427 | 6.31175829 | -12.811474 | 1.4987E-08 | 2.4401E-06 |
| CCDC90B | 1.20631563 | 5.3783851 | 12.7726938 | 1.5527E-08 | 2.501E-06 |
| GLS | -1.080384 | 6.58254974 | -12.732554 | 1.6107E-08 | 2.5593E-06 |
| TMEM64 | -1.0746271 | 6.04340307 | -12.724443 | 1.6227E-08 | 2.5593E-06 |
| KLHL18 | -1.1566344 | 5.72352004 | -12.688278 | 1.6774E-08 | 2.6183E-06 |
| TRIM47 | -1.4970949 | 4.05973153 | -12.649502 | 1.7383E-08 | 2.665E-06 |
| ANGPTL4 | -2.2319712 | 2.81396732 | -12.646837 | 1.7426E-08 | 2.665E-06 |
| NUCKS1 | -0.8375155 | 8.78192796 | -12.617899 | 1.7896E-08 | 2.6674E-06 |
| C12orf75 | -1.5584251 | 3.72553718 | -12.616283 | 1.7923E-08 | 2.6674E-06 |
| MAP2K4 | -1.1417417 | 5.44171893 | -12.610431 | 1.802E-08 | 2.6674E-06 |
| FBXO2 | -1.4765157 | 4.28012656 | -12.602929 | 1.8145E-08 | 2.6674E-06 |
| CT55 | -1.5262973 | 3.83924915 | -12.5794 | 1.8544E-08 | 2.6998E-06 |
| SHTN1 | -1.2240163 | 4.95036742 | -12.556087 | 1.8949E-08 | 2.7324E-06 |
| GFOD1 | -1.2079218 | 5.08810531 | -12.535617 | 1.9312E-08 | 2.7585E-06 |
| CDO1 | -1.3075802 | 4.60262643 | -12.511383 | 1.9751E-08 | 2.7949E-06 |
| MALT1 | -1.2045678 | 5.34981166 | -12.457572 | 2.0766E-08 | 2.9113E-06 |
| FSD1 | -1.0806064 | 6.25584759 | -12.43723 | 2.1164E-08 | 2.9399E-06 |
| GALE | -1.1155426 | 5.69140103 | -12.416481 | 2.1578E-08 | 2.9702E-06 |
| CFDP1 | 1.15343944 | 5.51753632 | 12.3748414 | 2.2437E-08 | 3.0605E-06 |
| AASDHPPT | -1.0280371 | 6.03045623 | -12.305682 | 2.3944E-08 | 3.2315E-06 |
| TOGARAM1 | 0.99802664 | 6.67085147 | 12.2980431 | 2.4117E-08 | 3.2315E-06 |
| LZTS3 | -1.8623413 | 3.32613882 | -12.222808 | 2.5896E-08 | 3.4237E-06 |
| LIN7C | -1.0777154 | 5.72250264 | -12.218405 | 2.6004E-08 | 3.4237E-06 |
| YARS | -0.8370277 | 8.4118637 | -12.193756 | 2.662E-08 | 3.4721E-06 |
| KCTD3 | 1.00639098 | 6.21038943 | 12.1727938 | 2.7155E-08 | 3.4721E-06 |
| GRPEL1 | -1.0553622 | 6.33302494 | -12.168025 | 2.7279E-08 | 3.4721E-06 |
| SEC23B | -0.9098568 | 7.38009519 | -12.167655 | 2.7289E-08 | 3.4721E-06 |
| RPA1 | -0.9557714 | 6.95669853 | -12.142099 | 2.7961E-08 | 3.528E-06 |
| ACAA1 | -1.4759914 | 4.27070979 | -12.042274 | 3.0763E-08 | 3.8235E-06 |
| SLC2A6 | -1.1848124 | 5.45584221 | -12.04077 | 3.0808E-08 | 3.8235E-06 |
| CTBP1 | -0.9059925 | 7.40679188 | -12.023835 | 3.1313E-08 | 3.8357E-06 |
| ACADSB | -1.3467221 | 4.20861604 | -12.020527 | 3.1413E-08 | 3.8357E-06 |
| PREX1 | 1.12565104 | 5.27427154 | 12.0048968 | 3.1889E-08 | 3.8624E-06 |
| CPNE3 | -0.9484404 | 6.67031781 | -11.996685 | 3.2142E-08 | 3.8624E-06 |
| RNF149 | -1.1047868 | 5.2899418 | -11.921484 | 3.4564E-08 | 4.1007E-06 |

| | | | | | |
|---|---|---|---|---|---|
| VGF | 1.35706155 | 6.2833442 | 11.9184192 | 3.4667E-08 | 4.1007E-06 |
| BZW1 | -0.9033137 | 7.1998302 | -11.895813 | 3.5435E-08 | 4.1591E-06 |
| APLP2 | -0.80404 | 8.62543542 | -11.88603 | 3.5773E-08 | 4.1664E-06 |
| CACYBP | -0.9137044 | 6.95569022 | -11.848432 | 3.7105E-08 | 4.2886E-06 |
| BTBD3 | -1.0629728 | 5.89571905 | -11.811197 | 3.8476E-08 | 4.3202E-06 |
| TMEM87B | -1.5092666 | 3.51838121 | -11.809128 | 3.8554E-08 | 4.3202E-06 |
| SPR | -0.9303759 | 7.05542061 | -11.80842 | 3.8581E-08 | 4.3202E-06 |
| CARS2 | -1.0898365 | 5.55326595 | -11.808265 | 3.8587E-08 | 4.3202E-06 |
| CDCA7 | 0.97093942 | 6.33115321 | 11.8024773 | 3.8805E-08 | 4.3202E-06 |
| MGAT4B | 0.93260447 | 6.75329155 | 11.7524458 | 4.0752E-08 | 4.5039E-06 |
| NIPA1 | -0.9573135 | 6.38482131 | -11.709883 | 4.2492E-08 | 4.6285E-06 |
| L1CAM | -2.4349604 | 1.88767791 | -11.706929 | 4.2615E-08 | 4.6285E-06 |
| MVP | -1.2233904 | 4.77514332 | -11.702589 | 4.2797E-08 | 4.6285E-06 |
| PARD3B | 1.09254715 | 5.19968524 | 11.6491693 | 4.5111E-08 | 4.7994E-06 |
| SMARCB1 | -0.9971766 | 6.384966 | -11.645263 | 4.5286E-08 | 4.7994E-06 |
| TROVE2 | -0.9763848 | 6.24538635 | -11.644308 | 4.5329E-08 | 4.7994E-06 |
| ATP10D | 0.9593237 | 6.3406159 | 11.6176387 | 4.654E-08 | 4.8935E-06 |
| AL365259.1 | 1.37208666 | 3.83525977 | 11.6101376 | 4.6887E-08 | 4.896E-06 |
| MYBBP1A | -0.8657729 | 7.65060566 | -11.56051 | 4.9254E-08 | 5.1079E-06 |
| MRPL18 | -1.120556 | 5.29658925 | -11.538713 | 5.0333E-08 | 5.179E-06 |
| EHD4 | -1.0082242 | 5.88195928 | -11.532618 | 5.064E-08 | 5.179E-06 |
| METAP1D | 1.08609389 | 5.07686242 | 11.5261837 | 5.0965E-08 | 5.179E-06 |
| FAM214A | 1.15457613 | 4.63103404 | 11.5121191 | 5.1685E-08 | 5.1873E-06 |
| OLFM1 | -1.0657651 | 5.40850603 | -11.5112 | 5.1732E-08 | 5.1873E-06 |
| SRP14 | 0.91955209 | 6.79210975 | 11.457191 | 5.4602E-08 | 5.4076E-06 |
| C1QL4 | -1.446951 | 3.9988902 | -11.456417 | 5.4644E-08 | 5.4076E-06 |
| PQLC3 | -1.5653209 | 3.43130728 | -11.444006 | 5.5328E-08 | 5.4398E-06 |
| BRI3BP | -0.975128 | 5.93195093 | -11.433076 | 5.5938E-08 | 5.4642E-06 |
| OPTN | 0.98533682 | 6.09078498 | 11.425701 | 5.6354E-08 | 5.4696E-06 |
| AKAP5 | -1.2590292 | 4.16777208 | -11.388111 | 5.8524E-08 | 5.644E-06 |
| NUDT2 | -1.2874092 | 4.18987692 | -11.359792 | 6.0218E-08 | 5.7706E-06 |
| PAN3 | 0.99605783 | 5.66755041 | 11.3274641 | 6.2216E-08 | 5.9246E-06 |
| BBC3 | 1.59228318 | 3.27612802 | 11.3057914 | 6.3596E-08 | 6.0101E-06 |
| WBP1L | -0.9607336 | 6.00222755 | -11.300972 | 6.3907E-08 | 6.0101E-06 |
| CRYZ | -0.9392101 | 6.11931108 | -11.237996 | 6.8128E-08 | 6.3313E-06 |
| FAM168B | -0.8424897 | 7.54517537 | -11.237233 | 6.8181E-08 | 6.3313E-06 |
| SRP68 | -0.902031 | 6.6043256 | -11.231525 | 6.8578E-08 | 6.3313E-06 |
| GTF2A1 | -1.096032 | 5.98519837 | -11.213337 | 6.9861E-08 | 6.4107E-06 |
| LIMA1 | -1.187007 | 5.07505628 | -11.202361 | 7.0648E-08 | 6.4438E-06 |
| ZDHHC13 | -1.182453 | 4.57390971 | -11.179815 | 7.2293E-08 | 6.5419E-06 |
| SLC7A2 | -0.8840826 | 6.81203583 | -11.175847 | 7.2587E-08 | 6.5419E-06 |
| FBXW4 | 1.02673301 | 5.33938352 | 11.1456127 | 7.4869E-08 | 6.7076E-06 |
| EPM2A | 1.26688002 | 4.17158447 | 11.0985572 | 7.8575E-08 | 6.9982E-06 |
| GLTPD2 | 1.97896582 | 2.2790152 | 11.0830103 | 7.9842E-08 | 7.0523E-06 |
| INTS14 | 0.98864721 | 6.06209713 | 11.0797206 | 8.0113E-08 | 7.0523E-06 |

| | | | | | |
|---|---|---|---|---|---|
| BNIP2 | -0.9156192 | 6.21212198 | -11.069377 | 8.0971E-08 | 7.0866E-06 |
| NEIL2 | -1.0157587 | 5.57731199 | -11.028538 | 8.4458E-08 | 7.3493E-06 |
| TMED3 | -0.9102075 | 6.61131745 | -10.982637 | 8.857E-08 | 7.6631E-06 |
| FOXA1 | 0.9196457 | 6.16838385 | 10.9732509 | 8.9437E-08 | 7.6941E-06 |
| ERO1A | -1.6846944 | 6.80277097 | -10.943042 | 9.2289E-08 | 7.8947E-06 |
| RAB3D | -1.0007262 | 5.41147461 | -10.932177 | 9.3339E-08 | 7.9373E-06 |
| CCNE1 | -1.1001078 | 4.80746193 | -10.927082 | 9.3836E-08 | 7.9373E-06 |
| CILP2 | -1.4710702 | 3.48800842 | -10.920556 | 9.4476E-08 | 7.947E-06 |
| PCMTD2 | 1.00168045 | 5.46226563 | 10.8993221 | 9.6592E-08 | 8.0801E-06 |
| SYT11 | -0.9288985 | 6.05138752 | -10.882228 | 9.8333E-08 | 8.1805E-06 |
| SUCLA2 | -0.925295 | 5.85765655 | -10.825687 | 1.0433E-07 | 8.6323E-06 |
| SYNJ2BP | 1.0586989 | 4.75602177 | 10.8201442 | 1.0494E-07 | 8.6355E-06 |
| AC144652.1 | 1.19039903 | 4.2657139 | 10.8042245 | 1.0671E-07 | 8.7337E-06 |
| ARPC5 | -1.0012385 | 6.54939591 | -10.790154 | 1.083E-07 | 8.8163E-06 |
| GPRIN1 | -0.9639956 | 6.4501369 | -10.771302 | 1.1048E-07 | 8.945E-06 |
| SC5D | -1.1179349 | 5.48039392 | -10.749893 | 1.13E-07 | 9.0618E-06 |
| TSPAN6 | 1.09133527 | 4.67482282 | 10.7489278 | 1.1312E-07 | 9.0618E-06 |
| HMOX2 | -0.9922834 | 5.48800716 | -10.72475 | 1.1604E-07 | 9.2474E-06 |
| NCOA2 | 0.8446095 | 7.1632795 | 10.6791842 | 1.2178E-07 | 9.6541E-06 |
| KATNB1 | -0.9765434 | 5.379183 | -10.672065 | 1.2271E-07 | 9.6767E-06 |
| FGFR4 | -0.9329039 | 5.85747446 | -10.661487 | 1.241E-07 | 9.6963E-06 |
| PPP1R3B | -1.6141261 | 3.06188453 | -10.660407 | 1.2424E-07 | 9.6963E-06 |
| CMPK1 | -0.8419169 | 6.64170566 | -10.644591 | 1.2635E-07 | 9.8103E-06 |
| ZFP91 | -0.8060446 | 7.10506428 | -10.61289 | 1.3069E-07 | 1.0096E-05 |
| RAB2A | -0.8454596 | 6.49527363 | -10.581619 | 1.3513E-07 | 1.0386E-05 |
| IL6R | -2.7625719 | 0.76955528 | -10.569714 | 1.3686E-07 | 1.0466E-05 |
| EXOSC3 | -1.0052914 | 4.9952047 | -10.562396 | 1.3794E-07 | 1.0495E-05 |
| RICTOR | 0.98931935 | 5.14806858 | 10.5386147 | 1.415E-07 | 1.0696E-05 |
| LIN54 | -0.9160427 | 5.72525827 | -10.535388 | 1.4199E-07 | 1.0696E-05 |
| PNKD | -0.976781 | 5.28074966 | -10.517721 | 1.4471E-07 | 1.0847E-05 |
| ANO10 | -1.0622568 | 4.79390317 | -10.511546 | 1.4567E-07 | 1.0865E-05 |
| CTR9 | -0.8787403 | 6.06005568 | -10.502277 | 1.4713E-07 | 1.0909E-05 |
| ZCRB1 | 0.97464169 | 5.92566459 | 10.4986626 | 1.4771E-07 | 1.0909E-05 |
| COMMD4 | -0.9331857 | 6.69891301 | -10.493973 | 1.4845E-07 | 1.0911E-05 |
| DNAJC22 | -1.5030188 | 3.1948608 | -10.482717 | 1.5027E-07 | 1.0991E-05 |
| RHBDF2 | -0.9336146 | 5.73325652 | -10.476581 | 1.5126E-07 | 1.1E-05 |
| SSC4D | 1.77275445 | 3.27778528 | 10.4730178 | 1.5185E-07 | 1.1E-05 |
| SURF4 | -0.7537553 | 7.81025324 | -10.461916 | 1.5368E-07 | 1.108E-05 |
| NT5DC3 | -0.9126317 | 5.73460414 | -10.456618 | 1.5456E-07 | 1.1091E-05 |
| APPL1 | -0.8300167 | 6.62762449 | -10.411588 | 1.6227E-07 | 1.159E-05 |
| KNOP1 | -0.9690518 | 5.55289829 | -10.406812 | 1.6312E-07 | 1.1595E-05 |
| B4GALT5 | 0.80298412 | 7.17425774 | 10.3866381 | 1.6673E-07 | 1.1743E-05 |
| CLIP4 | 0.9745809 | 5.2347929 | 10.3865161 | 1.6675E-07 | 1.1743E-05 |
| MTMR12 | -0.9234137 | 5.78012207 | -10.381342 | 1.6769E-07 | 1.1754E-05 |
| EBF3 | 1.20561652 | 4.159933 | 10.3727033 | 1.6927E-07 | 1.1811E-05 |

| | | | | | |
|---|---|---|---|---|---|
| SRSF4 | 0.83991444 | 7.1081224 | 10.3431864 | 1.7479E-07 | 1.214E-05 |
| MAPK1 | 0.72358146 | 8.23095985 | 10.3294006 | 1.7744E-07 | 1.2214E-05 |
| TNFAIP8L1 | -0.9559923 | 5.39059715 | -10.32928 | 1.7746E-07 | 1.2214E-05 |
| ITSN1 | -0.9266946 | 5.54454698 | -10.323023 | 1.7868E-07 | 1.2242E-05 |
| SLC39A11 | -1.0284734 | 4.9486687 | -10.302178 | 1.8279E-07 | 1.2415E-05 |
| MEIOC | -1.0960963 | 4.53253677 | -10.301893 | 1.8285E-07 | 1.2415E-05 |
| C1orf53 | 1.81271393 | 2.97519966 | 10.2934745 | 1.8454E-07 | 1.2474E-05 |
| MELTF | -1.0441913 | 4.88073663 | -10.283404 | 1.8659E-07 | 1.2556E-05 |
| NOL3 | -1.2385098 | 4.03112967 | -10.261751 | 1.9107E-07 | 1.2801E-05 |
| CTSL | -0.8601024 | 6.19796929 | -10.236861 | 1.9636E-07 | 1.2979E-05 |
| HBP1 | 1.06775607 | 5.8805481 | 10.2345448 | 1.9686E-07 | 1.2979E-05 |
| RAB18 | -0.8835747 | 5.76937712 | -10.234016 | 1.9697E-07 | 1.2979E-05 |
| AHNAK | -1.4784001 | 7.37490848 | -10.233143 | 1.9716E-07 | 1.2979E-05 |
| SPECC1L | 0.98540279 | 7.34597254 | 10.2258849 | 1.9874E-07 | 1.3027E-05 |
| PYCR1 | -0.7335019 | 7.94274768 | -10.207324 | 2.0284E-07 | 1.3233E-05 |
| C17orf51 | -0.8524239 | 6.19054853 | -10.203735 | 2.0365E-07 | 1.3233E-05 |
| OPA1 | -0.7704564 | 7.21290993 | -10.179407 | 2.0918E-07 | 1.3532E-05 |
| EIF4E2 | 0.92199537 | 6.14051235 | 10.1757832 | 2.1002E-07 | 1.3532E-05 |
| ZNF24 | -0.7821243 | 6.8904801 | -10.170009 | 2.1136E-07 | 1.356E-05 |
| MFSD6 | -1.3503748 | 3.31717204 | -10.148623 | 2.1642E-07 | 1.3826E-05 |
| ADGRL3 | 1.2461337 | 3.63866624 | 10.1124154 | 2.2528E-07 | 1.4332E-05 |
| RAD51D | -1.064499 | 4.42961124 | -10.090944 | 2.3071E-07 | 1.4606E-05 |
| RHOA | -0.7148812 | 7.93800217 | -10.087417 | 2.3162E-07 | 1.4606E-05 |
| PGGHG | -1.0275914 | 5.69221052 | -10.084047 | 2.3249E-07 | 1.4606E-05 |
| STS | -2.0799053 | 1.59168549 | -10.07485 | 2.3488E-07 | 1.4695E-05 |
| MITF | 0.94743316 | 5.4946517 | 10.0593213 | 2.3897E-07 | 1.489E-05 |
| TTC6 | 1.31206785 | 3.49237786 | 10.0548886 | 2.4016E-07 | 1.4903E-05 |
| ERVMER34-1 | -1.1264369 | 4.11188922 | -10.044731 | 2.4289E-07 | 1.5011E-05 |
| KATNA1 | -0.9749626 | 5.04133611 | -10.007509 | 2.5321E-07 | 1.5584E-05 |
| CDK6 | 0.94337192 | 7.94364534 | 9.99437 | 2.5696E-07 | 1.5669E-05 |
| FAM50B | 1.29437615 | 4.08573995 | 9.99209008 | 2.5761E-07 | 1.5669E-05 |
| WDR20 | 0.82116001 | 6.35977059 | 9.99185298 | 2.5768E-07 | 1.5669E-05 |
| SNX12 | -0.8648468 | 5.8899655 | -9.9874083 | 2.5897E-07 | 1.5684E-05 |
| ALS2CL | -1.75868 | 2.26311811 | -9.981615 | 2.6066E-07 | 1.5723E-05 |
| PAPLN | 2.81600751 | 0.41373612 | 9.97263255 | 2.6329E-07 | 1.582E-05 |
| MLF2 | -0.8009744 | 6.86402086 | -9.9677564 | 2.6474E-07 | 1.5844E-05 |
| ADORA2B | -1.0012598 | 5.21002273 | -9.9414901 | 2.7267E-07 | 1.6254E-05 |
| IGSF8 | -0.9423487 | 5.17369317 | -9.937875 | 2.7378E-07 | 1.6256E-05 |
| TRAK2 | -0.9063024 | 6.39841444 | -9.9121071 | 2.8184E-07 | 1.6669E-05 |
| SMAP2 | -0.9195063 | 5.73181567 | -9.8991663 | 2.8598E-07 | 1.6848E-05 |
| EGR1 | 1.37156257 | 6.34213728 | 9.86623954 | 2.9682E-07 | 1.7419E-05 |
| BEND4 | 0.84234772 | 5.91038207 | 9.84806069 | 3.0299E-07 | 1.7713E-05 |
| DUSP16 | 0.82441337 | 6.18508094 | 9.84028288 | 3.0568E-07 | 1.7744E-05 |
| AFAP1 | -1.0173554 | 4.60127588 | -9.8397482 | 3.0586E-07 | 1.7744E-05 |
| ZDHHC9 | -1.0128701 | 4.6481464 | -9.8325556 | 3.0837E-07 | 1.7791E-05 |

| | | | | | |
|---|---|---|---|---|---|
| VWA1 | -1.105663 | 4.35199692 | -9.8306443 | 3.0904E-07 | 1.7791E-05 |
| PRR5L | -1.1420768 | 4.04073756 | -9.820991 | 3.1244E-07 | 1.7919E-05 |
| DDX21 | -0.6153883 | 9.56811232 | -9.810511 | 3.1618E-07 | 1.8065E-05 |
| NAT9 | -0.8197356 | 6.17124892 | -9.8060376 | 3.178E-07 | 1.8089E-05 |
| CDKN1B | 0.8544321 | 5.76096389 | 9.80068175 | 3.1974E-07 | 1.8132E-05 |
| SMIM19 | 0.97220587 | 4.79094923 | 9.78727879 | 3.2465E-07 | 1.8342E-05 |
| SREK1 | 0.79082853 | 6.64116039 | 9.77644129 | 3.2868E-07 | 1.8488E-05 |
| KIAA1211 | -0.9359524 | 5.05755458 | -9.7737966 | 3.2968E-07 | 1.8488E-05 |
| SHC2 | 1.15341226 | 4.89426159 | 9.73954295 | 3.4282E-07 | 1.9034E-05 |
| FAM129A | -0.8179001 | 6.30933575 | -9.7392196 | 3.4295E-07 | 1.9034E-05 |
| TOM1L2 | 0.88356447 | 5.59569923 | 9.72906088 | 3.4695E-07 | 1.9034E-05 |
| EIF2S1 | -0.7022846 | 8.14937379 | -9.7288903 | 3.4702E-07 | 1.9034E-05 |
| CD9 | -1.0097432 | 4.52620934 | -9.7276487 | 3.4752E-07 | 1.9034E-05 |
| LINC01852 | 0.98350226 | 5.07636917 | 9.72752703 | 3.4756E-07 | 1.9034E-05 |
| RIMKLA | -1.9028217 | 4.89955399 | -9.7257589 | 3.4827E-07 | 1.9034E-05 |
| MYO1C | -0.739334 | 7.91071902 | -9.7220618 | 3.4974E-07 | 1.9034E-05 |
| SYTL2 | 1.45463569 | 2.81482553 | 9.71959462 | 3.5073E-07 | 1.9034E-05 |
| UFL1 | 0.83452035 | 5.77532162 | 9.71282057 | 3.5346E-07 | 1.9114E-05 |
| SCP2 | -0.8792452 | 6.35649572 | -9.7094344 | 3.5484E-07 | 1.912E-05 |
| B4GALT4 | -1.1576143 | 3.89717914 | -9.6889914 | 3.6325E-07 | 1.9504E-05 |
| SEC24D | -1.3682645 | 3.16339033 | -9.6805913 | 3.6677E-07 | 1.9623E-05 |
| AQR | 0.74899091 | 7.24836504 | 9.65348982 | 3.7838E-07 | 2.0173E-05 |
| KIDINS220 | -0.7949456 | 6.67061425 | -9.6401256 | 3.8425E-07 | 2.0414E-05 |
| UBL4A | -0.9408961 | 5.3767357 | -9.6328322 | 3.8749E-07 | 2.0514E-05 |
| HSP90AB1 | -0.559711 | 10.6321617 | -9.6271715 | 3.9003E-07 | 2.0577E-05 |
| GAS2L3 | -0.8200549 | 5.92407658 | -9.612761 | 3.9657E-07 | 2.0849E-05 |
| LZTFL1 | 1.19972032 | 3.71447848 | 9.59985518 | 4.0253E-07 | 2.1089E-05 |
| ZNF845 | -0.9736494 | 4.54787233 | -9.5902949 | 4.07E-07 | 2.125E-05 |
| PNMA2 | 0.90607228 | 5.57178028 | 9.5836306 | 4.1015E-07 | 2.1275E-05 |
| TSHZ1 | 0.90916911 | 5.11274009 | 9.58334172 | 4.1029E-07 | 2.1275E-05 |
| ENAH | 0.60117475 | 9.64858161 | 9.58012268 | 4.1182E-07 | 2.1281E-05 |
| RCBTB2 | 1.10605064 | 4.29504148 | 9.56710626 | 4.1808E-07 | 2.1531E-05 |
| ISM2 | -1.6668596 | 3.23242201 | -9.5623406 | 4.204E-07 | 2.1577E-05 |
| GPC4 | -1.4781503 | 2.65554719 | -9.5557541 | 4.2362E-07 | 2.1669E-05 |
| FOXF2 | -1.1554106 | 3.89116748 | -9.5524014 | 4.2527E-07 | 2.168E-05 |
| PRUNE1 | -0.9135559 | 5.05844194 | -9.5424075 | 4.3023E-07 | 2.1748E-05 |
| FANCC | -0.7729501 | 6.78476209 | -9.5411986 | 4.3084E-07 | 2.1748E-05 |
| SLC7A11 | -0.6521041 | 8.45397543 | -9.5410369 | 4.3092E-07 | 2.1748E-05 |
| HEY1 | 1.0941273 | 3.87687079 | 9.52891793 | 4.3703E-07 | 2.1967E-05 |
| PSMA5 | -0.6903802 | 7.71879391 | -9.5267087 | 4.3815E-07 | 2.1967E-05 |
| MOK | 1.05152205 | 4.58897035 | 9.5217625 | 4.4068E-07 | 2.2021E-05 |
| ACTR6 | -0.9795279 | 4.76260201 | -9.5038131 | 4.4999E-07 | 2.2412E-05 |
| G6PC3 | -0.9957835 | 4.54820644 | -9.4937884 | 4.5528E-07 | 2.2601E-05 |
| MYB | 1.19691801 | 3.53201446 | 9.49011388 | 4.5723E-07 | 2.2624E-05 |
| LRP8 | -0.8998149 | 5.61291668 | -9.4719383 | 4.6703E-07 | 2.3034E-05 |

| | | | | | |
|---|---|---|---|---|---|
| WASHC4 | -0.7614064 | 6.94513626 | -9.4685058 | 4.6891E-07 | 2.3051E-05 |
| MAST2 | -0.7777822 | 6.31077708 | -9.4475545 | 4.8054E-07 | 2.3547E-05 |
| ZNF518B | -0.7308488 | 6.98214037 | -9.4426935 | 4.8328E-07 | 2.3605E-05 |
| SH2B3 | -0.8854289 | 5.23832507 | -9.4392126 | 4.8526E-07 | 2.3625E-05 |
| GPM6A | 0.67376633 | 7.67689418 | 9.43164413 | 4.8958E-07 | 2.3759E-05 |
| TPM1 | -0.8954237 | 5.12103569 | -9.4040852 | 5.0567E-07 | 2.4402E-05 |
| POLR2G | -0.7512583 | 6.71783638 | -9.4034245 | 5.0606E-07 | 2.4402E-05 |
| CAMK2G | -0.8268496 | 6.0951352 | -9.3834174 | 5.181E-07 | 2.4904E-05 |
| ALDH1A3 | -1.284848 | 3.26691608 | -9.3632806 | 5.3054E-07 | 2.542E-05 |
| ABR | -0.800719 | 6.17925731 | -9.3563929 | 5.3486E-07 | 2.5547E-05 |
| FOS | 1.13385757 | 4.75941701 | 9.34283044 | 5.4349E-07 | 2.5877E-05 |
| GAN | -0.8454543 | 6.16399696 | -9.3401479 | 5.4521E-07 | 2.5878E-05 |
| C2orf72 | -1.2469628 | 3.59891463 | -9.3351212 | 5.4846E-07 | 2.5951E-05 |
| SH3GLB2 | -0.8497895 | 5.58271107 | -9.3117695 | 5.6381E-07 | 2.6594E-05 |
| SCAMP1-AS1 | 1.34334493 | 3.00052298 | 9.29174479 | 5.7735E-07 | 2.7148E-05 |
| NPEPPS | 0.78865896 | 6.18390168 | 9.28306865 | 5.8332E-07 | 2.7344E-05 |
| PEX11B | 1.20052155 | 3.56675735 | 9.27398623 | 5.8964E-07 | 2.7555E-05 |
| GK5 | -0.9033763 | 5.06881157 | -9.258453 | 6.0062E-07 | 2.7981E-05 |
| RGS5 | -0.7359045 | 6.8752768 | -9.2550381 | 6.0306E-07 | 2.8009E-05 |
| MYO19 | -0.7130715 | 7.15462725 | -9.2366985 | 6.1637E-07 | 2.854E-05 |
| SDC4 | 1.05644206 | 4.45245013 | 9.23239106 | 6.1954E-07 | 2.8599E-05 |
| TSPAN12 | 0.93242062 | 4.63945946 | 9.22581143 | 6.2442E-07 | 2.8657E-05 |
| INSM2 | 2.11473089 | 1.19745617 | 9.22558208 | 6.2459E-07 | 2.8657E-05 |
| RAB14 | -0.8291414 | 5.58139499 | -9.2015763 | 6.4273E-07 | 2.9401E-05 |
| LCA5 | -0.757647 | 6.24206544 | -9.1958598 | 6.4714E-07 | 2.9513E-05 |
| LINC00294 | -1.6131957 | 2.08662928 | -9.1890499 | 6.5243E-07 | 2.9665E-05 |
| TLDC1 | -0.8652874 | 5.1769086 | -9.1774068 | 6.6158E-07 | 2.995E-05 |
| MTR | -0.7341541 | 6.92992419 | -9.1743888 | 6.6397E-07 | 2.995E-05 |
| RBAK | -0.8581315 | 5.74138602 | -9.1711578 | 6.6654E-07 | 2.995E-05 |
| TRIM65 | -0.7191008 | 7.20537739 | -9.1710677 | 6.6662E-07 | 2.995E-05 |
| ERFE | -1.1487456 | 3.70628896 | -9.1612773 | 6.7448E-07 | 3.0075E-05 |
| RASSF8 | -0.7157123 | 6.74657025 | -9.1611866 | 6.7455E-07 | 3.0075E-05 |
| SH3BP2 | -1.5163766 | 2.98740429 | -9.1590435 | 6.7629E-07 | 3.0075E-05 |
| SLC25A24 | -0.7206983 | 6.94747993 | -9.1555221 | 6.7915E-07 | 3.0075E-05 |
| COG2 | 0.81115065 | 5.66758194 | 9.15530335 | 6.7933E-07 | 3.0075E-05 |
| BICC1 | -0.7557933 | 6.44168744 | -9.14326 | 6.8921E-07 | 3.0424E-05 |
| MVK | -0.8496478 | 5.76492599 | -9.1200367 | 7.0871E-07 | 3.1194E-05 |
| CPT1C | -0.7944063 | 5.81879063 | -9.1056578 | 7.2108E-07 | 3.1646E-05 |
| MAP3K7 | -0.7262741 | 6.73285993 | -9.0978561 | 7.2788E-07 | 3.1852E-05 |
| ITPR3 | -0.7374977 | 6.68154369 | -9.0662037 | 7.5621E-07 | 3.2997E-05 |
| FAM217B | -0.7533936 | 6.02801876 | -9.063555 | 7.5864E-07 | 3.3007E-05 |
| SAPCD2 | -0.8003195 | 5.98224754 | -9.0492671 | 7.7185E-07 | 3.3413E-05 |
| LRRC59 | -0.6088586 | 8.64428807 | -9.0487177 | 7.7237E-07 | 3.3413E-05 |
| RABEP1 | 0.84195035 | 6.9537544 | 9.03397284 | 7.8627E-07 | 3.3899E-05 |
| HIVEP2 | 0.98820849 | 5.09877701 | 9.03206202 | 7.881E-07 | 3.3899E-05 |

| | | | | | |
|---|---|---|---|---|---|
| NF2 | 0.76414268 | 7.07816477 | 9.02716525 | 7.9279E-07 | 3.4004E-05 |
| PDCD10 | -0.8272364 | 5.4144439 | -9.0216904 | 7.9806E-07 | 3.4041E-05 |
| NIN | -0.7313755 | 7.76095112 | -9.0213046 | 7.9844E-07 | 3.4041E-05 |
| IVD | -0.9431687 | 5.82759968 | -9.0192935 | 8.0039E-07 | 3.4041E-05 |
| ZNF28 | -0.8964048 | 4.67738867 | -9.007937 | 8.1149E-07 | 3.4417E-05 |
| PPP2R5A | 0.73850802 | 6.2963491 | 9.00326854 | 8.161E-07 | 3.4516E-05 |
| IFFO2 | -1.1514931 | 3.53225746 | -8.9936965 | 8.2565E-07 | 3.4784E-05 |
| RPS6KA1 | -0.7554272 | 6.23240522 | -8.9922996 | 8.2705E-07 | 3.4784E-05 |
| INF2 | -0.7793158 | 5.82679097 | -8.9875638 | 8.3182E-07 | 3.4888E-05 |
| SLC2A3 | -0.8060459 | 7.25710282 | -8.9738453 | 8.4582E-07 | 3.5305E-05 |
| GNPTAB | 0.80161562 | 5.69289021 | 8.97326543 | 8.4642E-07 | 3.5305E-05 |
| DEPDC1 | -0.7054362 | 6.78780788 | -8.9695569 | 8.5025E-07 | 3.5367E-05 |
| MAP4 | -0.7440586 | 6.37528528 | -8.9488821 | 8.7195E-07 | 3.617E-05 |
| MPI | -0.8243471 | 6.31047737 | -8.9435353 | 8.7765E-07 | 3.6307E-05 |
| CPSF2 | -0.6117905 | 8.30485266 | -8.9341946 | 8.8772E-07 | 3.6624E-05 |
| QSOX2 | -0.7225651 | 6.57290058 | -8.9291412 | 8.9321E-07 | 3.6712E-05 |
| LPIN3 | -1.5075617 | 3.1433597 | -8.9277651 | 8.9472E-07 | 3.6712E-05 |
| CLCN3 | 0.66217336 | 7.42320074 | 8.91426347 | 9.0961E-07 | 3.7196E-05 |
| B4GALNT4 | -0.9490192 | 5.39909815 | -8.912636 | 9.1142E-07 | 3.7196E-05 |
| ZC3H15 | -0.6477359 | 8.1208712 | -8.9103112 | 9.1402E-07 | 3.7202E-05 |
| LSM11 | -0.8762794 | 5.11340133 | -8.9060849 | 9.1876E-07 | 3.7221E-05 |
| CLDN11 | 0.72580615 | 6.48104122 | 8.90552158 | 9.1939E-07 | 3.7221E-05 |
| SYNGR3 | -1.0287015 | 4.38923267 | -8.8902422 | 9.3676E-07 | 3.7823E-05 |
| MEIS1 | 0.77837068 | 6.09723392 | 8.88470288 | 9.4315E-07 | 3.7979E-05 |
| DLX1 | 0.66400569 | 7.39988931 | 8.86192531 | 9.6989E-07 | 3.8952E-05 |
| IMPACT | 0.75973336 | 5.97215025 | 8.85195178 | 9.8185E-07 | 3.9329E-05 |
| A2M | -0.7774011 | 6.06546825 | -8.8484501 | 9.8609E-07 | 3.9394E-05 |
| PIK3R3 | 0.74585 | 6.57066853 | 8.84129449 | 9.9481E-07 | 3.9638E-05 |
| AL136964.1 | 2.01260702 | 0.78736573 | 8.83692098 | 1.0002E-06 | 3.9747E-05 |
| TXNDC15 | -0.8030862 | 5.423679 | -8.832418 | 1.0057E-06 | 3.9864E-05 |
| TRAF3IP2-AS1 | 1.20550714 | 3.23061686 | 8.82525907 | 1.0147E-06 | 4.0112E-05 |
| GALK1 | -0.7568833 | 6.15597013 | -8.8152459 | 1.0273E-06 | 4.0505E-05 |
| HK1 | -0.7073537 | 8.06190663 | -8.8112564 | 1.0323E-06 | 4.0599E-05 |
| ZCCHC14 | 0.69291695 | 6.77716074 | 8.80052757 | 1.0461E-06 | 4.1033E-05 |
| MFSD2A | -1.1175421 | 3.80794765 | -8.7931456 | 1.0557E-06 | 4.1302E-05 |
| STOX2 | 0.81698235 | 5.48488721 | 8.78554154 | 1.0656E-06 | 4.1585E-05 |
| LYRM7 | 0.79778351 | 5.20748122 | 8.77487786 | 1.0798E-06 | 4.2029E-05 |
| JAK3 | -1.2559551 | 3.24290087 | -8.7557056 | 1.1058E-06 | 4.2929E-05 |
| SLC6A6 | -0.8716163 | 6.14965186 | -8.7503528 | 1.1131E-06 | 4.3104E-05 |
| LMO4 | 0.70926726 | 6.50757215 | 8.7448651 | 1.1207E-06 | 4.3288E-05 |
| STX6 | 0.69819576 | 6.54050938 | 8.71457283 | 1.1637E-06 | 4.478E-05 |
| ALX4 | -0.8523516 | 5.00521109 | -8.7132035 | 1.1657E-06 | 4.478E-05 |
| SPATS2L | -0.8531468 | 4.84365123 | -8.7114646 | 1.1682E-06 | 4.478E-05 |
| SEMA3F | -0.974086 | 4.19990301 | -8.7042889 | 1.1787E-06 | 4.5067E-05 |
| SMO | 0.69971133 | 6.71598145 | 8.69494441 | 1.1925E-06 | 4.548E-05 |

| | | | | | |
|---|---|---|---|---|---|
| ZBTB7A | -0.8760275 | 4.84660303 | -8.6924677 | 1.1962E-06 | 4.5506E-05 |
| GOLM1 | -0.6866534 | 7.20021984 | -8.6882098 | 1.2026E-06 | 4.5535E-05 |
| MPZL1 | -0.6735278 | 7.0089458 | -8.6878303 | 1.2031E-06 | 4.5535E-05 |
| RFXANK | -0.7238396 | 6.33157459 | -8.6859421 | 1.206E-06 | 4.5535E-05 |
| FAM76A | 0.89286846 | 4.44553864 | 8.67629601 | 1.2206E-06 | 4.5971E-05 |
| PTBP2 | 0.80947045 | 5.27766523 | 8.66969408 | 1.2307E-06 | 4.6216E-05 |
| GFPT2 | -1.0804368 | 3.8169059 | -8.66807 | 1.2332E-06 | 4.6216E-05 |
| PIK3CB | -1.2078076 | 3.00393 | -8.6614565 | 1.2434E-06 | 4.6266E-05 |
| POFUT1 | -0.8077751 | 6.76586963 | -8.6606749 | 1.2446E-06 | 4.6266E-05 |
| CA12 | -0.9845505 | 4.39564115 | -8.6597508 | 1.246E-06 | 4.6266E-05 |
| IMPAD1 | -0.6713942 | 6.82284736 | -8.6591971 | 1.2469E-06 | 4.6266E-05 |
| USP32 | -0.650367 | 7.34175243 | -8.6573679 | 1.2498E-06 | 4.6266E-05 |
| LYN | -0.8300875 | 5.13240282 | -8.6487226 | 1.2633E-06 | 4.6655E-05 |
| BBX | 0.69832658 | 6.63682909 | 8.63574629 | 1.284E-06 | 4.7261E-05 |
| TNNT1 | -0.7576457 | 5.83400501 | -8.6345127 | 1.286E-06 | 4.7261E-05 |
| ZMPSTE24 | -0.686076 | 6.66463093 | -8.6201629 | 1.3094E-06 | 4.8003E-05 |
| EXOSC6 | -0.7099963 | 6.46604758 | -8.6151092 | 1.3177E-06 | 4.8129E-05 |
| ATXN3 | 0.78421328 | 5.27926346 | 8.61421452 | 1.3192E-06 | 4.8129E-05 |
| PRKD1 | 0.78108177 | 6.17504804 | 8.61126764 | 1.3241E-06 | 4.8191E-05 |
| TNRC6A | 0.84207086 | 6.69986692 | 8.60458853 | 1.3352E-06 | 4.848E-05 |
| CCND2 | -0.514104 | 10.370687 | -8.5995415 | 1.3437E-06 | 4.8672E-05 |
| MTDH | -0.5702959 | 8.63707167 | -8.5874136 | 1.3643E-06 | 4.9302E-05 |
| AAED1 | -0.7762652 | 5.40218539 | -8.5732776 | 1.3888E-06 | 5.0067E-05 |
| PBX1 | 1.24748247 | 3.04423675 | 8.56367405 | 1.4057E-06 | 5.0556E-05 |
| NACC2 | -0.7160018 | 6.16601302 | -8.5432346 | 1.4424E-06 | 5.1654E-05 |
| MNS1 | 0.94511169 | 4.25423017 | 8.54288549 | 1.4431E-06 | 5.1654E-05 |
| MAP1LC3B | -0.8299228 | 5.50206658 | -8.5391536 | 1.4499E-06 | 5.1776E-05 |
| ZDHHC22 | 1.41172802 | 2.39686399 | 8.51491255 | 1.495E-06 | 5.3196E-05 |
| H2AFV | 0.71662755 | 6.33660989 | 8.5132211 | 1.4982E-06 | 5.3196E-05 |
| AC141928.1 | 1.05640494 | 4.14729197 | 8.51216368 | 1.5002E-06 | 5.3196E-05 |
| PGF | -1.0561914 | 3.83099495 | -8.5065603 | 1.5109E-06 | 5.345E-05 |
| HPCAL4 | -1.2680786 | 3.06695743 | -8.494719 | 1.5337E-06 | 5.4131E-05 |
| AC009501.1 | 1.44243811 | 2.32324901 | 8.4896158 | 1.5437E-06 | 5.4231E-05 |
| CYP4F22 | -1.0264585 | 4.00903854 | -8.4896001 | 1.5437E-06 | 5.4231E-05 |
| XKR7 | 1.64996515 | 1.94334991 | 8.48737971 | 1.5481E-06 | 5.4258E-05 |
| SNRNP48 | 0.76346308 | 5.36402663 | 8.4810142 | 1.5606E-06 | 5.4572E-05 |
| EYA2 | -1.1917743 | 3.16183563 | -8.4767193 | 1.5692E-06 | 5.4744E-05 |
| SPART | -0.7203329 | 6.28688354 | -8.473162 | 1.5763E-06 | 5.4805E-05 |
| IQCE | -0.6824474 | 6.93579903 | -8.4722154 | 1.5782E-06 | 5.4805E-05 |
| CLK1 | 0.70268207 | 6.12990123 | 8.46632826 | 1.59E-06 | 5.492E-05 |
| CHGA | 1.00000377 | 4.44113124 | 8.46580689 | 1.5911E-06 | 5.492E-05 |
| ZDHHC23 | -0.9958413 | 4.3456593 | -8.4651693 | 1.5924E-06 | 5.492E-05 |
| ISCA1 | -0.794474 | 5.03224871 | -8.4519549 | 1.6193E-06 | 5.5724E-05 |
| CNTNAP4 | 0.69236794 | 7.23709098 | 8.447332 | 1.6289E-06 | 5.5868E-05 |
| PAQR7 | -0.8018462 | 5.07874507 | -8.4463648 | 1.6309E-06 | 5.5868E-05 |

| | | | | | |
|---|---|---|---|---|---|
| METAP2 | -0.5971629 | 7.84343394 | -8.43476 | 1.6552E-06 | 5.6571E-05 |
| ANO6 | 0.67871411 | 7.23015739 | 8.43075067 | 1.6637E-06 | 5.6697E-05 |
| ITGA5 | -0.7867447 | 6.00452941 | -8.4282926 | 1.6689E-06 | 5.6697E-05 |
| POLR3G | -0.9023444 | 4.273755 | -8.424897 | 1.6761E-06 | 5.6697E-05 |
| SGMS2 | -1.0728197 | 4.20035544 | -8.4243461 | 1.6773E-06 | 5.6697E-05 |
| CARHSP1 | 0.784909 | 5.17129696 | 8.42422113 | 1.6776E-06 | 5.6697E-05 |
| NUP210 | 0.59412578 | 8.35558555 | 8.41877348 | 1.6893E-06 | 5.6965E-05 |
| ID2 | 0.82474666 | 7.96099323 | 8.41612112 | 1.695E-06 | 5.6974E-05 |
| HPSE | -1.046124 | 3.83810713 | -8.4151725 | 1.6971E-06 | 5.6974E-05 |
| SHISA4 | -1.1414346 | 3.39525814 | -8.4117633 | 1.7045E-06 | 5.7095E-05 |
| RASSF7 | -0.8607104 | 4.75305263 | -8.406817 | 1.7153E-06 | 5.733E-05 |
| FAM114A1 | -0.8156075 | 5.12567245 | -8.3985311 | 1.7335E-06 | 5.7813E-05 |
| NUDT3 | -0.7789234 | 5.28940152 | -8.388137 | 1.7567E-06 | 5.8458E-05 |
| P4HA1 | -0.8118452 | 6.93568535 | -8.3858247 | 1.7619E-06 | 5.8502E-05 |
| CYB5A | 0.98053566 | 4.14488181 | 8.37756807 | 1.7806E-06 | 5.8995E-05 |
| KATNAL1 | -0.7733952 | 5.19608157 | -8.3714069 | 1.7947E-06 | 5.9245E-05 |
| C11orf96 | -1.0187685 | 4.38821826 | -8.3708569 | 1.796E-06 | 5.9245E-05 |
| MED1 | -0.8683631 | 6.27073658 | -8.3600516 | 1.821E-06 | 5.994E-05 |
| DUSP5 | -1.2994402 | 2.9253224 | -8.3580127 | 1.8258E-06 | 5.9967E-05 |
| GSE1 | 0.64573233 | 7.04620745 | 8.35127906 | 1.8417E-06 | 6.0356E-05 |
| GTF3C3 | -0.6766772 | 6.87174248 | -8.3407117 | 1.8668E-06 | 6.1049E-05 |
| BRIX1 | -0.6912051 | 6.1533443 | -8.3365216 | 1.8769E-06 | 6.1246E-05 |
| HIST1H2AC | 0.64912949 | 7.02721407 | 8.32652748 | 1.9012E-06 | 6.1904E-05 |
| SOGA1 | -0.6669545 | 6.62582353 | -8.3203467 | 1.9163E-06 | 6.2209E-05 |
| ZHX3 | -0.7539779 | 5.45061316 | -8.3185982 | 1.9207E-06 | 6.2209E-05 |
| IGF2BP3 | 0.80389074 | 5.40174529 | 8.31771434 | 1.9228E-06 | 6.2209E-05 |
| SYNM | -1.5452444 | 2.10655689 | -8.3145828 | 1.9306E-06 | 6.2327E-05 |
| CSRP1 | -0.8113168 | 4.86568627 | -8.3056366 | 1.953E-06 | 6.2915E-05 |
| SEMA4C | -0.6889589 | 6.99877068 | -8.2937226 | 1.9832E-06 | 6.3753E-05 |
| EPPK1 | -1.7155321 | 1.83748474 | -8.2840413 | 2.0081E-06 | 6.4418E-05 |
| VTI1B | 0.67959079 | 6.75656105 | 8.27254209 | 2.0382E-06 | 6.5243E-05 |
| KCTD18 | -0.9323937 | 4.07501505 | -8.2567829 | 2.0801E-06 | 6.6446E-05 |
| RNF6 | -0.7213349 | 6.04413754 | -8.2521592 | 2.0926E-06 | 6.6704E-05 |
| SGMS1 | -0.7019081 | 5.98351321 | -8.2399428 | 2.126E-06 | 6.7626E-05 |
| UTP14A | -0.9516151 | 5.90858853 | -8.2378637 | 2.1317E-06 | 6.7666E-05 |
| SECISBP2 | 0.71696023 | 5.69668105 | 8.22952535 | 2.1549E-06 | 6.8258E-05 |
| LARGE2 | -0.7485034 | 5.48186997 | -8.2148358 | 2.1964E-06 | 6.9373E-05 |
| RNF115 | 0.69802352 | 6.66433267 | 8.21320772 | 2.201E-06 | 6.9373E-05 |
| CHRAC1 | 0.81859568 | 6.47012954 | 8.21222599 | 2.2038E-06 | 6.9373E-05 |
| ATP2C1 | -0.7001816 | 6.03064996 | -8.2098576 | 2.2106E-06 | 6.9442E-05 |
| FBXO27 | -0.931491 | 4.13871326 | -8.1995291 | 2.2405E-06 | 7.0235E-05 |
| ALDH1A1 | -0.6782434 | 8.4723853 | -8.192284 | 2.2617E-06 | 7.0627E-05 |
| YAF2 | 0.85871127 | 5.8042221 | 8.19206813 | 2.2624E-06 | 7.0627E-05 |
| HIST1H4B | 0.68903966 | 6.85990826 | 8.18902074 | 2.2713E-06 | 7.0762E-05 |
| TIGD5 | -0.7282889 | 5.62614579 | -8.1842894 | 2.2854E-06 | 7.0958E-05 |

| | | | | | |
|---|---|---|---|---|---|
| PDCD4-AS1 | 1.55180939 | 1.9779932 | 8.18374544 | 2.287E-06 | 7.0958E-05 |
| GAB2 | 0.70176253 | 5.99435589 | 8.17963993 | 2.2993E-06 | 7.1192E-05 |
| PEX10 | -0.8141749 | 5.30657548 | -8.177154 | 2.3067E-06 | 7.1241E-05 |
| RLIM | -0.6587043 | 6.62871812 | -8.1759872 | 2.3102E-06 | 7.1241E-05 |
| CAT | -0.7325835 | 5.54765208 | -8.1669631 | 2.3376E-06 | 7.1876E-05 |
| MOCOS | -1.4199066 | 2.34151017 | -8.1660512 | 2.3403E-06 | 7.1876E-05 |
| WNT10B | -0.8822201 | 5.0246685 | -8.162877 | 2.3501E-06 | 7.2029E-05 |
| SESN3 | 0.78990599 | 4.78761336 | 8.15788367 | 2.3654E-06 | 7.2353E-05 |
| ETFRF1 | 0.77315767 | 4.68126862 | 8.14520126 | 2.4049E-06 | 7.3282E-05 |
| ZNF385D | -1.121596 | 3.4141772 | -8.1434016 | 2.4106E-06 | 7.3282E-05 |
| CDK7 | -0.8721357 | 4.30214849 | -8.1421837 | 2.4144E-06 | 7.3282E-05 |
| RXRB | 0.76790878 | 5.4526994 | 8.14034409 | 2.4202E-06 | 7.3282E-05 |
| B4GALNT1 | -0.6775251 | 6.29212223 | -8.1398841 | 2.4217E-06 | 7.3282E-05 |
| NACC1 | -0.737722 | 6.58324402 | -8.1388929 | 2.4248E-06 | 7.3282E-05 |
| TMEM56 | -1.091805 | 3.25485493 | -8.1369757 | 2.4309E-06 | 7.33E-05 |
| CECR2 | 0.90565663 | 5.28668761 | 8.13525782 | 2.4364E-06 | 7.33E-05 |
| PPP2R5C | 0.58778145 | 7.67018435 | 8.13413419 | 2.44E-06 | 7.33E-05 |
| ANKIB1 | 0.62227948 | 7.14972571 | 8.13093761 | 2.4502E-06 | 7.3462E-05 |
| RCOR3 | 0.73259656 | 5.59295006 | 8.12800987 | 2.4596E-06 | 7.3598E-05 |
| CSPG5 | -0.8685469 | 4.46817152 | -8.1229121 | 2.476E-06 | 7.3944E-05 |
| AP5Z1 | 0.6508314 | 6.71789669 | 8.11893546 | 2.489E-06 | 7.4184E-05 |
| PER3 | 0.77223955 | 5.3392201 | 8.11607108 | 2.4983E-06 | 7.4291E-05 |
| MORN2 | 0.85513076 | 4.70590215 | 8.11342357 | 2.507E-06 | 7.4291E-05 |
| DNAH7 | 0.77544604 | 5.00179047 | 8.11188429 | 2.5121E-06 | 7.4291E-05 |
| SLC43A3 | -1.219235 | 2.78729605 | -8.1118429 | 2.5122E-06 | 7.4291E-05 |
| AZI2 | 0.73533599 | 5.36374496 | 8.10605589 | 2.5313E-06 | 7.4499E-05 |
| TAF4B | 0.80291669 | 4.74645182 | 8.10570494 | 2.5325E-06 | 7.4499E-05 |
| KLC2 | -0.7043413 | 5.76619274 | -8.1052576 | 2.534E-06 | 7.4499E-05 |
| CIR1 | 0.86469734 | 4.30164912 | 8.10266362 | 2.5426E-06 | 7.4608E-05 |
| BCL6 | 1.04690656 | 3.5490624 | 8.09340999 | 2.5737E-06 | 7.5373E-05 |
| PLCB3 | -0.7361904 | 7.03013378 | -8.0901838 | 2.5846E-06 | 7.5481E-05 |
| TIMELESS | 0.56873464 | 8.17238164 | 8.0893774 | 2.5873E-06 | 7.5481E-05 |
| OLFM3 | 0.58475488 | 7.7207739 | 8.08394451 | 2.6058E-06 | 7.5875E-05 |
| CHAC2 | -0.9473993 | 4.05981504 | -8.0819943 | 2.6125E-06 | 7.5924E-05 |
| CAPN1 | -0.6648269 | 6.95327741 | -8.0698778 | 2.6545E-06 | 7.6995E-05 |
| COPS7A | 0.70827969 | 6.16012732 | 8.06658743 | 2.666E-06 | 7.7181E-05 |
| RSL24D1 | 0.63277663 | 6.70170161 | 8.05799749 | 2.6963E-06 | 7.7909E-05 |
| STARD4 | -0.625461 | 7.28312248 | -8.0541164 | 2.7101E-06 | 7.8061E-05 |
| SLC25A5 | -0.5690457 | 8.09671418 | -8.0536248 | 2.7119E-06 | 7.8061E-05 |
| LRIG1 | -1.0533226 | 3.5056543 | -8.0506947 | 2.7223E-06 | 7.8122E-05 |
| KCNA1 | 1.46566291 | 2.22702551 | 8.05015477 | 2.7243E-06 | 7.8122E-05 |
| IDE | -0.6326925 | 6.84767478 | -8.0448118 | 2.7435E-06 | 7.8525E-05 |
| PPP2R5B | -1.0062394 | 5.15571241 | -8.0384221 | 2.7667E-06 | 7.8953E-05 |
| ADAM19 | -0.7741375 | 5.21812429 | -8.0378235 | 2.7689E-06 | 7.8953E-05 |
| LINC00467 | 0.83191285 | 4.73966083 | 8.03585688 | 2.7761E-06 | 7.9009E-05 |

| | | | | | |
|---|---|---|---|---|---|
| ADD3 | 0.6256795 | 6.89647853 | 8.02846348 | 2.8033E-06 | 7.9574E-05 |
| PCYOX1L | -1.0067753 | 3.88544404 | -8.0276129 | 2.8065E-06 | 7.9574E-05 |
| OPRL1 | -1.0880627 | 4.21494581 | -8.0241622 | 2.8193E-06 | 7.9788E-05 |
| ERRFI1 | 0.75260133 | 8.68034164 | 8.02256971 | 2.8252E-06 | 7.9807E-05 |
| GCH1 | -0.8008964 | 4.72946532 | -8.0164846 | 2.848E-06 | 8.0301E-05 |
| CREBRF | 1.10706598 | 3.18413571 | 7.99719402 | 2.9216E-06 | 8.2163E-05 |
| AC009113.1 | 1.01516756 | 3.54045446 | 7.99540559 | 2.9285E-06 | 8.2163E-05 |
| SRPK2 | 0.66974249 | 6.08984589 | 7.99494558 | 2.9303E-06 | 8.2163E-05 |
| NOP14-AS1 | -0.7480237 | 5.30673949 | -7.9864404 | 2.9635E-06 | 8.2786E-05 |
| DNAL4 | 0.97067373 | 3.81302331 | 7.98380193 | 2.9739E-06 | 8.2923E-05 |
| TMEM256 | -1.3195699 | 2.53264875 | -7.9820319 | 2.9809E-06 | 8.2965E-05 |
| LDLR | -0.6098424 | 7.25124384 | -7.9622955 | 3.0599E-06 | 8.501E-05 |
| SRBD1 | 0.73842587 | 6.21266407 | 7.95274466 | 3.099E-06 | 8.5937E-05 |
| TYMP | -0.9137575 | 4.16795735 | -7.9388147 | 3.1569E-06 | 8.7383E-05 |
| ACVR2B | 0.77856111 | 5.73595668 | 7.93710224 | 3.1641E-06 | 8.7423E-05 |
| CNNM4 | -0.7047752 | 5.65170074 | -7.9314433 | 3.188E-06 | 8.7923E-05 |
| HSPA13 | -0.6885064 | 6.02684271 | -7.924751 | 3.2165E-06 | 8.8549E-05 |
| RAB12 | 0.82511768 | 4.48947992 | 7.90875243 | 3.2859E-06 | 9.0293E-05 |
| DISP3 | 1.97417579 | 1.09463524 | 7.90501823 | 3.3023E-06 | 9.0516E-05 |
| AMMECR1L | -0.694977 | 5.75782224 | -7.9038761 | 3.3073E-06 | 9.0516E-05 |
| DAPK1 | 0.68342673 | 6.33004601 | 7.90283533 | 3.3119E-06 | 9.0516E-05 |
| ZNF212 | 0.79761785 | 4.64454992 | 7.89106748 | 3.3644E-06 | 9.1776E-05 |
| NIPA2 | -0.6022162 | 7.31982301 | -7.8897787 | 3.3702E-06 | 9.1776E-05 |
| ATP6V0E1 | 0.62434673 | 6.78381648 | 7.88500901 | 3.3917E-06 | 9.2197E-05 |
| SPHK2 | -1.020212 | 3.50359323 | -7.8824575 | 3.4033E-06 | 9.2301E-05 |
| UTRN | 0.77471109 | 6.74511372 | 7.88148179 | 3.4077E-06 | 9.2301E-05 |
| MAPRE2 | 0.94661915 | 4.93921458 | 7.87956047 | 3.4165E-06 | 9.2363E-05 |
| ANP32E | -0.6381591 | 7.03192674 | -7.8783108 | 3.4222E-06 | 9.2363E-05 |
| COMT | -0.6636755 | 6.17787856 | -7.8765832 | 3.4301E-06 | 9.2412E-05 |
| GPR63 | 0.83824852 | 4.27779641 | 7.87002701 | 3.4603E-06 | 9.3061E-05 |
| TP53 | 1.01307935 | 3.63259195 | 7.86771742 | 3.4711E-06 | 9.3061E-05 |
| PVR | -0.6388531 | 7.10041354 | -7.8673736 | 3.4727E-06 | 9.3061E-05 |
| PRKACB | -0.9372949 | 3.6958924 | -7.8614605 | 3.5003E-06 | 9.3635E-05 |
| N4BP2 | 0.74090103 | 6.18100667 | 7.85133905 | 3.548E-06 | 9.4746E-05 |
| CUBN | 0.78690079 | 5.60798215 | 7.84656426 | 3.5708E-06 | 9.5187E-05 |
| ZNF888 | -0.9647076 | 3.66629572 | -7.8327574 | 3.6376E-06 | 9.6796E-05 |
| EEA1 | 0.71130084 | 6.09831844 | 7.83087455 | 3.6468E-06 | 9.6871E-05 |
| RAB3IL1 | -1.1286451 | 2.99847324 | -7.8253511 | 3.674E-06 | 9.7334E-05 |
| RARB | 0.61810339 | 7.30220343 | 7.82471504 | 3.6771E-06 | 9.7334E-05 |
| SEMA4F | -0.7655002 | 4.94898218 | -7.8202446 | 3.6993E-06 | 9.775E-05 |
| SMIM14 | 0.8589839 | 4.45785659 | 7.81325119 | 3.7342E-06 | 9.8476E-05 |
| HAPLN3 | -1.1763486 | 3.26835482 | -7.809047 | 3.7554E-06 | 9.8476E-05 |
| CRIM1 | -0.7034133 | 6.34702744 | -7.8089179 | 3.756E-06 | 9.8476E-05 |
| GPX8 | -0.6942842 | 5.59566712 | -7.8085117 | 3.7581E-06 | 9.8476E-05 |
| MKRN1 | 0.58180647 | 7.39103371 | 7.80828056 | 3.7593E-06 | 9.8476E-05 |

| | | | | | |
|---|---|---|---|---|---|
| PARD6B | -0.8194002 | 4.38545434 | -7.8054602 | 3.7736E-06 | 9.868E-05 |
| KCNK6 | -1.579402 | 1.96177626 | -7.800694 | 3.7979E-06 | 9.9041E-05 |
| PTP4A3 | -0.783151 | 4.94208419 | -7.7999361 | 3.8017E-06 | 9.9041E-05 |
| SMS | -0.7111428 | 5.48843836 | -7.7989087 | 3.807E-06 | 9.9041E-05 |
| TOMM70 | -0.5915663 | 7.12146215 | -7.7948136 | 3.8281E-06 | 9.9253E-05 |
| CHD6 | 0.65514593 | 6.47975622 | 7.79477168 | 3.8283E-06 | 9.9253E-05 |
| LLGL2 | -0.7947564 | 5.46888745 | -7.7919958 | 3.8426E-06 | 9.9343E-05 |
| SLC15A4 | -0.6958635 | 5.53931418 | -7.7915662 | 3.8448E-06 | 9.9343E-05 |
| COCH | -0.6747808 | 5.7664326 | -7.7882171 | 3.8622E-06 | 9.9622E-05 |
| MOSPD2 | -0.987741 | 3.45622895 | -7.7779278 | 3.9162E-06 | 0.00010084 |
| SGK1 | 1.03007932 | 3.06184077 | 7.77378767 | 3.9381E-06 | 0.00010123 |
| RHCG | 0.75393112 | 5.18911376 | 7.77012061 | 3.9577E-06 | 0.00010156 |
| ATG12 | -0.7525251 | 5.34468913 | -7.7588509 | 4.0184E-06 | 0.00010295 |
| RMI2 | 0.98545883 | 3.59806509 | 7.74492701 | 4.0948E-06 | 0.00010473 |
| KLRG2 | -1.5565561 | 1.81732351 | -7.7408576 | 4.1174E-06 | 0.00010513 |
| CCDC28A | 0.98238422 | 3.46570026 | 7.73703535 | 4.1387E-06 | 0.00010543 |
| EXOSC5 | -0.7625004 | 4.8267111 | -7.7362237 | 4.1433E-06 | 0.00010543 |
| ZMYND19 | -0.7158823 | 5.32146089 | -7.734696 | 4.1519E-06 | 0.00010548 |
| U2SURP | 0.55451384 | 7.90733486 | 7.73110199 | 4.1721E-06 | 0.00010581 |
| SLC2A1 | -1.4534129 | 5.15650236 | -7.7273624 | 4.1933E-06 | 0.00010617 |
| SLITRK5 | 0.66842758 | 6.070538 | 7.72123891 | 4.2283E-06 | 0.00010688 |
| REEP4 | -0.7044687 | 5.39368518 | -7.7174879 | 4.2498E-06 | 0.00010724 |
| JAG1 | -0.8381681 | 4.32556925 | -7.713209 | 4.2746E-06 | 0.00010769 |
| PCMTD1 | 0.88239294 | 4.38769822 | 7.71163319 | 4.2837E-06 | 0.00010774 |
| PSD | 0.86801993 | 4.64531787 | 7.70446819 | 4.3256E-06 | 0.00010849 |
| OLA1 | -0.5960328 | 6.95118935 | -7.7040885 | 4.3278E-06 | 0.00010849 |
| ALKBH8 | -0.7258562 | 5.47749746 | -7.7011074 | 4.3454E-06 | 0.00010875 |
| PAM | 0.5281039 | 8.26439251 | 7.69126631 | 4.4039E-06 | 0.00011003 |
| UPK1A-AS1 | -3.647838 | 0.00711598 | -7.689848 | 4.4124E-06 | 0.00011006 |
| WDTC1 | 0.76654337 | 4.91022811 | 7.67797927 | 4.4842E-06 | 0.00011167 |
| ATXN1 | 0.71872562 | 7.68090284 | 7.67388438 | 4.5093E-06 | 0.00011211 |
| DSTN | -0.6283443 | 6.93890888 | -7.659212 | 4.6003E-06 | 0.00011419 |
| SERPINE1 | -1.3758747 | 2.21604669 | -7.6403349 | 4.7203E-06 | 0.00011649 |
| MSRB1 | -0.8207773 | 4.36135347 | -7.6399592 | 4.7228E-06 | 0.00011649 |
| KDM1B | -0.7297107 | 5.71185943 | -7.6391976 | 4.7277E-06 | 0.00011649 |
| PHLDB1 | -0.8455471 | 4.43141395 | -7.6387991 | 4.7303E-06 | 0.00011649 |
| SHC1 | -0.5881643 | 7.31609264 | -7.6386009 | 4.7315E-06 | 0.00011649 |
| GIPC3 | -1.5450587 | 1.68347318 | -7.6286422 | 4.7964E-06 | 0.00011789 |
| RNF157 | 0.72968458 | 5.43341996 | 7.62349622 | 4.8302E-06 | 0.00011853 |
| CACNA2D1 | 0.70919412 | 5.0201647 | 7.61934912 | 4.8577E-06 | 0.00011883 |
| TIPARP | -0.6808002 | 5.41807245 | -7.6193142 | 4.8579E-06 | 0.00011883 |
| GADD45B | 0.7537691 | 4.73639246 | 7.6161284 | 4.8792E-06 | 0.00011915 |
| CENPH | 0.70589925 | 5.33837501 | 7.61461126 | 4.8893E-06 | 0.00011919 |
| OGA | 0.54920307 | 7.69928682 | 7.61324642 | 4.8985E-06 | 0.00011919 |
| ANKRD52 | -0.6260686 | 6.49102292 | -7.6123587 | 4.9044E-06 | 0.00011919 |

| | | | | | |
|---|---|---|---|---|---|
| TUBA3C | -0.6537506 | 6.6229614 | -7.6029154 | 4.9683E-06 | 0.00012055 |
| TMEM206 | -0.7212799 | 5.18847358 | -7.5930919 | 5.0356E-06 | 0.00012196 |
| CCDC30 | 1.35328147 | 2.04149236 | 7.59208304 | 5.0426E-06 | 0.00012196 |
| KLHL12 | 0.59431451 | 7.07240418 | 7.58910196 | 5.0632E-06 | 0.00012213 |
| DCAF16 | 0.57406554 | 7.13799591 | 7.58876104 | 5.0656E-06 | 0.00012213 |
| PYGO1 | -0.7223803 | 4.97920945 | -7.5874625 | 5.0746E-06 | 0.00012213 |
| ISCU | 0.62649437 | 6.2333048 | 7.58566341 | 5.0872E-06 | 0.00012213 |
| CPAMD8 | 1.99643686 | 0.8947045 | 7.58529625 | 5.0897E-06 | 0.00012213 |
| THUMPD1 | -0.6323583 | 6.11491147 | -7.5822335 | 5.1112E-06 | 0.00012234 |
| TPD52L1 | -0.6819844 | 6.18664933 | -7.5810009 | 5.1198E-06 | 0.00012234 |
| CHMP7 | -0.611141 | 6.54197333 | -7.577311 | 5.1458E-06 | 0.0001227 |
| AUH | 1.08036958 | 2.97225991 | 7.56806358 | 5.2116E-06 | 0.00012407 |
| DESI2 | -0.585332 | 7.49786651 | -7.5616764 | 5.2576E-06 | 0.00012497 |
| NDUFA10 | -0.5680733 | 7.71902997 | -7.5564243 | 5.2957E-06 | 0.00012568 |
| OSBPL3 | -0.7212154 | 5.0357512 | -7.5466535 | 5.3674E-06 | 0.000127 |
| CCDC85A | -0.6290384 | 6.13274487 | -7.5459153 | 5.3729E-06 | 0.000127 |
| LUM | -0.5831909 | 7.32170379 | -7.544676 | 5.3821E-06 | 0.000127 |
| CUL4B | -0.5772494 | 7.22717528 | -7.5442534 | 5.3852E-06 | 0.000127 |
| CDH12 | 0.80941169 | 4.34986959 | 7.54268023 | 5.3969E-06 | 0.00012708 |
| NDUFAF3 | -0.703052 | 5.42201076 | -7.5285405 | 5.5031E-06 | 0.00012918 |
| BTG2 | 1.01848921 | 3.40736253 | 7.52592915 | 5.5229E-06 | 0.00012945 |
| BMP6 | -1.3292402 | 2.31497932 | -7.5231875 | 5.5439E-06 | 0.00012955 |
| LTBP1 | 0.601792 | 7.14395757 | 7.52309029 | 5.5446E-06 | 0.00012955 |
| SNX21 | -0.7182514 | 5.11770818 | -7.5102555 | 5.6437E-06 | 0.00013125 |
| CCDC85B | -0.7187923 | 5.7789341 | -7.510129 | 5.6447E-06 | 0.00013125 |
| VIT | 1.06512193 | 4.2129112 | 7.50818041 | 5.6599E-06 | 0.00013125 |
| STARD8 | -1.2391011 | 2.54824545 | -7.5081052 | 5.6605E-06 | 0.00013125 |
| RAB11A | -0.5990366 | 7.08344231 | -7.5016069 | 5.7116E-06 | 0.00013223 |
| ZNF256 | 0.92762433 | 3.59780979 | 7.49795914 | 5.7405E-06 | 0.0001327 |
| SOWAHC | -0.7259304 | 5.4484225 | -7.4916113 | 5.7911E-06 | 0.00013366 |
| SSR1 | -0.5572398 | 7.3973734 | -7.4884356 | 5.8166E-06 | 0.00013405 |
| CHAC1 | -0.7179646 | 5.7490112 | -7.4801788 | 5.8835E-06 | 0.00013538 |
| PHACTR4 | 0.74796796 | 5.39661038 | 7.47910815 | 5.8922E-06 | 0.00013538 |
| PMEPA1 | -1.097148 | 2.99766436 | -7.4773735 | 5.9064E-06 | 0.00013546 |
| PIBF1 | 0.72435859 | 4.78506676 | 7.4764737 | 5.9138E-06 | 0.00013546 |
| FCHO1 | -0.8233606 | 4.33327177 | -7.4722692 | 5.9483E-06 | 0.00013605 |
| PHF6 | 0.63326076 | 6.16290791 | 7.46849045 | 5.9796E-06 | 0.00013656 |
| SLC49A3 | -1.7289654 | 1.1121094 | -7.4646838 | 6.0112E-06 | 0.00013707 |
| ENDOD1 | -0.6914821 | 5.72440151 | -7.461528 | 6.0376E-06 | 0.00013747 |
| AC006333.2 | 1.09878973 | 2.62478444 | 7.45975992 | 6.0524E-06 | 0.0001376 |
| AC006449.6 | 0.96318297 | 3.52054828 | 7.45538682 | 6.0893E-06 | 0.00013823 |
| FAP | 0.64045747 | 5.85664632 | 7.45244802 | 6.1142E-06 | 0.00013858 |
| MLKL | -0.7919873 | 4.49826513 | -7.4479939 | 6.1521E-06 | 0.00013924 |
| ETV7 | 2.33950674 | -0.1672783 | 7.44287415 | 6.196E-06 | 0.00014002 |
| SLC27A1 | -1.2200237 | 3.39467743 | -7.4412668 | 6.2099E-06 | 0.00014013 |

| | | | | | |
|---|---|---|---|---|---|
| OSBPL5 | -0.8413867 | 4.17779503 | -7.4377964 | 6.2399E-06 | 0.00014045 |
| CLOCK | -0.6877173 | 5.59780422 | -7.4374574 | 6.2429E-06 | 0.00014045 |
| ZNF512B | 0.59058134 | 6.92492686 | 7.43452403 | 6.2684E-06 | 0.0001407 |
| ZNF267 | -0.6327989 | 5.77662667 | -7.4340359 | 6.2726E-06 | 0.0001407 |
| SLC24A1 | 0.84194549 | 3.97920566 | 7.4263511 | 6.3401E-06 | 0.000142 |
| CTHRC1 | 0.63912185 | 6.20891363 | 7.4236178 | 6.3642E-06 | 0.00014233 |
| MAML1 | 0.62282678 | 6.61135927 | 7.42114478 | 6.3862E-06 | 0.00014262 |
| OAS3 | -0.8069824 | 4.42683118 | -7.416117 | 6.4311E-06 | 0.00014341 |
| EFCAB2 | 0.89070905 | 3.61191458 | 7.413748 | 6.4523E-06 | 0.00014367 |
| ZNF621 | -0.6474615 | 5.73957656 | -7.4119937 | 6.4681E-06 | 0.00014381 |
| HHEX | -0.9035993 | 3.62670627 | -7.408772 | 6.4972E-06 | 0.00014424 |
| SPINT1 | -2.1708803 | 0.36186848 | -7.4012367 | 6.5658E-06 | 0.00014533 |
| SNAPC2 | -0.6543568 | 5.7352509 | -7.4011061 | 6.567E-06 | 0.00014533 |
| DICER1 | 0.70009326 | 8.23077625 | 7.4002019 | 6.5753E-06 | 0.00014533 |
| MYCL | -1.0309456 | 3.42396757 | -7.3991904 | 6.5846E-06 | 0.00014533 |
| AC090114.2 | 0.85058155 | 3.91349963 | 7.397513 | 6.6E-06 | 0.00014546 |
| TMEM52 | -1.2794208 | 2.41670533 | -7.3932593 | 6.6393E-06 | 0.00014611 |
| ANP32A | 0.56704642 | 7.9089603 | 7.38891833 | 6.6797E-06 | 0.00014679 |
| SCYL3 | 0.91644148 | 4.32649287 | 7.38676694 | 6.6998E-06 | 0.00014702 |
| GABPB2 | 0.78609069 | 4.42831203 | 7.38401366 | 6.7256E-06 | 0.00014733 |
| TYSND1 | -0.6110053 | 6.79252722 | -7.3831495 | 6.7337E-06 | 0.00014733 |
| NCS1 | -0.6329213 | 6.40069995 | -7.3804284 | 6.7594E-06 | 0.00014768 |
| SLC22A18 | -1.047791 | 3.47683311 | -7.3761961 | 6.7995E-06 | 0.00014834 |
| FAR1 | -0.623759 | 5.88381152 | -7.3749609 | 6.8112E-06 | 0.00014839 |
| PLS3 | -0.5344363 | 8.02859977 | -7.3660271 | 6.8969E-06 | 0.00015004 |
| USH1C | -2.0771749 | 0.3910098 | -7.3607803 | 6.9477E-06 | 0.00015093 |
| WNK3 | -0.7985098 | 4.30571363 | -7.358374 | 6.9712E-06 | 0.00015109 |
| ZNF362 | 0.67958807 | 5.36174297 | 7.35794446 | 6.9754E-06 | 0.00015109 |
| AP1G2 | -0.8478657 | 4.8955248 | -7.3479193 | 7.074E-06 | 0.00015301 |
| RSPH3 | 0.7898624 | 4.3475011 | 7.34291875 | 7.1238E-06 | 0.00015387 |
| HIST2H2AC | 0.56102183 | 7.70278217 | 7.32229275 | 7.333E-06 | 0.00015816 |
| OTULIN | -0.648134 | 5.528843 | -7.3197069 | 7.3597E-06 | 0.00015851 |
| ADAMTSL5 | -1.0165562 | 3.34737254 | -7.310695 | 7.4535E-06 | 0.0001603 |
| CPLX2 | 1.68120769 | 1.09196589 | 7.30018834 | 7.5645E-06 | 0.00016246 |
| ZNF813 | -0.7857157 | 4.15450798 | -7.2939041 | 7.6317E-06 | 0.00016367 |
| MROH6 | -1.042397 | 3.32271354 | -7.2857625 | 7.7198E-06 | 0.00016516 |
| CHSY1 | -0.6562038 | 6.5109284 | -7.2854802 | 7.7228E-06 | 0.00016516 |
| RGS16 | -0.8307098 | 4.02895614 | -7.27689 | 7.8169E-06 | 0.00016693 |
| NELFB | -0.603721 | 6.30641356 | -7.2684783 | 7.9103E-06 | 0.00016866 |
| SLC16A13 | -1.442643 | 1.83991525 | -7.2671022 | 7.9256E-06 | 0.00016866 |
| PIAS2 | 0.6635181 | 5.29404593 | 7.26662798 | 7.931E-06 | 0.00016866 |
| GNAL | 0.76610185 | 4.45636617 | 7.26259327 | 7.9763E-06 | 0.00016938 |
| DYNC1I1 | 0.6529161 | 5.40591961 | 7.2568882 | 8.0408E-06 | 0.00017041 |
| GOT2 | -0.5428006 | 7.44860987 | -7.2563265 | 8.0472E-06 | 0.00017041 |
| TTC39B | -1.0712528 | 3.48939698 | -7.2499796 | 8.1197E-06 | 0.0001717 |

| | | | | | |
|---|---|---|---|---|---|
| RIOK1 | -0.6011146 | 6.52001206 | -7.2452114 | 8.1746E-06 | 0.00017235 |
| RPP25 | 0.58424738 | 6.81777062 | 7.2438137 | 8.1908E-06 | 0.00017235 |
| ARHGEF3 | 0.76172591 | 4.62374205 | 7.24351948 | 8.1942E-06 | 0.00017235 |
| AP002360.1 | 0.85127389 | 4.12369208 | 7.24337008 | 8.196E-06 | 0.00017235 |
| DISP2 | -0.7933832 | 4.5153281 | -7.2414846 | 8.2178E-06 | 0.00017257 |
| ZNF441 | 0.73194073 | 4.58728205 | 7.23911716 | 8.2454E-06 | 0.00017291 |
| FZD9 | -1.5096471 | 1.77658134 | -7.2300461 | 8.352E-06 | 0.00017491 |
| CPEB2 | 1.11641791 | 2.58162183 | 7.22894562 | 8.365E-06 | 0.00017494 |
| FAM49B | -0.5491183 | 7.21699356 | -7.2275462 | 8.3816E-06 | 0.00017504 |
| EPHB3 | -1.3664108 | 2.29589325 | -7.2243639 | 8.4195E-06 | 0.00017544 |
| KDM5A | 0.64342433 | 7.0787966 | 7.22398878 | 8.424E-06 | 0.00017544 |
| ARL4A | 0.54441847 | 7.56548233 | 7.22290885 | 8.4369E-06 | 0.00017547 |
| ZNF286A | 0.64912027 | 5.37334884 | 7.22060244 | 8.4645E-06 | 0.00017547 |
| DNAJC10 | 0.54572636 | 7.08699127 | 7.22004304 | 8.4712E-06 | 0.00017547 |
| SCFD1 | 0.58536285 | 6.69485191 | 7.22001605 | 8.4715E-06 | 0.00017547 |
| PHF21B | 0.67368455 | 5.26869707 | 7.21427392 | 8.5408E-06 | 0.00017666 |
| FAM189B | -0.6667005 | 5.65660794 | -7.2087688 | 8.6078E-06 | 0.0001778 |
| DCP2 | 0.51804153 | 7.78056989 | 7.20301345 | 8.6784E-06 | 0.0001789 |
| COQ7 | 0.77880984 | 4.64969192 | 7.20253065 | 8.6843E-06 | 0.0001789 |
| DUSP19 | 0.82331565 | 3.8117708 | 7.20102126 | 8.7029E-06 | 0.00017904 |
| TTYH2 | -0.7617179 | 4.50633824 | -7.1999993 | 8.7156E-06 | 0.00017905 |
| ETFBKMT | 1.45072388 | 1.60798421 | 7.19619123 | 8.7629E-06 | 0.00017966 |
| KLF13 | 0.61377799 | 7.45831714 | 7.19492161 | 8.7787E-06 | 0.00017966 |
| SLC22A31 | -1.4056462 | 2.98995917 | -7.1939915 | 8.7903E-06 | 0.00017966 |
| AK2 | -0.6138569 | 7.65906282 | -7.1938284 | 8.7923E-06 | 0.00017966 |
| TTC8 | 0.57503593 | 6.87244998 | 7.1889196 | 8.8539E-06 | 0.00018067 |
| CXXC1 | 0.60325236 | 6.15668301 | 7.18575566 | 8.8938E-06 | 0.00018124 |
| HSPA5 | -0.5936684 | 7.66958253 | -7.1820435 | 8.9409E-06 | 0.00018195 |
| SMURF2 | 0.5854366 | 6.44306064 | 7.17972163 | 8.9705E-06 | 0.00018231 |
| NECAB3 | -0.6738506 | 5.81975048 | -7.1776885 | 8.9964E-06 | 0.00018259 |
| LYAR | -0.646613 | 6.3230741 | -7.1738795 | 9.0453E-06 | 0.00018334 |
| CAMKK1 | -0.912276 | 3.64717916 | -7.1717387 | 9.0729E-06 | 0.00018365 |
| SENP2 | 0.61309919 | 6.06099845 | 7.1703229 | 9.0913E-06 | 0.00018378 |
| NPTN | -0.5907264 | 6.49746964 | -7.1649826 | 9.1607E-06 | 0.00018494 |
| ZNF844 | 0.75414352 | 4.44134648 | 7.16099261 | 9.2129E-06 | 0.00018574 |
| GOLIM4 | 0.69192984 | 6.89811362 | 7.15338505 | 9.3134E-06 | 0.00018752 |
| CALD1 | 0.81205319 | 4.94923942 | 7.14515197 | 9.4234E-06 | 0.0001893 |
| FUNDC2 | 0.93344339 | 4.11578629 | 7.14491145 | 9.4266E-06 | 0.0001893 |
| ARID4A | 0.60928054 | 5.87265219 | 7.13811395 | 9.5185E-06 | 0.00019082 |
| BICDL1 | -0.6243214 | 5.98872187 | -7.1374455 | 9.5276E-06 | 0.00019082 |
| RMND5A | -0.5525413 | 7.21155671 | -7.1294115 | 9.6376E-06 | 0.00019276 |
| RBP7 | -1.7891282 | 0.78683439 | -7.1283048 | 9.6529E-06 | 0.00019282 |
| SIAH1 | -0.9302907 | 3.30083854 | -7.1273645 | 9.6658E-06 | 0.00019282 |
| DPYSL5 | -0.6418558 | 5.95640017 | -7.1236811 | 9.7169E-06 | 0.00019346 |
| LIX1 | 0.59849816 | 6.02318529 | 7.12321698 | 9.7233E-06 | 0.00019346 |

| | | | | | |
|---|---|---|---|---|---|
| ALDH3A2 | 0.51741772 | 7.84478742 | 7.12217936 | 9.7378E-06 | 0.00019349 |
| MEIS2 | 0.52844548 | 7.86908741 | 7.11812249 | 9.7944E-06 | 0.0001942 |
| TBL1X | -0.7834952 | 4.32458 | -7.1157632 | 9.8276E-06 | 0.0001942 |
| RNF41 | -0.5915624 | 6.62510098 | -7.1153956 | 9.8327E-06 | 0.0001942 |
| IMPA1 | -0.640633 | 5.41836845 | -7.1150922 | 9.837E-06 | 0.0001942 |
| GBE1 | -0.7162434 | 7.31467102 | -7.1150629 | 9.8374E-06 | 0.0001942 |
| NFIB | -0.7663193 | 4.2243764 | -7.1103777 | 9.9036E-06 | 0.00019525 |
| C2CD2 | -0.7575806 | 4.91312972 | -7.1092721 | 9.9193E-06 | 0.0001953 |
| ALDOC | -2.9269975 | 2.6383058 | -7.1075784 | 9.9434E-06 | 0.00019552 |
| DOPEY1 | 0.76322224 | 4.45438499 | 7.10452873 | 9.9869E-06 | 0.00019612 |
| DNAH3 | 2.08818455 | 0.01681374 | 7.09510138 | 1.0123E-05 | 0.0001983 |
| H2AFY | -0.517829 | 7.70972327 | -7.0950304 | 1.0124E-05 | 0.0001983 |
| ESYT1 | -0.5160726 | 7.84158183 | -7.0937442 | 1.0142E-05 | 0.00019841 |
| IFIT1 | -1.5645334 | 1.76662831 | -7.0915484 | 1.0174E-05 | 0.00019878 |
| TMED7 | -0.7856983 | 5.19242028 | -7.0902468 | 1.0193E-05 | 0.00019889 |
| AL662844.4 | 0.85422453 | 3.68144722 | 7.08054791 | 1.0336E-05 | 0.00020142 |
| STAT6 | -0.5745469 | 6.6339486 | -7.0708835 | 1.0481E-05 | 0.00020397 |
| RASSF2 | -0.7153196 | 4.74329408 | -7.0682396 | 1.0521E-05 | 0.00020447 |
| AC008771.1 | 0.891698 | 3.55906037 | 7.06682981 | 1.0542E-05 | 0.00020447 |
| DRC3 | 1.16815672 | 2.41088794 | 7.06648356 | 1.0547E-05 | 0.00020447 |
| WNT2B | -1.3018009 | 2.43926485 | -7.0568129 | 1.0695E-05 | 0.00020707 |
| LZTR1 | -0.6588629 | 6.8627033 | -7.0530212 | 1.0753E-05 | 0.00020794 |
| KLF10 | 0.81101867 | 7.27678413 | 7.04960853 | 1.0806E-05 | 0.00020858 |
| LNPK | -0.6788783 | 5.99253823 | -7.0491 | 1.0814E-05 | 0.00020858 |
| CEP95 | 0.58932384 | 6.4578532 | 7.04767322 | 1.0836E-05 | 0.00020875 |
| NUMA1 | 0.55620513 | 7.68439759 | 7.04430874 | 1.0889E-05 | 0.00020949 |
| TGFB3 | -0.7243937 | 4.58402095 | -7.0418027 | 1.0928E-05 | 0.00020998 |
| RSU1 | -0.5570382 | 6.84083999 | -7.0404438 | 1.095E-05 | 0.00021013 |
| AL133367.1 | 1.08265139 | 2.75176037 | 7.03956341 | 1.0964E-05 | 0.00021013 |
| CREG1 | -0.7214078 | 4.75095618 | -7.0373332 | 1.0999E-05 | 0.00021054 |
| FAM166A | 1.69826579 | 1.30133892 | 7.03568794 | 1.1025E-05 | 0.00021077 |
| CEP170B | -0.5679994 | 6.63825673 | -7.032877 | 1.107E-05 | 0.00021136 |
| WDR48 | 0.57828533 | 6.29071616 | 7.03052365 | 1.1108E-05 | 0.00021181 |
| THAP9 | 1.06097905 | 2.73724855 | 7.02948279 | 1.1124E-05 | 0.00021186 |
| ASS1 | -0.933555 | 3.48166096 | -7.0284349 | 1.1141E-05 | 0.00021192 |
| HABP4 | 0.61070838 | 5.84956313 | 7.02395624 | 1.1213E-05 | 0.00021302 |
| AL645933.2 | 1.04280474 | 2.741923 | 7.01895156 | 1.1294E-05 | 0.0002143 |
| TBXA2R | -1.0622243 | 2.8914644 | -7.015919 | 1.1344E-05 | 0.00021497 |
| CNOT4 | 0.70205571 | 4.74273515 | 7.00785897 | 1.1477E-05 | 0.00021721 |
| PCYT1A | -0.5940179 | 6.25989859 | -7.0061884 | 1.1505E-05 | 0.00021747 |
| AADAT | -0.7525785 | 4.2195273 | -7.0052472 | 1.152E-05 | 0.00021749 |
| PNP | -0.563864 | 6.62882302 | -7.0016477 | 1.158E-05 | 0.00021795 |
| SLC9A2 | -0.7685008 | 4.11620139 | -7.0016045 | 1.1581E-05 | 0.00021795 |
| NKAP | 0.72364364 | 4.68373891 | 7.00119407 | 1.1588E-05 | 0.00021795 |
| TRMT11 | 0.67969996 | 5.61997184 | 6.99655416 | 1.1666E-05 | 0.00021915 |

| | | | | | |
|---|---|---|---|---|---|
| CBS | -0.8161037 | 4.38814019 | -6.9905839 | 1.1767E-05 | 0.00022073 |
| ZKSCAN1 | -0.6175617 | 6.31187596 | -6.9891134 | 1.1792E-05 | 0.00022073 |
| NID2 | -1.0903527 | 2.94768077 | -6.9877627 | 1.1815E-05 | 0.00022073 |
| KCNT2 | 1.11990938 | 2.5021037 | 6.98757188 | 1.1819E-05 | 0.00022073 |
| BEGAIN | 0.72386476 | 4.80861905 | 6.98732423 | 1.1823E-05 | 0.00022073 |
| HLA-E | -0.5755468 | 6.38470712 | -6.9850356 | 1.1862E-05 | 0.00022119 |
| NBEAL2 | -0.6957715 | 5.93183893 | -6.983747 | 1.1884E-05 | 0.00022133 |
| OPLAH | -0.8958682 | 4.32219971 | -6.981985 | 1.1915E-05 | 0.00022162 |
| ZNF76 | 0.63261672 | 5.72174653 | 6.97866749 | 1.1972E-05 | 0.00022241 |
| SERF2 | -0.5392806 | 7.73326606 | -6.9730295 | 1.207E-05 | 0.00022396 |
| ZBTB41 | -0.5758129 | 6.13714652 | -6.9697238 | 1.2128E-05 | 0.00022476 |
| SLC24A4 | -1.0146775 | 2.94743448 | -6.9686407 | 1.2147E-05 | 0.00022484 |
| CLN8 | -0.8194553 | 3.79170645 | -6.9634293 | 1.2239E-05 | 0.00022618 |
| C1orf43 | 0.5168858 | 8.11631054 | 6.96285638 | 1.2249E-05 | 0.00022618 |
| SLC8B1 | -0.7734823 | 4.45206238 | -6.9611899 | 1.2279E-05 | 0.00022645 |
| C22orf39 | 0.67090406 | 5.13518221 | 6.95920915 | 1.2314E-05 | 0.00022683 |
| HINT1 | -0.5955779 | 6.44942616 | -6.9572345 | 1.235E-05 | 0.0002272 |
| WDR5B | 0.93721358 | 3.11348612 | 6.94654682 | 1.2543E-05 | 0.00023048 |
| LMNB1 | -0.5096061 | 7.6167312 | -6.9442877 | 1.2584E-05 | 0.00023072 |
| ZNF345 | 1.11276682 | 2.41546214 | 6.94415329 | 1.2587E-05 | 0.00023072 |
| DNAJA2 | -0.5007938 | 7.74453652 | -6.9371509 | 1.2715E-05 | 0.0002328 |
| MEGF6 | -0.9319465 | 4.01303458 | -6.9326704 | 1.2798E-05 | 0.00023404 |
| AC007114.1 | 1.35877171 | 1.80583615 | 6.93164654 | 1.2818E-05 | 0.0002341 |
| FOXP1 | 0.64547663 | 5.21947265 | 6.92642566 | 1.2915E-05 | 0.0002356 |
| SLC30A8 | -0.5875096 | 6.05268369 | -6.925287 | 1.2937E-05 | 0.00023571 |
| KHNYN | -0.547578 | 7.1476232 | -6.9236763 | 1.2967E-05 | 0.00023572 |
| AGO2 | -0.5461074 | 7.56078002 | -6.9235991 | 1.2969E-05 | 0.00023572 |
| EIF4EBP2 | 0.5333001 | 7.23108384 | 6.92279234 | 1.2984E-05 | 0.00023572 |
| TMEM33 | -0.5230498 | 7.0781348 | -6.9219203 | 1.3E-05 | 0.00023573 |
| ZDHHC8 | -0.7467898 | 5.70581012 | -6.9129315 | 1.3172E-05 | 0.00023855 |
| HLA-A | -0.558294 | 7.03026123 | -6.9099318 | 1.3229E-05 | 0.00023881 |
| CLDND1 | -0.6988598 | 4.54996003 | -6.9095706 | 1.3236E-05 | 0.00023881 |
| CD82 | -1.6154874 | 1.08798303 | -6.9089254 | 1.3249E-05 | 0.00023881 |
| AMOTL2 | -0.6777888 | 6.3140876 | -6.9089116 | 1.3249E-05 | 0.00023881 |
| CIRBP | 0.53592791 | 7.10961404 | 6.9072061 | 1.3282E-05 | 0.00023912 |
| SORD | -0.5898154 | 5.95787471 | -6.9045716 | 1.3333E-05 | 0.00023976 |
| MCRIP1 | 0.56298021 | 6.57321519 | 6.90096553 | 1.3403E-05 | 0.00024057 |
| MT1X | 0.6337105 | 5.55207696 | 6.9006172 | 1.341E-05 | 0.00024057 |
| DAP | -0.7620264 | 4.1829187 | -6.8984163 | 1.3453E-05 | 0.00024106 |
| PEG3 | 0.76971196 | 6.53663409 | 6.89630203 | 1.3495E-05 | 0.00024141 |
| NOG | 1.16104992 | 2.95511893 | 6.89581459 | 1.3504E-05 | 0.00024141 |
| APPL2 | 0.65358607 | 5.14996111 | 6.89126616 | 1.3594E-05 | 0.00024247 |
| DPP4 | -0.9153515 | 3.67219607 | -6.8911731 | 1.3596E-05 | 0.00024247 |
| MUT | 0.69048241 | 4.75088426 | 6.88962137 | 1.3627E-05 | 0.00024274 |
| UBE2G2 | -0.6331468 | 6.59523577 | -6.8833492 | 1.3753E-05 | 0.00024414 |

| | | | | | |
|---|---|---|---|---|---|
| E2F1 | 0.53457638 | 6.97343779 | 6.88326091 | 1.3754E-05 | 0.00024414 |
| RGS10 | -0.7145689 | 4.54701204 | -6.8774675 | 1.3871E-05 | 0.00024593 |
| LRRC3 | -1.0009958 | 3.28758215 | -6.8721932 | 1.3979E-05 | 0.00024744 |
| SMARCC1 | 0.58374878 | 7.65966069 | 6.87168257 | 1.3989E-05 | 0.00024744 |
| TPMT | -0.6106003 | 5.46971062 | -6.862839 | 1.4171E-05 | 0.00025037 |
| SLC37A1 | -0.9413015 | 3.18948129 | -6.8600757 | 1.4229E-05 | 0.00025109 |
| F8 | -1.6386073 | 1.25212231 | -6.8545707 | 1.4344E-05 | 0.00025283 |
| SH2D3C | -1.1808605 | 2.40227163 | -6.8489355 | 1.4463E-05 | 0.00025435 |
| HNRNPLL | 0.5330273 | 7.04411021 | 6.84888642 | 1.4464E-05 | 0.00025435 |
| MFSD11 | 0.60463705 | 6.01126524 | 6.84775595 | 1.4488E-05 | 0.00025448 |
| IFIT5 | -0.9103616 | 3.4065238 | -6.8463459 | 1.4518E-05 | 0.00025471 |
| NAXE | -0.5324146 | 7.22311528 | -6.8452869 | 1.454E-05 | 0.00025481 |
| DDAH2 | -0.5752358 | 6.1514096 | -6.8392513 | 1.467E-05 | 0.00025678 |
| LRRC58 | 0.5288745 | 6.79424208 | 6.83706031 | 1.4717E-05 | 0.00025731 |
| FZD6 | -0.6581587 | 5.16709173 | -6.8353044 | 1.4755E-05 | 0.00025767 |
| KLHL24 | 0.71835842 | 5.37815205 | 6.83364524 | 1.4791E-05 | 0.00025774 |
| CPEB3 | -0.8970774 | 3.38105269 | -6.8335415 | 1.4793E-05 | 0.00025774 |
| HYAL2 | -0.6095425 | 5.85554872 | -6.826953 | 1.4937E-05 | 0.00025995 |
| B3GALT1 | 1.16004554 | 2.24198951 | 6.81806301 | 1.5133E-05 | 0.00026306 |
| SSPN | 1.08478168 | 2.44289058 | 6.81599642 | 1.5179E-05 | 0.00026352 |
| ZNF711 | 0.59840905 | 5.47432318 | 6.81531332 | 1.5194E-05 | 0.00026352 |
| EMB | -0.6503998 | 5.05161873 | -6.8073716 | 1.5373E-05 | 0.00026577 |
| CD59 | -0.6238079 | 5.26792461 | -6.807218 | 1.5376E-05 | 0.00026577 |
| C1orf198 | 0.64572783 | 5.25833596 | 6.80465006 | 1.5434E-05 | 0.00026635 |
| INAFM2 | 0.81331326 | 4.95053375 | 6.80417008 | 1.5445E-05 | 0.00026635 |
| CFAP54 | 1.13652737 | 2.42137649 | 6.80287831 | 1.5475E-05 | 0.00026656 |
| PELI3 | -0.8201743 | 4.10867392 | -6.8003535 | 1.5532E-05 | 0.00026698 |
| STAG1 | 0.60637243 | 6.43109365 | 6.80024906 | 1.5535E-05 | 0.00026698 |
| FBXO32 | 1.22020725 | 2.16538231 | 6.79808797 | 1.5584E-05 | 0.00026753 |
| KDELR3 | -0.5731807 | 6.19805279 | -6.7962177 | 1.5627E-05 | 0.00026794 |
| TCF15 | -1.2407027 | 2.1722704 | -6.7955072 | 1.5644E-05 | 0.00026794 |
| S100A10 | -1.3692222 | 1.59498311 | -6.792039 | 1.5724E-05 | 0.00026901 |
| PSPC1 | 0.59507481 | 6.6421647 | 6.78959574 | 1.578E-05 | 0.00026967 |
| MKX | -0.8256732 | 3.59346294 | -6.7887918 | 1.5799E-05 | 0.00026969 |
| MALRD1 | 1.14791709 | 2.39569468 | 6.78537745 | 1.5879E-05 | 0.00027074 |
| RTKN | -0.6000578 | 5.72827457 | -6.7825234 | 1.5946E-05 | 0.00027117 |
| FGFR1OP | 0.56568994 | 6.35331594 | 6.78244028 | 1.5948E-05 | 0.00027117 |
| PSD3 | 0.57512179 | 5.98369695 | 6.78201385 | 1.5958E-05 | 0.00027117 |
| NEMF | 0.53086071 | 6.90988691 | 6.77591007 | 1.6102E-05 | 0.0002731 |
| MRE11 | 0.54569477 | 6.41043205 | 6.77494 | 1.6125E-05 | 0.0002731 |
| SEPT3 | -0.7777303 | 4.07621035 | -6.7749376 | 1.6125E-05 | 0.0002731 |
| MAVS | -0.5285426 | 6.99288559 | -6.7707915 | 1.6224E-05 | 0.00027447 |
| MBD2 | 0.62458326 | 5.70315697 | 6.76625124 | 1.6333E-05 | 0.000276 |
| RAD51AP1 | 0.61172651 | 5.60034365 | 6.76503125 | 1.6363E-05 | 0.00027619 |
| AL050341.2 | 0.89449152 | 3.08000785 | 6.7609884 | 1.6461E-05 | 0.00027754 |

| | | | | | |
|---|---|---|---|---|---|
| WASL | 0.5533286 | 6.52653848 | 6.75804203 | 1.6532E-05 | 0.00027844 |
| FNIP1 | 0.52823599 | 6.86257884 | 6.75656852 | 1.6568E-05 | 0.00027874 |
| KCTD9 | -0.7682067 | 3.79841183 | -6.7510846 | 1.6703E-05 | 0.00028069 |
| PREB | -0.5418911 | 7.09083153 | -6.7499292 | 1.6732E-05 | 0.0002808 |
| FAM129B | -0.5790683 | 5.93700271 | -6.7485992 | 1.6765E-05 | 0.0002808 |
| AL031777.3 | 1.19917741 | 2.37650795 | 6.74791572 | 1.6782E-05 | 0.0002808 |
| OSTF1 | -0.7445772 | 4.19051074 | -6.7469266 | 1.6806E-05 | 0.0002808 |
| GTF2E2 | 0.59809934 | 5.79908882 | 6.74657438 | 1.6815E-05 | 0.0002808 |
| TOR1B | -0.6224641 | 5.23722209 | -6.7463343 | 1.6821E-05 | 0.0002808 |
| XYLT2 | -0.5805235 | 6.10516718 | -6.7444584 | 1.6868E-05 | 0.00028097 |
| NLGN2 | -0.9198226 | 3.36346075 | -6.7440657 | 1.6877E-05 | 0.00028097 |
| KCNJ11 | -0.8183159 | 3.86656403 | -6.7437009 | 1.6887E-05 | 0.00028097 |
| ARRB1 | -0.6102627 | 5.45344393 | -6.7416732 | 1.6937E-05 | 0.00028126 |
| TMTC4 | -0.701078 | 4.54760008 | -6.7415209 | 1.6941E-05 | 0.00028126 |
| C14orf132 | 0.64767437 | 6.08135044 | 6.74047726 | 1.6967E-05 | 0.00028138 |
| PLAG1 | 0.69685121 | 4.86478862 | 6.73935403 | 1.6996E-05 | 0.00028154 |
| CFLAR | 0.61136914 | 5.65260855 | 6.73672716 | 1.7062E-05 | 0.00028233 |
| TLNRD1 | -0.539052 | 6.7389926 | -6.7325485 | 1.7168E-05 | 0.00028377 |
| PSRC1 | -0.5729835 | 6.11748385 | -6.7310099 | 1.7207E-05 | 0.00028411 |
| KRIT1 | -0.5838362 | 6.29860354 | -6.7296135 | 1.7242E-05 | 0.00028439 |
| ACTL6B | 1.16598749 | 2.20803186 | 6.72796355 | 1.7285E-05 | 0.00028472 |
| PTPRJ | -0.6771549 | 4.7795633 | -6.7264118 | 1.7324E-05 | 0.00028472 |
| CCDC184 | -1.2609452 | 2.05164106 | -6.7261998 | 1.733E-05 | 0.00028472 |
| MYL5 | -0.7977661 | 3.88288092 | -6.7258889 | 1.7338E-05 | 0.00028472 |
| CLK4 | 0.63692023 | 4.95376493 | 6.7195602 | 1.7501E-05 | 0.00028704 |
| MAPKAPK5 | 0.56695715 | 6.51076413 | 6.71863167 | 1.7525E-05 | 0.00028704 |
| PNPLA6 | -0.5489975 | 6.86359619 | -6.7182322 | 1.7536E-05 | 0.00028704 |
| LINC01694 | -1.3346058 | 1.75615081 | -6.7174483 | 1.7556E-05 | 0.00028706 |
| CRK | -0.5214646 | 7.17083111 | -6.7127491 | 1.7679E-05 | 0.00028876 |
| SPC24 | -0.5931475 | 6.23862715 | -6.7096819 | 1.776E-05 | 0.00028947 |
| PCGF5 | -0.9649566 | 3.00699761 | -6.7096272 | 1.7761E-05 | 0.00028947 |
| TNFRSF11A | -1.8441241 | 0.36145527 | -6.7064673 | 1.7844E-05 | 0.00029052 |
| KLHL29 | -0.7137997 | 4.51206895 | -6.7053146 | 1.7875E-05 | 0.0002907 |
| RASSF1 | -0.6681045 | 4.80933992 | -6.7014798 | 1.7977E-05 | 0.00029205 |
| EXTL1 | 1.51045287 | 1.31014247 | 6.69651349 | 1.811E-05 | 0.0002939 |
| EP400 | -0.5786209 | 5.73235336 | -6.694144 | 1.8174E-05 | 0.00029462 |
| P4HA2 | -0.9353403 | 6.22326953 | -6.6933978 | 1.8194E-05 | 0.00029463 |
| DPYD | 0.87863045 | 3.52179739 | 6.69246903 | 1.8219E-05 | 0.00029472 |
| EPHA10 | -0.773991 | 4.19662179 | -6.6890848 | 1.8311E-05 | 0.00029589 |
| MIDN | 0.65320899 | 5.62556157 | 6.68175452 | 1.8512E-05 | 0.00029882 |
| PPIP5K2 | -0.5424056 | 6.39159062 | -6.6792602 | 1.8581E-05 | 0.00029928 |
| CISD3 | -0.7878361 | 4.48423213 | -6.6786852 | 1.8597E-05 | 0.00029928 |
| FSTL3 | -0.6459254 | 5.30284909 | -6.6784856 | 1.8602E-05 | 0.00029928 |
| SLC4A11 | -0.6959393 | 4.53090448 | -6.6778481 | 1.862E-05 | 0.00029928 |
| HOXC9 | 0.64838154 | 5.52443035 | 6.67608748 | 1.8669E-05 | 0.00029946 |

| | | | | | |
|---|---|---|---|---|---|
| ITGB4 | -0.9195056 | 3.36478625 | -6.676019 | 1.8671E-05 | 0.00029946 |
| DHFR | 0.65026932 | 5.64378425 | 6.67499501 | 1.8699E-05 | 0.0002996 |
| APPBP2 | -0.5412529 | 6.65758656 | -6.6693493 | 1.8857E-05 | 0.00030138 |
| RTN4R | -0.6823833 | 4.69926589 | -6.6688791 | 1.887E-05 | 0.00030138 |
| CBR1 | -0.55806 | 6.20534757 | -6.6634586 | 1.9023E-05 | 0.00030328 |
| RNF20 | -0.5490993 | 6.58061536 | -6.6632475 | 1.9029E-05 | 0.00030328 |
| HSBP1L1 | -0.8833405 | 3.41295441 | -6.6607067 | 1.9101E-05 | 0.00030395 |
| BSPRY | -2.0647049 | -0.004571 | -6.6599721 | 1.9122E-05 | 0.00030395 |
| SUN1 | -0.6071014 | 7.38869962 | -6.6596617 | 1.9131E-05 | 0.00030395 |
| HEBP1 | -0.9839419 | 2.8291684 | -6.6582523 | 1.9171E-05 | 0.00030427 |
| CDH15 | -1.8035286 | 0.40119325 | -6.6560417 | 1.9235E-05 | 0.00030495 |
| ADORA1 | -1.2016878 | 2.19668594 | -6.6531824 | 1.9317E-05 | 0.00030594 |
| MFSD9 | -0.7152071 | 4.54472941 | -6.6493339 | 1.9428E-05 | 0.00030738 |
| PELO | -0.5598619 | 6.08002329 | -6.6485986 | 1.9449E-05 | 0.00030739 |
| VSIR | -1.6217913 | 0.9668935 | -6.6448762 | 1.9558E-05 | 0.0003086 |
| INPP5J | -2.0779861 | 0.1385205 | -6.6445654 | 1.9567E-05 | 0.0003086 |
| LRP11 | -0.6346918 | 6.09273741 | -6.6386265 | 1.9741E-05 | 0.00031072 |
| NLE1 | -0.6413555 | 5.9755076 | -6.6385964 | 1.9742E-05 | 0.00031072 |
| SCML1 | -0.756405 | 5.2693769 | -6.6357243 | 1.9827E-05 | 0.00031173 |
| FXYD5 | -0.7022412 | 4.51826568 | -6.629637 | 2.0008E-05 | 0.00031425 |
| ZNF134 | 0.60637624 | 5.22130809 | 6.62863815 | 2.0038E-05 | 0.00031432 |
| TFDP2 | 0.55043394 | 6.38690267 | 6.62220386 | 2.0231E-05 | 0.00031678 |
| ZNF280B | -0.8318057 | 3.5192224 | -6.6163309 | 2.041E-05 | 0.00031905 |
| MARF1 | 0.66884523 | 4.91234243 | 6.61588958 | 2.0423E-05 | 0.00031905 |
| MORC3 | 0.54805497 | 6.08364227 | 6.60829429 | 2.0657E-05 | 0.00032211 |
| SLC29A1 | -0.5328577 | 6.6989639 | -6.6066986 | 2.0706E-05 | 0.00032255 |
| ABHD11 | -0.630596 | 5.09982316 | -6.6018327 | 2.0858E-05 | 0.00032457 |
| IFT46 | 0.65958499 | 4.99598377 | 6.59943285 | 2.0933E-05 | 0.00032541 |
| PBLD | 0.72403705 | 4.36244526 | 6.59699811 | 2.1009E-05 | 0.00032626 |
| KCND2 | 0.61338385 | 6.13286693 | 6.59580377 | 2.1047E-05 | 0.00032651 |
| FAM229B | 0.88958829 | 3.71558945 | 6.59500687 | 2.1072E-05 | 0.00032657 |
| FAM43B | -0.9439347 | 3.30957169 | -6.5929615 | 2.1137E-05 | 0.00032723 |
| CBFA2T3 | -1.8800001 | 2.0908499 | -6.5900147 | 2.1231E-05 | 0.00032806 |
| CCDC85C | 0.59214081 | 6.7607665 | 6.58896908 | 2.1264E-05 | 0.00032806 |
| AC108002.1 | -1.2654957 | 1.96075645 | -6.5886127 | 2.1275E-05 | 0.00032806 |
| CERS5 | -0.6098593 | 5.27758567 | -6.588559 | 2.1277E-05 | 0.00032806 |
| FAM234B | 0.76545027 | 4.44927409 | 6.58747411 | 2.1312E-05 | 0.00032826 |
| ARFGEF3 | -0.7640199 | 4.92090661 | -6.5820909 | 2.1485E-05 | 0.00033059 |
| NRXN2 | -1.7067777 | 0.89951883 | -6.5797124 | 2.1561E-05 | 0.00033143 |
| IL21R | -1.9626775 | 0.1707642 | -6.5781782 | 2.1611E-05 | 0.00033186 |
| AL512625.1 | 1.44470835 | 1.19574967 | 6.57664404 | 2.1661E-05 | 0.00033229 |
| STK3 | -0.5458668 | 6.23230802 | -6.5755463 | 2.1697E-05 | 0.0003325 |
| SKP2 | 0.55392161 | 6.68285974 | 6.57114615 | 2.1841E-05 | 0.00033437 |
| NLGN4X | -0.7422885 | 4.29078523 | -6.5700604 | 2.1876E-05 | 0.00033457 |
| HOXC10 | -0.6355036 | 5.5545567 | -6.5661637 | 2.2005E-05 | 0.0003362 |

| | | | | | |
|---|---|---|---|---|---|
| FBXO44 | -0.7165836 | 5.32069024 | -6.5647605 | 2.2051E-05 | 0.00033623 |
| FBXO33 | 0.54655122 | 6.09382109 | 6.56261208 | 2.2122E-05 | 0.00033698 |
| NR2F1 | 0.51735324 | 7.164505 | 6.55307746 | 2.2442E-05 | 0.00034082 |
| CORO1A | -0.8788059 | 3.31178389 | -6.5514601 | 2.2497E-05 | 0.00034131 |
| LRRC17 | 1.20056218 | 2.09471747 | 6.54905199 | 2.2578E-05 | 0.00034207 |
| GALNS | -0.5405412 | 6.5144015 | -6.548638 | 2.2593E-05 | 0.00034207 |
| NEDD8 | -0.5395614 | 6.44178182 | -6.5442854 | 2.2741E-05 | 0.00034398 |
| SNPH | -0.6774673 | 4.8048718 | -6.54288 | 2.2789E-05 | 0.00034402 |
| TUBA1A | -0.5472312 | 8.23582697 | -6.5390093 | 2.2923E-05 | 0.00034558 |
| PHF20 | -0.5724132 | 5.77501474 | -6.5380163 | 2.2957E-05 | 0.00034558 |
| SUSD1 | -1.1520093 | 2.12696974 | -6.5378888 | 2.2961E-05 | 0.00034558 |
| DOLPP1 | -0.7273413 | 4.93763023 | -6.53686 | 2.2997E-05 | 0.00034578 |
| TSPO | -0.5702207 | 6.35534291 | -6.5352582 | 2.3052E-05 | 0.00034603 |
| GAS2L1 | -0.5870897 | 6.18275085 | -6.5347877 | 2.3069E-05 | 0.00034603 |
| GPATCH2 | 0.61598095 | 5.34726191 | 6.53438916 | 2.3083E-05 | 0.00034603 |
| SLC30A1 | 0.51760482 | 6.54463567 | 6.53210367 | 2.3162E-05 | 0.00034689 |
| OSBPL8 | 0.51360622 | 7.53124137 | 6.52484106 | 2.3417E-05 | 0.00035036 |
| COL4A2 | -0.7737367 | 4.4183433 | -6.5194783 | 2.3608E-05 | 0.0003524 |
| WIPI1 | -0.7502457 | 4.67861182 | -6.5193074 | 2.3614E-05 | 0.0003524 |
| ATP7B | -0.7128135 | 4.31604066 | -6.5190363 | 2.3623E-05 | 0.0003524 |
| ELAVL3 | -1.5831473 | 0.90748497 | -6.5173928 | 2.3682E-05 | 0.00035292 |
| IFT122 | 0.58894601 | 5.52097156 | 6.51654591 | 2.3712E-05 | 0.00035303 |
| RAB11FIP3 | 0.55284451 | 6.05766379 | 6.51428466 | 2.3794E-05 | 0.00035389 |
| ZNF222 | 0.93512221 | 2.98618541 | 6.5102954 | 2.3937E-05 | 0.00035568 |
| ATG2B | 0.5763948 | 5.98663736 | 6.50898264 | 2.3985E-05 | 0.00035603 |
| NPTX1 | 0.7486519 | 4.05402219 | 6.50286366 | 2.4208E-05 | 0.00035864 |
| TBC1D19 | 1.22224067 | 1.99229787 | 6.50098764 | 2.4276E-05 | 0.00035895 |
| PDCD4 | 0.58883518 | 5.53863238 | 6.49935022 | 2.4337E-05 | 0.00035949 |
| INTS2 | -0.5404013 | 6.38339276 | -6.4983419 | 2.4374E-05 | 0.00035969 |
| HOXB6 | 0.90928679 | 3.38042831 | 6.49547937 | 2.4479E-05 | 0.00036056 |
| ETV1 | 0.55250228 | 6.26023202 | 6.49448049 | 2.4516E-05 | 0.00036074 |
| AL512791.2 | 0.95246579 | 2.76916796 | 6.48853864 | 2.4738E-05 | 0.00036365 |
| BPNT1 | -0.5771617 | 5.60619343 | -6.4857062 | 2.4844E-05 | 0.00036486 |
| LRFN5 | 0.53699455 | 6.64913682 | 6.48492225 | 2.4874E-05 | 0.00036493 |
| CLDN23 | -1.6692456 | 0.89934164 | -6.4822927 | 2.4973E-05 | 0.00036598 |
| CPTP | -0.5731452 | 5.98603129 | -6.4817589 | 2.4993E-05 | 0.00036598 |
| UNC119 | -0.6859391 | 5.29406399 | -6.4798046 | 2.5067E-05 | 0.00036671 |
| RILPL2 | -0.6348163 | 4.89009635 | -6.4773455 | 2.5161E-05 | 0.00036772 |
| BCAS2 | -0.5850307 | 5.80385955 | -6.4750275 | 2.5249E-05 | 0.00036831 |
| MEGF8 | 0.5744867 | 5.81709075 | 6.47393849 | 2.5291E-05 | 0.00036831 |
| DNAJC2 | -0.5260372 | 6.49589722 | -6.4731083 | 2.5323E-05 | 0.00036831 |
| DEPDC1B | 0.58898162 | 5.36087559 | 6.46968804 | 2.5454E-05 | 0.00036987 |
| BMP8B | -0.833416 | 3.67337782 | -6.4671901 | 2.5551E-05 | 0.00037092 |
| TUBA4A | -1.4078048 | 1.45922777 | -6.4664895 | 2.5578E-05 | 0.00037096 |
| ANKS6 | -0.6016428 | 5.20749763 | -6.4618466 | 2.5759E-05 | 0.00037322 |

| | | | | | |
|---|---|---|---|---|---|
| ETV4 | -0.6893195 | 5.62329443 | -6.4600563 | 2.5829E-05 | 0.00037388 |
| DLX2-DT | 0.96345792 | 2.86166491 | 6.45833834 | 2.5896E-05 | 0.00037405 |
| SLC25A21-AS1 | 1.16512395 | 1.78916794 | 6.45786004 | 2.5915E-05 | 0.00037405 |
| TLL2 | -0.8606649 | 3.36046558 | -6.4564181 | 2.5972E-05 | 0.00037427 |
| AKAP9 | 0.65455689 | 6.76503394 | 6.45621775 | 2.598E-05 | 0.00037427 |
| UBN2 | 0.60492594 | 5.09943093 | 6.44607985 | 2.6383E-05 | 0.00037861 |
| AC008124.1 | 0.77029389 | 4.11817006 | 6.44598584 | 2.6387E-05 | 0.00037861 |
| NUAK1 | -0.6987866 | 4.74197305 | -6.4455062 | 2.6406E-05 | 0.00037861 |
| HEY2 | 0.50257777 | 7.05962701 | 6.4437346 | 2.6477E-05 | 0.00037916 |
| MYD88 | -0.8924667 | 3.0871361 | -6.4433097 | 2.6494E-05 | 0.00037916 |
| KAZN | -0.8378127 | 3.45199009 | -6.4406335 | 2.6602E-05 | 0.00038034 |
| DOPEY2 | -0.7151555 | 4.1930333 | -6.4336892 | 2.6885E-05 | 0.00038294 |
| EML2 | -0.7199908 | 5.92550939 | -6.4313015 | 2.6982E-05 | 0.00038361 |
| TMC6 | -0.5670516 | 5.77411749 | -6.4269488 | 2.7162E-05 | 0.00038543 |
| SELENOH | -0.6101899 | 5.23322937 | -6.4228963 | 2.733E-05 | 0.00038745 |
| TGFBR1 | 0.54536208 | 5.85947017 | 6.4218368 | 2.7374E-05 | 0.00038772 |
| PLPP2 | -0.5154443 | 6.67708823 | -6.4199354 | 2.7453E-05 | 0.00038848 |
| MMP24 | 0.9554026 | 2.82689322 | 6.41924045 | 2.7482E-05 | 0.00038853 |
| GOLGA1 | 0.57234791 | 5.51202367 | 6.41823785 | 2.7524E-05 | 0.00038876 |
| AC091271.1 | 1.10874388 | 2.19855505 | 6.41708786 | 2.7573E-05 | 0.00038878 |
| STRBP | -0.5307716 | 6.19037331 | -6.4169811 | 2.7577E-05 | 0.00038878 |
| THEMIS2 | -0.8388931 | 3.49033029 | -6.4129386 | 2.7747E-05 | 0.00039081 |
| TNS2 | -0.9350306 | 3.13460501 | -6.4099858 | 2.7873E-05 | 0.00039221 |
| GCG | -0.777099 | 3.72712965 | -6.4083338 | 2.7943E-05 | 0.00039284 |
| PDPR | 0.54316447 | 5.98886971 | 6.40186844 | 2.822E-05 | 0.00039636 |
| GPR135 | -0.8104056 | 3.45677802 | -6.3977677 | 2.8397E-05 | 0.00039755 |
| NABP1 | -0.5966206 | 4.98512493 | -6.3974665 | 2.841E-05 | 0.00039755 |
| AC060780.1 | 1.06293979 | 2.73197998 | 6.39628204 | 2.8461E-05 | 0.00039791 |
| THY1 | -0.7606056 | 4.02819215 | -6.3955857 | 2.8492E-05 | 0.00039795 |
| PM20D2 | -0.5450071 | 6.01015943 | -6.394995 | 2.8517E-05 | 0.00039795 |
| AC084033.3 | 0.75529323 | 3.90122155 | 6.39297491 | 2.8605E-05 | 0.00039881 |
| FAM32A | -0.5310801 | 6.66604865 | -6.388073 | 2.882E-05 | 0.00040095 |
| IREB2 | -0.5282789 | 6.82559407 | -6.3879974 | 2.8824E-05 | 0.00040095 |
| SMPD1 | 0.59123338 | 7.08144776 | 6.38722699 | 2.8857E-05 | 0.00040095 |
| NEK10 | 1.08080386 | 2.27829686 | 6.38168377 | 2.9103E-05 | 0.00040389 |
| LPCAT2 | -0.9126379 | 2.99533676 | -6.3764018 | 2.9339E-05 | 0.00040679 |
| PAK2 | -0.552079 | 5.79592068 | -6.3734284 | 2.9472E-05 | 0.00040827 |
| TMEM203 | 0.60560422 | 5.10424538 | 6.37262301 | 2.9509E-05 | 0.0004083 |
| ZNF141 | 0.66248866 | 5.28149393 | 6.37218738 | 2.9528E-05 | 0.0004083 |
| ORAI2 | 0.50702845 | 6.85507567 | 6.36912246 | 2.9667E-05 | 0.00040985 |
| AL137077.2 | 1.25268859 | 1.4819292 | 6.3649684 | 2.9856E-05 | 0.00041208 |
| RPL26L1 | -0.5865487 | 5.30370506 | -6.3629106 | 2.9951E-05 | 0.00041301 |
| ZNF10 | 0.73097244 | 4.17786034 | 6.35730324 | 3.0209E-05 | 0.00041581 |
| ANK1 | 1.42292606 | 1.26655624 | 6.35500169 | 3.0315E-05 | 0.00041656 |
| AC024896.1 | 0.72423629 | 3.82333964 | 6.35494799 | 3.0318E-05 | 0.00041656 |

| | | | | | |
|---|---|---|---|---|---|
| SLCO4A1 | -0.5765569 | 5.79165375 | -6.3540133 | 3.0361E-05 | 0.00041677 |
| HAS2 | -0.7773928 | 3.78343202 | -6.3523202 | 3.044E-05 | 0.00041748 |
| MXD4 | 0.81651944 | 5.15235051 | 6.34884611 | 3.0603E-05 | 0.00041895 |
| UBA5 | -0.6291729 | 5.1434223 | -6.3452372 | 3.0772E-05 | 0.00042089 |
| CDKN2AIP | 0.6290001 | 5.16842101 | 6.34455735 | 3.0805E-05 | 0.00042095 |
| AL354920.1 | 1.15793323 | 1.99324968 | 6.34132875 | 3.0957E-05 | 0.00042266 |
| PPFIBP2 | -0.7741175 | 3.80456862 | -6.3396323 | 3.1038E-05 | 0.00042338 |
| XPR1 | -0.5046955 | 6.75646392 | -6.3388473 | 3.1075E-05 | 0.0004235 |
| SLC9A7 | -0.6808279 | 4.33085104 | -6.3349383 | 3.1262E-05 | 0.00042567 |
| SESN2 | 0.53579019 | 6.27700563 | 6.33317268 | 3.1347E-05 | 0.00042644 |
| LINC02393 | -1.7325586 | 0.32441895 | -6.3282636 | 3.1584E-05 | 0.00042889 |
| RANBP6 | 0.56313757 | 6.69470395 | 6.31550694 | 3.2209E-05 | 0.00043699 |
| CAVIN1 | -0.5362311 | 7.30582381 | -6.3127143 | 3.2348E-05 | 0.00043848 |
| DISC1 | 1.46521423 | 1.2372905 | 6.30085354 | 3.2944E-05 | 0.00044575 |
| ARG2 | -1.0873658 | 2.70709627 | -6.2982082 | 3.3078E-05 | 0.00044717 |
| ARPP21 | 1.21633464 | 1.90138846 | 6.28958674 | 3.352E-05 | 0.00045275 |
| CAB39 | -0.5529352 | 6.2485104 | -6.2861618 | 3.3697E-05 | 0.00045433 |
| RIMKLB | 0.56579099 | 5.47256316 | 6.28521583 | 3.3747E-05 | 0.00045442 |
| KIAA0825 | 0.95445705 | 2.52986899 | 6.284873 | 3.3764E-05 | 0.00045442 |
| ZSWIM2 | 0.78127095 | 3.47564496 | 6.28032489 | 3.4002E-05 | 0.00045722 |
| OSMR | -1.0179717 | 2.47500761 | -6.2728309 | 3.4397E-05 | 0.00046212 |
| LIMD1 | -0.5574414 | 6.32079396 | -6.2715502 | 3.4465E-05 | 0.00046245 |
| FAM204A | 0.52324044 | 6.52849928 | 6.27064801 | 3.4513E-05 | 0.00046245 |
| DIS3 | -0.5064832 | 6.57005638 | -6.2678678 | 3.4662E-05 | 0.00046403 |
| CACNB3 | -0.5923204 | 5.17778173 | -6.2642788 | 3.4854E-05 | 0.00046578 |
| ADIPOR1 | 0.5166921 | 6.93002983 | 6.26020025 | 3.5075E-05 | 0.00046831 |
| ST6GALNAC4 | -0.6155817 | 4.97107731 | -6.2591519 | 3.5131E-05 | 0.00046866 |
| AC124854.1 | 1.37679113 | 1.15065256 | 6.2559427 | 3.5306E-05 | 0.00047057 |
| EVI5L | 0.74452637 | 4.11162569 | 6.25440675 | 3.539E-05 | 0.0004708 |
| ZNF25 | -0.6052652 | 5.52750745 | -6.2541558 | 3.5404E-05 | 0.0004708 |
| ANO8 | -0.852052 | 4.43893216 | -6.2536722 | 3.543E-05 | 0.0004708 |
| CD40 | -0.9912393 | 2.75118051 | -6.2533575 | 3.5447E-05 | 0.0004708 |
| RAB8A | -0.5994217 | 5.97669953 | -6.2483918 | 3.5721E-05 | 0.00047292 |
| SLC39A4 | -0.6232273 | 4.90442081 | -6.24818 | 3.5732E-05 | 0.00047292 |
| ZNF397 | 0.62696676 | 4.98639779 | 6.24817662 | 3.5732E-05 | 0.00047292 |
| ACAD10 | 0.56178575 | 6.41858641 | 6.24659686 | 3.582E-05 | 0.00047325 |
| LGALSL | -0.6377597 | 5.10289084 | -6.2455622 | 3.5877E-05 | 0.0004736 |
| ENC1 | -0.5232732 | 6.27194259 | -6.2422804 | 3.606E-05 | 0.00047531 |
| KDM3A | 0.71191586 | 6.53036976 | 6.24209998 | 3.607E-05 | 0.00047531 |
| PDE4A | 0.67212112 | 4.99567752 | 6.24099479 | 3.6132E-05 | 0.00047571 |
| MICU3 | 0.67503763 | 4.05123376 | 6.23909102 | 3.6238E-05 | 0.0004763 |
| HLCS | -0.5821144 | 5.17966638 | -6.2389368 | 3.6247E-05 | 0.0004763 |
| ATF1 | -0.5938283 | 5.24714381 | -6.238513 | 3.6271E-05 | 0.0004763 |
| SAR1B | 0.50568216 | 6.48095293 | 6.2367658 | 3.6369E-05 | 0.000477 |
| CCDC24 | -0.8089394 | 4.01224142 | -6.2364467 | 3.6387E-05 | 0.000477 |

| | | | | | |
|---|---|---|---|---|---|
| IFNGR2 | -0.5752479 | 5.31941488 | -6.2357038 | 3.6429E-05 | 0.00047707 |
| TMEM178B | 0.52615579 | 6.10565194 | 6.23523099 | 3.6455E-05 | 0.00047707 |
| AKNA | -1.0458052 | 2.51720905 | -6.2331906 | 3.6571E-05 | 0.00047817 |
| SSH3 | -0.8331681 | 3.2770742 | -6.2298233 | 3.6762E-05 | 0.00048025 |
| AC096667.1 | 1.52182658 | 0.59118396 | 6.22750384 | 3.6894E-05 | 0.00048115 |
| MBOAT7 | -0.5699211 | 5.45303486 | -6.2261945 | 3.6969E-05 | 0.00048171 |
| RNPEP | -0.5330615 | 6.03114972 | -6.2234109 | 3.7129E-05 | 0.00048245 |
| ANKRD26 | 0.55226557 | 5.55210796 | 6.22310902 | 3.7146E-05 | 0.00048245 |
| CABLES2 | -0.6916443 | 4.26580314 | -6.2229965 | 3.7153E-05 | 0.00048245 |
| DOLK | 0.77088264 | 4.96127292 | 6.21951615 | 3.7354E-05 | 0.00048422 |
| LEF1 | 0.63528249 | 7.05954343 | 6.21852073 | 3.7412E-05 | 0.00048456 |
| PLOD1 | -0.5816114 | 6.76065165 | -6.216917 | 3.7505E-05 | 0.00048535 |
| OLFML2A | -0.9016489 | 3.0778483 | -6.2153089 | 3.7598E-05 | 0.00048614 |
| PEMT | -0.5498645 | 5.69183788 | -6.2131646 | 3.7724E-05 | 0.00048735 |
| PLPPR3 | 0.57225238 | 5.90973496 | 6.20890021 | 3.7974E-05 | 0.00049017 |
| GAPVD1 | -0.5137849 | 7.06754645 | -6.2067351 | 3.8102E-05 | 0.00049131 |
| TEAD1 | -0.5556759 | 6.74795347 | -6.2062995 | 3.8128E-05 | 0.00049131 |
| AF117829.1 | 0.93825647 | 3.25448701 | 6.20441171 | 3.8239E-05 | 0.00049191 |
| PIGM | 0.62642044 | 4.60543489 | 6.19580288 | 3.8754E-05 | 0.00049727 |
| PMS2 | -0.5265536 | 6.21412774 | -6.1938171 | 3.8874E-05 | 0.00049838 |
| C15orf65 | 1.26484448 | 1.74716794 | 6.19255347 | 3.895E-05 | 0.00049894 |
| TLE6 | -0.8368967 | 3.77278513 | -6.1916632 | 3.9004E-05 | 0.00049901 |
| NEURL1B | -0.5651197 | 6.13164243 | -6.1883102 | 3.9208E-05 | 0.00050055 |
| AL691432.2 | 0.78481768 | 3.46501581 | 6.18756251 | 3.9254E-05 | 0.00050071 |
| CRYBG2 | -1.1019296 | 2.35370507 | -6.1857254 | 3.9366E-05 | 0.00050172 |
| UGCG | -0.5601836 | 5.29091523 | -6.1838232 | 3.9482E-05 | 0.00050246 |
| CD24 | -1.1708374 | 1.8058417 | -6.1836864 | 3.9491E-05 | 0.00050246 |
| RAB20 | -2.0378812 | 0.78506623 | -6.1793235 | 3.976E-05 | 0.00050546 |
| GTPBP3 | -0.5421711 | 5.7718078 | -6.177377 | 3.988E-05 | 0.00050628 |
| ODF2L | -0.5681084 | 5.39977736 | -6.1768531 | 3.9913E-05 | 0.00050628 |
| INO80E | -0.5112518 | 6.1658871 | -6.1766703 | 3.9924E-05 | 0.00050628 |
| SNAPC1 | -0.5682319 | 5.1900279 | -6.1754297 | 4.0001E-05 | 0.00050683 |
| DCBLD2 | 0.5789576 | 5.74728513 | 6.17089503 | 4.0285E-05 | 0.00050999 |
| FIBCD1 | -0.9750052 | 2.7763429 | -6.1700003 | 4.0341E-05 | 0.00051028 |
| KCNH6 | -1.675503 | 0.65454974 | -6.1682139 | 4.0453E-05 | 0.00051085 |
| SCG3 | 0.94342172 | 2.58353561 | 6.16696451 | 4.0532E-05 | 0.00051133 |
| GLIDR | 1.13284712 | 2.21778529 | 6.16653524 | 4.0559E-05 | 0.00051133 |
| POLR3F | -0.5848775 | 5.39611246 | -6.162784 | 4.0797E-05 | 0.0005139 |
| LRRK1 | -0.6314938 | 4.62475648 | -6.1593207 | 4.1018E-05 | 0.00051625 |
| SAMD15 | 0.91209078 | 3.17852779 | 6.15867264 | 4.1059E-05 | 0.00051634 |
| EGR2 | 1.47338519 | 1.20324484 | 6.15267553 | 4.1445E-05 | 0.00052009 |
| TAF13 | -0.5837334 | 5.04999194 | -6.1524443 | 4.146E-05 | 0.00052009 |
| PAIP2B | 1.03865326 | 2.45126895 | 6.15003217 | 4.1616E-05 | 0.00052161 |
| ZNF140 | 0.5478931 | 5.43497519 | 6.14602857 | 4.1877E-05 | 0.00052445 |
| ZNF367 | 0.52107957 | 5.85202424 | 6.14549357 | 4.1912E-05 | 0.00052445 |

| | | | | | |
|---|---|---|---|---|---|
| DPH6 | 0.64846552 | 4.23726119 | 6.14017193 | 4.2261E-05 | 0.00052839 |
| DLX2 | 0.58710359 | 6.54756972 | 6.13826827 | 4.2387E-05 | 0.00052953 |
| BBS10 | 0.56470414 | 5.22025141 | 6.13703111 | 4.2469E-05 | 0.00053011 |
| VPS50 | 0.525972 | 5.87272933 | 6.13537697 | 4.2579E-05 | 0.00053105 |
| CLGN | 0.68585585 | 4.02403092 | 6.13431055 | 4.265E-05 | 0.0005312 |
| PEX1 | 0.52027602 | 5.93611869 | 6.13413902 | 4.2661E-05 | 0.0005312 |
| CES2 | -0.5152301 | 6.15198891 | -6.1330972 | 4.2731E-05 | 0.00053163 |
| SCRT1 | 0.99048737 | 2.6313607 | 6.12821645 | 4.3058E-05 | 0.00053526 |
| GPR137B | 0.59576855 | 4.91950018 | 6.12641825 | 4.3179E-05 | 0.00053632 |
| AC018647.2 | 0.69522135 | 3.88215784 | 6.12288262 | 4.3419E-05 | 0.00053823 |
| LDAH | 0.5201295 | 5.98422973 | 6.12209223 | 4.3472E-05 | 0.00053823 |
| HRAS | -0.5232946 | 6.1221365 | -6.1220514 | 4.3475E-05 | 0.00053823 |
| DGAT2 | 0.65911846 | 4.83861236 | 6.11884448 | 4.3694E-05 | 0.00054029 |
| HOOK3 | -0.5151603 | 6.27702677 | -6.1185591 | 4.3713E-05 | 0.00054029 |
| OCLN | -0.6242825 | 4.64859385 | -6.1129459 | 4.4099E-05 | 0.00054462 |
| ZER1 | 0.57617829 | 5.22133876 | 6.10854443 | 4.4404E-05 | 0.00054793 |
| ROBO3 | -0.6880614 | 5.09447986 | -6.1057038 | 4.4602E-05 | 0.00054993 |
| SNX10 | -1.311305 | 1.46456289 | -6.0971153 | 4.5206E-05 | 0.00055647 |
| BAIAP2-DT | -0.5413223 | 5.90558876 | -6.0950527 | 4.5352E-05 | 0.00055782 |
| NAGA | -0.5071036 | 6.2959958 | -6.0919762 | 4.5571E-05 | 0.00055961 |
| EPS8L1 | -0.7058759 | 4.01524985 | -6.090504 | 4.5677E-05 | 0.00056045 |
| MAP10 | 1.21795551 | 1.66225784 | 6.08979106 | 4.5728E-05 | 0.00056062 |
| ATP2A3 | -0.8655106 | 3.51768348 | -6.0892237 | 4.5769E-05 | 0.00056066 |
| LRP4 | 0.51269011 | 7.48677307 | 6.08828123 | 4.5836E-05 | 0.00056069 |
| KCNJ12 | -0.5974903 | 5.62550702 | -6.08406 | 4.6141E-05 | 0.00056352 |
| CEP162 | 0.63866778 | 4.360564 | 6.08341566 | 4.6187E-05 | 0.00056352 |
| MRPS9 | 0.5201762 | 6.25189728 | 6.08193937 | 4.6295E-05 | 0.00056437 |
| AC016394.1 | 1.00905628 | 2.45149508 | 6.08066841 | 4.6387E-05 | 0.00056504 |
| ZNF32 | -0.5763777 | 5.22306628 | -6.0793967 | 4.648E-05 | 0.00056571 |
| RASSF3 | -0.5034982 | 6.4166164 | -6.0780463 | 4.6578E-05 | 0.00056646 |
| DDIT3 | 0.64646524 | 6.2383835 | 6.07438107 | 4.6847E-05 | 0.00056927 |
| EMILIN3 | -1.0616092 | 2.28826009 | -6.0708183 | 4.711E-05 | 0.00057161 |
| ZNF333 | 0.84365963 | 3.36765713 | 6.0702872 | 4.7149E-05 | 0.00057161 |
| SLC4A8 | -0.6254571 | 4.60532926 | -6.0702392 | 4.7153E-05 | 0.00057161 |
| PLCG1 | -0.5620472 | 8.11719074 | -6.066191 | 4.7454E-05 | 0.00057434 |
| RAB3GAP1 | -0.5087476 | 6.40869851 | -6.0652165 | 4.7526E-05 | 0.00057476 |
| PGLS | -0.5606583 | 5.30962119 | -6.059937 | 4.7923E-05 | 0.00057909 |
| FGF2 | -0.8511931 | 3.43490165 | -6.0532223 | 4.8431E-05 | 0.00058384 |
| DUSP4 | 0.93264384 | 2.69198444 | 6.05235361 | 4.8498E-05 | 0.00058417 |
| AATF | 0.60377809 | 6.6677694 | 6.05138451 | 4.8572E-05 | 0.0005846 |
| TCEAL1 | -0.8814589 | 2.80548668 | -6.0485679 | 4.8787E-05 | 0.00058673 |
| STIM1 | -0.5316897 | 5.80117989 | -6.047254 | 4.8888E-05 | 0.00058747 |
| AC012360.3 | 0.82861905 | 3.40147841 | 6.04613209 | 4.8975E-05 | 0.00058805 |
| CGREF1 | -0.6651364 | 5.01120638 | -6.0442341 | 4.9121E-05 | 0.00058934 |
| MT-ATP6 | -0.584929 | 5.35439346 | -6.0429795 | 4.9218E-05 | 0.00059004 |

| | | | | | |
|---|---|---|---|---|---|
| MAK16 | -0.5460856 | 6.08512506 | -6.037121 | 4.9675E-05 | 0.00059504 |
| CRNKL1 | 0.51696951 | 6.41552634 | 6.036462 | 4.9726E-05 | 0.00059506 |
| MKRN3 | 0.5340711 | 5.87186355 | 6.03609192 | 4.9755E-05 | 0.00059506 |
| NOTUM | -0.6927079 | 4.11506939 | -6.0301892 | 5.022E-05 | 0.00059967 |
| C1GALT1 | -0.6427932 | 5.17859189 | -6.0288235 | 5.0329E-05 | 0.00060002 |
| SLC9A8 | 0.67855722 | 4.07674155 | 6.02417297 | 5.0699E-05 | 0.00060396 |
| CARF | 0.99463818 | 2.98796894 | 6.02136444 | 5.0924E-05 | 0.00060617 |
| ULK4 | 0.61535342 | 4.62020984 | 6.02028067 | 5.1011E-05 | 0.00060673 |
| HIST1H4H | 0.50438923 | 7.16594603 | 6.01928213 | 5.1092E-05 | 0.00060721 |
| BEND6 | 0.97302115 | 2.34546997 | 6.01851663 | 5.1154E-05 | 0.00060746 |
| UBAP1 | 0.52864312 | 5.78774764 | 6.01492329 | 5.1445E-05 | 0.00060996 |
| AL928921.2 | 1.19290154 | 1.60766589 | 6.01158167 | 5.1717E-05 | 0.00061271 |
| TMEM19 | -0.5983418 | 5.24619386 | -6.0089478 | 5.1932E-05 | 0.00061442 |
| USP30 | 0.68910388 | 4.13142388 | 6.00882371 | 5.1942E-05 | 0.00061442 |
| PARP3 | -0.6156666 | 4.62486632 | -6.0081817 | 5.1995E-05 | 0.00061456 |
| MT-ND6 | -1.0711315 | 4.40010603 | -6.0074497 | 5.2055E-05 | 0.0006148 |
| GALNT14 | -0.5301929 | 5.74516903 | -6.0023197 | 5.2479E-05 | 0.00061883 |
| PPFIA3 | -0.661605 | 6.00635821 | -5.9960136 | 5.3005E-05 | 0.00062406 |
| FLNB | -0.5313852 | 7.70837905 | -5.9943372 | 5.3145E-05 | 0.00062523 |
| AL137003.2 | 0.75629646 | 3.25406215 | 5.99371529 | 5.3198E-05 | 0.00062536 |
| PER2 | 0.60657871 | 5.56409227 | 5.98889463 | 5.3605E-05 | 0.00062966 |
| SMTN | -0.6088787 | 5.41047905 | -5.9882071 | 5.3663E-05 | 0.00062986 |
| IRF4 | -0.8893912 | 2.63010989 | -5.9863967 | 5.3817E-05 | 0.00063117 |
| RASSF5 | -1.1266813 | 2.0022812 | -5.9855871 | 5.3886E-05 | 0.00063149 |
| POMGNT1 | -0.5001171 | 6.81487278 | -5.9844965 | 5.3979E-05 | 0.0006321 |
| COA5 | 0.58466896 | 5.1457366 | 5.98024336 | 5.4344E-05 | 0.0006356 |
| MAT1A | -1.4308783 | 1.93971063 | -5.9773425 | 5.4594E-05 | 0.00063782 |
| FOSB | 1.0913216 | 2.16841517 | 5.97678119 | 5.4643E-05 | 0.00063789 |
| THNSL1 | 0.64088433 | 4.09926162 | 5.97504114 | 5.4794E-05 | 0.00063916 |
| PCDH15 | 0.70878981 | 4.29668855 | 5.97326014 | 5.4948E-05 | 0.00064047 |
| NCK1 | -0.6387917 | 4.3948771 | -5.9722989 | 5.5032E-05 | 0.00064096 |
| BCKDK | -0.5412362 | 5.51908014 | -5.9708535 | 5.5158E-05 | 0.00064193 |
| AC006058.3 | 1.04146751 | 2.20698469 | 5.97003904 | 5.523E-05 | 0.00064224 |
| NSMCE4A | -0.5320044 | 6.03163272 | -5.9691773 | 5.5305E-05 | 0.00064224 |
| ARID5B | 0.6311335 | 6.21704649 | 5.96898409 | 5.5322E-05 | 0.00064224 |
| YJEFN3 | -0.8801296 | 2.79969692 | -5.9686181 | 5.5354E-05 | 0.00064224 |
| FAM45A | -0.6245694 | 4.71740684 | -5.9628481 | 5.5863E-05 | 0.00064715 |
| ADAM23 | -0.6500314 | 4.19935541 | -5.9608086 | 5.6044E-05 | 0.00064875 |
| RFNG | -0.572394 | 5.37956911 | -5.9596758 | 5.6145E-05 | 0.00064913 |
| ZCCHC4 | -0.5689176 | 5.08162195 | -5.9594714 | 5.6163E-05 | 0.00064913 |
| OSCP1 | 0.98340423 | 2.46082417 | 5.95842412 | 5.6256E-05 | 0.00064971 |
| ZBTB18 | 0.63771734 | 4.25023951 | 5.95793462 | 5.63E-05 | 0.00064972 |
| BRD4 | 0.64074215 | 6.20864543 | 5.95183564 | 5.6847E-05 | 0.00065554 |
| CTTNBP2NL | -0.6041832 | 4.74300353 | -5.9503121 | 5.6985E-05 | 0.00065663 |
| SYT7 | -0.9166075 | 2.71657692 | -5.9477997 | 5.7213E-05 | 0.00065825 |

| | | | | | |
|---|---|---|---|---|---|
| B3GNT2 | -0.5340559 | 5.35791021 | -5.9440076 | 5.7559E-05 | 0.00066173 |
| MXD1 | 1.09498266 | 2.13299693 | 5.9401731 | 5.791E-05 | 0.00066476 |
| SNIP1 | 0.54058013 | 5.27395015 | 5.93755261 | 5.8152E-05 | 0.00066703 |
| SLC44A2 | -0.6502648 | 4.41605772 | -5.9360719 | 5.8289E-05 | 0.00066781 |
| BECN1 | -0.5274597 | 5.54553596 | -5.935865 | 5.8308E-05 | 0.00066781 |
| TMOD2 | -0.6853571 | 3.97305038 | -5.9318588 | 5.8681E-05 | 0.00067157 |
| MRPL48 | 0.58413559 | 5.37652909 | 5.93132858 | 5.873E-05 | 0.00067163 |
| AP4S1 | 0.66299408 | 4.49130423 | 5.93035703 | 5.8821E-05 | 0.00067216 |
| PANK1 | 0.52332446 | 5.67455101 | 5.9297775 | 5.8875E-05 | 0.00067227 |
| HIST1H4D | 0.56548715 | 6.51148315 | 5.92494812 | 5.9329E-05 | 0.00067643 |
| GABRD | -0.580518 | 4.98718868 | -5.9166027 | 6.0123E-05 | 0.00068496 |
| ZNF225 | 0.6211603 | 4.33200489 | 5.91077241 | 6.0684E-05 | 0.0006898 |
| USP12 | -0.5240869 | 5.63977637 | -5.9083669 | 6.0917E-05 | 0.00069193 |
| UIMC1 | 0.58192006 | 5.76269489 | 5.90500375 | 6.1244E-05 | 0.00069512 |
| RASGEF1A | -0.5683111 | 5.04168773 | -5.9036057 | 6.1381E-05 | 0.00069579 |
| GORAB | 0.67849149 | 4.36516613 | 5.90346755 | 6.1394E-05 | 0.00069579 |
| SEC24A | -0.5575066 | 5.07940825 | -5.902567 | 6.1482E-05 | 0.00069626 |
| MCOLN3 | 0.58628594 | 4.63725887 | 5.89805233 | 6.1927E-05 | 0.00069997 |
| MOB3C | -0.6692674 | 4.12848183 | -5.8980392 | 6.1928E-05 | 0.00069997 |
| TIMP2 | 0.65851411 | 4.15296516 | 5.89714371 | 6.2016E-05 | 0.00070022 |
| MVB12A | -0.5408184 | 5.44632182 | -5.8916822 | 6.2559E-05 | 0.00070477 |
| DENND1C | -1.2991087 | 1.75951057 | -5.8895246 | 6.2775E-05 | 0.00070667 |
| TC2N | -1.3410536 | 1.01632822 | -5.8856985 | 6.316E-05 | 0.00071004 |
| MED29 | 0.52724526 | 6.27586683 | 5.88561718 | 6.3168E-05 | 0.00071004 |
| HFE | -0.7654309 | 3.65504983 | -5.885015 | 6.3228E-05 | 0.00071019 |
| HNRNPD | 0.69617637 | 6.23165926 | 5.88289331 | 6.3443E-05 | 0.00071208 |
| NLGN1 | 0.57800178 | 4.73933616 | 5.88231219 | 6.3502E-05 | 0.00071221 |
| TMEM99 | 0.81085948 | 3.07153406 | 5.88091627 | 6.3644E-05 | 0.00071274 |
| FASTKD5 | -0.5463758 | 5.68388339 | -5.8796983 | 6.3768E-05 | 0.00071307 |
| SLC35G1 | 0.55897635 | 5.47347324 | 5.87952774 | 6.3785E-05 | 0.00071307 |
| DAOA-AS1 | -0.577403 | 4.7644891 | -5.8792411 | 6.3814E-05 | 0.00071307 |
| HES1 | 0.54509477 | 5.21005466 | 5.87731165 | 6.4011E-05 | 0.00071475 |
| LDHA | -0.5055307 | 8.941709 | -5.8753903 | 6.4208E-05 | 0.00071536 |
| ZNF16 | 0.60215095 | 5.25068372 | 5.87367958 | 6.4384E-05 | 0.00071679 |
| LRRIQ3 | 0.79954572 | 2.92208653 | 5.87240604 | 6.4515E-05 | 0.00071749 |
| CASC15 | 0.92079723 | 2.64873951 | 5.86646844 | 6.513E-05 | 0.00072351 |
| PATL1 | 0.51777933 | 5.71053963 | 5.8609474 | 6.5708E-05 | 0.00072886 |
| NAA20 | -0.5114643 | 5.80677941 | -5.8577646 | 6.6044E-05 | 0.00073204 |
| PFKFB4 | -1.785211 | 5.06691784 | -5.8567793 | 6.6148E-05 | 0.00073213 |
| TLE3 | 0.51416607 | 6.27371575 | 5.8533553 | 6.6512E-05 | 0.00073507 |
| LMNTD2 | -1.3023044 | 1.48728697 | -5.8501112 | 6.6858E-05 | 0.00073783 |
| JDP2 | -0.7221475 | 4.40328589 | -5.8469017 | 6.7203E-05 | 0.0007408 |
| FAM169A | -0.5731232 | 4.84460949 | -5.8453135 | 6.7374E-05 | 0.00074136 |
| HK2 | -0.642401 | 6.26141858 | -5.8407545 | 6.7868E-05 | 0.00074554 |
| ELOVL4 | -0.5570487 | 5.33697685 | -5.8362182 | 6.8363E-05 | 0.00074971 |

| | | | | | |
|---|---|---|---|---|---|
| RNF121 | 0.55756731 | 5.1719286 | 5.83221773 | 6.8803E-05 | 0.00075326 |
| GCA | -0.6721719 | 4.73842571 | -5.8314642 | 6.8887E-05 | 0.00075362 |
| CFAP298 | -0.6560958 | 4.30282812 | -5.8278504 | 6.9287E-05 | 0.00075746 |
| MEX3B | 0.54323244 | 5.15497413 | 5.8264329 | 6.9445E-05 | 0.00075816 |
| DOK4 | -0.7054293 | 3.78683461 | -5.8261434 | 6.9477E-05 | 0.00075816 |
| ZDHHC21 | -0.6374137 | 4.02376248 | -5.8243446 | 6.9678E-05 | 0.00075885 |
| AKAP12 | 0.51344789 | 7.94291096 | 5.8240128 | 6.9715E-05 | 0.00075885 |
| TOMM40L | -0.5807804 | 5.60979873 | -5.8215887 | 6.9987E-05 | 0.00076071 |
| PRKCB | -0.5324444 | 5.86788094 | -5.8184419 | 7.0342E-05 | 0.00076402 |
| TUB | -0.5763782 | 5.55878785 | -5.8160583 | 7.0611E-05 | 0.0007664 |
| CORO2A | -0.8823271 | 3.20915829 | -5.8133393 | 7.0921E-05 | 0.0007692 |
| ZBTB17 | 0.50800198 | 6.73564765 | 5.81103456 | 7.1184E-05 | 0.00077151 |
| KERA | -0.7329552 | 3.73593129 | -5.8103684 | 7.126E-05 | 0.00077178 |
| PYCR3 | -0.5657891 | 5.7985281 | -5.8073209 | 7.161E-05 | 0.00077501 |
| GBF1 | 0.50980325 | 7.7188996 | 5.80433524 | 7.1954E-05 | 0.00077763 |
| AC209154.2 | -0.7339825 | 3.53641224 | -5.7978353 | 7.2711E-05 | 0.00078468 |
| TESMIN | -0.9099599 | 2.6412628 | -5.7966763 | 7.2846E-05 | 0.00078559 |
| CGNL1 | -1.0057894 | 2.20350576 | -5.7947278 | 7.3075E-05 | 0.00078694 |
| TDRD12 | 1.24626366 | 1.36584532 | 5.79198557 | 7.3398E-05 | 0.00078971 |
| TAF12 | 0.55844033 | 5.02609306 | 5.79147032 | 7.3459E-05 | 0.00078971 |
| PLEKHG5 | -0.8970944 | 3.80174426 | -5.7912204 | 7.3489E-05 | 0.00078971 |
| HSD3B7 | -1.0114378 | 2.5401988 | -5.78857 | 7.3803E-05 | 0.00079252 |
| PHF8 | -0.5752977 | 5.19737515 | -5.7881307 | 7.3855E-05 | 0.00079252 |
| NDUFS4 | 0.51893654 | 5.46451088 | 5.78542897 | 7.4177E-05 | 0.00079541 |
| SLC25A44 | -0.5166114 | 5.45919783 | -5.7827164 | 7.4502E-05 | 0.00079833 |
| CCT6B | 1.26266289 | 1.13408109 | 5.77963741 | 7.4873E-05 | 0.00080173 |
| ATG16L2 | -0.721729 | 4.3326283 | -5.7788514 | 7.4968E-05 | 0.00080218 |
| MT-CO3 | -0.5415765 | 8.3729163 | -5.776281 | 7.5279E-05 | 0.00080437 |
| APBB1 | 0.50453736 | 6.11986918 | 5.77157691 | 7.5852E-05 | 0.00080993 |
| CHRNB1 | -0.9405285 | 2.69173685 | -5.7666532 | 7.6457E-05 | 0.00081579 |
| TCEANC | 1.22208386 | 1.3073378 | 5.76622821 | 7.6509E-05 | 0.00081579 |
| OGFRL1 | -0.5733474 | 4.59315261 | -5.7648204 | 7.6683E-05 | 0.00081707 |
| ID2-AS1 | 0.85318749 | 2.94393177 | 5.76365228 | 7.6828E-05 | 0.00081746 |
| BAIAP2 | 0.52540159 | 5.92302405 | 5.76263197 | 7.6955E-05 | 0.00081824 |
| TBRG1 | -0.5413369 | 5.33472781 | -5.7593037 | 7.7369E-05 | 0.00082149 |
| ATP6V0E2-AS1 | -0.5943176 | 5.11159541 | -5.7505802 | 7.8467E-05 | 0.00083098 |
| ZNF181 | 0.56411811 | 4.68568123 | 5.7483463 | 7.8751E-05 | 0.00083324 |
| KRBOX4 | -0.7031795 | 3.68243388 | -5.7453832 | 7.9129E-05 | 0.00083607 |
| CBX7 | -1.0023786 | 2.43535002 | -5.7423149 | 7.9522E-05 | 0.00083905 |
| LRRIQ1 | 1.17527781 | 1.2832586 | 5.74135059 | 7.9646E-05 | 0.00083936 |
| CDK18 | -0.8136244 | 3.45009058 | -5.7412252 | 7.9662E-05 | 0.00083936 |
| PNPLA8 | 0.55227847 | 4.94364089 | 5.73858776 | 8.0003E-05 | 0.00084178 |
| EIF3J-DT | 0.61921219 | 4.35720125 | 5.73779676 | 8.0105E-05 | 0.00084227 |
| SNX19 | -0.5224168 | 5.51776759 | -5.7360685 | 8.0329E-05 | 0.00084404 |
| MEX3A | -0.5334497 | 5.23652436 | -5.7348113 | 8.0493E-05 | 0.00084517 |

| | | | | | |
|---|---|---|---|---|---|
| PRR14 | -0.5169058 | 5.4817809 | -5.7311496 | 8.0971E-05 | 0.00084843 |
| LINC00648 | 0.95680165 | 2.79792259 | 5.72790537 | 8.1397E-05 | 0.00085207 |
| FAM161B | 0.79515884 | 3.42002637 | 5.7272263 | 8.1487E-05 | 0.00085207 |
| ACVR1B | 0.51492161 | 5.50307335 | 5.72365516 | 8.1959E-05 | 0.00085642 |
| ZCCHC17 | -0.5643959 | 5.14829189 | -5.7198001 | 8.2473E-05 | 0.00086119 |
| SELENOM | -0.631744 | 4.55570267 | -5.7178916 | 8.2728E-05 | 0.00086326 |
| MYH14 | -0.7088185 | 3.85144935 | -5.7162677 | 8.2946E-05 | 0.00086494 |
| DLGAP1-AS1 | 0.63798809 | 4.01141419 | 5.71400097 | 8.3251E-05 | 0.00086693 |
| TCP11L1 | -0.6214096 | 4.23454984 | -5.7130962 | 8.3374E-05 | 0.0008676 |
| SRSF12 | 0.72754377 | 3.47448881 | 5.71215931 | 8.35E-05 | 0.00086773 |
| UNC5B | -0.7034353 | 4.86421312 | -5.709566 | 8.3852E-05 | 0.00087019 |
| SYT3 | -1.1123096 | 1.90224789 | -5.7079829 | 8.4068E-05 | 0.00087162 |
| PGK1 | -0.6949011 | 9.04050813 | -5.7077077 | 8.4105E-05 | 0.00087162 |
| AC005076.1 | 1.18862853 | 1.63000385 | 5.70687315 | 8.4219E-05 | 0.00087179 |
| ULK2 | -0.554286 | 5.18901826 | -5.7067408 | 8.4237E-05 | 0.00087179 |
| ALG10B | 0.54522354 | 4.70359993 | 5.70476788 | 8.4507E-05 | 0.00087399 |
| KIF24 | 0.65468975 | 4.90749376 | 5.7032177 | 8.472E-05 | 0.00087559 |
| C5 | 0.68671012 | 3.80033221 | 5.70002425 | 8.516E-05 | 0.00087954 |
| FAH | -0.5063799 | 5.74496693 | -5.6936615 | 8.6044E-05 | 0.00088798 |
| PRICKLE3 | -0.6623149 | 4.10036458 | -5.6918302 | 8.63E-05 | 0.00088798 |
| AC008438.1 | 1.34074624 | 0.94049667 | 5.69112421 | 8.6399E-05 | 0.00088798 |
| LSAMP | 0.68671164 | 4.17172182 | 5.69104991 | 8.641E-05 | 0.00088798 |
| TNFRSF21 | -0.6167301 | 4.84697224 | -5.6903716 | 8.6505E-05 | 0.00088798 |
| ZNF610 | 0.84694524 | 2.80077486 | 5.6831057 | 8.7532E-05 | 0.00089554 |
| CD46 | -0.5282453 | 7.25037407 | -5.683071 | 8.7537E-05 | 0.00089554 |
| USP40 | 0.50486682 | 6.00617267 | 5.68103229 | 8.7827E-05 | 0.0008973 |
| NFIC | -0.5891707 | 4.54427774 | -5.6781207 | 8.8244E-05 | 0.00090034 |
| SLC44A1 | -0.5250546 | 5.34730523 | -5.6725038 | 8.9054E-05 | 0.00090738 |
| NXT2 | 0.62646991 | 3.87536253 | 5.66864125 | 8.9615E-05 | 0.00091126 |
| MFSD8 | 0.56137916 | 4.73098268 | 5.66433081 | 9.0246E-05 | 0.00091644 |
| NRBP2 | -1.049115 | 4.31575121 | -5.65925 | 9.0995E-05 | 0.00092343 |
| PDZD4 | -0.5666708 | 5.25125469 | -5.6570067 | 9.1328E-05 | 0.00092619 |
| FAM133A | 0.57396612 | 4.5433744 | 5.65641174 | 9.1417E-05 | 0.00092647 |
| ABCB9 | -0.5193321 | 5.31512254 | -5.6433981 | 9.3376E-05 | 0.00094506 |
| ATP6V0A2 | 0.51595635 | 6.71168219 | 5.64139207 | 9.3682E-05 | 0.00094632 |
| PLAUR | -0.5012672 | 5.64339294 | -5.6413566 | 9.3688E-05 | 0.00094632 |
| SLC4A3 | -0.7326601 | 3.46049777 | -5.6404715 | 9.3823E-05 | 0.00094705 |
| SLC6A9 | -0.6147174 | 4.38885939 | -5.6373929 | 9.4296E-05 | 0.00095118 |
| COLEC12 | -1.0324754 | 2.04857923 | -5.6339128 | 9.4832E-05 | 0.00095596 |
| BCL11A | 0.62672827 | 4.0512803 | 5.6327954 | 9.5006E-05 | 0.00095707 |
| RNF141 | 0.52690167 | 5.11334569 | 5.62851557 | 9.5672E-05 | 0.00096314 |
| MIR4500HG | 1.15097214 | 1.30384991 | 5.62688808 | 9.5926E-05 | 0.00096506 |
| DNAJC25 | -0.7765739 | 3.04148002 | -5.6245047 | 9.63E-05 | 0.00096754 |
| INAVA | -0.5988774 | 4.40888246 | -5.6238359 | 9.6406E-05 | 0.00096764 |
| GYS1 | -0.8067003 | 3.52040519 | -5.6219063 | 9.671E-05 | 0.00096972 |

| | | | | | |
|---|---|---|---|---|---|
| CXorf56 | -0.5780424 | 4.74267216 | -5.6191861 | 9.7141E-05 | 0.00097129 |
| EMP1 | -0.769167 | 3.08000556 | -5.6188987 | 9.7186E-05 | 0.00097129 |
| ZNF540 | 1.22787069 | 1.31914387 | 5.61847759 | 9.7253E-05 | 0.00097131 |
| LOXL2 | -0.7893863 | 3.85914978 | -5.6178834 | 9.7348E-05 | 0.00097162 |
| ASL | -0.6330499 | 4.56483961 | -5.6163549 | 9.7591E-05 | 0.0009734 |
| TRPC3 | 1.12939894 | 1.74168633 | 5.61232456 | 9.8236E-05 | 0.00097919 |
| CCDC88B | -0.9705325 | 3.9458363 | -5.6106714 | 9.8502E-05 | 0.0009812 |
| CPPED1 | -0.6586886 | 3.80515091 | -5.6102296 | 9.8573E-05 | 0.00098126 |
| CCDC113 | -0.669974 | 3.76404116 | -5.6095832 | 9.8678E-05 | 0.00098165 |
| LPIN1 | -0.6947604 | 3.71695563 | -5.6089974 | 9.8772E-05 | 0.00098195 |
| KAT14 | 0.58267362 | 4.98656899 | 5.60809616 | 9.8918E-05 | 0.00098275 |
| OXCT1 | -0.512265 | 5.42390949 | -5.6072729 | 9.9051E-05 | 0.00098343 |
| ARHGEF39 | -0.568554 | 4.7571467 | -5.6055077 | 9.9337E-05 | 0.00098563 |
| CLCN2 | -0.6168404 | 4.46860708 | -5.603357 | 9.9688E-05 | 0.00098845 |
| GNAZ | -0.7104058 | 4.55744617 | -5.5941914 | 0.00010119 | 0.00100208 |
| UBE2W | -0.616142 | 4.44835289 | -5.5904323 | 0.00010182 | 0.00100564 |
| ZNF394 | 0.51015901 | 5.61978341 | 5.58398951 | 0.0001029 | 0.00101565 |
| C11orf71 | 0.85409865 | 2.6124914 | 5.58316928 | 0.00010304 | 0.00101635 |
| PIK3C2G | 0.73690258 | 3.18532279 | 5.58268592 | 0.00010312 | 0.0010165 |
| GPR157 | -0.540567 | 5.34603177 | -5.5791332 | 0.00010372 | 0.00102177 |
| TCF7 | 0.63087126 | 5.3147285 | 5.57857121 | 0.00010382 | 0.00102204 |
| VPS37D | -0.9184892 | 2.94325558 | -5.5768686 | 0.00010411 | 0.00102204 |
| APCDD1 | 1.07499231 | 2.08356646 | 5.57473904 | 0.00010447 | 0.00102383 |
| COLGALT2 | -0.8956266 | 2.61309161 | -5.5741482 | 0.00010457 | 0.00102416 |
| FBXL20 | 0.67115979 | 3.79892579 | 5.57335729 | 0.00010471 | 0.00102483 |
| SLC26A6 | -0.5606569 | 6.24004061 | -5.5719214 | 0.00010496 | 0.00102592 |
| PPP3CB | -0.5024322 | 5.45489356 | -5.5704677 | 0.00010521 | 0.00102638 |
| SOWAHD | -1.7783685 | -0.3804871 | -5.5690078 | 0.00010546 | 0.00102818 |
| PCDHAC1 | -0.6570387 | 3.96278917 | -5.5672067 | 0.00010577 | 0.0010299 |
| AL022068.1 | 0.79690644 | 2.82526284 | 5.56499297 | 0.00010616 | 0.00103298 |
| OSBPL2 | 0.51398381 | 5.34433436 | 5.56208803 | 0.00010666 | 0.00103725 |
| MPG | -0.6101128 | 4.3226485 | -5.5608698 | 0.00010688 | 0.00103798 |
| LENG9 | -0.8997551 | 2.45039177 | -5.5604862 | 0.00010695 | 0.00103798 |
| DLX6-AS1 | 0.5126059 | 6.46227718 | 5.56003728 | 0.00010702 | 0.00103802 |
| USP51 | 0.70314678 | 3.95678839 | 5.55968534 | 0.00010709 | 0.00103802 |
| MAP3K14 | 0.88650131 | 4.79591553 | 5.55262556 | 0.00010834 | 0.00104745 |
| CEP83-DT | 1.10513398 | 1.57396512 | 5.55195376 | 0.00010845 | 0.00104793 |
| CARD9 | -0.7609969 | 3.05774668 | -5.5482533 | 0.00010912 | 0.00105164 |
| TBKBP1 | 0.51053616 | 5.68541133 | 5.54780901 | 0.0001092 | 0.00105174 |
| IGDCC3 | 0.83940552 | 3.02875406 | 5.54563138 | 0.00010959 | 0.00105417 |
| FMNL1 | -0.6841741 | 3.57385457 | -5.5415672 | 0.00011032 | 0.0010599 |
| ESX1 | 0.60469233 | 4.57760715 | 5.5233061 | 0.00011369 | 0.00108863 |
| LPXN | 0.96769878 | 2.06595745 | 5.52324575 | 0.0001137 | 0.00108863 |
| CRISPLD1 | 0.5640843 | 4.81532452 | 5.52046103 | 0.00011422 | 0.00109252 |
| LRRC8A | 0.52247774 | 5.81978227 | 5.51707279 | 0.00011486 | 0.00109692 |

| | | | | | |
|---|---|---|---|---|---|
| ZZEF1 | 0.54578207 | 6.54793269 | 5.51687243 | 0.0001149 | 0.00109692 |
| FRYL | 0.54594712 | 7.7117424 | 5.51626692 | 0.00011502 | 0.00109733 |
| ZNF530 | 0.71945639 | 3.52721774 | 5.51471118 | 0.00011531 | 0.00109876 |
| AIF1L | -0.5045034 | 5.3918751 | -5.5131206 | 0.00011561 | 0.00110081 |
| GSTM4 | -0.6313193 | 4.14909506 | -5.5128161 | 0.00011567 | 0.00110081 |
| PDK1 | -1.5582184 | 3.82785831 | -5.5110102 | 0.00011602 | 0.00110188 |
| TOMM5 | 0.69974978 | 3.5703106 | 5.50853384 | 0.00011649 | 0.00110514 |
| MAMSTR | -0.7687567 | 3.26200665 | -5.5068363 | 0.00011682 | 0.00110665 |
| HOXB8 | 0.83211172 | 2.87661927 | 5.50656632 | 0.00011687 | 0.00110665 |
| GNB3 | -0.9326271 | 2.6434183 | -5.5032838 | 0.00011751 | 0.00111197 |
| DLG3 | -0.5244254 | 5.0555955 | -5.5014661 | 0.00011786 | 0.00111392 |
| FAT4 | 0.5622205 | 5.71308148 | 5.50087395 | 0.00011797 | 0.00111431 |
| NUDT8 | -0.7724715 | 3.28549959 | -5.4993857 | 0.00011826 | 0.00111566 |
| CCDC148 | 1.34758454 | 0.76338091 | 5.49491153 | 0.00011914 | 0.00112323 |
| HAND2 | 0.56850081 | 4.50971141 | 5.49362662 | 0.00011939 | 0.00112492 |
| AL645608.3 | -0.7173306 | 3.58402402 | -5.4872492 | 0.00012066 | 0.00113542 |
| STAP2 | -0.7770639 | 3.22602077 | -5.481735 | 0.00012176 | 0.00114297 |
| KCNK15 | -1.1624252 | 1.6110823 | -5.4808043 | 0.00012195 | 0.00114334 |
| SMPD2 | -0.5987579 | 4.46453198 | -5.4800453 | 0.0001221 | 0.00114334 |
| ULBP2 | -0.6991734 | 3.54143591 | -5.4732751 | 0.00012348 | 0.00115193 |
| TFAP2E | -1.2250109 | 1.19540346 | -5.4728584 | 0.00012356 | 0.00115202 |
| TSPYL2 | 0.55751998 | 5.03828493 | 5.47151035 | 0.00012384 | 0.00115334 |
| PSD4 | -1.4635844 | 0.47731074 | -5.4703271 | 0.00012408 | 0.00115472 |
| MT1F | 0.67715496 | 3.59509768 | 5.46845141 | 0.00012447 | 0.00115618 |
| PTER | -0.5225438 | 4.91776289 | -5.4673851 | 0.00012469 | 0.00115685 |
| HDAC11 | -0.6104499 | 4.4643541 | -5.464724 | 0.00012524 | 0.00115907 |
| SNAI3-AS1 | -0.6965453 | 3.76894379 | -5.4602725 | 0.00012617 | 0.00116622 |
| SLC39A8 | -0.5527523 | 4.80313721 | -5.4588285 | 0.00012647 | 0.0011683 |
| RGS14 | -0.6050158 | 4.60769068 | -5.457191 | 0.00012681 | 0.00116949 |
| SIK1 | -1.3413086 | 0.73812961 | -5.4545511 | 0.00012737 | 0.00117374 |
| FAM172A | 0.55023816 | 5.03608215 | 5.45397409 | 0.00012749 | 0.00117415 |
| ADARB1 | -0.6288784 | 3.97720025 | -5.4523619 | 0.00012783 | 0.00117658 |
| CRAT | -1.4889817 | 0.76548673 | -5.4498947 | 0.00012835 | 0.00118068 |
| ABCD1 | -0.6746461 | 3.7424345 | -5.4485506 | 0.00012864 | 0.00118259 |
| AC107375.1 | 0.88448389 | 2.15994675 | 5.44479364 | 0.00012944 | 0.00118848 |
| ZNF214 | 0.94401816 | 2.16841024 | 5.44446052 | 0.00012952 | 0.00118848 |
| EPHA2 | -0.5012751 | 6.78769049 | -5.4404592 | 0.00013038 | 0.00119567 |
| LINC01018 | -1.5903711 | 0.13609864 | -5.4383519 | 0.00013083 | 0.00119768 |
| EMX2OS | 0.63188016 | 4.30436797 | 5.43664756 | 0.0001312 | 0.0011989 |
| SHROOM1 | -0.5826828 | 4.30907603 | -5.435081 | 0.00013155 | 0.00120129 |
| ADM2 | -0.8647447 | 2.59699353 | -5.4331581 | 0.00013197 | 0.00120441 |
| LIMD2 | -0.6320651 | 3.95969077 | -5.4316499 | 0.0001323 | 0.00120642 |
| USP43 | -0.6129837 | 4.17593146 | -5.4302103 | 0.00013261 | 0.00120642 |
| MESP1 | -0.88295 | 2.60574818 | -5.4301843 | 0.00013262 | 0.00120642 |
| AC074117.1 | 0.62151913 | 3.8710162 | 5.42997699 | 0.00013267 | 0.00120642 |

| | | | | | |
|---|---|---|---|---|---|
| HDHD3 | -0.6515887 | 4.06420812 | -5.4269849 | 0.00013333 | 0.00121108 |
| EMILIN1 | -1.0219503 | 1.95254825 | -5.4259095 | 0.00013356 | 0.00121108 |
| C1orf21 | 0.63556217 | 3.8982216 | 5.42585095 | 0.00013358 | 0.00121108 |
| EML3 | -0.5032876 | 6.27273283 | -5.4227102 | 0.00013428 | 0.00121617 |
| DTX4 | -0.5924784 | 4.8419841 | -5.4226049 | 0.0001343 | 0.00121617 |
| JOSD2 | -0.7170032 | 3.42704961 | -5.4205632 | 0.00013476 | 0.00121812 |
| CDC42EP2 | -0.5588237 | 4.65044869 | -5.4178339 | 0.00013537 | 0.00122293 |
| ALX1 | 0.65531152 | 3.76379827 | 5.4142553 | 0.00013618 | 0.00122838 |
| NOTCH3 | 0.5101284 | 6.2154692 | 5.41408084 | 0.00013622 | 0.00122838 |
| AL136962.1 | -1.4747355 | 0.31225278 | -5.412816 | 0.0001365 | 0.00123024 |
| BATF3 | -0.590939 | 4.46176058 | -5.4122375 | 0.00013663 | 0.00123069 |
| MFSD10 | -0.5440219 | 6.42551189 | -5.4113382 | 0.00013684 | 0.0012318 |
| LAMP3 | -0.9163669 | 2.40176867 | -5.4098961 | 0.00013717 | 0.0012338 |
| CCDC78 | -0.7234359 | 4.40362316 | -5.4070903 | 0.00013781 | 0.00123832 |
| MANEA | 0.56737074 | 4.46617481 | 5.40655366 | 0.00013793 | 0.00123869 |
| RARA | 0.55838086 | 5.05556373 | 5.40179058 | 0.00013903 | 0.00124707 |
| WIPF2 | 0.53630382 | 5.14189657 | 5.40026822 | 0.00013938 | 0.00124802 |
| PAXIP1-AS1 | 0.64305296 | 3.82780937 | 5.39817124 | 0.00013987 | 0.00125164 |
| NTS | -1.2330023 | 1.10255722 | -5.3958781 | 0.00014041 | 0.00125569 |
| AL109615.3 | 1.60750246 | -0.0482633 | 5.39425778 | 0.00014078 | 0.00125834 |
| GMEB2 | 0.51494931 | 6.03689318 | 5.39031021 | 0.00014171 | 0.00126354 |
| LPAR2 | -0.8415851 | 2.65687473 | -5.3896567 | 0.00014187 | 0.00126354 |
| C1orf216 | 0.68654939 | 3.41377772 | 5.38656663 | 0.0001426 | 0.00126932 |
| RNF215 | -0.6280749 | 4.17715653 | -5.3774037 | 0.0001448 | 0.00128583 |
| DNAJC3-DT | 1.08865285 | 1.34769842 | 5.37501086 | 0.00014537 | 0.00129006 |
| ZMAT1 | 1.60834977 | 0.03064968 | 5.37473335 | 0.00014544 | 0.00129006 |
| CTSV | -0.5294816 | 5.15135228 | -5.3634313 | 0.00014821 | 0.00130715 |
| SLC45A1 | -0.7123356 | 3.52849823 | -5.3632149 | 0.00014826 | 0.00130715 |
| RET | -1.2199464 | 1.68380022 | -5.362805 | 0.00014837 | 0.00130715 |
| CDC37L1 | 0.52294239 | 4.80790043 | 5.36265273 | 0.0001484 | 0.00130715 |
| AC007743.1 | -0.573654 | 4.23858837 | -5.3617524 | 0.00014863 | 0.00130835 |
| COL26A1 | 1.29464258 | 1.95727743 | 5.35956101 | 0.00014917 | 0.00131239 |
| TRMT13 | 0.58216258 | 4.92166846 | 5.35658596 | 0.00014992 | 0.00131643 |
| STC2 | -0.9899106 | 6.02261232 | -5.3558347 | 0.0001501 | 0.00131676 |
| C1GALT1C1 | -0.5531871 | 4.44052135 | -5.3529394 | 0.00015083 | 0.00132106 |
| ZMIZ1 | 0.63216802 | 3.84619004 | 5.35284342 | 0.00015086 | 0.00132106 |
| SPEG | -0.797096 | 3.337199 | -5.3435792 | 0.00015321 | 0.00133704 |
| ATP11A | -0.5236138 | 4.99350855 | -5.3420543 | 0.0001536 | 0.00133841 |
| TCEA3 | -0.7700714 | 3.11095302 | -5.3399022 | 0.00015416 | 0.00134143 |
| TCTN1 | 0.50841963 | 5.00277622 | 5.33437252 | 0.00015559 | 0.00135128 |
| ABCG4 | -1.4978774 | 0.44162658 | -5.3341564 | 0.00015565 | 0.00135128 |
| MDC1 | 0.6404047 | 5.73242298 | 5.33177151 | 0.00015627 | 0.00135453 |
| KLF12 | 0.67608606 | 3.5442634 | 5.32794268 | 0.00015727 | 0.00136229 |
| CLIP2 | -0.504636 | 5.46027264 | -5.3248133 | 0.0001581 | 0.00136866 |
| PHC3 | 0.51581282 | 5.59236394 | 5.32360831 | 0.00015842 | 0.0013696 |

| | | | | | |
|---|---|---|---|---|---|
| WHAMM | 0.68754966 | 4.03923987 | 5.32341942 | 0.00015847 | 0.0013696 |
| CCDC28B | -0.5536432 | 4.66972008 | -5.3233816 | 0.00015848 | 0.0013696 |
| BARX1 | -0.5005625 | 5.3165485 | -5.3192905 | 0.00015957 | 0.00137745 |
| SEC14L5 | -1.5730529 | 0.15084298 | -5.3187695 | 0.00015971 | 0.00137787 |
| BOLA3-AS1 | -0.6363266 | 3.90997119 | -5.3177603 | 0.00015998 | 0.00137941 |
| FAM13B | 0.51409408 | 5.52029181 | 5.31717346 | 0.00016014 | 0.00137998 |
| HYAL3 | -0.8261838 | 3.20213217 | -5.3131074 | 0.00016123 | 0.00138784 |
| MYLK2 | 1.37564697 | 0.67674787 | 5.31136579 | 0.0001617 | 0.00139111 |
| SLC25A30 | -0.5692329 | 4.2663536 | -5.3108297 | 0.00016185 | 0.00139157 |
| ZNF618 | -0.5721973 | 4.34840661 | -5.3080907 | 0.00016259 | 0.00139719 |
| OTX1 | 0.68922847 | 3.42308925 | 5.30624319 | 0.0001631 | 0.00139994 |
| IL17RC | -0.6179112 | 4.17697515 | -5.2963065 | 0.00016584 | 0.00142026 |
| DRAM1 | 0.56801694 | 4.82909865 | 5.29520282 | 0.00016615 | 0.00142209 |
| WDR78 | 1.01973808 | 1.6842743 | 5.29368071 | 0.00016658 | 0.00142462 |
| MLXIPL | -1.2375113 | 1.8562423 | -5.293417 | 0.00016665 | 0.00142462 |
| OSER1-DT | 1.0109051 | 1.89920367 | 5.29313372 | 0.00016673 | 0.00142462 |
| PHF21A | 0.50839062 | 4.94970191 | 5.28488733 | 0.00016905 | 0.00144043 |
| TANGO2 | -0.5840477 | 4.19975654 | -5.2786989 | 0.00017082 | 0.00145141 |
| BRSK1 | -0.7665386 | 3.24394483 | -5.2783248 | 0.00017093 | 0.00145151 |
| ZKSCAN8 | 0.55361629 | 5.06169841 | 5.27673842 | 0.00017139 | 0.00145213 |
| GSDMB | -1.3323063 | 1.83659753 | -5.2762826 | 0.00017152 | 0.00145243 |
| TMEM79 | -0.6384392 | 4.07222343 | -5.2754997 | 0.00017174 | 0.00145272 |
| ACOT4 | -1.1638098 | 1.68971566 | -5.2748144 | 0.00017194 | 0.00145315 |
| TMIE | -1.2927357 | 0.97997442 | -5.274661 | 0.00017199 | 0.00145315 |
| ING3 | 0.5343388 | 4.71891384 | 5.27420739 | 0.00017212 | 0.00145345 |
| RAB9B | 1.16704003 | 1.06832094 | 5.26681724 | 0.00017427 | 0.00147082 |
| MMP17 | -0.5500408 | 4.53957257 | -5.2660039 | 0.00017451 | 0.00147201 |
| RABGGTB | -0.5118878 | 6.31354483 | -5.2640572 | 0.00017508 | 0.0014752 |
| ANKRD10 | 0.54016304 | 5.83440752 | 5.2609451 | 0.000176 | 0.00147998 |
| PSTK | 0.50017863 | 4.9618724 | 5.26047288 | 0.00017614 | 0.00148001 |
| MICAL2 | -0.8856217 | 2.62659462 | -5.2472519 | 0.00018011 | 0.00150574 |
| BMP5 | -0.5574491 | 4.32056292 | -5.2401422 | 0.00018228 | 0.00152064 |
| STOX1 | -0.548099 | 4.44253542 | -5.2386968 | 0.00018273 | 0.00152351 |
| MTHFSD | 0.50225235 | 4.95486168 | 5.23772073 | 0.00018303 | 0.00152518 |
| THUMPD2 | 0.54433425 | 5.4222841 | 5.23647626 | 0.00018341 | 0.00152671 |
| FAM131C | -1.1638753 | 1.38665126 | -5.2324621 | 0.00018466 | 0.00153624 |
| TAF1A-AS1 | 1.2532969 | 0.68203409 | 5.22841236 | 0.00018593 | 0.00154423 |
| NYAP1 | -1.0789442 | 2.40254947 | -5.2277379 | 0.00018614 | 0.00154515 |
| OSR1 | -0.8216017 | 2.68694898 | -5.2264467 | 0.00018655 | 0.00154767 |
| SERINC5 | -0.6142359 | 3.83211858 | -5.2242412 | 0.00018724 | 0.00155259 |
| ONECUT2 | 0.73364057 | 3.2818227 | 5.22282994 | 0.00018769 | 0.0015546 |
| C2CD4C | -0.7183435 | 3.3227419 | -5.221088 | 0.00018824 | 0.00155833 |
| ZNF98 | -0.6136204 | 3.73548833 | -5.2197932 | 0.00018865 | 0.00156003 |
| FRMPD1 | -1.2892168 | 0.91597787 | -5.21293 | 0.00019086 | 0.00157394 |
| TMEM102 | -0.7836503 | 2.84359574 | -5.2122501 | 0.00019108 | 0.00157489 |

| | | | | | |
|---|---|---|---|---|---|
| ZFHX4-AS1 | 0.60263613 | 3.88586069 | 5.20985231 | 0.00019185 | 0.00157871 |
| MYO15B | -0.8880745 | 4.53872146 | -5.209101 | 0.0001921 | 0.00157986 |
| BX664718.1 | 1.30117542 | 0.7920144 | 5.20712489 | 0.00019274 | 0.00158343 |
| TTC9 | -0.7589472 | 2.92566274 | -5.204325 | 0.00019365 | 0.00158757 |
| ENTPD3-AS1 | 1.06877738 | 1.64774504 | 5.20403786 | 0.00019375 | 0.00158757 |
| SPSB4 | 1.22913974 | 1.08138984 | 5.19748619 | 0.00019591 | 0.00160251 |
| DOK3 | -1.0384614 | 2.83230071 | -5.1959845 | 0.00019641 | 0.00160572 |
| MAFG-DT | -0.5811577 | 4.38182943 | -5.1943513 | 0.00019695 | 0.00160775 |
| SDK2 | 0.71802011 | 3.95387041 | 5.19316899 | 0.00019735 | 0.00160847 |
| TCL1B | -1.2939482 | 0.93831097 | -5.191515 | 0.0001979 | 0.00161096 |
| EPC1 | 0.50283474 | 5.44076865 | 5.18854505 | 0.0001989 | 0.00161648 |
| PARP16 | 0.62102782 | 4.33259557 | 5.18703477 | 0.00019941 | 0.00161975 |
| TMEM141 | -0.6347392 | 3.77915496 | -5.1843932 | 0.0002003 | 0.00162527 |
| SPAG1 | -0.5367111 | 4.78082249 | -5.1840449 | 0.00020042 | 0.00162536 |
| IPP | 0.50465063 | 4.75590525 | 5.1808157 | 0.00020152 | 0.00162994 |
| IRF1 | 0.61542938 | 3.98022964 | 5.18080729 | 0.00020152 | 0.00162994 |
| LRRN3 | 0.64666871 | 3.36626337 | 5.17874775 | 0.00020223 | 0.0016339 |
| SNX22 | -0.7088521 | 3.13325263 | -5.1742868 | 0.00020376 | 0.00164542 |
| MT-ATP8 | -1.1006522 | 2.54612245 | -5.1730356 | 0.0002042 | 0.00164804 |
| TMEM44 | -0.6059019 | 4.11947995 | -5.1678313 | 0.00020601 | 0.00166177 |
| ZNF549 | 0.63898216 | 3.59619208 | 5.16345739 | 0.00020754 | 0.00167237 |
| NUMBL | -0.5602193 | 4.29304457 | -5.1627793 | 0.00020778 | 0.00167341 |
| AL133338.1 | 0.9816146 | 1.72681007 | 5.15889208 | 0.00020916 | 0.0016827 |
| CREBBP | 0.50951447 | 5.91945165 | 5.15345156 | 0.0002111 | 0.00169472 |
| SPSB2 | -0.544035 | 4.37050851 | -5.1508969 | 0.00021202 | 0.00169939 |
| COQ5 | 0.5070828 | 5.68825572 | 5.14585249 | 0.00021384 | 0.0017113 |
| AC093484.4 | -1.0015346 | 1.5856573 | -5.143314 | 0.00021477 | 0.00171683 |
| MAP1B | -0.5673026 | 6.29879575 | -5.1398302 | 0.00021604 | 0.00172254 |
| SDCBP2-AS1 | 0.61872152 | 3.64859668 | 5.13065729 | 0.00021944 | 0.00174596 |
| RASSF4 | -0.8112558 | 2.81938609 | -5.1270247 | 0.0002208 | 0.00175334 |
| SULT2B1 | -1.6902332 | 0.02981031 | -5.1265891 | 0.00022097 | 0.00175334 |
| LINC02345 | 1.10368915 | 1.28893561 | 5.12489506 | 0.0002216 | 0.00175499 |
| ZNF347 | -0.5251715 | 4.5687426 | -5.1243737 | 0.0002218 | 0.00175499 |
| WDFY2 | 0.57724468 | 4.10279737 | 5.12374635 | 0.00022204 | 0.00175555 |
| ZC2HC1C | 0.61189195 | 3.91742226 | 5.11938006 | 0.0002237 | 0.00176681 |
| FAT1 | 0.51643789 | 7.06181466 | 5.11245632 | 0.00022635 | 0.00178406 |
| UTP25 | -0.5026183 | 5.10817883 | -5.1083983 | 0.00022792 | 0.00179364 |
| ANXA4 | -0.5498874 | 4.79861497 | -5.1072106 | 0.00022838 | 0.00179635 |
| NBPF4 | 0.92028725 | 1.98111667 | 5.10365799 | 0.00022977 | 0.00180632 |
| AL450332.1 | 1.32251814 | 0.317971 | 5.10259986 | 0.00023019 | 0.00180771 |
| POLR3GL | 0.53720107 | 4.39834508 | 5.09847193 | 0.00023181 | 0.00181954 |
| MYOF | -0.9803951 | 2.20221684 | -5.0979703 | 0.00023201 | 0.00182016 |
| DDX59 | 0.5010154 | 4.97591263 | 5.09763728 | 0.00023214 | 0.00182025 |
| ASPHD2 | -0.8377857 | 2.67099224 | -5.0949553 | 0.00023321 | 0.00182638 |
| SPTBN5 | -0.9436268 | 1.91392345 | -5.0947588 | 0.00023329 | 0.00182638 |

| | | | | | |
|---|---|---|---|---|---|
| SLC38A9 | 0.56445698 | 4.05202545 | 5.09283419 | 0.00023406 | 0.00183049 |
| AC005288.1 | -0.7094616 | 2.97316868 | -5.085684 | 0.00023693 | 0.00185107 |
| BBS7 | -0.5441475 | 4.35475957 | -5.0849829 | 0.00023721 | 0.00185233 |
| ABHD15 | -1.4606186 | 0.21522034 | -5.0845695 | 0.00023738 | 0.00185269 |
| PEA15 | -0.5035215 | 6.40669595 | -5.0820352 | 0.00023841 | 0.00185727 |
| AC116565.1 | 1.3451081 | 0.58915813 | 5.08191888 | 0.00023846 | 0.00185727 |
| AC010422.6 | 1.37114206 | 0.59038533 | 5.0801241 | 0.00023919 | 0.00186106 |
| MYORG | -0.922749 | 2.09455698 | -5.0787461 | 0.00023976 | 0.00186353 |
| TMEM42 | 0.52877441 | 4.52575877 | 5.07056354 | 0.00024313 | 0.00188688 |
| ZFP69B | 0.51084004 | 4.74859386 | 5.07024731 | 0.00024327 | 0.00188693 |
| EFEMP2 | -0.8423895 | 3.44315149 | -5.0619392 | 0.00024675 | 0.00191198 |
| LACTB | -0.567793 | 4.06979626 | -5.0608801 | 0.0002472 | 0.00191218 |
| AL512625.2 | 0.76771183 | 2.80213599 | 5.06041115 | 0.00024739 | 0.00191218 |
| SH3RF2 | 0.58311165 | 4.82151933 | 5.06038469 | 0.00024741 | 0.00191218 |
| DNHD1 | -0.8902906 | 2.75425182 | -5.0595828 | 0.00024775 | 0.00191286 |
| PPP1R3G | -1.7151052 | 0.75770203 | -5.0553542 | 0.00024955 | 0.00192186 |
| UNC93B1 | -0.752648 | 3.34223558 | -5.0530888 | 0.00025052 | 0.00192835 |
| ANGPTL1 | 0.70610208 | 3.51244381 | 5.05149579 | 0.0002512 | 0.00193165 |
| ZFR2 | 0.52317733 | 4.74376044 | 5.05094543 | 0.00025144 | 0.00193227 |
| ENO2 | -0.8690826 | 6.21212869 | -5.0485318 | 0.00025248 | 0.00193755 |
| TJP3 | -1.0169333 | 2.35734979 | -5.046142 | 0.00025351 | 0.00194451 |
| ZBED5-AS1 | 0.86324735 | 2.23772172 | 5.04157521 | 0.00025551 | 0.00195682 |
| PLD2 | -0.5656866 | 4.45305918 | -5.0381455 | 0.00025701 | 0.00196538 |
| ARL14EP | -0.5167261 | 4.43345482 | -5.0369678 | 0.00025753 | 0.00196835 |
| MTMR10 | 0.59840927 | 4.30703891 | 5.0249007 | 0.00026292 | 0.00200245 |
| AC012414.5 | 1.4047254 | 0.30309454 | 5.02312753 | 0.00026372 | 0.00200654 |
| PLEKHM3 | 0.75029426 | 2.86939191 | 5.02076943 | 0.00026479 | 0.00201367 |
| ARL6IP6 | 0.51469764 | 4.47986155 | 5.01910665 | 0.00026555 | 0.0020174 |
| DACT3 | 0.84027236 | 2.4285658 | 5.01414234 | 0.00026782 | 0.00203264 |
| ASAH2B | -0.6958952 | 2.94893817 | -5.0115337 | 0.00026903 | 0.00204074 |
| CHRM4 | 0.89472211 | 2.41572161 | 5.00988312 | 0.00026979 | 0.00204552 |
| GPR27 | -0.5678435 | 4.03538331 | -5.0074091 | 0.00027094 | 0.00205218 |
| ATP8B3 | -0.7686482 | 3.76359652 | -5.0059486 | 0.00027162 | 0.00205528 |
| PANDAR | -1.0137956 | 1.51999527 | -5.0038223 | 0.00027262 | 0.00206178 |
| THAP7-AS1 | 0.65694221 | 3.25414454 | 4.99600491 | 0.00027631 | 0.00208344 |
| AC009005.1 | 0.69177035 | 3.16439707 | 4.98884863 | 0.00027973 | 0.00210506 |
| DOC2A | -0.5854923 | 4.50164872 | -4.9877273 | 0.00028027 | 0.00210808 |
| SMAD9 | 0.67878027 | 3.16621507 | 4.98508368 | 0.00028155 | 0.00211665 |
| FGFBP3 | 0.72319697 | 2.87422011 | 4.98413824 | 0.00028201 | 0.00211806 |
| CCDC40 | 0.62489517 | 3.78139496 | 4.96772314 | 0.0002901 | 0.00217229 |
| PCOLCE2 | 0.54756695 | 4.1891037 | 4.96662191 | 0.00029065 | 0.00217427 |
| AC060765.1 | 1.52376985 | -0.17616 | 4.96580277 | 0.00029106 | 0.00217627 |
| CLMN | -0.7763038 | 2.84942848 | -4.9647309 | 0.0002916 | 0.00217922 |
| ZNF713 | 0.57383316 | 3.84632531 | 4.96426287 | 0.00029183 | 0.0021799 |
| IFT74 | 0.57540492 | 4.12051474 | 4.96236413 | 0.00029279 | 0.00218598 |

| | | | | | |
|---|---|---|---|---|---|
| SLC35D3 | -0.7881956 | 2.67309878 | -4.9604859 | 0.00029374 | 0.00219199 |
| LIPG | -1.0400558 | 1.53577472 | -4.9543625 | 0.00029686 | 0.00221091 |
| ZNRF1 | -0.5034069 | 4.5806838 | -4.9528354 | 0.00029765 | 0.00221457 |
| TMEM151A | -0.7101152 | 3.28739207 | -4.952159 | 0.00029799 | 0.00221549 |
| UBE2F | -0.6116321 | 3.60182645 | -4.9512223 | 0.00029848 | 0.00221748 |
| SERINC2 | -0.7739696 | 2.84395874 | -4.9496033 | 0.00029931 | 0.0022226 |
| AKAP6 | 0.65802096 | 3.4261811 | 4.94827485 | 0.0003 | 0.00222661 |
| PPP1R16B | -0.5684321 | 4.13574817 | -4.9447837 | 0.00030181 | 0.00223517 |
| RASD2 | -1.2722033 | 0.57117106 | -4.9441472 | 0.00030215 | 0.00223517 |
| PCDHGB3 | 1.30391603 | 0.72504431 | 4.93854039 | 0.00030509 | 0.00225332 |
| PRPH | -1.4412799 | 0.31575615 | -4.936164 | 0.00030634 | 0.00226039 |
| GLRA3 | 1.17005987 | 0.86890348 | 4.93241963 | 0.00030833 | 0.00227396 |
| NTRK1 | -1.1317038 | 1.24550985 | -4.931319 | 0.00030892 | 0.00227718 |
| KCNMB4 | 0.55979426 | 4.00947174 | 4.93014344 | 0.00030955 | 0.0022796 |
| RERGL | 0.76858833 | 2.50922532 | 4.92661214 | 0.00031144 | 0.00228942 |
| RUNDC3A | 0.78142096 | 3.08347843 | 4.92653358 | 0.00031149 | 0.00228942 |
| GRK5 | -0.5717745 | 3.93296827 | -4.9243326 | 0.00031267 | 0.00229526 |
| COL6A1 | -0.7280541 | 4.6178731 | -4.9197903 | 0.00031514 | 0.00230956 |
| DCN | -0.5051388 | 4.41044342 | -4.9159311 | 0.00031725 | 0.00232055 |
| SLC9A3R2 | -0.5301947 | 4.34955029 | -4.9147253 | 0.00031792 | 0.00232316 |
| B4GALNT3 | -0.6100291 | 3.66746217 | -4.9120212 | 0.00031941 | 0.00232904 |
| NFKBID | -0.7621079 | 2.74529112 | -4.9119522 | 0.00031945 | 0.00232904 |
| MT-ND2 | -0.8659835 | 5.50744124 | -4.9059239 | 0.0003228 | 0.00234864 |
| OLMALINC | 0.51169711 | 4.44353409 | 4.90369007 | 0.00032405 | 0.00235371 |
| AC016355.1 | 1.16014881 | 0.7201123 | 4.90357116 | 0.00032412 | 0.00235371 |
| CPED1 | 0.71361945 | 3.73860854 | 4.90073639 | 0.00032572 | 0.00236303 |
| FBLL1 | 0.62682357 | 4.05723405 | 4.89852685 | 0.00032696 | 0.00237097 |
| ZDHHC11 | -0.732454 | 2.83295239 | -4.8910443 | 0.00033123 | 0.00239619 |
| WDR63 | 0.87819951 | 1.96984821 | 4.88823628 | 0.00033285 | 0.0024033 |
| CRELD1 | -0.5595032 | 4.424924 | -4.8869798 | 0.00033358 | 0.00240429 |
| ARHGEF19 | -0.652619 | 3.52438902 | -4.886901 | 0.00033362 | 0.00240429 |
| IL10RA | -1.0652394 | 1.53570057 | -4.8860593 | 0.00033411 | 0.00240655 |
| CD3EAP | -0.5207567 | 4.41569297 | -4.885333 | 0.00033453 | 0.0024074 |
| KNDC1 | -0.7694123 | 3.06010874 | -4.8847173 | 0.00033489 | 0.00240883 |
| AC015712.2 | -0.8823822 | 2.00020594 | -4.8807197 | 0.00033722 | 0.002421 |
| SLC10A4 | -1.1212338 | 1.08491475 | -4.8784284 | 0.00033857 | 0.00242835 |
| BFSP1 | 0.85305453 | 2.25140509 | 4.87565124 | 0.0003402 | 0.00243893 |
| HOXA5 | 0.68129401 | 3.37489789 | 4.87156861 | 0.00034262 | 0.00245164 |
| TUBG2 | -0.6260222 | 3.71506158 | -4.8705958 | 0.0003432 | 0.00245393 |
| CCDC15 | 0.54680076 | 4.00780741 | 4.87048905 | 0.00034327 | 0.00245393 |
| CBWD2 | 0.52151337 | 4.27790427 | 4.86931608 | 0.00034397 | 0.0024547 |
| SLC45A3 | -0.5414349 | 4.4266307 | -4.8692204 | 0.00034403 | 0.0024547 |
| RBKS | 0.77693724 | 2.52023759 | 4.86707155 | 0.00034531 | 0.00246157 |
| ZNF614 | 0.51984068 | 4.252439 | 4.86635337 | 0.00034574 | 0.00246348 |
| MYT1 | -0.9623934 | 2.38207987 | -4.8608868 | 0.00034904 | 0.00247806 |

| | | | | | |
|---|---|---|---|---|---|
| PDLIM1 | -0.5124175 | 4.50192032 | -4.8606215 | 0.00034921 | 0.00247806 |
| CFAP69 | 0.68715789 | 3.12823641 | 4.8603232 | 0.00034939 | 0.00247806 |
| BEND7 | 0.52080599 | 4.47828316 | 4.85962335 | 0.00034981 | 0.00247964 |
| MATN2 | -0.6796094 | 3.99967787 | -4.8560845 | 0.00035197 | 0.00249028 |
| JMJD4 | -0.5996927 | 4.37046234 | -4.85472 | 0.00035281 | 0.00249387 |
| EPN3 | -0.8950102 | 1.78635693 | -4.8543825 | 0.00035301 | 0.00249416 |
| CRLF1 | -0.6540596 | 3.62679572 | -4.8531425 | 0.00035378 | 0.00249722 |
| ABHD4 | 0.52539661 | 4.33882667 | 4.85152372 | 0.00035477 | 0.0025031 |
| SQOR | -0.7099963 | 3.01364433 | -4.8503186 | 0.00035552 | 0.00250718 |
| RELB | 0.87518509 | 4.51613199 | 4.84996901 | 0.00035574 | 0.00250754 |
| C21orf91 | -0.5255563 | 4.40032692 | -4.8470318 | 0.00035756 | 0.00251687 |
| MMP10 | -1.3773601 | 0.24004232 | -4.8446349 | 0.00035905 | 0.00252622 |
| OTUB2 | -0.6215775 | 3.58491323 | -4.8439275 | 0.0003595 | 0.00252713 |
| DLL3 | -0.6391246 | 3.41307781 | -4.8438212 | 0.00035956 | 0.00252713 |
| TMEM255B | -1.1279816 | 1.11477817 | -4.8385356 | 0.00036289 | 0.00254491 |
| TMEM121B | 0.86176236 | 2.73102974 | 4.83779713 | 0.00036335 | 0.00254701 |
| SLC9A5 | -0.5373055 | 4.20764585 | -4.8349345 | 0.00036517 | 0.00255737 |
| LINC01140 | -0.8783468 | 2.20527037 | -4.8346453 | 0.00036535 | 0.00255748 |
| PTPRU | -0.537312 | 5.06396533 | -4.8324853 | 0.00036673 | 0.00256058 |
| ASAP2 | -0.5058483 | 4.82425964 | -4.832114 | 0.00036697 | 0.00256058 |
| GVQW3 | 0.83970917 | 3.14501203 | 4.82093209 | 0.00037419 | 0.00260489 |
| WASF1 | 0.56924268 | 3.73213186 | 4.81939363 | 0.0003752 | 0.00260949 |
| AUTS2 | 0.58646914 | 4.16264511 | 4.81388825 | 0.00037882 | 0.00263029 |
| ZNF79 | 0.55577811 | 4.4060954 | 4.81379097 | 0.00037888 | 0.00263029 |
| ZNF425 | 0.64661244 | 3.30296917 | 4.81295616 | 0.00037943 | 0.00263291 |
| LINC00858 | 0.86471951 | 2.15064037 | 4.80970769 | 0.00038159 | 0.00264667 |
| TTI2 | 0.53440673 | 4.29699001 | 4.80596711 | 0.00038409 | 0.00265914 |
| AC004969.1 | 0.82815434 | 2.12399983 | 4.80407671 | 0.00038536 | 0.00266427 |
| CEBPB-AS1 | 1.13590601 | 0.90326893 | 4.80269168 | 0.00038629 | 0.0026695 |
| ANAPC15 | -0.6417106 | 3.27967938 | -4.8022192 | 0.00038661 | 0.00267049 |
| BCL2L11 | 0.62025919 | 3.31653864 | 4.80188698 | 0.00038684 | 0.00267082 |
| N4BP2L1 | 1.21596981 | 0.59890442 | 4.80099329 | 0.00038744 | 0.0026725 |
| HFM1 | 1.41467445 | -0.29545 | 4.80074382 | 0.00038761 | 0.0026725 |
| MYBPH | -1.0662519 | 1.38933267 | -4.7970987 | 0.00039009 | 0.00268713 |
| PRRT4 | -0.8612322 | 2.59154405 | -4.7937617 | 0.00039237 | 0.0026967 |
| KIFC3 | -0.6039218 | 3.57687476 | -4.77883 | 0.00040275 | 0.00275428 |
| GALNT6 | -0.531504 | 4.27689889 | -4.7738767 | 0.00040625 | 0.00277342 |
| KCNN1 | -0.6780716 | 3.91679088 | -4.7720373 | 0.00040756 | 0.00278094 |
| KLC3 | -1.3190856 | 0.58066091 | -4.7710974 | 0.00040823 | 0.00278301 |
| ATG9A | -0.5277089 | 5.52902679 | -4.7625589 | 0.00041438 | 0.00282112 |
| LINC02241 | 1.53611208 | -0.2761446 | 4.75889774 | 0.00041705 | 0.00283417 |
| SCN5A | -0.9910052 | 1.77731166 | -4.7580541 | 0.00041767 | 0.00283582 |
| CALCOCO1 | 0.57331758 | 4.19717163 | 4.75776028 | 0.00041788 | 0.00283601 |
| GABRG3 | 0.65587114 | 3.09634519 | 4.75562662 | 0.00041945 | 0.00284408 |
| GABRG1 | -1.1588084 | 0.65064678 | -4.754615 | 0.00042019 | 0.00284785 |

| | | | | | |
|---|---|---|---|---|---|
| HAGH | -0.6166573 | 4.63588993 | -4.7519348 | 0.00042217 | 0.00285487 |
| EEF1AKMT4 | -0.6419479 | 3.50925793 | -4.7505356 | 0.00042321 | 0.00286061 |
| TASP1 | -0.5088065 | 4.31218102 | -4.7492485 | 0.00042416 | 0.00286579 |
| ARHGDIG | -0.9686452 | 1.72437155 | -4.7487939 | 0.0004245 | 0.00286679 |
| KLLN | 0.65219379 | 3.15014844 | 4.74441032 | 0.00042778 | 0.00288506 |
| AC040162.1 | -0.6864008 | 2.91076741 | -4.7440863 | 0.00042802 | 0.00288541 |
| AC004943.2 | 0.54278323 | 4.66160255 | 4.74366333 | 0.00042834 | 0.00288627 |
| ZNF423 | 0.60024054 | 3.58462961 | 4.74279917 | 0.00042899 | 0.00288936 |
| PRDM11 | 0.5590786 | 4.07186186 | 4.73941423 | 0.00043154 | 0.00290528 |
| CRACR2B | -0.7118789 | 3.05493247 | -4.7388805 | 0.00043195 | 0.00290671 |
| AKAP3 | 0.95563302 | 1.57085271 | 4.73574658 | 0.00043433 | 0.00292144 |
| HIRIP3 | 0.61595343 | 4.74512218 | 4.73445477 | 0.00043531 | 0.00292677 |
| AL450998.2 | 0.58224443 | 3.68499323 | 4.73339609 | 0.00043612 | 0.00293091 |
| ZDHHC1 | -0.9073496 | 1.94248263 | -4.7322486 | 0.000437 | 0.00293552 |
| CDON | -0.5114949 | 4.42973754 | -4.7293981 | 0.0004392 | 0.00294763 |
| ARL10 | -0.631312 | 3.29035713 | -4.7288899 | 0.00043959 | 0.00294895 |
| CARD19 | -0.5343812 | 4.1051591 | -4.722169 | 0.00044481 | 0.0029803 |
| LIF | -1.0454101 | 1.60137702 | -4.719329 | 0.00044703 | 0.00299095 |
| PAOX | -0.7584398 | 2.83127826 | -4.7183341 | 0.00044781 | 0.00299486 |
| SHC4 | 1.51130602 | -0.1049806 | 4.7174212 | 0.00044853 | 0.00299834 |
| AC136475.3 | -1.3080967 | 0.27922424 | -4.7167124 | 0.00044909 | 0.0030006 |
| AL136379.1 | -1.3258386 | 0.12302212 | -4.7144819 | 0.00045086 | 0.00300723 |
| FAM227B | 0.94089732 | 1.48780646 | 4.71138129 | 0.00045332 | 0.00301835 |
| BORA | -0.5058693 | 4.31758103 | -4.7109926 | 0.00045363 | 0.00301909 |
| ZNF730 | -0.5181658 | 4.00210324 | -4.7067702 | 0.00045701 | 0.00303492 |
| AP003390.1 | -0.81574 | 2.42273515 | -4.7054077 | 0.00045811 | 0.00304087 |
| GLI3 | 0.63665004 | 3.32248784 | 4.70479977 | 0.0004586 | 0.00304279 |
| LINC01770 | -1.3557418 | 0.57267179 | -4.7014406 | 0.00046132 | 0.00305547 |
| METTL18 | 0.58861436 | 3.38515632 | 4.69913377 | 0.00046319 | 0.00306655 |
| ZNF211 | 0.53984348 | 3.97760141 | 4.69495307 | 0.00046661 | 0.00308246 |
| GLI2 | -0.6761996 | 3.0223288 | -4.6913503 | 0.00046958 | 0.00310072 |
| ZNF600 | -0.5543988 | 3.80793506 | -4.6908983 | 0.00046996 | 0.00310184 |
| VIM-AS1 | 0.80990849 | 2.01394956 | 4.68888402 | 0.00047163 | 0.00311015 |
| NALT1 | -1.4281901 | 0.00348576 | -4.6881133 | 0.00047227 | 0.00311302 |
| DYNC2LI1 | 0.50417151 | 4.22558932 | 4.68627381 | 0.0004738 | 0.00312176 |
| ZNF572 | 0.79707434 | 2.16574885 | 4.67740664 | 0.00048126 | 0.00316541 |
| C18orf54 | -0.5317486 | 3.75179185 | -4.6735854 | 0.00048451 | 0.00317957 |
| SMIM1 | -0.8812206 | 2.06105776 | -4.6672658 | 0.00048994 | 0.00320666 |
| KHK | -0.6256543 | 3.56744283 | -4.6619471 | 0.00049456 | 0.00322873 |
| CHST2 | -0.8118984 | 2.24647303 | -4.6617593 | 0.00049473 | 0.00322873 |
| ICOSLG | -1.1119047 | 0.89706691 | -4.6577642 | 0.00049823 | 0.00324458 |
| PPP4R4 | -0.6611193 | 3.44735112 | -4.6568748 | 0.00049901 | 0.00324828 |
| CD70 | -0.5495233 | 4.14721672 | -4.656246 | 0.00049956 | 0.00325049 |
| TRPM4 | -0.5728247 | 3.87067187 | -4.653828 | 0.0005017 | 0.0032602 |
| AL163051.1 | 0.77768642 | 2.31066708 | 4.65202017 | 0.00050331 | 0.00326615 |

| | | | | | |
|---|---|---|---|---|---|
| ROGDI | -0.5265654 | 4.19522956 | -4.647938 | 0.00050695 | 0.00328583 |
| SNAI3 | -0.7548982 | 2.59156635 | -4.647582 | 0.00050727 | 0.00328648 |
| ZNF18 | 0.52001335 | 4.08178031 | 4.6453198 | 0.0005093 | 0.00329401 |
| KCNC2 | 0.64579331 | 3.09973449 | 4.64050803 | 0.00051365 | 0.0033193 |
| MATN1-AS1 | -1.0444074 | 1.28342782 | -4.6402277 | 0.0005139 | 0.00331953 |
| TRAM2-AS1 | 0.5054031 | 4.14403463 | 4.63994444 | 0.00051416 | 0.00331977 |
| HOXC4 | -0.5080421 | 4.21895949 | -4.6390574 | 0.00051496 | 0.00332215 |
| OLFML2B | -1.3088091 | 0.53226102 | -4.6375211 | 0.00051637 | 0.00332766 |
| PSD2 | -1.102741 | 0.90632749 | -4.6373975 | 0.00051648 | 0.00332766 |
| AL162411.1 | 0.78891221 | 2.70813453 | 4.6345224 | 0.00051911 | 0.00334035 |
| SYNE2 | 0.52614312 | 8.00984902 | 4.63402262 | 0.00051957 | 0.00334126 |
| HSD17B6 | -0.9828547 | 1.44010536 | -4.6335226 | 0.00052003 | 0.003342 |
| NR4A2 | -0.7695481 | 2.80094189 | -4.6323666 | 0.00052109 | 0.00334742 |
| GPCPD1 | 0.52691021 | 3.91410379 | 4.62375686 | 0.00052909 | 0.00339364 |
| ZNF792 | -0.5519613 | 3.96593736 | -4.6231686 | 0.00052964 | 0.00339364 |
| METTL22 | 0.50024809 | 4.31559372 | 4.62316146 | 0.00052965 | 0.00339364 |
| MAMDC4 | -0.6462402 | 4.47232586 | -4.6226287 | 0.00053015 | 0.00339406 |
| MT-ND1 | -0.6579745 | 5.83335799 | -4.6218251 | 0.0005309 | 0.00339745 |
| PLA2G15 | -0.5004523 | 4.88287196 | -4.6201869 | 0.00053244 | 0.00340588 |
| PYM1 | 0.55008164 | 5.07309749 | 4.61563488 | 0.00053675 | 0.0034232 |
| JHY | 0.81161377 | 2.57626726 | 4.61542512 | 0.00053695 | 0.0034232 |
| ADAMTS10 | 0.83913747 | 2.77966587 | 4.61332324 | 0.00053895 | 0.00343301 |
| SMAGP | -0.6043475 | 3.48559752 | -4.6121474 | 0.00054008 | 0.00343727 |
| SETD9 | 0.76516696 | 2.17229707 | 4.61045526 | 0.0005417 | 0.0034418 |
| LINC02197 | 1.14662661 | 0.51775426 | 4.60935813 | 0.00054275 | 0.00344561 |
| CXorf40B | -0.5314941 | 4.02371696 | -4.6088927 | 0.0005432 | 0.003447 |
| AC069224.1 | 1.3491972 | -0.0201623 | 4.60846271 | 0.00054361 | 0.00344818 |
| SLC39A13 | -0.5002752 | 4.44060018 | -4.6051378 | 0.00054682 | 0.00346565 |
| PLOD2 | -0.6428859 | 5.91014645 | -4.6037483 | 0.00054817 | 0.00347274 |
| MAPK15 | -0.9967966 | 1.63406093 | -4.5978093 | 0.00055397 | 0.00350509 |
| TMEM121 | -0.5419087 | 4.02923748 | -4.5956992 | 0.00055605 | 0.00351529 |
| FAM89A | -0.6360624 | 3.19386564 | -4.5929867 | 0.00055873 | 0.00352488 |
| AL358472.2 | 0.89689268 | 1.92087496 | 4.58940974 | 0.00056228 | 0.00354141 |
| LOXL3 | 0.51066459 | 4.11223895 | 4.58678327 | 0.00056491 | 0.00355454 |
| LINC00508 | -1.3023511 | 0.36574384 | -4.5866068 | 0.00056509 | 0.00355454 |
| ZNF285 | 0.58775485 | 3.44145853 | 4.57707643 | 0.00057473 | 0.00360777 |
| TESC | -0.557066 | 3.72937255 | -4.5766446 | 0.00057517 | 0.00360904 |
| AL135925.1 | 0.80286924 | 2.16174969 | 4.57531744 | 0.00057652 | 0.00361456 |
| ARMCX5 | -0.5303383 | 3.88753016 | -4.5740151 | 0.00057786 | 0.00362142 |
| ATP2B4 | -1.2105966 | 0.60353642 | -4.5733482 | 0.00057854 | 0.00362422 |
| COL2A1 | 0.95861702 | 2.14735413 | 4.56886626 | 0.00058317 | 0.00364714 |
| ZNF34 | 0.58241881 | 4.36603423 | 4.56808657 | 0.00058398 | 0.00365069 |
| GVQW2 | 0.9561082 | 1.50309711 | 4.56537591 | 0.00058679 | 0.00365925 |
| A4GALT | -0.6645057 | 2.99015103 | -4.5621001 | 0.00059022 | 0.00367758 |
| NUDT18 | -0.764962 | 2.66229341 | -4.5577551 | 0.0005948 | 0.00369975 |

| | | | | | |
|---|---|---|---|---|---|
| ELF4 | -0.6001229 | 4.3578199 | -4.5575626 | 0.000595 | 0.00369975 |
| MARK4 | -0.5266998 | 4.10579212 | -4.5569006 | 0.0005957 | 0.00370259 |
| GLMP | -0.5834143 | 3.50874205 | -4.5545221 | 0.00059823 | 0.00371371 |
| TFCP2L1 | -1.0176044 | 1.10445833 | -4.5542372 | 0.00059853 | 0.00371407 |
| FBXO36 | 0.72936981 | 3.24219281 | 4.55145643 | 0.0006015 | 0.00372942 |
| Z95115.1 | 0.78927747 | 2.11791156 | 4.55006653 | 0.00060298 | 0.00373249 |
| TMEM169 | -0.8579168 | 2.38338323 | -4.5494535 | 0.00060364 | 0.00373279 |
| SLC46A1 | -0.5060638 | 4.09569137 | -4.5465515 | 0.00060677 | 0.00374829 |
| ENPP2 | 1.42762172 | 0.10161852 | 4.5458837 | 0.00060749 | 0.00375122 |
| SYP | -0.5189151 | 4.27946269 | -4.5454206 | 0.00060799 | 0.00375278 |
| SPIN4 | -0.6363285 | 3.02381824 | -4.5450812 | 0.00060836 | 0.00375351 |
| SCN3A | -0.5697151 | 4.01760617 | -4.5443703 | 0.00060913 | 0.0037552 |
| MVB12B | -0.5831737 | 3.6766251 | -4.5379303 | 0.00061615 | 0.00379078 |
| TLX2 | -1.1346878 | 0.87602484 | -4.5372579 | 0.00061689 | 0.00379374 |
| P2RY11 | -0.7298612 | 2.96597177 | -4.5370351 | 0.00061713 | 0.00379374 |
| RERG | -0.8705948 | 1.81171968 | -4.5337295 | 0.00062077 | 0.00381149 |
| AL137003.1 | 0.6629682 | 2.88865743 | 4.53332337 | 0.00062122 | 0.00381271 |
| SP7 | 0.97963965 | 1.40609708 | 4.5329132 | 0.00062168 | 0.0038139 |
| DDIT4 | -1.2993039 | 4.91730096 | -4.5326927 | 0.00062192 | 0.0038139 |
| LINC01793 | -0.7857084 | 2.02055372 | -4.5311123 | 0.00062367 | 0.0038231 |
| SLC12A9 | -0.5072074 | 4.97631178 | -4.5287947 | 0.00062625 | 0.00383249 |
| NRN1 | -0.630797 | 5.59389071 | -4.5250818 | 0.00063041 | 0.00385347 |
| DNAJB4 | -0.5902926 | 3.24328148 | -4.5225095 | 0.00063331 | 0.00386338 |
| LSM12 | -0.6378777 | 2.89629004 | -4.5183704 | 0.000638 | 0.00388572 |
| BAMBI | 0.53514922 | 4.04009229 | 4.51724283 | 0.00063928 | 0.00389198 |
| ZNF492 | -0.5927997 | 3.26394291 | -4.5150921 | 0.00064174 | 0.00390379 |
| SURF2 | -0.5264889 | 5.00565162 | -4.5125056 | 0.0006447 | 0.00391869 |
| EID2B | 0.71907394 | 2.71974277 | 4.50505161 | 0.00065333 | 0.00396001 |
| ZSCAN5A | 0.56525961 | 3.77759852 | 4.50461206 | 0.00065385 | 0.00396154 |
| AL513318.2 | 0.89005079 | 2.85490707 | 4.50438292 | 0.00065411 | 0.00396157 |
| FZD2 | -0.5883665 | 3.66637416 | -4.5019471 | 0.00065696 | 0.00397315 |
| C18orf21 | 0.53074879 | 4.25332414 | 4.5016453 | 0.00065732 | 0.00397315 |
| RCN3 | -0.9224925 | 1.59993527 | -4.5016284 | 0.00065734 | 0.00397315 |
| LGI4 | -0.9828683 | 1.66753137 | -4.5012211 | 0.00065781 | 0.00397445 |
| SPATA13 | -0.5103748 | 3.99383325 | -4.4991092 | 0.0006603 | 0.00398787 |
| LZTS1 | -0.8768127 | 1.98512221 | -4.4983499 | 0.00066119 | 0.00399168 |
| ANKRD34B | -0.9277719 | 1.67980187 | -4.4958064 | 0.0006642 | 0.00400666 |
| PIM1 | 0.55179636 | 4.32345292 | 4.49006812 | 0.00067104 | 0.00403186 |
| AP006284.1 | -1.39943 | 0.6587582 | -4.4877676 | 0.00067381 | 0.00404525 |
| AF111167.2 | -0.6756541 | 2.90450703 | -4.4853034 | 0.00067678 | 0.0040583 |
| DNAJC6 | -0.6587914 | 3.051168 | -4.4852836 | 0.00067681 | 0.0040583 |
| DISP1 | 0.61660368 | 3.1981767 | 4.48507844 | 0.00067705 | 0.0040583 |
| ZNF418 | 0.64987316 | 3.00859364 | 4.48467816 | 0.00067754 | 0.00405959 |
| CYP4V2 | -0.67014 | 2.67609685 | -4.4841872 | 0.00067813 | 0.00406106 |
| ZNF821 | 0.54308224 | 3.88896506 | 4.48403295 | 0.00067832 | 0.00406106 |

| | | | | | |
|---|---|---|---|---|---|
| NEXMIF | -1.0318114 | 1.2948166 | -4.4808645 | 0.00068217 | 0.00407928 |
| AL133215.2 | 0.69538802 | 2.59600513 | 4.4773407 | 0.00068648 | 0.0041002 |
| MYH3 | -0.7827004 | 2.01828113 | -4.4769264 | 0.00068699 | 0.00410162 |
| TCF7L2 | 0.65724912 | 2.88197001 | 4.47069984 | 0.00069468 | 0.00413692 |
| SLC25A21 | 0.61087766 | 3.29359355 | 4.46925321 | 0.00069648 | 0.00414521 |
| AC092329.1 | 0.799152 | 2.20193857 | 4.46460407 | 0.0007023 | 0.00417491 |
| AC113346.1 | 1.27220374 | 0.24614439 | 4.45412154 | 0.0007156 | 0.00424565 |
| COQ10A | -0.5453567 | 4.63602998 | -4.4503109 | 0.0007205 | 0.00426732 |
| WDPCP | 0.50328804 | 4.24494585 | 4.44550375 | 0.00072673 | 0.00429583 |
| SLC32A1 | 1.25411845 | 0.28369207 | 4.44536127 | 0.00072692 | 0.00429583 |
| RELL2 | -0.6581636 | 3.09606358 | -4.4451592 | 0.00072718 | 0.00429583 |
| AL513550.1 | 0.52365708 | 4.05140126 | 4.43780439 | 0.00073683 | 0.00434266 |
| GPM6B | 0.58874626 | 3.39893495 | 4.43704059 | 0.00073784 | 0.00434692 |
| KLHL3 | 1.12846107 | 0.83571905 | 4.43652667 | 0.00073852 | 0.00434754 |
| AC026801.2 | 1.37700808 | -0.3520228 | 4.42845246 | 0.00074929 | 0.00440079 |
| AL138762.1 | 0.91829713 | 1.36584573 | 4.42653746 | 0.00075186 | 0.00441069 |
| LRRC6 | 1.06750721 | 0.81397242 | 4.42614252 | 0.0007524 | 0.0044121 |
| TAGLN | -0.8321952 | 2.54884728 | -4.4247097 | 0.00075433 | 0.00442175 |
| SYT16 | 1.48554619 | -0.4264345 | 4.42056158 | 0.00075997 | 0.00444446 |
| ERVMER61-1 | 0.53296847 | 4.05200525 | 4.41911265 | 0.00076195 | 0.00445259 |
| MAFB | 1.40145967 | -0.4869153 | 4.41252628 | 0.00077101 | 0.00449687 |
| COMMD3 | 0.56562517 | 3.56138273 | 4.41223715 | 0.00077141 | 0.00449741 |
| SLC35D2 | 0.56858487 | 3.44799482 | 4.41194629 | 0.00077181 | 0.00449741 |
| AC012377.1 | 1.19020222 | 0.11787276 | 4.409581 | 0.0007751 | 0.00451203 |
| AL136295.6 | 1.19234773 | 0.17108844 | 4.40912486 | 0.00077573 | 0.00451399 |
| AC003986.2 | 0.86887523 | 1.68701749 | 4.4078936 | 0.00077745 | 0.00451704 |
| DET1 | 0.60033375 | 3.41920325 | 4.40406121 | 0.00078282 | 0.00454025 |
| MANSC1 | -1.3715755 | -0.2796038 | -4.4039736 | 0.00078295 | 0.00454025 |
| SFRP5 | -0.8332056 | 2.07128901 | -4.397092 | 0.00079269 | 0.00458795 |
| HBEGF | -0.6078997 | 3.15959308 | -4.3954775 | 0.00079499 | 0.00459508 |
| OSBP2 | 0.50151151 | 4.14303569 | 4.39537809 | 0.00079513 | 0.00459508 |
| CYP2J2 | -0.712333 | 2.45071287 | -4.3918602 | 0.00080018 | 0.00461719 |
| AC068338.2 | 0.88879056 | 1.59889694 | 4.38873451 | 0.00080469 | 0.00463967 |
| ZSCAN16-AS1 | 0.75763759 | 2.32809196 | 4.38820964 | 0.00080545 | 0.00464229 |
| NME9 | 1.26664724 | 0.13289734 | 4.38798164 | 0.00080578 | 0.00464242 |
| CAMK2N1 | 0.67709296 | 2.77079456 | 4.38726351 | 0.00080682 | 0.00464312 |
| IDNK | -1.0507349 | 0.84168356 | -4.3863313 | 0.00080817 | 0.00464738 |
| MIR29B2CHG | -0.9110052 | 1.35055066 | -4.3771357 | 0.00082166 | 0.0047106 |
| EPOR | -0.5775645 | 3.67594315 | -4.3750584 | 0.00082473 | 0.00472645 |
| MPP7 | -0.7777408 | 1.93750591 | -4.3738334 | 0.00082655 | 0.00473174 |
| GDF9 | 1.11178529 | 0.81060698 | 4.37380671 | 0.00082659 | 0.00473174 |
| KCNN4 | 0.75032071 | 2.24321281 | 4.37325459 | 0.00082742 | 0.00473287 |
| AL365205.1 | -0.9091882 | 1.74188417 | -4.3669225 | 0.0008369 | 0.00477451 |
| COL1A1 | -0.9060303 | 2.62112385 | -4.3654067 | 0.00083919 | 0.00478576 |
| MBNL1-AS1 | 1.38323128 | -0.6350372 | 4.36504145 | 0.00083974 | 0.00478711 |

| | | | | | |
|---|---|---|---|---|---|
| RWDD4 | 0.56699884 | 3.2873542 | 4.36196315 | 0.00084441 | 0.00480856 |
| AJM1 | -0.8302 | 2.67685791 | -4.3619322 | 0.00084446 | 0.00480856 |
| ANKRD1 | -1.9374211 | -0.1485011 | -4.3560302 | 0.00085349 | 0.00484721 |
| AC034231.1 | 1.15379644 | 0.46550474 | 4.35380253 | 0.00085692 | 0.00485291 |
| ADAMTS2 | -0.5358288 | 4.74804588 | -4.3536073 | 0.00085722 | 0.00485291 |
| ENHO | -0.7910082 | 2.1493733 | -4.3535091 | 0.00085738 | 0.00485291 |
| PPM1K | 0.62562157 | 3.02626014 | 4.34646733 | 0.00086833 | 0.00490391 |
| WAC-AS1 | -0.6559263 | 2.63459145 | -4.344867 | 0.00087084 | 0.00491625 |
| PLAC8 | -0.9356041 | 1.27608043 | -4.3436088 | 0.00087282 | 0.00492558 |
| VPS9D1-AS1 | -0.7804909 | 1.98999717 | -4.3418916 | 0.00087553 | 0.00493902 |
| AC106820.2 | -0.8913416 | 1.51724459 | -4.341574 | 0.00087603 | 0.00494001 |
| KIF9 | 0.63473197 | 2.8394876 | 4.33968587 | 0.00087902 | 0.00495318 |
| AATK | -0.5188263 | 4.48144716 | -4.3373924 | 0.00088266 | 0.00497002 |
| AL138976.2 | 1.23872983 | -0.3656491 | 4.32451222 | 0.00090342 | 0.00505871 |
| MYPN | 1.30458947 | 0.11477097 | 4.32308495 | 0.00090576 | 0.00506802 |
| TYW1B | 0.70038994 | 2.54587987 | 4.31726344 | 0.00091533 | 0.00511127 |
| PLEKHH2 | -0.7148967 | 2.57882291 | -4.3171536 | 0.00091551 | 0.00511127 |
| MCEE | 0.63563295 | 3.28356058 | 4.31599668 | 0.00091743 | 0.00512006 |
| ADPRM | 0.79788102 | 2.3451947 | 4.3145076 | 0.0009199 | 0.0051301 |
| BHLHA15 | -0.884073 | 1.60689723 | -4.3128352 | 0.00092268 | 0.00514184 |
| SP9 | 0.54396662 | 3.89397509 | 4.31231647 | 0.00092355 | 0.00514476 |
| EME2 | -0.5418268 | 5.50040451 | -4.3065363 | 0.00093325 | 0.00518735 |
| PHLDA2 | -0.757359 | 2.52131419 | -4.3062458 | 0.00093374 | 0.00518817 |
| SYN2 | -0.6350833 | 3.05891292 | -4.3040901 | 0.00093739 | 0.0052027 |
| CRIP3 | -1.7353018 | -0.4946508 | -4.3038787 | 0.00093775 | 0.00520279 |
| AC008966.1 | 0.76701416 | 2.25279401 | 4.3022152 | 0.00094057 | 0.0052153 |
| ADCY7 | -0.5419833 | 3.62894807 | -4.302145 | 0.00094069 | 0.0052153 |
| ITGA2 | -0.5946818 | 3.06628115 | -4.3009533 | 0.00094272 | 0.00522115 |
| AC092958.1 | 0.71482708 | 2.60895778 | 4.30071647 | 0.00094312 | 0.00522115 |
| SEC14L2 | -0.607315 | 3.04799847 | -4.2982067 | 0.00094741 | 0.00524298 |
| ZNF611 | -0.512289 | 3.7207806 | -4.29399 | 0.00095467 | 0.00527927 |
| SLCO3A1 | -0.8067277 | 1.94885041 | -4.2925922 | 0.00095709 | 0.00528441 |
| MPV17L | -1.2644557 | -0.1709366 | -4.2924449 | 0.00095734 | 0.00528441 |
| SPATA17 | 1.21666532 | 0.07376881 | 4.29062358 | 0.0009605 | 0.00529992 |
| LATS2 | 0.59111721 | 5.80260621 | 4.28725518 | 0.00096638 | 0.00532651 |
| RTN4RL2 | -1.1218815 | 0.64093109 | -4.2848514 | 0.00097059 | 0.00534002 |
| SAXO2 | 0.97659483 | 0.94688543 | 4.28257866 | 0.00097459 | 0.0053562 |
| JPH1 | -1.1283355 | 0.34075997 | -4.2820553 | 0.00097552 | 0.00535739 |
| SEMA7A | -0.7206775 | 2.73551703 | -4.2801655 | 0.00097886 | 0.0053738 |
| HLA-F | -0.5519105 | 3.66292143 | -4.273227 | 0.00099124 | 0.00543389 |
| COL7A1 | -0.857288 | 4.13273822 | -4.2646835 | 0.00100671 | 0.00549678 |
| NXPH3 | -1.1223811 | 0.65694658 | -4.2642421 | 0.00100751 | 0.00549719 |
| ST6GALNAC6 | -1.126848 | 0.54814965 | -4.2640461 | 0.00100787 | 0.00549719 |
| ZSWIM3 | 0.59745783 | 3.22956871 | 4.26125272 | 0.00101299 | 0.00552112 |
| AL136040.1 | 0.90125639 | 1.34180552 | 4.25869737 | 0.00101769 | 0.00554477 |

| | | | | | |
|---|---|---|---|---|---|
| FHDC1 | -1.2399227 | -0.1167674 | -4.2571134 | 0.00102062 | 0.00555672 |
| NR2F2 | -0.5162752 | 3.74782046 | -4.2566749 | 0.00102143 | 0.00555739 |
| EXOC3-AS1 | 0.56497383 | 3.29824086 | 4.25429618 | 0.00102585 | 0.00557516 |
| NMNAT3 | -0.7699201 | 2.17091455 | -4.2540004 | 0.0010264 | 0.00557615 |
| BAALC-AS1 | 0.72264349 | 2.26501511 | 4.25362294 | 0.0010271 | 0.0055763 |
| TOR4A | -0.97743 | 1.28966259 | -4.2533251 | 0.00102766 | 0.00557698 |
| AL031058.1 | 0.99681448 | 0.97123724 | 4.24980678 | 0.00103424 | 0.00560666 |
| AC009506.1 | 0.70446465 | 2.43704616 | 4.24850265 | 0.00103669 | 0.00561392 |
| KANK3 | -1.1070022 | 1.159397 | -4.245448 | 0.00104245 | 0.00563303 |
| ZNF548 | 0.54866366 | 3.33639568 | 4.23828278 | 0.0010561 | 0.00568446 |
| TTPA | 0.54757953 | 3.42234894 | 4.23629396 | 0.00105992 | 0.00570098 |
| CPNE7 | -0.5198799 | 5.87359954 | -4.2324036 | 0.00106744 | 0.0057329 |
| HSD11B2 | -1.0094573 | 1.01909646 | -4.2250037 | 0.00108188 | 0.00579646 |
| GBX2 | -0.5246349 | 4.12359954 | -4.2225861 | 0.00108665 | 0.00581375 |
| SLC26A10 | -1.1468334 | 0.83877718 | -4.221392 | 0.00108901 | 0.00582021 |
| CD83 | 0.64378414 | 3.22680687 | 4.21896373 | 0.00109382 | 0.00584183 |
| RALGPS1 | -0.6935296 | 3.0120709 | -4.2167681 | 0.0010982 | 0.00586106 |
| NAGS | -0.649349 | 2.7822166 | -4.2158292 | 0.00110008 | 0.00586901 |
| SPATA7 | 0.50910153 | 3.7456858 | 4.21356988 | 0.0011046 | 0.0058828 |
| SLC27A3 | -0.5993561 | 2.97907555 | -4.2096961 | 0.00111241 | 0.00591607 |
| KCNAB2 | -0.7676698 | 2.75316393 | -4.2055425 | 0.00112085 | 0.00595049 |
| MAP3K21 | -0.9179494 | 1.22878376 | -4.1963048 | 0.00113985 | 0.00603022 |
| EREG | 0.8802145 | 1.45890085 | 4.19318637 | 0.00114634 | 0.00606032 |
| PLEKHA7 | 0.52615922 | 3.5648106 | 4.19270252 | 0.00114735 | 0.00606202 |
| RTBDN | -0.8716813 | 1.53379463 | -4.1900075 | 0.001153 | 0.00608488 |
| KCNJ15 | 0.62556024 | 2.93693683 | 4.18948179 | 0.0011541 | 0.00608806 |
| AL355001.2 | 0.72245882 | 2.30434253 | 4.18933785 | 0.0011544 | 0.00608806 |
| SYBU | -1.1766213 | 0.27027243 | -4.1872281 | 0.00115885 | 0.0061046 |
| FAM227A | 0.59016152 | 3.47077446 | 4.18389166 | 0.00116591 | 0.00613508 |
| ZNF688 | 0.53446377 | 3.71908544 | 4.18377664 | 0.00116616 | 0.00613508 |
| SMTNL2 | -0.8849447 | 1.51850624 | -4.1825681 | 0.00116873 | 0.00614646 |
| ZC4H2 | -0.5263633 | 3.47416679 | -4.1787188 | 0.00117695 | 0.00617685 |
| CDK20 | 0.51250771 | 3.77158028 | 4.17551913 | 0.00118383 | 0.00620652 |
| SLC7A10 | -1.214125 | 0.2798119 | -4.1746465 | 0.00118572 | 0.00620856 |
| C17orf97 | 1.00093251 | 0.80721439 | 4.17457907 | 0.00118586 | 0.00620856 |
| ADPRH | -0.9733044 | 0.8668205 | -4.1695065 | 0.00119688 | 0.00625975 |
| AC021054.1 | -0.6048391 | 2.96809312 | -4.167851 | 0.0012005 | 0.00627217 |
| ECHDC3 | -0.8404425 | 1.76028613 | -4.1674468 | 0.00120138 | 0.00627463 |
| B9D2 | 0.61812328 | 2.99972134 | 4.1667372 | 0.00120294 | 0.00627842 |
| DEXI | 0.51143137 | 3.84328899 | 4.16575811 | 0.00120509 | 0.00628531 |
| LRRC37B | 0.51822694 | 3.7659608 | 4.16536778 | 0.00120595 | 0.00628546 |
| CPA4 | -0.9118064 | 1.35882177 | -4.1650413 | 0.00120667 | 0.00628704 |
| GPER1 | -0.5142565 | 3.90087914 | -4.1614017 | 0.0012147 | 0.00632057 |
| ATP6AP1L | 0.96074587 | 0.95875068 | 4.16121913 | 0.00121511 | 0.00632057 |
| JAKMIP1 | -0.6270818 | 3.24774869 | -4.1581388 | 0.00122195 | 0.00635358 |

| | | | | | |
|---|---|---|---|---|---|
| IL1R1 | -1.2006054 | 0.90245945 | -4.1576641 | 0.00122301 | 0.0063569 |
| TTC32 | 0.69855184 | 2.73403775 | 4.15319301 | 0.00123303 | 0.00639799 |
| LINGO1 | -0.9150415 | 1.33183831 | -4.1526964 | 0.00123415 | 0.00639941 |
| GSDME | -0.5026832 | 3.81329182 | -4.1519216 | 0.0012359 | 0.00640188 |
| URB1-AS1 | 0.58742543 | 3.05365108 | 4.1485732 | 0.00124347 | 0.00642004 |
| RRAGB | 0.51919611 | 3.49644993 | 4.14850258 | 0.00124363 | 0.00642004 |
| CACNA1A | 1.36831159 | -0.4293756 | 4.14757436 | 0.00124574 | 0.00642869 |
| C8orf58 | -0.5002259 | 3.93241137 | -4.1429479 | 0.00125631 | 0.00647 |
| SLC1A1 | -0.9253433 | 2.13344953 | -4.1423066 | 0.00125778 | 0.00647196 |
| PSMA8 | 0.66027076 | 2.31999529 | 4.13954547 | 0.00126414 | 0.00649486 |
| RASIP1 | -0.7212134 | 2.26384822 | -4.1373837 | 0.00126914 | 0.00651172 |
| LMTK3 | -1.1057738 | 0.88099587 | -4.1352451 | 0.00127411 | 0.00652682 |
| LNP1 | 0.54158667 | 3.26712781 | 4.13491304 | 0.00127488 | 0.00652682 |
| LINC01003 | 0.74520088 | 2.18822718 | 4.13481844 | 0.0012751 | 0.00652682 |
| PRKCD | -0.5236851 | 3.40753117 | -4.123617 | 0.00130148 | 0.00662975 |
| AP001972.1 | -0.7119167 | 2.35248178 | -4.1234891 | 0.00130178 | 0.00662975 |
| SMIM37 | 0.5054038 | 3.76641393 | 4.12175379 | 0.00130592 | 0.00663563 |
| AC068888.1 | -0.8218668 | 1.99198208 | -4.1215344 | 0.00130644 | 0.00663563 |
| AC092329.3 | 0.75063182 | 1.98934775 | 4.11760944 | 0.00131585 | 0.00666866 |
| SLC51A | -0.9422702 | 1.00867098 | -4.1097108 | 0.001335 | 0.00674766 |
| SARM1 | -0.5338824 | 3.70296164 | -4.1096358 | 0.00133518 | 0.00674766 |
| CHIC2 | 0.54817716 | 3.30254183 | 4.10748082 | 0.00134046 | 0.00676303 |
| SLC22A23 | -0.5301776 | 3.53804883 | -4.1072339 | 0.00134106 | 0.00676383 |
| CORO7 | -0.6314352 | 3.04895266 | -4.1059935 | 0.00134411 | 0.00677137 |
| ELAC1 | 0.64384154 | 2.64981188 | 4.10589776 | 0.00134435 | 0.00677137 |
| AC017100.1 | 0.86217539 | 1.2922159 | 4.10331812 | 0.00135071 | 0.00679437 |
| MST1 | -0.5159616 | 3.80336113 | -4.1003559 | 0.00135805 | 0.00682225 |
| HOOK1 | 0.64350502 | 2.86014603 | 4.08913662 | 0.00138624 | 0.00694478 |
| PIH1D2 | 0.90784831 | 1.1102429 | 4.08842602 | 0.00138804 | 0.00694757 |
| UBAC2-AS1 | 0.7274962 | 2.04630638 | 4.08651504 | 0.00139291 | 0.00696963 |
| SPATC1 | -1.3211134 | -0.4591195 | -4.0846624 | 0.00139765 | 0.0069887 |
| ANKRD31 | 0.88095245 | 1.20531006 | 4.07961498 | 0.00141064 | 0.00704666 |
| GPLD1 | -0.5933973 | 3.14498957 | -4.0769529 | 0.00141753 | 0.00707412 |
| FBLN2 | -0.5426614 | 3.57937423 | -4.0741178 | 0.00142492 | 0.00710162 |
| SHROOM3 | -0.7042581 | 2.16698519 | -4.0698558 | 0.0014361 | 0.00714322 |
| AC025181.2 | 0.80757766 | 1.47925701 | 4.06476297 | 0.00144958 | 0.00719843 |
| ARHGAP8 | -0.857114 | 1.39741544 | -4.060056 | 0.00146215 | 0.00723949 |
| NPAS3 | 0.68839021 | 2.43596737 | 4.05816423 | 0.00146723 | 0.00725517 |
| AC068870.2 | 0.82065802 | 1.5075451 | 4.05478307 | 0.00147636 | 0.00728367 |
| PALLD | -0.8539762 | 1.22363417 | -4.050606 | 0.00148772 | 0.00732303 |
| ZNF419 | 0.68729175 | 2.61303452 | 4.0498973 | 0.00148966 | 0.00732779 |
| ACOT1 | 0.53371241 | 3.39449247 | 4.04855071 | 0.00149335 | 0.00734116 |
| NFASC | -0.5456044 | 3.36723457 | -4.048338 | 0.00149393 | 0.00734142 |
| AANAT | -1.155337 | 0.12613829 | -4.046549 | 0.00149885 | 0.00735625 |
| KDM6B | 0.54075514 | 3.48265051 | 4.03908786 | 0.00151952 | 0.00744326 |

| | | | | | |
|---|---|---|---|---|---|
| NTNG2 | -0.6794063 | 2.5378685 | -4.037509 | 0.00152394 | 0.00746246 |
| GALNT12 | -0.6356183 | 3.0518223 | -4.0346201 | 0.00153205 | 0.00748803 |
| TFPI2 | -0.815144 | 2.78924123 | -4.0330709 | 0.00153641 | 0.00750172 |
| TRNP1 | -0.7133242 | 2.16699444 | -4.0303053 | 0.00154424 | 0.00753508 |
| B4GALT1 | 0.54791632 | 3.74883601 | 4.02747002 | 0.00155231 | 0.00756957 |
| ALDH2 | -0.5985628 | 3.48957277 | -4.025598 | 0.00155766 | 0.00758956 |
| PIK3R5 | -1.0534239 | 0.6187158 | -4.0255099 | 0.00155791 | 0.00758956 |
| AC015712.4 | -1.2993557 | -0.3915963 | -4.0241704 | 0.00156175 | 0.00760582 |
| LINC01122 | 0.52936093 | 3.28700637 | 4.02247804 | 0.00156662 | 0.00762462 |
| MAP3K13 | 0.65700417 | 2.33816054 | 4.02141907 | 0.00156967 | 0.00763702 |
| SSH2 | -0.5541759 | 3.11209225 | -4.0209329 | 0.00157108 | 0.00763901 |
| ZNF571 | 0.59095694 | 2.82722323 | 4.02016215 | 0.0015733 | 0.00764486 |
| CAPS2 | 0.87381622 | 1.7746624 | 4.01912238 | 0.00157631 | 0.00765458 |
| PLLP | -0.6329863 | 2.91388019 | -4.0137803 | 0.00159188 | 0.00772025 |
| AC016717.2 | 0.79409125 | 1.35345442 | 4.0132261 | 0.0015935 | 0.00772565 |
| NRARP | -0.5368014 | 3.3802228 | -4.0127902 | 0.00159478 | 0.00772937 |
| ANKDD1A | -0.5913125 | 3.22723114 | -4.0124434 | 0.0015958 | 0.00773183 |
| CMTM1 | 0.61792286 | 2.65593777 | 4.0119567 | 0.00159723 | 0.00773627 |
| TNNI3 | -1.2684975 | -0.1859994 | -4.0109802 | 0.0016001 | 0.00774771 |
| TCTE3 | 0.68238646 | 2.19013855 | 4.00586652 | 0.00161522 | 0.00779104 |
| CXCL16 | -0.5499688 | 3.22267454 | -3.9990216 | 0.0016357 | 0.00786841 |
| AL359258.2 | 0.72039378 | 1.84562665 | 3.99894174 | 0.00163594 | 0.00786841 |
| JMJD1C-AS1 | 0.72873592 | 1.99846413 | 3.99686657 | 0.0016422 | 0.00789101 |
| NT5M | -0.5453143 | 3.62873562 | -3.9908444 | 0.00166051 | 0.00796383 |
| ARMC2 | 0.68246524 | 2.16014227 | 3.98889033 | 0.0016665 | 0.00798748 |
| BEX4 | 0.51975047 | 3.47596879 | 3.98703113 | 0.00167221 | 0.00801234 |
| FOXD2-AS1 | 0.59711477 | 2.75957991 | 3.98562226 | 0.00167656 | 0.00802301 |
| PRSS53 | -1.2349204 | 1.55494764 | -3.9850129 | 0.00167844 | 0.00802948 |
| MAP6 | -1.2291777 | 0.07513059 | -3.9840116 | 0.00168154 | 0.00804082 |
| TAPT1-AS1 | 0.8256727 | 1.51526177 | 3.98169362 | 0.00168874 | 0.00806261 |
| LINC01521 | 0.5004027 | 3.87706027 | 3.97701014 | 0.00170337 | 0.00812247 |
| GPRASP2 | 0.53763196 | 4.13112783 | 3.97688068 | 0.00170378 | 0.00812247 |
| PRL | -0.7299066 | 1.79867027 | -3.9766555 | 0.00170449 | 0.00812328 |
| SEMA4D | 0.51812122 | 3.51979726 | 3.97098973 | 0.00172238 | 0.00818794 |
| RBMS2 | -0.5057525 | 3.78925764 | -3.9701515 | 0.00172505 | 0.00819545 |
| NKILA | 0.86411327 | 1.37610835 | 3.96788716 | 0.00173226 | 0.00822716 |
| NPR2 | -0.7340081 | 2.1308833 | -3.9673571 | 0.00173396 | 0.00823004 |
| AC012615.1 | 0.81660001 | 1.42861271 | 3.96573701 | 0.00173914 | 0.00824949 |
| NHLRC1 | 0.52233759 | 3.40625341 | 3.96527516 | 0.00174063 | 0.00825135 |
| AC002467.1 | 0.76069547 | 1.73235601 | 3.96452432 | 0.00174304 | 0.00826019 |
| ROPN1L | 1.10402952 | 0.26793593 | 3.96327476 | 0.00174706 | 0.00827148 |
| TENT5B | -1.1468674 | 0.37379286 | -3.9550065 | 0.00177391 | 0.00838042 |
| PPM1J | -0.7384787 | 2.18111892 | -3.9549961 | 0.00177394 | 0.00838042 |
| KIAA1549L | -0.8700603 | 1.37704505 | -3.9521638 | 0.00178324 | 0.00840859 |
| BNIP3 | -0.7694295 | 6.11098714 | -3.9513757 | 0.00178583 | 0.00841821 |

| | | | | | |
|---|---|---|---|---|---|
| IPPK | 0.52825599 | 3.91307475 | 3.94951817 | 0.00179196 | 0.00844185 |
| KDM4D | 0.63174004 | 2.6423207 | 3.94791432 | 0.00179728 | 0.00845636 |
| COL9A3 | -1.3073665 | -0.4907477 | -3.9460288 | 0.00180354 | 0.0084832 |
| FGF14-AS2 | 1.06957159 | 0.34874462 | 3.94484144 | 0.0018075 | 0.00849917 |
| AP002360.3 | 1.25710676 | -0.9639464 | 3.94290834 | 0.00181396 | 0.00851897 |
| CUX2 | 1.18232916 | 0.15136185 | 3.94149266 | 0.00181871 | 0.00853333 |
| ZNF782 | 0.51596424 | 3.44896979 | 3.93758426 | 0.00183188 | 0.00857739 |
| MIR9-3HG | -0.5120711 | 3.45776927 | -3.9370554 | 0.00183367 | 0.00857739 |
| PRDM16 | -0.5302813 | 3.23196091 | -3.932205 | 0.00185017 | 0.00863546 |
| GEMIN8 | 0.60464142 | 2.74181271 | 3.92625781 | 0.00187061 | 0.00870402 |
| NBPF6 | 0.70893611 | 2.11306708 | 3.92077689 | 0.00188965 | 0.00878453 |
| CAHM | 1.13719691 | -0.073119 | 3.91714941 | 0.00190236 | 0.00882736 |
| ITGB3 | -1.1738829 | -0.0047416 | -3.9156504 | 0.00190764 | 0.00884915 |
| RAP1GAP2 | -0.5547816 | 3.21375488 | -3.9148848 | 0.00191034 | 0.00885897 |
| LINC01579 | 0.50453138 | 3.43010326 | 3.914367 | 0.00191217 | 0.00886095 |
| PTPRO | 1.00855661 | 0.65040348 | 3.90709682 | 0.00193805 | 0.00896002 |
| ZNF213 | 0.51196095 | 3.53119868 | 3.90658412 | 0.00193989 | 0.00896578 |
| ASAH2 | -0.7544915 | 1.66996193 | -3.9045039 | 0.00194737 | 0.00899211 |
| NIFK-AS1 | 0.59168684 | 2.80711664 | 3.90405023 | 0.001949 | 0.00899692 |
| FILIP1L | 0.75500909 | 1.59270143 | 3.90264655 | 0.00195407 | 0.00901229 |
| MALINC1 | 0.81029533 | 1.38821109 | 3.90229556 | 0.00195534 | 0.00901497 |
| KLF2 | -0.7251137 | 1.94758593 | -3.8998627 | 0.00196416 | 0.00905033 |
| PIK3CD-AS2 | -0.5529248 | 3.13333628 | -3.8970335 | 0.00197447 | 0.00908797 |
| ANKRD29 | -0.9487725 | 0.95813014 | -3.8915282 | 0.00199469 | 0.0091631 |
| LINC02506 | 0.61305146 | 2.56521419 | 3.8907317 | 0.00199763 | 0.00917384 |
| LINC01977 | 0.87520315 | 1.02289141 | 3.88372961 | 0.00202369 | 0.00927945 |
| COL23A1 | -0.9584912 | 1.7113422 | -3.8813563 | 0.00203261 | 0.00930904 |
| ZEB1-AS1 | 1.18400171 | -0.03519 | 3.87979318 | 0.0020385 | 0.00933038 |
| COL4A6 | -0.8256876 | 1.65135078 | -3.8794513 | 0.00203979 | 0.00933347 |
| PLEKHF1 | -0.7031764 | 2.21297148 | -3.8739212 | 0.00206079 | 0.00940966 |
| PRRG4 | -0.9113115 | 0.87998573 | -3.8710516 | 0.00207177 | 0.00944557 |
| AFAP1L2 | -0.8417013 | 1.43441662 | -3.8675851 | 0.00208512 | 0.00949785 |
| FAM160A1 | -1.2680313 | -0.1370269 | -3.8644882 | 0.00209712 | 0.00954964 |
| LINC00886 | 1.24941288 | -0.167418 | 3.86368949 | 0.00210023 | 0.00955804 |
| ZNF486 | -0.5104677 | 3.18059241 | -3.8618719 | 0.00210732 | 0.00958384 |
| C3orf52 | -0.5321934 | 3.08897383 | -3.8617486 | 0.0021078 | 0.00958384 |
| PARVA | -0.8248999 | 1.30997878 | -3.8612964 | 0.00210957 | 0.00958612 |
| FHOD3 | -0.5023402 | 3.50807969 | -3.856157 | 0.00212976 | 0.00966338 |
| FAM162A | -0.643757 | 5.6023189 | -3.8555807 | 0.00213204 | 0.00967081 |
| ASB2 | -1.0775624 | 0.2991307 | -3.8548982 | 0.00213474 | 0.00968016 |
| AC007881.3 | 1.02231222 | 0.33070614 | 3.85338682 | 0.00214073 | 0.0096974 |
| RAB6D | -1.1691991 | -0.1352603 | -3.8529705 | 0.00214238 | 0.0096974 |
| CDS1 | -0.9590712 | 0.66262419 | -3.8517827 | 0.0021471 | 0.00971136 |
| CA13 | -0.5762039 | 2.72408673 | -3.8513526 | 0.00214882 | 0.00971201 |
| RASL11A | -0.7263867 | 1.87702155 | -3.8477919 | 0.00216305 | 0.00976179 |

| | | | | | |
|---|---|---|---|---|---|
| C21orf62-AS1 | 0.76658684 | 1.62648793 | 3.84706897 | 0.00216596 | 0.0097663 |
| RGS11 | -0.9550298 | 1.2203811 | -3.8410966 | 0.00219009 | 0.00986027 |
| IGLV5-52 | -0.7485912 | 1.73188138 | -3.83888 | 0.00219911 | 0.00989502 |
| MCF2L | -0.542334 | 3.13708026 | -3.8386039 | 0.00220024 | 0.00989715 |
| PLCH1 | -1.213191 | -0.4963947 | -3.8384228 | 0.00220098 | 0.00989754 |
| KANK4 | 0.77946725 | 1.77657102 | 3.83811551 | 0.00220223 | 0.00990024 |
| HIST1H3E | 0.57460151 | 2.87693998 | 3.83675955 | 0.00220778 | 0.00991734 |
| FAM167A | 0.50611282 | 3.51236065 | 3.83664651 | 0.00220824 | 0.00991734 |
| ZNF596 | 0.57302878 | 2.69762689 | 3.83499782 | 0.00221501 | 0.00993409 |
| GCNT1 | -0.8662286 | 1.12031027 | -3.8330242 | 0.00222314 | 0.0099558 |
| BCAN | 1.15594497 | 0.05543423 | 3.82943153 | 0.00223801 | 0.01001352 |
| TNFRSF25 | -0.8877779 | 1.79738501 | -3.8284112 | 0.00224226 | 0.01002362 |
| RAB26 | -0.5025145 | 3.97520391 | -3.8257826 | 0.00225322 | 0.01005484 |
| UBALD1 | 0.61439031 | 2.83400556 | 3.82364762 | 0.00226217 | 0.01008883 |
| ANO7 | -0.6910058 | 2.3580127 | -3.8229163 | 0.00226525 | 0.01009659 |
| CCDC38 | 0.85698951 | 0.82082999 | 3.82201525 | 0.00226904 | 0.01009838 |
| FOXO1 | 0.55202498 | 3.6960752 | 3.82151368 | 0.00227116 | 0.0101021 |
| CSF1 | -0.9811192 | 0.67481044 | -3.817799 | 0.00228688 | 0.0101601 |
| CYP1A1 | -0.5973765 | 2.72452111 | -3.8120194 | 0.00231156 | 0.01025471 |
| AC012306.2 | 0.78787093 | 1.42091556 | 3.8107407 | 0.00231706 | 0.01027335 |
| PAQR6 | -0.6637799 | 2.70822045 | -3.8097292 | 0.00232142 | 0.01028036 |
| RETREG1 | -1.3198626 | -0.6801899 | -3.8065062 | 0.00233536 | 0.01033305 |
| RIN2 | 1.16703119 | 0.00847783 | 3.80394112 | 0.00234652 | 0.01037332 |
| PORCN | -0.6501742 | 2.39818522 | -3.8017428 | 0.00235612 | 0.01040364 |
| MBLAC2 | -0.529458 | 3.00551442 | -3.8009437 | 0.00235963 | 0.01041607 |
| ITGA2B | -0.5361802 | 3.66996145 | -3.7993383 | 0.00236668 | 0.01043806 |
| LLPH-DT | 1.01188428 | 0.51106626 | 3.79715079 | 0.00237632 | 0.01047449 |
| AC138904.1 | 0.61861624 | 2.40087911 | 3.79679973 | 0.00237787 | 0.01047828 |
| AC009133.1 | -0.5066713 | 3.74763263 | -3.7963835 | 0.00237971 | 0.01048334 |
| U91328.1 | 0.85896894 | 1.62785006 | 3.79620133 | 0.00238052 | 0.01048384 |
| PITX3 | -1.2537863 | -0.5082855 | -3.7939956 | 0.0023903 | 0.0105156 |
| CDH8 | 0.91400475 | 0.89550116 | 3.79396159 | 0.00239045 | 0.0105156 |
| AOX1 | -1.1507925 | -0.0325837 | -3.7933397 | 0.00239322 | 0.01051972 |
| DEPDC1-AS1 | 1.04882353 | 0.21091395 | 3.79192008 | 0.00239954 | 0.01054309 |
| STK32B | -0.7058092 | 2.0980842 | -3.791389 | 0.00240191 | 0.01054657 |
| LINC01291 | 0.98087957 | 0.38816281 | 3.78865629 | 0.00241415 | 0.0105919 |
| PINLYP | 1.21734999 | -0.333272 | 3.78544397 | 0.00242862 | 0.0106492 |
| FAM149A | 0.7699027 | 1.69256084 | 3.7844276 | 0.00243321 | 0.01066317 |
| FBXO48 | 0.92666879 | 1.14333928 | 3.78346389 | 0.00243758 | 0.01067612 |
| SH3BGR | 0.83310071 | 1.65012465 | 3.77951376 | 0.00245556 | 0.01073003 |
| AC016065.1 | 0.53378105 | 2.97061159 | 3.7789727 | 0.00245803 | 0.01073774 |
| ATP2B1-AS1 | 0.77950587 | 1.42737957 | 3.7767349 | 0.00246828 | 0.01077011 |
| RGS17 | 0.96425667 | 0.35571663 | 3.77623334 | 0.00247059 | 0.01077706 |
| PTGER2 | -0.6156407 | 2.44796762 | -3.7741579 | 0.00248015 | 0.01081253 |
| PSMG3-AS1 | -0.5765666 | 3.08337564 | -3.7736111 | 0.00248267 | 0.01081968 |

| | | | | | |
|---|---|---|---|---|---|
| AC062029.1 | 0.6514818 | 2.5618358 | 3.77349349 | 0.00248322 | 0.01081968 |
| NAP1L5 | 0.53163864 | 2.90765047 | 3.77285031 | 0.00248619 | 0.01082952 |
| VWCE | 0.52176716 | 3.12900504 | 3.76205785 | 0.00253664 | 0.0110081 |
| SLC16A3 | -1.2511924 | 1.04574667 | -3.7527717 | 0.00258089 | 0.01115217 |
| FUT11 | -0.5037424 | 6.45246696 | -3.7503313 | 0.00259264 | 0.01119658 |
| PEX12 | 0.50752913 | 3.24359363 | 3.74913064 | 0.00259845 | 0.01121526 |
| AQP11 | 0.53446694 | 3.38182923 | 3.7479311 | 0.00260426 | 0.01123074 |
| NATD1 | 0.570431 | 2.90652639 | 3.74348117 | 0.00262594 | 0.01130492 |
| KIF5C | -0.6033732 | 2.50259639 | -3.7415023 | 0.00263564 | 0.01134023 |
| AC103808.3 | 0.79047845 | 1.31754022 | 3.74083787 | 0.00263891 | 0.01135035 |
| CH25H | -1.1362372 | -0.1959974 | -3.7356588 | 0.0026645 | 0.01144814 |
| ZNF416 | 0.52603612 | 3.19044036 | 3.72850846 | 0.00270026 | 0.01156238 |
| AC093512.2 | 0.57090178 | 2.7005833 | 3.72686125 | 0.00270856 | 0.01159139 |
| VAX2 | 0.56744925 | 2.81208451 | 3.72538222 | 0.00271604 | 0.01161355 |
| AC107068.1 | -0.7981436 | 1.13936786 | -3.7230242 | 0.00272801 | 0.01165815 |
| PRSS12 | -1.0254871 | 0.0314011 | -3.7226846 | 0.00272974 | 0.01166224 |
| GRIP1 | 0.85633887 | 1.09828677 | 3.72096153 | 0.00273853 | 0.0116866 |
| SLC27A2 | -0.5108805 | 3.26054779 | -3.7204599 | 0.00274109 | 0.01169423 |
| RBM24 | -1.1983166 | -0.262087 | -3.7190273 | 0.00274842 | 0.01172221 |
| AL157400.4 | 1.00545191 | -0.0784752 | 3.71790082 | 0.0027542 | 0.01173365 |
| AL135902.1 | 1.03702306 | 0.39362477 | 3.71660591 | 0.00276086 | 0.01175872 |
| PLA2G4A | -0.5601132 | 2.76652033 | -3.7133882 | 0.00277749 | 0.01180626 |
| GPC2 | -0.6096333 | 3.02889545 | -3.7118473 | 0.00278548 | 0.01183693 |
| KLF4 | 0.84191079 | 0.95675703 | 3.70836809 | 0.00280362 | 0.01190066 |
| SEMA3D | 0.6251575 | 2.2980607 | 3.70388579 | 0.00282717 | 0.01198047 |
| STYK1 | -0.8001551 | 1.34002556 | -3.7009634 | 0.00284263 | 0.01203335 |
| HOTAIR | 0.66908339 | 2.81156224 | 3.6983357 | 0.00285661 | 0.01208155 |
| AL356488.3 | 0.6905154 | 2.08677873 | 3.69730482 | 0.00286211 | 0.01209806 |
| GAREM1 | -0.6012304 | 2.61963743 | -3.6968776 | 0.0028644 | 0.01210096 |
| BBS12 | 0.67392953 | 2.17701836 | 3.69650658 | 0.00286638 | 0.01210255 |
| CISH | -0.7121553 | 1.68542642 | -3.6953203 | 0.00287274 | 0.01211827 |
| AP000866.1 | 0.71770814 | 1.67605489 | 3.69511578 | 0.00287383 | 0.01211827 |
| LINC00337 | -0.6542968 | 2.41596714 | -3.6950667 | 0.0028741 | 0.01211827 |
| AC025165.1 | -0.8091648 | 1.54718649 | -3.6845458 | 0.00293112 | 0.01231275 |
| AL645608.8 | -0.8303589 | 1.80847558 | -3.6843082 | 0.00293242 | 0.01231275 |
| NRTN | -0.6754096 | 2.27635761 | -3.6827307 | 0.00294107 | 0.01232516 |
| DMBX1 | -0.8681575 | 0.9079342 | -3.6778879 | 0.0029678 | 0.01241616 |
| LINC00621 | 0.98675262 | 0.46975689 | 3.67738607 | 0.00297058 | 0.01241616 |
| AC148477.2 | -0.793959 | 2.60431383 | -3.6772482 | 0.00297135 | 0.01241616 |
| MIR210HG | -1.1702651 | 1.19978722 | -3.6769559 | 0.00297297 | 0.01241759 |
| SMKR1 | -0.6706931 | 1.97109394 | -3.6745864 | 0.00298616 | 0.01246581 |
| PPFIA4 | -1.2300585 | 3.59210754 | -3.6717907 | 0.0030018 | 0.01251729 |
| PRR35 | -0.9055274 | 0.88401484 | -3.6715074 | 0.00300339 | 0.01252047 |
| AC068768.1 | -0.6636685 | 2.1084674 | -3.6698383 | 0.00301277 | 0.01255267 |
| MLPH | -1.2467126 | -0.3332538 | -3.668857 | 0.0030183 | 0.01255843 |

| | | | | | |
|---|---|---|---|---|---|
| CKMT2-AS1 | 0.50198316 | 3.12338716 | 3.66709569 | 0.00302826 | 0.01256876 |
| AC006148.1 | 0.62972292 | 2.03520968 | 3.66626006 | 0.00303299 | 0.0125815 |
| AC025171.2 | 0.62273214 | 2.31172699 | 3.66093734 | 0.00306332 | 0.01268298 |
| HAND2-AS1 | 0.66515455 | 2.62095427 | 3.66045484 | 0.00306608 | 0.01269095 |
| NHLRC3 | -0.502373 | 3.85058545 | -3.6458957 | 0.0031507 | 0.01297735 |
| OLIG2 | -0.6847298 | 2.0661337 | -3.6449914 | 0.00315603 | 0.01299225 |
| RAB30-AS1 | -0.5142237 | 2.82286823 | -3.6446348 | 0.00315814 | 0.01299738 |
| AL662796.1 | 0.64012134 | 2.54057378 | 3.64435263 | 0.00315981 | 0.01299788 |
| ZNF284 | 0.58268186 | 2.30781817 | 3.64432387 | 0.00315998 | 0.01299788 |
| RFTN2 | 0.98914621 | 0.91878109 | 3.64094052 | 0.00318004 | 0.0130733 |
| OVGP1 | 0.78097417 | 1.4776189 | 3.63982793 | 0.00318666 | 0.01308988 |
| AP001453.4 | -0.8598126 | 1.00503803 | -3.6368322 | 0.00320458 | 0.01314918 |
| ZNF470 | -0.5499815 | 3.72254455 | -3.6364064 | 0.00320713 | 0.0131561 |
| AL645728.1 | 0.75443524 | 1.70528668 | 3.63471939 | 0.00321727 | 0.01319054 |
| STUM | 1.6133746 | -0.4904047 | 3.62894723 | 0.00325221 | 0.01331217 |
| MAP1LC3B2 | -0.5045805 | 3.1341311 | -3.6287032 | 0.0032537 | 0.01331465 |
| CFAP47 | -0.9305562 | 0.38397301 | -3.6278235 | 0.00325906 | 0.01333299 |
| MYO7A | -0.7523493 | 2.29518276 | -3.626541 | 0.00326689 | 0.01335782 |
| TMEM86A | 1.04357474 | 0.18627263 | 3.62601781 | 0.00327009 | 0.01336729 |
| TSPAN18 | -1.050879 | 0.25004002 | -3.6256232 | 0.00327251 | 0.01337356 |
| TIGD3 | -0.5689139 | 2.76432963 | -3.6226172 | 0.00329097 | 0.01343612 |
| KIAA0040 | -1.2609568 | -0.6056681 | -3.6156612 | 0.00333411 | 0.01357396 |
| EDNRA | 0.81876639 | 1.24787057 | 3.61446559 | 0.00334159 | 0.01358984 |
| CHODL | -1.2144326 | -0.2463694 | -3.6133412 | 0.00334863 | 0.01361482 |
| LTBP2 | -0.6413374 | 2.42350094 | -3.6100588 | 0.00336928 | 0.01369142 |
| PSCA | -0.9691063 | 0.29029007 | -3.6081823 | 0.00338114 | 0.01372139 |
| C1orf226 | 1.23428941 | -0.6407413 | 3.60809241 | 0.00338171 | 0.01372139 |
| OXCT2 | -1.0457907 | 0.14186333 | -3.6080163 | 0.00338219 | 0.01372139 |
| AC016590.3 | 0.88341998 | 0.78035706 | 3.60788912 | 0.003383 | 0.01372139 |
| AL096865.1 | 0.58675519 | 2.86808428 | 3.60478839 | 0.0034027 | 0.01378654 |
| ENKUR | 0.95634221 | 0.32145213 | 3.60371965 | 0.00340952 | 0.0137994 |
| HOTAIRM1 | 0.56019326 | 2.67323136 | 3.59748844 | 0.00344956 | 0.01393164 |
| KLC4 | 0.5058473 | 3.31576459 | 3.59320719 | 0.00347734 | 0.01401943 |
| AC093157.1 | 0.50548562 | 3.22576317 | 3.59285656 | 0.00347963 | 0.01401943 |
| RFLNA | -0.8015717 | 1.23375555 | -3.590509 | 0.00349497 | 0.01405533 |
| LINC00320 | 0.64493947 | 2.15069895 | 3.58914462 | 0.00350392 | 0.01407609 |
| RHBDL1 | -0.620873 | 3.18053493 | -3.5870364 | 0.00351779 | 0.01412433 |
| PTK2B | 0.58147873 | 2.44842318 | 3.57522826 | 0.00359653 | 0.01437282 |
| TRAPPC6A | 0.56821199 | 2.79190882 | 3.57482288 | 0.00359926 | 0.01437876 |
| FTCDNL1 | 0.61992201 | 2.12722193 | 3.57465194 | 0.00360042 | 0.01437876 |
| AL132639.2 | 0.89366249 | 0.5666149 | 3.57435507 | 0.00360242 | 0.01438025 |
| TMEM74B | -0.701508 | 2.7983288 | -3.5668783 | 0.00365329 | 0.01455261 |
| THSD1 | -0.6805568 | 2.29719762 | -3.5636075 | 0.00367577 | 0.01462293 |
| CDKN2D | -0.5867235 | 2.60550494 | -3.563289 | 0.00367797 | 0.01462782 |
| AC148477.3 | -0.8565439 | 1.60253624 | -3.5626341 | 0.00368249 | 0.01463811 |

| | | | | | |
|---|---|---|---|---|---|
| ADGRB3 | 0.6788336 | 1.70374704 | 3.55155767 | 0.00375982 | 0.01489466 |
| ZNF677 | 0.62960207 | 2.02458561 | 3.54647153 | 0.00379588 | 0.01499827 |
| GATA2-AS1 | -0.5708026 | 2.40676925 | -3.5458193 | 0.00380053 | 0.0150088 |
| LINC01089 | 0.56855536 | 4.14573394 | 3.54527394 | 0.00380442 | 0.01501634 |
| KIF26B | 0.86115261 | 1.02761995 | 3.54293542 | 0.00382115 | 0.01506668 |
| KLHDC1 | 1.07748222 | 0.50541742 | 3.54105431 | 0.00383467 | 0.01510424 |
| CCDC149 | 0.63589396 | 1.98076915 | 3.53409498 | 0.0038851 | 0.01524339 |
| FOXG1-AS1 | 0.76963017 | 1.24439377 | 3.53186197 | 0.00390142 | 0.01528475 |
| SMPDL3B | -0.8822196 | 0.93293431 | -3.5318243 | 0.0039017 | 0.01528475 |
| ZNF703 | 0.5322413 | 3.19807121 | 3.52947337 | 0.00391896 | 0.01534357 |
| STRA6 | -0.8924796 | 0.89925117 | -3.529228 | 0.00392076 | 0.01534357 |
| NPEPL1 | -0.5615334 | 3.8957528 | -3.5285437 | 0.0039258 | 0.01535788 |
| WDR25 | 0.54606676 | 3.83363496 | 3.52666508 | 0.00393968 | 0.01539372 |
| ZNF862 | -0.5744009 | 2.71940589 | -3.5257975 | 0.0039461 | 0.01541485 |
| AC007750.1 | 0.93696322 | -0.0038406 | 3.52182745 | 0.00397564 | 0.0155102 |
| LOH12CR2 | 0.8828006 | 0.85809867 | 3.5189107 | 0.00399748 | 0.01556733 |
| AL121906.2 | -0.8244225 | 1.00852444 | -3.5169779 | 0.00401202 | 0.0156079 |
| SARDH | -0.8299475 | 1.28012355 | -3.5158016 | 0.00402089 | 0.01562774 |
| DICER1-AS1 | 0.88733964 | 1.11132451 | 3.51578639 | 0.00402101 | 0.01562774 |
| ATOH8 | 0.56340364 | 2.62810151 | 3.51575467 | 0.00402125 | 0.01562774 |
| WDR88 | 0.61072744 | 2.09800289 | 3.50647003 | 0.00409201 | 0.01586203 |
| RARA-AS1 | 0.70237097 | 1.73137082 | 3.50051602 | 0.00413805 | 0.01600497 |
| ASB3 | 0.58221093 | 2.34747202 | 3.49779423 | 0.00415927 | 0.01606929 |
| DNASE1L1 | -0.590876 | 2.42246292 | -3.4935993 | 0.0041922 | 0.01618411 |
| VDR | -0.5551101 | 2.49232747 | -3.4920061 | 0.00420477 | 0.01622025 |
| AL035563.1 | 0.96020299 | 0.08065835 | 3.49088591 | 0.00421364 | 0.01624616 |
| NMNAT2 | -0.6747373 | 1.9486663 | -3.4878428 | 0.00423781 | 0.01632275 |
| C12orf76 | 0.5013757 | 3.66679443 | 3.48215411 | 0.00428338 | 0.01646894 |
| FST | 0.99497218 | 0.10124172 | 3.48110348 | 0.00429185 | 0.01648477 |
| GRIP2 | -0.9573234 | 0.83332264 | -3.4783989 | 0.00431374 | 0.01655622 |
| PLCH2 | -0.5565282 | 2.43324874 | -3.4755847 | 0.00433663 | 0.01662722 |
| FEZ1 | -1.0476711 | 0.17086654 | -3.4716876 | 0.00436853 | 0.01670847 |
| AC005670.2 | 0.59276492 | 2.19266276 | 3.47057666 | 0.00437767 | 0.0167295 |
| LINC02327 | 0.51176629 | 2.75746579 | 3.47040243 | 0.0043791 | 0.01673076 |
| AC147651.1 | -0.7757773 | 1.59971619 | -3.4699783 | 0.0043826 | 0.01673989 |
| TNKS2-AS1 | 1.15298554 | -0.098135 | 3.46887481 | 0.00439171 | 0.0167663 |
| CXorf40A | -0.5487337 | 2.7389157 | -3.4621336 | 0.00444775 | 0.01692897 |
| LINC00663 | 0.76069773 | 1.88660517 | 3.4602544 | 0.00446351 | 0.01698466 |
| ELOVL7 | -0.9489527 | 0.26942506 | -3.4528672 | 0.00452598 | 0.01717975 |
| THBS4 | 0.86236858 | 0.46113932 | 3.45013429 | 0.00454932 | 0.01724187 |
| AL118558.4 | 1.05967474 | -0.5060263 | 3.4499838 | 0.00455061 | 0.01724244 |
| RAP2C-AS1 | 1.04335229 | -0.1154776 | 3.44940297 | 0.00455559 | 0.01725698 |
| AC092803.2 | 0.70233834 | 1.48574801 | 3.44251889 | 0.004615 | 0.01744712 |
| ESAM | -0.5449045 | 2.64511974 | -3.4422596 | 0.00461725 | 0.01745128 |
| AZU1 | -1.3680392 | -0.2110035 | -3.4377893 | 0.00465627 | 0.01756805 |

| | | | | | |
|---|---|---|---|---|---|
| MESP2 | -0.5901012 | 2.54522546 | -3.437266 | 0.00466086 | 0.01757223 |
| SNCA-AS1 | 0.91516925 | 0.24553914 | 3.43539022 | 0.00467735 | 0.01761247 |
| AC067930.5 | -0.8479923 | 0.59546184 | -3.4342083 | 0.00468777 | 0.01764731 |
| FAM84B | -0.8633209 | 0.62009615 | -3.4294232 | 0.00473019 | 0.0177653 |
| PDE2A | -0.934318 | 0.47098203 | -3.4284363 | 0.00473899 | 0.01778267 |
| LINC02249 | 0.88208097 | 0.4935859 | 3.42663597 | 0.00475509 | 0.0178298 |
| HOXB-AS3 | 0.67254933 | 1.74573081 | 3.42170381 | 0.00479946 | 0.01793843 |
| AC007388.1 | 0.6974777 | 1.58303331 | 3.42097604 | 0.00480604 | 0.01795037 |
| AC106744.2 | 0.82943445 | 0.67231965 | 3.41967272 | 0.00481785 | 0.01797612 |
| AC090515.2 | 0.92714648 | 0.35766122 | 3.41876508 | 0.00482609 | 0.017998 |
| AC011603.2 | 1.03966766 | -0.1961834 | 3.41783235 | 0.00483458 | 0.01802521 |
| EHD2 | -1.0244324 | 0.17119243 | -3.4148307 | 0.00486199 | 0.01809177 |
| PRRT2 | -0.5893747 | 2.57311929 | -3.4143183 | 0.00486669 | 0.01810479 |
| AC079949.1 | -1.0691313 | -0.2821984 | -3.4114458 | 0.00489309 | 0.01817622 |
| NRCAM | 1.31265737 | -0.7611976 | 3.41069529 | 0.00490001 | 0.01819747 |
| AC107032.2 | 1.1082588 | -0.524118 | 3.40994206 | 0.00490697 | 0.01821438 |
| SYNPO2 | -0.5942913 | 2.10997867 | -3.4094335 | 0.00491167 | 0.0182229 |
| SLC9B1 | 0.86604652 | 0.82196343 | 3.40768067 | 0.00492792 | 0.01827422 |
| HEYL | -0.7687923 | 1.39487947 | -3.4039168 | 0.00496299 | 0.01839525 |
| RBMS1 | -0.7247793 | 1.19842858 | -3.4017041 | 0.00498372 | 0.01845403 |
| MYOM2 | 0.79487149 | 1.10059083 | 3.40044102 | 0.0049956 | 0.01847993 |
| ACHE | 0.63953512 | 2.23786384 | 3.39763196 | 0.00502211 | 0.01857192 |
| AZIN1-AS1 | 0.58513886 | 2.13614306 | 3.39754652 | 0.00502292 | 0.01857192 |
| IGFL4 | 1.17182686 | -0.5865059 | 3.3950182 | 0.00504691 | 0.01861619 |
| SYNPO | -0.9868707 | 0.47607402 | -3.394056 | 0.00505607 | 0.01863532 |
| SH2D2A | -0.8540057 | 0.8381307 | -3.3898073 | 0.00509671 | 0.01873497 |
| HLF | 0.87848475 | 0.50885174 | 3.3884539 | 0.00510973 | 0.0187737 |
| Z97989.1 | 0.92304271 | 0.2511601 | 3.38405549 | 0.00515227 | 0.01889003 |
| HOXA2 | 0.75473841 | 1.22288667 | 3.38223176 | 0.00517001 | 0.01893086 |
| ENOX1 | 0.64624411 | 1.85367383 | 3.38197963 | 0.00517247 | 0.01893528 |
| AC005332.6 | 0.52212173 | 2.75823131 | 3.37908548 | 0.00520076 | 0.01901129 |
| ZNF185 | -0.789481 | 1.22175964 | -3.3788485 | 0.00520309 | 0.01901519 |
| CYP26B1 | -0.5516443 | 2.57605295 | -3.377447 | 0.00521685 | 0.01905171 |
| ICAM1 | -1.1021606 | -0.349346 | -3.3732511 | 0.00525828 | 0.01914759 |
| MANEA-DT | 0.95168687 | -0.4981978 | 3.37178459 | 0.00527284 | 0.01918215 |
| ING4 | 0.50256077 | 3.46669256 | 3.36935666 | 0.00529704 | 0.01925166 |
| AC024909.2 | -0.5677485 | 2.23449557 | -3.3690329 | 0.00530027 | 0.01925879 |
| FOXA3 | -1.0629366 | -0.2840249 | -3.3678838 | 0.00531177 | 0.01929271 |
| CCDC157 | 0.61223485 | 1.93833317 | 3.36736112 | 0.005317 | 0.0193057 |
| SLIT1 | -0.5416694 | 2.82601724 | -3.3662938 | 0.00532771 | 0.01933531 |
| ANKRD37 | -1.0396966 | 1.52934737 | -3.3660577 | 0.00533009 | 0.01933929 |
| NPR1 | -0.9562229 | 0.25663207 | -3.3639951 | 0.00535086 | 0.01940536 |
| RNF165 | -0.6955232 | 2.14911988 | -3.3630913 | 0.00535999 | 0.01942435 |
| ZBTB32 | -0.5618735 | 2.35111669 | -3.3627148 | 0.00536379 | 0.01942435 |
| AC026250.1 | -0.7970407 | 0.97800967 | -3.3602067 | 0.00538922 | 0.01948382 |

| | | | | | |
|---|---|---|---|---|---|
| AC005837.1 | 0.77411009 | 0.9152075 | 3.35005913 | 0.00549336 | 0.01981299 |
| AK1 | -0.5596083 | 2.44555365 | -3.349878 | 0.00549524 | 0.01981504 |
| ICA1 | -0.8322614 | 0.80252607 | -3.34933 | 0.00550092 | 0.01982609 |
| RASGRP3 | 0.50948397 | 3.04544827 | 3.3488669 | 0.00550573 | 0.0198387 |
| EFNA3 | -0.7098669 | 2.91646055 | -3.3416028 | 0.00558169 | 0.02005991 |
| AL118558.3 | 0.53050017 | 2.5822739 | 3.3378068 | 0.00562181 | 0.02017535 |
| ZNF19 | 0.57702703 | 2.20440445 | 3.33528476 | 0.00564862 | 0.02023241 |
| GTF2IRD2B | 0.87831233 | 0.86848391 | 3.33518043 | 0.00564973 | 0.02023241 |
| LRRC37A3 | 0.61470026 | 2.07334415 | 3.33354452 | 0.0056672 | 0.02028056 |
| KCNS3 | -1.0644023 | -0.3130396 | -3.3330047 | 0.00567297 | 0.02029331 |
| BCAM | -0.7636782 | 2.53351521 | -3.3292268 | 0.00571356 | 0.02039791 |
| CCDC163 | -0.5092237 | 2.70072985 | -3.328953 | 0.00571651 | 0.02039791 |
| AL135999.1 | -0.8242852 | 0.74893194 | -3.3268287 | 0.00573947 | 0.02045823 |
| HESX1 | 1.06848528 | 0.09021494 | 3.3264448 | 0.00574363 | 0.02045823 |
| GPRC5C | -1.2045785 | -0.4748546 | -3.3262836 | 0.00574538 | 0.02045823 |
| SPOCK1 | 1.21409068 | -0.874548 | 3.32592439 | 0.00574927 | 0.02045823 |
| NT5E | 0.51469772 | 2.69644405 | 3.32089462 | 0.00580411 | 0.02060492 |
| AC244517.1 | 1.18510655 | -0.55395 | 3.31651485 | 0.00585228 | 0.02074676 |
| AL031123.2 | 0.7928327 | 0.93265348 | 3.31527565 | 0.00586599 | 0.0207856 |
| CHSY3 | 0.54493531 | 2.35547975 | 3.31470415 | 0.00587232 | 0.02079344 |
| MANCR | 0.81470898 | 0.60438909 | 3.31201775 | 0.00590217 | 0.0208864 |
| PLEKHG6 | -1.3374192 | -0.8281819 | -3.3119691 | 0.00590271 | 0.0208864 |
| SLC10A7 | 0.52382418 | 2.34396501 | 3.31170573 | 0.00590564 | 0.02089191 |
| TP53I13 | -1.0353296 | -0.0705115 | -3.3069266 | 0.00595916 | 0.02102719 |
| IQCN | 0.84637062 | 0.44585436 | 3.30586996 | 0.00597106 | 0.02105935 |
| DAPK2 | -0.7174438 | 1.54476464 | -3.3048655 | 0.00598239 | 0.02108459 |
| CYP26A1 | 0.73813274 | 1.46127201 | 3.30442817 | 0.00598733 | 0.02109218 |
| AC109587.1 | 0.80598873 | 0.59924565 | 3.30270022 | 0.0060069 | 0.02113651 |
| FGD3 | -0.6400574 | 1.81855849 | -3.2973284 | 0.00606812 | 0.02128764 |
| PTGS2 | -0.9008307 | 0.23528227 | -3.2949721 | 0.00609518 | 0.02135894 |
| MMP24OS | -0.5503914 | 2.24189508 | -3.2949441 | 0.0060955 | 0.02135894 |
| SEPT4 | -0.6581267 | 1.83563256 | -3.2890371 | 0.00616387 | 0.02153868 |
| MAP7 | 0.98481013 | -0.1306931 | 3.28860648 | 0.00616888 | 0.02155122 |
| DRAXIN | -0.6371767 | 1.83200859 | -3.2882338 | 0.00617322 | 0.02156004 |
| MTMR11 | -0.6476586 | 2.33278489 | -3.2871623 | 0.00618573 | 0.02158517 |
| AGAP2-AS1 | -0.5529475 | 2.29265616 | -3.283832 | 0.00622475 | 0.02170134 |
| FCRLB | 0.54671833 | 2.30434705 | 3.28326485 | 0.00623142 | 0.02171459 |
| AC008060.2 | 1.1105408 | -0.4294462 | 3.28221423 | 0.00624379 | 0.02174403 |
| LAMA3 | 0.67612376 | 1.58445781 | 3.28198925 | 0.00624645 | 0.02174694 |
| STAM-AS1 | 0.7961399 | 0.57541475 | 3.2756104 | 0.00632215 | 0.02195497 |
| AC246817.1 | 0.68090283 | 1.48461294 | 3.27429332 | 0.00633789 | 0.02199451 |
| DOK6 | -0.8252672 | 1.05101302 | -3.2741075 | 0.00634012 | 0.0219959 |
| FBXO15 | 0.83378066 | 0.65324971 | 3.27277846 | 0.00635605 | 0.02202458 |
| AL441992.1 | -1.0071307 | -0.1628163 | -3.2700889 | 0.00638842 | 0.022104 |
| CTSO | 0.62074853 | 1.82015488 | 3.26825019 | 0.00641065 | 0.02216188 |

| | | | | | |
|---|---|---|---|---|---|
| FIGN | 0.8703223 | 0.42539373 | 3.26822056 | 0.00641101 | 0.02216188 |
| AL353759.1 | 1.14234284 | -0.2527263 | 3.2667734 | 0.00642855 | 0.02220144 |
| RAB6C | -0.9601053 | 0.00022898 | -3.2661657 | 0.00643594 | 0.02220473 |
| GALNT9 | -0.6136224 | 2.40007331 | -3.2660217 | 0.00643769 | 0.02220473 |
| BNC2 | 0.61020167 | 1.98355777 | 3.26566347 | 0.00644204 | 0.02221305 |
| SPATA3-AS1 | -0.8243365 | 0.61866504 | -3.2623695 | 0.00648225 | 0.0223408 |
| NAALADL2 | 0.68959359 | 1.50333747 | 3.26151075 | 0.00649277 | 0.02235777 |
| RASGEF1C | -0.7099641 | 1.49376295 | -3.2594268 | 0.00651838 | 0.02241537 |
| PURG | 0.79043438 | 0.75993454 | 3.25845365 | 0.00653038 | 0.02245151 |
| MFSD4A | -0.8048083 | 0.96730203 | -3.2541542 | 0.00658363 | 0.02260353 |
| AC246817.2 | 0.53342273 | 2.37329572 | 3.24708482 | 0.00667215 | 0.02284555 |
| AC027682.6 | -0.8417157 | 0.7526417 | -3.2443975 | 0.00670611 | 0.02294627 |
| GPR89B | 0.54920172 | 2.57607071 | 3.24280893 | 0.00672627 | 0.02300538 |
| RAB24 | -0.5326772 | 2.78381236 | -3.2420832 | 0.0067355 | 0.02303121 |
| USP46-AS1 | -0.8366381 | 0.19047434 | -3.2415297 | 0.00674255 | 0.02305011 |
| HOXA7 | 0.89182914 | 0.46541987 | 3.2401656 | 0.00675995 | 0.02309395 |
| GREM1 | -0.7492818 | 1.14481803 | -3.2397701 | 0.006765 | 0.02310079 |
| MAML3 | 0.5853852 | 2.07003741 | 3.23920838 | 0.00677219 | 0.0231201 |
| LYRM9 | 0.97951336 | 0.10299732 | 3.23868182 | 0.00677893 | 0.02313269 |
| AC064807.1 | 0.61436405 | 1.77757194 | 3.23581226 | 0.00681578 | 0.02322298 |
| HMGCLL1 | -0.7070788 | 1.72993597 | -3.2350551 | 0.00682554 | 0.02323851 |
| DLL1 | -0.5236642 | 2.95906014 | -3.2145962 | 0.0070946 | 0.0239508 |
| AC093326.1 | 0.85591632 | 0.45253909 | 3.21414115 | 0.00710071 | 0.02396073 |
| ABCC8 | -0.5489994 | 2.66334969 | -3.2046015 | 0.0072299 | 0.02430998 |
| AC125611.4 | -1.2160944 | -0.6169185 | -3.2041232 | 0.00723643 | 0.02432117 |
| GAL3ST4 | -0.8421067 | 0.60286919 | -3.2030647 | 0.00725093 | 0.02434826 |
| SPIN2B | 0.65270671 | 1.49380772 | 3.20287153 | 0.00725358 | 0.02435175 |
| AC009163.7 | 0.69539841 | 1.37700983 | 3.20147771 | 0.00727271 | 0.02439435 |
| FLT3 | 0.8706142 | 0.3218495 | 3.19719348 | 0.00733185 | 0.0245492 |
| KCTD21 | -0.5800329 | 2.09819142 | -3.1969301 | 0.0073355 | 0.02455599 |
| LINC02482 | 0.8890258 | 0.45221508 | 3.19613861 | 0.00734648 | 0.02458189 |
| AC011448.1 | -0.8052788 | 0.53959445 | -3.1955226 | 0.00735504 | 0.02460509 |
| C2orf50 | -0.9078004 | 0.31572601 | -3.1944799 | 0.00736955 | 0.02462643 |
| SEMA3E | -0.532846 | 2.44444707 | -3.1937379 | 0.00737989 | 0.02465304 |
| AL391095.1 | -0.7714063 | 0.54488536 | -3.1936751 | 0.00738077 | 0.02465304 |
| AL355488.1 | 0.60242564 | 2.62675842 | 3.19262168 | 0.00739548 | 0.0246804 |
| SCARF1 | -1.025419 | -0.0502522 | -3.1920355 | 0.00740368 | 0.02469688 |
| AC116565.2 | 0.97379619 | -0.1489036 | 3.191819 | 0.00740671 | 0.02470155 |
| GAD1 | 0.5050392 | 3.41891306 | 3.1914057 | 0.0074125 | 0.02470997 |
| TGM2 | -0.7219568 | 1.50122736 | -3.191009 | 0.00741806 | 0.02472306 |
| ZNF513 | -0.5299377 | 2.19868665 | -3.1885317 | 0.00745288 | 0.0248118 |
| PLA2G4B | -0.8927902 | 0.2584808 | -3.1870452 | 0.00747386 | 0.02484885 |
| TCHH | 0.95322287 | -0.2860656 | 3.18152743 | 0.00755222 | 0.02506537 |
| AC083843.3 | 0.54610155 | 2.73646744 | 3.18115622 | 0.00755753 | 0.02507747 |
| JAKMIP2 | 0.89938569 | 0.58283645 | 3.17903756 | 0.00758786 | 0.02515056 |

| | | | | | |
|---|---|---|---|---|---|
| NKX2-8 | -0.6667558 | 1.72977354 | -3.1764895 | 0.0076245 | 0.02525542 |
| HSF4 | -0.7065936 | 2.31460241 | -3.1727181 | 0.00767906 | 0.02540834 |
| TMEM88 | -0.6501464 | 1.73729332 | -3.170913 | 0.00770531 | 0.02545071 |
| VSTM4 | -0.7205481 | 1.12172521 | -3.1707791 | 0.00770726 | 0.0254516 |
| GDPD3 | -0.7908587 | 0.93608632 | -3.1673711 | 0.00775708 | 0.0255938 |
| RASGRF1 | -1.5556257 | 0.67200328 | -3.1650196 | 0.00779164 | 0.02569664 |
| SATB2-AS1 | 0.77783928 | 0.80319978 | 3.16215611 | 0.00783394 | 0.02581365 |
| TRAPPC2B | 0.56235482 | 1.96103517 | 3.16168147 | 0.00784098 | 0.02582309 |
| HHLA3 | 0.76489063 | 0.89000897 | 3.15630151 | 0.00792114 | 0.02604431 |
| DEGS2 | -1.0411769 | 1.75796875 | -3.155644 | 0.007931 | 0.02606873 |
| MUC1 | -1.0137442 | -0.2921558 | -3.1540206 | 0.00795538 | 0.02612523 |
| KRBA2 | 0.79953377 | 0.63936011 | 3.14842143 | 0.00804005 | 0.02637227 |
| CCDC181 | 0.76555617 | 0.90164964 | 3.14774228 | 0.00805038 | 0.02638807 |
| SYT5 | -0.7325698 | 1.09464823 | -3.1476995 | 0.00805103 | 0.02638807 |
| C6orf52 | 0.91557309 | -0.0123942 | 3.14566079 | 0.00808213 | 0.02644727 |
| SCN4B | -0.820751 | 0.77760561 | -3.1451743 | 0.00808957 | 0.02646018 |
| P4HA3 | -0.9993626 | -0.4378821 | -3.1437423 | 0.0081115 | 0.02652046 |
| KIF6 | 0.8861255 | 0.04842562 | 3.14283395 | 0.00812545 | 0.02654886 |
| MATN3 | -0.6223009 | 1.5770485 | -3.1420809 | 0.00813703 | 0.02657112 |
| FAM110C | -0.676332 | 1.75606965 | -3.1420305 | 0.0081378 | 0.02657112 |
| SSPO | -0.9651033 | 0.89921046 | -3.1391803 | 0.00818178 | 0.02666286 |
| BHLHE40-AS1 | 0.7770863 | 0.7404004 | 3.13855323 | 0.00819149 | 0.02667319 |
| AC009404.1 | 0.65846742 | 1.70434707 | 3.13582647 | 0.00823383 | 0.02677012 |
| GRIN1 | -0.7140678 | 1.34690199 | -3.1354984 | 0.00823894 | 0.02678098 |
| CALB2 | -0.8545386 | 0.29101501 | -3.1352773 | 0.00824239 | 0.02678285 |
| OR2B6 | 1.04557614 | -0.3858423 | 3.13362595 | 0.00826817 | 0.02684747 |
| AC007952.4 | 0.62339569 | 2.95778644 | 3.13122355 | 0.00830581 | 0.02692212 |
| AL121821.1 | 0.52613729 | 2.14472933 | 3.13112379 | 0.00830738 | 0.02692212 |
| DSCR8 | 0.54670899 | 2.09538751 | 3.13035334 | 0.00831949 | 0.02693939 |
| FAM124A | -0.5935262 | 2.03967611 | -3.1288759 | 0.00834277 | 0.02700831 |
| ZNF585A | 0.57637027 | 1.82122995 | 3.12585178 | 0.00839061 | 0.02715159 |
| SIM2 | -0.6032123 | 1.7210337 | -3.1204508 | 0.00847675 | 0.02738351 |
| ACTL10 | -0.5195697 | 2.34585036 | -3.1142123 | 0.00857735 | 0.02764948 |
| SHISA8 | -0.8612863 | 0.65104272 | -3.1090772 | 0.00866105 | 0.02783042 |
| PCAT6 | 0.91838934 | 0.23968277 | 3.10705431 | 0.00869424 | 0.02790747 |
| MXI1 | -0.5090749 | 4.77299962 | -3.1006709 | 0.00879983 | 0.02817473 |
| WDR86 | 0.81020789 | 0.80336943 | 3.09975923 | 0.00881502 | 0.02820546 |
| ERBB3 | -0.5036186 | 2.90737968 | -3.0970773 | 0.00885984 | 0.0283297 |
| LRRC10B | -0.6247855 | 1.81044126 | -3.0965189 | 0.0088692 | 0.02834657 |
| GTSE1-DT | 0.62599177 | 1.60342295 | 3.09488707 | 0.00889662 | 0.02840052 |
| ME1 | -0.6995552 | 0.96607149 | -3.0899744 | 0.00897966 | 0.02862065 |
| AL162171.1 | 0.71217392 | 1.07636709 | 3.08924519 | 0.00899205 | 0.0286508 |
| LINC00526 | 0.73554256 | 0.88837671 | 3.08747367 | 0.00902222 | 0.02872881 |
| GMPR | -0.5506944 | 2.10355917 | -3.0857418 | 0.00905182 | 0.02881094 |
| BX088651.4 | 0.57688568 | 1.86108983 | 3.08369735 | 0.00908689 | 0.02888005 |

| | | | | | |
|---|---|---|---|---|---|
| LINC01560 | 0.62246944 | 1.33961963 | 3.08193659 | 0.00911719 | 0.02896421 |
| CNTF | -0.7401144 | 0.79259995 | -3.081669 | 0.00912181 | 0.02897279 |
| MIMT1 | 0.94378656 | 0.09388511 | 3.08053275 | 0.00914143 | 0.02901003 |
| LINC02021 | 0.88644547 | -0.0887605 | 3.07952646 | 0.00915884 | 0.02902826 |
| SMIM10L2A | -0.7346886 | 0.94033279 | -3.0786879 | 0.00917338 | 0.02905735 |
| AHRR | -0.8385739 | 0.43671447 | -3.0694253 | 0.00933548 | 0.02946603 |
| AC008687.4 | -0.8896845 | -0.1949872 | -3.0683786 | 0.00935398 | 0.02951826 |
| C18orf65 | 0.73793263 | 0.82285476 | 3.06745947 | 0.00937025 | 0.02953883 |
| LINC00703 | 0.50408178 | 2.17239783 | 3.06214591 | 0.00946488 | 0.02981232 |
| TBC1D10C | -0.8697559 | 0.2141003 | -3.0547544 | 0.00959811 | 0.03019427 |
| GPR61 | 0.7185336 | 0.81129803 | 3.0537057 | 0.00961716 | 0.03023536 |
| SLC10A5 | 0.97481089 | -0.6057741 | 3.05189144 | 0.00965022 | 0.03032667 |
| SH3GL3 | 0.97520897 | -0.5002561 | 3.0491981 | 0.00969949 | 0.03044993 |
| COBL | -0.7480943 | 0.76154716 | -3.0463073 | 0.00975266 | 0.03058984 |
| C17orf113 | 0.59900149 | 2.07991185 | 3.0459939 | 0.00975844 | 0.03059692 |
| ABCA1 | -0.9397607 | 0.22807488 | -3.0454886 | 0.00976777 | 0.03061983 |
| AL645608.7 | -0.769816 | 0.78318623 | -3.0441503 | 0.00979252 | 0.03068843 |
| FGD5 | -0.5680209 | 2.01957578 | -3.0439732 | 0.0097958 | 0.03068843 |
| AC133785.1 | -0.9265947 | 0.32911788 | -3.0420456 | 0.00983157 | 0.0307816 |
| CALML6 | -0.7185385 | 1.68571645 | -3.0398212 | 0.00987301 | 0.03089219 |
| LINC00896 | -1.1293259 | -0.9366454 | -3.0380067 | 0.00990695 | 0.03096636 |
| AC008738.2 | 0.80865671 | 0.30644917 | 3.03787106 | 0.00990949 | 0.03096791 |
| CCDC170 | 0.99441767 | -0.3571264 | 3.03773395 | 0.00991206 | 0.03096817 |
| AC011815.1 | 0.6597939 | 1.33304681 | 3.03387319 | 0.00998468 | 0.03113864 |
| GAS6-DT | -0.9896648 | -0.408543 | -3.0294563 | 0.01006842 | 0.03129665 |
| PLAU | -0.935563 | -0.0117262 | -3.0274693 | 0.01010632 | 0.03140156 |
| NTN3 | -0.5747569 | 1.83144111 | -3.0257002 | 0.01014018 | 0.03146803 |
| IL1RAP | -0.9217602 | -0.120043 | -3.0251808 | 0.01015015 | 0.0314925 |
| RN7SL832P | 0.79796144 | 0.25603746 | 3.02455042 | 0.01016225 | 0.03151714 |
| ST8SIA6 | 0.59092852 | 1.70582023 | 3.02443121 | 0.01016454 | 0.03151779 |
| AL138724.1 | 0.60581833 | 1.4081164 | 3.02419656 | 0.01016905 | 0.03152532 |
| LINC02282 | 0.68199877 | 1.13926205 | 3.02252664 | 0.01020121 | 0.0316056 |
| TMEM132B | 1.16591084 | -1.0421823 | 3.02122161 | 0.01022642 | 0.03166425 |
| LINC00535 | 0.82938264 | 0.30278082 | 3.02005129 | 0.01024907 | 0.03170865 |
| AL592494.3 | 0.64142244 | 1.30425997 | 3.01828536 | 0.01028335 | 0.03178852 |
| SENP8 | 0.53667262 | 2.10121722 | 3.01511366 | 0.0103452 | 0.03194711 |
| ITGBL1 | 0.81643794 | 0.50712966 | 3.01445749 | 0.01035804 | 0.03198025 |
| AC116351.2 | -0.9299439 | 0.05881465 | -3.0117798 | 0.01041062 | 0.03206947 |
| LINC01551 | 0.51352849 | 2.52025475 | 3.00679015 | 0.01050929 | 0.03231036 |
| TMEM171 | -0.9503413 | 0.02483017 | -3.0051828 | 0.01054128 | 0.03239325 |
| AC137767.1 | 0.57077858 | 1.80470406 | 3.00418948 | 0.01056109 | 0.03242203 |
| AC234582.1 | -0.6925601 | 1.38047533 | -3.0012017 | 0.01062091 | 0.03256607 |
| PABPC4L | -0.5582556 | 2.32345084 | -2.9969141 | 0.01070735 | 0.03278463 |
| PHOSPHO2 | 0.54719174 | 1.52061752 | 2.99422269 | 0.01076197 | 0.03288536 |
| HOXD1 | -0.5863682 | 1.91387698 | -2.9939177 | 0.01076818 | 0.03289769 |

| | | | | | |
|---|---|---|---|---|---|
| AL451085.2 | 0.88259305 | 0.07280389 | 2.99135679 | 0.01082043 | 0.03303068 |
| AC110285.2 | 0.65069906 | 2.11971655 | 2.98708026 | 0.01090826 | 0.03324516 |
| PEX11G | 0.65965401 | 1.24445214 | 2.98422714 | 0.01096725 | 0.03340478 |
| ZNF404 | -0.5678588 | 2.60540264 | -2.9791479 | 0.01107305 | 0.03365933 |
| NXPH1 | 0.62325158 | 1.41913914 | 2.97527793 | 0.01115435 | 0.0338385 |
| CNFN | -0.7744195 | 0.64412982 | -2.9744341 | 0.01117215 | 0.03388572 |
| DENND2A | 0.9930452 | -0.3596562 | 2.97024088 | 0.01126105 | 0.03411698 |
| HOXC13-AS | 0.67583701 | 1.07687978 | 2.97019996 | 0.01126192 | 0.03411698 |
| AC025031.4 | 0.79278429 | 0.54198586 | 2.96604445 | 0.01135071 | 0.03435162 |
| HIST2H2AA4 | 0.67112438 | 1.16002544 | 2.96496062 | 0.01137399 | 0.03440133 |
| HOXC-AS1 | 0.67127262 | 1.20489637 | 2.96493085 | 0.01137463 | 0.03440133 |
| VSTM2L | -0.633606 | 2.16058962 | -2.9572221 | 0.01154155 | 0.03477624 |
| RASAL2-AS1 | 0.66091007 | 1.12357604 | 2.95175935 | 0.01166131 | 0.03505338 |
| EPHA8 | -0.6791722 | 1.06531528 | -2.94932 | 0.01171518 | 0.0351907 |
| RGMA | 0.67904595 | 1.07576617 | 2.94927011 | 0.01171629 | 0.0351907 |
| AC245297.3 | -0.6424087 | 1.33380819 | -2.9473795 | 0.01175822 | 0.03526767 |
| MSS51 | -0.6809559 | 1.57530443 | -2.9443304 | 0.01182615 | 0.03545039 |
| TP53TG3D | 0.82526541 | 0.11712717 | 2.94102885 | 0.01190016 | 0.03563934 |
| ABTB1 | -0.6180937 | 1.37833056 | -2.9376016 | 0.01197746 | 0.03584007 |
| LINC01473 | 0.79215012 | 0.24368027 | 2.93749433 | 0.01197989 | 0.03584025 |
| ST6GALNAC3 | -0.7435837 | 1.31250789 | -2.9355962 | 0.01202293 | 0.0359406 |
| PLEKHN1 | -0.7351756 | 0.5164043 | -2.9351338 | 0.01203344 | 0.03596492 |
| EPHA5 | 0.88498282 | 0.10531254 | 2.93441525 | 0.01204978 | 0.03599214 |
| ITGB7 | -1.0143164 | -0.7969642 | -2.9310055 | 0.01212765 | 0.03618223 |
| AL354892.2 | 0.85154782 | -0.0083793 | 2.93074889 | 0.01213353 | 0.03619264 |
| DLGAP4-AS1 | 0.64550343 | 1.35710201 | 2.92996708 | 0.01215147 | 0.0362192 |
| NOXRED1 | 0.90409791 | -0.1296827 | 2.92509171 | 0.0122639 | 0.03647371 |
| AC106739.1 | 0.80851061 | 0.04050589 | 2.92420465 | 0.01228446 | 0.03652053 |
| GGACT | -0.668336 | 1.0269253 | -2.9214818 | 0.01234781 | 0.03668003 |
| AC104073.4 | 0.71302643 | 0.9191997 | 2.92087492 | 0.01236197 | 0.03670594 |
| ARHGAP23 | -0.5231957 | 2.12526595 | -2.9200787 | 0.01238057 | 0.03674854 |
| AC022400.6 | 0.59269332 | 1.50967409 | 2.91966391 | 0.01239028 | 0.03676292 |
| TBX5 | 0.51534416 | 2.21030576 | 2.91892909 | 0.01240749 | 0.03680677 |
| SALRNA1 | 0.50760806 | 2.14598164 | 2.91559162 | 0.01248595 | 0.03701051 |
| OOEP | 0.8023957 | 0.25605346 | 2.9145243 | 0.01251114 | 0.03705262 |
| SEPT1 | -0.7375335 | 1.23294053 | -2.9144909 | 0.01251193 | 0.03705262 |
| AC004803.1 | 0.6643407 | 1.10959538 | 2.91381303 | 0.01252796 | 0.03708423 |
| SUSD2 | -0.945928 | 0.01910686 | -2.9113783 | 0.0125857 | 0.03721149 |
| LINC01719 | -0.7237621 | 0.8536256 | -2.910879 | 0.01259757 | 0.03723933 |
| IRF6 | -0.7399991 | 0.88168195 | -2.9081124 | 0.01266356 | 0.0373906 |
| NMRK1 | -0.7968407 | 0.92358297 | -2.9028291 | 0.01279054 | 0.03767064 |
| CRIP1 | -0.9169329 | -0.480605 | -2.9027173 | 0.01279324 | 0.03767064 |
| AC090061.1 | 0.75251853 | 0.39780365 | 2.89895296 | 0.0128845 | 0.03790251 |
| OPN3 | -0.8900133 | 0.01535574 | -2.8962069 | 0.01295148 | 0.03806257 |
| ODF3B | -0.636226 | 1.22192807 | -2.8926029 | 0.01303991 | 0.03827013 |

| | | | | | |
|---|---|---|---|---|---|
| AC004805.1 | -0.930713 | -0.516461 | -2.8925923 | 0.01304017 | 0.03827013 |
| AC005726.1 | -0.8866229 | -0.1693008 | -2.8919343 | 0.01305638 | 0.03828999 |
| NKX2-3 | -0.7480106 | 0.85048524 | -2.8916267 | 0.01306396 | 0.03830393 |
| RANBP3L | 0.70435639 | 1.01444556 | 2.89135051 | 0.01307078 | 0.03831324 |
| CYBA | -0.9417073 | -0.1875075 | -2.8908325 | 0.01308356 | 0.03833171 |
| FN3K | -0.5665474 | 1.72532869 | -2.8901991 | 0.01309922 | 0.03836273 |
| HAL | -0.8751328 | -0.2431364 | -2.8835608 | 0.01326441 | 0.038779 |
| KCNK13 | -0.731424 | 0.57625687 | -2.8832019 | 0.0132734 | 0.03879779 |
| AC007541.1 | 0.55625669 | 2.0103591 | 2.87935649 | 0.0133701 | 0.03904275 |
| PPP1R32 | 1.05947938 | -0.6064984 | 2.87843467 | 0.01339338 | 0.03909471 |
| TOX2 | -0.5669987 | 1.69201503 | -2.8781412 | 0.0134008 | 0.03909471 |
| SCN1B | 0.55875453 | 2.03764946 | 2.87558043 | 0.01346573 | 0.03923876 |
| RBM26-AS1 | 0.65876583 | 0.99349864 | 2.87268765 | 0.01353945 | 0.03940808 |
| NEUROG2 | 1.0845192 | -0.2832586 | 2.87090283 | 0.01358513 | 0.03949549 |
| RPS6KA2 | -0.647225 | 1.59588998 | -2.8691073 | 0.01363125 | 0.03960674 |
| AL021368.2 | 0.75237649 | 0.49974464 | 2.8686356 | 0.01364339 | 0.0396344 |
| B3GNT4 | -0.5640569 | 2.11777305 | -2.8681277 | 0.01365647 | 0.03966479 |
| TSNAXIP1 | 0.57359511 | 1.50336144 | 2.86740871 | 0.01367501 | 0.03970342 |
| KCNN2 | 0.54854986 | 1.680117 | 2.86708257 | 0.01368343 | 0.03971659 |
| AC067735.1 | -0.5282698 | 2.09580055 | -2.8670297 | 0.01368479 | 0.03971659 |
| AL356489.2 | 0.51937769 | 1.92856156 | 2.86665655 | 0.01369443 | 0.03973695 |
| LDHD | -0.9924095 | -0.585529 | -2.8656127 | 0.01372144 | 0.03980768 |
| S1PR3 | -0.575891 | 1.78029948 | -2.8638669 | 0.01376671 | 0.03990081 |
| SLITRK1 | 0.57852279 | 1.53756699 | 2.85971025 | 0.01387511 | 0.04016885 |
| IL11RA | -0.8290627 | 1.48599016 | -2.8595269 | 0.01387991 | 0.0401697 |
| ASIC3 | -0.543002 | 2.22431003 | -2.8562311 | 0.01396649 | 0.04035187 |
| SH2D3A | -0.9744427 | -0.315658 | -2.8560861 | 0.01397031 | 0.04035187 |
| ZBTB20 | 0.66041473 | 0.9711001 | 2.85285426 | 0.01405575 | 0.04051365 |
| AC145285.2 | 0.87105599 | -0.2576689 | 2.85108013 | 0.01410288 | 0.040634 |
| CXCR4 | -1.5799691 | 1.18480272 | -2.8491366 | 0.01415468 | 0.04075157 |
| RAB43 | -0.8803779 | -0.0242932 | -2.8471368 | 0.01420818 | 0.04085188 |
| NLRP6 | -0.9521627 | -0.5958341 | -2.8445828 | 0.01427679 | 0.04101021 |
| PPIL6 | 0.58071678 | 1.38805101 | 2.84308938 | 0.01431706 | 0.04110251 |
| HIST1H1T | 0.54215234 | 1.90341472 | 2.84144428 | 0.01436155 | 0.0412068 |
| SIX6 | 0.54460623 | 1.94827567 | 2.8412135 | 0.01436781 | 0.04121693 |
| AC044802.1 | 0.73334448 | 0.20248465 | 2.84072136 | 0.01438115 | 0.04123177 |
| BATF2 | -0.8656956 | 0.15143966 | -2.8387799 | 0.0144339 | 0.04132879 |
| SLC3A1 | 0.67134497 | 1.97948105 | 2.83841443 | 0.01444386 | 0.04134892 |
| AC004943.1 | 0.71022057 | 0.55521584 | 2.83645389 | 0.01449736 | 0.04146288 |
| FAM92B | -0.8222757 | 0.39694078 | -2.8343118 | 0.01455604 | 0.04161499 |
| PRSS23 | -0.5357981 | 2.18226678 | -2.8312134 | 0.01464134 | 0.04181148 |
| GAP43 | -0.5739748 | 1.37606484 | -2.8296084 | 0.01468571 | 0.04190659 |
| DNAJC27-AS1 | 0.73795909 | 0.4201952 | 2.82893513 | 0.01470437 | 0.04192822 |
| NOXA1 | -0.6374162 | 1.77809343 | -2.8257943 | 0.0147917 | 0.04213757 |
| AL136038.3 | 0.8095792 | -0.3217624 | 2.82539875 | 0.01480274 | 0.04216108 |

| | | | | | |
|---|---|---|---|---|---|
| AL359711.2 | 0.67439979 | 0.67642089 | 2.82350535 | 0.01485568 | 0.04226415 |
| SRGAP1 | -0.5010424 | 2.13126566 | -2.8231874 | 0.01486458 | 0.04228155 |
| FERMT3 | -0.5081836 | 1.97508079 | -2.8225818 | 0.01488157 | 0.04231395 |
| RUSC1-AS1 | -0.5735116 | 3.34133982 | -2.8216403 | 0.014908 | 0.04238116 |
| PIK3IP1 | -0.610363 | 2.54330992 | -2.8208156 | 0.0149312 | 0.04243117 |
| CD4 | -0.6541648 | 1.05810866 | -2.8194099 | 0.01497082 | 0.04252483 |
| NUPR1 | 1.02081981 | -0.8493713 | 2.81666161 | 0.01504858 | 0.04270492 |
| AL161421.1 | 0.57117907 | 1.50660054 | 2.81446472 | 0.01511102 | 0.04283767 |
| SHLD1 | 0.66119119 | 1.06456099 | 2.81086885 | 0.01521378 | 0.04307414 |
| FAM78B | 0.85736198 | -0.2743735 | 2.80906232 | 0.01526567 | 0.04320328 |
| ZNF385A | -0.6258933 | 1.26609605 | -2.8072523 | 0.01531783 | 0.0433266 |
| AC079949.2 | -0.580502 | 1.45637268 | -2.8067924 | 0.01533112 | 0.04334798 |
| LEKR1 | 0.68459266 | 0.62864188 | 2.80660593 | 0.0153365 | 0.04335512 |
| AC015922.4 | -0.8097499 | -0.1265368 | -2.8039741 | 0.01541275 | 0.04353746 |
| PSMB8 | -0.8648195 | 0.013155 | -2.7997121 | 0.01553703 | 0.04378522 |
| SEPSECS-AS1 | 0.85602961 | -0.5617009 | 2.79474747 | 0.01568304 | 0.04407143 |
| GRASP | -0.5964701 | 1.61305508 | -2.7933737 | 0.01572368 | 0.04417743 |
| PTPRH | -0.6925975 | 0.84990096 | -2.7932612 | 0.01572701 | 0.0441786 |
| MAGEE1 | -0.5957595 | 1.39571156 | -2.7921814 | 0.01575903 | 0.04423573 |
| DOCK8 | -0.9169542 | -0.3959616 | -2.7913447 | 0.01578389 | 0.04428088 |
| AXDND1 | 0.82710338 | -0.1353278 | 2.79007257 | 0.01582176 | 0.04434603 |
| STX11 | -0.9959574 | -0.6347328 | -2.7883979 | 0.01587175 | 0.044445 |
| AC026780.2 | 0.95067178 | -0.5776906 | 2.78586883 | 0.01594753 | 0.04459949 |
| CARD8-AS1 | 0.6949614 | 0.77073189 | 2.78412232 | 0.01600008 | 0.0446969 |
| NTN4 | 0.57763245 | 1.5343765 | 2.77585607 | 0.01625109 | 0.04528115 |
| IFNLR1 | -0.6387036 | 1.20953606 | -2.77302 | 0.0163381 | 0.04547339 |
| TENT5C | 0.76028009 | 0.09262841 | 2.76918252 | 0.01645657 | 0.04571906 |
| DRAIC | 0.70192639 | 0.91383363 | 2.76868767 | 0.01647191 | 0.04574489 |
| CTSK | 0.51433744 | 2.10186324 | 2.76406206 | 0.01661596 | 0.04607732 |
| LINC00511 | 0.70821313 | 0.48231069 | 2.76338242 | 0.01663723 | 0.04611235 |
| AC007292.1 | -0.65169 | 0.99621131 | -2.760975 | 0.01671278 | 0.04626956 |
| YPEL2 | 0.59555936 | 1.44797925 | 2.75836277 | 0.01679515 | 0.0464721 |
| STBD1 | -0.6503947 | 1.04571707 | -2.756264 | 0.01686161 | 0.04663045 |
| HLA-DMA | -0.9220744 | 0.09017816 | -2.7559183 | 0.01687259 | 0.04665227 |
| AP001107.4 | -0.6705957 | 0.50905719 | -2.7553619 | 0.01689026 | 0.04669261 |
| AC009121.1 | 0.89884547 | -0.4903868 | 2.75506836 | 0.01689959 | 0.04669668 |
| AC004825.2 | 0.76578171 | 0.30535296 | 2.75325315 | 0.01695742 | 0.04682696 |
| AL391069.3 | 0.75934752 | 0.29125702 | 2.75112859 | 0.01702534 | 0.04698882 |
| DSC2 | -0.8673882 | -0.6603035 | -2.7441784 | 0.01724941 | 0.04747744 |
| VWA5B2 | -0.5991256 | 1.60994706 | -2.7433802 | 0.01727534 | 0.04754014 |
| RAPGEF4 | 0.71009786 | 0.62799826 | 2.73929291 | 0.01740866 | 0.04782011 |
| EDA | -0.5246238 | 1.93038038 | -2.7366679 | 0.01749481 | 0.04798429 |
| WEE2-AS1 | 0.69973409 | 0.40413666 | 2.72922974 | 0.01774123 | 0.0484786 |
| JUP | -0.5235085 | 1.8716825 | -2.7267533 | 0.01782402 | 0.04859959 |
| TCN2 | 0.62871369 | 1.36864295 | 2.72501927 | 0.01788222 | 0.04870565 |

| | | | | | |
|---|---|---|---|---|---|
| PIP5KL1 | -0.7000275 | 0.76374036 | -2.7098416 | 0.01839965 | 0.04977473 |
| LINC01876 | 0.58807821 | 1.75108975 | 2.70905369 | 0.01842691 | 0.04983956 |
| AC011287.1 | 0.69663786 | 1.02202698 | 2.70749073 | 0.0184811 | 0.04995044 |
