## Supplementary Tables for "Zinc finger protein SALL4 functions through an AT-rich motif to regulate gene expression": Suppl_Table5 Sheet1.pdf

**Supplementary Table 5. List of differentially expressed genes bound by SALL4 in CUT&RUN.**

| gene_id | logFC | AveExpr | t | P.Value | adj.P.Val |
|---|---|---|---|---|---|
| GNB4 | -2.00967685 | 5.40672299 | -21.9137179 | 2.4296E-11 | 1.5462E-07 |
| KTN1 | -1.39685187 | 8.37472703 | -20.445608 | 5.6498E-11 | 1.6555E-07 |
| SALL4 | -1.62951354 | 5.62655801 | -18.2045898 | 2.3039E-10 | 2.1802E-07 |
| DIXDC1 | -1.41234877 | 6.13750561 | -15.9946007 | 1.0896E-09 | 5.2546E-07 |
| ITGAV | -1.01356443 | 8.93550924 | -15.5016215 | 1.5829E-09 | 6.979E-07 |
| CACUL1 | 1.16564962 | 7.39983835 | 15.4769181 | 1.6133E-09 | 6.979E-07 |
| TXNRD1 | -1.04104161 | 8.47228229 | -14.9534382 | 2.4287E-09 | 9.4291E-07 |
| BNIP3L | -2.8868756 | 4.09147179 | -14.4347887 | 3.6901E-09 | 1.2146E-06 |
| BCL11B | 1.77976061 | 5.37960507 | 14.3286728 | 4.0265E-09 | 1.2587E-06 |
| OCA2 | 1.66096816 | 4.84491764 | 14.2949636 | 4.1401E-09 | 1.2587E-06 |
| ADAR | -0.98340412 | 8.46985341 | -14.0185953 | 5.213E-09 | 1.4351E-06 |
| SAMD8 | -1.03057199 | 7.79544348 | -13.8736277 | 5.8922E-09 | 1.5191E-06 |
| CASC1 | 1.40183266 | 4.79848259 | 13.862841 | 5.9465E-09 | 1.5191E-06 |
| ATXN7L3B | 1.05847432 | 7.04940712 | 13.7988148 | 6.2795E-09 | 1.5578E-06 |
| ABCD3 | -1.09099032 | 6.73349843 | -13.7559488 | 6.5137E-09 | 1.5578E-06 |
| CEND1 | -1.96107303 | 3.0666186 | -13.626231 | 7.2814E-09 | 1.6704E-06 |
| DUSP22 | 1.18709585 | 5.73990306 | 13.4792359 | 8.2706E-09 | 1.8415E-06 |
| DHRS2 | 1.99946634 | 3.86639432 | 13.4046622 | 8.8269E-09 | 1.8505E-06 |
| PSAP | -0.90990069 | 9.23057814 | -13.3924387 | 8.9219E-09 | 1.8505E-06 |
| MTMR2 | -0.9925992 | 7.06076327 | -13.058718 | 1.1989E-08 | 2.267E-06 |
| SLC7A1 | -1.11446657 | 6.9692252 | -12.948504 | 1.3237E-08 | 2.3038E-06 |
| GLS | -1.08038398 | 6.58254974 | -12.732554 | 1.6107E-08 | 2.5593E-06 |
| TMEM64 | -1.07462706 | 6.04340307 | -12.724443 | 1.6227E-08 | 2.5593E-06 |
| TOGARAM1 | 0.99802664 | 6.67085147 | 12.2980431 | 2.4117E-08 | 3.2315E-06 |
| LIN7C | -1.07771539 | 5.72250264 | -12.2184051 | 2.6004E-08 | 3.4237E-06 |
| KCTD3 | 1.00639098 | 6.21038943 | 12.1727938 | 2.7155E-08 | 3.4721E-06 |
| SEC23B | -0.90985678 | 7.38009519 | -12.1676551 | 2.7289E-08 | 3.4721E-06 |
| ACADSB | -1.34672211 | 4.20861604 | -12.020527 | 3.1413E-08 | 3.8357E-06 |
| PREX1 | 1.12565104 | 5.27427154 | 12.0048968 | 3.1889E-08 | 3.8624E-06 |
| CACYBP | -0.91370443 | 6.95569022 | -11.8484325 | 3.7105E-08 | 4.2886E-06 |
| BTBD3 | -1.06297281 | 5.89571905 | -11.8111965 | 3.8476E-08 | 4.3202E-06 |
| SPR | -0.93037587 | 7.05542061 | -11.80842 | 3.8581E-08 | 4.3202E-06 |
| CDCA7 | 0.97093942 | 6.33115321 | 11.8024773 | 3.8805E-08 | 4.3202E-06 |
| PARD3B | 1.09254715 | 5.19968524 | 11.6491693 | 4.5111E-08 | 4.7994E-06 |
| SRP14 | 0.91955209 | 6.79210975 | 11.457191 | 5.4602E-08 | 5.4076E-06 |
| CRYZ | -0.93921013 | 6.11931108 | -11.2379962 | 6.8128E-08 | 6.3313E-06 |
| GTF2A1 | -1.09603201 | 5.98519837 | -11.2133373 | 6.9861E-08 | 6.4107E-06 |
| ZDHHC13 | -1.18245296 | 4.57390971 | -11.1798149 | 7.2293E-08 | 6.5419E-06 |
| INTS14 | 0.98864721 | 6.06209713 | 11.0797206 | 8.0113E-08 | 7.0523E-06 |
| BNIP2 | -0.91561918 | 6.21212198 | -11.0693768 | 8.0971E-08 | 7.0866E-06 |
| FOXA1 | 0.9196457 | 6.16838385 | 10.9732509 | 8.9437E-08 | 7.6941E-06 |
| ERO1A | -1.68469438 | 6.80277097 | -10.9430419 | 9.2289E-08 | 7.8947E-06 |

| | | | | | |
|---|---|---|---|---|---|
| RICTOR | 0.98931935 | 5.14806858 | 10.5386147 | 1.415E-07 | 1.0696E-05 |
| APPL1 | -0.83001668 | 6.62762449 | -10.411588 | 1.6227E-07 | 1.159E-05 |
| HBP1 | 1.06775607 | 5.8805481 | 10.2345448 | 1.9686E-07 | 1.2979E-05 |
| SPECC1L | 0.98540279 | 7.34597254 | 10.2258849 | 1.9874E-07 | 1.3027E-05 |
| OPA1 | -0.77045637 | 7.21290993 | -10.1794071 | 2.0918E-07 | 1.3532E-05 |
| TTC6 | 1.31206785 | 3.49237786 | 10.0548886 | 2.4016E-07 | 1.4903E-05 |
| CDK6 | 0.94337192 | 7.94364534 | 9.99437 | 2.5696E-07 | 1.5669E-05 |
| WDR20 | 0.82116001 | 6.35977059 | 9.99185298 | 2.5768E-07 | 1.5669E-05 |
| EIF2S1 | -0.70228463 | 8.14937379 | -9.72889034 | 3.4702E-07 | 1.9034E-05 |
| LINC01852 | 0.98350226 | 5.07636917 | 9.72752703 | 3.4756E-07 | 1.9034E-05 |
| UFL1 | 0.83452035 | 5.77532162 | 9.71282057 | 3.5346E-07 | 1.9114E-05 |
| KIDINS220 | -0.79494562 | 6.67061425 | -9.64012558 | 3.8425E-07 | 2.0414E-05 |
| ISM2 | -1.66685961 | 3.23242201 | -9.56234061 | 4.204E-07 | 2.1577E-05 |
| FANCC | -0.77295005 | 6.78476209 | -9.5411986 | 4.3084E-07 | 2.1748E-05 |
| SLC7A11 | -0.65210415 | 8.45397543 | -9.54103694 | 4.3092E-07 | 2.1748E-05 |
| MOK | 1.05152205 | 4.58897035 | 9.5217625 | 4.4068E-07 | 2.2021E-05 |
| ACTR6 | -0.9795279 | 4.76260201 | -9.50381307 | 4.4999E-07 | 2.2412E-05 |
| MYB | 1.19691801 | 3.53201446 | 9.49011388 | 4.5723E-07 | 2.2624E-05 |
| WASHC4 | -0.76140637 | 6.94513626 | -9.46850582 | 4.6891E-07 | 2.3051E-05 |
| SH2B3 | -0.88542889 | 5.23832507 | -9.43921258 | 4.8526E-07 | 2.3625E-05 |
| GPM6A | 0.67376633 | 7.67689418 | 9.43164413 | 4.8958E-07 | 2.3759E-05 |
| RGS5 | -0.73590447 | 6.8752768 | -9.25503808 | 6.0306E-07 | 2.8009E-05 |
| INSM2 | 2.11473089 | 1.19745617 | 9.22558208 | 6.2459E-07 | 2.8657E-05 |
| LCA5 | -0.75764701 | 6.24206544 | -9.19585975 | 6.4714E-07 | 2.9513E-05 |
| RASSF8 | -0.7157123 | 6.74657025 | -9.16118661 | 6.7455E-07 | 3.0075E-05 |
| BICC1 | -0.75579334 | 6.44168744 | -9.14325998 | 6.8921E-07 | 3.0424E-05 |
| MAP3K7 | -0.72627411 | 6.73285993 | -9.09785612 | 7.2788E-07 | 3.1852E-05 |
| HIVEP2 | 0.98820849 | 5.09877701 | 9.03206202 | 7.881E-07 | 3.3899E-05 |
| NIN | -0.73137547 | 7.76095112 | -9.02130462 | 7.9844E-07 | 3.4041E-05 |
| CPSF2 | -0.61179055 | 8.30485266 | -8.93419462 | 8.8772E-07 | 3.6624E-05 |
| ZC3H15 | -0.64773595 | 8.1208712 | -8.91031121 | 9.1402E-07 | 3.7202E-05 |
| LSM11 | -0.87627941 | 5.11340133 | -8.90608494 | 9.1876E-07 | 3.7221E-05 |
| DLX1 | 0.66400569 | 7.39988931 | 8.86192531 | 9.6989E-07 | 3.8952E-05 |
| ZCCHC14 | 0.69291695 | 6.77716074 | 8.80052757 | 1.0461E-06 | 4.1033E-05 |
| SPATS2L | -0.8531468 | 4.84365123 | -8.71146461 | 1.1682E-06 | 4.478E-05 |
| PTBP2 | 0.80947045 | 5.27766523 | 8.66969408 | 1.2307E-06 | 4.6216E-05 |
| GFPT2 | -1.08043681 | 3.8169059 | -8.66807004 | 1.2332E-06 | 4.6216E-05 |
| EXOSC6 | -0.7099963 | 6.46604758 | -8.61510922 | 1.3177E-06 | 4.8129E-05 |
| PRKD1 | 0.78108177 | 6.17504804 | 8.61126764 | 1.3241E-06 | 4.8191E-05 |
| CCND2 | -0.51410398 | 10.370687 | -8.59954154 | 1.3437E-06 | 4.8672E-05 |
| PBX1 | 1.24748247 | 3.04423675 | 8.56367405 | 1.4057E-06 | 5.0556E-05 |
| MNS1 | 0.94511169 | 4.25423017 | 8.54288549 | 1.4431E-06 | 5.1654E-05 |
| PGF | -1.0561914 | 3.83099495 | -8.50656034 | 1.5109E-06 | 5.345E-05 |
| SNRNP48 | 0.76346308 | 5.36402663 | 8.4810142 | 1.5606E-06 | 5.4572E-05 |
| ZDHHC23 | -0.99584126 | 4.3456593 | -8.46516928 | 1.5924E-06 | 5.492E-05 |
| CNTNAP4 | 0.69236794 | 7.23709098 | 8.447332 | 1.6289E-06 | 5.5868E-05 |
| ANO6 | 0.67871411 | 7.23015739 | 8.43075067 | 1.6637E-06 | 5.6697E-05 |
| DUSP5 | -1.29944019 | 2.9253224 | -8.35801275 | 1.8258E-06 | 5.9967E-05 |
| GSE1 | 0.64573233 | 7.04620745 | 8.35127906 | 1.8417E-06 | 6.0356E-05 |

| | | | | | |
|---|---|---|---|---|---|
| ZHX3 | -0.75397792 | 5.45061316 | -8.31859816 | 1.9207E-06 | 6.2209E-05 |
| KCTD18 | -0.93239375 | 4.07501505 | -8.25678292 | 2.0801E-06 | 6.6446E-05 |
| SGMS1 | -0.70190813 | 5.98351321 | -8.23994279 | 2.126E-06 | 6.7626E-05 |
| ALDH1A1 | -0.6782434 | 8.4723853 | -8.192284 | 2.2617E-06 | 7.0627E-05 |
| SESN3 | 0.78990599 | 4.78761336 | 8.15788367 | 2.3654E-06 | 7.2353E-05 |
| ETFRF1 | 0.77315767 | 4.68126862 | 8.14520126 | 2.4049E-06 | 7.3282E-05 |
| PPP2R5C | 0.58778145 | 7.67018435 | 8.13413419 | 2.44E-06 | 7.33E-05 |
| DNAH7 | 0.77544604 | 5.00179047 | 8.11188429 | 2.5121E-06 | 7.4291E-05 |
| BCL6 | 1.04690656 | 3.5490624 | 8.09340999 | 2.5737E-06 | 7.5373E-05 |
| OLFM3 | 0.58475488 | 7.7207739 | 8.08394451 | 2.6058E-06 | 7.5875E-05 |
| CHAC2 | -0.94739926 | 4.05981504 | -8.08199433 | 2.6125E-06 | 7.5924E-05 |
| STARD4 | -0.62546102 | 7.28312248 | -8.05411638 | 2.7101E-06 | 7.8061E-05 |
| OPRL1 | -1.08806272 | 4.21494581 | -8.02416219 | 2.8193E-06 | 7.9788E-05 |
| ERRFI1 | 0.75260133 | 8.68034164 | 8.02256971 | 2.8252E-06 | 7.9807E-05 |
| SRBD1 | 0.73842587 | 6.21266407 | 7.95274466 | 3.099E-06 | 8.5937E-05 |
| PRKACB | -0.93729493 | 3.6958924 | -7.86146045 | 3.5003E-06 | 9.3635E-05 |
| CUBN | 0.78690079 | 5.60798215 | 7.84656426 | 3.5708E-06 | 9.5187E-05 |
| EEA1 | 0.71130084 | 6.09831844 | 7.83087455 | 3.6468E-06 | 9.6871E-05 |
| RARB | 0.61810339 | 7.30220343 | 7.82471504 | 3.6771E-06 | 9.7334E-05 |
| HAPLN3 | -1.17634856 | 3.26835482 | -7.80904696 | 3.7554E-06 | 9.8476E-05 |
| CRIM1 | -0.70341326 | 6.34702744 | -7.80891786 | 3.756E-06 | 9.8476E-05 |
| GPX8 | -0.69428417 | 5.59566712 | -7.80851174 | 3.7581E-06 | 9.8476E-05 |
| PARD6B | -0.81940023 | 4.38545434 | -7.80546024 | 3.7736E-06 | 9.868E-05 |
| CHD6 | 0.65514593 | 6.47975622 | 7.79477168 | 3.8283E-06 | 9.9253E-05 |
| COCH | -0.67478084 | 5.7664326 | -7.78821708 | 3.8622E-06 | 9.9622E-05 |
| SLITRK5 | 0.66842758 | 6.070538 | 7.72123891 | 4.2283E-06 | 0.00010688 |
| PSD | 0.86801993 | 4.64531787 | 7.70446819 | 4.3256E-06 | 0.00010849 |
| OLA1 | -0.59603284 | 6.95118935 | -7.70408855 | 4.3278E-06 | 0.00010849 |
| ALKBH8 | -0.72585621 | 5.47749746 | -7.70110736 | 4.3454E-06 | 0.00010875 |
| ATXN1 | 0.71872562 | 7.68090284 | 7.67388438 | 4.5093E-06 | 0.00011211 |
| DSTN | -0.62834432 | 6.93890888 | -7.65921205 | 4.6003E-06 | 0.00011419 |
| KDM1B | -0.72971073 | 5.71185943 | -7.6391976 | 4.7277E-06 | 0.00011649 |
| CACNA2D1 | 0.70919412 | 5.0201647 | 7.61934912 | 4.8577E-06 | 0.00011883 |
| OGA | 0.54920307 | 7.69928682 | 7.61324642 | 4.8985E-06 | 0.00011919 |
| PYGO1 | -0.72238026 | 4.97920945 | -7.58746247 | 5.0746E-06 | 0.00012213 |
| TPD52L1 | -0.68198441 | 6.18664933 | -7.58100086 | 5.1198E-06 | 0.00012234 |
| DESI2 | -0.58533196 | 7.49786651 | -7.56167638 | 5.2576E-06 | 0.00012497 |
| CDH12 | 0.80941169 | 4.34986959 | 7.54268023 | 5.3969E-06 | 0.00012708 |
| LTBP1 | 0.601792 | 7.14395757 | 7.52309029 | 5.5446E-06 | 0.00012955 |
| VIT | 1.06512193 | 4.2129112 | 7.50818041 | 5.6599E-06 | 0.00013125 |
| SSR1 | -0.55723977 | 7.3973734 | -7.48843564 | 5.8166E-06 | 0.00013405 |
| PHACTR4 | 0.74796796 | 5.39661038 | 7.47910815 | 5.8922E-06 | 0.00013538 |
| PMEPA1 | -1.09714801 | 2.99766436 | -7.47737349 | 5.9064E-06 | 0.00013546 |
| FAP | 0.64045747 | 5.85664632 | 7.45244802 | 6.1142E-06 | 0.00013858 |
| DICER1 | 0.70009326 | 8.23077625 | 7.4002019 | 6.5753E-06 | 0.00014533 |
| SCYL3 | 0.91644148 | 4.32649287 | 7.38676694 | 6.6998E-06 | 0.00014702 |
| GABPB2 | 0.78609069 | 4.42831203 | 7.38401366 | 6.7256E-06 | 0.00014733 |
| DYNC1I1 | 0.6529161 | 5.40591961 | 7.2568882 | 8.0408E-06 | 0.00017041 |
| CPEB2 | 1.11641791 | 2.58162183 | 7.22894562 | 8.365E-06 | 0.00017494 |

| | | | | | |
|---|---|---|---|---|---|
| ARL4A | 0.54441847 | 7.56548233 | 7.22290885 | 8.4369E-06 | 0.00017547 |
| DNAJC10 | 0.54572636 | 7.08699127 | 7.22004304 | 8.4712E-06 | 0.00017547 |
| SCFD1 | 0.58536285 | 6.69485191 | 7.22001605 | 8.4715E-06 | 0.00017547 |
| TTC8 | 0.57503593 | 6.87244998 | 7.1889196 | 8.8539E-06 | 0.00018067 |
| ARID4A | 0.60928054 | 5.87265219 | 7.13811395 | 9.5185E-06 | 0.00019082 |
| MEIS2 | 0.52844548 | 7.86908741 | 7.11812249 | 9.7944E-06 | 0.0001942 |
| GBE1 | -0.71624337 | 7.31467102 | -7.11506292 | 9.8374E-06 | 0.0001942 |
| NFIB | -0.76631931 | 4.2243764 | -7.1103777 | 9.9036E-06 | 0.00019525 |
| ESYT1 | -0.51607256 | 7.84158183 | -7.09374422 | 1.0142E-05 | 0.00019841 |
| IFIT1 | -1.56453336 | 1.76662831 | -7.09154845 | 1.0174E-05 | 0.00019878 |
| RASSF2 | -0.71531962 | 4.74329408 | -7.06823957 | 1.0521E-05 | 0.00020447 |
| KLF10 | 0.81101867 | 7.27678413 | 7.04960853 | 1.0806E-05 | 0.00020858 |
| LNPK | -0.67887825 | 5.99253823 | -7.0491 | 1.0814E-05 | 0.00020858 |
| CEP95 | 0.58932384 | 6.4578532 | 7.04767322 | 1.0836E-05 | 0.00020875 |
| RSU1 | -0.55703817 | 6.84083999 | -7.04044375 | 1.095E-05 | 0.00021013 |
| CREG1 | -0.72140777 | 4.75095618 | -7.03733315 | 1.0999E-05 | 0.00021054 |
| SLC9A2 | -0.76850084 | 4.11620139 | -7.00160447 | 1.1581E-05 | 0.00021795 |
| TRMT11 | 0.67969996 | 5.61997184 | 6.99655416 | 1.1666E-05 | 0.00021915 |
| CBS | -0.81610365 | 4.38814019 | -6.99058389 | 1.1767E-05 | 0.00022073 |
| NID2 | -1.0903527 | 2.94768077 | -6.98776272 | 1.1815E-05 | 0.00022073 |
| BEGAIN | 0.72386476 | 4.80861905 | 6.98732423 | 1.1823E-05 | 0.00022073 |
| SLC24A4 | -1.01467748 | 2.94743448 | -6.96864073 | 1.2147E-05 | 0.00022484 |
| HINT1 | -0.59557792 | 6.44942616 | -6.9572345 | 1.235E-05 | 0.0002272 |
| SLC30A8 | -0.58750964 | 6.05268369 | -6.92528698 | 1.2937E-05 | 0.00023571 |
| TMEM33 | -0.52304976 | 7.0781348 | -6.92192026 | 1.3E-05 | 0.00023573 |
| DAP | -0.76202643 | 4.1829187 | -6.89841626 | 1.3453E-05 | 0.00024106 |
| PEG3 | 0.76971196 | 6.53663409 | 6.89630203 | 1.3495E-05 | 0.00024141 |
| DPP4 | -0.91535154 | 3.67219607 | -6.89117309 | 1.3596E-05 | 0.00024247 |
| HNRNPLL | 0.5330273 | 7.04411021 | 6.84888642 | 1.4464E-05 | 0.00025435 |
| B3GALT1 | 1.16004554 | 2.24198951 | 6.81806301 | 1.5133E-05 | 0.00026306 |
| SSPN | 1.08478168 | 2.44289058 | 6.81599642 | 1.5179E-05 | 0.00026352 |
| EMB | -0.65039975 | 5.05161873 | -6.80737159 | 1.5373E-05 | 0.00026577 |
| CFAP54 | 1.13652737 | 2.42137649 | 6.80287831 | 1.5475E-05 | 0.00026656 |
| NEMF | 0.53086071 | 6.90988691 | 6.77591007 | 1.6102E-05 | 0.0002731 |
| WASL | 0.5533286 | 6.52653848 | 6.75804203 | 1.6532E-05 | 0.00027844 |
| KCNJ11 | -0.81831587 | 3.86656403 | -6.74370092 | 1.6887E-05 | 0.00028097 |
| KRIT1 | -0.58383621 | 6.29860354 | -6.72961355 | 1.7242E-05 | 0.00028439 |
| DPYD | 0.87863045 | 3.52179739 | 6.69246903 | 1.8219E-05 | 0.00029472 |
| DHFR | 0.65026932 | 5.64378425 | 6.67499501 | 1.8699E-05 | 0.0002996 |
| RTN4R | -0.68238332 | 4.69926589 | -6.66887908 | 1.887E-05 | 0.00030138 |
| PELO | -0.55986191 | 6.08002329 | -6.64859864 | 1.9449E-05 | 0.00030739 |
| PBLD | 0.72403705 | 4.36244526 | 6.59699811 | 2.1009E-05 | 0.00032626 |
| STK3 | -0.54586682 | 6.23230802 | -6.57554629 | 2.1697E-05 | 0.0003325 |
| FBXO33 | 0.54655122 | 6.09382109 | 6.56261208 | 2.2122E-05 | 0.00033698 |
| NR2F1 | 0.51735324 | 7.164505 | 6.55307746 | 2.2442E-05 | 0.00034082 |
| GPATCH2 | 0.61598095 | 5.34726191 | 6.53438916 | 2.3083E-05 | 0.00034603 |
| SLC30A1 | 0.51760482 | 6.54463567 | 6.53210367 | 2.3162E-05 | 0.00034689 |
| OSBPL8 | 0.51360622 | 7.53124137 | 6.52484106 | 2.3417E-05 | 0.00035036 |
| ATG2B | 0.5763948 | 5.98663736 | 6.50898264 | 2.3985E-05 | 0.00035603 |

| | | | | | |
|---|---|---|---|---|---|
| ETV1 | 0.55250228 | 6.26023202 | 6.49448049 | 2.4516E-05 | 0.00036074 |
| LRFN5 | 0.53699455 | 6.64913682 | 6.48492225 | 2.4874E-05 | 0.00036493 |
| SLC25A21-AS1 | 1.16512395 | 1.78916794 | 6.45786004 | 2.5915E-05 | 0.00037405 |
| TLL2 | -0.86066494 | 3.36046558 | -6.45641806 | 2.5972E-05 | 0.00037427 |
| EML2 | -0.71999076 | 5.92550939 | -6.43130148 | 2.6982E-05 | 0.00038361 |
| GCG | -0.77709903 | 3.72712965 | -6.40833377 | 2.7943E-05 | 0.00039284 |
| PDPR | 0.54316447 | 5.98886971 | 6.40186844 | 2.822E-05 | 0.00039636 |
| GPR135 | -0.81040562 | 3.45677802 | -6.39776766 | 2.8397E-05 | 0.00039755 |
| ANK1 | 1.42292606 | 1.26655624 | 6.35500169 | 3.0315E-05 | 0.00041656 |
| SLCO4A1 | -0.57655694 | 5.79165375 | -6.35401333 | 3.0361E-05 | 0.00041677 |
| XPR1 | -0.50469551 | 6.75646392 | -6.33884727 | 3.1075E-05 | 0.0004235 |
| ARPP21 | 1.21633464 | 1.90138846 | 6.28958674 | 3.352E-05 | 0.00045275 |
| KIAA0825 | 0.95445705 | 2.52986899 | 6.284873 | 3.3764E-05 | 0.00045442 |
| ZSWIM2 | 0.78127095 | 3.47564496 | 6.28032489 | 3.4002E-05 | 0.00045722 |
| ADIPOR1 | 0.5166921 | 6.93002983 | 6.26020025 | 3.5075E-05 | 0.00046831 |
| ZNF25 | -0.60526523 | 5.52750745 | -6.25415579 | 3.5404E-05 | 0.0004708 |
| KDM3A | 0.71191586 | 6.53036976 | 6.24209998 | 3.607E-05 | 0.00047531 |
| MICU3 | 0.67503763 | 4.05123376 | 6.23909102 | 3.6238E-05 | 0.0004763 |
| ODF2L | -0.56810842 | 5.39977736 | -6.17685307 | 3.9913E-05 | 0.00050628 |
| GLIDR | 1.13284712 | 2.21778529 | 6.16653524 | 4.0559E-05 | 0.00051133 |
| EGR2 | 1.47338519 | 1.20324484 | 6.15267553 | 4.1445E-05 | 0.00052009 |
| PAIP2B | 1.03865326 | 2.45126895 | 6.15003217 | 4.1616E-05 | 0.00052161 |
| DPH6 | 0.64846552 | 4.23726119 | 6.14017193 | 4.2261E-05 | 0.00052839 |
| DLX2 | 0.58710359 | 6.54756972 | 6.13826827 | 4.2387E-05 | 0.00052953 |
| BBS10 | 0.56470414 | 5.22025141 | 6.13703111 | 4.2469E-05 | 0.00053011 |
| VPS50 | 0.525972 | 5.87272933 | 6.13537697 | 4.2579E-05 | 0.00053105 |
| PEX1 | 0.52027602 | 5.93611869 | 6.13413902 | 4.2661E-05 | 0.0005312 |
| KCNJ12 | -0.59749035 | 5.62550702 | -6.08405998 | 4.6141E-05 | 0.00056352 |
| CEP162 | 0.63866778 | 4.360564 | 6.08341566 | 4.6187E-05 | 0.00056352 |
| ZNF32 | -0.57637765 | 5.22306628 | -6.07939669 | 4.648E-05 | 0.00056571 |
| DDIT3 | 0.64646524 | 6.2383835 | 6.07438107 | 4.6847E-05 | 0.00056927 |
| MKRN3 | 0.5340711 | 5.87186355 | 6.03609192 | 4.9755E-05 | 0.00059506 |
| C1GALT1 | -0.64279316 | 5.17859189 | -6.02882352 | 5.0329E-05 | 0.00060002 |
| BEND6 | 0.97302115 | 2.34546997 | 6.01851663 | 5.1154E-05 | 0.00060746 |
| IRF4 | -0.88939122 | 2.63010989 | -5.98639674 | 5.3817E-05 | 0.00063117 |
| PCDH15 | 0.70878981 | 4.29668855 | 5.97326014 | 5.4948E-05 | 0.00064047 |
| ARID5B | 0.6311335 | 6.21704649 | 5.96898409 | 5.5322E-05 | 0.00064224 |
| ZBTB18 | 0.63771734 | 4.25023951 | 5.95793462 | 5.63E-05 | 0.00064972 |
| CTTNBP2NL | -0.60418323 | 4.74300353 | -5.95031209 | 5.6985E-05 | 0.00065663 |
| AP4S1 | 0.66299408 | 4.49130423 | 5.93035703 | 5.8821E-05 | 0.00067216 |
| RASGEF1A | -0.56831109 | 5.04168773 | -5.90360573 | 6.1381E-05 | 0.00069579 |
| NLGN1 | 0.57800178 | 4.73933616 | 5.88231219 | 6.3502E-05 | 0.00071221 |
| DAOA-AS1 | -0.577403 | 4.7644891 | -5.87924109 | 6.3814E-05 | 0.00071307 |
| LRRIQ3 | 0.79954572 | 2.92208653 | 5.87240604 | 6.4515E-05 | 0.00071749 |
| NAA20 | -0.51146427 | 5.80677941 | -5.85776456 | 6.6044E-05 | 0.00073204 |
| ELOVL4 | -0.55704867 | 5.33697685 | -5.83621819 | 6.8363E-05 | 0.00074971 |
| GCA | -0.67217195 | 4.73842571 | -5.83146417 | 6.8887E-05 | 0.00075362 |
| GBF1 | 0.50980325 | 7.7188996 | 5.80433524 | 7.1954E-05 | 0.00077763 |
| OGFRL1 | -0.57334738 | 4.59315261 | -5.76482037 | 7.6683E-05 | 0.00081707 |

| | | | | | |
|---|---|---|---|---|---|
| ID2-AS1 | 0.85318749 | 2.94393177 | 5.76365228 | 7.6828E-05 | 0.00081746 |
| KRBOX4 | -0.7031795 | 3.68243388 | -5.74538322 | 7.9129E-05 | 0.00083607 |
| LRRIQ1 | 1.17527781 | 1.2832586 | 5.74135059 | 7.9646E-05 | 0.00083936 |
| CDK18 | -0.81362443 | 3.45009058 | -5.74122522 | 7.9662E-05 | 0.00083936 |
| PNPLA8 | 0.55227847 | 4.94364089 | 5.73858776 | 8.0003E-05 | 0.00084178 |
| EIF3J-DT | 0.61921219 | 4.35720125 | 5.73779676 | 8.0105E-05 | 0.00084227 |
| LINC00648 | 0.95680165 | 2.79792259 | 5.72790537 | 8.1397E-05 | 0.00085207 |
| UNC5B | -0.7034353 | 4.86421312 | -5.709566 | 8.3852E-05 | 0.00087019 |
| ALG10B | 0.54522354 | 4.70359993 | 5.70476788 | 8.4507E-05 | 0.00087399 |
| USP40 | 0.50486682 | 6.00617267 | 5.68103229 | 8.7827E-05 | 0.0008973 |
| NFIC | -0.5891707 | 4.54427774 | -5.67812068 | 8.8244E-05 | 0.00090034 |
| SLC4A3 | -0.73266006 | 3.46049777 | -5.64047149 | 9.3823E-05 | 0.00094705 |
| BCL11A | 0.62672827 | 4.0512803 | 5.6327954 | 9.5006E-05 | 0.00095707 |
| MIR4500HG | 1.15097214 | 1.30384991 | 5.62688808 | 9.5926E-05 | 0.00096506 |
| PIK3C2G | 0.73690258 | 3.18532279 | 5.58268592 | 0.00010312 | 0.0010165 |
| PPP3CB | -0.50243217 | 5.45489356 | -5.57046773 | 0.00010521 | 0.00102638 |
| DLX6-AS1 | 0.5126059 | 6.46227718 | 5.56003728 | 0.00010702 | 0.00103802 |
| LPXN | 0.96769878 | 2.06595745 | 5.52324575 | 0.0001137 | 0.00108863 |
| CRISPLD1 | 0.5640843 | 4.81532452 | 5.52046103 | 0.00011422 | 0.00109252 |
| PDK1 | -1.55821837 | 3.82785831 | -5.51101024 | 0.00011602 | 0.00110188 |
| FAT4 | 0.5622205 | 5.71308148 | 5.50087395 | 0.00011797 | 0.00111431 |
| CCDC148 | 1.34758454 | 0.76338091 | 5.49491153 | 0.00011914 | 0.00112323 |
| SLC39A8 | -0.55275227 | 4.80313721 | -5.45882852 | 0.00012647 | 0.0011683 |
| FAM172A | 0.55023816 | 5.03608215 | 5.45397409 | 0.00012749 | 0.00117415 |
| C1orf21 | 0.63556217 | 3.8982216 | 5.42585095 | 0.00013358 | 0.00121108 |
| ALX1 | 0.65531152 | 3.76379827 | 5.4142553 | 0.00013618 | 0.00122838 |
| MANEA | 0.56737074 | 4.46617481 | 5.40655366 | 0.00013793 | 0.00123869 |
| NTS | -1.23300235 | 1.10255722 | -5.39587807 | 0.00014041 | 0.00125569 |
| RET | -1.21994641 | 1.68380022 | -5.36280497 | 0.00014837 | 0.00130715 |
| CDC37L1 | 0.52294239 | 4.80790043 | 5.36265273 | 0.0001484 | 0.00130715 |
| COL26A1 | 1.29464258 | 1.95727743 | 5.35956101 | 0.00014917 | 0.00131239 |
| ZMIZ1 | 0.63216802 | 3.84619004 | 5.35284342 | 0.00015086 | 0.00132106 |
| KLF12 | 0.67608606 | 3.5442634 | 5.32794268 | 0.00015727 | 0.00136229 |
| OTX1 | 0.68922847 | 3.42308925 | 5.30624319 | 0.0001631 | 0.00139994 |
| OSER1-DT | 1.0109051 | 1.89920367 | 5.29313372 | 0.00016673 | 0.00142462 |
| ING3 | 0.5343388 | 4.71891384 | 5.27420739 | 0.00017212 | 0.00145345 |
| BMP5 | -0.55744907 | 4.32056292 | -5.24014224 | 0.00018228 | 0.00152064 |
| THUMPD2 | 0.54433425 | 5.4222841 | 5.23647626 | 0.00018341 | 0.00152671 |
| FAM131C | -1.16387528 | 1.38665126 | -5.23246206 | 0.00018466 | 0.00153624 |
| MYO15B | -0.88807446 | 4.53872146 | -5.20910098 | 0.0001921 | 0.00157986 |
| SPAG1 | -0.53671106 | 4.78082249 | -5.18404486 | 0.00020042 | 0.00162536 |
| IRF1 | 0.61542938 | 3.98022964 | 5.18080729 | 0.00020152 | 0.00162994 |
| NBPF4 | 0.92028725 | 1.98111667 | 5.10365799 | 0.00022977 | 0.00180632 |
| MYOF | -0.9803951 | 2.20221684 | -5.09797028 | 0.00023201 | 0.00182016 |
| DDX59 | 0.5010154 | 4.97591263 | 5.09763728 | 0.00023214 | 0.00182025 |
| UNC93B1 | -0.75264796 | 3.34223558 | -5.05308875 | 0.00025052 | 0.00192835 |
| ANGPTL1 | 0.70610208 | 3.51244381 | 5.05149579 | 0.0002512 | 0.00193165 |
| PLEKHM3 | 0.75029426 | 2.86939191 | 5.02076943 | 0.00026479 | 0.00201367 |
| FGFBP3 | 0.72319697 | 2.87422011 | 4.98413824 | 0.00028201 | 0.00211806 |

| | | | | | |
|---|---|---|---|---|---|
| SLC35D3 | -0.78819558 | 2.67309878 | -4.96048586 | 0.00029374 | 0.00219199 |
| AKAP6 | 0.65802096 | 3.4261811 | 4.94827485 | 0.0003 | 0.00222661 |
| PPP1R16B | -0.5684321 | 4.13574817 | -4.94478373 | 0.00030181 | 0.00223517 |
| GLRA3 | 1.17005987 | 0.86890348 | 4.93241963 | 0.00030833 | 0.00227396 |
| KCNMB4 | 0.55979426 | 4.00947174 | 4.93014344 | 0.00030955 | 0.0022796 |
| RERGL | 0.76858833 | 2.50922532 | 4.92661214 | 0.00031144 | 0.00228942 |
| DCN | -0.50513882 | 4.41044342 | -4.9159311 | 0.00031725 | 0.00232055 |
| ZDHHC11 | -0.73245403 | 2.83295239 | -4.89104433 | 0.00033123 | 0.00239619 |
| BFSP1 | 0.85305453 | 2.25140509 | 4.87565124 | 0.0003402 | 0.00243893 |
| CFAP69 | 0.68715789 | 3.12823641 | 4.8603232 | 0.00034939 | 0.00247806 |
| MATN2 | -0.67960938 | 3.99967787 | -4.85608453 | 0.00035197 | 0.00249028 |
| LINC01140 | -0.87834684 | 2.20527037 | -4.83464531 | 0.00036535 | 0.00255748 |
| AUTS2 | 0.58646914 | 4.16264511 | 4.81388825 | 0.00037882 | 0.00263029 |
| LINC00858 | 0.86471951 | 2.15064037 | 4.80970769 | 0.00038159 | 0.00264667 |
| TTI2 | 0.53440673 | 4.29699001 | 4.80596711 | 0.00038409 | 0.00265914 |
| CEBPB-AS1 | 1.13590601 | 0.90326893 | 4.80269168 | 0.00038629 | 0.0026695 |
| ANAPC15 | -0.64171057 | 3.27967938 | -4.80221923 | 0.00038661 | 0.00267049 |
| GABRG3 | 0.65587114 | 3.09634519 | 4.75562662 | 0.00041945 | 0.00284408 |
| GABRG1 | -1.15880838 | 0.65064678 | -4.75461501 | 0.00042019 | 0.00284785 |
| TASP1 | -0.50880646 | 4.31218102 | -4.74924849 | 0.00042416 | 0.00286579 |
| KLLN | 0.65219379 | 3.15014844 | 4.74441032 | 0.00042778 | 0.00288506 |
| VIM-AS1 | 0.80990849 | 2.01394956 | 4.68888402 | 0.00047163 | 0.00311015 |
| DYNC2LI1 | 0.50417151 | 4.22558932 | 4.68627381 | 0.0004738 | 0.00312176 |
| KCNC2 | 0.64579331 | 3.09973449 | 4.64050803 | 0.00051365 | 0.0033193 |
| OLFML2B | -1.30880914 | 0.53226102 | -4.63752107 | 0.00051637 | 0.00332766 |
| SYNE2 | 0.52614312 | 8.00984902 | 4.63402262 | 0.00051957 | 0.00334126 |
| GPCPD1 | 0.52691021 | 3.91410379 | 4.62375686 | 0.00052909 | 0.00339364 |
| LINC02197 | 1.14662661 | 0.51775426 | 4.60935813 | 0.00054275 | 0.00344561 |
| TMEM121 | -0.54190874 | 4.02923748 | -4.59569919 | 0.00055605 | 0.00351529 |
| LOXL3 | 0.51066459 | 4.11223895 | 4.58678327 | 0.00056491 | 0.00355454 |
| ENPP2 | 1.42762172 | 0.10161852 | 4.5458837 | 0.00060749 | 0.00375122 |
| SCN3A | -0.56971507 | 4.01760617 | -4.54437034 | 0.00060913 | 0.0037552 |
| DDIT4 | -1.29930393 | 4.91730096 | -4.53269271 | 0.00062192 | 0.0038139 |
| LINC01793 | -0.78570842 | 2.02055372 | -4.53111233 | 0.00062367 | 0.0038231 |
| DNAJB4 | -0.59029261 | 3.24328148 | -4.52250948 | 0.00063331 | 0.00386338 |
| DNAJC6 | -0.65879139 | 3.051168 | -4.48528355 | 0.00067681 | 0.0040583 |
| DISP1 | 0.61660368 | 3.1981767 | 4.48507844 | 0.00067705 | 0.0040583 |
| TCF7L2 | 0.65724912 | 2.88197001 | 4.47069984 | 0.00069468 | 0.00413692 |
| WDPCP | 0.50328804 | 4.24494585 | 4.44550375 | 0.00072673 | 0.00429583 |
| SYT16 | 1.48554619 | -0.42643453 | 4.42056158 | 0.00075997 | 0.00444446 |
| MAFB | 1.40145967 | -0.48691532 | 4.41252628 | 0.00077101 | 0.00449687 |
| MPP7 | -0.7777408 | 1.93750591 | -4.37383344 | 0.00082655 | 0.00473174 |
| ADAMTS2 | -0.53582878 | 4.74804588 | -4.35360733 | 0.00085722 | 0.00485291 |
| SP9 | 0.54396662 | 3.89397509 | 4.31231647 | 0.00092355 | 0.00514476 |
| SLCO3A1 | -0.80672773 | 1.94885041 | -4.29259218 | 0.00095709 | 0.00528441 |
| SPATA17 | 1.21666532 | 0.07376881 | 4.29062358 | 0.0009605 | 0.00529992 |
| JPH1 | -1.12833553 | 0.34075997 | -4.28205526 | 0.00097552 | 0.00535739 |
| SEMA7A | -0.7206775 | 2.73551703 | -4.28016547 | 0.00097886 | 0.0053738 |
| NR2F2 | -0.51627518 | 3.74782046 | -4.25667488 | 0.00102143 | 0.00555739 |

| | | | | | |
|---|---|---|---|---|---|
| HSD11B2 | -1.00945728 | 1.01909646 | -4.22500366 | 0.00108188 | 0.00579646 |
| SPATA7 | 0.50910153 | 3.7456858 | 4.21356988 | 0.0011046 | 0.0058828 |
| SYBU | -1.17662132 | 0.27027243 | -4.18722812 | 0.00115885 | 0.0061046 |
| IL1R1 | -1.20060543 | 0.90245945 | -4.15766406 | 0.00122301 | 0.0063569 |
| TTC32 | 0.69855184 | 2.73403775 | 4.15319301 | 0.00123303 | 0.00639799 |
| HOOK1 | 0.64350502 | 2.86014603 | 4.08913662 | 0.00138624 | 0.00694478 |
| NPAS3 | 0.68839021 | 2.43596737 | 4.05816423 | 0.00146723 | 0.00725517 |
| NFASC | -0.54560439 | 3.36723457 | -4.04833801 | 0.00149393 | 0.00734142 |
| TFPI2 | -0.81514403 | 2.78924123 | -4.03307086 | 0.00153641 | 0.00750172 |
| LINC01122 | 0.52936093 | 3.28700637 | 4.02247804 | 0.00156662 | 0.00762462 |
| CAPS2 | 0.87381622 | 1.7746624 | 4.01912238 | 0.00157631 | 0.00765458 |
| PRL | -0.7299066 | 1.79867027 | -3.97665552 | 0.00170449 | 0.00812328 |
| TENT5B | -1.14686737 | 0.37379286 | -3.95500653 | 0.00177391 | 0.00838042 |
| PPM1J | -0.73847866 | 2.18111892 | -3.9549961 | 0.00177394 | 0.00838042 |
| BNIP3 | -0.7694295 | 6.11098714 | -3.95137566 | 0.00178583 | 0.00841821 |
| PTPRO | 1.00855661 | 0.65040348 | 3.90709682 | 0.00193805 | 0.00896002 |
| ASAH2 | -0.75449154 | 1.66996193 | -3.9045039 | 0.00194737 | 0.00899211 |
| ANKRD29 | -0.9487725 | 0.95813014 | -3.89152819 | 0.00199469 | 0.0091631 |
| LINC02506 | 0.61305146 | 2.56521419 | 3.8907317 | 0.00199763 | 0.00917384 |
| PRRG4 | -0.91131154 | 0.87998573 | -3.87105155 | 0.00207177 | 0.00944557 |
| AFAP1L2 | -0.84170132 | 1.43441662 | -3.86758515 | 0.00208512 | 0.00949785 |
| MCF2L | -0.54233397 | 3.13708026 | -3.83860388 | 0.00220024 | 0.00989715 |
| KANK4 | 0.77946725 | 1.77657102 | 3.83811551 | 0.00220223 | 0.00990024 |
| FOXO1 | 0.55202498 | 3.6960752 | 3.82151368 | 0.00227116 | 0.0101021 |
| CDH8 | 0.91400475 | 0.89550116 | 3.79396159 | 0.00239045 | 0.0105156 |
| FBXO48 | 0.92666879 | 1.14333928 | 3.78346389 | 0.00243758 | 0.01067612 |
| RGS17 | 0.96425667 | 0.35571663 | 3.77623334 | 0.00247059 | 0.01077706 |
| PTGER2 | -0.61564067 | 2.44796762 | -3.77415788 | 0.00248015 | 0.01081253 |
| GRIP1 | 0.85633887 | 1.09828677 | 3.72096153 | 0.00273853 | 0.0116866 |
| PLA2G4A | -0.56011317 | 2.76652033 | -3.71338823 | 0.00277749 | 0.01180626 |
| SEMA3D | 0.6251575 | 2.2980607 | 3.70388579 | 0.00282717 | 0.01198047 |
| BBS12 | 0.67392953 | 2.17701836 | 3.69650658 | 0.00286638 | 0.01210255 |
| PPFIA4 | -1.23005855 | 3.59210754 | -3.67179071 | 0.0030018 | 0.01251729 |
| CKMT2-AS1 | 0.50198316 | 3.12338716 | 3.66709569 | 0.00302826 | 0.01256876 |
| OVGP1 | 0.78097417 | 1.4776189 | 3.63982793 | 0.00318666 | 0.01308988 |
| LINC00320 | 0.64493947 | 2.15069895 | 3.58914462 | 0.00350392 | 0.01407609 |
| FTCDNL1 | 0.61992201 | 2.12722193 | 3.57465194 | 0.00360042 | 0.01437876 |
| TMEM74B | -0.701508 | 2.7983288 | -3.56687831 | 0.00365329 | 0.01455261 |
| ADGRB3 | 0.6788336 | 1.70374704 | 3.55155767 | 0.00375982 | 0.01489466 |
| FOXG1-AS1 | 0.76963017 | 1.24439377 | 3.53186197 | 0.00390142 | 0.01528475 |
| NPEPL1 | -0.56153335 | 3.8957528 | -3.52854371 | 0.0039258 | 0.01535788 |
| WDR25 | 0.54606676 | 3.83363496 | 3.52666508 | 0.00393968 | 0.01539372 |
| ATOH8 | 0.56340364 | 2.62810151 | 3.51575467 | 0.00402125 | 0.01562774 |
| LINC00663 | 0.76069773 | 1.88660517 | 3.4602544 | 0.00446351 | 0.01698466 |
| SNCA-AS1 | 0.91516925 | 0.24553914 | 3.43539022 | 0.00467735 | 0.01761247 |
| LINC02249 | 0.88208097 | 0.4935859 | 3.42663597 | 0.00475509 | 0.0178298 |
| CYP26B1 | -0.55164434 | 2.57605295 | -3.37744696 | 0.00521685 | 0.01905171 |
| MANEA-DT | 0.95168687 | -0.49819777 | 3.37178459 | 0.00527284 | 0.01918215 |
| ICA1 | -0.83226145 | 0.80252607 | -3.34932999 | 0.00550092 | 0.01982609 |

| | | | | | |
|---|---|---|---|---|---|
| KCNS3 | -1.06440227 | -0.31303956 | -3.33300471 | 0.00567297 | 0.02029331 |
| NT5E | 0.51469772 | 2.69644405 | 3.32089462 | 0.00580411 | 0.02060492 |
| CHSY3 | 0.54493531 | 2.35547975 | 3.31470415 | 0.00587232 | 0.02079344 |
| MANCR | 0.81470898 | 0.60438909 | 3.31201775 | 0.00590217 | 0.0208864 |
| PTGS2 | -0.90083068 | 0.23528227 | -3.29497209 | 0.00609518 | 0.02135894 |
| AGAP2-AS1 | -0.55294752 | 2.29265616 | -3.28383197 | 0.00622475 | 0.02170134 |
| CTSO | 0.62074853 | 1.82015488 | 3.26825019 | 0.00641065 | 0.02216188 |
| FIGN | 0.8703223 | 0.42539373 | 3.26822056 | 0.00641101 | 0.02216188 |
| NAALADL2 | 0.68959359 | 1.50333747 | 3.26151075 | 0.00649277 | 0.02235777 |
| PURG | 0.79043438 | 0.75993454 | 3.25845365 | 0.00653038 | 0.02245151 |
| HMGCLL1 | -0.70707877 | 1.72993597 | -3.23505515 | 0.00682554 | 0.02323851 |
| SEMA3E | -0.53284598 | 2.44444707 | -3.1937379 | 0.00737989 | 0.02465304 |
| NKX2-8 | -0.66675585 | 1.72977354 | -3.1764895 | 0.0076245 | 0.02525542 |
| VSTM4 | -0.72054814 | 1.12172521 | -3.1707791 | 0.00770726 | 0.0254516 |
| CCDC181 | 0.76555617 | 0.90164964 | 3.14774228 | 0.00805038 | 0.02638807 |
| MXI1 | -0.50907491 | 4.77299962 | -3.10067093 | 0.00879983 | 0.02817473 |
| C18orf65 | 0.73793263 | 0.82285476 | 3.06745947 | 0.00937025 | 0.02953883 |
| LINC00703 | 0.50408178 | 2.17239783 | 3.06214591 | 0.00946488 | 0.02981232 |
| COBL | -0.74809433 | 0.76154716 | -3.04630731 | 0.00975266 | 0.03058984 |
| ST8SIA6 | 0.59092852 | 1.70582023 | 3.02443121 | 0.01016454 | 0.03151779 |
| LINC02282 | 0.68199877 | 1.13926205 | 3.02252664 | 0.01020121 | 0.0316056 |
| LINC00535 | 0.82938264 | 0.30278082 | 3.02005129 | 0.01024907 | 0.03170865 |
| ITGBL1 | 0.81643794 | 0.50712966 | 3.01445749 | 0.01035804 | 0.03198025 |
| LINC01551 | 0.51352849 | 2.52025475 | 3.00679015 | 0.01050929 | 0.03231036 |
| NXPH1 | 0.62325158 | 1.41913914 | 2.97527793 | 0.01115435 | 0.0338385 |
| RGMA | 0.67904595 | 1.07576617 | 2.94927011 | 0.01171629 | 0.0351907 |
| MSS51 | -0.68095589 | 1.57530443 | -2.94433041 | 0.01182615 | 0.03545039 |
| TP53TG3D | 0.82526541 | 0.11712717 | 2.94102885 | 0.01190016 | 0.03563934 |
| LINC01473 | 0.79215012 | 0.24368027 | 2.93749433 | 0.01197989 | 0.03584025 |
| ST6GALNAC3 | -0.74358366 | 1.31250789 | -2.93559623 | 0.01202293 | 0.0359406 |
| DLGAP4-AS1 | 0.64550343 | 1.35710201 | 2.92996708 | 0.01215147 | 0.0362192 |
| TBX5 | 0.51534416 | 2.21030576 | 2.91892909 | 0.01240749 | 0.03680677 |
| CRIP1 | -0.91693295 | -0.48060505 | -2.90271734 | 0.01279324 | 0.03767064 |
| RANBP3L | 0.70435639 | 1.01444556 | 2.89135051 | 0.01307078 | 0.03831324 |
| KCNN2 | 0.54854986 | 1.680117 | 2.86708257 | 0.01368343 | 0.03971659 |
| SLITRK1 | 0.57852279 | 1.53756699 | 2.85971025 | 0.01387511 | 0.04016885 |
| ZBTB20 | 0.66041473 | 0.9711001 | 2.85285426 | 0.01405575 | 0.04051365 |
| CXCR4 | -1.5799691 | 1.18480272 | -2.8491366 | 0.01415468 | 0.04075157 |
| SIX6 | 0.54460623 | 1.94827567 | 2.8412135 | 0.01436781 | 0.04121693 |
| SRGAP1 | -0.50104241 | 2.13126566 | -2.82318739 | 0.01486458 | 0.04228155 |
| AXDND1 | 0.82710338 | -0.13532778 | 2.79007257 | 0.01582176 | 0.04434603 |
| NTN4 | 0.57763245 | 1.5343765 | 2.77585607 | 0.01625109 | 0.04528115 |
| TENT5C | 0.76028009 | 0.09262841 | 2.76918252 | 0.01645657 | 0.04571906 |
| RAPGEF4 | 0.71009786 | 0.62799826 | 2.73929291 | 0.01740866 | 0.04782011 |
| LINC01876 | 0.58807821 | 1.75108975 | 2.70905369 | 0.01842691 | 0.04983956 |
