## Supplementary Tables for "Zinc finger protein SALL4 functions through an AT-rich motif to regulate gene expression": Supplementary Table 1.pdf

**Supplementary Table 1.** Oligonucleotide sequences used in EMSA and ITC assays

| | |
|---|---|
| PBM WT oligo 1 | GTGAAAAAAAATATTAACGTACAGCGGGGAGGCGGC |
| PBM Mutant oligo 1 | AAAAGCGCAGGCATTAAAGGTATACGTGTGAAAAGA |
| PBM WT oligo 2 | TTAAGCAGAAATATTACGGTCTCCGGATTTGGCGCT |
| PBM Mutant oligo 2 | ATTTACAACAGGCCAGAAGTTCTTTGGCTTATCCAT |
| ITC WT oligo 1 | GAGTTATTAATG |
| ITC Mutant oligo 1 | GAGTCGCTAATG |
| ITC WT oligo 2 | GATAAATATTTG |
| ITC Mutant oligo 2 | GATAAACGCTTG |
| *KDM3A* WT oligo | TCTTCATTTATCCTTCAAAA |
| *KDM3A* Mutant oligo | TCTTCTTTTAACCTTCAAAA |
