## Supplementary Tables for "Zinc finger protein SALL4 functions through an AT-rich motif to regulate gene expression": Supplementary Table 3.pdf

**Supplementary Table 3.** shSALL4 target sequences CRISPR/Cas9 gRNA sequences

| | |
|---|---|
| shSALL4-1 target region | GCGTTGAAACAGGCCAAGCTG |
| shSALL4-2 target region | CTATTTAGCCAAAGGCAAA |
| CRISPR/Cas9 targeted region | ATCATTC<u>ATTATG</u>GCCTTCAACTACT |
