## Supplementary Tables for "Zinc finger protein SALL4 functions through an AT-rich motif to regulate gene expression": Supplementary Table 6.pdf

**Supplementary Table 6.** List of ChIP-qPCR and qPCR primers

| | |
|---|---|
| *KDM3A* ChIP primers | |
| Positive forward | CCTCACCCTTTCCTGTGAGA |
| Positive reverse | CGCGAAATCGGTTATCAACT |
| Negative forward | AACGGAGACCAGAAAGTTGG |
| Negative reverse | TGAAGCGTGTCTGAACAACC |

| | |
|---|---|
| *KDM3A* qPCR primers | |
| Forward_1 | GGAGTTCAAGGCTGGGCTAT |
| Reverse_1 | TCCTGAGTAAGCCAGAAGCAG |
| Forward_2 | ATTTGCAGGTGCTGCTTACA |
| Reverse_2 | CCTGTTTGAACAAGGGCAGT |
| *KDM4C* qPCR primers | |
| Forward | CACCTGCTGAGGGAGAAGTC |
| Reverse | GCATCTGCCAGCACTTACAA |
| *RCOR3* qPCR primers | |
| Forward | TCCAGATAAGACAATTGCAAGC |
| Reverse | TCACTATCATTCCCATCCATTG |
| h18S rRNA control qPCR primers | |
| Forward | GTAACCCGTTGAACCCCATT |
| Reverse | CCATCCAATCGGTAGTAGCG |
| *GAPDH* control qPCR primers | |
| Forward | GAAGGTGAAGGTCGGAGTCAAC |
| Reverse | TGGAAGATGGTGATGGGATTTC |
