## Supplementary Tables for "Zinc finger protein SALL4 functions through an AT-rich motif to regulate gene expression": SupplTable2 SupplTable2.pdf

Supplementary Table 2. SALL4 CUT&RUN peaks in SNU398 liver cancer cells common among three biological triplicates.

| | | |
|---|---|---|
| chr6 | 72363826 | 72366664 |
| chr5 | 117460026 | 117462489 |
| chr7 | 84270226 | 84273278 |
| chr9 | 63801469 | 63804125 |
| chr15 | 86387083 | 86388654 |
| chr10 | 108343491 | 108345735 |
| chr16 | 36225509 | 36229271 |
| chr15 | 97654709 | 97656285 |
| chr12 | 44054474 | 44057463 |
| chr16 | 34746660 | 34748464 |
| chr10 | 99241529 | 99242862 |
| chr7 | 82829868 | 82831989 |
| chr10 | 124724314 | 124725639 |
| chr14 | 66322607 | 66325834 |
| chr7 | 80466781 | 80470112 |
| chr4 | 93048615 | 93050693 |
| chr6 | 48190210 | 48192671 |
| chr16 | 74109473 | 74110862 |
| chr4 | 30959309 | 30961013 |
| chr12 | 77834122 | 77836118 |
| chr10 | 35545799 | 35547835 |
| chr2 | 163740871 | 163743529 |
| chr14 | 38503300 | 38507553 |
| chr8_KI270810v1_a | 309773 | 311065 |
| chr2 | 83174405 | 83176962 |
| chr7 | 88824811 | 88827310 |
| chr1 | 90700136 | 90702988 |
| chr15 | 51494647 | 51496403 |
| chr7 | 96219473 | 96220781 |
| chr12 | 25319420 | 25321907 |
| chr10 | 21530304 | 21532084 |
| chr12 | 63783290 | 63786123 |
| chr17_KI270857v1_ | 2164184 | 2167280 |
| chr7 | 9431108 | 9433174 |
| chr10 | 38961409 | 38962971 |
| chr10 | 35072554 | 35073776 |
| chr1 | 213946068 | 213948045 |
| chr1 | 100466843 | 100470001 |
| chr2 | 185314057 | 185317581 |
| chr1 | 195218880 | 195220460 |
| chr18 | 68960868 | 68962563 |

| | | |
|---|---|---|
| chr2 | 168169895 | 168172706 |
| chr3 | 76968955 | 76971502 |
| chr8 | 82458161 | 82459733 |
| chr4 | 156656040 | 156658143 |
| chr14 | 96852220 | 96854308 |
| chr10 | 36905003 | 36906693 |
| chr4 | 35515246 | 35518754 |
| chr13 | 66910663 | 66914950 |
| chr14 | 82962669 | 82964129 |
| chr21 | 15893142 | 15896016 |
| chr2 | 56131049 | 56132907 |
| chr10 | 87284127 | 87285859 |
| chr9 | 73136229 | 73138644 |
| chr22 | 11577540 | 11580541 |
| chr10 | 113160686 | 113162650 |
| chr15 | 40106707 | 40108468 |
| chr1 | 237429906 | 237431967 |
| chr15 | 52350567 | 52352564 |
| chr6 | 98841474 | 98844581 |
| chr22 | 12609364 | 12611770 |
| chr1 | 178126293 | 178128425 |
| chr2 | 62884287 | 62886438 |
| chr2 | 17425330 | 17427930 |
| chr12 | 46675810 | 46677723 |
| chr18 | 66706782 | 66709463 |
| chr22 | 10735132 | 10737121 |
| chr18 | 21116616 | 21119651 |
| chr1 | 144327383 | 144330687 |
| chr14 | 101491601 | 101492893 |
| chr7 | 77255100 | 77256664 |
| chr1 | 80553567 | 80556622 |
| chr10 | 116674180 | 116676327 |
| chr8 | 76019866 | 76021429 |
| chr4 | 123402461 | 123405899 |
| chr4 | 175826461 | 175829075 |
| chr1 | 190119664 | 190122291 |
| chr13 | 105772394 | 105775865 |
| chr9 | 13442854 | 13445026 |
| chr20 | 30870702 | 30873103 |
| chr1 | 103838952 | 103840869 |
| chr2 | 177145378 | 177151112 |
| chr9 | 67293720 | 67295599 |
| chr14 | 88568973 | 88571014 |
| chr22 | 50397092 | 50398414 |
| chr9 | 6411881 | 6413352 |
| chr21 | 9964379 | 9968288 |
| chr1 | 187794205 | 187796192 |
| chr2 | 102341220 | 102343009 |
| chr12 | 76877055 | 76880346 |

| | | |
|---|---|---|
| chr1 | 205527411 | 205529496 |
| chr6 | 52995885 | 52997149 |
| chr4 | 30470255 | 30474605 |
| chr6 | 80004895 | 80007553 |
| chr2 | 53568611 | 53570431 |
| chr12 | 84624308 | 84626725 |
| chr15 | 37526910 | 37529015 |
| chr12 | 17372467 | 17375103 |
| chr11 | 23726584 | 23729122 |
| chr21 | 22789796 | 22792509 |
| chr2 | 192065062 | 192067428 |
| chr6 | 7593497 | 7594841 |
| chr6 | 53350240 | 53352202 |
| chr1 | 200128409 | 200131734 |
| chr1 | 187454253 | 187456889 |
| chr1 | 190695087 | 190697450 |
| chr1 | 195464007 | 195465501 |
| chr4 | 90861629 | 90863107 |
| chr10 | 54627636 | 54629438 |
| chr21 | 10415817 | 10416909 |
| chr6 | 22595628 | 22597593 |
| chr6 | 153463762 | 153465066 |
| chr1 | 104077827 | 104080388 |
| chr11 | 92222723 | 92224150 |
| chr1 | 107809270 | 107811039 |
| chr12 | 84695880 | 84699285 |
| chr1 | 172357421 | 172360518 |
| chr2 | 71226716 | 71228659 |
| chr1_KI270765v1_a | 105037 | 106205 |
| chr14 | 101425658 | 101428285 |
| chr5 | 129242321 | 129244164 |
| chr2 | 193189173 | 193190390 |
| chr6 | 62730950 | 62733622 |
| chr7 | 82047771 | 82050030 |
| chr7 | 80496593 | 80500586 |
| chr1 | 225084772 | 225086560 |
| chr12 | 76122368 | 76124554 |
| chr2 | 163244618 | 163245881 |
| chr22 | 12604010 | 12605563 |
| chr11 | 104688948 | 104691219 |
| chr14 | 47467461 | 47469538 |
| chr14 | 26824533 | 26826436 |
| chr2 | 57620061 | 57622808 |
| chr1 | 152559429 | 152561984 |
| chr2 | 195455934 | 195458313 |
| chr12 | 103161101 | 103163646 |
| chr16 | 76928879 | 76931400 |
| chr21 | 16426550 | 16428229 |
| chr6 | 124182275 | 124183552 |

| | | |
|---|---|---|
| chr14 | 57289475 | 57291554 |
| chr2 | 50456147 | 50457790 |
| chr2 | 57461031 | 57463320 |
| chr14 | 19617384 | 19620962 |
| chr22 | 12629300 | 12632517 |
| chr10 | 115435733 | 115439886 |
| chr9 | 62475201 | 62477041 |
| chr12 | 67087015 | 67088595 |
| chr8 | 75973750 | 75977668 |
| chr7 | 84652976 | 84654713 |
| chr7 | 61070116 | 61073551 |
| chr16 | 55138009 | 55139855 |
| chr20 | 46401864 | 46403719 |
| chr14 | 41984187 | 41986972 |
| chr16 | 34751223 | 34753697 |
| chr15 | 36642654 | 36646353 |
| chr2 | 95934782 | 95938069 |
| chr21 | 10461402 | 10465193 |
| chr1 | 97571589 | 97573607 |
| chr3 | 174533358 | 174534468 |
| chr2 | 199860731 | 199862514 |
| chr10 | 55016869 | 55018627 |
| chr2 | 31872002 | 31875835 |
| chr2 | 35084784 | 35086118 |
| chr2 | 83068595 | 83071885 |
| chr15 | 68290678 | 68293026 |
| chr7 | 127347721 | 127350647 |
| chr5 | 172764349 | 172766000 |
| chr12 | 89460843 | 89462539 |
| chr2 | 219168565 | 219170455 |
| chr16 | 29592724 | 29594918 |
| chr1 | 102946505 | 102950968 |
| chr7 | 57707727 | 57709026 |
| chr3 | 81587612 | 81590290 |
| chr10 | 91123371 | 91125323 |
| chr2 | 169544265 | 169546138 |
| chr8 | 76194676 | 76197960 |
| chr5 | 93084203 | 93086599 |
| chr9 | 73085341 | 73087475 |
| chr2 | 51846480 | 51848011 |
| chr10 | 65888830 | 65890965 |
| chr11 | 24918290 | 24920348 |
| chr14 | 53541252 | 53545101 |
| chr2 | 215973856 | 215976188 |
| chr2 | 206000507 | 206001941 |
| chr6 | 130794434 | 130796025 |
| chr1 | 150066954 | 150069386 |
| chr2 | 83034122 | 83036706 |
| chr2 | 144997047 | 145001148 |

| | | |
|---|---|---|
| chr2 | 226969128 | 226971080 |
| chr15 | 47840753 | 47842951 |
| chr3 | 120093478 | 120096145 |
| chr7 | 82681385 | 82683374 |
| chr2 | 34987655 | 34990611 |
| chr18 | 25634277 | 25636066 |
| chr1 | 97566418 | 97570533 |
| chr10 | 49990133 | 49993008 |
| chr16 | 63639164 | 63640213 |
| chr11 | 127613706 | 127616362 |
| chr16 | 5153381 | 5154820 |
| chr6 | 92326228 | 92328208 |
| chr8 | 128872981 | 128876040 |
| chr14 | 100267647 | 100269141 |
| chr10 | 59109270 | 59110612 |
| chr21 | 9325260 | 9327767 |
| chr4 | 31417155 | 31419658 |
| chr11 | 16737127 | 16739379 |
| chr2 | 58426430 | 58428437 |
| chr1 | 16740304 | 16742175 |
| chr6 | 60417772 | 60419467 |
| chr14 | 88694889 | 88696437 |
| chr9 | 62807932 | 62810813 |
| chr9 | 41847374 | 41849198 |
| chr12 | 44256404 | 44258712 |
| chr19 | 9911405 | 9913344 |
| chr7 | 156700690 | 156701695 |
| chr5 | 52782519 | 52784889 |
| chr7 | 83832305 | 83835131 |
| chr15 | 20408209 | 20410032 |
| chr1 | 195505142 | 195507475 |
| chr10 | 55945657 | 55948332 |
| chr2 | 186659251 | 186660452 |
| chr1 | 105517056 | 105518533 |
| chr5 | 144176826 | 144179105 |
| chr1 | 38884713 | 38885897 |
| chr2 | 179678617 | 179679964 |
| chr4 | 20486496 | 20489619 |
| chr17 | 59962253 | 59965139 |
| chr1 | 187035544 | 187038219 |
| chr15 | 38066589 | 38068659 |
| chr20 | 29349831 | 29351683 |
| chr6 | 92037131 | 92039080 |
| chr1 | 180691039 | 180693881 |
| chr1 | 218705417 | 218707574 |
| chr2 | 162332392 | 162334704 |
| chr7 | 80095241 | 80097681 |
| chr15 | 75389010 | 75390159 |
| chr9 | 68243781 | 68245555 |

| | | |
|---|---|---|
| chr12 | 99411870 | 99413888 |
| chr20 | 29501599 | 29503592 |
| chr3 | 76454443 | 76456591 |
| chr21 | 10359738 | 10361812 |
| chr6 | 127439750 | 127442114 |
| chr17 | 20800933 | 20802718 |
| chr16 | 87118948 | 87121332 |
| chr10 | 54648272 | 54650498 |
| chr7 | 95475980 | 95480079 |
| chr14 | 33102221 | 33104825 |
| chr16 | 76280232 | 76282934 |
| chr6 | 71293783 | 71295862 |
| chr9 | 73154539 | 73157338 |
| chr11 | 102805853 | 102807577 |
| chr1 | 65418515 | 65421688 |
| chr1 | 219321205 | 219324357 |
| chr6 | 136288354 | 136290996 |
| chr2 | 57446879 | 57449853 |
| chr1 | 179021108 | 179023380 |
| chr12 | 90675659 | 90677343 |
| chr2 | 56763058 | 56764243 |
| chr15 | 23559958 | 23561975 |
| chr18 | 14178376 | 14180663 |
| chr15_KI270850v1_ | 291251 | 293372 |
| chr5 | 52727194 | 52728310 |
| chr1 | 99011444 | 99015367 |
| chr12 | 96864245 | 96866200 |
| chr9 | 82154998 | 82156625 |
| chr1 | 242421073 | 242423231 |
| chr10 | 64339788 | 64341200 |
| chr10 | 50575794 | 50577350 |
| chr14 | 32600186 | 32602410 |
| chr12 | 58188461 | 58191050 |
| chr1 | 176467440 | 176470316 |
| chr2 | 68523046 | 68524370 |
| chr4 | 34716284 | 34719746 |
| chr14 | 56278660 | 56279699 |
| chr2 | 144519711 | 144521935 |
| chr13 | 106349624 | 106351637 |
| chr4 | 37686169 | 37687744 |
| chr8 | 111071308 | 111073068 |
| chr9 | 65805045 | 65809056 |
| chr5 | 27821315 | 27824699 |
| chr1 | 191495065 | 191498290 |
| chr10 | 110572831 | 110574321 |
| chr6 | 76357109 | 76358360 |
| chr4 | 34092670 | 34095500 |
| chr14 | 84627840 | 84628909 |
| chr7 | 62437154 | 62440819 |

| | | |
|---|---|---|
| chr10 | 100546676 | 100548302 |
| chr2 | 184953985 | 184955744 |
| chr10 | 34769787 | 34772203 |
| chr3 | 38159615 | 38162449 |
| chr10 | 61245666 | 61247251 |
| chr8 | 106656404 | 106659511 |
| chr2 | 197000034 | 197002531 |
| chr14 | 27345987 | 27348086 |
| chr11 | 62727379 | 62729533 |
| chr10 | 61041704 | 61043236 |
| chr22 | 10933591 | 10936187 |
| chr7 | 94396351 | 94399165 |
| chr4 | 34104747 | 34107308 |
| chr7 | 83495391 | 83498196 |
| chr7 | 82701888 | 82703515 |
| chr14 | 19925114 | 19926523 |
| chr4 | 78871401 | 78872712 |
| chr2 | 51293151 | 51294306 |
| chr1 | 90712254 | 90715337 |
| chr10 | 85727051 | 85729943 |
| chrY | 6251618 | 6252903 |
| chr11 | 89758406 | 89760534 |
| chr12 | 65469540 | 65471417 |
| chr14 | 58229949 | 58232330 |
| chr2 | 177420345 | 177421903 |
| chr13 | 41810766 | 41814212 |
| chr20 | 10404343 | 10405784 |
| chr6 | 151944823 | 151946509 |
| chr1 | 98384365 | 98387276 |
| chr6 | 1795853 | 1797951 |
| chr10 | 65365659 | 65367754 |
| chr1 | 90918116 | 90919638 |
| chr1 | 119954278 | 119957245 |
| chr20 | 64301406 | 64303442 |
| chr4 | 90922762 | 90925065 |
| chr9 | 62369142 | 62371264 |
| chr9 | 67830698 | 67832582 |
| chr14 | 40979402 | 40981995 |
| chr1 | 192002795 | 192005788 |
| chr11 | 66907676 | 66909942 |
| chr7 | 18842706 | 18844935 |
| chr4 | 156614348 | 156616353 |
| chr1 | 218817525 | 218819938 |
| chr12 | 84582663 | 84584960 |
| chr4 | 31396680 | 31397732 |
| chr12 | 77946417 | 77948994 |
| chr5 | 26575718 | 26578125 |
| chr3 | 85454535 | 85455979 |
| chr2 | 55693188 | 55694904 |

| | | |
|---|---|---|
| chr14 | 88503532 | 88506306 |
| chr10 | 53326877 | 53329135 |
| chr16 | 76559818 | 76561444 |
| chr11 | 87289814 | 87291291 |
| chr14 | 43892185 | 43893785 |
| chr2 | 55292584 | 55295239 |
| chr5 | 7856082 | 7857713 |
| chr6 | 27471768 | 27473507 |
| chr3 | 95277097 | 95278440 |
| chr5 | 120753554 | 120755432 |
| chr5 | 118901501 | 118904499 |
| chr1 | 200668662 | 200671008 |
| chr8 | 123396245 | 123397407 |
| chr14 | 31026020 | 31028425 |
| chr6 | 705486 | 709063 |
| chr8 | 76566623 | 76568457 |
| chr11 | 87970772 | 87973725 |
| chr13 | 106160780 | 106163015 |
| chr11 | 5787380 | 5790332 |
| chr6 | 53067936 | 53069016 |
| chr1 | 97833924 | 97836175 |
| chr11 | 89908107 | 89910367 |
| chr2 | 53307458 | 53309188 |
| chr10 | 88270082 | 88272073 |
| chr8 | 27918100 | 27919611 |
| chr14 | 89698636 | 89701072 |
| chr5 | 16477191 | 16479006 |
| chr14 | 87484281 | 87486523 |
| chr14 | 39117318 | 39118587 |
| chr10 | 116333622 | 116334757 |
| chr11 | 97254313 | 97257216 |
| chr2 | 170791387 | 170793548 |
| chr3 | 14199325 | 14200592 |
| chr1 | 193027750 | 193030667 |
| chr7 | 89478625 | 89480023 |
| chr14 | 49861020 | 49862959 |
| chr6 | 103956245 | 103957813 |
| chr2 | 113597386 | 113599813 |
| chr11 | 103065194 | 103068016 |
| chr6 | 72222198 | 72224009 |
| chr22 | 10684442 | 10685832 |
| chr11 | 124064390 | 124066595 |
| chr1 | 102280252 | 102282596 |
| chr5 | 27688246 | 27691173 |
| chr19 | 44447917 | 44450123 |
| chr7 | 9742789 | 9744754 |
| chr1 | 189656158 | 189658579 |
| chr8 | 16753213 | 16754369 |
| chr7 | 84337250 | 84341802 |

| | | |
|---|---|---|
| chr2 | 195999614 | 196001607 |
| chr2 | 57328518 | 57332598 |
| chr12 | 103965485 | 103967729 |
| chr1 | 193223749 | 193225294 |
| chr1 | 198125373 | 198127412 |
| chr6 | 80771821 | 80774580 |
| chr9 | 13089110 | 13091281 |
| chr1 | 71188455 | 71192778 |
| chr2 | 192421219 | 192423935 |
| chr22 | 15559349 | 15560769 |
| chr8 | 89994082 | 89996503 |
| chr2 | 185833323 | 185835504 |
| chr10 | 100912408 | 100914366 |
| chr2 | 168973337 | 168976441 |
| chr10 | 17234468 | 17240193 |
| chr7 | 90116103 | 90118692 |
| chr9 | 31497617 | 31499691 |
| chr12 | 48779167 | 48780757 |
| chr14 | 19760554 | 19762524 |
| chr10 | 108290705 | 108293295 |
| chr15 | 93080623 | 93083447 |
| chr14 | 100789792 | 100791650 |
| chrUn_GL000218v1 | 20869 | 24480 |
| chr14 | 19613204 | 19615224 |
| chr8 | 93024522 | 93027002 |
| chr12 | 77970910 | 77973544 |
| chr14 | 52549660 | 52553013 |
| chr2 | 67182494 | 67184669 |
| chr2 | 53170123 | 53171558 |
| chr3 | 168509763 | 168511809 |
| chr11 | 95321967 | 95323704 |
| chr14 | 68190072 | 68191807 |
| chr7 | 87714630 | 87715646 |
| chr10 | 52927507 | 52929590 |
| chr14 | 86927054 | 86929006 |
| chr8 | 92962170 | 92963650 |
| chr10 | 110640391 | 110642566 |
| chr19 | 19872607 | 19874193 |
| chr2 | 162300100 | 162303444 |
| chr12 | 97835727 | 97837607 |
| chr8 | 47733205 | 47735138 |
| chr1 | 99440869 | 99444461 |
| chr2 | 88363929 | 88366023 |
| chr14 | 63641350 | 63643569 |
| chr2 | 58857513 | 58859611 |
| chr1 | 161468474 | 161471554 |
| chr15_KI270850v1_ | 259093 | 260628 |
| chr12 | 95761677 | 95764423 |
| chr1 | 104925680 | 104927139 |

| | | |
|---|---|---|
| chr7 | 82843894 | 82846206 |
| chr12 | 16510484 | 16511939 |
| chr18 | 24161736 | 24163703 |
| chr14 | 40802336 | 40805181 |
| chr5 | 130389001 | 130391209 |
| chr2 | 161956072 | 161959031 |
| chr7 | 80215915 | 80218640 |
| chr2 | 22291802 | 22293511 |
| chr14 | 106258852 | 106260857 |
| chr4 | 8969850 | 8970853 |
| chr14 | 89953894 | 89955593 |
| chr16 | 33732869 | 33736374 |
| chr3 | 102929598 | 102931887 |
| chr15 | 20365866 | 20367926 |
| chr21 | 28570252 | 28572772 |
| chr7 | 104154378 | 104157592 |
| chr3 | 147513182 | 147515983 |
| chr9 | 31541931 | 31543387 |
| chr6 | 31619430 | 31621180 |
| chr6 | 81084552 | 81087871 |
| chr17 | 79178398 | 79179724 |
| chr14 | 59468520 | 59470490 |
| chr10 | 68333370 | 68335266 |
| chr14 | 37420528 | 37424052 |
| chr1 | 149389729 | 149390976 |
| chr2 | 135984342 | 135985503 |
| chr7 | 96783836 | 96785379 |
| chr10 | 115228071 | 115230007 |
| chr5 | 123344568 | 123346197 |
| chr14 | 92603181 | 92604935 |
| chr1 | 103897586 | 103898898 |
| chr8 | 115453362 | 115455058 |
| chr10 | 54642107 | 54645067 |
| chr11 | 99302549 | 99304579 |
| chr12 | 77923437 | 77926294 |
| chr12 | 25243525 | 25246426 |
| chr1 | 107776920 | 107779318 |
| chr1 | 101110190 | 101112078 |
| chr10 | 104208372 | 104210069 |
| chr8 | 115545199 | 115547468 |
| chr9 | 68247834 | 68250403 |
| chr16 | 79150975 | 79154770 |
| chr1 | 110618811 | 110621195 |
| chr4 | 122731686 | 122733721 |
| chr14 | 27497241 | 27500008 |
| chr3 | 107231140 | 107232614 |
| chr2 | 57693458 | 57695907 |
| chr12 | 85589700 | 85590794 |
| chr15 | 49434106 | 49437541 |

| | | |
|---|---|---|
| chr10 | 82826926 | 82829047 |
| chr2 | 187598836 | 187601441 |
| chr6 | 17609300 | 17610517 |
| chr15 | 88900486 | 88901836 |
| chr10 | 84890057 | 84892070 |
| chr6 | 103275095 | 103276659 |
| chr12 | 85503874 | 85506267 |
| chr14 | 26828573 | 26831329 |
| chr1 | 102596875 | 102599191 |
| chr2 | 187406340 | 187409073 |
| chr13 | 106154608 | 106157529 |
| chr14 | 32497657 | 32500825 |
| chr8 | 40807114 | 40808726 |
| chr5_KI270897v1_a | 3883 | 7091 |
| chr20 | 29044278 | 29045926 |
| chr2 | 186522187 | 186524950 |
| chr13 | 87155261 | 87157158 |
| chr7 | 81121862 | 81123142 |
| chr12 | 18253179 | 18254392 |
| chr7 | 80461189 | 80462936 |
| chr12 | 65532090 | 65535427 |
| chr14 | 19306887 | 19309002 |
| chr10 | 60500014 | 60501463 |
| chr4 | 124511558 | 124513690 |
| chr14 | 29239362 | 29242042 |
| chr5 | 107761909 | 107764113 |
| chr2 | 115217353 | 115219308 |
| chr13 | 106171964 | 106174350 |
| chr6 | 112560412 | 112563002 |
| chr5_KI270897v1_a | 32624 | 34527 |
| chr7 | 80035655 | 80038318 |
| chr10 | 57203709 | 57205776 |
| chr10 | 58147363 | 58148522 |
| chr2 | 172051722 | 172056042 |
| chr12 | 87211601 | 87213262 |
| chr4 | 34962970 | 34966946 |
| chr16 | 85640063 | 85641666 |
| chr2 | 44678934 | 44681160 |
| chr7 | 81678071 | 81679157 |
| chr14 | 74737974 | 74739085 |
| chr4 | 170948322 | 170950672 |
| chr2 | 187876885 | 187879339 |
| chr10 | 79641995 | 79643197 |
| chr14 | 27141194 | 27142977 |
| chr8 | 17158114 | 17160447 |
| chr14 | 39433649 | 39437398 |
| chr10 | 57524967 | 57527497 |
| chr11 | 23039725 | 23043257 |
| chr12 | 86660076 | 86663378 |

| | | |
|---|---|---|
| chr2 | 43541906 | 43543035 |
| chr4 | 115713454 | 115715309 |
| chr2 | 76756716 | 76758958 |
| chr1 | 186106844 | 186109139 |
| chr12 | 65334198 | 65336285 |
| chr10 | 56084475 | 56087231 |
| chr12 | 25246500 | 25247931 |
| chr13 | 71603876 | 71606209 |
| chr8 | 7341293 | 7342449 |
| chr15_KI270905v1_ | 618304 | 619856 |
| chr6 | 75112276 | 75117861 |
| chr2 | 83845538 | 83847111 |
| chr17_KI270909v1_ | 309519 | 312233 |
| chr20 | 53308246 | 53309742 |
| chr2 | 184353903 | 184356840 |
| chr6 | 64072387 | 64075746 |
| chr4 | 69187762 | 69189741 |
| chr11 | 82036557 | 82039502 |
| chr4 | 34953498 | 34956509 |
| chr1 | 102798839 | 102800907 |
| chr14 | 41826749 | 41828744 |
| chr5 | 124740542 | 124742543 |
| chr9 | 18970315 | 18973079 |
| chr12 | 65846830 | 65848742 |
| chr10 | 112182987 | 112184573 |
| chr13 | 84550476 | 84551806 |
| chr8 | 16898581 | 16901393 |
| chr7 | 82905630 | 82909229 |
| chr14 | 47263186 | 47264857 |
| chr2 | 60443541 | 60444879 |
| chr9 | 67807869 | 67809685 |
| chr11 | 110625520 | 110627236 |
| chr14 | 71006006 | 71008446 |
| chr11 | 30594468 | 30597331 |
| chr14 | 74492114 | 74494108 |
| chr8 | 71049362 | 71050865 |
| chr1 | 143268065 | 143269977 |
| chr14 | 83756096 | 83757953 |
| chr5 | 16464534 | 16467502 |
| chr9 | 82597852 | 82600319 |
| chr1 | 98034527 | 98037704 |
| chr2 | 55734393 | 55736500 |
| chr21 | 8220601 | 8222530 |
| chr1 | 113414404 | 113417287 |
| chr10 | 50622283 | 50624940 |
| chr19 | 36238747 | 36240582 |
| chr4 | 33728095 | 33729831 |
| chr1 | 104027192 | 104031428 |
| chr14 | 55028833 | 55030659 |

| | | |
|---|---|---|
| chr11 | 90006465 | 90009601 |
| chr7 | 90854210 | 90856124 |
| chr4 | 78918985 | 78920765 |
| chr12 | 88110411 | 88112459 |
| chr14 | 60974608 | 60978183 |
| chr7 | 13043803 | 13048723 |
| chr15_KI270852v1_ | 6561 | 11877 |
| chr13 | 67067556 | 67071251 |
| chr3 | 83132526 | 83134765 |
| chr21 | 21352928 | 21354733 |
| chr11 | 16174459 | 16176340 |
| chr14 | 27863787 | 27865904 |
| chr12 | 77855937 | 77859935 |
| chr12 | 99017974 | 99019885 |
| chr10 | 63216291 | 63219086 |
| chr14 | 36991503 | 36996692 |
| chr6 | 63059174 | 63060742 |
| chr14 | 38816620 | 38818109 |
| chr7 | 18845975 | 18847677 |
| chr1 | 190302674 | 190304144 |
| chr3 | 75434244 | 75437392 |
| chr7 | 50146776 | 50148386 |
| chr12 | 43680618 | 43683069 |
| chr17 | 65696489 | 65700052 |
| chr10 | 130106666 | 130108802 |
| chr14 | 105527255 | 105530141 |
| chr2 | 166132171 | 166133851 |
| chr5 | 106089435 | 106090570 |
| chr12 | 53499506 | 53502601 |
| chr14 | 27646082 | 27647320 |
| chr14 | 48308716 | 48311323 |
| chr1 | 97147873 | 97149592 |
| chr7 | 83437959 | 83439169 |
| chr21 | 10537976 | 10540803 |
| chr1 | 32071828 | 32073430 |
| chr7 | 88110640 | 88112329 |
| chr2 | 176812813 | 176815458 |
| chr2 | 225667041 | 225671174 |
| chr14 | 86324811 | 86327897 |
| chr4 | 90124745 | 90127998 |
| chr13 | 66581376 | 66582598 |
| chr4 | 136966197 | 136968106 |
| chr12 | 46469982 | 46472888 |
| chr11 | 5885125 | 5887463 |
| chr14 | 90892502 | 90895333 |
| chr4 | 46500162 | 46501714 |
| chr11 | 96241930 | 96244850 |
| chr1 | 196391077 | 196393067 |
| chr2 | 181176522 | 181179310 |

| | | |
|---|---|---|
| chr10 | 88915707 | 88917882 |
| chr12 | 84791849 | 84793785 |
| chr15 | 41330454 | 41334039 |
| chr2 | 173664971 | 173667059 |
| chr1 | 219151449 | 219154381 |
| chr2 | 146221621 | 146224077 |
| chr9 | 64846883 | 64849346 |
| chr10 | 117254736 | 117257608 |
| chr13 | 82507366 | 82509968 |
| chr12 | 41322681 | 41324654 |
| chr6 | 66974191 | 66976296 |
| chr4 | 116767094 | 116768709 |
| chr2 | 105054078 | 105056041 |
| chr13 | 59720027 | 59721558 |
| chr1 | 103734392 | 103735584 |
| chr14 | 51742038 | 51743188 |
| chr1 | 103736074 | 103737867 |
| chr8 | 102237629 | 102240480 |
| chr6 | 83098763 | 83100510 |
| chr1 | 171588139 | 171589729 |
| chr17 | 14299403 | 14301460 |
| chr3 | 20263089 | 20264984 |
| chr1 | 28647512 | 28649568 |
| chr6 | 121853877 | 121855698 |
| chr14 | 85956862 | 85958812 |
| chr14 | 30361458 | 30363443 |
| chr12 | 124417695 | 124420386 |
| chr2 | 145130714 | 145133442 |
| chr10 | 66350488 | 66353018 |
| chr3 | 155083705 | 155085101 |
| chr3 | 19064473 | 19067561 |
| chr2 | 195488268 | 195490048 |
| chr7 | 93854545 | 93856644 |
| chr14 | 30676846 | 30679056 |
| chr1 | 172268497 | 172271343 |
| chr3 | 76012669 | 76014360 |
| chr4 | 30404901 | 30406262 |
| chr12 | 85757004 | 85759345 |
| chr10 | 35015075 | 35017780 |
| chr15 | 48446977 | 48449532 |
| chr12 | 91440231 | 91443039 |
| chr4 | 92987295 | 92990547 |
| chr8 | 99520256 | 99522140 |
| chr7 | 20277091 | 20278839 |
| chr17 | 21833560 | 21835718 |
| chr3 | 86073321 | 86075258 |
| chr2 | 91551388 | 91554053 |
| chr11 | 100931487 | 100933694 |
| chr3 | 164792377 | 164794579 |

| | | |
|---|---|---|
| chr10 | 65361454 | 65363423 |
| chr16 | 31457865 | 31461335 |
| chr10 | 16942587 | 16945484 |
| chr10 | 75008230 | 75010105 |
| chr22 | 15899939 | 15901687 |
| chr6 | 45002731 | 45004383 |
| chr5 | 166631464 | 166634536 |
| chr7 | 49943235 | 49944865 |
| chr1 | 83288260 | 83290251 |
| chrY | 26435707 | 26437444 |
| chr9 | 66222983 | 66226957 |
| chr10 | 125888039 | 125889666 |
| chrUn_GL000195v1 | 43408 | 46486 |
| chr4 | 34125933 | 34127695 |
| chr22 | 10959901 | 10961503 |
| chr2 | 196112760 | 196114617 |
| chr1 | 192188172 | 192189683 |
| chr10 | 108989022 | 108990890 |
| chr10 | 38483166 | 38485953 |
| chr1 | 119546710 | 119547911 |
| chr15 | 61619974 | 61623022 |
| chr22 | 15647456 | 15648963 |
| chr11 | 60647737 | 60648991 |
| chr14 | 46254592 | 46256301 |
| chr16 | 34608732 | 34611200 |
| chr7 | 85136966 | 85138719 |
| chr11 | 31162017 | 31164057 |
| chr13 | 104236895 | 104242053 |
| chr12 | 85216412 | 85219638 |
| chr1 | 85634918 | 85636453 |
| chr2 | 181290712 | 181293205 |
| chr1 | 202161662 | 202163619 |
| chr6 | 70167310 | 70170703 |
| chr20 | 60149308 | 60151306 |
| chr2 | 41060537 | 41062490 |
| chr15 | 22702377 | 22704723 |
| chr7 | 70593084 | 70594679 |
| chr8 | 111742669 | 111743981 |
| chr4 | 22710642 | 22712620 |
| chr10 | 72273903 | 72276520 |
| chr3 | 152522984 | 152526539 |
| chr3 | 29285177 | 29289537 |
| chr20 | 29082659 | 29083820 |
| chr7 | 72827002 | 72830671 |
| chr2 | 107032578 | 107035441 |
| chr2 | 195324875 | 195328488 |
| chr10 | 64500918 | 64503337 |
| chr11 | 22586180 | 22588487 |
| chr5 | 90614790 | 90616916 |

| | | |
|---|---|---|
| chr7 | 89847320 | 89849012 |
| chr21 | 5243754 | 5245497 |
| chr19 | 50328954 | 50331066 |
| chr2 | 80412107 | 80414633 |
| chr7 | 57180920 | 57183470 |
| chr9 | 10164393 | 10165628 |
| chr2 | 164718796 | 164721380 |
| chr11 | 96237291 | 96241494 |
| chr13 | 105187629 | 105191025 |
| chr4 | 90690378 | 90692718 |
| chr15 | 37735952 | 37739579 |
| chr12 | 40062752 | 40065298 |
| chr10 | 56993321 | 56995086 |
| chr15_KI270905v1_ | 2661919 | 2664000 |
| chr17 | 65740805 | 65743908 |
| chr21 | 13561549 | 13562856 |
| chr10 | 99429915 | 99431859 |
| chr12 | 81322658 | 81325313 |
| chr4 | 115792323 | 115794783 |
| chr14 | 85602507 | 85603936 |
| chr2 | 194496145 | 194498227 |
| chr7 | 80117670 | 80118917 |
| chr2 | 39334979 | 39337984 |
| chr12 | 67037496 | 67041043 |
| chr2 | 208460419 | 208462530 |
| chr1 | 104182671 | 104184525 |
| chr1 | 16901888 | 16906425 |
| chr8 | 128984792 | 128986788 |
| chr14 | 22956315 | 22957814 |
| chr14 | 28383755 | 28385470 |
| chr1 | 217245908 | 217248402 |
| chr3 | 174283577 | 174286088 |
| chr1 | 178453539 | 178456621 |
| chr13 | 71800517 | 71803174 |
| chr15 | 35475579 | 35478218 |
| chr4 | 107253232 | 107254980 |
| chr13 | 80334476 | 80338820 |
| chr4 | 107127077 | 107132173 |
| chr6 | 74753699 | 74754901 |
| chr12 | 62545706 | 62547687 |
| chr1 | 143261434 | 143263738 |
| chr2 | 80381044 | 80383266 |
| chr5 | 129951186 | 129952812 |
| chr9 | 29866356 | 29868163 |
| chr9 | 40148941 | 40153056 |
| chr1 | 244835391 | 244837683 |
| chr1 | 102980601 | 102983235 |
| chr7 | 94781568 | 94783627 |
| chr1 | 170398816 | 170400260 |

| | | |
|---|---|---|
| chr7 | 90624163 | 90626594 |
| chr1 | 188272869 | 188274934 |
| chr6 | 70208652 | 70210941 |
| chr10 | 99695302 | 99696993 |
| chr16 | 79012912 | 79014811 |
| chr1 | 186569676 | 186570806 |
| chr7 | 93942923 | 93944565 |
| chr10 | 124469594 | 124471815 |
| chr12 | 74414484 | 74416326 |
| chr8 | 60649936 | 60653763 |
| chr7 | 79319862 | 79322527 |
| chr18 | 72672308 | 72673507 |
| chr1 | 125147993 | 125151053 |
| chr1 | 173823480 | 173825232 |
| chr10 | 124445119 | 124446923 |
| chr10 | 48807889 | 48810134 |
| chr7 | 47875499 | 47876765 |
| chr14 | 25008622 | 25011134 |
| chr12 | 50085333 | 50087220 |
| chr3 | 174494520 | 174496669 |
| chr9 | 66301014 | 66303092 |
| chr12 | 82966013 | 82968125 |
| chr17 | 10688273 | 10689982 |
| chr2 | 186701384 | 186703814 |
| chr7 | 79439245 | 79442726 |
| chr13 | 80234041 | 80236488 |
| chr1 | 102361141 | 102363625 |
| chr6 | 64207746 | 64210580 |
| chr6 | 94178175 | 94180642 |
| chr14 | 26858883 | 26861818 |
| chr9 | 31416180 | 31419223 |
| chr19 | 7506014 | 7507585 |
| chr2 | 229459917 | 229462260 |
| chr14 | 65584100 | 65586327 |
| chr7 | 84676253 | 84678223 |
| chr1 | 87681737 | 87684044 |
| chr6 | 132922172 | 132925711 |
| chr14 | 36822589 | 36826009 |
| chr15 | 37084985 | 37086835 |
| chr5 | 87542343 | 87544671 |
| chr1 | 191575645 | 191578403 |
| chr2 | 191811701 | 191813404 |
| chr17 | 63854457 | 63855843 |
| chr22 | 10718323 | 10720492 |
| chr2 | 192617340 | 192619545 |
| chr10 | 62437068 | 62440558 |
| chr11 | 71786107 | 71788424 |
| chr7 | 98547426 | 98548941 |
| chr4 | 107257649 | 107259219 |

| | | |
|---|---|---|
| chr20 | 20001621 | 20004525 |
| chr2 | 195362881 | 195365141 |
| chr20 | 5988966 | 5991187 |
| chr7 | 100313887 | 100315258 |
| chr6 | 73394343 | 73396318 |
| chr8 | 7365404 | 7366514 |
| chr7 | 93990034 | 93992867 |
| chr5 | 168578299 | 168581017 |
| chr10 | 92992881 | 92994772 |
| chr10 | 55195300 | 55197221 |
| chr10 | 74861213 | 74863294 |
| chr22 | 50163167 | 50164166 |
| chr2 | 229860270 | 229862102 |
| chr4 | 66493030 | 66495253 |
| chr13 | 106371610 | 106373667 |
| chr15 | 81657716 | 81660037 |
| chr5 | 27791304 | 27794080 |
| chr4 | 12144794 | 12146687 |
| chr14 | 25952794 | 25954778 |
| chr7 | 83410148 | 83412435 |
| chr21 | 16450723 | 16453174 |
| chr2 | 41840236 | 41842649 |
| chr5 | 27807547 | 27809586 |
| chr2 | 192381598 | 192382960 |
| chr12 | 85417058 | 85418816 |
| chr16 | 77483136 | 77485035 |
| chr8 | 91454436 | 91456194 |
| chr5 | 126162266 | 126164942 |
| chr1 | 191645255 | 191650182 |
| chr7 | 96478952 | 96480740 |
| chr6 | 24424770 | 24426289 |
| chr2 | 51218466 | 51220574 |
| chrUn_GL000195v1 | 40095 | 42627 |
| chr12 | 70327604 | 70329872 |
| chr2 | 78719775 | 78722024 |
| chr16 | 34700109 | 34705780 |
| chr10 | 46253638 | 46255582 |
| chr18 | 40426563 | 40427941 |
| chr2 | 159508065 | 159511304 |
| chr7 | 90469082 | 90471631 |
| chr3 | 173798307 | 173800295 |
| chr11 | 106472866 | 106474454 |
| chr1 | 99571941 | 99573885 |
| chr4 | 42616241 | 42618510 |
| chr2 | 57636861 | 57639431 |
| chr2 | 172091156 | 172092485 |
| chr10 | 65192233 | 65193773 |
| chr10 | 36614050 | 36616557 |
| chr10 | 104980295 | 104982200 |

| | | |
|---|---|---|
| chr20 | 60491695 | 60493678 |
| chr6 | 82943200 | 82944447 |
| chr21 | 9279673 | 9281145 |
| chr11 | 88595290 | 88598049 |
| chr2 | 184716619 | 184718011 |
| chr5 | 110420369 | 110422600 |
| chr2 | 58715209 | 58717255 |
| chr7 | 158949929 | 158951234 |
| chr1 | 190997451 | 191000441 |
| chr8 | 77932390 | 77934907 |
| chr6 | 78603495 | 78605583 |
| chr2 | 211998276 | 212000120 |
| chr15 | 63281351 | 63283595 |
| chr10 | 31759667 | 31761002 |
| chr2 | 178911945 | 178913798 |
| chr7 | 85034717 | 85039172 |
| chr14 | 42149224 | 42151296 |
| chr3 | 169393867 | 169395301 |
| chr11 | 20995515 | 20997812 |
| chr16_KI270853v1_ | 1030248 | 1032071 |
| chr8 | 68051794 | 68054192 |
| chr1 | 185350851 | 185352512 |
| chr14 | 19928237 | 19930147 |
| chr10 | 81677001 | 81678437 |
| chr7 | 77601627 | 77603902 |
| chr17 | 65543365 | 65544468 |
| chr2 | 84278462 | 84280352 |
| chr10 | 37118741 | 37120044 |
| chr9 | 73163083 | 73165898 |
| chr6 | 56325654 | 56327618 |
| chr8 | 114486506 | 114489128 |
| chr2 | 213124235 | 213126333 |
| chr3 | 85909470 | 85911027 |
| chr15 | 77069421 | 77071656 |
| chr2 | 39609127 | 39610436 |
| chr12 | 18461968 | 18464436 |
| chr1 | 192376617 | 192378437 |
| chr2 | 57342160 | 57344002 |
| chr2 | 215867912 | 215870110 |
| chr17 | 26725248 | 26726411 |
| chr6 | 91877552 | 91879680 |
| chr5 | 145223493 | 145224775 |
| chr3 | 21258796 | 21261774 |
| chr15 | 96028440 | 96032268 |
| chr8 | 105910677 | 105915245 |
| chr2 | 165843863 | 165847106 |
| chr1 | 84549725 | 84551709 |
| chr10 | 61898008 | 61901712 |
| chr13 | 91452417 | 91453715 |

| | | |
|---|---|---|
| chr3 | 187739930 | 187743366 |
| chr1 | 106253532 | 106255371 |
| chr1 | 202942853 | 202944362 |
| chr10 | 104121298 | 104123916 |
| chr1 | 192699232 | 192701589 |
| chr12 | 104523989 | 104525880 |
| chr12 | 81254289 | 81255705 |
| chr2 | 200392582 | 200395628 |
| chr7 | 144239164 | 144241877 |
| chr1 | 16366181 | 16367525 |
| chr12 | 63684105 | 63685908 |
| chr1 | 107428870 | 107431612 |
| chr3 | 25056542 | 25059037 |
| chr13 | 82121896 | 82123864 |
| chr10 | 113354487 | 113356596 |
| chr1 | 101142515 | 101144053 |
| chr6 | 74671076 | 74673625 |
| chr6 | 93369966 | 93371230 |
| chr10 | 51418180 | 51419880 |
| chr2 | 59366429 | 59368588 |
| chr1 | 191993286 | 191996229 |
| chr20 | 25779141 | 25780191 |
| chr12 | 85638653 | 85641502 |
| chr14 | 27686001 | 27689137 |
| chr2 | 192952775 | 192953895 |
| chr2 | 163941952 | 163943142 |
| chr13 | 83430055 | 83432668 |
| chr15 | 92886690 | 92890168 |
| chr13 | 71725997 | 71727932 |
| chr5 | 127693699 | 127695277 |
| chr10 | 119596700 | 119598562 |
| chr12 | 16125431 | 16127179 |
| chr11 | 107041709 | 107044194 |
| chr2 | 56930289 | 56933228 |
| chr20 | 29394464 | 29396758 |
| chr12 | 57424104 | 57426812 |
| chr16 | 32325073 | 32326303 |
| chr8 | 110614989 | 110617064 |
| chr1 | 72124768 | 72126535 |
| chr12 | 85149123 | 85151818 |
| chr6 | 56637375 | 56639565 |
| chr10 | 57297141 | 57298857 |
| chr10 | 17226863 | 17228787 |
| chr7 | 84226358 | 84228568 |
| chr10 | 95064580 | 95066869 |
| chr13 | 19137349 | 19138843 |
| chr1 | 100519849 | 100522168 |
| chr10 | 71074771 | 71077102 |
| chr6 | 79492444 | 79494800 |

| | | |
|---|---|---|
| chr7 | 75184242 | 75187626 |
| chr15 | 82223559 | 82226470 |
| chr2 | 102049411 | 102050846 |
| chr12 | 76026017 | 76030378 |
| chr12 | 104286323 | 104288014 |
| chr9 | 73073597 | 73074911 |
| chr2 | 169463109 | 169465750 |
| chr14 | 59581773 | 59583885 |
| chr12 | 77672903 | 77674652 |
| chr1 | 196032826 | 196035335 |
| chr2 | 225324685 | 225327188 |
| chr14 | 94800233 | 94801760 |
| chr5 | 27486904 | 27490001 |
| chr6 | 71375437 | 71378359 |
| chr5 | 114184805 | 114186536 |
| chr4 | 35120303 | 35121906 |
| chr1 | 117067215 | 117068425 |
| chr2 | 46697939 | 46701281 |
| chr14 | 35761307 | 35763620 |
| chr22 | 12286427 | 12288160 |
| chr1 | 149409775 | 149411810 |
| chr1 | 224458653 | 224460344 |
| chr1 | 61452684 | 61455008 |
| chr7 | 65772933 | 65774625 |
| chr10 | 57762282 | 57765006 |
| chr7 | 75342039 | 75344090 |
| chr12 | 90159378 | 90160648 |
| chr3 | 104713601 | 104716007 |
| chr13 | 81512564 | 81514235 |
| chr1 | 83390215 | 83391950 |
| chr4 | 138447777 | 138449445 |
| chr1 | 86969042 | 86970811 |
| chr6 | 125659742 | 125661171 |
| chr14 | 26775098 | 26776549 |
| chr10 | 87362980 | 87365042 |
| chr1 | 67698213 | 67700082 |
| chr5 | 114697539 | 114699872 |
| chr1 | 102126249 | 102127689 |
| chr14 | 102137576 | 102140337 |
| chr10 | 102641708 | 102644164 |
| chr14 | 49189071 | 49191673 |
| chr2 | 55014462 | 55015700 |
| chr6 | 94018751 | 94019763 |
| chr1 | 174497663 | 174499616 |
| chr3 | 80437791 | 80439058 |
| chr3 | 83921646 | 83922778 |
| chr12 | 27651554 | 27654177 |
| chr2 | 51242820 | 51245229 |
| chr4 | 34890301 | 34891527 |

| | | |
|---|---|---|
| chr2 | 81011697 | 81013048 |
| chr15_KI270852v1_ | 257872 | 259638 |
| chr9 | 40989070 | 40992619 |
| chr2 | 83075303 | 83077861 |
| chr1 | 194409790 | 194412170 |
| chr12 | 57519598 | 57521594 |
| chrUn_GL000195v1 | 167733 | 169125 |
| chr15 | 40037589 | 40038831 |
| chr4 | 35811368 | 35812718 |
| chr9 | 41120243 | 41123566 |
| chr7 | 80505596 | 80508437 |
| chr5 | 165457174 | 165458649 |
| chr2 | 178734082 | 178737506 |
| chr7 | 76358901 | 76360050 |
| chr1 | 61850995 | 61854399 |
| chr10 | 50502290 | 50503771 |
| chr7 | 79208873 | 79211627 |
| chr1 | 103460830 | 103463246 |
| chr7 | 81564115 | 81565973 |
| chr20 | 47897950 | 47900036 |
| chr1 | 178852182 | 178854238 |
| chr10_GL383545v1_ | 66679 | 69267 |
| chr14 | 60643609 | 60645419 |
| chr5 | 17631482 | 17633390 |
| chr10 | 66283634 | 66286832 |
| chr6 | 90184540 | 90186911 |
| chr2 | 191942960 | 191945349 |
| chr1 | 195759933 | 195763090 |
| chr21 | 26715771 | 26718403 |
| chr3 | 110099684 | 110101862 |
| chr7 | 94428196 | 94433485 |
| chrX | 57715733 | 57716851 |
| chr6 | 123316318 | 123318019 |
| chr10 | 50937338 | 50938785 |
| chr14 | 82838119 | 82840434 |
| chr9 | 66121017 | 66123256 |
| chr5 | 41008872 | 41010518 |
| chr6 | 95141051 | 95143098 |
| chr14 | 19359619 | 19363731 |
| chr11 | 34266778 | 34268169 |
| chr4 | 125313139 | 125315327 |
| chr1 | 184506991 | 184509090 |
| chr12 | 74773565 | 74775218 |
| chr14 | 48709598 | 48711504 |
| chr3 | 63458827 | 63460111 |
| chr3 | 5857785 | 5859729 |
| chr12 | 58625572 | 58627215 |
| chr14 | 82281251 | 82283125 |
| chr2 | 68514662 | 68516823 |

| | | |
|---|---|---|
| chr12 | 83794294 | 83795340 |
| chr16 | 70130773 | 70132422 |
| chr10 | 86469939 | 86472057 |
| chr14 | 63373714 | 63376273 |
| chr4 | 29067028 | 29068850 |
| chr11 | 97106188 | 97107330 |
| chr10 | 76285540 | 76288140 |
| chr10 | 38391906 | 38394356 |
| chr4 | 36147432 | 36150011 |
| chr7 | 7664062 | 7665780 |
| chr2 | 52086357 | 52088659 |
| chr4 | 46439474 | 46441461 |
| chr4 | 57996176 | 57998272 |
| chr1 | 215368205 | 215369762 |
| chr12 | 75345450 | 75348744 |
| chr18 | 14226260 | 14228208 |
| chr1 | 44728934 | 44731927 |
| chr7 | 50919073 | 50921080 |
| chr11 | 108811211 | 108813069 |
| chr12 | 81556033 | 81558479 |
| chr15 | 34094614 | 34096247 |
| chr7 | 82168794 | 82170656 |
| chr7 | 84527808 | 84530833 |
| chr5 | 79555154 | 79557284 |
| chr6 | 32175695 | 32178797 |
| chr10 | 119824977 | 119827032 |
| chr21 | 9935582 | 9938225 |
| chr12 | 43406898 | 43409135 |
| chr12 | 72543414 | 72546093 |
| chr5 | 87666501 | 87668930 |
| chr9 | 67889541 | 67891469 |
| chr13 | 65654962 | 65657910 |
| chr6 | 13711770 | 13713403 |
| chr5 | 138429474 | 138432027 |
| chr5 | 34193263 | 34194540 |
| chr12 | 84159077 | 84160653 |
| chr17 | 81719504 | 81721246 |
| chr6 | 127034999 | 127036902 |
| chr7 | 76783758 | 76785284 |
| chr6 | 126131505 | 126132584 |
| chr2 | 27991899 | 27992936 |
| chr9 | 32835919 | 32837898 |
| chr16_KI270728v1_ | 1470629 | 1472636 |
| chr14 | 47611566 | 47614335 |
| chr13 | 89484146 | 89486540 |
| chr3 | 21062837 | 21065450 |
| chr2 | 51353568 | 51355534 |
| chr14 | 52720076 | 52721480 |
| chr8 | 105878937 | 105880149 |

| | | |
|---|---|---|
| chr14 | 27723921 | 27725227 |
| chr1 | 73674634 | 73676768 |
| chr12 | 100215490 | 100217175 |
| chr6 | 85542791 | 85544622 |
| chr1 | 186482352 | 186484350 |
| chr17 | 65527882 | 65530001 |
| chr1 | 194692455 | 194696264 |
| chr19 | 6373050 | 6374529 |
| chr12 | 110402968 | 110405259 |
| chrUn_GL000220v1 | 10325 | 12836 |
| chr11 | 23988088 | 23989987 |
| chr11 | 105892128 | 105893976 |
| chr3 | 24626531 | 24628100 |
| chr3 | 171393652 | 171394710 |
| chr14 | 86940310 | 86943708 |
| chr7 | 13982326 | 13983450 |
| chr16 | 35384988 | 35386883 |
| chr1 | 23844619 | 23845735 |
| chr7 | 12685730 | 12688107 |
| chr4 | 147782818 | 147784178 |
| chr7 | 105695104 | 105696367 |
| chr19 | 1303173 | 1304177 |
| chr2 | 209036262 | 209038980 |
| chr6 | 75816887 | 75819735 |
| chr10 | 4785537 | 4787893 |
| chr14 | 18661631 | 18662838 |
| chr7 | 93526630 | 93527907 |
| chr4 | 64370326 | 64371991 |
| chr12 | 84488126 | 84489947 |
| chr2 | 85432704 | 85435376 |
| chr15 | 37817821 | 37819849 |
| chr20 | 29121899 | 29125999 |
| chr1 | 221572466 | 221574397 |
| chr14 | 78245560 | 78246604 |
| chr8 | 91188606 | 91191683 |
| chr5 | 161487596 | 161490306 |
| chr2 | 66252121 | 66253919 |
| chr8 | 86836188 | 86838157 |
| chr1 | 104689434 | 104690870 |
| chr8 | 82718562 | 82720753 |
| chr2 | 166478694 | 166481220 |
| chr10 | 56589270 | 56591285 |
| chr13 | 62674422 | 62676583 |
| chr18 | 68842657 | 68846635 |
| chr13 | 37654754 | 37656948 |
| chr3 | 94896969 | 94898877 |
| chr20 | 25781450 | 25785275 |
| chr19 | 16777841 | 16779812 |
| chr6 | 10923693 | 10925274 |

| | | |
|---|---|---|
| chr14 | 68089202 | 68090623 |
| chr3 | 168574583 | 168577814 |
| chr1 | 190309064 | 190312031 |
| chr6 | 79779433 | 79780997 |
| chr6 | 141400712 | 141403014 |
| chr14 | 87866523 | 87867701 |
| chr12 | 65239324 | 65241010 |
| chr20 | 7144111 | 7146548 |
| chr12 | 66023888 | 66026336 |
| chr14 | 84841633 | 84844399 |
| chr2 | 229086469 | 229088622 |
| chr13 | 71872266 | 71874187 |
| chr3 | 174093382 | 174095462 |
| chr4 | 142839437 | 142841412 |
| chr14 | 39113854 | 39115614 |
| chr2 | 91560597 | 91564154 |
| chr5 | 59177380 | 59180237 |
| chr11 | 31090507 | 31092130 |
| chr20 | 29061910 | 29063984 |
| chr5 | 22324035 | 22325187 |
| chr1 | 15882279 | 15885962 |
| chr16_KI270728v1_ | 1000998 | 1002475 |
| chr1 | 193564237 | 193565430 |
| chr6 | 49798134 | 49800011 |
| chr8 | 64058326 | 64059334 |
| chr1 | 148602608 | 148605127 |
| chr14 | 31563630 | 31565851 |
| chr7 | 7639748 | 7642542 |
| chr10 | 56303942 | 56305950 |
| chr1 | 216029351 | 216031502 |
| chr1 | 238968045 | 238969915 |
| chr15 | 47599898 | 47602768 |
| chr1 | 120559191 | 120561337 |
| chr13 | 105461734 | 105464230 |
| chr11 | 67730546 | 67731682 |
| chr13 | 105616042 | 105617851 |
| chr6 | 101080164 | 101082060 |
| chr1 | 97114260 | 97116400 |
| chr4 | 171455913 | 171457376 |
| chr8 | 91198266 | 91199703 |
| chr2 | 211926789 | 211929094 |
| chr15 | 35205054 | 35207151 |
| chr14 | 41893742 | 41895366 |
| chr10 | 67113353 | 67116354 |
| chr9 | 31424780 | 31426243 |
| chr13 | 105508234 | 105512348 |
| chr13 | 104021492 | 104024086 |
| chr14 | 24006304 | 24007922 |
| chr8 | 35898511 | 35900280 |

| | | |
|---|---|---|
| chr2 | 227113407 | 227114970 |
| chr10 | 68874659 | 68876343 |
| chr5 | 177431409 | 177434081 |
| chr12 | 43547842 | 43550553 |
| chr14 | 47977401 | 47979982 |
| chr1 | 81943880 | 81947170 |
| chr1 | 111761775 | 111763344 |
| chr10 | 123157925 | 123159561 |
| chr2 | 63034084 | 63038012 |
| chr2 | 189048680 | 189050388 |
| chr11 | 118496512 | 118497511 |
| chr7 | 57834866 | 57839769 |
| chr15 | 20275338 | 20277601 |
| chr16 | 76425894 | 76428552 |
| chr4 | 150858546 | 150860817 |
| chr2 | 50964174 | 50966212 |
| chr7 | 79810010 | 79811831 |
| chr12 | 46764148 | 46767068 |
| chr3 | 173836452 | 173837917 |
| chr10 | 108359757 | 108362564 |
| chr11 | 22824775 | 22828570 |
| chr7 | 8953068 | 8957171 |
| chr2 | 78876227 | 78878781 |
| chr10 | 130697401 | 130698983 |
| chr15 | 20649359 | 20651716 |
| chr5 | 847098 | 848156 |
| chr7 | 81741749 | 81743339 |
| chr1 | 144336909 | 144339167 |
| chr21 | 10431832 | 10433604 |
| chr10 | 95445392 | 95448060 |
| chr16 | 76219042 | 76221860 |
| chr6 | 120127775 | 120129978 |
| chr2 | 196658030 | 196659520 |
| chr1 | 104530231 | 104532673 |
| chr20 | 39795477 | 39797141 |
| chr12 | 80845006 | 80847762 |
| chr4 | 83649145 | 83650974 |
| chr20 | 17970695 | 17972800 |
| chr5 | 21599428 | 21600939 |
| chr15 | 97847431 | 97849739 |
| chr14 | 40480106 | 40482148 |
| chr1 | 62190046 | 62191645 |
| chr7 | 89903871 | 89905771 |
| chr1 | 207357640 | 207360657 |
| chr1 | 145123375 | 145124943 |
| chr1 | 186395690 | 186400290 |
| chr14 | 26512773 | 26515905 |
| chr3 | 75669305 | 75670632 |
| chr15 | 53164851 | 53167557 |

| | | |
|---|---|---|
| chr8 | 113918421 | 113919899 |
| chr12 | 70362501 | 70365134 |
| chr10 | 45129790 | 45132050 |
| chr1 | 98315048 | 98317213 |
| chr2 | 162822980 | 162825026 |
| chr16 | 76317132 | 76319525 |
| chr1 | 71229158 | 71231223 |
| chr2 | 200383526 | 200386011 |
| chr20 | 7392466 | 7393561 |
| chr2 | 188083099 | 188086154 |
| chr4 | 35110085 | 35111571 |
| chr1 | 105185667 | 105189658 |
| chr2 | 194401002 | 194402766 |
| chr20 | 30423977 | 30425487 |
| chr10 | 4662517 | 4664893 |
| chr11 | 35084985 | 35086239 |
| chr14 | 39022932 | 39024880 |
| chr14 | 100217622 | 100219961 |
| chr11 | 124669140 | 124670806 |
| chr14 | 19710670 | 19713199 |
| chr1 | 97086305 | 97088102 |
| chr2 | 208344089 | 208345762 |
| chr16 | 30063533 | 30066889 |
| chr10 | 65375961 | 65378114 |
| chr10 | 65744565 | 65746899 |
| chr2 | 131565072 | 131566121 |
| chr8 | 80237577 | 80240106 |
| chr10 | 88597829 | 88600267 |
| chr10 | 56853761 | 56855259 |
| chr1 | 106640209 | 106642875 |
| chr1 | 239320664 | 239322536 |
| chr6 | 72925650 | 72927919 |
| chr13 | 71551796 | 71554635 |
| chr10 | 22002321 | 22004494 |
| chr7 | 88762973 | 88766015 |
| chr1 | 85993386 | 85995006 |
| chr14 | 27856878 | 27858443 |
| chr14 | 31632428 | 31634556 |
| chr2 | 192735775 | 192737075 |
| chr1 | 105537499 | 105539422 |
| chr2 | 182904026 | 182905684 |
| chr2 | 61537326 | 61539845 |
| chr1 | 84076885 | 84080633 |
| chr4 | 47469030 | 47471916 |
| chr12 | 62313572 | 62315830 |
| chr6 | 92347991 | 92349855 |
| chr2 | 197480635 | 197482257 |
| chr1 | 206227755 | 206229722 |
| chr7 | 86035597 | 86037334 |

| | | |
|---|---|---|
| chr6 | 49908567 | 49910458 |
| chr2 | 205808198 | 205809576 |
| chr15_KI270852v1_ | 281036 | 283100 |
| chr2 | 57599504 | 57602127 |
| chr12 | 22357502 | 22359370 |
| chrUn_GL000195v1 | 126322 | 128662 |
| chr1 | 103005496 | 103006884 |
| chr5 | 173615742 | 173617305 |
| chr8 | 51817144 | 51820012 |
| chr4 | 59093683 | 59095521 |
| chr20 | 30418956 | 30421646 |
| chr1 | 163729188 | 163731517 |
| chr2 | 57674926 | 57676996 |
| chr14 | 44016241 | 44018342 |
| chr20 | 22627797 | 22630353 |
| chr2 | 82965245 | 82967552 |
| chr17 | 65883847 | 65888354 |
| chr4 | 176749124 | 176751057 |
| chr6 | 87510845 | 87512804 |
| chr11 | 96135628 | 96137567 |
| chr7 | 92690911 | 92694227 |
| chr7 | 82768179 | 82771149 |
| chr14 | 85272111 | 85274652 |
| chr2 | 84543258 | 84545222 |
| chr6 | 70450943 | 70452756 |
| chr6 | 153291113 | 153292568 |
| chr4 | 34365570 | 34367535 |
| chr4 | 91869002 | 91871550 |
| chr9 | 40959824 | 40962952 |
| chr10 | 112497310 | 112499169 |
| chr3 | 87620155 | 87622292 |
| chr7 | 90209743 | 90212504 |
| chr14 | 99229631 | 99232353 |
| chr3 | 105962903 | 105965250 |
| chr12 | 1078368 | 1080349 |
| chr20 | 8337616 | 8339097 |
| chr14 | 103079539 | 103081408 |
| chr1 | 195300947 | 195302730 |
| chr14 | 32958634 | 32961153 |
| chr6 | 50520015 | 50522029 |
| chr11 | 95840144 | 95842638 |
| chr20 | 14535228 | 14536772 |
| chr4 | 32701801 | 32703371 |
| chr10 | 91996713 | 91998147 |
| chr1 | 102470991 | 102472873 |
| chr14 | 67005771 | 67007724 |
| chr14 | 41593997 | 41595340 |
| chr3 | 169616217 | 169618237 |
| chr5 | 51535156 | 51536875 |

| | | |
|---|---|---|
| chr20 | 29358755 | 29360524 |
| chr20 | 6105919 | 6107374 |
| chr2 | 51597783 | 51599707 |
| chr11 | 74630652 | 74633186 |
| chr1 | 98478966 | 98480582 |
| chr2 | 29629552 | 29631036 |
| chr9 | 100217417 | 100219211 |
| chr2 | 194517319 | 194519244 |
| chr2 | 166822212 | 166824883 |
| chr2 | 179965350 | 179967267 |
| chr14 | 41071945 | 41074350 |
| chr2 | 63502820 | 63505870 |
| chr2 | 177489435 | 177490814 |
| chr1 | 103283522 | 103285856 |
| chr2 | 44836178 | 44837362 |
| chr16 | 75981572 | 75985103 |
| chr9 | 110401432 | 110403174 |
| chr19 | 36112294 | 36116045 |
| chr20 | 29495765 | 29500283 |
| chr11 | 102827612 | 102829317 |
| chr21 | 9885912 | 9888472 |
| chr1 | 218472667 | 218477001 |
| chr12 | 17748500 | 17749970 |
| chr6 | 99979453 | 99981435 |
| chr12 | 83112060 | 83114386 |
| chr4 | 30162891 | 30164411 |
| chr10 | 49057712 | 49060135 |
| chr7 | 82395912 | 82397157 |
| chr20 | 26227901 | 26230926 |
| chr1 | 219159672 | 219163075 |
| chr14 | 83913588 | 83915400 |
| chrUn_GL000219v1 | 104111 | 105729 |
| chr2 | 186991977 | 186994102 |
| chr12 | 84675737 | 84678378 |
| chr14 | 99908985 | 99910820 |
| chr9 | 43282532 | 43283562 |
| chr12 | 105167978 | 105169611 |
| chr2 | 66603578 | 66604887 |
| chr5_KI270897v1_a | 747495 | 749550 |
| chr2 | 85749245 | 85751857 |
| chr1 | 193735109 | 193736803 |
| chr10 | 115169942 | 115171982 |
| chr7 | 81287519 | 81289731 |
| chr16 | 53222807 | 53225215 |
| chr2 | 195319686 | 195320902 |
| chr2 | 83541325 | 83543184 |
| chr10 | 63719391 | 63720756 |
| chr1 | 190750384 | 190752396 |
| chr8 | 93739432 | 93741657 |

| | | |
|---|---|---|
| chr8 | 82691851 | 82695723 |
| chr7 | 57729606 | 57732099 |
| chr12 | 81267048 | 81269814 |
| chr14 | 100334346 | 100335726 |
| chr1 | 215901182 | 215902652 |
| chr21 | 10365592 | 10369043 |
| chr15 | 36413655 | 36415396 |
| chr7 | 85597497 | 85599913 |
| chr12 | 114680163 | 114682525 |
| chr16 | 34083680 | 34085348 |
| chr5 | 87286480 | 87288379 |
| chr4 | 174751160 | 174753312 |
| chr12 | 88270134 | 88271935 |
| chr14 | 33260880 | 33262743 |
| chr2 | 52584988 | 52587306 |
| chr6 | 140710529 | 140712543 |
| chr1 | 81905429 | 81907630 |
| chr2 | 80228061 | 80229072 |
| chr9 | 10273728 | 10274995 |
| chr7 | 89835783 | 89837917 |
| chr10 | 55501028 | 55503742 |
| chr20 | 29513714 | 29515984 |
| chr5 | 117024522 | 117026125 |
| chr11 | 22812556 | 22814916 |
| chr6 | 152231287 | 152233294 |
| chr2 | 186574877 | 186576565 |
| chr20 | 54146045 | 54147390 |
| chr10 | 36883693 | 36885653 |
| chr14 | 19665579 | 19667744 |
| chr2 | 148508716 | 148510915 |
| chr7 | 84665651 | 84668132 |
| chr12 | 25293449 | 25296294 |
| chr6 | 81303637 | 81307228 |
| chr20 | 17476172 | 17477472 |
| chr14 | 88626255 | 88628065 |
| chr14 | 24335914 | 24337895 |
| chr14 | 27244093 | 27245724 |
| chr9 | 65795785 | 65798483 |
| chr16 | 28843844 | 28847345 |
| chr14 | 86708258 | 86710615 |
| chr17 | 62967083 | 62969147 |
| chr7 | 88170326 | 88172487 |
| chr7 | 86664093 | 86665538 |
| chr12 | 114740627 | 114742584 |
| chr7 | 11008900 | 11011074 |
| chr20 | 542994 | 544961 |
| chr14 | 38737443 | 38739793 |
| chr6 | 20999898 | 21001831 |
| chr10 | 91384569 | 91387779 |

| | | |
|---|---|---|
| chr7 | 25843337 | 25844752 |
| chr18 | 68832455 | 68835705 |
| chr14 | 37498421 | 37501862 |
| chr3 | 77860395 | 77862016 |
| chr10 | 120320993 | 120323416 |
| chr1 | 149103050 | 149104588 |
| chr15 | 36954094 | 36955622 |
| chr7 | 84131040 | 84133014 |
| chr22 | 10700695 | 10704360 |
| chr6 | 56482918 | 56485715 |
| chr8 | 96942766 | 96945234 |
| chr13 | 61578855 | 61580425 |
| chr8 | 81013639 | 81015655 |
| chr1 | 197768631 | 197771920 |
| chr6 | 80942980 | 80945293 |
| chr16 | 34073526 | 34076946 |
| chr14 | 32426443 | 32430316 |
| chr1 | 99273239 | 99274265 |
| chr14 | 29922946 | 29926543 |
| chr10 | 115121333 | 115123322 |
| chr5 | 17920871 | 17922055 |
| chr10 | 28179217 | 28180709 |
| chr3 | 85420479 | 85422171 |
| chr4 | 29990275 | 29992043 |
| chr5 | 12403431 | 12404515 |
| chr11 | 107010637 | 107012197 |
| chr1 | 100423669 | 100426219 |
| chr7 | 146911658 | 146913140 |
| chr16 | 89910373 | 89911896 |
| chr12 | 84872655 | 84876958 |
| chr7 | 90244602 | 90245780 |
| chr6 | 26198969 | 26201067 |
| chr2 | 34786111 | 34787597 |
| chr10 | 17459196 | 17461060 |
| chr14 | 69765419 | 69769973 |
| chr11 | 98114528 | 98116729 |
| chr12 | 80316843 | 80319979 |
| chr2 | 210166920 | 210167995 |
| chr16 | 53670874 | 53674318 |
| chr2 | 38075165 | 38076841 |
| chr10 | 118007898 | 118009609 |
| chr12 | 63046649 | 63047769 |
| chr15 | 57133744 | 57136638 |
| chr2 | 103160196 | 103162777 |
| chr13 | 82077601 | 82080728 |
| chr6 | 70738685 | 70739939 |
| chr10 | 114144187 | 114145988 |
| chrX | 56768440 | 56771105 |
| chr6 | 61811336 | 61813526 |

| | | |
|---|---|---|
| chr7 | 7241392 | 7243102 |
| chr8 | 111606229 | 111608755 |
| chr7 | 121944778 | 121946143 |
| chr8 | 76225443 | 76231829 |
| chr1 | 102569015 | 102572057 |
| chr7 | 8212189 | 8214149 |
| chr9 | 41416682 | 41419811 |
| chr2 | 213250296 | 213253246 |
| chr2 | 8677293 | 8679852 |
| chr12 | 80648587 | 80649753 |
| chr6 | 125017092 | 125019044 |
| chr8 | 133806373 | 133808594 |
| chr2 | 60782376 | 60784418 |
| chr12 | 120290424 | 120293926 |
| chr5 | 126137380 | 126140454 |
| chr2 | 82110555 | 82112456 |
| chr10 | 52061336 | 52063419 |
| chr7 | 100948576 | 100952255 |
| chr2 | 183651462 | 183653975 |
| chr7 | 62385933 | 62387366 |
| chr13 | 91346226 | 91348639 |
| chr12 | 84690049 | 84692194 |
| chr13 | 96314017 | 96316189 |
| chr12 | 65311480 | 65316402 |
| chr6 | 119975938 | 119978852 |
| chr14 | 21475178 | 21477836 |
| chr7 | 83974578 | 83978024 |
| chr2 | 85353637 | 85355267 |
| chr13 | 67301237 | 67303440 |
| chr17 | 63338546 | 63340406 |
| chr2 | 183531760 | 183533551 |
| chr14 | 74549864 | 74551438 |
| chr2 | 169582885 | 169584762 |
| chr14 | 41143695 | 41144743 |
| chr8 | 134239284 | 134241248 |
| chr14 | 21457774 | 21460417 |
| chr3 | 34130577 | 34132847 |
| chr1 | 72101027 | 72103178 |
| chr10 | 87262319 | 87264033 |
| chr11 | 74801522 | 74802709 |
| chr19 | 40011755 | 40013267 |
| chr21 | 5220329 | 5222523 |
| chr6 | 119429051 | 119431011 |
| chr3 | 81914433 | 81916551 |
| chr15 | 20410475 | 20411924 |
| chr20 | 29867976 | 29869446 |
| chr1 | 86501700 | 86503445 |
| chr1 | 104480369 | 104482477 |
| chr9 | 40974950 | 40977291 |

| | | |
|---|---|---|
| chr15 | 63644348 | 63646808 |
| chr14 | 53813604 | 53815260 |
| chr1 | 172443376 | 172445248 |
| chr1 | 172406632 | 172409461 |
| chr2 | 186028813 | 186030787 |
| chr2 | 59072731 | 59074794 |
| chr10 | 21462974 | 21465271 |
| chr14 | 29547446 | 29549893 |
| chr12 | 44222102 | 44224841 |
| chr1 | 103974501 | 103976987 |
| chr1 | 86648781 | 86649829 |
| chr22 | 15816237 | 15819178 |
| chr2 | 214992446 | 214994038 |
| chr2 | 80265031 | 80267779 |
| chr6 | 14642893 | 14644598 |
| chr4 | 74997829 | 74999750 |
| chr11 | 104485960 | 104487862 |
| chr1 | 201954736 | 201957600 |
| chr6 | 19753192 | 19756705 |
| chr4 | 34957930 | 34962274 |
| chr13 | 71733750 | 71736315 |
| chr10 | 55206878 | 55208782 |
| chr4 | 156389424 | 156392095 |
| chr10 | 54905271 | 54906482 |
| chr16 | 30656974 | 30659642 |
| chr12 | 63770988 | 63772485 |
| chr12 | 42882081 | 42883676 |
| chr1 | 224966761 | 224969293 |
| chr2 | 79729995 | 79731800 |
| chr10 | 64481024 | 64482752 |
| chr12 | 84559940 | 84564782 |
| chr7 | 82012410 | 82014464 |
| chr6 | 79809118 | 79810747 |
| chr14 | 42423763 | 42426144 |
| chr12 | 74256420 | 74259452 |
| chr1 | 125174672 | 125177325 |
| chr4 | 31090098 | 31092463 |
| chr1 | 788415 | 792292 |
| chr11 | 107961188 | 107962571 |
| chr2 | 212290511 | 212293947 |
| chr2 | 184313488 | 184315271 |
| chr2 | 91572003 | 91573334 |
| chr13 | 72295016 | 72297192 |
| chr3 | 76684961 | 76687568 |
| chr5 | 153112135 | 153113809 |
| chr18 | 66463166 | 66464327 |
| chr14 | 57490287 | 57491807 |
| chr9 | 72995805 | 72998204 |
| chr2 | 54103359 | 54105390 |

| | | |
|---|---|---|
| chr5 | 123261077 | 123262379 |
| chr11 | 109127019 | 109128176 |
| chr2 | 53770161 | 53772574 |
| chr5 | 3677733 | 3679552 |
| chr14 | 38023295 | 38024735 |
| chr5 | 28093370 | 28095898 |
| chr11 | 26742448 | 26745597 |
| chr4 | 156622630 | 156625270 |
| chr2 | 195406057 | 195408989 |
| chr10 | 91552546 | 91554450 |
| chr16 | 34760267 | 34762221 |
| chr5 | 17597892 | 17599743 |
| chr13 | 69700908 | 69702801 |
| chr7 | 81182361 | 81184405 |
| chr6 | 141116936 | 141119610 |
| chr12 | 12660770 | 12662391 |
| chr2 | 104159691 | 104161707 |
| chr5 | 113190453 | 113191962 |
| chr7 | 13030855 | 13033170 |
| chr6 | 80302031 | 80303468 |
| chr12 | 77820906 | 77822844 |
| chr6 | 16073098 | 16075618 |
| chr1 | 104971149 | 104973324 |
| chr2 | 94691403 | 94693048 |
| chr1 | 237589880 | 237591628 |
| chr1 | 210372717 | 210374627 |
| chr10 | 102150993 | 102153186 |
| chr22 | 11820618 | 11821830 |
| chr11 | 103529674 | 103532291 |
| chr1 | 88142684 | 88144684 |
| chr14 | 97146565 | 97147591 |
| chr17 | 63842102 | 63844318 |
| chr11 | 77815352 | 77817536 |
| chr4 | 32352256 | 32355411 |
| chr14 | 26235544 | 26237613 |
| chr5 | 131634853 | 131636967 |
| chr4 | 116289654 | 116291236 |
| chr15 | 37794247 | 37796467 |
| chr6 | 22259477 | 22261531 |
| chr1 | 95231944 | 95234983 |
| chr6 | 149349996 | 149352459 |
| chr1 | 91642829 | 91644776 |
| chr1 | 196026743 | 196028560 |
| chr22 | 15728532 | 15730657 |
| chr10 | 36181842 | 36183226 |
| chr7 | 57490125 | 57491257 |
| chr7 | 86327598 | 86330331 |
| chr7 | 125669164 | 125671219 |
| chr11 | 23142637 | 23144798 |

| | | |
|---|---|---|
| chr1 | 8027248 | 8028974 |
| chr10 | 68979271 | 68981560 |
| chr2 | 206764923 | 206765958 |
| chr4 | 75513642 | 75515201 |
| chr1 | 218338401 | 218341014 |
| chr2 | 187457591 | 187458857 |
| chr10 | 43454942 | 43456500 |
| chr1 | 99221370 | 99224977 |
| chr8 | 80071339 | 80073267 |
| chr1 | 190050091 | 190051969 |
| chr2 | 162416755 | 162418724 |
| chr1 | 89913252 | 89914619 |
| chr14 | 81034235 | 81035918 |
| chr8_KI270813v1_a | 25649 | 27562 |
| chr12 | 105644224 | 105645610 |
| chr1 | 83235508 | 83237346 |
| chr12 | 71616210 | 71618190 |
| chr14 | 98072448 | 98074948 |
| chr1 | 117685883 | 117687485 |
| chr6 | 16712521 | 16713993 |
| chr16 | 34756224 | 34759975 |
| chr7 | 124929357 | 124931080 |
| chr12 | 84598603 | 84601860 |
| chr10 | 90111270 | 90113317 |
| chr14 | 87174140 | 87177013 |
| chr4 | 132310919 | 132312580 |
| chr7 | 58072022 | 58074121 |
| chr12 | 81149506 | 81151401 |
| chr2 | 185577299 | 185579845 |
| chr12 | 9991855 | 9993394 |
| chr15_KI270905v1_ | 629717 | 632212 |
| chr1 | 190444409 | 190446187 |
| chr12 | 82898584 | 82901125 |
| chr14 | 24779734 | 24781326 |
| chr16 | 70563129 | 70564401 |
| chr2 | 213106503 | 213108333 |
| chr19 | 41362905 | 41364782 |
| chr2 | 59727794 | 59729644 |
| chr2 | 52926085 | 52928082 |
| chr7 | 81114747 | 81116826 |
| chr11 | 23838184 | 23840434 |
| chr11 | 96566453 | 96568249 |
| chrX | 32341780 | 32342955 |
| chr2 | 184907448 | 184909934 |
| chr3 | 181053811 | 181056914 |
| chr14 | 46998534 | 47001490 |
| chr1 | 104265863 | 104267811 |
| chr1 | 90114001 | 90116691 |
| chr10 | 116278702 | 116280568 |

| | | |
|---|---|---|
| chr9 | 13449815 | 13451595 |
| chr2 | 211404974 | 211406426 |
| chr2 | 35634567 | 35637295 |
| chr14 | 26069901 | 26072404 |
| chr15 | 100860657 | 100862007 |
| chr2 | 50423972 | 50426639 |
| chr8 | 113162435 | 113164169 |
| chr20 | 29745902 | 29748346 |
| chr9 | 66097589 | 66099512 |
| chr1 | 78676049 | 78677595 |
| chr2 | 76190600 | 76192753 |
| chr5 | 111550066 | 111552707 |
| chr14 | 49012689 | 49015265 |
| chr11 | 40007382 | 40009133 |
| chr9 | 41187966 | 41189554 |
| chr7 | 53808163 | 53810011 |
| chr2 | 63193559 | 63195680 |
| chr3 | 85430849 | 85434815 |
| chr2 | 79947869 | 79950360 |
| chr2 | 192173492 | 192175231 |
| chr2 | 198350086 | 198351634 |
| chr12 | 96825735 | 96829517 |
| chr7 | 86489224 | 86491273 |
| chr3 | 29334523 | 29336580 |
| chr14 | 27048276 | 27051874 |
| chr6 | 64274418 | 64276755 |
| chr10 | 94406455 | 94409552 |
| chr7 | 84656308 | 84659707 |
| chr2 | 193490263 | 193492245 |
| chrX | 46470136 | 46471136 |
| chr2 | 207097978 | 207099864 |
| chr12_GL877875v1_ | 20365 | 22200 |
| chr2 | 159113736 | 159115422 |
| chr14 | 99389501 | 99393008 |
| chr1 | 150628300 | 150630958 |
| chr3 | 112166358 | 112168455 |
| chr5 | 9098408 | 9099627 |
| chr15 | 55402411 | 55404214 |
| chrY | 11331145 | 11333207 |
| chr14 | 41993883 | 41997255 |
| chr5 | 93161480 | 93164354 |
| chr1 | 216233653 | 216235650 |
| chr2 | 64981411 | 64983723 |
| chr11 | 88662794 | 88664135 |
| chr11 | 67984676 | 67985849 |
| chr14 | 19151819 | 19153014 |
| chr10 | 57312408 | 57314747 |
| chr14 | 44122176 | 44124216 |
| chr1 | 65298301 | 65299756 |

| | | |
|---|---|---|
| chr16 | 71807100 | 71810260 |
| chr1 | 189632076 | 189633643 |
| chr1 | 146471474 | 146472795 |
| chr2 | 187842305 | 187844592 |
| chr17 | 61316768 | 61318630 |
| chr4 | 31189894 | 31191770 |
| chr14 | 52541175 | 52543344 |
| chr2 | 233564021 | 233565699 |
| chr3 | 34535885 | 34537027 |
| chr1 | 90937433 | 90938875 |
| chr14 | 38768738 | 38772214 |
| chr2 | 122962939 | 122964336 |
| chr4 | 32267998 | 32269538 |
| chr4 | 102931223 | 102932973 |
| chr10 | 65158984 | 65162107 |
| chr10 | 49143727 | 49146857 |
| chr2 | 205858511 | 205860792 |
| chr7 | 83266455 | 83268957 |
| chr1 | 210051399 | 210053424 |
| chr12 | 84225510 | 84228013 |
| chr1 | 199774698 | 199776735 |
| chr2 | 57498955 | 57500492 |
| chr13 | 22852166 | 22853855 |
| chr4 | 138076179 | 138077290 |
| chr1 | 172397052 | 172398870 |
| chr1 | 247796241 | 247799227 |
| chr2 | 171046716 | 171049083 |
| chr15_KI270905v1_ | 724700 | 727492 |
| chr14 | 47894604 | 47896714 |
| chr15 | 66502657 | 66505997 |
| chr22 | 15783839 | 15785727 |
| chr7 | 86515697 | 86518370 |
| chr10 | 53385645 | 53388878 |
| chr9 | 17300967 | 17302516 |
| chr11 | 92702530 | 92704343 |
| chr1 | 225050904 | 225052261 |
| chr6 | 156782386 | 156784529 |
| chr14 | 98270268 | 98273046 |
| chr12 | 44106273 | 44108125 |
| chr4 | 13480072 | 13482997 |
| chr14 | 36875263 | 36877378 |
| chr6 | 49949916 | 49951874 |
| chr7 | 90151285 | 90153822 |
| chr8 | 122847557 | 122849265 |
| chr7 | 149628891 | 149630872 |
| chr1 | 170532084 | 170533913 |
| chr10 | 61704317 | 61707580 |
| chr22 | 15527039 | 15530614 |
| chr9 | 102922599 | 102924381 |

| | | |
|---|---|---|
| chr22 | 11583913 | 11586591 |
| chr8 | 90953757 | 90956921 |
| chr1 | 16613525 | 16616171 |
| chr7 | 89729931 | 89732674 |
| chr2 | 162343218 | 162345255 |
| chr2 | 100196731 | 100197981 |
| chr6 | 79631560 | 79634232 |
| chr10 | 70917271 | 70919039 |
| chr7 | 65240121 | 65241918 |
| chr4 | 144142372 | 144144222 |
| chr14 | 44957514 | 44959590 |
| chr2 | 40748350 | 40751119 |
| chr5 | 20533412 | 20534951 |
| chr14 | 37328409 | 37331368 |
| chr13 | 57710574 | 57712674 |
| chr10 | 88622601 | 88624048 |
| chr7 | 80307138 | 80309356 |
| chr14 | 32091869 | 32092976 |
| chr2 | 162346879 | 162349480 |
| chr7 | 89606048 | 89608310 |
| chr1 | 93732199 | 93734655 |
| chr10 | 62238923 | 62241677 |
| chr7 | 12964731 | 12967477 |
| chrUn_GL000195v1 | 133850 | 135930 |
| chr1 | 87494392 | 87497902 |
| chr10 | 12563859 | 12565355 |
| chr2 | 226028225 | 226033409 |
| chr7 | 90871995 | 90874087 |
| chr16 | 86039620 | 86040789 |
| chr9 | 62322817 | 62324884 |
| chr1 | 104636204 | 104638022 |
| chr6 | 95226761 | 95228403 |
| chr2 | 43943343 | 43945516 |
| chr1 | 96806449 | 96809158 |
| chr5 | 175217592 | 175218731 |
| chr1 | 101135837 | 101137928 |
| chr2 | 68255283 | 68257185 |
| chr6 | 63752778 | 63753982 |
| chr10 | 75724462 | 75726820 |
| chr3 | 31438487 | 31440394 |
| chr8 | 110976252 | 110978851 |
| chr1 | 192485743 | 192487120 |
| chr2 | 184744853 | 184746050 |
| chr4 | 34186555 | 34188020 |
| chr15_KI270905v1_ | 809680 | 811715 |
| chr1 | 159006749 | 159009337 |
| chr10 | 62383144 | 62384773 |
| chr13 | 82995530 | 82998113 |
| chr14 | 37726505 | 37727665 |

| | | |
|---|---|---|
| chr13 | 67198161 | 67199600 |
| chr14 | 40901204 | 40903202 |
| chr11 | 27705646 | 27708551 |
| chr10 | 88491588 | 88493799 |
| chr2 | 54980704 | 54982177 |
| chr6 | 14795467 | 14797196 |
| chr12 | 10523105 | 10525507 |
| chr21 | 10575883 | 10578779 |
| chr1 | 206199379 | 206202960 |
| chr7 | 136841993 | 136844081 |
| chr21 | 10400557 | 10403658 |
| chr21 | 13276851 | 13279337 |
| chr1 | 104879690 | 104881229 |
| chr13 | 34926348 | 34928082 |
| chr14 | 33388569 | 33390062 |
| chr3 | 169767853 | 169770575 |
| chr10 | 64148799 | 64150878 |
| chr16 | 34696918 | 34699591 |
| chr5 | 16159090 | 16162125 |
| chr1 | 50424919 | 50426151 |
| chr2 | 15165417 | 15167678 |
| chr2 | 162364238 | 162367905 |
| chr8 | 7918576 | 7920500 |
| chr2 | 144497690 | 144499556 |
| chr3 | 34993695 | 34995590 |
| chr1 | 218768009 | 218770027 |
| chr10 | 66826106 | 66828447 |
| chr7 | 63963355 | 63965163 |
| chr20 | 7828764 | 7830667 |
| chr10 | 113793640 | 113796388 |
| chr6 | 83597785 | 83600668 |
| chr10 | 89565563 | 89567735 |
| chr1 | 79160565 | 79162769 |
| chr6 | 75095626 | 75098708 |
| chr14 | 35883362 | 35885130 |
| chr7 | 85850014 | 85852544 |
| chr17 | 20338962 | 20341023 |
| chr16 | 34855617 | 34860156 |
| chr1 | 98457036 | 98459310 |
| chr10 | 115888612 | 115891360 |
| chr12 | 18563360 | 18564944 |
| chr1 | 120353910 | 120356407 |
| chr14 | 41950840 | 41952925 |
| chr14 | 49007382 | 49009150 |
| chr12 | 56314707 | 56316679 |
| chr20 | 58891815 | 58894225 |
| chr18 | 47830424 | 47832176 |
| chr7 | 77758562 | 77760999 |
| chr6 | 132233968 | 132236006 |

| | | |
|---|---|---|
| chr7 | 9819386 | 9821687 |
| chr9 | 67333990 | 67335690 |
| chr2 | 220844515 | 220847026 |
| chr3 | 76975298 | 76978714 |
| chr14 | 58299219 | 58302267 |
| chr15 | 47370826 | 47372904 |
| chr14 | 41296055 | 41298158 |
| chr15 | 48166671 | 48168815 |
| chr12 | 69568888 | 69571161 |
| chr9 | 63821348 | 63823354 |
| chr12 | 18294383 | 18296383 |
| chr1 | 93215552 | 93217856 |
| chr7 | 54091509 | 54093720 |
| chr2 | 37094695 | 37097526 |
| chr2 | 188746167 | 188749767 |
| chr1 | 189079242 | 189081696 |
| chr1 | 21891520 | 21892521 |
| chr9 | 72974034 | 72975318 |
| chr1 | 169472082 | 169473206 |
| chr1 | 217678227 | 217679380 |
| chr14 | 24973716 | 24975572 |
| chr14 | 35404654 | 35406131 |
| chr4 | 175303201 | 175305614 |
| chr10 | 50872136 | 50874211 |
| chr12 | 65903069 | 65904521 |
| chr1 | 64176797 | 64178203 |
| chr6 | 121930991 | 121933393 |
| chr14 | 26740545 | 26744487 |
| chr12 | 87925713 | 87928579 |
| chr1 | 98375863 | 98378219 |
| chr7 | 84140108 | 84142845 |
| chr2 | 188695396 | 188697327 |
| chr5 | 149035322 | 149036394 |
| chr2 | 140749855 | 140752131 |
| chr1 | 227120887 | 227122211 |
| chr14 | 42555484 | 42557249 |
| chr6 | 90411042 | 90413726 |
| chr4 | 35938322 | 35941363 |
| chr2 | 65324423 | 65325611 |
| chr2 | 57390223 | 57392265 |
| chr4 | 33427207 | 33429193 |
| chr4 | 49256966 | 49259069 |
| chr10 | 43110237 | 43111392 |
| chr2 | 4686862 | 4688606 |
| chr6 | 100703391 | 100706432 |
| chr12 | 85522655 | 85524137 |
| chr5_KI270897v1_a | 791460 | 793373 |
| chr7 | 79297562 | 79299613 |
| chr1 | 103606821 | 103608485 |

| | | |
|---|---|---|
| chr14 | 27372499 | 27374552 |
| chr2 | 162380145 | 162382959 |
| chr21 | 10330761 | 10333408 |
| chr7 | 81664742 | 81668467 |
| chr4 | 107220576 | 107223061 |
| chr10 | 65703566 | 65704984 |
| chr1 | 105834654 | 105836755 |
| chr15 | 57491603 | 57493458 |
| chr2 | 17765399 | 17767468 |
| chr7 | 13020491 | 13022672 |
| chr10 | 95237085 | 95238526 |
| chr10 | 102244158 | 102246113 |
| chr14 | 35122821 | 35125327 |
| chr16 | 87486362 | 87488011 |
| chr7 | 83615799 | 83619227 |
| chr5_KI270897v1_a | 139978 | 141934 |
| chr6 | 72237392 | 72238859 |
| chr12 | 124010025 | 124011689 |
| chr2 | 171920957 | 171922993 |
| chr16 | 87483833 | 87485521 |
| chr12 | 74979385 | 74980896 |
| chrUn_GL000195v1 | 95919 | 98272 |
| chr2 | 193094963 | 193098125 |
| chr2 | 41719987 | 41722412 |
| chr2 | 102868182 | 102869229 |
| chr7 | 76914650 | 76917743 |
| chrY | 11340104 | 11342025 |
| chr1 | 103756316 | 103759025 |
| chr2 | 188731941 | 188733200 |
| chr8 | 109814150 | 109815920 |
| chr10 | 67240503 | 67242450 |
| chr4 | 107277706 | 107281300 |
| chr3 | 153160260 | 153162822 |
| chr6 | 72713994 | 72715558 |
| chr2 | 165000385 | 165002295 |
| chr5 | 171128962 | 171131223 |
| chr10 | 133262473 | 133264177 |
| chr2 | 184756161 | 184759405 |
| chr2 | 164537251 | 164539059 |
| chr10 | 84216298 | 84218302 |
| chr12 | 102060714 | 102062629 |
| chrY | 6248937 | 6250793 |
| chr2 | 20050412 | 20053527 |
| chr16 | 33873560 | 33877364 |
| chr10 | 62087902 | 62091190 |
| chr4 | 114826094 | 114829053 |
| chr7 | 65635037 | 65636725 |
| chr18 | 9583645 | 9585936 |
| chr7 | 102660281 | 102662809 |

| | | |
|---|---|---|
| chr13 | 71876440 | 71880593 |
| chr14 | 27960699 | 27961859 |
| chr4 | 156376453 | 156378951 |
| chr7 | 84756747 | 84758610 |
| chr21 | 33321411 | 33322700 |
| chr2 | 104245648 | 104247923 |
| chr13 | 105473321 | 105476831 |
| chr14 | 92630140 | 92631853 |
| chr10 | 124774009 | 124775726 |
| chr1 | 161669480 | 161672097 |
| chr12 | 74738426 | 74739763 |
| chr2 | 196255712 | 196257071 |
| chr8 | 98581720 | 98583675 |
| chr1 | 219877520 | 219879642 |
| chr12 | 98221116 | 98222910 |
| chr2 | 187249709 | 187251143 |
| chr1 | 101049535 | 101052199 |
| chr2 | 63884622 | 63887109 |
| chr2 | 188962652 | 188964023 |
| chr16 | 34732297 | 34736161 |
| chr1 | 227903354 | 227906164 |
| chr2 | 195198538 | 195200468 |
| chr15 | 46638779 | 46640553 |
| chr9 | 31393347 | 31396581 |
| chr13 | 38473376 | 38475400 |
| chr10 | 97548682 | 97550756 |
| chr17 | 59785670 | 59788656 |
| chr5 | 163855168 | 163857125 |
| chr4 | 38455590 | 38458010 |
| chr1_KI270765v1_a | 33170 | 35491 |
| chr6 | 123001150 | 123004269 |
| chr4 | 115729746 | 115730903 |
| chr1 | 161111249 | 161112378 |
| chr6 | 89784479 | 89786236 |
| chr7 | 80645605 | 80647818 |
| chr7 | 77897800 | 77898991 |
| chr5 | 28497104 | 28499018 |
| chr15 | 96529475 | 96530792 |
| chr10 | 105162100 | 105163669 |
| chr3 | 174566658 | 174568605 |
| chr7 | 91717239 | 91719219 |
| chr3 | 174202412 | 174206968 |
| chr10 | 46848296 | 46851284 |
| chr6 | 74524793 | 74526253 |
| chr12 | 25193777 | 25197809 |
| chr7 | 57804647 | 57806423 |
| chr1 | 159777653 | 159779253 |
| chr10 | 84688133 | 84689790 |
| chr2 | 102804540 | 102806804 |

| | | |
|---|---|---|
| chr14 | 28667475 | 28669709 |
| chr10 | 109648599 | 109650647 |
| chr7 | 77543922 | 77546935 |
| chr13 | 45193978 | 45195613 |
| chr14 | 27298264 | 27299818 |
| chr2 | 73805767 | 73807848 |
| chr12 | 28363500 | 28366444 |
| chr4 | 13976556 | 13978486 |
| chr10 | 61754751 | 61757375 |
| chr12 | 62593571 | 62595109 |
| chr2 | 76931845 | 76933235 |
| chr1 | 189837542 | 189840634 |
| chr6 | 54769430 | 54771631 |
| chr9 | 67825699 | 67828944 |
| chr12 | 97539244 | 97541342 |
| chr13 | 67043023 | 67044400 |
| chr21 | 37613877 | 37616097 |
| chr9 | 22208152 | 22209945 |
| chr6 | 160640576 | 160642543 |
| chr8 | 115451056 | 115453287 |
| chr14 | 45555824 | 45557166 |
| chr10 | 61050316 | 61051937 |
| chr11 | 85317523 | 85319973 |
| chr7 | 12817142 | 12820814 |
| chr20 | 45976971 | 45979194 |
| chr2 | 83079708 | 83081493 |
| chr7 | 87640790 | 87643030 |
| chr4 | 163894078 | 163895622 |
| chrX | 80809478 | 80810571 |
| chr4 | 176740658 | 176742237 |
| chr11 | 101299005 | 101300908 |
| chr17_KI270909v1_ | 317418 | 318987 |
| chr14 | 34982137 | 34983755 |
| chr12 | 44150411 | 44152203 |
| chr20 | 6959777 | 6962699 |
| chr1 | 143274797 | 143277001 |
| chr8 | 86162581 | 86164610 |
| chr16 | 72948340 | 72950941 |
| chr1 | 86335004 | 86336890 |
| chr1 | 174454648 | 174457271 |
| chr13 | 89780734 | 89782371 |
| chr11 | 37348073 | 37349408 |
| chr4 | 176777894 | 176779166 |
| chr1 | 148451263 | 148453006 |
| chr12 | 65448359 | 65452299 |
| chr2 | 103881822 | 103884405 |
| chr12 | 77836351 | 77839086 |
| chr12 | 53937216 | 53940164 |
| chr14 | 46724818 | 46726261 |

| | | |
|---|---|---|
| chr14 | 49894982 | 49897105 |
| chr2 | 162269597 | 162271641 |
| chr14 | 28338406 | 28339749 |
| chr14 | 77022923 | 77025896 |
| chr3 | 95664497 | 95666589 |
| chr12 | 44348497 | 44349764 |
| chr6 | 39106348 | 39108494 |
| chr12 | 14366539 | 14368667 |
| chr1 | 113812243 | 113815296 |
| chr11 | 17387713 | 17389522 |
| chr15 | 56243788 | 56246815 |
| chr13 | 81380849 | 81382737 |
| chr7 | 89824863 | 89826533 |
| chr18 | 959559 | 961570 |
| chr9 | 65301503 | 65304467 |
| chr11 | 121426482 | 121428221 |
| chr11 | 3410115 | 3412278 |
| chr2 | 64538987 | 64541890 |
| chr14 | 57173184 | 57175759 |
| chr2 | 53525697 | 53526928 |
| chr17 | 47894687 | 47897364 |
| chr14 | 75340073 | 75341959 |
| chr10 | 109677458 | 109679675 |
| chr2 | 54654011 | 54655704 |
| chr4 | 34849490 | 34850672 |
| chr10 | 53788550 | 53790231 |
| chr10 | 108913247 | 108916618 |
| chr12 | 99981925 | 99983372 |
| chr2 | 58428631 | 58430336 |
| chr18 | 38415087 | 38416114 |
| chr10 | 54351566 | 54354268 |
| chr6 | 68934341 | 68937381 |
| chr11 | 16431911 | 16433340 |
| chr1 | 15524862 | 15528406 |
| chr6 | 578938 | 581145 |
| chr2 | 172101210 | 172103992 |
| chr12 | 68332065 | 68334104 |
| chr12 | 21161605 | 21163242 |
| chr17 | 43315178 | 43316908 |
| chr2 | 72860600 | 72861786 |
| chr10 | 44934714 | 44937220 |
| chr12 | 98288877 | 98292939 |
| chr14 | 26175714 | 26178759 |
| chr14 | 86332759 | 86335485 |
| chr10 | 46854891 | 46857745 |
| chr2 | 119757915 | 119760475 |
| chr21 | 16309310 | 16311674 |
| chr5 | 44807835 | 44811206 |
| chr5 | 125757974 | 125759838 |

| | | |
|---|---|---|
| chr12 | 84330107 | 84332356 |
| chr2 | 65263700 | 65266438 |
| chr10 | 35030506 | 35033670 |
| chr2 | 156908942 | 156912903 |
| chr1 | 98537221 | 98539029 |
| chr4 | 120361214 | 120363070 |
| chr1 | 188167009 | 188169072 |
| chr10 | 87424978 | 87426545 |
| chr6 | 40956689 | 40958367 |
| chr13 | 70889401 | 70891618 |
| chr2 | 140393703 | 140395926 |
| chr15 | 27046410 | 27047860 |
| chr2 | 67648657 | 67651391 |
| chr14 | 19304505 | 19306623 |
| chr1 | 102705096 | 102706714 |
| chr1 | 194301162 | 194303755 |
| chr2 | 144769637 | 144772707 |
| chr1 | 191764909 | 191768142 |
| chr1 | 148685636 | 148687040 |
| chr4 | 19024976 | 19027694 |
| chr14 | 20612400 | 20615475 |
| chr10 | 4728679 | 4730709 |
| chr2 | 195682885 | 195684402 |
| chr12 | 55991670 | 55994293 |
| chr3 | 105424860 | 105426455 |
| chr6 | 64221699 | 64225577 |
| chr1 | 200554588 | 200556827 |
| chr2 | 82841870 | 82844252 |
| chr4 | 43764178 | 43765599 |
| chr10 | 51727120 | 51728877 |
| chr1 | 102909057 | 102910970 |
| chr21 | 10465663 | 10468551 |
| chr6 | 56534606 | 56536113 |
| chr6 | 156902071 | 156903640 |
| chr2 | 37215899 | 37217410 |
| chr22 | 10777669 | 10781187 |
| chr14 | 41065890 | 41067858 |
| chr10 | 83412105 | 83414891 |
| chr16 | 75897090 | 75899825 |
| chr13 | 87266176 | 87268019 |
| chr14 | 27177742 | 27179326 |
| chr22 | 17033076 | 17034800 |
| chr2 | 193387001 | 193390643 |
| chr1 | 105512220 | 105514084 |
| chr14 | 28579384 | 28580411 |
| chr13 | 20437810 | 20440261 |
| chr11 | 23864993 | 23867106 |
| chr17 | 60268519 | 60269617 |
| chr7 | 84759105 | 84761262 |

| | | |
|---|---|---|
| chr2 | 57345042 | 57347048 |
| chr10 | 56298966 | 56303194 |
| chr11 | 22602210 | 22604645 |
| chr2 | 166082559 | 166084456 |
| chr10 | 37150388 | 37152856 |
| chr14 | 36240866 | 36242116 |
| chr18 | 2562479 | 2564360 |
| chr15 | 20890543 | 20893875 |
| chr11 | 90035012 | 90036916 |
| chr5 | 26700073 | 26701415 |
| chr4 | 156685129 | 156687313 |
| chr14 | 90482258 | 90484324 |
| chr12 | 17522019 | 17523719 |
| chr7 | 81481610 | 81483259 |
| chr1 | 86974949 | 86976745 |
| chr12 | 33004892 | 33006380 |
| chr1 | 146376056 | 146378213 |
| chr14 | 25696652 | 25699584 |
| chr1 | 109089923 | 109091753 |
| chr6 | 81744428 | 81747310 |
| chr7 | 32554976 | 32557429 |
| chr10 | 110136413 | 110138663 |
| chr2 | 176304587 | 176306309 |
| chr13 | 72289279 | 72292062 |
| chr12 | 97549476 | 97552037 |
| chr11 | 96104748 | 96106709 |
| chr6 | 101252728 | 101254613 |
| chr10 | 61970798 | 61972017 |
| chr5 | 20678 | 22929 |
| chr9 | 82148586 | 82151237 |
| chr20 | 61378873 | 61380864 |
| chr2 | 185536536 | 185538466 |
| chr12 | 90013275 | 90014606 |
| chr14 | 22375262 | 22377037 |
| chr1 | 95237862 | 95239752 |
| chr7 | 83040714 | 83042731 |
| chr14 | 26657140 | 26659368 |
| chr11 | 97894340 | 97896156 |
| chr4 | 29691148 | 29692459 |
| chr14 | 47266685 | 47269681 |
| chr5 | 117186508 | 117187848 |
| chr4 | 115666414 | 115669420 |
| chr2 | 51673186 | 51676110 |
| chr14 | 27072729 | 27074647 |
| chr14 | 27108641 | 27110864 |
| chr1 | 149270361 | 149271919 |
| chr7 | 54298190 | 54300454 |
| chr2 | 188751721 | 188753065 |
| chr3 | 46976031 | 46977309 |

| | | |
|---|---|---|
| chr5 | 108886285 | 108888187 |
| chr7 | 82411736 | 82413651 |
| chr3 | 105485523 | 105486631 |
| chr8 | 117749577 | 117752264 |
| chr6 | 57969956 | 57972229 |
| chr1 | 104144591 | 104147193 |
| chr4 | 115661522 | 115665752 |
| chr15 | 49156517 | 49157795 |
| chr7 | 146402004 | 146403606 |
| chr12 | 85954154 | 85956564 |
| chr14 | 52679917 | 52681320 |
| chr1 | 172285621 | 172288188 |
| chr10 | 112537953 | 112541072 |
| chr7 | 81354953 | 81357693 |
| chr21 | 8994836 | 9000203 |
| chr13 | 72277634 | 72279682 |
| chr9 | 6954689 | 6956187 |
| chr20 | 14346547 | 14348320 |
| chr7 | 83130430 | 83132381 |
| chr2 | 196558328 | 196561198 |
| chr1 | 189263552 | 189265601 |
| chr15 | 44536117 | 44537573 |
| chr10 | 81226028 | 81228054 |
| chr1 | 98101972 | 98103503 |
| chr10 | 104541797 | 104543249 |
| chr2 | 214584440 | 214586508 |
| chr7 | 79027963 | 79029696 |
| chr5 | 26448226 | 26450374 |
| chr1 | 195571088 | 195572552 |
| chr9 | 4712510 | 4714845 |
| chr14 | 74325979 | 74328833 |
| chr6 | 63697584 | 63699281 |
| chr13 | 70793775 | 70796357 |
| chr20 | 20072675 | 20074009 |
| chr14 | 30768903 | 30771133 |
| chr7 | 85920494 | 85922571 |
| chr9 | 66141242 | 66144515 |
| chr11 | 95067659 | 95070270 |
| chr7 | 64503348 | 64505488 |
| chr7 | 57845860 | 57848546 |
| chrY | 9535520 | 9537232 |
| chr2 | 59899364 | 59901045 |
| chr12 | 34366231 | 34367552 |
| chr14 | 39184922 | 39186549 |
| chr13 | 73985203 | 73986839 |
| chr15 | 20387667 | 20389223 |
| chr10 | 88330880 | 88333446 |
| chr1 | 103176033 | 103178842 |
| chr21 | 10457514 | 10459975 |

| | | |
|---|---|---|
| chr6 | 45481707 | 45485218 |
| chr2 | 47344596 | 47346468 |
| chr2 | 103015025 | 103017114 |
| chr10 | 54910036 | 54912495 |
| chr5 | 93164852 | 93166607 |
| chr2 | 201853890 | 201855249 |
| chr1 | 88863400 | 88864875 |
| chr6 | 94649077 | 94651729 |
| chr2 | 53776398 | 53778002 |
| chr20 | 16432442 | 16435341 |
| chr2 | 189596045 | 189598015 |
| chr1 | 193867562 | 193869730 |
| chr1 | 197091377 | 197093462 |
| chr6 | 90612331 | 90614121 |
| chr10 | 62236981 | 62238794 |
| chr5 | 28810194 | 28812653 |
| chr1 | 107141528 | 107144391 |
| chr18 | 23643404 | 23645473 |
| chr6 | 87701862 | 87703200 |
| chr14 | 67694361 | 67696456 |
| chr14 | 36604123 | 36607171 |
| chr1 | 243254757 | 243256575 |
| chr14 | 36909251 | 36911551 |
| chr2 | 144027578 | 144029316 |
| chr1 | 169495032 | 169496906 |
| chr20 | 4687039 | 4688472 |
| chr1 | 153932542 | 153934211 |
| chr2 | 187016111 | 187018620 |
| chr1 | 186005748 | 186008332 |
| chr8 | 31109422 | 31111074 |
| chr10 | 36145940 | 36147883 |
| chr2 | 198326822 | 198330211 |
| chr12 | 63035421 | 63037535 |
| chr10 | 62156630 | 62159033 |
| chr6 | 77416119 | 77417594 |
| chr2 | 44775039 | 44776882 |
| chr2 | 52823687 | 52825129 |
| chr2 | 225353983 | 225355805 |
| chr14 | 86406187 | 86408071 |
| chr5 | 128180474 | 128183614 |
| chr2 | 193699844 | 193702044 |
| chr9 | 63043666 | 63045474 |
| chr17 | 82236047 | 82237996 |
| chr13 | 86017068 | 86018288 |
| chr14 | 40978004 | 40979235 |
| chr7 | 80501383 | 80504318 |
| chr1 | 177306518 | 177307994 |
| chr6 | 129505962 | 129508623 |
| chr14 | 85392421 | 85394807 |

| | | |
|---|---|---|
| chr16 | 32123888 | 32125663 |
| chr7 | 87032498 | 87034870 |
| chr7 | 49140880 | 49142170 |
| chr9 | 62888199 | 62889874 |
| chr6 | 82767542 | 82769495 |
| chr11 | 16618055 | 16620198 |
| chr1 | 146961328 | 146963352 |
| chr17_JH159146v1_ | 133203 | 135498 |
| chr14 | 42442560 | 42444066 |
| chr5 | 108726327 | 108730123 |
| chr14 | 42632007 | 42633939 |
| chr1 | 97498730 | 97501418 |
| chr12 | 84060426 | 84062387 |
| chr9 | 61075080 | 61076460 |
| chr2 | 63691602 | 63693807 |
| chr2 | 34562514 | 34565032 |
| chr7 | 79193510 | 79195730 |
| chr1 | 102886198 | 102887943 |
| chr9 | 66109611 | 66111469 |
| chr11 | 103188171 | 103190202 |
| chr5 | 109773476 | 109774905 |
| chr14 | 33133804 | 33135431 |
| chr17 | 75574122 | 75575628 |
| chr5 | 140799058 | 140801306 |
| chr18_GL383571v1_ | 95018 | 96655 |
| chr1_GL383520v2_a | 280442 | 282177 |
| chr14 | 47074335 | 47077382 |
| chr10 | 38954369 | 38957972 |
| chr11 | 89993554 | 89995430 |
| chr6 | 19378339 | 19379550 |
| chr4 | 41206154 | 41208527 |
| chr4 | 115066531 | 115068018 |
| chr20 | 61027103 | 61029631 |
| chr11 | 121968334 | 121969530 |
| chr22 | 42855949 | 42857795 |
| chr14 | 52238360 | 52240573 |
| chr10 | 56885655 | 56887443 |
| chr16 | 32454014 | 32456908 |
| chr17 | 80543491 | 80545298 |
| chr1 | 241519412 | 241521155 |
| chr3 | 125739012 | 125740043 |
| chr1 | 79259522 | 79261003 |
| chr16 | 80824477 | 80825598 |
| chr7 | 78676725 | 78678505 |
| chr10 | 87072293 | 87073389 |
| chr20 | 56280894 | 56282455 |
| chr2 | 210932948 | 210935248 |
| chr10 | 52099137 | 52101184 |
| chr14 | 58960877 | 58963336 |

| | | |
|---|---|---|
| chr2 | 109856032 | 109858246 |
| chr5 | 132292902 | 132295748 |
| chr14 | 22925755 | 22927427 |
| chr2_KI270768v1_a | 12389 | 14797 |
| chr10 | 61901734 | 61906710 |
| chr9 | 63679089 | 63681397 |
| chr10 | 52596887 | 52600836 |
| chr12 | 17135342 | 17137361 |
| chr7 | 80424634 | 80426887 |
| chr2 | 194233615 | 194236866 |
| chr21 | 7612255 | 7614428 |
| chr6 | 90753962 | 90755159 |
| chr11 | 24590929 | 24593502 |
| chr7 | 53506689 | 53508493 |
| chr7 | 25600207 | 25602840 |
| chr7 | 99593773 | 99595788 |
| chr6 | 63724770 | 63727047 |
| chr2 | 66114359 | 66116537 |
| chr4 | 19399364 | 19401391 |
| chr2 | 82906968 | 82909782 |
| chr2 | 58495217 | 58496845 |
| chr15_KI270905v1_ | 2853860 | 2855252 |
| chr9 | 65785178 | 65788850 |
| chr2 | 58616465 | 58618001 |
| chr5 | 161789140 | 161791026 |
| chr7 | 3752933 | 3754959 |
| chr1 | 98481341 | 98484039 |
| chr5 | 52553147 | 52555562 |
| chr3 | 166879684 | 166881587 |
| chr6 | 133060849 | 133062688 |
| chr12 | 96396188 | 96399567 |
| chr14 | 100864334 | 100866402 |
| chr2 | 212358952 | 212360350 |
| chr2 | 65129685 | 65131530 |
| chr12 | 83888920 | 83891139 |
| chr4 | 36295924 | 36298965 |
| chr20 | 25390641 | 25391929 |
| chr2 | 85189348 | 85192031 |
| chr7 | 80671467 | 80675291 |
| chr1 | 106365593 | 106366753 |
| chr17 | 61090093 | 61091849 |
| chr14 | 68086733 | 68088856 |
| chr14 | 88241298 | 88244082 |
| chr12 | 89016804 | 89020583 |
| chr10 | 63162246 | 63164072 |
| chr10 | 67260983 | 67263472 |
| chr4 | 126601495 | 126603302 |
| chr6 | 75061239 | 75064382 |
| chr7 | 88692699 | 88694535 |

| | | |
|---|---|---|
| chr2 | 75786307 | 75788197 |
| chrY | 25349544 | 25350770 |
| chr10 | 119892172 | 119893902 |
| chr20 | 54199232 | 54200909 |
| chr1 | 108921402 | 108923227 |
| chr14 | 105486790 | 105488733 |
| chr2 | 91426390 | 91429976 |
| chr10 | 66675776 | 66676909 |
| chr10 | 113680208 | 113682890 |
| chr1 | 108754368 | 108757781 |
| chr6 | 63740194 | 63741991 |
| chr5 | 160418247 | 160420745 |
| chr18 | 3458891 | 3460392 |
| chr12 | 17449582 | 17451716 |
| chr2 | 164875768 | 164877767 |
| chr12 | 63780828 | 63782476 |
| chr10 | 36902959 | 36904287 |
| chr14 | 88828640 | 88830498 |
| chr6 | 8608273 | 8609977 |
| chr14 | 43857196 | 43858630 |
| chr6 | 90073372 | 90074956 |
| chr16 | 61834217 | 61837356 |
| chr15 | 20240069 | 20241322 |
| chr15 | 93708757 | 93710215 |
| chr12 | 14256627 | 14258803 |
| chr13 | 63072925 | 63075546 |
| chr9 | 72975393 | 72981701 |
| chr1 | 175635548 | 175638161 |
| chr11 | 133067441 | 133070069 |
| chr7 | 93903462 | 93905695 |
| chr11 | 5860685 | 5862176 |
| chr14 | 46875431 | 46879436 |
| chr6 | 124494571 | 124496599 |
| chr13 | 79547268 | 79549337 |
| chrY | 11278104 | 11280492 |
| chr3 | 95418780 | 95422506 |
| chr14 | 45907055 | 45908650 |
| chr11 | 106683732 | 106686115 |
| chr2 | 187139932 | 187141424 |
| chr20 | 55606884 | 55608198 |
| chr16 | 76521149 | 76522815 |
| chr12 | 80369168 | 80372888 |
| chr12 | 116061623 | 116064141 |
| chr6 | 9577205 | 9579501 |
| chr14 | 38972287 | 38973644 |
| chr12 | 78193717 | 78195297 |
| chr2 | 187422596 | 187423826 |
| chr14 | 83587776 | 83589256 |
| chr5 | 123755315 | 123757393 |

| | | |
|---|---|---|
| chr11 | 25969487 | 25971920 |
| chr5 | 20937916 | 20939659 |
| chr1 | 172342945 | 172346025 |
| chr2 | 208412233 | 208414172 |
| chr13 | 102426994 | 102428091 |
| chr2 | 57515856 | 57518646 |
| chr8 | 14944341 | 14945352 |
| chr13 | 101745390 | 101747410 |
| chr10 | 65566280 | 65569265 |
| chr10 | 120573993 | 120577146 |
| chr10 | 44059851 | 44061685 |
| chr14 | 27268074 | 27269472 |
| chr14 | 38278938 | 38280792 |
| chr2 | 182317388 | 182319809 |
| chr2 | 185261206 | 185262848 |
| chr10 | 62267365 | 62270922 |
| chr2 | 167728814 | 167730767 |
| chr6 | 9785807 | 9787080 |
| chr2 | 113582389 | 113585168 |
| chr20 | 51932603 | 51935401 |
| chr14 | 20453874 | 20455936 |
| chr19 | 19775089 | 19777547 |
| chr2 | 71512504 | 71513833 |
| chr1 | 110206155 | 110208253 |
| chr16 | 75566368 | 75568908 |
| chr2 | 198352856 | 198354966 |
| chr12 | 104740836 | 104743481 |
| chr14 | 84058347 | 84059911 |
| chr22 | 10715335 | 10718215 |
| chr14 | 86763986 | 86767666 |
| chr2 | 103288768 | 103292747 |
| chr1 | 191798222 | 191800769 |
| chr2 | 130461774 | 130463404 |
| chr12 | 91882864 | 91884607 |
| chr15 | 20774556 | 20776107 |
| chr1 | 87939385 | 87941736 |
| chr9 | 62374883 | 62376424 |
| chr2 | 189156294 | 189159884 |
| chr11 | 100355508 | 100357333 |
| chr11 | 26835042 | 26837604 |
| chr5 | 59104512 | 59106438 |
| chr7 | 82995616 | 82998041 |
| chr11 | 100238217 | 100241042 |
| chr3 | 86859473 | 86861464 |
| chr15 | 96657079 | 96659004 |
| chr10 | 115144280 | 115146736 |
| chr12 | 75333025 | 75335532 |
| chr7 | 91424635 | 91425976 |
| chr20 | 13819775 | 13820853 |

| | | |
|---|---|---|
| chr10 | 37394006 | 37396985 |
| chr21 | 16278362 | 16280891 |
| chr12 | 88945175 | 88948938 |
| chr8 | 109535169 | 109538200 |
| chr12 | 85510807 | 85513157 |
| chr11 | 23795625 | 23797462 |
| chr8 | 63168776 | 63170941 |
| chr10 | 63416319 | 63419238 |
| chr7 | 25861278 | 25863747 |
| chr13 | 108406141 | 108408524 |
| chr8_KI270810v1_a | 318277 | 320420 |
| chr8 | 110493796 | 110495353 |
| chr2 | 4429594 | 4430922 |
| chr7 | 78808992 | 78811783 |
| chr1 | 86429616 | 86431283 |
| chr1 | 195988277 | 195990676 |
| chr2 | 180130768 | 180132584 |
| chr20 | 32206396 | 32207803 |
| chr7 | 84249508 | 84252398 |
| chr2 | 53273916 | 53275464 |
| chr7 | 125607754 | 125609903 |
| chr12 | 86186433 | 86191034 |
| chr10 | 114461548 | 114463487 |
| chr8 | 65822778 | 65824211 |
| chr2 | 58750386 | 58752089 |
| chr14 | 73695436 | 73697575 |
| chr13 | 67060359 | 67063095 |
| chr12 | 40147656 | 40150090 |
| chr2 | 59531514 | 59532807 |
| chr16 | 77033165 | 77035079 |
| chr1 | 148688370 | 148691294 |
| chr14 | 90501675 | 90503567 |
| chr6 | 82616091 | 82617816 |
| chr17 | 52896792 | 52898227 |
| chr11 | 23687282 | 23690755 |
| chr12 | 9749804 | 9751892 |
| chr6 | 73725859 | 73728233 |
| chr5 | 35120901 | 35122609 |
| chr6 | 120745990 | 120748001 |
| chr4 | 49295455 | 49296738 |
| chr13 | 89401374 | 89403446 |
| chr6 | 56222036 | 56224893 |
| chr2 | 94948069 | 94949199 |
| chr14 | 87944767 | 87947149 |
| chr10 | 50813082 | 50814882 |
| chr14 | 81398227 | 81400762 |
| chr9 | 63880491 | 63883222 |
| chr2 | 193808570 | 193809862 |
| chr11 | 22821413 | 22823724 |

| | | |
|---|---|---|
| chr20 | 29048890 | 29050529 |
| chr12_KI270834v1_ | 28723 | 29794 |
| chr1 | 61089392 | 61091038 |
| chr1 | 180613720 | 180615620 |
| chr6 | 64307635 | 64309761 |
| chr11 | 18632911 | 18635423 |
| chr1 | 195882270 | 195884179 |
| chr11 | 94602335 | 94604375 |
| chr8 | 115960864 | 115965100 |
| chr2 | 233062240 | 233064620 |
| chr1 | 96796359 | 96797879 |
| chr18 | 25194352 | 25196723 |
| chr1 | 108222059 | 108224081 |
| chr21 | 7419153 | 7421780 |
| chr13 | 70962698 | 70965327 |
| chr4 | 36095073 | 36097369 |
| chr14 | 87532218 | 87533592 |
| chr6 | 71671573 | 71672991 |
| chr2 | 163199111 | 163200450 |
| chr12 | 79196172 | 79197996 |
| chr7 | 82097719 | 82099630 |
| chr1 | 27366177 | 27368889 |
| chr2 | 65015884 | 65018640 |
| chr20 | 14084756 | 14086595 |
| chr1 | 219064502 | 219066462 |
| chr10 | 63627334 | 63630710 |
| chr10 | 47971299 | 47973394 |
| chr7 | 83585714 | 83589308 |
| chr7 | 55149915 | 55151754 |
| chr8 | 121613521 | 121615231 |
| chr5 | 90408575 | 90410665 |
| chr6 | 136888072 | 136889874 |
| chr10 | 23775674 | 23778308 |
| chr6 | 149762149 | 149763682 |
| chr11 | 319359 | 322802 |
| chr13 | 87713364 | 87715665 |
| chr20 | 26190361 | 26192063 |
| chr2 | 55307958 | 55311277 |
| chr4 | 23319817 | 23322190 |
| chr10 | 86452784 | 86454808 |
| chr10 | 109957093 | 109959668 |
| chr7 | 149741017 | 149742519 |
| chr11 | 32966109 | 32967188 |
| chr8 | 78012521 | 78015001 |
| chr11 | 76015452 | 76017698 |
| chr5 | 80691552 | 80694005 |
| chr14 | 30004403 | 30006317 |
| chr6 | 19346718 | 19349113 |
| chr19 | 49812914 | 49814134 |

| | | |
|---|---|---|
| chr1 | 85986368 | 85988939 |
| chr9 | 66116885 | 66118761 |
| chr19 | 37370282 | 37372303 |
| chr12 | 44118348 | 44120989 |
| chr9 | 138261045 | 138262076 |
| chr15 | 63698197 | 63700092 |
| chr2 | 56020045 | 56021718 |
| chr11 | 81504401 | 81506288 |
| chr7 | 12731068 | 12732893 |
| chr2 | 69474574 | 69475817 |
| chr16 | 76522995 | 76525908 |
| chr2 | 226211062 | 226214383 |
| chr9 | 6931339 | 6933478 |
| chr2 | 80030001 | 80032348 |
| chr2 | 105435869 | 105437443 |
| chr6 | 91694677 | 91696283 |
| chr21 | 9919595 | 9921090 |
| chr4 | 116105892 | 116107564 |
| chr13 | 83827969 | 83830808 |
| chr14 | 77456344 | 77458951 |
| chr18 | 25095062 | 25096379 |
| chr10 | 74826747 | 74829868 |
| chr9 | 31169207 | 31171311 |
| chr2 | 171703169 | 171706809 |
| chr15_KI270905v1_ | 747696 | 750128 |
| chr4 | 37238254 | 37239493 |
| chr11 | 23745025 | 23747319 |
| chr17 | 22114010 | 22115062 |
| chr2 | 193274733 | 193276003 |
| chr10 | 61762735 | 61764500 |
| chr2 | 170932429 | 170934133 |
| chr2 | 177702532 | 177704953 |
| chr1 | 97827259 | 97829291 |
| chr4 | 92995269 | 92997084 |
| chr14 | 42873842 | 42876057 |
| chr1 | 199228675 | 199231729 |
| chr20 | 14122353 | 14124779 |
| chr5 | 10627486 | 10628526 |
| chr14 | 77761019 | 77762056 |
| chr19 | 42659122 | 42660318 |
| chr2 | 189066566 | 189069512 |
| chr7 | 100992706 | 100995843 |
| chr14 | 32625063 | 32631119 |
| chr12 | 56125946 | 56129225 |
| chr22 | 10732313 | 10734780 |
| chr14 | 40840863 | 40843118 |
| chr16 | 76944918 | 76946356 |
| chr6 | 61645210 | 61647009 |
| chr12 | 80856519 | 80858956 |

| | | |
|---|---|---|
| chr10 | 90666197 | 90668033 |
| chr20 | 3918171 | 3920531 |
| chr14 | 66036043 | 66037096 |
| chr16 | 76594825 | 76596043 |
| chr4 | 175823131 | 175825720 |
| chr1 | 87895170 | 87897341 |
| chr14 | 29596591 | 29600771 |
| chr7 | 81209598 | 81213074 |
| chr2 | 225793711 | 225797590 |
| chr13 | 93547140 | 93549303 |
| chr7 | 93254387 | 93257691 |
| chr11 | 21511890 | 21514085 |
| chr2 | 229528965 | 229530745 |
| chr2 | 182994488 | 182995577 |
| chr9 | 62345642 | 62347488 |
| chr7 | 63029741 | 63030776 |
| chr4 | 34347532 | 34348715 |
| chrY | 11843730 | 11844737 |
| chr12 | 84313322 | 84316749 |
| chr10 | 45951035 | 45952879 |
| chr14 | 41055342 | 41057207 |
| chr9 | 30865109 | 30867192 |
| chr10 | 110277868 | 110280340 |
| chr10 | 56490730 | 56492472 |
| chr12 | 75292322 | 75294251 |
| chr4 | 114972497 | 114974445 |
| chr16 | 65211485 | 65213626 |
| chr10 | 57673628 | 57675838 |
| chr6 | 72579891 | 72581353 |
| chr8 | 16187496 | 16188902 |
| chr2 | 213620382 | 213621997 |
| chr13 | 106076652 | 106079150 |
| chr12 | 120437491 | 120438918 |
| chr11 | 99255928 | 99257226 |
| chr10 | 55284028 | 55287402 |
| chr4 | 107139643 | 107141823 |
| chr1 | 67528004 | 67529336 |
| chr2 | 39770458 | 39773099 |
| chr20 | 53827428 | 53830180 |
| chr7 | 87269177 | 87270501 |
| chr1 | 190640194 | 190641833 |
| chr10 | 79546677 | 79548294 |
| chr14 | 38654625 | 38657056 |
| chr14 | 29113522 | 29116190 |
| chr1 | 79912317 | 79915353 |
| chr20 | 14651179 | 14653351 |
| chr2 | 95571965 | 95573675 |
| chr14 | 53305366 | 53307294 |
| chr12 | 72233158 | 72234792 |

| | | |
|---|---|---|
| chr11 | 22361495 | 22364364 |
| chr14 | 45693233 | 45695900 |
| chr10 | 110226545 | 110228231 |
| chr1 | 98208453 | 98211238 |
| chr6 | 73515309 | 73517735 |
| chr2 | 164116251 | 164117859 |
| chr11 | 27108015 | 27109740 |
| chr12 | 119790954 | 119792369 |
| chr8 | 111492611 | 111495915 |
| chr1 | 102964668 | 102967092 |
| chr1 | 105759927 | 105762096 |
| chr20 | 58650320 | 58652588 |
| chr2 | 50159001 | 50160604 |
| chr20 | 30287993 | 30290555 |
| chr6 | 23026810 | 23028332 |
| chr20 | 53586850 | 53588710 |
| chr4 | 28310825 | 28312172 |
| chr12 | 81976974 | 81978449 |
| chr16_KI270728v1_ | 1269706 | 1271020 |
| chr4 | 66507474 | 66509228 |
| chr7 | 84797883 | 84799492 |
| chr1 | 99395163 | 99397079 |
| chr1 | 243026907 | 243027949 |
| chr8 | 110916310 | 110919587 |
| chr7 | 86267334 | 86268534 |
| chr12 | 27649833 | 27650952 |
| chr14 | 40701691 | 40703093 |
| chr8 | 58009925 | 58011580 |
| chr4 | 107210189 | 107214062 |
| chr14 | 100776989 | 100779539 |
| chr13 | 63057497 | 63059752 |
| chr10 | 55085253 | 55087001 |
| chr1 | 104098982 | 104103142 |
| chr10 | 50450064 | 50452409 |
| chr1 | 66925429 | 66927227 |
| chr1 | 100919945 | 100923963 |
| chr10 | 107425952 | 107427629 |
| chr13 | 22230290 | 22232379 |
| chr14 | 40815328 | 40817067 |
| chr16 | 80246391 | 80247791 |
| chr22 | 15779946 | 15781813 |
| chr1 | 240755208 | 240756887 |
| chr4 | 35649212 | 35650521 |
| chr10 | 32958829 | 32960034 |
| chr1 | 51354080 | 51355787 |
| chr8 | 118189013 | 118191295 |
| chr7 | 61033404 | 61034913 |
| chr14 | 64522829 | 64524614 |
| chr12 | 6904171 | 6905234 |

| | | |
|---|---|---|
| chr22 | 12898864 | 12900401 |
| chr1 | 236546457 | 236549878 |
| chr3 | 8392154 | 8394125 |
| chr20 | 16442447 | 16444539 |
| chr7 | 85938016 | 85940065 |
| chr10 | 62232571 | 62234367 |
| chr22 | 16308859 | 16311282 |
| chr1 | 90007614 | 90009917 |
| chr1 | 102790306 | 102793247 |
| chr6 | 120790513 | 120791677 |
| chr2 | 64645949 | 64648419 |
| chr8 | 140763837 | 140766374 |
| chr16 | 76431777 | 76434082 |
| chr8 | 139378509 | 139380863 |
| chr10 | 17058080 | 17060291 |
| chr21 | 9916552 | 9919039 |
| chr12 | 65509676 | 65511520 |
| chr2 | 80144221 | 80147123 |
| chr12 | 23631084 | 23632838 |
| chr5 | 92559088 | 92562340 |
| chr1 | 75911495 | 75913092 |
| chr10 | 57360033 | 57362742 |
| chr15 | 92394222 | 92396630 |
| chr5 | 28220839 | 28222864 |
| chr2 | 225932326 | 225934875 |
| chr1 | 104434993 | 104437577 |
| chr15_KI270905v1_ | 928128 | 929554 |
| chr4 | 107166543 | 107168085 |
| chr1 | 175024684 | 175027834 |
| chr22 | 12283922 | 12285324 |
| chr2 | 184286022 | 184288198 |
| chr1 | 218199238 | 218202268 |
| chr6 | 78383293 | 78386220 |
| chr5 | 32405720 | 32409241 |
| chr10 | 84297699 | 84299055 |
| chr14 | 95135155 | 95137120 |
| chr9 | 65991953 | 65993512 |
| chr17 | 49190954 | 49193492 |
| chr9 | 66353108 | 66355475 |
| chr4 | 157366753 | 157370448 |
| chr12 | 18310141 | 18314470 |
| chr4 | 29504785 | 29506359 |
| chr19 | 37076463 | 37078169 |
| chr10 | 82379525 | 82380668 |
| chr14 | 46249534 | 46251418 |
| chr4 | 175851674 | 175854893 |
| chr8 | 108914539 | 108917726 |
| chr7 | 82052873 | 82055827 |
| chr9 | 11867355 | 11869652 |

| | | |
|---|---|---|
| chr8 | 43018308 | 43020198 |
| chr7 | 70003570 | 70005022 |
| chr6 | 120287743 | 120289352 |
| chr18 | 58315260 | 58317228 |
| chr1 | 125167666 | 125171007 |
| chr21 | 10497960 | 10500364 |
| chr14 | 105491618 | 105493049 |
| chr1 | 75787695 | 75791168 |
| chr4 | 8535075 | 8536887 |
| chr1 | 245058032 | 245060128 |
| chr10 | 16949499 | 16951676 |
| chr10 | 35872679 | 35875115 |
| chr1 | 9306295 | 9308528 |
| chr22 | 12277945 | 12280797 |
| chr7 | 103614434 | 103615741 |
| chr3 | 110235510 | 110237390 |
| chr10 | 75746214 | 75749337 |
| chr7 | 60950935 | 60952587 |
| chr7 | 10575035 | 10577086 |
| chr2 | 50887036 | 50891993 |
| chr20 | 44160193 | 44161785 |
| chr5 | 140281835 | 140283688 |
| chr2 | 181821647 | 181823123 |
| chr1 | 103030289 | 103033229 |
| chr7 | 76869667 | 76872387 |
| chr20 | 29321589 | 29323681 |
| chrUn_GL000219v1 | 90285 | 91342 |
| chr9 | 62864892 | 62867954 |
| chr7 | 89756191 | 89758925 |
| chr1 | 103781193 | 103782548 |
| chr2 | 204346108 | 204348251 |
| chr7 | 51276754 | 51278405 |
| chr1 | 62599307 | 62603422 |
| chr17 | 65779475 | 65781357 |
| chr14 | 99438529 | 99440005 |
| chr10 | 79860821 | 79862463 |
| chr2 | 212223243 | 212224778 |
| chr2 | 58139811 | 58141739 |
| chr2 | 114369998 | 114372427 |
| chr9 | 73131194 | 73135018 |
| chr9 | 66670044 | 66671807 |
| chr14 | 64296803 | 64299589 |
| chr10 | 114457903 | 114460281 |
| chr2 | 34816134 | 34817906 |
| chr1 | 108856855 | 108858614 |
| chr1 | 103524760 | 103527938 |
| chr10 | 8949873 | 8951650 |
| chr4 | 107153406 | 107155765 |
| chr16 | 34081331 | 34082975 |

| | | |
|---|---|---|
| chr13 | 65050999 | 65053970 |
| chr3 | 59590479 | 59592648 |
| chr5 | 81640223 | 81642418 |
| chr14 | 43882079 | 43884937 |
| chr16 | 34946079 | 34948458 |
| chr2 | 59055072 | 59056318 |
| chr2 | 82408880 | 82410938 |
| chr4 | 35536751 | 35538640 |
| chr1 | 168062214 | 168064379 |
| chr15_KI270850v1_ | 129404 | 131528 |
| chr14 | 47361687 | 47364508 |
| chr14 | 81213236 | 81214752 |
| chr2 | 66272697 | 66274575 |
| chr1 | 102738204 | 102741129 |
| chr22 | 12596243 | 12598012 |
| chrUn_GL000218v1 | 135087 | 138120 |
| chr2 | 78175170 | 78179081 |
| chr2 | 58573552 | 58575431 |
| chr7 | 79212571 | 79214733 |
| chr8 | 90197002 | 90199285 |
| chr4 | 90838870 | 90841755 |
| chr1 | 104699953 | 104702382 |
| chr10 | 123503918 | 123505467 |
| chr6 | 77816952 | 77818938 |
| chr6 | 93719631 | 93722783 |
| chr21 | 24495461 | 24498038 |
| chr2 | 215607057 | 215609981 |
| chr14 | 27691625 | 27695654 |
| chr21 | 10347192 | 10348404 |
| chr13 | 86295091 | 86296457 |
| chr16 | 30664978 | 30666084 |
| chr6 | 88210249 | 88212228 |
| chr3 | 54261738 | 54263351 |
| chr13 | 102595720 | 102597285 |
| chr12 | 82478481 | 82480524 |
| chr12 | 86379667 | 86381139 |
| chr1 | 66981334 | 66983328 |
| chr1 | 193104547 | 193107024 |
| chrUn_GL000195v1 | 109841 | 111413 |
| chr1 | 172281111 | 172283160 |
| chr1 | 189343273 | 189345729 |
| chr5 | 107671964 | 107674504 |
| chr8 | 114089421 | 114091842 |
| chr10 | 54560592 | 54564992 |
| chr12 | 78715809 | 78717363 |
| chr10 | 119969906 | 119972093 |
| chr10 | 115446211 | 115447575 |
| chr1 | 117499945 | 117502240 |
| chr4 | 9790304 | 9791747 |

| | | |
|---|---|---|
| chr6 | 6174957 | 6176989 |
| chr15 | 51981735 | 51983305 |
| chr6 | 1004459 | 1005728 |
| chr10 | 66375857 | 66380741 |
| chr1 | 95992286 | 95993993 |
| chr2 | 167179619 | 167181879 |
| chr2 | 207160264 | 207162626 |
| chr2 | 36416334 | 36418113 |
| chr10 | 59329572 | 59331562 |
| chr4 | 34722355 | 34724408 |
| chr1 | 107325568 | 107328311 |
| chr15 | 20261426 | 20263471 |
| chr2 | 59212469 | 59214703 |
| chr22 | 15746667 | 15748546 |
| chr10 | 21517988 | 21521284 |
| chr11 | 72111143 | 72114036 |
| chr7 | 107167132 | 107170241 |
| chr20 | 50193904 | 50195306 |
| chr15 | 35302422 | 35303577 |
| chr7 | 92521724 | 92523695 |
| chr14 | 48358013 | 48360440 |
| chr14 | 98047076 | 98049094 |
| chr1 | 66928758 | 66931314 |
| chrUn_GL000195v1 | 130273 | 132303 |
| chr9 | 138255770 | 138258260 |
| chr1 | 168808591 | 168811693 |
| chr2 | 132148419 | 132149568 |
| chr17 | 43169127 | 43172632 |
| chr16 | 30921183 | 30924143 |
| chr2 | 144942125 | 144945380 |
| chr16 | 73629469 | 73631524 |
| chr20 | 62611713 | 62613730 |
| chr2 | 21429265 | 21430780 |
| chr6 | 79556317 | 79558727 |
| chr7 | 54988972 | 54991914 |
| chr6 | 73951467 | 73953379 |
| chr14 | 62486117 | 62488657 |
| chr11 | 23644107 | 23645760 |
| chr11 | 119312116 | 119313271 |
| chr6 | 73507634 | 73510283 |
| chr8 | 133644892 | 133647123 |
| chr8 | 16884684 | 16886572 |
| chr3 | 110770194 | 110772112 |
| chr9 | 68234875 | 68236403 |
| chr13 | 71955875 | 71957577 |
| chr14 | 70978906 | 70980578 |
| chr9 | 41224978 | 41226022 |
| chr11 | 105454140 | 105456085 |
| chr12 | 72617956 | 72621209 |

| | | |
|---|---|---|
| chr7 | 76792179 | 76794231 |
| chr2 | 56894544 | 56897130 |
| chr14 | 95086674 | 95089615 |
| chr1 | 84163986 | 84165782 |
| chr7 | 149090329 | 149091496 |
| chr7 | 83151911 | 83154660 |
| chr14 | 49585290 | 49587684 |
| chr10 | 75233158 | 75234973 |
| chr2 | 82584030 | 82587283 |
| chr2 | 132013389 | 132016215 |
| chr2 | 183424560 | 183425908 |
| chr12 | 84846208 | 84847909 |
| chr10 | 97277512 | 97279331 |
| chr1 | 104566441 | 104568548 |
| chr14 | 25656102 | 25659076 |
| chr16 | 30813678 | 30814875 |
| chr9 | 63783902 | 63785869 |
| chr13 | 87099673 | 87101231 |
| chr14 | 85834924 | 85837080 |
| chr14 | 28616823 | 28621363 |
| chr14 | 32071693 | 32074386 |
| chr12 | 21227072 | 21228316 |
| chr5 | 91670044 | 91671794 |
| chr1 | 101945504 | 101947581 |
| chr5 | 39107383 | 39109060 |
| chr12 | 21023593 | 21026364 |
| chr11 | 23532828 | 23534868 |
| chr14 | 30183453 | 30185657 |
| chr14 | 58265908 | 58267010 |
| chr5 | 162095047 | 162097035 |
| chr22 | 41443501 | 41445688 |
| chr14 | 63201037 | 63202756 |
| chr2 | 86137667 | 86139586 |
| chr14 | 37023986 | 37025934 |
| chr4 | 156301455 | 156304248 |
| chr8 | 98825612 | 98826902 |
| chr4 | 137207799 | 137210135 |
| chr13 | 70120848 | 70123028 |
| chr1 | 143252356 | 143253997 |
| chr10 | 65339430 | 65344316 |
| chr12 | 86057478 | 86058667 |
| chr9 | 20610950 | 20612530 |
| chr13 | 79543269 | 79545554 |
| chr2 | 99968003 | 99970418 |
| chr7 | 84839666 | 84840828 |
| chr2 | 198940850 | 198942986 |
| chr2 | 66430744 | 66432862 |
| chr2 | 76585638 | 76587593 |
| chr14 | 16034633 | 16036535 |

| | | |
|---|---|---|
| chr1 | 149622714 | 149625484 |
| chr13 | 79914676 | 79916235 |
| chr2 | 49112278 | 49113979 |
| chr7 | 127619557 | 127621665 |
| chr10 | 65611073 | 65612608 |
| chr1 | 194182506 | 194184121 |
| chr16 | 36248227 | 36251306 |
| chr9 | 72558327 | 72560796 |
| chr6 | 79152283 | 79154228 |
| chr4 | 175881632 | 175883489 |
| chr14 | 44303553 | 44305867 |
| chr12 | 77151444 | 77154037 |
| chr7 | 80643505 | 80645292 |
| chr2 | 104444222 | 104445950 |
| chr4 | 34401320 | 34403211 |
| chr13 | 29590011 | 29591880 |
| chr7 | 76047463 | 76049070 |
| chr12 | 74649006 | 74651490 |
| chr10 | 35933736 | 35934997 |
| chr2 | 61047674 | 61050405 |
| chr8 | 99496284 | 99499285 |
| chr8 | 17697061 | 17699084 |
| chr14 | 53787912 | 53790319 |
| chr10 | 45647850 | 45650423 |
| chr16 | 36109879 | 36111000 |
| chr12 | 90709020 | 90710715 |
| chr14 | 91005105 | 91006204 |
| chr12 | 90543367 | 90546413 |
| chr12 | 88615946 | 88617186 |
| chr5 | 138183302 | 138185412 |
| chr14 | 54101266 | 54103252 |
| chr7 | 96899350 | 96901779 |
| chr7 | 80971843 | 80973955 |
| chr10 | 85721646 | 85723701 |
| chr1 | 119274874 | 119275979 |
| chr13 | 64092563 | 64093890 |
| chr2 | 191051632 | 191054037 |
| chr2 | 103297757 | 103300202 |
| chr7 | 77011054 | 77012639 |
| chr11 | 110653931 | 110656282 |
| chr8 | 6115052 | 6116348 |
| chr11 | 80650500 | 80652456 |
| chr13 | 50264231 | 50266118 |
| chr20 | 30921940 | 30924671 |
| chr2 | 204832243 | 204833655 |
| chr7 | 92840619 | 92843045 |
| chr2 | 49606939 | 49609647 |
| chr22 | 15743660 | 15745658 |
| chr2 | 195409221 | 195413552 |

| | | |
|---|---|---|
| chr1 | 99444948 | 99447238 |
| chr11 | 60853671 | 60855862 |
| chr20 | 20051987 | 20054124 |
| chr19 | 40593687 | 40595545 |
| chr1 | 89646781 | 89648047 |
| chr14 | 48154935 | 48156403 |
| chr12 | 88094696 | 88097476 |
| chr22 | 10634143 | 10636005 |
| chr7 | 79349644 | 79353739 |
| chr6 | 92802938 | 92805045 |
| chr18 | 63469200 | 63470496 |
| chr4 | 89904903 | 89907750 |
| chr5 | 19525556 | 19527461 |
| chr12 | 61303301 | 61304839 |
| chr1 | 92079401 | 92080734 |
| chr12 | 55453292 | 55455525 |
| chr1 | 167540059 | 167542174 |
| chr14 | 67764201 | 67768349 |
| chr6 | 26103438 | 26104919 |
| chr14 | 37900844 | 37904080 |
| chr14 | 43764291 | 43765623 |
| chr4 | 43676258 | 43678991 |
| chr14 | 66511030 | 66514522 |
| chr9 | 62350144 | 62351673 |
| chr10 | 127986321 | 127988275 |
| chr15 | 101640974 | 101643213 |
| chr1 | 199722559 | 199725474 |
| chr1 | 194164589 | 194166215 |
| chr12 | 84736994 | 84740064 |
| chr10 | 54239232 | 54240474 |
| chr12 | 96367613 | 96369819 |
| chr11 | 80488360 | 80490515 |
| chr12 | 65169684 | 65171512 |
| chr20 | 30394189 | 30396532 |
| chr1 | 97552485 | 97554790 |
| chr8 | 107303232 | 107305538 |
| chr12 | 70355339 | 70357142 |
| chr17 | 21757093 | 21758110 |
| chr1 | 148527910 | 148530068 |
| chr17 | 51207213 | 51209388 |
| chr2 | 165381512 | 165383613 |
| chr15 | 69391258 | 69393248 |
| chr2 | 160323516 | 160326771 |
| chr2 | 40755138 | 40756598 |
| chr2 | 60577486 | 60579292 |
| chr11 | 26562502 | 26565405 |
| chr8 | 63620179 | 63622884 |
| chr2 | 39017460 | 39018948 |
| chr2 | 99887074 | 99888619 |

| | | |
|---|---|---|
| chr15 | 50738522 | 50740534 |
| chr1 | 54600246 | 54602043 |
| chr14 | 45640669 | 45642699 |
| chrX | 29536336 | 29537388 |
| chr5 | 120790969 | 120792569 |
| chr13 | 81969354 | 81971499 |
| chr10 | 116436648 | 116438810 |
| chr1 | 101973420 | 101975477 |
| chr6 | 101456375 | 101458690 |
| chr10 | 93372257 | 93374013 |
| chr17 | 59753127 | 59755243 |
| chr10 | 48038455 | 48039819 |
| chr2 | 140278787 | 140280223 |
| chr6 | 26044752 | 26046160 |
| chr20 | 30820323 | 30822757 |
| chr1 | 213819742 | 213822032 |
| chr10 | 65327521 | 65331170 |
| chr4 | 115248123 | 115249335 |
| chr12 | 80610121 | 80612075 |
| chr15 | 44517973 | 44519549 |
| chr6 | 122769060 | 122771133 |
| chr8 | 109380726 | 109384689 |
| chr10 | 65134953 | 65138281 |
| chr10 | 76738529 | 76740629 |
| chr16 | 22195038 | 22196594 |
| chr10 | 54744697 | 54746363 |
| chr11 | 31653023 | 31656482 |
| chr5 | 27825556 | 27830012 |
| chr5_KI270897v1_a | 789073 | 791255 |
| chr4 | 34918495 | 34919611 |
| chr8 | 82049486 | 82051615 |
| chr12 | 90481487 | 90483035 |
| chr1 | 65371221 | 65373408 |
| chr14 | 38900735 | 38901822 |
| chr2 | 35543492 | 35545262 |
| chr11 | 98784264 | 98786438 |
| chr2 | 57424349 | 57426648 |
| chr3 | 161608876 | 161611250 |
| chr16 | 79932818 | 79935112 |
| chr6 | 77602770 | 77604943 |
| chr19 | 39411300 | 39412422 |
| chr15_KI270852v1_ | 196125 | 197681 |
| chr1 | 8046829 | 8047972 |
| chr5 | 121792945 | 121794111 |
| chr12 | 39683844 | 39685074 |
| chr11 | 88754984 | 88756685 |
| chr10 | 50846039 | 50848076 |
| chr4 | 70051523 | 70054531 |
| chr10 | 65344703 | 65347124 |

| | | |
|---|---|---|
| chr1 | 171840393 | 171841463 |
| chr6 | 120494760 | 120496214 |
| chr15 | 41695430 | 41696731 |
| chr12 | 84292664 | 84296073 |
| chr9 | 83247825 | 83249833 |
| chr12 | 18738257 | 18740987 |
| chr1 | 190538115 | 190540248 |
| chr8 | 104018760 | 104021883 |
| chr14 | 51354561 | 51357085 |
| chr16 | 35505688 | 35507348 |
| chr1 | 72010322 | 72011904 |
| chr16 | 76273208 | 76277594 |
| chr14 | 89040115 | 89042645 |
| chr16 | 90180556 | 90181974 |
| chr5 | 149854910 | 149856855 |
| chr1 | 98860076 | 98862368 |
| chr2 | 189439131 | 189441203 |
| chr5 | 120301295 | 120302685 |
| chr1 | 99528681 | 99532122 |
| chr1 | 148714713 | 148716641 |
| chr10 | 101814353 | 101820067 |
| chr10 | 91035024 | 91037614 |
| chr11 | 28184259 | 28186943 |
| chr12 | 50504223 | 50506736 |
| chr10 | 105744852 | 105747214 |
| chr19 | 14749137 | 14750208 |
| chr3 | 96421248 | 96422744 |
| chr1 | 193088815 | 193091308 |
| chr6 | 122725173 | 122727910 |
| chr5 | 74395464 | 74397356 |
| chr12 | 80595383 | 80597060 |
| chr2 | 131857684 | 131859652 |
| chr3 | 24724165 | 24726083 |
| chr1 | 102751082 | 102753036 |
| chr4 | 117049511 | 117051701 |
| chr8 | 115658644 | 115662070 |
| chr4 | 29654602 | 29656007 |
| chr12 | 45498702 | 45500177 |
| chr2 | 84576812 | 84578079 |
| chr2 | 92078221 | 92080907 |
| chr14 | 44960705 | 44965081 |
| chr11 | 46786468 | 46787474 |
| chr1 | 143264136 | 143266823 |
| chr8 | 105750384 | 105754446 |
| chr3 | 104790987 | 104792802 |
| chr6 | 47741965 | 47745475 |
| chr2 | 50081610 | 50083640 |
| chr20 | 41315775 | 41318369 |
| chr10 | 52620614 | 52623688 |

| | | |
|---|---|---|
| chr12 | 45960226 | 45962089 |
| chr15 | 94814991 | 94816835 |
| chr12 | 66035001 | 66036993 |
| chr14 | 27557187 | 27559908 |
| chr2 | 24705873 | 24707657 |
| chr16 | 34853417 | 34855012 |
| chr6 | 120754166 | 120755779 |
| chr12 | 18343595 | 18346144 |
| chr2 | 185475130 | 185477292 |
| chr7 | 102667278 | 102668816 |
| chr1 | 220191512 | 220194183 |
| chr2 | 204905237 | 204908712 |
| chr9 | 23076496 | 23077813 |
| chr1 | 101986641 | 101988062 |
| chr1 | 76493166 | 76495217 |
| chr14 | 63983324 | 63985229 |
| chr10 | 108004116 | 108006253 |
| chr1 | 68127981 | 68129922 |
| chr4 | 35091382 | 35095539 |
| chr2 | 97396757 | 97398215 |
| chr11 | 134826970 | 134828558 |
| chr7 | 79075517 | 79076978 |
| chr6 | 104943003 | 104945672 |
| chr2 | 165698734 | 165700104 |
| chr14 | 33304407 | 33307127 |
| chr10 | 46550271 | 46552474 |
| chr14 | 59006043 | 59008339 |
| chr10 | 112156418 | 112158381 |
| chr18 | 53691540 | 53693477 |
| chr22_KI270733v1_ | 142791 | 144520 |
| chr6 | 26021082 | 26022806 |
| chr19 | 39308116 | 39309353 |
| chr10 | 37222180 | 37224815 |
| chr6 | 133923002 | 133924431 |
| chr16 | 53207408 | 53210552 |
| chr11 | 87287677 | 87289559 |
| chr7 | 79279751 | 79282782 |
| chr2 | 161803599 | 161805309 |
| chr1 | 82988057 | 82990913 |
| chr11 | 108466712 | 108468135 |
| chr7 | 5230469 | 5232227 |
| chr20 | 11873419 | 11874593 |
| chr7 | 102650958 | 102652610 |
| chr7 | 82152296 | 82154600 |
| chr12 | 84912370 | 84914621 |
| chr2 | 44840858 | 44842931 |
| chr15 | 27967400 | 27968484 |
| chr14 | 46831082 | 46832945 |
| chr3 | 83352025 | 83353800 |

| | | |
|---|---|---|
| chr6 | 82841949 | 82843708 |
| chr12 | 102762504 | 102765259 |
| chr12 | 75570789 | 75572457 |
| chr6 | 45440681 | 45442458 |
| chr11 | 26557257 | 26559208 |
| chr16 | 61896680 | 61898906 |
| chr17_KI270908v1_ | 220904 | 222589 |
| chr2 | 206566002 | 206568032 |
| chr2 | 222237453 | 222239527 |
| chr11 | 39108346 | 39110195 |
| chr14 | 38332800 | 38335216 |
| chr1 | 86437498 | 86439286 |
| chr20 | 33444038 | 33445480 |
| chr4 | 142525693 | 142526842 |
| chr2 | 178661489 | 178663354 |
| chr14 | 65413988 | 65415445 |
| chr4 | 22915978 | 22917407 |
| chr8 | 76139877 | 76141262 |
| chr11 | 106589757 | 106591880 |
| chr7 | 85450626 | 85452341 |
| chr10 | 104356817 | 104358703 |
| chr15 | 51573615 | 51576075 |
| chr7 | 80601452 | 80602904 |
| chr12 | 17383848 | 17385445 |
| chr8 | 99708801 | 99710041 |
| chr12 | 41298969 | 41303774 |
| chr14 | 26820069 | 26822341 |
| chr3 | 23926060 | 23927590 |
| chr2 | 155372623 | 155375041 |
| chr12 | 85292051 | 85294066 |
| chr12 | 89352011 | 89354309 |
| chr10 | 92985053 | 92987414 |
| chr1 | 102113041 | 102115496 |
| chr21 | 16992845 | 16994245 |
| chr10 | 55434516 | 55437731 |
| chr14 | 74022284 | 74023788 |
| chr10 | 24620232 | 24621382 |
| chr1 | 173329120 | 173330371 |
| chr14 | 54451906 | 54454840 |
| chr16 | 35479912 | 35482259 |
| chr5 | 121960843 | 121962619 |
| chr6 | 141347641 | 141349410 |
| chr12 | 87084647 | 87086804 |
| chr11 | 21647995 | 21650268 |
| chr15 | 34009933 | 34012024 |
| chr14 | 45014966 | 45017031 |
| chr5 | 176057565 | 176058570 |
| chr6 | 51624935 | 51627731 |
| chr2 | 183155480 | 183158199 |

| | | |
|---|---|---|
| chr12 | 71245754 | 71248389 |
| chr11 | 27472265 | 27474205 |
| chr4 | 161159811 | 161162078 |
| chr13 | 105415222 | 105417147 |
| chr15 | 32346230 | 32347932 |
| chr10 | 89772331 | 89774625 |
| chr12 | 79165210 | 79166477 |
| chr6 | 160611206 | 160614782 |
| chr14 | 43169124 | 43171222 |
| chr20 | 58716797 | 58718147 |
| chr4 | 78880279 | 78882800 |
| chr1 | 104149501 | 104151257 |
| chr1 | 27044209 | 27046507 |
| chr20 | 53385642 | 53387199 |
| chr10 | 121056122 | 121058740 |
| chr7 | 82413701 | 82415601 |
| chr10 | 55741450 | 55744445 |
| chr12 | 81450510 | 81453787 |
| chr1 | 166388857 | 166391706 |
| chr2 | 45582881 | 45586140 |
| chr21 | 9009304 | 9011089 |
| chr2 | 185470471 | 185473002 |
| chr1 | 193158614 | 193160598 |
| chr14 | 25833444 | 25838914 |
| chr6 | 101371399 | 101373838 |
| chr19 | 39325732 | 39327803 |
| chr1 | 215292356 | 215293571 |
| chr13 | 66445813 | 66447491 |
| chr2 | 95623290 | 95624994 |
| chr10 | 54833485 | 54835557 |
| chr10 | 105210215 | 105212855 |
| chr6 | 94844544 | 94846973 |
| chr1 | 186962768 | 186964714 |
| chr6 | 142941264 | 142943654 |
| chr20 | 19174225 | 19175226 |
| chr6 | 71998380 | 72000391 |
| chr11 | 112980477 | 112981902 |
| chr4 | 179449943 | 179451301 |
| chr14 | 56997589 | 56999886 |
| chr8 | 111443595 | 111445678 |
| chr14 | 65850322 | 65853259 |
| chr5 | 88827630 | 88829586 |
| chr10 | 92874939 | 92877521 |
| chr10 | 50499211 | 50501482 |
| chr11 | 99548830 | 99550398 |
| chr6 | 131060770 | 131063832 |
| chr6 | 79042248 | 79044136 |
| chr2 | 61259790 | 61261207 |
| chr10 | 90038073 | 90039466 |

| | | |
|---|---|---|
| chr3 | 117678136 | 117680076 |
| chr11 | 101789530 | 101791888 |
| chr14 | 47427764 | 47429008 |
| chr14 | 33466464 | 33468697 |
| chr2 | 97584109 | 97585642 |
| chr2 | 82461674 | 82463089 |
| chr14 | 89917345 | 89918626 |
| chr20 | 35732251 | 35733443 |
| chr5 | 127031073 | 127034197 |
| chr4 | 73068439 | 73070802 |
| chr14 | 49827851 | 49829978 |
| chr12 | 44735954 | 44738006 |
| chr2 | 189087621 | 189089589 |
| chr7 | 83904614 | 83906494 |
| chr8 | 104906745 | 104909306 |
| chr21 | 10427745 | 10431249 |
| chr15 | 101379139 | 101381349 |
| chr14 | 29038809 | 29040773 |
| chr3 | 110579121 | 110581427 |
| chr9 | 138311383 | 138312919 |
| chr11 | 104055248 | 104056654 |
| chr14 | 46402936 | 46405864 |
| chr6 | 121922988 | 121926679 |
| chr14 | 103972127 | 103973602 |
| chr15 | 23431373 | 23433062 |
| chr8 | 111558241 | 111562015 |
| chr2 | 75689795 | 75691278 |
| chr1 | 211295970 | 211297392 |
| chr4 | 19829362 | 19831654 |
| chr2 | 192995240 | 192996883 |
| chr1 | 62810481 | 62811531 |
| chr7 | 85001260 | 85003767 |
| chr6 | 101622347 | 101624030 |
| chr6 | 77630677 | 77632751 |
| chr13 | 64604914 | 64606536 |
| chr1 | 149234931 | 149237223 |
| chr11 | 99699323 | 99701287 |
| chr1 | 92623784 | 92626017 |
| chr10 | 46310106 | 46313397 |
| chr17_KI270908v1_ | 1160719 | 1163746 |
| chr21 | 10569283 | 10571038 |
| chr15 | 36265529 | 36267400 |
| chr6 | 98162618 | 98164817 |
| chr7 | 84852733 | 84854357 |
| chr16 | 59300966 | 59302595 |
| chr3 | 179895562 | 179898756 |
| chrY | 25379825 | 25381693 |
| chr5 | 109464192 | 109465870 |
| chr14 | 44285261 | 44286970 |

| | | |
|---|---|---|
| chr19 | 50139603 | 50140817 |
| chr17 | 11577575 | 11578594 |
| chr12 | 82932540 | 82934273 |
| chr6 | 94687988 | 94690372 |
| chr1 | 107746716 | 107749853 |
| chr5_KI270897v1_a | 79262 | 81718 |
| chr10 | 124453607 | 124455258 |
| chr2 | 189284875 | 189288151 |
| chr1 | 65845234 | 65848444 |
| chr2 | 55049103 | 55051969 |
| chr21 | 9650351 | 9653090 |
| chr1 | 147598885 | 147600902 |
| chr14 | 90399109 | 90400518 |
| chr1 | 151070418 | 151072301 |
| chr16 | 1969481 | 1970483 |
| chr13 | 73995591 | 73997372 |
| chr1 | 103315297 | 103317809 |
| chr16 | 33740557 | 33742981 |
| chr10 | 4719279 | 4722241 |
| chr7 | 93818394 | 93820578 |
| chr8 | 101631099 | 101632983 |
| chr10 | 121879475 | 121881200 |
| chr10 | 115892134 | 115893262 |
| chr10 | 66732827 | 66734556 |
| chr10 | 82982884 | 82986094 |
| chr2 | 224692444 | 224694110 |
| chr12 | 83676248 | 83678010 |
| chr4 | 156670760 | 156675794 |
| chr7 | 94434990 | 94436354 |
| chr10 | 4241222 | 4244618 |
| chr19 | 9818351 | 9820753 |
| chr6 | 56084405 | 56086578 |
| chr11 | 6353072 | 6354455 |
| chr8 | 126216748 | 126218006 |
| chr4 | 168092198 | 168093662 |
| chr14 | 51241241 | 51242869 |
| chr14 | 67322004 | 67324681 |
| chr10 | 116286472 | 116288730 |
| chr5 | 111192974 | 111194101 |
| chr2 | 113594093 | 113595197 |
| chr10 | 55449371 | 55451157 |
| chr6 | 73128500 | 73130784 |
| chr10 | 55959012 | 55962710 |
| chr2 | 86333110 | 86334801 |
| chr2 | 182017405 | 182020302 |
| chr2 | 104273211 | 104274937 |
| chr7_KI270809v1_a | 20581 | 21657 |
| chr13 | 28003149 | 28004525 |
| chr14 | 102458521 | 102460320 |

| | | |
|---|---|---|
| chr9 | 6937031 | 6939512 |
| chr15 | 20303765 | 20305865 |
| chr1 | 101755869 | 101758098 |
| chr1 | 85530582 | 85531610 |
| chr5 | 84308191 | 84309332 |
| chr12 | 111409000 | 111411006 |
| chr15 | 53833351 | 53835987 |
| chr1 | 167989524 | 167991764 |
| chr13 | 71664139 | 71666913 |
| chr12 | 51047694 | 51049380 |
| chr2 | 58761589 | 58765774 |
| chr16_KI270728v1_ | 17250 | 19847 |
| chr2 | 197828196 | 197830357 |
| chr7 | 88909743 | 88912095 |
| chr14 | 19970910 | 19973981 |
| chr10 | 55376405 | 55378118 |
| chr12 | 71523587 | 71525813 |
| chr13 | 66655222 | 66658047 |
| chr6 | 123664809 | 123666507 |
| chr7 | 81079381 | 81081349 |
| chr13 | 71710610 | 71715030 |
| chr14 | 24677704 | 24679615 |
| chr9 | 66273623 | 66276761 |
| chr13 | 63730856 | 63732774 |
| chr4 | 175418097 | 175420684 |
| chr7 | 25849539 | 25851021 |
| chr12 | 43400128 | 43401900 |
| chr2 | 187112206 | 187114240 |
| chr6 | 10597410 | 10599069 |
| chr14 | 83679181 | 83680904 |
| chr14 | 26380417 | 26381848 |
| chr1 | 172254385 | 172256149 |
| chr6 | 120259902 | 120261894 |
| chr11 | 86306636 | 86308993 |
| chr6 | 94487092 | 94488421 |
| chr1 | 86893243 | 86895443 |
| chr12 | 18217068 | 18218221 |
| chr10 | 50848552 | 50851531 |
| chr14 | 36323396 | 36325089 |
| chr6 | 9013702 | 9015079 |
| chr2 | 159877528 | 159879817 |
| chr6 | 8063781 | 8065032 |
| chr4 | 30192212 | 30195885 |
| chr11 | 23105295 | 23107117 |
| chr12 | 74890005 | 74892285 |
| chr12 | 76559297 | 76560994 |
| chr14 | 19477279 | 19478818 |
| chr10 | 113560614 | 113562545 |
| chr14 | 50144428 | 50146052 |

| | | |
|---|---|---|
| chr22 | 11043626 | 11044797 |
| chr9 | 35115406 | 35117308 |
| chr2 | 28011490 | 28013307 |
| chr9 | 62836993 | 62838749 |
| chr15 | 76347434 | 76348952 |
| chr10 | 63232264 | 63234499 |
| chr14 | 36492978 | 36495774 |
| chr14 | 28817311 | 28819314 |
| chr7 | 89241224 | 89243576 |
| chr10 | 115298913 | 115301066 |
| chr8 | 24297089 | 24301330 |
| chr14 | 80279081 | 80280984 |
| chr6 | 135193911 | 135196101 |
| chr9 | 128066729 | 128069159 |
| chr14 | 27087743 | 27089673 |
| chr14 | 40867307 | 40869308 |
| chr10 | 64910721 | 64912694 |
| chr4 | 66848594 | 66851014 |
| chr3 | 89183388 | 89184826 |
| chr2 | 165564196 | 165566652 |
| chr14 | 47288582 | 47290304 |
| chr11 | 113573519 | 113574940 |
| chr2 | 40974774 | 40976350 |
| chr22 | 12039480 | 12041032 |
| chr1 | 87128159 | 87130040 |
| chr20 | 324284 | 325781 |
| chr21 | 7469316 | 7471832 |
| chr19 | 182304 | 183814 |
| chr10 | 117309777 | 117311722 |
| chr13 | 41700528 | 41702631 |
| chr8 | 15352170 | 15354071 |
| chr13 | 66659806 | 66662023 |
| chr9 | 73022574 | 73024077 |
| chr6 | 63579421 | 63583535 |
| chr10 | 86263587 | 86267212 |
| chr2 | 190087780 | 190089680 |
| chr10 | 110459516 | 110462676 |
| chr4 | 176789608 | 176790979 |
| chr5_KI270897v1_a | 779620 | 781871 |
| chr10 | 115566901 | 115568920 |
| chr1 | 215540064 | 215541971 |
| chr6 | 72390514 | 72392195 |
| chr1 | 103144182 | 103146248 |
| chr2 | 60552482 | 60554291 |
| chr7 | 96931450 | 96933923 |
| chr1 | 88786675 | 88788437 |
| chrUn_GL000195v1 | 57294 | 59273 |
| chr12 | 126840027 | 126842138 |
| chr4 | 29437042 | 29438882 |

| | | |
|---|---|---|
| chr14 | 21658031 | 21659824 |
| chr1 | 61724359 | 61725920 |
| chr13 | 87667800 | 87671753 |
| chr1 | 176132071 | 176133570 |
| chr2 | 52968439 | 52970918 |
| chr2 | 171045537 | 171046702 |
| chr3 | 174183683 | 174185706 |
| chr7 | 95288335 | 95291569 |
| chr15 | 50144871 | 50146129 |
| chr2 | 188505190 | 188506854 |
| chr1 | 196719148 | 196720689 |
| chr1 | 69640377 | 69642067 |
| chr6 | 64077191 | 64079297 |
| chr1 | 208246253 | 208247688 |
| chr5 | 27280094 | 27281106 |
| chr2 | 177418221 | 177420014 |
| chr6 | 97208264 | 97210991 |
| chr15_KI270905v1_ | 826880 | 828419 |
| chr10 | 63192500 | 63195321 |
| chr6 | 47857485 | 47859143 |
| chr16 | 79449471 | 79451793 |
| chr3 | 84422784 | 84425688 |
| chr10 | 4368881 | 4371876 |
| chr6 | 121474735 | 121475973 |
| chr5 | 153155725 | 153157370 |
| chr16 | 72902171 | 72905218 |
| chr1 | 111437013 | 111439086 |
| chr1 | 215779966 | 215781694 |
| chr14 | 40502037 | 40503189 |
| chrUn_GL000195v1 | 62732 | 64516 |
| chr14 | 41240275 | 41242250 |
| chr2 | 39182903 | 39185062 |
| chr13 | 79918072 | 79920148 |
| chr14 | 20049100 | 20050717 |
| chr5 | 125527935 | 125529612 |
| chr10 | 70231945 | 70234293 |
| chr2 | 163344147 | 163348093 |
| chrY | 11707376 | 11710259 |
| chr4 | 32778673 | 32780839 |
| chr11 | 89673838 | 89675798 |
| chr21 | 5224989 | 5226606 |
| chr2 | 55591365 | 55593768 |
| chr2 | 81684395 | 81687874 |
| chr6 | 141482455 | 141484703 |
| chr7 | 84682611 | 84685381 |
| chr14 | 52329982 | 52331575 |
| chr1 | 107974580 | 107976088 |
| chr1 | 185342496 | 185344580 |
| chr4 | 126399956 | 126401708 |

| | | |
|---|---|---|
| chr15 | 95468289 | 95470150 |
| chr10 | 79894884 | 79896832 |
| chr1 | 238641875 | 238644819 |
| chr7 | 93874765 | 93876410 |
| chr2 | 163653587 | 163655682 |
| chr14 | 89278182 | 89280489 |
| chr12 | 86258098 | 86259494 |
| chr7 | 152263965 | 152266560 |
| chr2 | 213310221 | 213312284 |
| chr7 | 81466815 | 81468360 |
| chr2 | 213284539 | 213285829 |
| chr12 | 27687716 | 27688993 |
| chr9 | 5755945 | 5758063 |
| chr14 | 40759726 | 40761528 |
| chr14 | 36051961 | 36054451 |
| chr12 | 77479550 | 77481530 |
| chr7 | 89582346 | 89583910 |
| chr6 | 71661851 | 71663100 |
| chr12 | 84889410 | 84892265 |
| chr8 | 85991305 | 85993983 |
| chr13 | 69860072 | 69861562 |
| chr7 | 83935790 | 83937550 |
| chr1 | 230101898 | 230103453 |
| chr3 | 87688687 | 87690382 |
| chr2 | 63465732 | 63467533 |
| chr8 | 93670696 | 93672171 |
| chr12 | 94548527 | 94550767 |
| chr10 | 59003099 | 59004672 |
| chr14 | 24435746 | 24437257 |
| chr6 | 120557449 | 120558768 |
| chr16 | 70521851 | 70524299 |
| chr11 | 22271550 | 22273503 |
| chr2 | 209149303 | 209152580 |
| chr4 | 11966179 | 11968259 |
| chr12 | 46237898 | 46239187 |
| chr14 | 46655116 | 46656548 |
| chr1 | 87983655 | 87986412 |
| chr1 | 163297503 | 163299588 |
| chr6 | 79544614 | 79546362 |
| chr10 | 37196165 | 37197705 |
| chr1 | 77942305 | 77944890 |
| chr2 | 160123450 | 160125965 |
| chr12 | 25200815 | 25208335 |
| chr10 | 38649513 | 38651731 |
| chr6 | 135794684 | 135795796 |
| chr22 | 11047329 | 11049345 |
| chr2 | 13817854 | 13819987 |
| chr4 | 138196636 | 138198522 |
| chr15 | 87029936 | 87032560 |

| | | |
|---|---|---|
| chr12 | 90027846 | 90031310 |
| chr2 | 39436100 | 39437938 |
| chr2 | 189025262 | 189026639 |
| chr1 | 103994337 | 103995443 |
| chr6 | 53652386 | 53655500 |
| chr11 | 24841035 | 24843016 |
| chr11 | 59100091 | 59101129 |
| chr6 | 7150853 | 7152752 |
| chr21 | 32910351 | 32912039 |
| chr8 | 46931176 | 46932702 |
| chr1 | 104482817 | 104486586 |
| chr4 | 175799089 | 175803095 |
| chr21 | 9781235 | 9783400 |
| chr6 | 99746267 | 99747840 |
| chr1 | 95345012 | 95348262 |
| chr1 | 86347958 | 86349915 |
| chr4 | 126680101 | 126683362 |
| chr14 | 57006423 | 57009307 |
| chr5 | 767607 | 769083 |
| chr1 | 143257684 | 143259843 |
| chr12 | 71193593 | 71196073 |
| chr1 | 195764696 | 195766545 |
| chr3 | 146639857 | 146642419 |
| chr10 | 58145260 | 58146681 |
| chr9 | 61824463 | 61826239 |
| chr12 | 82997710 | 82999138 |
| chr16 | 33783731 | 33786095 |
| chr17 | 65711758 | 65713140 |
| chrUn_GL000220v1 | 96094 | 97992 |
| chr2 | 155695230 | 155697377 |
| chr7 | 82189431 | 82192033 |
| chr10 | 107935801 | 107938089 |
| chr14 | 37243392 | 37249640 |
| chr10 | 100563316 | 100564815 |
| chr1 | 192235060 | 192237055 |
| chr10 | 55168327 | 55170364 |
| chr10 | 15878674 | 15880029 |
| chr14 | 88258253 | 88259853 |
| chr11 | 32266596 | 32269000 |
| chr7 | 76441361 | 76442901 |
| chr18 | 2607310 | 2609183 |
| chr7 | 64827421 | 64828486 |
| chr7 | 7474063 | 7476320 |
| chr6 | 41218697 | 41219714 |
| chr2 | 63447530 | 63449775 |
| chr14 | 87345822 | 87348667 |
| chr10 | 44764160 | 44766607 |
| chr16 | 76649365 | 76650807 |
| chr6 | 155313335 | 155316441 |

| | | |
|---|---|---|
| chr2 | 189253997 | 189255125 |
| chr8 | 116779195 | 116782229 |
| chr6 | 94665443 | 94667494 |
| chr2 | 68906430 | 68909357 |
| chrY | 9521615 | 9523083 |
| chr8 | 112945543 | 112947229 |
| chr2 | 194065012 | 194068354 |
| chr16 | 46393117 | 46394544 |
| chr4 | 158072979 | 158074323 |
| chr10 | 77391406 | 77393586 |
| chr14 | 25899455 | 25900734 |
| chr10 | 71595472 | 71598889 |
| chr5 | 118259437 | 118261410 |
| chr2 | 179395520 | 179398270 |
| chr1 | 101208031 | 101209235 |
| chr14 | 97400241 | 97401786 |
| chr9 | 74569714 | 74572353 |
| chr10 | 54619898 | 54622368 |
| chr2 | 35663335 | 35665308 |
| chr1 | 92598236 | 92599690 |
| chr6 | 13927156 | 13928798 |
| chr7 | 145292504 | 145294338 |
| chr14 | 61720311 | 61722474 |
| chr5 | 51489316 | 51491296 |
| chr7 | 7704670 | 7705924 |
| chr2 | 162223115 | 162225881 |
| chr10 | 65479707 | 65481838 |
| chr2 | 205826630 | 205828243 |
| chr11 | 43311152 | 43313049 |
| chr20 | 44460431 | 44461862 |
| chr1 | 196673755 | 196676254 |
| chr3 | 77125304 | 77127099 |
| chr1 | 103090742 | 103092631 |
| chr7 | 83312420 | 83315853 |
| chr14 | 54188699 | 54190666 |
| chr10 | 62019528 | 62022344 |
| chr2 | 195725899 | 195727025 |
| chr6 | 49028318 | 49030041 |
| chr2 | 177749293 | 177750956 |
| chr2 | 69957397 | 69958859 |
| chr2 | 88786820 | 88788883 |
| chr2 | 220542823 | 220543824 |
| chr16 | 61652061 | 61654470 |
| chr14 | 19731778 | 19736414 |
| chrX | 128748683 | 128749710 |
| chr2 | 193368143 | 193369908 |
| chr5 | 9533038 | 9534566 |
| chr14 | 32059704 | 32061118 |
| chr16 | 34919029 | 34920754 |

| | | |
|---|---|---|
| chr12 | 77433319 | 77435856 |
| chr11 | 129211742 | 129213192 |
| chr15 | 66497388 | 66498989 |
| chr2 | 103991600 | 103994051 |
| chrY | 9516280 | 9519420 |
| chr10 | 66488689 | 66490854 |
| chr6 | 79079493 | 79081886 |
| chr3 | 95956920 | 95958247 |
| chr1 | 72156920 | 72159289 |
| chr19 | 21379274 | 21380856 |
| chr11 | 4338198 | 4340493 |
| chr13 | 67201678 | 67203598 |
| chr2 | 47906370 | 47908633 |
| chr12 | 61648868 | 61650323 |
| chr4 | 131920658 | 131922790 |
| chr1 | 105545506 | 105547209 |
| chr14 | 39640541 | 39643048 |
| chr19 | 49641820 | 49643658 |
| chr2 | 32553786 | 32555620 |
| chr14 | 50774358 | 50777233 |
| chr2 | 163410720 | 163412941 |
| chr15 | 20137854 | 20138870 |
| chr14 | 35336627 | 35338410 |
| chr2 | 35639018 | 35641058 |
| chrUn_GL000218v1 | 129431 | 131834 |
| chr20 | 29738902 | 29741964 |
| chr5 | 110501095 | 110502492 |
| chr5 | 120253098 | 120256487 |
| chr10 | 102350909 | 102353723 |
| chr7 | 84843641 | 84845899 |
| chr1 | 61835254 | 61836879 |
| chr12 | 75964782 | 75966927 |
| chr14 | 52848159 | 52849681 |
| chr2 | 192175679 | 192178875 |
| chr7 | 94546017 | 94547782 |
| chr12 | 102712733 | 102714821 |
| chr8 | 111768899 | 111772168 |
| chr14 | 70907771 | 70910442 |
| chr9 | 95203543 | 95208193 |
| chr3 | 79036076 | 79038679 |
| chr6 | 74575909 | 74579498 |
| chr2 | 55882469 | 55884062 |
| chr3 | 162318466 | 162319976 |
| chr10 | 79703586 | 79707402 |
| chr16_KI270728v1_ | 1623439 | 1625813 |
| chr10 | 37166190 | 37168315 |
| chr10 | 87770422 | 87772602 |
| chr11 | 80542688 | 80543719 |
| chr22 | 11574431 | 11576157 |

| | | |
|---|---|---|
| chr7 | 89338996 | 89341901 |
| chr14 | 52641252 | 52643351 |
| chr1 | 74197026 | 74200102 |
| chr3 | 110352442 | 110354262 |
| chrY | 9538522 | 9539876 |
| chr10 | 35020958 | 35022033 |
| chr11 | 108331524 | 108334189 |
| chr14 | 42444877 | 42448170 |
| chr2 | 185008452 | 185009528 |
| chr13 | 67017223 | 67018734 |
| chr7 | 79488638 | 79491516 |
| chr20 | 13515126 | 13516496 |
| chr20 | 17989863 | 17991333 |
| chr8 | 127742075 | 127743536 |
| chr12 | 80607926 | 80609900 |
| chr4 | 180048768 | 180050507 |
| chr12 | 74522511 | 74524210 |
| chr7 | 84234542 | 84235884 |
| chr1 | 144723166 | 144724885 |
| chr14 | 37170884 | 37172958 |
| chr1 | 92065521 | 92067146 |
| chr13 | 96678988 | 96681088 |
| chr4 | 34943329 | 34945476 |
| chr5 | 166300753 | 166303467 |
| chr11 | 4902219 | 4904479 |
| chr20 | 29825002 | 29827278 |
| chr7 | 95247150 | 95249051 |
| chr10 | 55229946 | 55232468 |
| chr7 | 91108789 | 91111994 |
| chr12 | 84603557 | 84610780 |
| chr10 | 81328697 | 81330929 |
| chr2 | 47821729 | 47823287 |
| chr14 | 46471177 | 46473204 |
| chr2 | 74995925 | 74997277 |
| chr5 | 81208516 | 81210316 |
| chr10 | 65737966 | 65740237 |
| chr11 | 48972323 | 48973830 |
| chr7 | 82589162 | 82591050 |
| chr9 | 93539882 | 93540881 |
| chr14 | 19345262 | 19347111 |
| chr5 | 67899146 | 67902656 |
| chr6 | 75891734 | 75894492 |
| chr2 | 59258759 | 59261224 |
| chr14 | 26982408 | 26985429 |
| chr14 | 27509346 | 27511415 |
| chr1 | 89640057 | 89642404 |
| chr2 | 194411058 | 194414583 |
| chr10 | 102982219 | 102983586 |
| chr12 | 46555375 | 46558241 |

| | | |
|---|---|---|
| chr11 | 31029627 | 31032367 |
| chr1 | 102906523 | 102907831 |
| chr12 | 81927014 | 81928156 |
| chr3 | 184176483 | 184178074 |
| chr4 | 98142878 | 98145748 |
| chr10 | 101567961 | 101569448 |
| chr1 | 238475027 | 238476632 |
| chr7 | 84026728 | 84030387 |
| chr2 | 224777492 | 224779292 |
| chr3 | 22309891 | 22312155 |
| chr12 | 44296720 | 44299526 |
| chr1 | 149456096 | 149457860 |
| chr2 | 167342225 | 167343707 |
| chr11 | 24537609 | 24541319 |
| chr14 | 33773131 | 33774537 |
| chr19 | 18519621 | 18521350 |
| chr1 | 194317073 | 194319156 |
| chr5 | 28271372 | 28273690 |
| chr1 | 7939537 | 7941794 |
| chr6 | 55724514 | 55726318 |
| chr13 | 83653047 | 83654944 |
| chr5 | 144483870 | 144487222 |
| chr5 | 108613491 | 108615762 |
| chr14 | 54666274 | 54668263 |
| chr20 | 54869265 | 54870455 |
| chr10 | 130109513 | 130111208 |
| chr2 | 179121010 | 179123037 |
| chr6 | 33578938 | 33581495 |
| chr20 | 57332170 | 57333648 |
| chrUn_GL000220v1 | 28365 | 29684 |
| chr14 | 86072065 | 86074751 |
| chr13 | 72303558 | 72305000 |
| chr10 | 65830939 | 65832605 |
| chr1 | 101527043 | 101528233 |
| chr14 | 23296826 | 23299830 |
| chr2 | 103796825 | 103798642 |
| chr4 | 34733040 | 34734817 |
| chr14 | 69397892 | 69399320 |
| chr1 | 72079723 | 72082789 |
| chr6 | 73621513 | 73623181 |
| chr13 | 93885553 | 93887076 |
| chr3 | 147307939 | 147309139 |
| chr7 | 86766364 | 86770553 |
| chr12 | 25224859 | 25227687 |
| chr7 | 81365355 | 81368595 |
| chr2 | 35837051 | 35838934 |
| chr18 | 60612343 | 60613417 |
| chr2 | 101427740 | 101428740 |
| chrY | 24290806 | 24292619 |

| | | |
|---|---|---|
| chr6 | 102219190 | 102220759 |
| chr11 | 113773476 | 113774874 |
| chr2 | 57745547 | 57748759 |
| chr1_KI270765v1_a | 136404 | 140491 |
| chr8 | 75700462 | 75702369 |
| chr10 | 87200948 | 87202248 |
| chr12 | 39491254 | 39493766 |
| chr9 | 62267715 | 62269811 |
| chr10 | 20573552 | 20575059 |
| chr12 | 25985558 | 25988815 |
| chr5 | 58871472 | 58874234 |
| chr11 | 89811839 | 89813929 |
| chr7 | 89841992 | 89844918 |
| chr1 | 92996452 | 92999168 |
| chr12 | 87855222 | 87856590 |
| chr8 | 33512430 | 33513777 |
| chr1 | 99426641 | 99428631 |
| chr13 | 72137889 | 72139539 |
| chr6 | 7239062 | 7240511 |
| chr2 | 19955298 | 19956766 |
| chr1 | 63586812 | 63588730 |
| chr4 | 137998451 | 138000935 |
| chr2 | 189084461 | 189086324 |
| chr15 | 37857018 | 37859480 |
| chr1 | 244407735 | 244409486 |
| chr3 | 167552453 | 167553766 |
| chr10 | 54801839 | 54805471 |
| chr3 | 84955626 | 84956992 |
| chr1 | 157929398 | 157930535 |
| chr1 | 172349104 | 172353320 |
| chr13 | 86578310 | 86580306 |
| chr7 | 89906556 | 89908839 |
| chr12 | 84120609 | 84121979 |
| chr12 | 18139482 | 18141825 |
| chr7 | 85448725 | 85450537 |
| chr14 | 52877267 | 52880413 |
| chr1 | 144767436 | 144768878 |
| chr6 | 51649851 | 51652359 |
| chr9 | 65931592 | 65933511 |
| chr20 | 36310984 | 36312118 |
| chr14 | 83270066 | 83273199 |
| chr18 | 26951365 | 26952541 |
| chr13 | 72252718 | 72255722 |
| chr14 | 68884366 | 68886885 |
| chr10 | 119680883 | 119683416 |
| chr4 | 38856313 | 38858257 |
| chr4 | 111641774 | 111644004 |
| chr2 | 186852565 | 186854953 |
| chr2 | 19901167 | 19903226 |

| | | |
|---|---|---|
| chr7 | 82432850 | 82434662 |
| chr2 | 188934337 | 188936143 |
| chr7 | 76165399 | 76167670 |
| chr12 | 71295198 | 71296977 |
| chr19 | 36139152 | 36141010 |
| chr1 | 218219198 | 218221361 |
| chr14 | 60504313 | 60507281 |
| chr1 | 105507562 | 105509942 |
| chr11 | 89997083 | 89999842 |
| chr1 | 119613483 | 119615502 |
| chr7 | 82111350 | 82112849 |
| chr19 | 50656880 | 50659815 |
| chr6 | 51258959 | 51260942 |
| chr20 | 38520425 | 38521884 |
| chr13 | 35109844 | 35112144 |
| chr2 | 78578194 | 78579851 |
| chr2 | 230869283 | 230871253 |
| chr7 | 79677941 | 79679773 |
| chr16 | 25156619 | 25157743 |
| chr12 | 62293539 | 62294836 |
| chr8 | 65679203 | 65680604 |
| chr2 | 189209833 | 189211633 |
| chr12 | 65952208 | 65954974 |
| chr12 | 71456683 | 71458166 |
| chr12 | 43929216 | 43930220 |
| chr22 | 15792286 | 15796409 |
| chr5 | 3587783 | 3590277 |
| chr3 | 95895290 | 95897338 |
| chr7 | 84539826 | 84541738 |
| chr1 | 219233859 | 219236552 |
| chr2 | 178612156 | 178613237 |
| chr2 | 137152780 | 137154267 |
| chr1 | 101603754 | 101607172 |
| chr6 | 63150387 | 63153087 |
| chr2 | 162563707 | 162566176 |
| chr8 | 32072953 | 32074367 |
| chr9 | 42725185 | 42727871 |
| chr1 | 213302090 | 213304390 |
| chr5 | 18817005 | 18818271 |
| chr1 | 118053598 | 118055338 |
| chr11 | 22729598 | 22731643 |
| chr1 | 157634700 | 157636872 |
| chr8 | 103428526 | 103429610 |
| chr11 | 27675542 | 27676950 |
| chr14 | 89648158 | 89649595 |
| chr5 | 30851612 | 30852694 |
| chr14 | 97569617 | 97571548 |
| chr1 | 154219885 | 154222935 |
| chr10 | 125759907 | 125762260 |

| | | |
|---|---|---|
| chr5 | 111861359 | 111862503 |
| chr4 | 45617724 | 45619259 |
| chr12 | 33708360 | 33710846 |
| chr6 | 100334767 | 100337735 |
| chr14 | 87181123 | 87182350 |
| chr1 | 77348141 | 77350899 |
| chr5 | 143675959 | 143677703 |
| chr14 | 27381092 | 27383126 |
| chr20 | 30398439 | 30400606 |
| chr10 | 91393002 | 91394952 |
| chr3 | 146584576 | 146586534 |
| chr7 | 82068301 | 82070756 |
| chr9 | 31681433 | 31682551 |
| chr11 | 17956797 | 17959068 |
| chr14 | 30213711 | 30215705 |
| chr2 | 83223544 | 83226168 |
| chr15 | 23564478 | 23566551 |
| chr7 | 84513523 | 84514927 |
| chr10 | 63132332 | 63134053 |
| chr13 | 78908337 | 78910198 |
| chr21 | 9940951 | 9942197 |
| chr7 | 80610913 | 80613066 |
| chr2 | 53792677 | 53794979 |
| chr11 | 27498188 | 27499832 |
| chr10 | 121537326 | 121539886 |
| chr17 | 68046787 | 68048722 |
| chr20 | 28575201 | 28577700 |
| chr12 | 85281347 | 85284599 |
| chr9 | 62829914 | 62832045 |
| chr10 | 60274428 | 60276972 |
| chr5 | 27931640 | 27932714 |
| chr4 | 68284416 | 68286222 |
| chr10 | 52258686 | 52260133 |
| chr2 | 37581534 | 37584406 |
| chr1 | 219441136 | 219443859 |
| chr14 | 39973172 | 39975051 |
| chr4 | 35637657 | 35640101 |
| chr14 | 43219826 | 43221471 |
| chr1 | 222618985 | 222620784 |
| chr12 | 801601 | 804409 |
| chr4 | 22131197 | 22132355 |
| chr7 | 83092552 | 83093864 |
| chr8 | 76157943 | 76160711 |
| chr14 | 50472133 | 50473819 |
| chr14 | 89255407 | 89257401 |
| chr4 | 41146398 | 41147646 |
| chr1 | 99408859 | 99411035 |
| chr1 | 62036709 | 62038431 |
| chr16 | 33041460 | 33042540 |

| | | |
|---|---|---|
| chr12 | 45727811 | 45730007 |
| chr10 | 46286554 | 46288066 |
| chr1 | 217884020 | 217886131 |
| chr1 | 91200162 | 91202304 |
| chr2 | 65874157 | 65877201 |
| chr10 | 65893539 | 65896372 |
| chr12 | 121406722 | 121409955 |
| chr14 | 19822095 | 19824880 |
| chr1 | 195939202 | 195941436 |
| chr4 | 8999873 | 9001320 |
| chr10 | 85791586 | 85793308 |
| chr7 | 78199320 | 78200858 |
| chr10 | 71211698 | 71214038 |
| chr13 | 112521585 | 112523690 |
| chr2 | 64533002 | 64534892 |
| chr21 | 31400543 | 31402102 |
| chr20 | 54649046 | 54651322 |
| chr2 | 225673369 | 225675602 |
| chr15 | 37764344 | 37766907 |
| chr14 | 35291776 | 35293082 |
| chr14 | 86702945 | 86704597 |
| chr16 | 77403866 | 77405939 |
| chr2 | 159610757 | 159614776 |
| chr5 | 122401654 | 122404777 |
| chr11 | 30230535 | 30232671 |
| chr4 | 90667569 | 90669317 |
| chr8 | 98509375 | 98511315 |
| chr1 | 86456311 | 86458404 |
| chr10 | 104800329 | 104802263 |
| chr12 | 46370312 | 46372577 |
| chr14 | 48141592 | 48144463 |
| chr2 | 38584536 | 38586245 |
| chr1 | 68219707 | 68222660 |
| chr10 | 34811144 | 34813394 |
| chr3 | 84502373 | 84503898 |
| chr16 | 87302538 | 87304835 |
| chr10 | 86710562 | 86712848 |
| chr10 | 18665622 | 18668633 |
| chr8 | 41305569 | 41308025 |
| chr21 | 10417644 | 10420070 |
| chr2 | 182626167 | 182628611 |
| chr21 | 9018895 | 9021052 |
| chr4 | 32547823 | 32549799 |
| chr16 | 53215344 | 53216427 |
| chr1 | 38012091 | 38013693 |
| chr1 | 104124051 | 104125875 |
| chr11 | 120333786 | 120337075 |
| chr5 | 68214055 | 68216139 |
| chr8 | 128008235 | 128009954 |

| | | |
|---|---|---|
| chr2 | 164231961 | 164234681 |
| chr6 | 80961900 | 80963954 |
| chr9 | 65960572 | 65962791 |
| chr9 | 40970386 | 40971950 |
| chr5 | 138361325 | 138363837 |
| chr1 | 89526457 | 89528159 |
| chr10 | 66499926 | 66501781 |
| chr9 | 10439717 | 10441328 |
| chr3 | 14947691 | 14950111 |
| chr7 | 19113947 | 19115417 |
| chr3 | 94929835 | 94931520 |
| chr10 | 68659477 | 68660658 |
| chr10 | 127424473 | 127426647 |
| chr12 | 76852551 | 76854828 |
| chr2 | 177116729 | 177118588 |
| chr12 | 59201763 | 59203460 |
| chr10 | 114602337 | 114603965 |
| chr5 | 16879779 | 16881569 |
| chr10 | 26944584 | 26946654 |
| chr2 | 185349114 | 185350256 |
| chr20 | 60457038 | 60458442 |
| chr16 | 76978627 | 76982614 |
| chr21 | 16375601 | 16377596 |
| chr14 | 43286113 | 43288264 |
| chr3 | 67309480 | 67311083 |
| chr2 | 164991142 | 164993844 |
| chr10 | 63824527 | 63826994 |
| chr1 | 195879644 | 195881228 |
| chr4 | 176751506 | 176753872 |
| chr1 | 218495616 | 218500052 |
| chr4 | 33361546 | 33363849 |
| chr14 | 92311072 | 92314997 |
| chr20 | 14306784 | 14308792 |
| chr9 | 65814773 | 65818776 |
| chr3 | 114647451 | 114650327 |
| chr13 | 84481865 | 84484754 |
| chr5 | 57220471 | 57221898 |
| chr20 | 20446433 | 20448919 |
| chr3 | 15797472 | 15799941 |
| chr12 | 87217453 | 87220197 |
| chr2 | 57514313 | 57515577 |
| chr10 | 59834150 | 59837297 |
| chr17 | 70768352 | 70770368 |
| chr12 | 42434741 | 42436126 |
| chr14 | 99262754 | 99263927 |
| chr2 | 33277781 | 33280109 |
| chr19 | 9913736 | 9915137 |
| chr6 | 133550609 | 133552741 |
| chr12 | 71918939 | 71921631 |

| | | |
|---|---|---|
| chr2 | 41503885 | 41507632 |
| chr1 | 219403994 | 219406866 |
| chr7 | 6505483 | 6507061 |
| chr12 | 15334289 | 15336288 |
| chr7 | 13990086 | 13992093 |
| chr11 | 24578876 | 24580171 |
| chr6 | 56141622 | 56143348 |
| chr7 | 82029510 | 82031995 |
| chr14 | 19264147 | 19266867 |
| chr9 | 40648752 | 40649959 |
| chr12 | 65492326 | 65495824 |
| chr5 | 113796280 | 113799046 |
| chr1 | 103188016 | 103192319 |
| chr8 | 17905425 | 17908475 |
| chr2 | 51299314 | 51303592 |
| chr14 | 20608486 | 20610005 |
| chr14 | 102200370 | 102202019 |
| chr7 | 66450733 | 66451940 |
| chr5 | 113718082 | 113719421 |
| chr1 | 50043986 | 50045100 |
| chr7 | 93909092 | 93911572 |
| chr1 | 178857686 | 178860141 |
| chr10 | 44378575 | 44380894 |
| chr1 | 191522009 | 191523519 |
| chr14 | 81174453 | 81177111 |
| chr15 | 57486601 | 57488046 |
| chr20 | 38962820 | 38964150 |
| chr14 | 20311246 | 20312779 |
| chr2 | 161765261 | 161767376 |
| chr2 | 197208602 | 197211180 |
| chr2 | 184804219 | 184807140 |
| chr7 | 63721782 | 63723372 |
| chr14 | 43924987 | 43927517 |
| chr12 | 85219655 | 85220875 |
| chr4 | 115937859 | 115939107 |
| chr4 | 156980983 | 156984049 |
| chr21 | 7433489 | 7435935 |
| chr7 | 94403341 | 94405876 |
| chr2 | 70292512 | 70295000 |
| chr1 | 98020207 | 98022285 |
| chr2 | 97710604 | 97712124 |
| chr2 | 36775178 | 36776362 |
| chr6 | 95431441 | 95432944 |
| chr7 | 81268748 | 81270794 |
| chr5 | 160138522 | 160139964 |
| chr6 | 91037412 | 91038672 |
| chr1 | 159369912 | 159371043 |
| chr7 | 61031542 | 61033316 |
| chr22 | 12576593 | 12579204 |

| | | |
|---|---|---|
| chr3 | 83489519 | 83490628 |
| chr2 | 43854696 | 43856928 |
| chr14 | 96519154 | 96520325 |
| chr1 | 87140216 | 87142517 |
| chr12 | 72522294 | 72525261 |
| chr4 | 34950448 | 34952806 |
| chr4 | 156662147 | 156664637 |
| chr10 | 41847911 | 41850787 |
| chr7 | 84436342 | 84439048 |
| chr8 | 82316275 | 82319131 |
| chr2 | 236807885 | 236809884 |
| chr2 | 194534933 | 194536332 |
| chr6 | 51602623 | 51604160 |
| chr2 | 57860389 | 57861969 |
| chr16 | 6556598 | 6557748 |
| chr8 | 102225724 | 102226925 |
| chr19 | 11424048 | 11426038 |
| chr6 | 157310000 | 157311994 |
| chr7 | 77351161 | 77354243 |
| chr10 | 42168957 | 42172208 |
| chr13 | 82186009 | 82188700 |
| chr10 | 51552016 | 51554929 |
| chr8 | 110050333 | 110051787 |
| chr14 | 47870333 | 47873103 |
| chr14 | 62882001 | 62883815 |
| chr11 | 23310460 | 23313129 |
| chr1 | 219616557 | 219618173 |
| chr12 | 75431277 | 75432795 |
| chr3 | 35688544 | 35691384 |
| chr5 | 93584121 | 93587282 |
| chr6 | 49440539 | 49442785 |
| chr12 | 85158621 | 85164487 |
| chr1_KI270765v1_a | 69647 | 73138 |
| chr1 | 102723720 | 102725026 |
| chr14 | 47037162 | 47039703 |
| chr6 | 83197334 | 83198717 |
| chr2 | 205140160 | 205142090 |
| chr2 | 185164662 | 185166405 |
| chr1 | 105224157 | 105226106 |
| chr1 | 8095637 | 8098390 |
| chr14 | 36668623 | 36670700 |
| chr6 | 81160548 | 81163647 |
| chr5 | 51266087 | 51267553 |
| chr8 | 111356238 | 111358267 |
| chr2 | 162479107 | 162481009 |
| chr6 | 81164348 | 81166752 |
| chr6 | 73634934 | 73637176 |
| chr13 | 35471430 | 35473341 |
| chr9 | 40154054 | 40155963 |

| | | |
|---|---|---|
| chr20 | 32108983 | 32110469 |
| chr2 | 164902338 | 164905286 |
| chr7 | 77322359 | 77323824 |
| chr2 | 32492985 | 32494646 |
| chr14 | 46216116 | 46217628 |
| chr12 | 28489549 | 28492096 |
| chr8 | 112959218 | 112962810 |
| chr14 | 86161714 | 86164370 |
| chr13 | 105009038 | 105010495 |
| chr13 | 71534680 | 71536903 |
| chr6 | 45324348 | 45326860 |
| chr10 | 87551231 | 87552965 |
| chr10 | 110557916 | 110559216 |
| chr18 | 62714391 | 62715817 |
| chr1 | 74691521 | 74693460 |
| chr20 | 29483754 | 29485218 |
| chr13 | 69653057 | 69654850 |
| chr4 | 34010351 | 34012574 |
| chr6 | 56219151 | 56221817 |
| chr7 | 91166182 | 91167575 |
| chr10 | 63263037 | 63266673 |
| chr11 | 25690562 | 25692638 |
| chr1 | 179351731 | 179353919 |
| chr16 | 76408487 | 76412087 |
| chr4 | 107255163 | 107257645 |
| chr4 | 35905806 | 35907827 |
| chr14 | 29351597 | 29353454 |
| chr7 | 87809213 | 87812277 |
| chr2 | 83672621 | 83674959 |
| chr19 | 19649257 | 19650856 |
| chr1 | 195728245 | 195729642 |
| chr6 | 133016998 | 133018737 |
| chr9 | 40986929 | 40987984 |
| chr5_KI270897v1_a | 580770 | 582180 |
| chr10 | 64612060 | 64613425 |
| chr12 | 67325391 | 67326491 |
| chr12 | 68110584 | 68112535 |
| chr2 | 165879245 | 165881279 |
| chr14 | 41346491 | 41348647 |
| chr10 | 60352386 | 60355213 |
| chr6 | 83588882 | 83590965 |
| chr14 | 62469888 | 62472026 |
| chr3 | 179449924 | 179452241 |
| chr15 | 37813852 | 37816067 |
| chr6 | 100607246 | 100608473 |
| chr2 | 181171721 | 181172940 |
| chr15 | 37009040 | 37011321 |
| chr5 | 111090489 | 111093279 |
| chr20 | 16572079 | 16573898 |

| | | |
|---|---|---|
| chr9 | 13321602 | 13324941 |
| chr2 | 51160243 | 51162597 |
| chr1 | 92303755 | 92304992 |
| chr14 | 26662630 | 26664456 |
| chr3 | 158643679 | 158645413 |
| chr2 | 160396850 | 160398225 |
| chr12 | 100317699 | 100320580 |
| chr10 | 57600840 | 57602643 |
| chr2 | 58306907 | 58308598 |
| chr14 | 46853365 | 46855442 |
| chr14 | 55321237 | 55323291 |
| chr19 | 46600469 | 46602224 |
| chr10 | 95217276 | 95219239 |
| chr17 | 26769606 | 26772108 |
| chr2 | 182068082 | 182069768 |
| chr6 | 56643018 | 56644714 |
| chr9 | 65324318 | 65325458 |
| chr7 | 83126994 | 83128553 |
| chr10 | 62318228 | 62320090 |
| chr10 | 58775737 | 58777634 |
| chr2 | 204265174 | 204267396 |
| chr11 | 89915774 | 89918400 |
| chr2 | 164909420 | 164914319 |
| chr2 | 186663475 | 186665361 |
| chr3 | 111978321 | 111981304 |
| chr4 | 126727689 | 126730702 |
| chr2 | 158463528 | 158467308 |
| chr11 | 33287354 | 33289304 |
| chr1 | 195079101 | 195080832 |
| chr2 | 187417053 | 187419050 |
| chr16 | 34717523 | 34719722 |
| chr10 | 64302181 | 64303920 |
| chr7 | 85205577 | 85209491 |
| chr10 | 91059387 | 91060462 |
| chr6 | 295604 | 297247 |
| chr7 | 74965111 | 74967501 |
| chr7 | 82604753 | 82606321 |
| chr1 | 189433230 | 189435238 |
| chr14 | 91481033 | 91485486 |
| chr6 | 101894148 | 101895333 |
| chr5 | 150946953 | 150948154 |
| chr7 | 92291582 | 92293099 |
| chr2 | 81372283 | 81374190 |
| chr6 | 44931458 | 44933634 |
| chr5 | 27708275 | 27710920 |
| chr16 | 89121163 | 89123437 |
| chr3 | 145968010 | 145969278 |
| chr8 | 105161378 | 105163341 |
| chr2 | 194866956 | 194870319 |

| | | |
|---|---|---|
| chr8 | 107965002 | 107967235 |
| chr14 | 40580170 | 40583583 |
| chr14 | 25517390 | 25519167 |
| chr13 | 68044399 | 68047496 |
| chr1 | 77218892 | 77221015 |
| chr2 | 177210378 | 177212651 |
| chr5 | 88790993 | 88792198 |
| chr14 | 29057401 | 29059734 |
| chr4 | 49515930 | 49517033 |
| chr2 | 74552221 | 74553758 |
| chr2 | 49382587 | 49384749 |
| chr13 | 28462896 | 28465379 |
| chr7 | 79153670 | 79155491 |
| chr5 | 123916073 | 123918060 |
| chr16 | 46380678 | 46382538 |
| chr14 | 48951121 | 48953142 |
| chr13 | 70991116 | 70994337 |
| chr14 | 86519211 | 86521448 |
| chr1 | 98091081 | 98092837 |
| chr20 | 51801529 | 51804610 |
| chr2 | 42766871 | 42769263 |
| chr2 | 163725727 | 163727345 |
| chr1 | 97840787 | 97844056 |
| chr14 | 35670846 | 35673560 |
| chr14 | 24137458 | 24140710 |
| chr7 | 83808027 | 83810006 |
| chr14 | 30135755 | 30137790 |
| chr7 | 73005400 | 73007455 |
| chr11 | 55382619 | 55384546 |
| chr3 | 74999843 | 75002071 |
| chr1 | 98948836 | 98950260 |
| chr9 | 13269474 | 13271461 |
| chr2 | 164914739 | 164916434 |
| chr17 | 65744090 | 65745712 |
| chr2 | 196455447 | 196457374 |
| chr20 | 63266424 | 63268567 |
| chr12 | 77377924 | 77379934 |
| chr16 | 73745019 | 73747394 |
| chr13 | 64009053 | 64010996 |
| chr2 | 197192541 | 197193720 |
| chr2 | 145025152 | 145029180 |
| chr2 | 194300361 | 194302054 |
| chr2 | 77011717 | 77013788 |
| chr2 | 58698607 | 58701533 |
| chrUn_GL000195v1 | 52596 | 54214 |
| chr7 | 82242601 | 82245795 |
| chr2 | 65690107 | 65691997 |
| chr1 | 102661990 | 102664881 |
| chr16 | 32389887 | 32392155 |

| | | |
|---|---|---|
| chr2 | 66653876 | 66656269 |
| chr4 | 110651525 | 110653030 |
| chr4 | 115100977 | 115102753 |
| chr15 | 27132989 | 27134909 |
| chr7 | 124154508 | 124156534 |
| chr10 | 131175460 | 131178799 |
| chr13 | 38861800 | 38863146 |
| chr2 | 34773665 | 34776591 |
| chr20 | 50930917 | 50932225 |
| chr2 | 98641767 | 98644482 |
| chr12 | 57757446 | 57759458 |
| chr17 | 59892367 | 59893954 |
| chr11 | 38587458 | 38589122 |
| chr1 | 219328293 | 219330719 |
| chr1 | 77681544 | 77684222 |
| chr8 | 135486179 | 135488091 |
| chr10 | 46187897 | 46189794 |
| chr7 | 14393525 | 14395041 |
| chr1 | 106612394 | 106613534 |
| chr14 | 80630116 | 80631304 |
| chr8 | 115621989 | 115624288 |
| chr12 | 89318540 | 89320741 |
| chr2 | 225661899 | 225663939 |
| chr14 | 37735230 | 37739470 |
| chr2 | 34409378 | 34412799 |
| chr1 | 187680855 | 187683828 |
| chr10 | 55937920 | 55940285 |
| chr12 | 106199766 | 106202147 |
| chr7 | 85694423 | 85695798 |
| chr1 | 95792763 | 95794214 |
| chr20 | 36569264 | 36571396 |
| chr12 | 78120597 | 78122071 |
| chr12 | 101615309 | 101617127 |
| chr1 | 95143137 | 95144703 |
| chr2 | 177480767 | 177483364 |
| chr3 | 33797937 | 33799592 |
| chr14 | 43109866 | 43111038 |
| chr5 | 125497251 | 125498730 |
| chr1 | 195378232 | 195379908 |
| chr12 | 26027889 | 26030743 |
| chr4 | 115670870 | 115673612 |
| chr1 | 86710874 | 86714012 |
| chr1 | 187126099 | 187127778 |
| chr16 | 76283688 | 76285298 |
| chr5 | 41152981 | 41154826 |
| chr12 | 71636352 | 71637781 |
| chr6 | 122026311 | 122027764 |
| chr2 | 100932135 | 100934477 |
| chr2 | 50575705 | 50577261 |

| | | |
|---|---|---|
| chr2 | 193949796 | 193953002 |
| chr15 | 22239319 | 22240618 |
| chr20 | 31546632 | 31549390 |
| chr5 | 20810613 | 20812047 |
| chr2 | 77120798 | 77122851 |
| chr2 | 163063935 | 163066044 |
| chr6 | 61907551 | 61909477 |
| chr12 | 29380057 | 29383100 |
| chr1 | 218329520 | 218331926 |
| chr15 | 72116296 | 72119230 |
| chr20 | 59838983 | 59840748 |
| chr5 | 52346072 | 52347818 |
| chr3 | 81870514 | 81872928 |
| chr15 | 40593232 | 40595891 |
| chr1 | 109604345 | 109606373 |
| chr10 | 47885941 | 47887224 |
| chr1 | 28926444 | 28927692 |
| chr2 | 189248327 | 189249853 |
| chr2 | 166127316 | 166130449 |
| chr2 | 33145032 | 33146645 |
| chr1 | 210071990 | 210074967 |
| chr7 | 94540148 | 94542112 |
| chr2 | 59835165 | 59838896 |
| chr15 | 61132765 | 61134807 |
| chr5 | 128445222 | 128446921 |
| chr14 | 98040145 | 98042491 |
| chr16 | 34637946 | 34639111 |
| chr1 | 148596042 | 148599506 |
| chr1 | 33471926 | 33473679 |
| chr13 | 70494429 | 70497204 |
| chr2 | 185637561 | 185639594 |
| chr2 | 199691632 | 199692985 |
| chr10 | 130171108 | 130173803 |
| chr12 | 71791995 | 71794350 |
| chr4 | 20308512 | 20309889 |
| chr8 | 99475186 | 99476705 |
| chr22_KI270733v1_ | 7640 | 8791 |
| chr1 | 109992493 | 109994302 |
| chr1 | 62746191 | 62747342 |
| chr12 | 75039054 | 75041731 |
| chr3 | 130029498 | 130030689 |
| chr12 | 39596893 | 39599323 |
| chr6 | 78900005 | 78901587 |
| chr2 | 82995883 | 82998528 |
| chr4 | 70347257 | 70348588 |
| chr1 | 149479695 | 149482816 |
| chr4 | 137388875 | 137391595 |
| chr10 | 94278804 | 94280848 |
| chr9 | 13350722 | 13354190 |

| | | |
|---|---|---|
| chr2 | 162225899 | 162228217 |
| chr5 | 167103966 | 167106017 |
| chr11 | 97148621 | 97150101 |
| chr6 | 92064397 | 92066722 |
| chr6 | 129428854 | 129429982 |
| chr6 | 57476950 | 57478109 |
| chr4 | 70343482 | 70345360 |
| chr9 | 21554338 | 21557038 |
| chr12 | 80514343 | 80517010 |
| chr10 | 62844622 | 62845959 |
| chr7 | 93464250 | 93465549 |
| chr14 | 20559231 | 20561680 |
| chr14 | 50906060 | 50907596 |
| chr13 | 46438260 | 46440598 |
| chr1 | 99635630 | 99637368 |
| chr10 | 51626375 | 51627933 |
| chr20 | 30217892 | 30219804 |
| chr2 | 208712866 | 208714749 |
| chr10 | 51206469 | 51208363 |
| chr5 | 180351438 | 180353069 |
| chr1 | 83799187 | 83800757 |
| chr5 | 129967461 | 129969697 |
| chr1 | 191885710 | 191887610 |
| chr10 | 57866408 | 57867950 |
| chr2 | 188521960 | 188526719 |
| chr10 | 58651776 | 58653441 |
| chr14 | 90627374 | 90630853 |
| chr9 | 62871232 | 62874055 |
| chr14 | 42356029 | 42359310 |
| chr2 | 187462946 | 187465600 |
| chr2 | 103413422 | 103414718 |
| chr11 | 16403589 | 16406520 |
| chr18 | 65068047 | 65070138 |
| chr2 | 92081205 | 92084381 |
| chr1 | 179503413 | 179507543 |
| chr10 | 57261465 | 57263200 |
| chr6 | 20657049 | 20660606 |
| chr12 | 81037955 | 81040321 |
| chr3 | 22590790 | 22592680 |
| chr2 | 166473576 | 166478358 |
| chr1 | 15571203 | 15573253 |
| chr2 | 40459023 | 40462060 |
| chr6 | 64426474 | 64427971 |
| chr7 | 89612552 | 89614926 |
| chr12 | 66901096 | 66903142 |
| chr14 | 56834634 | 56837747 |
| chr14 | 46833562 | 46835545 |
| chr12 | 85297353 | 85299773 |
| chr2 | 182354738 | 182356801 |

| | | |
|---|---|---|
| chr9 | 12963490 | 12966539 |
| chr19 | 35930937 | 35932972 |
| chr5 | 119915438 | 119917698 |
| chr5 | 29011102 | 29014387 |
| chr12 | 26068167 | 26070646 |
| chr11 | 31159286 | 31161820 |
| chr2 | 79979671 | 79982082 |
| chr1 | 186080768 | 186084906 |
| chr4 | 36386283 | 36388550 |
| chr14 | 27439994 | 27442025 |
| chr12 | 98921171 | 98923380 |
| chr10 | 37171065 | 37173675 |
| chr10 | 79253523 | 79255337 |
| chr7 | 79865397 | 79868245 |
| chr14 | 43762444 | 43764163 |
| chr14 | 47067210 | 47069523 |
| chr10 | 126059254 | 126061442 |
| chr14 | 96975594 | 96977461 |
| chr17 | 22099763 | 22102463 |
| chr7 | 77041524 | 77044244 |
| chr2 | 195130280 | 195131842 |
| chr2 | 63434817 | 63436205 |
| chr18 | 71611881 | 71613409 |
| chr3 | 114962076 | 114963869 |
| chr2 | 180502428 | 180504613 |
| chr9 | 19722775 | 19725171 |
| chr1 | 189447938 | 189449628 |
| chr14 | 49798805 | 49801138 |
| chr2 | 213236277 | 213238101 |
| chr1 | 194904246 | 194905860 |
| chr10 | 66948023 | 66949609 |
| chr21 | 16503529 | 16506645 |
| chr2 | 57173549 | 57175477 |
| chr2 | 80705006 | 80706952 |
| chr6 | 121435827 | 121437029 |
| chr4 | 30964466 | 30967145 |
| chr20 | 33816815 | 33818070 |
| chr2 | 171878814 | 171880145 |
| chr1 | 154944599 | 154946012 |
| chr4 | 43222853 | 43224494 |
| chr14 | 27987960 | 27990353 |
| chr12 | 65957329 | 65959970 |
| chr12 | 20124359 | 20127986 |
| chr16 | 53501508 | 53505175 |
| chr12 | 65565184 | 65567388 |
| chr7 | 84122065 | 84126420 |
| chr1 | 177486565 | 177488200 |
| chr1 | 85196186 | 85197896 |
| chr6 | 94787673 | 94789960 |

| | | |
|---|---|---|
| chr10 | 79666944 | 79670060 |
| chr9 | 4664982 | 4666611 |
| chr7 | 93726148 | 93727420 |
| chr1 | 100452170 | 100453765 |
| chr12 | 59188511 | 59190809 |
| chr14 | 60276638 | 60277886 |
| chr5 | 179209722 | 179212411 |
| chr7 | 81730068 | 81733320 |
| chr8 | 99701565 | 99703199 |
| chr1 | 221030674 | 221033299 |
| chr1 | 238590066 | 238591609 |
| chr12 | 75197336 | 75199114 |
| chr6 | 71264588 | 71265672 |
| chr1 | 173661116 | 173663821 |
| chr2 | 187309309 | 187311116 |
| chr16 | 75966357 | 75969120 |
| chr4 | 157410902 | 157412276 |
| chr18 | 12189315 | 12190713 |
| chr21 | 10542417 | 10544480 |
| chr15 | 37120237 | 37122016 |
| chr15 | 91543480 | 91545202 |
| chr14 | 106800124 | 106802683 |
| chr5 | 60565613 | 60567001 |
| chr14 | 86578910 | 86581913 |
| chr11 | 96011593 | 96013797 |
| chr21 | 16118082 | 16119871 |
| chr13 | 104495985 | 104498239 |
| chr4 | 62825729 | 62827760 |
| chr14 | 80940533 | 80942522 |
| chr6 | 63634959 | 63637262 |
| chr2 | 34066528 | 34068788 |
| chr2 | 194003677 | 194005981 |
| chr10 | 55647461 | 55650274 |
| chr2 | 58537788 | 58539814 |
| chr2 | 56865143 | 56866646 |
| chr15 | 53699166 | 53701628 |
| chr2 | 57740792 | 57742322 |
| chr2 | 104738455 | 104741434 |
| chr2 | 76004435 | 76005953 |
| chr10 | 69966295 | 69970201 |
| chr3 | 140597378 | 140599271 |
| chr10 | 56498992 | 56502577 |
| chr11 | 24174459 | 24176116 |
| chr7 | 100317800 | 100318865 |
| chr8 | 60038870 | 60040360 |
| chr12 | 63428402 | 63430222 |
| chr22 | 10740930 | 10743266 |
| chr14 | 81923999 | 81925749 |
| chr1 | 70615466 | 70617464 |

| | | |
|---|---|---|
| chr14 | 32543303 | 32547072 |
| chr22 | 24284280 | 24286064 |
| chr1 | 191161295 | 191164199 |
| chr2 | 97149388 | 97151669 |
| chr10 | 61216473 | 61218067 |
| chr2 | 193974969 | 193976284 |
| chr13 | 98605818 | 98606892 |
| chr18 | 27431391 | 27432910 |
| chr22 | 11600538 | 11603244 |
| chr2 | 55281759 | 55283884 |
| chr3 | 15813780 | 15815497 |
| chr20 | 44091035 | 44092281 |
| chr22 | 15796849 | 15800263 |
| chr2 | 55893835 | 55895459 |
| chr4 | 59246743 | 59251126 |
| chr12 | 65499547 | 65501512 |
| chr11 | 31262836 | 31264282 |
| chr2 | 71545368 | 71547132 |
| chr2 | 187429276 | 187432724 |
| chr9 | 10119581 | 10121645 |
| chr15 | 53881695 | 53884087 |
| chr10 | 55090423 | 55093362 |
| chr9 | 65163327 | 65165721 |
| chr6 | 73519403 | 73522877 |
| chr4 | 115946505 | 115947957 |
| chr10 | 74290990 | 74292099 |
| chr6 | 54828674 | 54831566 |
| chr8 | 120452589 | 120455988 |
| chr6 | 137203085 | 137204868 |
| chr1 | 85357224 | 85358668 |
| chr15 | 97101415 | 97102692 |
| chr1 | 96621554 | 96623510 |
| chr2 | 189865869 | 189868312 |
| chr14 | 102907048 | 102908498 |
| chr14 | 44927995 | 44929548 |
| chr11 | 23376998 | 23379513 |
| chr1 | 191314240 | 191315775 |
| chr9 | 42444531 | 42445728 |
| chr12 | 41569770 | 41572268 |
| chr16 | 78743726 | 78745834 |
| chr4 | 21031785 | 21035020 |
| chr12 | 89345752 | 89347079 |
| chr7 | 103803628 | 103806317 |
| chr10 | 36997201 | 36999361 |
| chr4 | 115575027 | 115577435 |
| chr14 | 19206897 | 19208956 |
| chr17 | 62377244 | 62378425 |
| chr2 | 111803002 | 111804543 |
| chr2 | 145020978 | 145024527 |

| | | |
|---|---|---|
| chr10 | 78046311 | 78048671 |
| chr1 | 72042272 | 72043551 |
| chr7 | 85535899 | 85538128 |
| chr13 | 101002002 | 101004206 |
| chr2 | 224856902 | 224858574 |
| chr7 | 84006493 | 84009188 |
| chr2 | 176321616 | 176324305 |
| chr2 | 86903763 | 86905138 |
| chr2 | 50585189 | 50587357 |
| chr1 | 241859789 | 241860985 |
| chr10 | 75182079 | 75183608 |
| chr12 | 80250206 | 80252863 |
| chr22 | 16313845 | 16315729 |
| chr9 | 75503337 | 75505362 |
| chr9 | 63011193 | 63012290 |
| chr6 | 55081911 | 55084285 |
| chr8 | 92510287 | 92512404 |
| chr14 | 64025674 | 64027669 |
| chr5 | 123508762 | 123510165 |
| chr16_KI270853v1_ | 2044076 | 2045529 |
| chr8 | 39465481 | 39468238 |
| chr22 | 15843664 | 15845682 |
| chr8 | 116660813 | 116663330 |
| chr5 | 98410397 | 98412477 |
| chr7 | 97571037 | 97573956 |
| chr6 | 92356908 | 92358975 |
| chr7 | 78457912 | 78460531 |
| chr14 | 33926196 | 33928911 |
| chr22 | 15813788 | 15815519 |
| chr6 | 26018445 | 26020856 |
| chr6 | 126658799 | 126660657 |
| chr12 | 19392726 | 19394575 |
| chr20 | 52230185 | 52232653 |
| chr1 | 83225014 | 83227301 |
| chr4 | 36763714 | 36765515 |
| chr6 | 125780136 | 125782121 |
| chr22 | 16136920 | 16138036 |
| chr7 | 83396887 | 83398710 |
| chr10 | 91632140 | 91633772 |
| chr10 | 74969436 | 74975456 |
| chr12 | 43347588 | 43348984 |
| chr14 | 59483787 | 59485404 |
| chr4 | 78136813 | 78138913 |
| chr6 | 21371940 | 21373510 |
| chr4 | 114363340 | 114364344 |
| chr4 | 91263867 | 91266245 |
| chr13 | 64844314 | 64846666 |
| chr14 | 29951763 | 29953092 |
| chr1 | 190038292 | 190040459 |

| | | |
|---|---|---|
| chr1 | 190042568 | 190043925 |
| chr3 | 29328858 | 29331375 |
| chr8 | 84445004 | 84446592 |
| chr16 | 32336302 | 32338991 |
| chr20 | 2745416 | 2748846 |
| chr14 | 87016347 | 87018635 |
| chr3 | 85176923 | 85179895 |
| chr10 | 47629539 | 47632785 |
| chr14 | 38172958 | 38174574 |
| chr4 | 47902231 | 47904265 |
| chr14 | 27935480 | 27937300 |
| chr2 | 56021982 | 56023641 |
| chr16 | 34602432 | 34604272 |
| chr6 | 316387 | 321218 |
| chr8 | 84196387 | 84198710 |
| chr16 | 69625094 | 69627505 |
| chr6 | 55536109 | 55538736 |
| chr2 | 50652544 | 50654263 |
| chr1 | 90852595 | 90853982 |
| chr2 | 189275864 | 189278481 |
| chr6 | 26597979 | 26599993 |
| chrX | 132514170 | 132515182 |
| chr8 | 115999571 | 116001103 |
| chr3 | 85235818 | 85238283 |
| chr10 | 87317317 | 87319173 |
| chr4 | 107242432 | 107245166 |
| chr6 | 83054889 | 83056738 |
| chr2 | 82303607 | 82304875 |
| chr6 | 77292290 | 77294752 |
| chr4 | 156856846 | 156858966 |
| chr5 | 144219490 | 144221074 |
| chr10 | 14994436 | 14995669 |
| chr2 | 161372739 | 161374063 |
| chr12 | 45214991 | 45216486 |
| chr7 | 108524693 | 108526737 |
| chr2 | 157585889 | 157587052 |
| chr14 | 103070064 | 103071845 |
| chr4 | 34289503 | 34291496 |
| chr21 | 9959890 | 9963233 |
| chr1 | 186671835 | 186673421 |
| chr10 | 63918087 | 63921052 |
| chr21 | 9646707 | 9648540 |
| chr7 | 7189218 | 7191398 |
| chr18 | 74875532 | 74877166 |
| chr1 | 192919907 | 192921794 |
| chr1 | 5007841 | 5010440 |
| chr6 | 112801480 | 112803125 |
| chr1 | 203795034 | 203796227 |
| chr2 | 59132434 | 59134219 |

| | | |
|---|---|---|
| chr10 | 59324336 | 59327823 |
| chr15 | 55674270 | 55676168 |
| chr2 | 75500819 | 75502679 |
| chr10 | 50001639 | 50003643 |
| chr20 | 30504465 | 30505466 |
| chr15 | 56470139 | 56471226 |
| chr17 | 7003496 | 7005368 |
| chr17 | 2354532 | 2356144 |
| chr10 | 35198174 | 35201468 |
| chr19 | 39362989 | 39365389 |
| chr1 | 110482960 | 110485155 |
| chr6 | 154099896 | 154101736 |
| chr15_KI270850v1_ | 135465 | 136997 |
| chr20 | 59891914 | 59893487 |
| chr1 | 25997220 | 26000520 |
| chr12 | 90097503 | 90099109 |
| chr15_KI270852v1_ | 443485 | 445452 |
| chr2 | 162037549 | 162039366 |
| chr16 | 19701964 | 19703133 |
| chr1 | 119759299 | 119761229 |
| chr22 | 10965780 | 10967010 |
| chr4 | 45113937 | 45115929 |
| chr3 | 174190129 | 174192419 |
| chr1 | 63009162 | 63012149 |
| chr7 | 46354495 | 46357210 |
| chr12 | 80402144 | 80405907 |
| chr2 | 203066843 | 203068768 |
| chr3 | 147574011 | 147576920 |
| chr9 | 31389261 | 31391697 |
| chr13 | 25336023 | 25337908 |
| chr14 | 33585180 | 33587068 |
| chr10 | 99773778 | 99776362 |
| chr10 | 105329112 | 105330479 |
| chr12 | 79128373 | 79130178 |
| chr10 | 109161455 | 109162609 |
| chr14 | 29096196 | 29097950 |
| chr1 | 161661255 | 161663935 |
| chr4 | 35097111 | 35099644 |
| chr15 | 96691180 | 96693178 |
| chr3 | 21266634 | 21268267 |
| chr11 | 85595161 | 85596355 |
| chr1 | 98189888 | 98191369 |
| chr6 | 92496569 | 92498199 |
| chr18 | 72106498 | 72108199 |
| chr16 | 35488749 | 35493142 |
| chr7 | 84182409 | 84185042 |
| chr1 | 90045922 | 90047250 |
| chr4 | 139943064 | 139944247 |
| chr5 | 27949898 | 27951445 |

| | | |
|---|---|---|
| chr12 | 68562531 | 68563977 |
| chr16 | 65127466 | 65129775 |
| chr16 | 73886411 | 73888794 |
| chr2 | 90397343 | 90399253 |
| chr16 | 84069567 | 84072289 |
| chr4 | 29223321 | 29224854 |
| chr1 | 178703176 | 178705216 |
| chr7 | 92797739 | 92799251 |
| chr12 | 84951115 | 84953344 |
| chr6 | 99755812 | 99757370 |
| chr1 | 196372114 | 196373677 |
| chr12 | 45122736 | 45125253 |
| chr5 | 165808006 | 165810554 |
| chr1 | 79739569 | 79740950 |
| chr1 | 60024980 | 60026279 |
| chr6 | 166020534 | 166021545 |
| chr10 | 31926894 | 31928370 |
| chr15 | 93768652 | 93769866 |
| chr2 | 103021445 | 103023669 |
| chr16 | 53914167 | 53915710 |
| chr3 | 83629530 | 83630908 |
| chr2 | 179236810 | 179239108 |
| chr20 | 18552832 | 18554914 |
| chr1 | 161205746 | 161207527 |
| chr9 | 41229957 | 41232118 |
| chr2 | 191385555 | 191387713 |
| chr9 | 39998645 | 39999665 |
| chr11 | 16434459 | 16435876 |
| chr20 | 25142179 | 25143960 |
| chr10 | 65576063 | 65579638 |
| chr10 | 91924207 | 91926539 |
| chr12 | 85265425 | 85267447 |
| chr11 | 19142433 | 19144078 |
| chr1 | 94473914 | 94477040 |
| chr5 | 34934988 | 34936514 |
| chr14 | 50727001 | 50729242 |
| chr14 | 55381537 | 55383146 |
| chr11 | 845537 | 846939 |
| chr2 | 203393269 | 203394955 |
| chr2 | 49455475 | 49457385 |
| chr14 | 88657420 | 88659639 |
| chr2 | 59462314 | 59463698 |
| chr6 | 41117915 | 41119468 |
| chr4 | 47785 | 50726 |
| chr14 | 20114294 | 20116188 |
| chr10 | 74896254 | 74897644 |
| chr8 | 94248641 | 94251124 |
| chr10 | 88594857 | 88597529 |
| chr6 | 73133213 | 73135107 |

| | | |
|---|---|---|
| chr7 | 91896482 | 91898258 |
| chr6 | 23676956 | 23678932 |
| chr5 | 148133415 | 148135465 |
| chr8 | 77343666 | 77345458 |
| chr7 | 88681425 | 88683059 |
| chr2 | 177896440 | 177897944 |
| chr2 | 207972882 | 207975093 |
| chr2 | 162889309 | 162891993 |
| chr1 | 103157995 | 103163179 |
| chr13 | 64580961 | 64583964 |
| chr7 | 40764601 | 40765678 |
| chr2 | 44877809 | 44880786 |
| chr20 | 17871834 | 17873646 |
| chr2 | 37390221 | 37391851 |
| chr7 | 52451081 | 52453832 |
| chr9 | 66018265 | 66019994 |
| chr1 | 199078672 | 199081014 |
| chr2 | 146035366 | 146037771 |
| chr7 | 112205970 | 112207904 |
| chr9 | 20993838 | 20995572 |
| chr11 | 106109900 | 106112106 |
| chr14 | 30123870 | 30126222 |
| chr12 | 90815611 | 90817810 |
| chr2 | 87182324 | 87184720 |
| chr4 | 36584694 | 36586336 |
| chr7 | 92213751 | 92215105 |
| chr22 | 15645540 | 15647170 |
| chr8 | 82731827 | 82733864 |
| chr1 | 149406873 | 149409512 |
| chr9 | 63864004 | 63865145 |
| chr9 | 66314852 | 66316763 |
| chr3 | 80716155 | 80717917 |
| chr1 | 79399661 | 79401087 |
| chr5 | 129554458 | 129556827 |
| chr1 | 172311396 | 172313893 |
| chr20 | 30867666 | 30868880 |
| chr6 | 48970059 | 48972153 |
| chr15 | 75283023 | 75284188 |
| chr5 | 123752889 | 123755250 |
| chr15 | 74433771 | 74436960 |
| chr14 | 74969300 | 74971237 |
| chr4 | 93788344 | 93790219 |
| chr3 | 101675834 | 101678184 |
| chr3 | 163365077 | 163367334 |
| chr2 | 178589033 | 178591260 |
| chr2 | 62641388 | 62643965 |
| chr3 | 28425908 | 28427423 |
| chr2 | 229442473 | 229444387 |
| chr14 | 33791405 | 33794651 |

| | | |
|---|---|---|
| chr2 | 194953137 | 194954771 |
| chr10 | 58513268 | 58515094 |
| chr2 | 70682095 | 70683268 |
| chr1 | 247673404 | 247675493 |
| chr12 | 84862675 | 84864224 |
| chr6 | 132852341 | 132854207 |
| chr1 | 98766640 | 98768398 |
| chr2 | 180334808 | 180336702 |
| chr10 | 65630567 | 65631831 |
| chr2 | 39900910 | 39903056 |
| chr14 | 46664758 | 46666160 |
| chr2 | 207196699 | 207197976 |
| chr1 | 62866842 | 62869135 |
| chr20 | 11038266 | 11040056 |
| chr2 | 209138721 | 209141318 |
| chr22 | 11296788 | 11298787 |
| chr19 | 42422641 | 42424141 |
| chr3 | 110772789 | 110774635 |
| chr11 | 101125265 | 101127283 |
| chr12 | 82988934 | 82992200 |
| chr10 | 94576879 | 94578041 |
| chr2_GL383522v1_a | 41887 | 43802 |
| chr10 | 48521015 | 48523997 |
| chr11 | 49844430 | 49846178 |
| chr21 | 20951616 | 20953212 |
| chr1 | 232514560 | 232516091 |
| chr2 | 53432796 | 53435928 |
| chr1 | 219278746 | 219280920 |
| chr1 | 199127644 | 199129540 |
| chr8 | 38968844 | 38971473 |
| chr7 | 90163634 | 90166773 |
| chr1 | 180631215 | 180632697 |
| chr8 | 86813159 | 86817076 |
| chr6 | 126545764 | 126547632 |
| chr10 | 52332126 | 52336245 |
| chr5 | 27700204 | 27701258 |
| chr10 | 16938056 | 16939520 |
| chr16 | 74873436 | 74874918 |
| chr2 | 42285634 | 42287206 |
| chr6 | 15547028 | 15549188 |
| chr7 | 77409103 | 77411350 |
| chr13 | 91509048 | 91510580 |
| chr2 | 154709035 | 154710462 |
| chr4 | 46039920 | 46041844 |
| chr14 | 88990854 | 88993108 |
| chr4 | 71932845 | 71935141 |
| chr4 | 27647021 | 27650044 |
| chr11 | 67507469 | 67509300 |
| chr1 | 185153319 | 185155279 |

| | | |
|---|---|---|
| chr9 | 65312017 | 65315650 |
| chr1 | 103844426 | 103846259 |
| chr2 | 203013910 | 203016335 |
| chr1 | 86359680 | 86361288 |
| chr2 | 55864984 | 55867182 |
| chr4 | 23630298 | 23631816 |
| chr10 | 53964155 | 53965419 |
| chr14 | 50462590 | 50463639 |
| chr12 | 66203489 | 66205049 |
| chr1 | 8142478 | 8143616 |
| chr3 | 147086118 | 147089128 |
| chr11 | 108565367 | 108567508 |
| chr6 | 132912165 | 132913969 |
| chrUn_GL000195v1 | 28576 | 30522 |
| chr10 | 75629391 | 75632080 |
| chr15 | 38250675 | 38252657 |
| chr6 | 57717508 | 57718677 |
| chr14 | 25419024 | 25422441 |
| chr15 | 98922194 | 98924611 |
| chr15 | 20288936 | 20290190 |
| chr3 | 9730184 | 9731832 |
| chr14 | 40641427 | 40645106 |
| chr4 | 28718299 | 28719406 |
| chr14 | 84422869 | 84424092 |
| chr17 | 78359224 | 78361451 |
| chr10 | 61685052 | 61687417 |
| chr6 | 45491009 | 45493853 |
| chrY | 56877860 | 56880147 |
| chr11 | 23543646 | 23546358 |
| chr14 | 48896935 | 48898749 |
| chr7 | 138537881 | 138539509 |
| chr4 | 9209507 | 9213410 |
| chr10 | 116559627 | 116561509 |
| chr2 | 211670345 | 211672123 |
| chr14 | 41870403 | 41871740 |
| chr3 | 81315001 | 81317778 |
| chr4 | 116135635 | 116137587 |
| chr5 | 59655999 | 59658508 |
| chr20 | 48744803 | 48746235 |
| chr2 | 172098270 | 172101079 |
| chr1 | 216938122 | 216940094 |
| chr14 | 21602120 | 21603781 |
| chr4 | 154026717 | 154027731 |
| chr1 | 111448145 | 111450596 |
| chr2 | 184295681 | 184297631 |
| chr5 | 40305698 | 40307563 |
| chr14 | 32445647 | 32447633 |
| chr10 | 70169066 | 70171367 |
| chr20 | 43147861 | 43149455 |

| | | |
|---|---|---|
| chr12 | 83465377 | 83467683 |
| chr2 | 132275608 | 132277821 |
| chr8 | 84472558 | 84475288 |
| chr22 | 12177400 | 12179305 |
| chr2 | 50634345 | 50638351 |
| chr2 | 234591347 | 234592933 |
| chr16 | 74622528 | 74624778 |
| chr8 | 92805350 | 92807309 |
| chr4 | 34311297 | 34312811 |
| chr5 | 129391052 | 129394487 |
| chr6 | 70164114 | 70166055 |
| chr7 | 89742190 | 89743461 |
| chr15 | 20645478 | 20646783 |
| chr2 | 50645307 | 50647694 |
| chr6 | 74062773 | 74065507 |
| chr18 | 14351592 | 14354098 |
| chr6 | 153002230 | 153004850 |
| chr2 | 187983774 | 187984795 |
| chr2 | 184825863 | 184828367 |
| chr1 | 176620505 | 176622991 |
| chr14 | 80698979 | 80701613 |
| chr2 | 57760348 | 57761481 |
| chr4 | 35289440 | 35292053 |
| chr14 | 32858093 | 32860200 |
| chr2 | 41869396 | 41871506 |
| chr4 | 36722159 | 36723594 |
| chr7 | 94123352 | 94125156 |
| chr11 | 101035403 | 101037541 |
| chr16_KI270728v1_ | 219025 | 220419 |
| chr2 | 65296145 | 65297302 |
| chr14 | 21434644 | 21436940 |
| chr14 | 28984605 | 28986366 |
| chr4 | 126694250 | 126695850 |
| chr1 | 101981325 | 101984390 |
| chr11 | 24501734 | 24504078 |
| chr14 | 92307725 | 92309370 |
| chr17 | 48245748 | 48248258 |
| chr10 | 115431299 | 115432456 |
| chr12 | 124446843 | 124448143 |
| chr1 | 101963137 | 101966527 |
| chr2 | 213371166 | 213373658 |
| chr2 | 211497255 | 211498492 |
| chr8 | 76171979 | 76176019 |
| chr14 | 47780530 | 47783435 |
| chr7 | 79840900 | 79843848 |
| chr10 | 38350620 | 38352980 |
| chr2 | 39306783 | 39308136 |
| chr12 | 86392553 | 86394175 |
| chr4 | 189932493 | 189933514 |

| | | |
|---|---|---|
| chr7 | 78614741 | 78616691 |
| chr15 | 85381585 | 85383284 |
| chr1 | 104985462 | 104987530 |
| chr2 | 188734104 | 188735768 |
| chr2 | 103737141 | 103739582 |
| chr6 | 121888634 | 121891552 |
| chr15_KI270905v1_ | 838944 | 840687 |
| chr5 | 132706399 | 132707540 |
| chr5 | 175533700 | 175534938 |
| chr4 | 20544139 | 20545784 |
| chr2 | 176668275 | 176669538 |
| chr2 | 39249176 | 39250326 |
| chr6 | 62655675 | 62657671 |
| chr4 | 168960065 | 168961081 |
| chr10 | 105062018 | 105064462 |
| chr20 | 40929220 | 40930402 |
| chr7 | 54075236 | 54076713 |
| chr13 | 83103131 | 83104870 |
| chr10 | 59190530 | 59192139 |
| chr12 | 76826583 | 76828228 |
| chr7 | 78797375 | 78799990 |
| chr17_GL000258v2_ | 320731 | 323304 |
| chr7 | 65653739 | 65655325 |
| chr4 | 70361655 | 70364490 |
| chr2 | 50143943 | 50145298 |
| chr6 | 2104009 | 2108308 |
| chr1 | 191101573 | 191103436 |
| chr1 | 233984633 | 233985666 |
| chr2 | 45642633 | 45644571 |
| chr7 | 95909885 | 95912390 |
| chr14 | 47614998 | 47616119 |
| chr10 | 129657277 | 129659398 |
| chr7 | 147698472 | 147699665 |
| chr12 | 91345630 | 91348126 |
| chr5_KI270897v1_a | 68643 | 70232 |
| chr10 | 99435679 | 99437386 |
| chr22 | 15525399 | 15526953 |
| chr20 | 52038852 | 52040405 |
| chr11 | 18525172 | 18527081 |
| chr20 | 44905298 | 44907531 |
| chr2 | 31881867 | 31883749 |
| chr16 | 76552016 | 76555186 |
| chr6 | 80195585 | 80196964 |
| chr2 | 201732474 | 201733722 |
| chr14 | 36311261 | 36313055 |
| chr5_KI270897v1_a | 94779 | 98753 |
| chr2 | 177140627 | 177144504 |
| chr2 | 64653610 | 64655718 |
| chr6 | 64310475 | 64311976 |

| | | |
|---|---|---|
| chr2 | 53504868 | 53506084 |
| chr2 | 194791667 | 194793961 |
| chr12 | 68659935 | 68661462 |
| chr7 | 94407548 | 94409945 |
| chr12 | 44301653 | 44304216 |
| chr2 | 32863866 | 32865340 |
| chr15 | 64424121 | 64426751 |
| chr2 | 179599453 | 179603406 |
| chr6 | 79233471 | 79235317 |
| chr14 | 86626157 | 86628277 |
| chr1 | 10905135 | 10906331 |
| chr15 | 36170929 | 36173054 |
| chr19 | 8741885 | 8743766 |
| chrY | 56864667 | 56866729 |
| chr20 | 59907335 | 59909722 |
| chr22 | 11036381 | 11038440 |
| chr11 | 82105291 | 82107446 |
| chr10 | 115276002 | 115278442 |
| chr3 | 147413355 | 147415408 |
| chr9 | 66260385 | 66262236 |
| chr12 | 85364072 | 85365708 |
| chr7 | 88587441 | 88589154 |
| chr12 | 57724444 | 57726511 |
| chr12 | 78123016 | 78124759 |
| chr1 | 239688519 | 239690017 |
| chr21 | 9730185 | 9734527 |
| chr16 | 70534041 | 70536010 |
| chr2 | 188875972 | 188878980 |
| chr8 | 109432409 | 109433806 |
| chr16 | 75885063 | 75886983 |
| chr14 | 27480322 | 27482353 |
| chr1 | 189510056 | 189512845 |
| chr14 | 38008975 | 38010343 |
| chr5 | 170248993 | 170250362 |
| chr2 | 178972744 | 178974642 |
| chr7 | 101805629 | 101806890 |
| chr1 | 16702482 | 16705429 |
| chr2 | 78512262 | 78515120 |
| chr2 | 178741137 | 178742637 |
| chr7 | 85586028 | 85588172 |
| chr1 | 90378470 | 90380143 |
| chr6 | 132434094 | 132435475 |
| chr7 | 72121068 | 72122297 |
| chr8 | 12020220 | 12022325 |
| chr2 | 172093241 | 172095150 |
| chr2 | 91411056 | 91423045 |
| chr1 | 169958417 | 169960646 |
| chr3 | 211929 | 214914 |
| chr3 | 36990864 | 36994470 |

| | | |
|---|---|---|
| chr7 | 53418172 | 53421412 |
| chr10 | 21491147 | 21493087 |
| chr8 | 112294999 | 112296608 |
| chr11 | 90217679 | 90219156 |
| chr5 | 11601224 | 11603411 |
| chr12 | 84860016 | 84862051 |
| chr1 | 149106636 | 149108809 |
| chr13 | 71959346 | 71962084 |
| chrY | 9542170 | 9543212 |
| chr1 | 190151610 | 190153593 |
| chr8 | 114885111 | 114887355 |
| chr4 | 136058032 | 136060047 |
| chr6 | 87339396 | 87341957 |
| chr2 | 39134216 | 39137344 |
| chr10 | 67254958 | 67257881 |
| chr14 | 27093247 | 27096002 |
| chr7 | 8684021 | 8685273 |
| chr1 | 98378306 | 98380488 |
| chr2 | 60438390 | 60440619 |
| chr2 | 103446902 | 103449258 |
| chr2 | 100993189 | 100995173 |
| chr7 | 64410125 | 64411308 |
| chr2 | 52562171 | 52564083 |
| chr11 | 105613323 | 105616262 |
| chr1 | 181288876 | 181291510 |
| chr2 | 161942890 | 161945428 |
| chr2 | 217694450 | 217696814 |
| chr4 | 67669948 | 67671845 |
| chr1 | 98292331 | 98293697 |
| chr9 | 39702881 | 39704993 |
| chr1 | 149810601 | 149813012 |
| chr10 | 124986792 | 124988750 |
| chr14 | 77706421 | 77708154 |
| chr10 | 73680965 | 73683442 |
| chr4 | 27662884 | 27665537 |
| chr10 | 87783283 | 87786022 |
| chr5 | 28673484 | 28675313 |
| chr14 | 63437222 | 63439182 |
| chr9 | 12972415 | 12975073 |
| chr11 | 49591879 | 49595228 |
| chr17 | 68191355 | 68193300 |
| chr1 | 86732314 | 86734095 |
| chr9 | 68263360 | 68265032 |
| chr2 | 136167749 | 136169680 |
| chr6 | 93337889 | 93339830 |
| chr11 | 33165891 | 33168086 |
| chr2 | 50713458 | 50715588 |
| chr10 | 56613112 | 56614170 |
| chr7 | 80577340 | 80579310 |

| | | |
|---|---|---|
| chr4 | 46762218 | 46763898 |
| chr15_KI270905v1_ | 601010 | 602713 |
| chr11 | 87052774 | 87054562 |
| chr10 | 67047926 | 67051264 |
| chr2 | 241351823 | 241353600 |
| chr15 | 36438358 | 36441001 |
| chr1 | 214910871 | 214911904 |
| chr2 | 212182837 | 212185452 |
| chr12 | 85774102 | 85776776 |
| chr10 | 130480185 | 130482225 |
| chr11 | 91264392 | 91266623 |
| chr1 | 244049494 | 244051350 |
| chr1 | 9909918 | 9911557 |
| chr14 | 19798772 | 19800010 |
| chr10 | 46526795 | 46529214 |
| chr6 | 153031281 | 153032888 |
| chr14 | 39136083 | 39137868 |
| chr10 | 65235839 | 65237237 |
| chr5 | 65983285 | 65984650 |
| chr1 | 103104330 | 103107074 |
| chr6 | 2550223 | 2552213 |
| chr2 | 58872485 | 58874619 |
| chr12 | 43796440 | 43798041 |
| chr1 | 149612410 | 149613880 |
| chr15 | 88454736 | 88456330 |
| chr2 | 226123034 | 226126015 |
| chr10 | 45891056 | 45893340 |
| chr10 | 67662901 | 67666959 |
| chr1 | 103982801 | 103984500 |
| chr22 | 15782112 | 15783404 |
| chr1 | 233546992 | 233548939 |
| chr14 | 88181857 | 88185013 |
| chr1 | 90667363 | 90669394 |
| chr14 | 42389002 | 42391774 |
| chr9 | 62890635 | 62892630 |
| chr7 | 79659452 | 79661451 |
| chr7 | 75344468 | 75345978 |
| chr14 | 33352479 | 33355368 |
| chr2 | 204970294 | 204972555 |
| chr11 | 26597588 | 26599131 |
| chr11 | 119014319 | 119016137 |
| chr20 | 3933007 | 3934784 |
| chr6 | 72200046 | 72204582 |
| chr14 | 48857835 | 48859623 |
| chr2 | 233417553 | 233419455 |
| chr5 | 92400366 | 92402316 |
| chr14 | 19757300 | 19760240 |
| chr9 | 138238363 | 138241436 |
| chr15 | 95248709 | 95250770 |

| | | |
|---|---|---|
| chr7 | 90206995 | 90208273 |
| chr2 | 78329299 | 78330425 |
| chr2 | 103294648 | 103297065 |
| chr2 | 193193643 | 193196246 |
| chr12 | 81768170 | 81770248 |
| chr17 | 65676604 | 65680632 |
| chr13 | 38094770 | 38096480 |
| chr22 | 16426633 | 16428866 |
| chr4 | 125949862 | 125951998 |
| chr10 | 58626648 | 58628045 |
| chr10 | 53671462 | 53674257 |
| chr8 | 105637181 | 105639584 |
| chr2 | 91441795 | 91445603 |
| chr6 | 905547 | 907450 |
| chr1 | 219209199 | 219212476 |
| chr6 | 96533479 | 96535123 |
| chr7 | 88463252 | 88465146 |
| chr4 | 109126227 | 109127721 |
| chr1 | 182081383 | 182082517 |
| chr7 | 67290651 | 67291659 |
| chr10 | 82872716 | 82874133 |
| chr22 | 10944486 | 10946448 |
| chr2 | 64083466 | 64085046 |
| chr2 | 162228539 | 162233888 |
| chr14 | 39489424 | 39491009 |
| chr13 | 92339327 | 92341687 |
| chr3 | 78670235 | 78672215 |
| chr13 | 87806119 | 87810213 |
| chr14 | 32150334 | 32151664 |
| chr6 | 46128817 | 46130992 |
| chr8 | 111121332 | 111122693 |
| chr7 | 141797962 | 141800070 |
| chr7 | 85696353 | 85697859 |
| chr10 | 47604262 | 47605516 |
| chr2 | 55172162 | 55174431 |
| chr9 | 65809682 | 65811115 |
| chr14 | 43381450 | 43384768 |
| chr7 | 8723577 | 8726052 |
| chr1 | 104641524 | 104644190 |
| chr4 | 32794321 | 32796304 |
| chr12 | 109017892 | 109020076 |
| chr14 | 48397856 | 48399852 |
| chr10 | 84896185 | 84897654 |
| chr14 | 20970182 | 20971404 |
| chr7 | 84522775 | 84525517 |
| chr20 | 16878631 | 16881085 |
| chr9 | 80710341 | 80712272 |
| chr10 | 63175841 | 63178089 |
| chr14 | 22509032 | 22510670 |

| | | |
|---|---|---|
| chr7 | 80267103 | 80269106 |
| chr16 | 74407999 | 74410951 |
| chr20 | 61603067 | 61604877 |
| chr1 | 27487631 | 27488964 |
| chr10 | 52993496 | 52995677 |
| chr8 | 106509640 | 106511502 |
| chr14 | 41272755 | 41274009 |
| chr12 | 78918604 | 78921079 |
| chr14 | 49870754 | 49871851 |
| chr2 | 48620693 | 48622595 |
| chr20 | 31541266 | 31543674 |
| chr8 | 90802354 | 90804614 |
| chr14 | 37003460 | 37007474 |
| chr3 | 85094190 | 85097112 |
| chr17 | 29431484 | 29432484 |
| chr2 | 161554382 | 161555708 |
| chr2 | 18559319 | 18561301 |
| chr10 | 22322554 | 22325671 |
| chr14 | 85545205 | 85547789 |
| chr1 | 219339935 | 219342563 |
| chr17 | 22372224 | 22373238 |
| chr2 | 58524276 | 58526574 |
| chr10 | 65033264 | 65034364 |
| chr1 | 157042934 | 157045822 |
| chr10 | 113905969 | 113908845 |
| chr20 | 10739733 | 10741484 |
| chr8 | 84723855 | 84726268 |
| chr7_KI270809v1_a | 24259 | 25974 |
| chr8 | 105772228 | 105774719 |
| chr2 | 145489871 | 145491668 |
| chr1 | 109104731 | 109107953 |
| chr2 | 36379309 | 36380737 |
| chr12 | 79684705 | 79687467 |
| chr1 | 101812210 | 101813841 |
| chr2 | 87170157 | 87172217 |
| chr1 | 93909039 | 93910952 |
| chr2 | 57123147 | 57124625 |
| chr5 | 107646325 | 107647794 |
| chr17 | 18616713 | 18618242 |
| chr2 | 46298597 | 46301512 |
| chr1 | 185770094 | 185772292 |
| chr4 | 29797785 | 29800105 |
| chr1 | 198756315 | 198759122 |
| chr15 | 60202648 | 60204978 |
| chr4 | 35690387 | 35692589 |
| chr6 | 60904688 | 60906235 |
| chr15 | 35998151 | 36000599 |
| chr15 | 49179838 | 49181469 |
| chr14 | 26464384 | 26465634 |

| | | |
|---|---|---|
| chr5_KI270897v1_a | 45542 | 47950 |
| chr3 | 115812429 | 115815128 |
| chr12 | 30685568 | 30687655 |
| chr13 | 81975227 | 81977344 |
| chr2 | 174591849 | 174592869 |
| chr10 | 62000918 | 62002940 |
| chr10 | 96192341 | 96193987 |
| chr1 | 192826556 | 192828322 |
| chr10 | 64976531 | 64979231 |
| chr10 | 89754059 | 89755821 |
| chr14 | 44362845 | 44364757 |
| chr6 | 54643108 | 54644433 |
| chr16 | 32127016 | 32128501 |
| chr11 | 21183037 | 21185425 |
| chr2 | 65086104 | 65088545 |
| chr9 | 4639063 | 4640859 |
| chr21 | 7404592 | 7405799 |
| chr11 | 111601562 | 111604685 |
| chr10 | 53046254 | 53049195 |
| chr1 | 215878403 | 215880268 |
| chr9 | 62798553 | 62800999 |
| chr6 | 54452824 | 54455489 |
| chr11 | 96391523 | 96394175 |
| chr14 | 44802955 | 44804298 |
| chr9 | 64490329 | 64492486 |
| chr5 | 142324618 | 142327410 |
| chr4 | 156596240 | 156600570 |
| chr14 | 39445365 | 39447234 |
| chr12 | 39966190 | 39967690 |
| chr6 | 44867933 | 44870356 |
| chr20 | 56975785 | 56977450 |
| chr1 | 185823824 | 185825547 |
| chr3 | 148001485 | 148002726 |
| chr9 | 12978565 | 12981370 |
| chr1 | 217470073 | 217473321 |
| chr5_KI270897v1_a | 749820 | 751614 |
| chr10 | 75020960 | 75022507 |
| chr4 | 79569179 | 79571748 |
| chr10 | 92007646 | 92009350 |
| chr10 | 54085058 | 54087589 |
| chr1 | 161906793 | 161909218 |
| chr4 | 1519340 | 1520915 |
| chr2 | 35782035 | 35783457 |
| chr11 | 35280828 | 35283435 |
| chr5 | 108541319 | 108543506 |
| chr2 | 57725523 | 57727705 |
| chr1 | 194167754 | 194170003 |
| chr7 | 134487769 | 134488973 |
| chr6 | 57361009 | 57363242 |

| | | |
|---|---|---|
| chr14 | 104767294 | 104770035 |
| chr6 | 5470271 | 5471897 |
| chr2 | 94714535 | 94716483 |
| chr1 | 195199908 | 195201748 |
| chr8 | 112836466 | 112840373 |
| chr14 | 36470722 | 36474557 |
| chr3 | 94458847 | 94461569 |
| chr12 | 18341844 | 18343512 |
| chr1 | 72264331 | 72265847 |
| chr1 | 145078365 | 145080239 |
| chr4 | 31648534 | 31650090 |
| chr20 | 30826669 | 30828177 |
| chr16 | 34093421 | 34095321 |
| chr1 | 187809325 | 187811532 |
| chr3 | 187655984 | 187657809 |
| chr17 | 21827622 | 21829802 |
| chr8 | 68762032 | 68763523 |
| chr1 | 70841259 | 70844124 |
| chr1 | 87049952 | 87051927 |
| chr3 | 2314702 | 2317027 |
| chr17 | 65920987 | 65922921 |
| chr10 | 85576113 | 85577178 |
| chr14 | 21433249 | 21434418 |
| chr11 | 105228455 | 105230382 |
| chr2 | 34999322 | 35001351 |
| chr16 | 54923365 | 54925001 |
| chr1 | 195223633 | 195225833 |
| chr5 | 116989542 | 116990599 |
| chr10 | 123152504 | 123154790 |
| chr11 | 8454447 | 8455901 |
| chr12 | 15430656 | 15432265 |
| chr11 | 102951550 | 102953885 |
| chr16 | 71021313 | 71023319 |
| chr12 | 74397467 | 74399120 |
| chr20 | 30213105 | 30214492 |
| chr10 | 78541549 | 78544347 |
| chr1 | 82361869 | 82364565 |
| chr15_KI270905v1_ | 2696790 | 2700194 |
| chr6 | 87655647 | 87657422 |
| chr17 | 31230954 | 31231970 |
| chr6 | 101485287 | 101487140 |
| chr5 | 93800783 | 93802732 |
| chr8 | 106778254 | 106780155 |
| chr9 | 73686448 | 73687920 |
| chr14 | 61794744 | 61797080 |
| chr13 | 101514164 | 101515760 |
| chr2 | 199620551 | 199622643 |
| chr10 | 61727033 | 61728874 |
| chr2 | 64848609 | 64850235 |

| | | |
|---|---|---|
| chr7 | 53948092 | 53950682 |
| chr7 | 61019491 | 61021604 |
| chr7 | 96074915 | 96077942 |
| chr2 | 42040130 | 42042061 |
| chr3 | 93828119 | 93830549 |
| chr12 | 98742772 | 98744909 |
| chr2 | 58919148 | 58921036 |
| chr1 | 222137819 | 222139320 |
| chr4 | 86092565 | 86094712 |
| chr14 | 55598937 | 55600833 |
| chr2 | 98395723 | 98398377 |
| chr2 | 207552945 | 207554275 |
| chr3 | 173416427 | 173418802 |
| chr17 | 64419676 | 64421608 |
| chr10 | 55294137 | 55295926 |
| chr16 | 74382808 | 74384751 |
| chr1 | 218672683 | 218675209 |
| chr6 | 68836600 | 68838116 |
| chr4 | 47170309 | 47172835 |
| chr12 | 21317532 | 21319827 |
| chr20 | 62200659 | 62202121 |
| chr2 | 50639836 | 50642872 |
| chr9 | 66377696 | 66379443 |
| chr1 | 239559380 | 239560916 |
| chr20 | 54722621 | 54723886 |
| chr5 | 165656361 | 165657453 |
| chr6 | 75524253 | 75525469 |
| chr1 | 161161738 | 161164051 |
| chr6 | 74924662 | 74926035 |
| chr7 | 74877909 | 74879015 |
| chr7 | 79623314 | 79625047 |
| chr2 | 48929422 | 48931310 |
| chr5 | 110763652 | 110765163 |
| chr15 | 96999478 | 97001330 |
| chr6 | 93860547 | 93863301 |
| chr3 | 77014344 | 77015828 |
| chr11 | 31599931 | 31603002 |
| chr2 | 69456606 | 69458331 |
| chr8 | 76369785 | 76371762 |
| chr10 | 66670854 | 66673347 |
| chr5 | 121291064 | 121292243 |
| chr11 | 89986791 | 89989564 |
| chr2 | 183213998 | 183216296 |
| chr11 | 23150241 | 23152586 |
| chr10 | 63524874 | 63527487 |
| chr2 | 101005495 | 101008355 |
| chr13 | 71572090 | 71573610 |
| chr11 | 27366071 | 27367834 |
| chr1 | 178192513 | 178194652 |

| | | |
|---|---|---|
| chr2 | 37048619 | 37051179 |
| chr14 | 44752429 | 44754777 |
| chr14 | 36087881 | 36090159 |
| chr21 | 9763022 | 9764293 |
| chr14 | 33131342 | 33133189 |
| chr7 | 11211295 | 11212951 |
| chr6 | 56089125 | 56091668 |
| chr10 | 67082275 | 67087190 |
| chr15 | 54551193 | 54552680 |
| chr12 | 71307213 | 71310659 |
| chr6 | 91083415 | 91085467 |
| chr6 | 120592334 | 120594305 |
| chr15_KI270852v1_ | 127166 | 129745 |
| chr20 | 52002013 | 52005109 |
| chr10 | 52877519 | 52878681 |
| chr2 | 44758015 | 44760083 |
| chr9 | 131585854 | 131587816 |
| chr17 | 68128463 | 68131114 |
| chr1_KI270765v1_a | 163773 | 165997 |
| chr2 | 179174983 | 179178041 |
| chr1 | 102195121 | 102198717 |
| chr4 | 30714375 | 30716774 |
| chr7 | 96720709 | 96722763 |
| chr16 | 73148906 | 73150401 |
| chr14 | 102418992 | 102421521 |
| chr7 | 85472463 | 85475122 |
| chr1 | 70520651 | 70521696 |
| chr17 | 18246932 | 18248771 |
| chr2 | 208357526 | 208358995 |
| chr6 | 98427155 | 98429045 |
| chr6 | 72030003 | 72031902 |
| chr12 | 110467979 | 110469702 |
| chr14 | 32728460 | 32729916 |
| chr2 | 64320891 | 64322646 |
| chr15 | 62059207 | 62061142 |
| chr14 | 30077266 | 30079257 |
| chr10 | 72718984 | 72720156 |
| chr10 | 100152551 | 100155465 |
| chr9 | 66680216 | 66682495 |
| chr9 | 17332344 | 17334313 |
| chr12 | 86663620 | 86665641 |
| chr10 | 74310403 | 74311953 |
| chr1 | 108770422 | 108773309 |
| chr1 | 229617382 | 229618974 |
| chr2 | 158967241 | 158969801 |
| chr11 | 112465083 | 112466986 |
| chr6 | 900857 | 902250 |
| chr7 | 158233883 | 158235074 |
| chr20 | 4873176 | 4875204 |

| | | |
|---|---|---|
| chr2 | 82163151 | 82164316 |
| chr11 | 47425598 | 47427056 |
| chr6 | 78989492 | 78991890 |
| chr13 | 70590668 | 70593076 |
| chr4 | 115429263 | 115431088 |
| chr20 | 29619023 | 29620830 |
| chr2 | 209671153 | 209673392 |
| chr2 | 43761886 | 43764283 |
| chr7 | 117863866 | 117866326 |
| chr6 | 6261152 | 6262744 |
| chr2 | 48329140 | 48330755 |
| chr2 | 80371163 | 80372831 |
| chr3 | 147239260 | 147240526 |
| chr11 | 90002512 | 90004023 |
| chr10 | 86103473 | 86105537 |
| chr9 | 41180954 | 41184137 |
| chr7 | 83846507 | 83848158 |
| chr1 | 164557863 | 164559932 |
| chr11 | 95039885 | 95041803 |
| chr20 | 56383114 | 56385274 |
| chr9 | 65287243 | 65289327 |
| chr10 | 123043226 | 123045131 |
| chr7 | 93231310 | 93233370 |
| chr10 | 67379807 | 67381259 |
| chr14 | 87070833 | 87073356 |
| chr15 | 22297467 | 22301809 |
| chr1 | 104973912 | 104976796 |
| chr1 | 216825216 | 216826960 |
| chr14 | 41847054 | 41849766 |
| chr14 | 45531610 | 45533093 |
| chr1 | 81992052 | 81994225 |
| chr11 | 26064750 | 26066313 |
| chr6 | 38366779 | 38368240 |
| chr2 | 57176655 | 57179144 |
| chr3 | 133572234 | 133573794 |
| chr6 | 123805056 | 123807824 |
| chr2 | 43420180 | 43422408 |
| chr5 | 27780057 | 27783236 |
| chr4 | 32886827 | 32889467 |
| chr9 | 104091881 | 104094331 |
| chr6 | 93117259 | 93118780 |
| chr3 | 129038933 | 129041170 |
| chr14 | 77614880 | 77617718 |
| chr2 | 211044880 | 211047245 |
| chr11 | 80255267 | 80257697 |
| chr7 | 9156337 | 9158004 |
| chr7 | 104766579 | 104769064 |
| chr14 | 90290000 | 90292247 |
| chr6 | 123574447 | 123576407 |

| | | |
|---|---|---|
| chr7 | 64540303 | 64541859 |
| chr10 | 35363775 | 35365485 |
| chr7 | 82042908 | 82047547 |
| chr5 | 64567750 | 64570132 |
| chr14 | 41010545 | 41012812 |
| chr13 | 92954959 | 92957151 |
| chr8 | 47131798 | 47132967 |
| chr14 | 90906386 | 90907713 |
| chr10 | 45838588 | 45839848 |
| chr7 | 49253360 | 49255328 |
| chr15 | 55912336 | 55914965 |
| chr13 | 86068608 | 86070300 |
| chr7 | 83071181 | 83072699 |
| chr4 | 33032229 | 33033676 |
| chr2 | 39974898 | 39976623 |
| chr12 | 75256320 | 75259161 |
| chr3 | 81843899 | 81846650 |
| chr14 | 19454435 | 19456826 |
| chr20 | 30878353 | 30880171 |
| chr1 | 66652239 | 66655414 |
| chr6 | 101814909 | 101816772 |
| chr12 | 67270516 | 67273874 |
| chr7 | 89427251 | 89428936 |
| chr14 | 42537454 | 42541600 |
| chr2 | 231513960 | 231515791 |
| chr9 | 68258732 | 68263087 |
| chr2 | 65231549 | 65233365 |
| chr14 | 19400701 | 19405224 |
| chr5 | 67121816 | 67123605 |
| chr2 | 163189208 | 163193496 |
| chr6 | 381161 | 383284 |
| chr14 | 33780403 | 33782019 |
| chr10 | 55140865 | 55142309 |
| chr1 | 151249224 | 151251040 |
| chr3 | 57526700 | 57527702 |
| chr9 | 64789982 | 64791136 |
| chr12 | 40364073 | 40367418 |
| chr20 | 35708562 | 35711564 |
| chr2 | 62891056 | 62893718 |
| chr5 | 129487192 | 129488614 |
| chr11 | 22873935 | 22876940 |
| chr10 | 50085028 | 50086670 |
| chr9 | 13369093 | 13371066 |
| chr1 | 104521135 | 104522267 |
| chr2 | 66544353 | 66546886 |
| chr15 | 99387934 | 99389719 |
| chr16 | 80732228 | 80733702 |
| chr2 | 75304361 | 75305784 |
| chr2 | 95947804 | 95949040 |

| | | |
|---|---|---|
| chr19_KI270866v1_ | 40253 | 43135 |
| chr10 | 61452178 | 61454462 |
| chr20 | 53964658 | 53965974 |
| chr10 | 127756807 | 127758007 |
| chr2 | 41644325 | 41646803 |
| chr2 | 212710101 | 212712656 |
| chr8 | 67489433 | 67491453 |
| chr5 | 151252161 | 151254277 |
| chr7 | 84294642 | 84297305 |
| chr1 | 96718982 | 96720158 |
| chr6 | 149320067 | 149322500 |
| chr8 | 103502349 | 103505245 |
| chr12 | 68326230 | 68328167 |
| chr7 | 21255263 | 21256516 |
| chr7 | 78689451 | 78692049 |
| chr4 | 89111228 | 89113434 |
| chr1 | 97109123 | 97110477 |
| chr6 | 72275944 | 72277340 |
| chr12 | 9416606 | 9419058 |
| chr10 | 53865817 | 53867261 |
| chr10 | 114065332 | 114067749 |
| chr7 | 83954438 | 83956183 |
| chr14 | 52553881 | 52555650 |
| chr10 | 46749016 | 46751109 |
| chr10 | 65312953 | 65320890 |
| chr7 | 80988098 | 80989705 |
| chr9 | 4978838 | 4980525 |
| chr12 | 77010173 | 77011569 |
| chr11 | 58576739 | 58578886 |
| chr14 | 81218332 | 81219995 |
| chr9 | 31426438 | 31430325 |
| chr10 | 90729241 | 90731053 |
| chr11 | 96213236 | 96214492 |
| chr1 | 194528280 | 194529468 |
| chr11 | 89797264 | 89798913 |
| chr1 | 104712336 | 104716036 |
| chr7 | 90261155 | 90264921 |
| chr2 | 155678748 | 155680925 |
| chr10 | 125732453 | 125735456 |
| chr4 | 69421402 | 69423280 |
| chr13 | 64848720 | 64850569 |
| chr2 | 68831966 | 68834235 |
| chr1 | 28414912 | 28417428 |
| chr2 | 91788446 | 91789456 |
| chr2 | 185761482 | 185763937 |
| chr20 | 31738418 | 31740112 |
| chr1 | 90717893 | 90721546 |
| chr7 | 84477689 | 84479601 |
| chr7 | 92821262 | 92822857 |

| | | |
|---|---|---|
| chr15 | 65577511 | 65579486 |
| chr2 | 164726702 | 164728662 |
| chr14 | 48633489 | 48635364 |
| chr1 | 90463387 | 90464983 |
| chr3 | 116615398 | 116616801 |
| chr14 | 85749840 | 85751717 |
| chr20 | 28607095 | 28608957 |
| chr2 | 174247094 | 174248802 |
| chr14 | 73106175 | 73108323 |
| chr1 | 85894242 | 85896677 |
| chr3 | 176429425 | 176432412 |
| chr6 | 25990732 | 25992783 |
| chr9 | 82487879 | 82489916 |
| chr2 | 157668377 | 157670788 |
| chr2 | 219218553 | 219220157 |
| chr7 | 90331115 | 90333238 |
| chr2 | 36999519 | 37000528 |
| chr6 | 98201294 | 98203080 |
| chr8 | 39555605 | 39556831 |
| chr14 | 60983165 | 60985141 |
| chr1 | 235327421 | 235329815 |
| chr6 | 121442833 | 121444887 |
| chr1 | 98049824 | 98053345 |
| chr10 | 87087186 | 87088980 |
| chr16 | 33796955 | 33798907 |
| chr20 | 24967604 | 24970492 |
| chr6 | 99093642 | 99095005 |
| chr14 | 26979778 | 26981911 |
| chr5 | 166990399 | 166992113 |
| chr12 | 74180398 | 74182269 |
| chr11 | 99945881 | 99948489 |
| chr1 | 148651180 | 148653534 |
| chr10 | 65142329 | 65145350 |
| chr14 | 99497244 | 99500091 |
| chr3 | 144272181 | 144273996 |
| chr14 | 20325735 | 20327893 |
| chr10 | 87411440 | 87413106 |
| chr14 | 52062286 | 52063906 |
| chr10 | 21524173 | 21526376 |
| chr2 | 181890076 | 181892291 |
| chr15 | 30079751 | 30080967 |
| chr1 | 102676667 | 102678720 |
| chr2 | 43691849 | 43693277 |
| chr22 | 11939569 | 11940568 |
| chr22 | 15548809 | 15552411 |
| chr19 | 27931955 | 27933744 |
| chr15_KI270850v1_ | 96610 | 98507 |
| chr10 | 123363132 | 123364269 |
| chr10 | 59321254 | 59323248 |

| | | |
|---|---|---|
| chr1 | 102905157 | 102906383 |
| chr10 | 111318259 | 111320311 |
| chr5 | 119115553 | 119116992 |
| chr14 | 30726738 | 30728864 |
| chr14 | 46901361 | 46903245 |
| chr7 | 18834311 | 18836282 |
| chr14 | 40055740 | 40057584 |
| chr12 | 26419944 | 26422392 |
| chr7 | 61578637 | 61580552 |
| chr2 | 91845729 | 91847146 |
| chr6 | 56963152 | 56964205 |
| chr10 | 131971989 | 131974811 |
| chr2 | 54458080 | 54460423 |
| chr13 | 86878617 | 86880407 |
| chr16 | 33841086 | 33845200 |
| chr6 | 75933852 | 75936108 |
| chr5 | 27143409 | 27144856 |
| chr7 | 105093597 | 105096276 |
| chr12 | 122398891 | 122401215 |
| chr7 | 52187643 | 52189473 |
| chr1 | 117405520 | 117407524 |
| chr12 | 63958364 | 63959843 |
| chr10 | 90399685 | 90402501 |
| chr3 | 111099837 | 111100840 |
| chr7 | 82825651 | 82829560 |
| chr10 | 63256187 | 63257842 |
| chr13 | 80132378 | 80133982 |
| chr2 | 87782776 | 87784677 |
| chr4 | 131645116 | 131646657 |
| chr2 | 157442371 | 157444478 |
| chr1 | 99110944 | 99112246 |
| chr8 | 111345020 | 111348007 |
| chr2 | 193614841 | 193616631 |
| chr15 | 38128826 | 38131972 |
| chr8 | 128883471 | 128885055 |
| chr10 | 59574383 | 59577105 |
| chr14 | 96880382 | 96882458 |
| chr5 | 41012616 | 41013793 |
| chr7 | 90986851 | 90987883 |
| chr4 | 93278822 | 93280583 |
| chr17 | 8225563 | 8227978 |
| chr1 | 66990160 | 66991218 |
| chr16 | 76024058 | 76025490 |
| chrX | 108090756 | 108092803 |
| chr14 | 50860507 | 50861687 |
| chr1 | 144894251 | 144896996 |
| chr5 | 125115640 | 125118361 |
| chr5 | 164059293 | 164060711 |
| chr1 | 104742056 | 104744959 |

| | | |
|---|---|---|
| chr2 | 88626526 | 88628659 |
| chr6 | 94076187 | 94078679 |
| chr6 | 122028218 | 122031131 |
| chr10 | 16891598 | 16892804 |
| chr7 | 123747673 | 123750525 |
| chr1 | 192807942 | 192810808 |
| chr16 | 80704959 | 80707234 |
| chr10 | 85041786 | 85043147 |
| chr10 | 66966226 | 66968749 |
| chr17 | 42608634 | 42610078 |
| chr12 | 65112116 | 65114516 |
| chr7 | 85223524 | 85227263 |
| chr2 | 76534566 | 76536027 |
| chr14 | 47749712 | 47751457 |
| chr8 | 16996912 | 16999737 |
| chr1 | 83745213 | 83746967 |
| chr9 | 66147632 | 66149969 |
| chr7 | 9847765 | 9850102 |
| chr13 | 61883200 | 61884485 |
| chr1 | 90081810 | 90083462 |
| chr10 | 87669634 | 87672522 |
| chr21 | 9950328 | 9952208 |
| chr15 | 37642287 | 37643658 |
| chr1 | 81953951 | 81955399 |
| chr1 | 184279328 | 184280852 |
| chr7 | 113850408 | 113852387 |
| chr10 | 88801911 | 88803509 |
| chr14 | 32676220 | 32677744 |
| chr9 | 4338083 | 4340719 |
| chr9 | 66671925 | 66673926 |
| chr10 | 53838175 | 53839999 |
| chr7 | 85481906 | 85484821 |
| chr6 | 45547773 | 45549167 |
| chr2 | 186205492 | 186207040 |
| chr2 | 167298614 | 167299872 |
| chr2 | 242163260 | 242164654 |
| chr14 | 85576247 | 85578448 |
| chr11 | 106846581 | 106848965 |
| chr7 | 86417695 | 86419755 |
| chr12 | 70915305 | 70918654 |
| chr1 | 87450395 | 87453179 |
| chr2 | 77873970 | 77877391 |
| chr14 | 38604775 | 38606894 |
| chr9 | 103943754 | 103945690 |
| chr14 | 43235251 | 43237924 |
| chr8 | 89318144 | 89320442 |
| chr7 | 101450254 | 101452140 |
| chr12 | 55737342 | 55739084 |
| chr10 | 106387768 | 106389075 |

| | | |
|---|---|---|
| chr15 | 56920899 | 56922897 |
| chr10 | 121181545 | 121183672 |
| chr14 | 48626144 | 48628345 |
| chr3 | 73064967 | 73066636 |
| chr14 | 59226156 | 59228743 |
| chr12 | 39441308 | 39444236 |
| chr12 | 44057480 | 44060324 |
| chr15 | 93048394 | 93051169 |
| chr5 | 26345367 | 26347391 |
| chr16 | 55969074 | 55970950 |
| chr22 | 15705008 | 15706610 |
| chr11 | 127271102 | 127272356 |
| chr7 | 65253401 | 65256017 |
| chr7 | 79838607 | 79840662 |
| chr2 | 15046688 | 15047977 |
| chr4 | 126677789 | 126679354 |
| chr9 | 14610285 | 14612214 |
| chr2 | 195111242 | 195112707 |
| chr17 | 41687647 | 41689104 |
| chr7 | 147351549 | 147352769 |
| chr10 | 115725644 | 115729473 |
| chr12 | 120249363 | 120251097 |
| chr9 | 72982479 | 72985556 |
| chr16 | 46415569 | 46417974 |
| chr10 | 63863540 | 63864568 |
| chr7 | 3694425 | 3697364 |
| chr10 | 35038160 | 35039513 |
| chr15 | 44496332 | 44497349 |
| chr8 | 110828769 | 110829993 |
| chr14 | 52852636 | 52853757 |
| chr7 | 89614976 | 89617184 |
| chr14 | 38749153 | 38751154 |
| chr15 | 21290884 | 21292905 |
| chr14 | 26501002 | 26503644 |
| chr5_KI270897v1_a | 87782 | 89789 |
| chr1 | 86653881 | 86655838 |
| chr5 | 123750100 | 123752304 |
| chr13 | 92746875 | 92748653 |
| chr2 | 184641295 | 184645930 |
| chr2 | 185608314 | 185609923 |
| chr2 | 188590412 | 188593124 |
| chr7 | 105706355 | 105708183 |
| chr7 | 82559778 | 82562012 |
| chr10 | 79091482 | 79093483 |
| chr21 | 16500660 | 16503293 |
| chr3 | 152075399 | 152077019 |
| chr2 | 199667858 | 199669567 |
| chr21 | 10392175 | 10394238 |
| chr12 | 65322903 | 65325862 |

| | | |
|---|---|---|
| chr16 | 65080562 | 65082754 |
| chr11 | 56644787 | 56646944 |
| chr2 | 231462923 | 231464883 |
| chrUn_GL000218v1 | 35805 | 37647 |
| chr7 | 84784637 | 84786410 |
| chr2 | 72007207 | 72009004 |
| chr11 | 35415299 | 35418305 |
| chr9 | 97203286 | 97204304 |
| chr7 | 47443011 | 47445013 |
| chr2 | 35260128 | 35261775 |
| chr14 | 31673062 | 31675016 |
| chr20 | 18506819 | 18508198 |
| chr21 | 23469688 | 23471638 |
| chr14 | 52693887 | 52695452 |
| chr1 | 190734842 | 190736027 |
| chr2 | 223146729 | 223148409 |
| chr10 | 131360325 | 131362292 |
| chr6 | 102506530 | 102508140 |
| chr14 | 48327782 | 48329471 |
| chr2 | 178504442 | 178507191 |
| chr16 | 33110656 | 33112445 |
| chr6 | 18154426 | 18157711 |
| chr6 | 57504710 | 57507357 |
| chr1 | 994539 | 996033 |
| chr11 | 26348556 | 26350703 |
| chr1 | 186310605 | 186312898 |
| chr6 | 135052559 | 135054647 |
| chr4 | 65038921 | 65040724 |
| chr1 | 247990921 | 247992303 |
| chr2 | 27568955 | 27571569 |
| chr10 | 118442444 | 118444918 |
| chr6 | 80247642 | 80250360 |
| chr5 | 52732684 | 52734405 |
| chr2 | 188760451 | 188762855 |
| chr6 | 152466403 | 152468007 |
| chr14 | 23390703 | 23392761 |
| chr3 | 152063076 | 152065828 |
| chr11 | 45917042 | 45919396 |
| chr21 | 20597212 | 20598910 |
| chr14 | 41038688 | 41039946 |
| chr1 | 195606215 | 195607868 |
| chr8 | 68072493 | 68073957 |
| chr14 | 33005545 | 33008057 |
| chr9 | 134191650 | 134192651 |
| chr16_KI270728v1_ | 1178381 | 1181231 |
| chr22 | 42613843 | 42616065 |
| chrY | 10974642 | 10976596 |
| chr2 | 104341300 | 104343259 |
| chr3 | 109926970 | 109928422 |

| | | |
|---|---|---|
| chr20 | 35975432 | 35977171 |
| chr10 | 115685360 | 115687171 |
| chr6 | 72875250 | 72877397 |
| chr18 | 44465584 | 44468760 |
| chr13 | 63189739 | 63191250 |
| chr1 | 44720934 | 44722645 |
| chr8 | 85579544 | 85580713 |
| chr11 | 16354466 | 16356466 |
| chr4 | 90143725 | 90145932 |
| chr10 | 112139413 | 112141899 |
| chr1 | 101405950 | 101408026 |
| chr2 | 27439440 | 27441569 |
| chr3 | 83381346 | 83382616 |
| chr10 | 62035753 | 62038112 |
| chr2 | 186830229 | 186831708 |
| chr13 | 18421246 | 18423243 |
| chr12 | 84425962 | 84427734 |
| chr1 | 103365458 | 103366815 |
| chr4 | 65005901 | 65007946 |
| chr1 | 148639985 | 148641815 |
| chr6 | 151988582 | 151990109 |
| chr7 | 89029631 | 89030990 |
| chr2 | 193554031 | 193557761 |
| chr14 | 97762619 | 97764628 |
| chr12 | 58851345 | 58852346 |
| chr12 | 89608181 | 89610501 |
| chr10 | 115164876 | 115166084 |
| chr12 | 99649229 | 99651449 |
| chr14 | 50466593 | 50468309 |
| chr2 | 52626421 | 52627860 |
| chr8 | 7762734 | 7765174 |
| chr20 | 5585322 | 5588215 |
| chr10 | 88300620 | 88302789 |
| chr7 | 53452412 | 53454407 |
| chr2 | 50702264 | 50704123 |
| chr6 | 132821861 | 132824518 |
| chr12 | 80265909 | 80268426 |
| chr8 | 116336202 | 116339634 |
| chr4 | 82428640 | 82430314 |
| chr10 | 51468710 | 51471421 |
| chr13 | 70220939 | 70223115 |
| chr1 | 87073036 | 87074311 |
| chr3 | 130097616 | 130098759 |
| chr13 | 87693355 | 87695657 |
| chr2 | 178742753 | 178744844 |
| chr14 | 41610812 | 41612254 |
| chr1 | 116380870 | 116383117 |
| chr1 | 22094327 | 22095732 |
| chr7 | 8435108 | 8436648 |

| | | |
|---|---|---|
| chr2 | 159368530 | 159369780 |
| chr10 | 64817923 | 64820091 |
| chr10 | 117336728 | 117339017 |
| chr14 | 59359716 | 59360993 |
| chr10 | 53868946 | 53871306 |
| chr2 | 169560923 | 169562663 |
| chr1 | 196296433 | 196298583 |
| chr4 | 46918876 | 46921040 |
| chr4 | 34101524 | 34103779 |
| chr7 | 82923865 | 82926790 |
| chr2 | 72770300 | 72771616 |
| chrUn_GL000195v1 | 106826 | 108398 |
| chr3 | 195503683 | 195504769 |
| chr4 | 41946901 | 41949616 |
| chr4 | 45337412 | 45339791 |
| chr12 | 99424904 | 99426993 |
| chr1 | 93905480 | 93906948 |
| chr11 | 3378542 | 3380569 |
| chr3 | 80628153 | 80629407 |
| chr20 | 8751354 | 8752606 |
| chr7 | 49583308 | 49586867 |
| chr7 | 84800955 | 84802394 |
| chr10 | 57812042 | 57813944 |
| chr10 | 62399029 | 62400854 |
| chr12 | 69286244 | 69287487 |
| chr1 | 112713869 | 112716061 |
| chr20 | 8964573 | 8966526 |
| chr7 | 82204208 | 82206921 |
| chr1 | 153139587 | 153141380 |
| chr14 | 39627931 | 39629508 |
| chr10 | 121741511 | 121743149 |
| chr20 | 11045035 | 11047827 |
| chr12 | 69584237 | 69586568 |
| chr21 | 8394195 | 8396081 |
| chr2 | 67604201 | 67605769 |
| chr4 | 15477437 | 15479560 |
| chr5 | 59761897 | 59763318 |
| chr8 | 104901172 | 104902305 |
| chrY | 11103278 | 11104308 |
| chr13 | 67204139 | 67206194 |
| chr12 | 24747094 | 24749472 |
| chr7 | 78706155 | 78708921 |
| chr1 | 98387910 | 98389330 |
| chr12 | 77962668 | 77965467 |
| chr1 | 192306543 | 192308425 |
| chr2 | 41304795 | 41307342 |
| chr11 | 110969222 | 110971187 |
| chr21 | 10602866 | 10606009 |
| chr10 | 61689953 | 61691652 |

| | | |
|---|---|---|
| chrX | 148889992 | 148892452 |
| chr2 | 187435537 | 187437990 |
| chr8 | 50315369 | 50316861 |
| chr10 | 56414478 | 56417027 |
| chr1 | 104432115 | 104433573 |
| chr7 | 8509591 | 8510725 |
| chr10 | 54459608 | 54462562 |
| chr9 | 64495293 | 64497280 |
| chr11 | 21902261 | 21904275 |
| chr14 | 96560848 | 96562041 |
| chr7 | 132450336 | 132451583 |
| chr15_KI270905v1_ | 816452 | 817938 |
| chr8 | 26449865 | 26453276 |
| chr3 | 185819352 | 185821739 |
| chr2 | 39784226 | 39787609 |
| chr7 | 81292018 | 81293928 |
| chr1 | 94850530 | 94852494 |
| chr3 | 35803382 | 35804777 |
| chr12 | 43064081 | 43066104 |
| chr12 | 75542991 | 75544284 |
| chr21 | 5781249 | 5783493 |
| chr4 | 124261419 | 124263078 |
| chr1 | 36284295 | 36287120 |
| chr20 | 8091498 | 8093699 |
| chr2 | 40121249 | 40124045 |
| chr12 | 82963257 | 82964800 |
| chr2 | 57311638 | 57313935 |
| chr8 | 112651853 | 112655933 |
| chr6 | 64505265 | 64506969 |
| chr2 | 225269862 | 225271376 |
| chr1 | 218304015 | 218305541 |
| chr14 | 24658132 | 24659753 |
| chr14 | 39664306 | 39666521 |
| chr9 | 42489339 | 42492722 |
| chr3 | 95722840 | 95726109 |
| chr5 | 173857736 | 173859177 |
| chr19 | 38336221 | 38338067 |
| chr6 | 28335666 | 28337309 |
| chr1 | 52280026 | 52281438 |
| chr1 | 218930134 | 218932274 |
| chr2 | 97116436 | 97119150 |
| chr12 | 47700793 | 47702019 |
| chr15 | 58184346 | 58186199 |
| chr1 | 94184326 | 94187041 |
| chr1 | 103980207 | 103982530 |
| chr12 | 77874554 | 77877575 |
| chr8 | 40102538 | 40104530 |
| chr20 | 26206536 | 26208899 |
| chr1 | 199132835 | 199135076 |

| | | |
|---|---|---|
| chr6 | 16717313 | 16720040 |
| chr16 | 12455995 | 12457038 |
| chr7 | 10938519 | 10940645 |
| chr4 | 107027794 | 107030186 |
| chr12 | 71531945 | 71533334 |
| chr11 | 1893606 | 1894691 |
| chr2 | 192080796 | 192082048 |
| chr7 | 78490889 | 78492935 |
| chr16 | 66753247 | 66754450 |
| chr7 | 89510268 | 89512702 |
| chr6 | 1436307 | 1437755 |
| chr14 | 19659747 | 19660829 |
| chr5 | 78928609 | 78929746 |
| chr8 | 91039190 | 91040836 |
| chr4 | 11398661 | 11400116 |
| chr2 | 66111807 | 66113303 |
| chr13 | 70283115 | 70286055 |
| chr13 | 71691761 | 71693508 |
| chr15 | 21270410 | 21273419 |
| chr15 | 38196014 | 38197697 |
| chr2 | 207602389 | 207603974 |
| chr14 | 19365865 | 19367917 |
| chr2 | 219734469 | 219737714 |
| chr12 | 25240533 | 25242730 |
| chr3 | 181299457 | 181300824 |
| chr6 | 117320735 | 117323132 |
| chr12 | 89948435 | 89950136 |
| chr4 | 96242421 | 96245080 |
| chr1 | 40690256 | 40692362 |
| chr10 | 118754677 | 118755888 |
| chr10 | 52445244 | 52446409 |
| chr1 | 101762409 | 101765422 |
| chr10 | 103060537 | 103062501 |
| chr3 | 77533977 | 77535468 |
| chr2 | 40320528 | 40325676 |
| chr18 | 32983055 | 32984833 |
| chr7 | 58081166 | 58085868 |
| chr14 | 19475259 | 19476656 |
| chr17 | 22055261 | 22056789 |
| chr2 | 675501 | 678827 |
| chr1 | 111388735 | 111390267 |
| chr2 | 129996666 | 129997693 |
| chr6 | 162011120 | 162012543 |
| chr2 | 165358969 | 165363140 |
| chr17 | 75782121 | 75785727 |
| chr10 | 78932203 | 78934189 |
| chr6 | 95350819 | 95353017 |
| chr2 | 159137572 | 159139322 |
| chr5 | 118357071 | 118359696 |

| | | |
|---|---|---|
| chr4 | 42776060 | 42777901 |
| chr2 | 8823233 | 8824771 |
| chr14 | 77334711 | 77338326 |
| chr7 | 80492702 | 80493915 |
| chr14 | 86251711 | 86253921 |
| chr7 | 81462044 | 81463939 |
| chr10 | 4277128 | 4278833 |
| chr10 | 23439895 | 23441630 |
| chr6 | 55519269 | 55520855 |
| chr10 | 64254266 | 64256324 |
| chr15 | 90679385 | 90680889 |
| chr1 | 89102071 | 89104153 |
| chr2 | 51760519 | 51762353 |
| chr3 | 29431739 | 29433919 |
| chr20 | 22055085 | 22057467 |
| chr6 | 160167678 | 160170009 |
| chr18 | 71703551 | 71706607 |
| chr2 | 211827259 | 211829052 |
| chr10 | 66504254 | 66505799 |
| chr14 | 44164033 | 44166039 |
| chr1 | 195076472 | 195078331 |
| chr11 | 101263656 | 101265114 |
| chr14 | 83469357 | 83471937 |
| chr14 | 28358434 | 28360824 |
| chr4 | 108034352 | 108036232 |
| chr10 | 4764992 | 4772710 |
| chr10 | 57131507 | 57133370 |
| chr7 | 95819347 | 95822591 |
| chr15 | 84586819 | 84588322 |
| chr14 | 88709420 | 88711622 |
| chr12 | 74666949 | 74669922 |
| chr2 | 81690187 | 81692202 |
| chr1 | 94467567 | 94468988 |
| chr1 | 64649572 | 64651146 |
| chr6 | 55970191 | 55972866 |
| chr7 | 81694962 | 81696714 |
| chr10 | 69178215 | 69180416 |
| chr17 | 56110903 | 56112338 |
| chr6 | 54144485 | 54146423 |
| chr1 | 218635676 | 218638215 |
| chr5 | 88140782 | 88142825 |
| chr10 | 84588255 | 84590315 |
| chr11 | 25478961 | 25480400 |
| chr10 | 92933884 | 92935910 |
| chr6 | 133754932 | 133756726 |
| chr2 | 105082034 | 105083943 |
| chr2 | 88739615 | 88741469 |
| chr21 | 7585354 | 7588002 |
| chr8 | 114699401 | 114701566 |

| | | |
|---|---|---|
| chr2 | 84714557 | 84717182 |
| chr14 | 27725413 | 27727055 |
| chr10 | 57620742 | 57623248 |
| chr1 | 170038706 | 170040870 |
| chr2 | 94979913 | 94981929 |
| chr1 | 189942583 | 189944470 |
| chr14 | 30264593 | 30267073 |
| chr1 | 169347072 | 169349945 |
| chr8 | 92964035 | 92967993 |
| chr1 | 244652082 | 244653858 |
| chr14 | 78263721 | 78266628 |
| chr21 | 28879422 | 28880693 |
| chr8 | 52080077 | 52082255 |
| chr21 | 10027993 | 10030083 |
| chr12 | 6613098 | 6615067 |
| chr2 | 97612585 | 97613964 |
| chr1 | 195831569 | 195832727 |
| chr7 | 83901290 | 83904493 |
| chr10 | 64444423 | 64445823 |
| chr2 | 87325292 | 87327267 |
| chr1 | 196653409 | 196655242 |
| chr1 | 196261338 | 196262817 |
| chr2 | 72516508 | 72518752 |
| chr1 | 238640075 | 238641444 |
| chr17 | 21581913 | 21583527 |
| chr2 | 193761782 | 193764263 |
| chr10 | 37793391 | 37794562 |
| chr6 | 74312940 | 74314421 |
| chr10 | 66111203 | 66112837 |
| chr5 | 44939784 | 44943749 |
| chr14 | 41625128 | 41627525 |
| chr8 | 79057533 | 79059403 |
| chr20 | 53222038 | 53223707 |
| chr6 | 54149518 | 54151948 |
| chr14 | 64546687 | 64548630 |
| chr2 | 34740525 | 34742696 |
| chr11 | 40393836 | 40395569 |
| chr7 | 81506848 | 81509574 |
| chr10 | 89767815 | 89770385 |
| chr1 | 47558205 | 47559976 |
| chr2 | 169071880 | 169074102 |
| chr2 | 102958603 | 102960239 |
| chr10 | 79676439 | 79679348 |
| chr14 | 50722897 | 50724221 |
| chr11 | 96305637 | 96308903 |
| chrX | 106309267 | 106310297 |
| chr2 | 195574982 | 195576233 |
| chr9 | 65933525 | 65937396 |
| chr4 | 29567638 | 29569423 |

| | | |
|---|---|---|
| chr10 | 66891983 | 66894495 |
| chr10 | 114855064 | 114856438 |
| chr11 | 31431414 | 31432817 |
| chr14 | 47193841 | 47195950 |
| chr4 | 107282385 | 107284380 |
| chr11 | 62994557 | 62996534 |
| chr20 | 8043469 | 8045313 |
| chr8 | 41872444 | 41874374 |
| chr14 | 49124546 | 49125783 |
| chr8 | 116083169 | 116085103 |
| chr1 | 116795584 | 116797611 |
| chr13 | 71645143 | 71647800 |
| chr7 | 81657049 | 81658897 |
| chr1 | 148495453 | 148497441 |
| chr9 | 13273564 | 13276709 |
| chr13 | 62702525 | 62705408 |
| chr2 | 185277936 | 185281286 |
| chr21 | 9898076 | 9901107 |
| chr14 | 18561373 | 18564264 |
| chr10 | 55863199 | 55864424 |
| chr2 | 58041731 | 58043395 |
| chr1 | 73649113 | 73651166 |
| chr5 | 52597896 | 52600255 |
| chr11 | 109677954 | 109679551 |
| chr14 | 105255780 | 105257792 |
| chr7 | 79451317 | 79452970 |
| chr21 | 10034345 | 10036289 |
| chr4 | 27923620 | 27925065 |
| chr6 | 71911192 | 71913813 |
| chr8 | 104880034 | 104882114 |
| chr2 | 46156212 | 46158209 |
| chr2 | 172990494 | 172992402 |
| chr4 | 47182090 | 47184526 |
| chr13 | 37764822 | 37767451 |
| chr11 | 103693939 | 103695575 |
| chr2 | 39297344 | 39300449 |
| chr12 | 131371702 | 131373505 |
| chr12 | 71663490 | 71665460 |
| chr8 | 85649071 | 85650410 |
| chr9 | 63611309 | 63614480 |
| chr14 | 41049692 | 41052085 |
| chr6 | 69961412 | 69963407 |
| chr7 | 85950924 | 85953393 |
| chr1 | 148662770 | 148665029 |
| chr11 | 33890733 | 33892719 |
| chr2 | 39721942 | 39723478 |
| chr6 | 77326943 | 77329096 |
| chr19 | 21194391 | 21196000 |
| chr8 | 7765565 | 7772221 |

| | | |
|---|---|---|
| chr1 | 107781137 | 107783683 |
| chr2 | 184108434 | 184110377 |
| chr6 | 132827249 | 132828468 |
| chr1 | 107394470 | 107397562 |
| chr3 | 113952783 | 113954291 |
| chr2 | 217572054 | 217573931 |
| chr11 | 96745082 | 96746825 |
| chr8 | 17048681 | 17049682 |
| chr11 | 36165729 | 36166967 |
| chr2 | 66172752 | 66174458 |
| chr19 | 18432902 | 18434851 |
| chr15 | 56241725 | 56243714 |
| chr1 | 198931183 | 198938657 |
| chr4 | 37073137 | 37075329 |
| chr1 | 213554624 | 213556985 |
| chr22 | 16267307 | 16268505 |
| chr10 | 84145960 | 84148635 |
| chr1 | 118502968 | 118504930 |
| chr8 | 79614353 | 79616516 |
| chr21 | 22576638 | 22577956 |
| chr22 | 12618666 | 12621370 |
| chr7 | 78997212 | 79001037 |
| chr6 | 74983772 | 74985538 |
| chr10 | 55762702 | 55764668 |
| chr11 | 43828621 | 43830481 |
| chr2 | 38601519 | 38604706 |
| chr8 | 103594949 | 103596596 |
| chr15 | 44426767 | 44429081 |
| chr4 | 174826285 | 174828373 |
| chr10 | 35725974 | 35727866 |
| chr16 | 86587144 | 86588791 |
| chr7 | 99713201 | 99714813 |
| chr2 | 154771093 | 154773292 |
| chr5 | 77557753 | 77561358 |
| chr10 | 54606420 | 54608250 |
| chr12 | 62089326 | 62092129 |
| chr14 | 68167038 | 68170899 |
| chr6 | 84404606 | 84406007 |
| chr6 | 55515780 | 55517600 |
| chr2 | 171005315 | 171007598 |
| chr21 | 13836426 | 13837465 |
| chr1 | 147845902 | 147847186 |
| chr6 | 79820642 | 79822359 |
| chr1 | 218808990 | 218811416 |
| chr4 | 33098150 | 33100761 |
| chr2 | 242161428 | 242162744 |
| chr2 | 195015795 | 195017698 |
| chr7 | 82888338 | 82891428 |
| chr1 | 220293844 | 220295664 |

| | | |
|---|---|---|
| chr13 | 54314309 | 54315526 |
| chr14 | 59843228 | 59844975 |
| chr16 | 46443171 | 46444567 |
| chr17 | 60743170 | 60744768 |
| chr11 | 24246624 | 24248621 |
| chrUn_GL000219v1 | 98419 | 100081 |
| chr1 | 248873012 | 248875320 |
| chr5_KI270897v1_a | 709565 | 710996 |
| chr14 | 105300910 | 105302277 |
| chr2 | 141030218 | 141031564 |
| chr2 | 94292278 | 94293295 |
| chr10 | 73413464 | 73415083 |
| chr14 | 88870813 | 88872146 |
| chr2 | 56922265 | 56925229 |
| chr20 | 23416453 | 23418277 |
| chr2 | 176078965 | 176081248 |
| chr2 | 57800204 | 57804313 |
| chr6 | 72852631 | 72854924 |
| chr1 | 214970931 | 214973206 |
| chr1 | 182670062 | 182671984 |
| chr10 | 51794645 | 51795974 |
| chr5 | 85215234 | 85217159 |
| chr20 | 55627881 | 55629058 |
| chr9 | 63764054 | 63765484 |
| chr10 | 73679527 | 73680788 |
| chr3 | 158821878 | 158824341 |
| chr6 | 26095411 | 26097333 |
| chr5 | 81300592 | 81302094 |
| chr10 | 48438259 | 48440881 |
| chr1 | 189692815 | 189694405 |
| chr10 | 87861915 | 87863798 |
| chr12 | 25187617 | 25189629 |
| chr15 | 95263905 | 95265312 |
| chr2 | 210918812 | 210920606 |
| chr2 | 84792161 | 84793949 |
| chr1 | 216075786 | 216080178 |
| chr1 | 217846039 | 217848113 |
| chr14 | 40892969 | 40894348 |
| chr15 | 72300979 | 72302911 |
| chr5 | 73217642 | 73219411 |
| chr5 | 52133901 | 52135063 |
| chr6 | 81747441 | 81749509 |
| chr20 | 53736668 | 53739122 |
| chr6 | 57188418 | 57190090 |
| chr12 | 56102638 | 56104801 |
| chr14 | 45286516 | 45288209 |
| chr1 | 104710040 | 104711432 |
| chr1 | 90505805 | 90507279 |
| chr2 | 103265018 | 103266282 |

| | | |
|---|---|---|
| chr6 | 75699872 | 75701888 |
| chr12 | 61818180 | 61820758 |
| chr2 | 219571673 | 219573602 |
| chr6 | 122059498 | 122061068 |
| chr10 | 65218908 | 65221734 |
| chr2 | 148495276 | 148496995 |
| chr6 | 73669126 | 73670866 |
| chr1 | 197391595 | 197394323 |
| chr1 | 193202228 | 193203526 |
| chr1 | 145063666 | 145065773 |
| chr6 | 53981560 | 53983427 |
| chr6 | 77141956 | 77145846 |
| chr17 | 75670282 | 75671896 |
| chr9 | 39552486 | 39553654 |
| chr7 | 81497650 | 81500193 |
| chr10 | 59808706 | 59810711 |
| chr4 | 32253484 | 32256272 |
| chr20 | 4016849 | 4019106 |
| chr16 | 77721096 | 77723199 |
| chr10 | 35582501 | 35585264 |
| chr10 | 84468914 | 84471674 |
| chr12 | 19519827 | 19521137 |
| chr6 | 78160388 | 78162510 |
| chr7 | 83542240 | 83545687 |
| chr6 | 94619979 | 94622246 |
| chr14 | 52939299 | 52941168 |
| chr7 | 85032671 | 85034069 |
| chr14 | 26572394 | 26573996 |
| chrY | 11196596 | 11197970 |
| chr15 | 96932573 | 96933915 |
| chr14 | 32937258 | 32938679 |
| chr1 | 245850329 | 245853604 |
| chr11 | 77994436 | 77995920 |
| chr14 | 37874193 | 37875815 |
| chr1 | 50708917 | 50710668 |
| chr6 | 8940532 | 8942339 |
| chr7 | 56063173 | 56064823 |
| chr16 | 80592679 | 80593935 |
| chr2 | 82601599 | 82603290 |
| chr12 | 19255045 | 19256469 |
| chr8 | 99037066 | 99039651 |
| chr15 | 35117166 | 35119247 |
| chr7 | 78992219 | 78993354 |
| chr14 | 79948277 | 79951576 |
| chr10 | 105514451 | 105516515 |
| chr6 | 133612666 | 133614398 |
| chr2 | 13396351 | 13397602 |
| chr8 | 140990474 | 140991752 |
| chr11 | 18934825 | 18937378 |

| | | |
|---|---|---|
| chr12 | 77547862 | 77553874 |
| chr1 | 195664915 | 195666033 |
| chr7 | 10604259 | 10605920 |
| chr2 | 216856903 | 216858831 |
| chr4 | 157164708 | 157166639 |
| chr14 | 60651133 | 60653110 |
| chr14 | 28574975 | 28576320 |
| chr7 | 93423475 | 93426752 |
| chr1 | 103670205 | 103671641 |
| chr2 | 172555482 | 172556498 |
| chr6 | 19291429 | 19293313 |
| chr2 | 51272533 | 51273764 |
| chr10 | 79899757 | 79901793 |
| chr1 | 143249229 | 143250766 |
| chr7 | 79904386 | 79906644 |
| chr2 | 228565837 | 228567384 |
| chr10 | 124622179 | 124625336 |
| chr3 | 60111510 | 60114037 |
| chr6 | 95848974 | 95850972 |
| chr19 | 6839092 | 6840095 |
| chr11 | 96688343 | 96689497 |
| chr2 | 157438614 | 157440461 |
| chr14 | 44056728 | 44058379 |
| chr14 | 47856488 | 47858465 |
| chrUn_GL000195v1 | 139731 | 141846 |
| chr5 | 27408143 | 27411639 |
| chr14 | 41274071 | 41275605 |
| chr3 | 36599960 | 36601676 |
| chr16 | 76651770 | 76653290 |
| chr7 | 81511708 | 81513075 |
| chr16 | 70172943 | 70175074 |
| chr13 | 69673114 | 69675102 |
| chrY | 10983658 | 10985318 |
| chr1 | 104354687 | 104356954 |
| chr2 | 188968776 | 188970938 |
| chr1 | 103169634 | 103173775 |
| chr20 | 17568851 | 17571538 |
| chr1 | 71902635 | 71905234 |
| chr11 | 32838279 | 32839443 |
| chr11 | 90039553 | 90041141 |
| chr15 | 36891643 | 36894940 |
| chr2 | 54557035 | 54559144 |
| chr20 | 41821169 | 41823133 |
| chr1 | 102531398 | 102532902 |
| chr13 | 53482280 | 53483406 |
| chr10 | 89014401 | 89016159 |
| chr11 | 69651716 | 69654362 |
| chr13 | 72726517 | 72728761 |
| chr1 | 163854607 | 163855977 |

| | | |
|---|---|---|
| chr1 | 97836389 | 97838208 |
| chr10 | 17199029 | 17201903 |
| chr10 | 79276431 | 79278521 |
| chr9 | 43238810 | 43240751 |
| chr2 | 200031946 | 200033566 |
| chr11 | 112830114 | 112831762 |
| chr2 | 94585052 | 94586707 |
| chr12 | 55373492 | 55375468 |
| chr12 | 55494831 | 55497585 |
| chr11 | 28105590 | 28109697 |
| chr11 | 24552524 | 24554480 |
| chr14 | 46921692 | 46923228 |
| chr8 | 17685410 | 17687050 |
| chr1 | 217866881 | 217868169 |
| chr7 | 80550405 | 80551660 |
| chr2 | 193510079 | 193513302 |
| chr2 | 95497391 | 95499203 |
| chr20 | 29882275 | 29884280 |
| chr7 | 63762936 | 63765158 |
| chr14 | 36072734 | 36076502 |
| chr2 | 62962489 | 62965266 |
| chr2 | 79926415 | 79927876 |
| chr13 | 114340733 | 114342967 |
| chr12 | 98766705 | 98769363 |
| chr1 | 196735907 | 196737757 |
| chr1 | 185933597 | 185936177 |
| chr12 | 44805595 | 44808926 |
| chr9 | 64555552 | 64558717 |
| chr1 | 104087138 | 104091011 |
| chr10 | 127796072 | 127798036 |
| chr2 | 50213880 | 50215899 |
| chr2 | 31869574 | 31871554 |
| chr1 | 97905147 | 97906284 |
| chr6 | 73047983 | 73050258 |
| chr10 | 60644235 | 60645847 |
| chr2 | 51546942 | 51548737 |
| chr20 | 42634996 | 42636645 |
| chr2 | 199686114 | 199687970 |
| chr6 | 74970138 | 74972826 |
| chr6 | 118962052 | 118964102 |
| chr10 | 83708057 | 83710246 |
| chr9 | 62331763 | 62334412 |
| chr17 | 69253627 | 69255072 |
| chr16 | 77398703 | 77400567 |
| chr14 | 95951697 | 95953365 |
| chr2 | 75846793 | 75848516 |
| chr4 | 21270846 | 21272270 |
| chr6 | 160880850 | 160883229 |
| chr1 | 188302194 | 188303635 |

| | | |
|---|---|---|
| chr2 | 182986200 | 182989194 |
| chr1 | 225265321 | 225267218 |
| chr10 | 110644811 | 110649581 |
| chr14 | 56795087 | 56796982 |
| chr13 | 69955256 | 69956769 |
| chr2 | 8679926 | 8682591 |
| chr7 | 57714961 | 57716246 |
| chr9 | 66123732 | 66125493 |
| chr3 | 76814461 | 76816954 |
| chr7 | 84998163 | 85000984 |
| chr12 | 56928919 | 56930321 |
| chr2 | 37334970 | 37338096 |
| chr4 | 176375521 | 176377084 |
| chr1 | 102990005 | 102993456 |
| chr12 | 90977642 | 90979931 |
| chr11 | 45244849 | 45247170 |
| chr6 | 132082139 | 132084162 |
| chr2 | 158454536 | 158457092 |
| chr2 | 42360190 | 42363251 |
| chr2 | 113965318 | 113966872 |
| chr2 | 35878391 | 35880260 |
| chr7 | 70545103 | 70546318 |
| chr1 | 77120326 | 77123082 |
| chr2 | 225786003 | 225787834 |
| chr10 | 55984080 | 55986813 |
| chr15 | 20226047 | 20229088 |
| chr1 | 197807940 | 197810655 |
| chr14 | 39647697 | 39649882 |
| chr14 | 82709591 | 82711596 |
| chr1 | 244846758 | 244849015 |
| chr21 | 20799767 | 20801295 |
| chr5 | 92705690 | 92707071 |
| chr6 | 8101191 | 8103764 |
| chr6 | 80466850 | 80469500 |
| chr12 | 83943439 | 83944608 |
| chr1 | 86394157 | 86397354 |
| chr4 | 158958685 | 158960491 |
| chrY | 6246278 | 6248759 |
| chr7 | 85160358 | 85162137 |
| chr1 | 245873577 | 245875757 |
| chr8 | 110412494 | 110414272 |
| chr5 | 86333469 | 86336047 |
| chr14 | 36042448 | 36045379 |
| chr10 | 86924767 | 86926578 |
| chr20 | 6938374 | 6940896 |
| chr4 | 49232765 | 49236271 |
| chr15 | 36403067 | 36404792 |
| chr12 | 89465432 | 89467501 |
| chr6 | 93289656 | 93290766 |

| | | |
|---|---|---|
| chr13 | 106492914 | 106495214 |
| chr20 | 50729585 | 50731620 |
| chr6 | 95315968 | 95317561 |
| chr4 | 34766242 | 34767512 |
| chr8 | 91970640 | 91972876 |
| chr6 | 70852803 | 70856607 |
| chr6 | 26216292 | 26218213 |
| chr13 | 105175193 | 105177078 |
| chr2 | 197862978 | 197864364 |
| chr2 | 206555334 | 206557352 |
| chr15 | 57213061 | 57215923 |
| chr5 | 165010753 | 165012404 |
| chr14 | 40673610 | 40676016 |
| chr1 | 192337793 | 192339694 |
| chr6 | 57735052 | 57738141 |
| chr2 | 203329825 | 203332225 |
| chr16 | 81576580 | 81578001 |
| chr2 | 105329608 | 105331355 |
| chr20 | 16361005 | 16362990 |
| chr14 | 52729308 | 52731107 |
| chr10 | 59139308 | 59141568 |
| chr21 | 16047370 | 16051021 |
| chr4 | 176902063 | 176903781 |
| chr10 | 65688360 | 65691523 |
| chr1 | 113763874 | 113765229 |
| chr12 | 77393550 | 77396673 |
| chr5 | 168976172 | 168978749 |
| chr13 | 81418100 | 81419532 |
| chr2 | 3461824 | 3463205 |
| chr16_KI270728v1_ | 26508 | 28107 |
| chr15 | 35518635 | 35522225 |
| chr5 | 166261734 | 166265142 |
| chr2 | 161910684 | 161912069 |
| chr9 | 63823485 | 63825650 |
| chr13 | 93519863 | 93521708 |
| chr2 | 82846566 | 82849189 |
| chr8 | 86373863 | 86374941 |
| chr14 | 65580917 | 65582849 |
| chr8 | 12132501 | 12135231 |
| chr15 | 39679276 | 39680653 |
| chr2 | 193084634 | 193086575 |
| chr6 | 64212928 | 64216182 |
| chr4 | 3887550 | 3888570 |
| chr5 | 145182621 | 145184380 |
| chr12 | 77712287 | 77714067 |
| chr10 | 132652955 | 132655202 |
| chr20 | 53592791 | 53595720 |
| chr2 | 61190816 | 61194243 |
| chr2 | 188382564 | 188384371 |

| | | |
|---|---|---|
| chr20 | 25244237 | 25246743 |
| chr2 | 64586313 | 64588266 |
| chr1 | 238345824 | 238348744 |
| chr6 | 56048353 | 56051661 |
| chr21 | 23123109 | 23125274 |
| chr10 | 65216847 | 65218166 |
| chr6 | 47593892 | 47596224 |
| chr10 | 53191844 | 53193004 |
| chr9 | 63860899 | 63862679 |
| chr2 | 185672386 | 185675378 |
| chr1 | 93623534 | 93625377 |
| chr2 | 57833496 | 57835207 |
| chr8 | 92506120 | 92507770 |
| chr1 | 98763469 | 98765932 |
| chr2 | 63419007 | 63420411 |
| chr14 | 27258245 | 27260140 |
| chr11 | 37993788 | 37995735 |
| chr12 | 41128837 | 41130116 |
| chr13 | 85928426 | 85930505 |
| chr3 | 172748846 | 172751218 |
| chr9 | 3359045 | 3361618 |
| chr1 | 148676235 | 148678609 |
| chr1 | 103429298 | 103431457 |
| chr12 | 78806161 | 78808523 |
| chr6 | 45461635 | 45463286 |
| chr2 | 48110467 | 48112512 |
| chr1 | 89915103 | 89917398 |
| chr4 | 59765196 | 59767618 |
| chr6 | 75494049 | 75497003 |
| chr2 | 47462712 | 47464088 |
| chr6 | 72939235 | 72941553 |
| chr6 | 74853517 | 74855738 |
| chr7 | 82592332 | 82594360 |
| chr20 | 32129952 | 32131118 |
| chr7 | 104266742 | 104268657 |
| chr7 | 102748067 | 102749582 |
| chr1 | 195827066 | 195828389 |
| chr16 | 46465032 | 46467126 |
| chr2 | 154243072 | 154245790 |
| chr1 | 79684906 | 79686589 |
| chr16 | 46882993 | 46884963 |
| chr10 | 114969849 | 114972977 |
| chr2 | 184457801 | 184459562 |
| chr12 | 84243600 | 84245844 |
| chr8 | 91313688 | 91315603 |
| chr7 | 79580493 | 79582336 |
| chr10 | 65573211 | 65574360 |
| chr8 | 77465886 | 77468694 |
| chr10 | 89392401 | 89393844 |

| | | |
|---|---|---|
| chr1 | 210858146 | 210860512 |
| chr3 | 80432585 | 80434642 |
| chr2 | 59439086 | 59440424 |
| chr7 | 1566361 | 1567639 |
| chr8 | 39705726 | 39707311 |
| chr14 | 42885633 | 42888040 |
| chr1 | 164781958 | 164784403 |
| chr12 | 84259389 | 84262458 |
| chr2 | 41588409 | 41589939 |
| chr7 | 63094445 | 63095594 |
| chr7 | 53849102 | 53852081 |
| chr1 | 188632118 | 188634058 |
| chr2 | 168960938 | 168963382 |
| chr2 | 35703755 | 35705863 |
| chr7 | 81921094 | 81924152 |
| chr7 | 67042725 | 67045081 |
| chr2 | 186744938 | 186746834 |
| chr22 | 12621412 | 12624026 |
| chr17 | 60907939 | 60910496 |
| chr2 | 66521178 | 66523126 |
| chr16 | 53244205 | 53246511 |
| chr6 | 75002286 | 75003819 |
| chr13 | 65989121 | 65990656 |
| chr14 | 29516684 | 29518512 |
| chr1 | 23018533 | 23020543 |
| chr11 | 95460464 | 95462212 |
| chr1 | 92460685 | 92463189 |
| chr9 | 65152978 | 65154462 |
| chr7 | 89106761 | 89108293 |
| chr13 | 74281136 | 74282850 |
| chr7 | 84740338 | 84742273 |
| chr4 | 19350488 | 19352247 |
| chr13 | 80039470 | 80042711 |
| chr20 | 3208663 | 3210440 |
| chr14 | 38089042 | 38090131 |
| chr6 | 110179560 | 110180875 |
| chr1 | 199910549 | 199913870 |
| chr2 | 53649566 | 53651468 |
| chr2 | 164487690 | 164490009 |
| chrY | 56854846 | 56857897 |
| chr10 | 88934334 | 88935934 |
| chr14 | 46299581 | 46301071 |
| chr10 | 37144107 | 37145500 |
| chr2 | 35276397 | 35278240 |
| chr2 | 164821407 | 164823162 |
| chr2 | 70549188 | 70550441 |
| chr14 | 57190411 | 57191625 |
| chr14 | 26459173 | 26462955 |
| chr16 | 46384524 | 46386086 |

| | | |
|---|---|---|
| chr2 | 55215217 | 55216924 |
| chr10 | 62491555 | 62493859 |
| chr2 | 184807179 | 184809951 |
| chr11 | 16615038 | 16617786 |
| chr12 | 33581385 | 33582835 |
| chr20 | 1158731 | 1160505 |
| chr14 | 53623272 | 53624844 |
| chr2 | 58193944 | 58195744 |
| chr9 | 6985586 | 6988069 |
| chr16 | 77175978 | 77177539 |
| chr14 | 89922670 | 89925321 |
| chr3 | 80557172 | 80558495 |
| chr12 | 88362606 | 88364519 |
| chr6 | 99328429 | 99330987 |
| chr2 | 30642783 | 30644791 |
| chr11 | 105829489 | 105831251 |
| chr14 | 49925940 | 49928257 |
| chr9 | 39668561 | 39671061 |
| chr14 | 60557456 | 60560152 |
| chr2 | 226861052 | 226863105 |
| chr1 | 77509133 | 77510739 |
| chr2 | 189727601 | 189729696 |
| chr2 | 213074137 | 213075269 |
| chr22 | 15886620 | 15889909 |
| chr4 | 29080671 | 29082638 |
| chr8 | 17843438 | 17846373 |
| chr14 | 24942565 | 24944673 |
| chr2 | 67693336 | 67695579 |
| chr2 | 166334286 | 166336233 |
| chr2 | 95938316 | 95939655 |
| chr14 | 29796741 | 29798244 |
| chr15 | 90793418 | 90795604 |
| chr14 | 41355125 | 41357837 |
| chr14 | 36793494 | 36794880 |
| chr9 | 4695689 | 4697503 |
| chr2 | 224947912 | 224949027 |
| chr1 | 198245949 | 198247307 |
| chr2 | 42315880 | 42317573 |
| chr12 | 98535476 | 98538318 |
| chr1 | 20941499 | 20944172 |
| chr1 | 157670317 | 157672667 |
| chr12 | 38680421 | 38681483 |
| chr10 | 71836593 | 71838463 |
| chr14 | 101761924 | 101763615 |
| chr5 | 124111977 | 124113570 |
| chr5 | 109811200 | 109813003 |
| chr10 | 50833245 | 50836240 |
| chr3 | 86772658 | 86775157 |
| chr13 | 104137647 | 104140371 |

| | | |
|---|---|---|
| chr20 | 21386127 | 21390039 |
| chr2 | 164864038 | 164866392 |
| chr10 | 46514983 | 46518571 |
| chr10 | 46629659 | 46632164 |
| chr1 | 167225846 | 167227923 |
| chr12 | 83938370 | 83941135 |
| chr17 | 74512172 | 74514117 |
| chr6 | 80666485 | 80668305 |
| chr5 | 88872514 | 88874859 |
| chr7 | 94015617 | 94016780 |
| chr20 | 41612448 | 41614773 |
| chr9 | 39781956 | 39783453 |
| chr5 | 121700278 | 121702893 |
| chr14 | 99015239 | 99017766 |
| chr6 | 55401995 | 55403882 |
| chr2 | 57430649 | 57432033 |
| chr1 | 203025214 | 203027244 |
| chr7 | 62363623 | 62364640 |
| chr6 | 140713303 | 140714461 |
| chr12 | 79404063 | 79405815 |
| chr2 | 103308611 | 103310115 |
| chr2 | 184969522 | 184972308 |
| chr14 | 26648495 | 26650757 |
| chr7 | 8760774 | 8762967 |
| chr14_GL000194v1_ | 187915 | 190015 |
| chr5 | 59033262 | 59035235 |
| chr20 | 30720611 | 30722491 |
| chr2 | 68185166 | 68186980 |
| chr8 | 116234337 | 116239793 |
| chr10 | 63228244 | 63231347 |
| chr1 | 219448949 | 219450550 |
| chr1 | 168470550 | 168471952 |
| chr7 | 80259183 | 80261427 |
| chr14 | 28484881 | 28486813 |
| chr1 | 94237307 | 94240247 |
| chr7 | 52485642 | 52487908 |
| chr10 | 52515158 | 52516560 |
| chr2 | 76973921 | 76975485 |
| chr21 | 14273892 | 14275658 |
| chr1 | 89633893 | 89635262 |
| chrX | 128666673 | 128668076 |
| chr6 | 135737464 | 135739289 |
| chr14 | 51079201 | 51081507 |
| chr10 | 59258261 | 59260010 |
| chr15 | 27563499 | 27565535 |
| chr9 | 65693440 | 65694656 |
| chr12 | 77977742 | 77979436 |
| chr3 | 51597490 | 51598489 |
| chr4 | 34828052 | 34830565 |

| | | |
|---|---|---|
| chr22 | 15859160 | 15862064 |
| chr10 | 112666790 | 112668685 |
| chr5 | 128995778 | 128997476 |
| chr10 | 96533017 | 96534894 |
| chr2 | 94271110 | 94273275 |
| chr20 | 4366522 | 4368243 |
| chr7 | 11776068 | 11777683 |
| chr10 | 64531018 | 64532430 |
| chr14 | 50438054 | 50440363 |
| chr14 | 40647972 | 40650143 |
| chr6 | 55985223 | 55987186 |
| chr2 | 159129182 | 159131893 |
| chr11 | 79139427 | 79140493 |
| chr17 | 26734285 | 26735495 |
| chr12 | 38222575 | 38224339 |
| chr16 | 46412595 | 46414993 |
| chr3 | 29181714 | 29184517 |
| chr10 | 58363789 | 58365352 |
| chr5 | 113975238 | 113977499 |
| chr4 | 169146709 | 169148125 |
| chr1 | 6820777 | 6822614 |
| chr4 | 35371617 | 35373474 |
| chr21 | 20812494 | 20816060 |
| chr22_KI270734v1_ | 84146 | 85777 |
| chr12 | 21710988 | 21713368 |
| chr5 | 156025489 | 156027178 |
| chr5 | 90887562 | 90890224 |
| chr8 | 74327207 | 74328776 |
| chr4 | 166591428 | 166593838 |
| chr10 | 64652944 | 64655211 |
| chr14 | 41957823 | 41959075 |
| chrX | 12012668 | 12014369 |
| chr1 | 72282793 | 72284652 |
| chr20 | 8882322 | 8884614 |
| chr2 | 185250149 | 185252046 |
| chr10 | 75110784 | 75113375 |
| chr14 | 33569769 | 33572428 |
| chr14 | 31456092 | 31459211 |
| chr7 | 84355768 | 84356987 |
| chr14 | 43681870 | 43684037 |
| chr1 | 89907281 | 89908436 |
| chr4 | 107249926 | 107253165 |
| chr2 | 219723849 | 219725932 |
| chr1 | 98026189 | 98029819 |
| chr14 | 99382068 | 99383282 |
| chr14 | 36817013 | 36819922 |
| chr4 | 39063256 | 39065227 |
| chr2 | 55998992 | 56000023 |
| chr1 | 195514851 | 195516829 |

| | | |
|---|---|---|
| chr7 | 85593760 | 85597245 |
| chr1 | 68382595 | 68383606 |
| chr6 | 121032898 | 121034574 |
| chr6 | 24708520 | 24710570 |
| chr16 | 75763384 | 75765022 |
| chr12 | 66103192 | 66105023 |
| chr17 | 69703084 | 69705584 |
| chr10 | 68900314 | 68902082 |
| chr15 | 65598358 | 65600271 |
| chr2 | 41561127 | 41562563 |
| chr1 | 191952875 | 191954274 |
| chr11 | 93735943 | 93737966 |
| chr14 | 88593501 | 88595555 |
| chr11 | 18802204 | 18804052 |
| chr8 | 76673735 | 76674977 |
| chr1 | 74071653 | 74073267 |
| chr1 | 92486511 | 92488321 |
| chr2 | 193109726 | 193112894 |
| chr14 | 83921907 | 83923357 |
| chr5 | 117541211 | 117542985 |
| chr15 | 94627366 | 94628852 |
| chr9 | 64546111 | 64549509 |
| chr11 | 21986575 | 21989405 |
| chr7 | 65547485 | 65549013 |
| chr14 | 41028992 | 41031467 |
| chr1 | 99300354 | 99302147 |
| chr7 | 82001111 | 82005047 |
| chr4 | 30724924 | 30727249 |
| chr15 | 35564740 | 35567352 |
| chr2 | 57125680 | 57127641 |
| chr2 | 187440836 | 187443343 |
| chr8 | 97036258 | 97038291 |
| chr20 | 49101935 | 49103121 |
| chr2 | 57428929 | 57430235 |
| chr12 | 68686281 | 68687795 |
| chr10 | 102420063 | 102422667 |
| chr17 | 4795655 | 4797997 |
| chr2 | 58811133 | 58814146 |
| chr1 | 78916129 | 78919350 |
| chr10 | 74850488 | 74852771 |
| chr10 | 57684628 | 57688139 |
| chr2 | 161159172 | 161161054 |
| chr6 | 81786317 | 81788853 |
| chr4 | 93023738 | 93025883 |
| chr9 | 22427216 | 22429309 |
| chr14 | 26916380 | 26917779 |
| chr20 | 29876836 | 29879739 |
| chr14 | 87639109 | 87643288 |
| chr14 | 85285176 | 85286404 |

| | | |
|---|---|---|
| chr18 | 25286997 | 25288099 |
| chr17 | 75224705 | 75226193 |
| chr16 | 65000984 | 65002813 |
| chr8 | 98657685 | 98660123 |
| chr16 | 63347947 | 63349603 |
| chr14 | 105058529 | 105060439 |
| chr21 | 34936943 | 34937949 |
| chr11 | 43837563 | 43838828 |
| chr14 | 83473675 | 83475936 |
| chr2 | 99986080 | 99987527 |
| chr14 | 94752467 | 94754415 |
| chr6 | 122019944 | 122021194 |
| chr10 | 4544770 | 4546354 |
| chr4 | 35067216 | 35069325 |
| chr2 | 181565302 | 181568073 |
| chr2 | 188891892 | 188893741 |
| chr17 | 21341857 | 21345765 |
| chr1 | 157182808 | 157184542 |
| chr4 | 90758523 | 90760807 |
| chr6 | 56956548 | 56958184 |
| chr1_KI270763v1_a | 49461 | 51910 |
| chr13 | 67328635 | 67331790 |
| chr2 | 226129404 | 226131339 |
| chr10 | 67008990 | 67011956 |
| chrUn_GL000195v1 | 154600 | 155841 |
| chr13 | 75591015 | 75593181 |
| chr3 | 174097605 | 174099904 |
| chr12 | 4085532 | 4087348 |
| chrY | 24289225 | 24290511 |
| chr8 | 115931479 | 115933102 |
| chr10 | 102833731 | 102835490 |
| chr10 | 60982757 | 60985100 |
| chr9 | 71663467 | 71664669 |
| chr12 | 54027589 | 54028903 |
| chr10 | 120656960 | 120658997 |
| chr3 | 114444648 | 114447503 |
| chr12 | 83664973 | 83668109 |
| chr5 | 55146245 | 55148205 |
| chr12 | 28385050 | 28387706 |
| chr1 | 75122062 | 75124425 |
| chr13 | 25978756 | 25979968 |
| chr1 | 101971239 | 101972382 |
| chr11 | 26238426 | 26240999 |
| chr3 | 158297299 | 158299465 |
| chr1 | 79586774 | 79588759 |
| chr12 | 73126607 | 73129185 |
| chr2 | 186043861 | 186045489 |
| chr10 | 61748462 | 61750012 |
| chr6 | 100591945 | 100594204 |

| | | |
|---|---|---|
| chr1 | 104823791 | 104826298 |
| chr3 | 105925938 | 105928443 |
| chr14 | 62700556 | 62702803 |
| chr10 | 3171894 | 3173386 |
| chr6 | 135534060 | 135535962 |
| chr9 | 4210525 | 4212754 |
| chr2 | 95961926 | 95964238 |
| chr9 | 65855517 | 65856730 |
| chr1 | 104781972 | 104783999 |
| chr14 | 48499772 | 48501804 |
| chr6 | 161273696 | 161276242 |
| chr14 | 68293941 | 68296008 |
| chr1 | 104400343 | 104401527 |
| chr7 | 149142461 | 149143720 |
| chr4 | 104409536 | 104411238 |
| chr2 | 103024780 | 103026769 |
| chr10 | 64859342 | 64860916 |
| chr14 | 25893129 | 25896319 |
| chr17 | 57604077 | 57605749 |
| chr4 | 89724723 | 89726831 |
| chr9 | 30763031 | 30764824 |
| chr8 | 81455366 | 81456578 |
| chr14 | 19937971 | 19939204 |
| chr1 | 143204299 | 143207568 |
| chr11 | 22693280 | 22696003 |
| chr14 | 30306682 | 30310208 |
| chr14 | 101724000 | 101725166 |
| chr4 | 30981057 | 30983016 |
| chr3 | 174084948 | 174087030 |
| chr14 | 28765938 | 28767609 |
| chr15 | 37692577 | 37694646 |
| chr6 | 123380726 | 123385234 |
| chr9 | 66601880 | 66603120 |
| chr1 | 112644135 | 112645797 |
| chr2 | 43941501 | 43942925 |
| chr16 | 87822573 | 87825783 |
| chr13 | 59772018 | 59773185 |
| chr14 | 53383576 | 53386048 |
| chr14 | 99938082 | 99940009 |
| chr7 | 10385037 | 10388618 |
| chr11 | 100914850 | 100916621 |
| chr4 | 9223404 | 9225362 |
| chr14 | 71697315 | 71699763 |
| chr9 | 63559172 | 63560864 |
| chr12 | 38618186 | 38619841 |
| chr1 | 185686247 | 185689498 |
| chr8 | 25215687 | 25217531 |
| chr5 | 88824617 | 88826998 |
| chr9 | 20565054 | 20566799 |

| | | |
|---|---|---|
| chr7 | 10766189 | 10767473 |
| chr2 | 49522418 | 49523620 |
| chr1 | 106380457 | 106383210 |
| chr5 | 111097822 | 111098945 |
| chr2 | 226053592 | 226055451 |
| chr2 | 103113124 | 103115688 |
| chr2 | 164599516 | 164602550 |
| chr7 | 13186703 | 13188199 |
| chr2 | 70247310 | 70248540 |
| chr15 | 92902675 | 92905218 |
| chr6 | 95783708 | 95785824 |
| chr10 | 61460699 | 61462038 |
| chr14 | 86489018 | 86491704 |
| chr2 | 181123155 | 181125011 |
| chr1 | 204922750 | 204923904 |
| chr7 | 85713769 | 85716600 |
| chr10 | 129289115 | 129290947 |
| chr1 | 248238043 | 248239659 |
| chr18 | 15183870 | 15185144 |
| chr12 | 75057589 | 75060217 |
| chr4 | 113318065 | 113319998 |
| chr9 | 37079278 | 37081005 |
| chr2 | 205414069 | 205416009 |
| chr2 | 194052717 | 194054986 |
| chr14 | 28875606 | 28877391 |
| chr3 | 146072801 | 146074830 |
| chr7 | 6714087 | 6715318 |
| chr4 | 29502143 | 29504350 |
| chr5 | 44866266 | 44868126 |
| chr8 | 66608886 | 66610982 |
| chr1 | 87011111 | 87012646 |
| chr8 | 95447681 | 95448681 |
| chr9 | 68241025 | 68243306 |
| chr2 | 82074654 | 82076946 |
| chr2 | 228141125 | 228142602 |
| chr8 | 98704058 | 98706311 |
| chr2 | 52994416 | 52997207 |
| chr1 | 154337580 | 154340116 |
| chr12 | 85366956 | 85368731 |
| chr8 | 115625477 | 115627723 |
| chr8 | 41063745 | 41065660 |
| chr18 | 56862837 | 56864662 |
| chr2 | 88015723 | 88017456 |
| chr1 | 98045796 | 98049813 |
| chr19 | 3306849 | 3308009 |
| chr2 | 161198668 | 161201227 |
| chr4 | 126991876 | 126993116 |
| chr7 | 100328533 | 100330426 |
| chr1 | 97697015 | 97698920 |

| | | |
|---|---|---|
| chr22 | 16311793 | 16313489 |
| chr6 | 9257009 | 9258589 |
| chr2 | 186862550 | 186864036 |
| chr15 | 20187143 | 20188856 |
| chr20 | 7938127 | 7939706 |
| chr1 | 190490834 | 190493486 |
| chr6 | 64081253 | 64082840 |
| chr7 | 85290404 | 85291712 |
| chr12 | 83952695 | 83954640 |
| chr6 | 121810470 | 121812629 |
| chr4 | 34903377 | 34906076 |
| chr7 | 21481730 | 21482919 |
| chr13 | 67112864 | 67113892 |
| chr10 | 128696660 | 128698582 |
| chr14 | 45027626 | 45029156 |
| chr1 | 16878729 | 16880824 |
| chr10 | 87333417 | 87334431 |
| chr11 | 55900307 | 55902849 |
| chr12 | 40513185 | 40515140 |
| chr1 | 160575988 | 160577607 |
| chr13 | 111939374 | 111940911 |
| chr4 | 37432409 | 37434402 |
| chr7 | 83270804 | 83272823 |
| chr13 | 69930926 | 69932483 |
| chr7 | 104126285 | 104128451 |
| chr1 | 170542278 | 170544409 |
| chr14 | 42079519 | 42081934 |
| chr10 | 60453860 | 60455529 |
| chr10 | 56348978 | 56350944 |
| chr7 | 23682988 | 23684219 |
| chr1 | 119007591 | 119009272 |
| chr14 | 90288060 | 90289756 |
| chr14 | 104250580 | 104252402 |
| chr15 | 83256012 | 83258277 |
| chr10 | 5489906 | 5491716 |
| chr2 | 220587972 | 220590159 |
| chr7 | 77604071 | 77605486 |
| chr14 | 29832743 | 29834902 |
| chr12 | 15178670 | 15182968 |
| chr4 | 28109352 | 28111721 |
| chr2 | 78184911 | 78186354 |
| chr2 | 216000699 | 216002471 |
| chr2 | 20726999 | 20728982 |
| chr14 | 59253794 | 59255775 |
| chr11 | 94134783 | 94136267 |
| chr14 | 37987817 | 37989984 |
| chr20 | 55435861 | 55438523 |
| chr10 | 42786883 | 42788530 |
| chr5 | 27770812 | 27771879 |

| | | |
|---|---|---|
| chr8 | 111909899 | 111910998 |
| chr2 | 19193713 | 19195519 |
| chr2 | 58227581 | 58230014 |
| chr2 | 161945531 | 161947347 |
| chr11 | 28465780 | 28467641 |
| chr11 | 106520240 | 106521755 |
| chr4 | 176390616 | 176392588 |
| chr1 | 16622849 | 16624463 |
| chr7 | 104152620 | 104154358 |
| chr7 | 79786241 | 79789577 |
| chr2 | 207038588 | 207041030 |
| chr13 | 96220402 | 96221613 |
| chr20 | 44211884 | 44213730 |
| chr1 | 87804111 | 87805650 |
| chr5 | 5121326 | 5124511 |
| chr2 | 170013776 | 170017770 |
| chr12 | 85138803 | 85140555 |
| chr11 | 3653361 | 3655531 |
| chr12 | 100338068 | 100340315 |
| chr12 | 77952274 | 77954100 |
| chr10 | 56346011 | 56347672 |
| chr2 | 97642284 | 97643612 |
| chr12 | 104386985 | 104388395 |
| chr12 | 41449663 | 41452330 |
| chr12 | 84742080 | 84746139 |
| chr6 | 83573592 | 83576450 |
| chr5 | 81307693 | 81309740 |
| chr4 | 93076588 | 93079082 |
| chr14 | 43379655 | 43381117 |
| chr6 | 81996602 | 81998413 |
| chr1 | 99900290 | 99902667 |
| chr8 | 7779412 | 7781955 |
| chr14 | 75175482 | 75177458 |
| chr2 | 193186148 | 193187715 |
| chr15 | 92090922 | 92092343 |
| chr20 | 13424754 | 13426515 |
| chr1 | 206270361 | 206272545 |
| chr10 | 99443871 | 99445719 |
| chr6 | 122824579 | 122826517 |
| chr10 | 44462879 | 44464517 |
| chr2 | 31902377 | 31904051 |
| chr1 | 220569843 | 220572083 |
| chr1 | 93576260 | 93579303 |
| chr14 | 53513535 | 53515761 |
| chr2 | 65222219 | 65223858 |
| chr10 | 58805559 | 58808054 |
| chr11 | 25471994 | 25475010 |
| chr3 | 110808369 | 110810946 |
| chr14 | 96564125 | 96566759 |

| | | |
|---|---|---|
| chr8 | 138721964 | 138724346 |
| chr6 | 75491198 | 75493825 |
| chr6 | 93328173 | 93330415 |
| chr4 | 28929549 | 28931468 |
| chr10 | 108878953 | 108880833 |
| chr14 | 39106194 | 39109210 |
| chr7 | 114662465 | 114664206 |
| chr7 | 91833853 | 91835476 |
| chr14 | 26770466 | 26772997 |
| chr1 | 219994830 | 219995994 |
| chr11 | 81324956 | 81326438 |
| chr2 | 185003140 | 185005225 |
| chr3 | 98248973 | 98249996 |
| chr14 | 23364171 | 23367320 |
| chr1 | 50557311 | 50558934 |
| chr11 | 20915694 | 20917742 |
| chr20 | 62058215 | 62059715 |
| chr6 | 78563839 | 78566455 |
| chr7 | 93957568 | 93959351 |
| chr11 | 103232842 | 103234108 |
| chr20 | 63740435 | 63742217 |
| chr14 | 55648184 | 55650220 |
| chr12 | 44567421 | 44569042 |
| chr17 | 21632906 | 21634218 |
| chr12 | 78771066 | 78772785 |
| chr7 | 158915073 | 158917749 |
| chr10 | 76646992 | 76648763 |
| chr2 | 162216757 | 162218819 |
| chr10 | 72215848 | 72217318 |
| chr1 | 97630086 | 97633241 |
| chr1 | 150512989 | 150516247 |
| chr11 | 22663252 | 22665112 |
| chr17 | 58833601 | 58834705 |
| chr1 | 98009420 | 98011442 |
| chr5 | 27416694 | 27418864 |
| chr10 | 66796476 | 66798104 |
| chr10 | 81657304 | 81659223 |
| chr5 | 45826323 | 45828699 |
| chr4 | 48569687 | 48570838 |
| chr11 | 99555495 | 99557996 |
| chr1 | 103202568 | 103204545 |
| chr5 | 27862097 | 27863890 |
| chr7 | 49073659 | 49075818 |
| chr12 | 44267598 | 44268604 |
| chr12 | 63515099 | 63517463 |
| chrY | 10978245 | 10979253 |
| chr15 | 39230507 | 39232498 |
| chr14 | 99793928 | 99795407 |
| chr3 | 193216317 | 193217442 |

| | | |
|---|---|---|
| chr10 | 17071207 | 17072702 |
| chr14 | 54578415 | 54579955 |
| chr8 | 71289736 | 71291086 |
| chr20 | 13343677 | 13345975 |
| chr6 | 82450501 | 82452853 |
| chr22 | 12615643 | 12618626 |
| chr7 | 80369696 | 80373940 |
| chr22 | 12295002 | 12296839 |
| chr10 | 89710154 | 89711897 |
| chr14 | 84171970 | 84173866 |
| chr12 | 24234674 | 24235818 |
| chr7 | 82392327 | 82393864 |
| chr12 | 75098902 | 75100710 |
| chr9 | 68341607 | 68344925 |
| chr14 | 105476187 | 105478197 |
| chr7 | 80590268 | 80592373 |
| chr10 | 43204237 | 43207772 |
| chr2 | 226792898 | 226794627 |
| chr2 | 52645130 | 52647566 |
| chr2 | 88955435 | 88958573 |
| chr3 | 62461697 | 62463788 |
| chr15 | 33691929 | 33693215 |
| chr4 | 62114434 | 62115764 |
| chr3 | 193605406 | 193606593 |
| chr6 | 76317593 | 76319333 |
| chr1 | 198863890 | 198866424 |
| chr12 | 72033646 | 72035169 |
| chr2 | 161827654 | 161828945 |
| chr14 | 79943752 | 79945507 |
| chr4 | 151949680 | 151953214 |
| chr1 | 143501349 | 143504459 |
| chr5 | 154754316 | 154755982 |
| chr4 | 131526410 | 131527416 |
| chr16 | 31699222 | 31700772 |
| chr1 | 93896810 | 93898748 |
| chr7 | 53087652 | 53089369 |
| chr9 | 63551573 | 63552672 |
| chr7 | 88344594 | 88346243 |
| chr2 | 79664644 | 79666403 |
| chr14 | 26969774 | 26973989 |
| chr6 | 62231980 | 62235481 |
| chr2 | 213051724 | 213054411 |
| chr15 | 49641670 | 49644486 |
| chr2 | 36520836 | 36522054 |
| chr14 | 68178216 | 68180773 |
| chr10 | 56147595 | 56149120 |
| chr2 | 145659474 | 145662795 |
| chr10 | 55619897 | 55621714 |
| chr2 | 63584390 | 63586859 |

| | | |
|---|---|---|
| chr12 | 83046494 | 83048970 |
| chr14 | 90370713 | 90373389 |
| chr8 | 65115721 | 65117654 |
| chr2 | 214236375 | 214239220 |
| chr5 | 156584286 | 156586067 |
| chr7 | 78734754 | 78736641 |
| chr13 | 91904633 | 91905670 |
| chr20 | 11891067 | 11894065 |
| chr4 | 32771602 | 32772678 |
| chr12 | 41248108 | 41250076 |
| chr1 | 100505892 | 100507138 |
| chr2 | 41732899 | 41735801 |
| chr11 | 96015023 | 96016877 |
| chr3 | 142999991 | 143001968 |
| chr15 | 25364775 | 25367626 |
| chr21 | 9982367 | 9985453 |
| chr1 | 227597950 | 227599525 |
| chr15 | 23346851 | 23348194 |
| chr2 | 226079250 | 226080804 |
| chr15 | 35416073 | 35418646 |
| chr1 | 85156232 | 85160181 |
| chr2 | 206115735 | 206117090 |
| chr12 | 78985006 | 78986657 |
| chr14 | 41795465 | 41798842 |
| chr2 | 197719306 | 197721411 |
| chr1 | 148661130 | 148662715 |
| chr20 | 24378595 | 24380189 |
| chr2 | 67790152 | 67792582 |
| chr12 | 18136550 | 18138616 |
| chr2 | 57774776 | 57777903 |
| chr7 | 75352605 | 75354215 |
| chr10 | 96405293 | 96407248 |
| chr9 | 67803282 | 67805230 |
| chr1 | 165597668 | 165600064 |
| chr2 | 95980773 | 95982557 |
| chr14 | 104424079 | 104425887 |
| chr4 | 156568213 | 156569843 |
| chr1 | 103667641 | 103669459 |
| chr15 | 47515588 | 47517183 |
| chr1 | 117948746 | 117951432 |
| chr2 | 224519012 | 224520260 |
| chr22 | 16437709 | 16439535 |
| chr14 | 42769973 | 42771703 |
| chr5 | 38441386 | 38443016 |
| chr10 | 56724630 | 56726745 |
| chr15_KI270905v1_ | 806300 | 809256 |
| chr16 | 67427906 | 67430361 |
| chr14 | 101899163 | 101900915 |
| chr14 | 48518302 | 48521528 |

| | | |
|---|---|---|
| chr22 | 15708500 | 15710395 |
| chr4 | 63789423 | 63791937 |
| chr1 | 180271138 | 180272613 |
| chr2 | 84611243 | 84612964 |
| chr6 | 98040638 | 98041747 |
| chr9 | 67847533 | 67849005 |
| chr11 | 97892553 | 97894164 |
| chr14 | 35277600 | 35280028 |
| chr6 | 324269 | 328611 |
| chr14 | 79688660 | 79690652 |
| chr20 | 25996791 | 25998612 |
| chr20 | 56864749 | 56866009 |
| chr7 | 112110687 | 112112154 |
| chr3 | 107091187 | 107092629 |
| chr3 | 83174147 | 83176214 |
| chr12 | 90712164 | 90715101 |
| chr15_KI270905v1_ | 603854 | 607677 |
| chr12 | 85055775 | 85058483 |
| chr9 | 39039225 | 39040963 |
| chr13 | 22028238 | 22029360 |
| chr11 | 24523603 | 24525474 |
| chr8 | 28888644 | 28890532 |
| chr20 | 52045989 | 52049156 |
| chr2 | 41146806 | 41149154 |
| chr4 | 135212897 | 135215431 |
| chr16 | 73713723 | 73715782 |
| chr9 | 66023560 | 66024682 |
| chr1 | 195298148 | 195300043 |
| chr20 | 35363030 | 35364329 |
| chr1 | 230684592 | 230685892 |
| chr10 | 35127814 | 35130952 |
| chr20 | 41171729 | 41174326 |
| chr2 | 172064542 | 172067836 |
| chr17 | 75977504 | 75979770 |
| chr14 | 30178957 | 30181242 |
| chr1 | 196414588 | 196416351 |
| chr12 | 122507126 | 122508820 |
| chr1 | 97726197 | 97728989 |
| chr2 | 57405493 | 57407528 |
| chr11 | 47500957 | 47503497 |
| chr2 | 188783525 | 188785309 |
| chr15 | 37076348 | 37078792 |
| chr2 | 192334280 | 192335954 |
| chr14 | 30699628 | 30701287 |
| chr7 | 77111619 | 77113875 |
| chr7 | 120950203 | 120952696 |
| chr1 | 190105888 | 190108522 |
| chr2 | 18073272 | 18076488 |
| chr2 | 205390182 | 205394034 |

| | | |
|---|---|---|
| chr11 | 101745177 | 101747040 |
| chr16_KI270728v1_ | 1852988 | 1855241 |
| chr17 | 81859246 | 81860445 |
| chr3 | 95337353 | 95339252 |
| chr1 | 185274503 | 185277148 |
| chr1 | 159982849 | 159985400 |
| chr2 | 97011913 | 97013742 |
| chr7 | 61036126 | 61038509 |
| chr13 | 82253092 | 82255069 |
| chr10 | 75030094 | 75033609 |
| chr15 | 51497376 | 51499873 |
| chr1 | 117440629 | 117442260 |
| chr2 | 172088510 | 172090710 |
| chr6 | 75338102 | 75340505 |
| chr1 | 83934629 | 83937009 |
| chr2 | 198994835 | 198996029 |
| chr2 | 86914790 | 86916937 |
| chr21 | 41609532 | 41611936 |
| chr5_KI270897v1_a | 754498 | 755867 |
| chr12 | 79424670 | 79426335 |
| chr1 | 104167995 | 104170164 |
| chr15_KI270905v1_ | 4792264 | 4794013 |
| chr2 | 178530473 | 178533100 |
| chr14 | 46768798 | 46769982 |
| chr5 | 128132143 | 128133523 |
| chr10 | 110066461 | 110068460 |
| chr21 | 29024412 | 29026261 |
| chr14 | 19419541 | 19420873 |
| chr2 | 185175405 | 185177291 |
| chr3 | 88228860 | 88232734 |
| chr4 | 115525834 | 115527076 |
| chr1 | 196676782 | 196678829 |
| chr5 | 119632845 | 119636260 |
| chr2 | 207369496 | 207371333 |
| chr10 | 38161201 | 38162934 |
| chr7 | 105017492 | 105019696 |
| chr3 | 89059037 | 89061054 |
| chr12 | 86905394 | 86907705 |
| chr13 | 78981057 | 78983007 |
| chr15 | 57920740 | 57922644 |
| chr14 | 58138015 | 58139998 |
| chr15 | 36473810 | 36476285 |
| chr10 | 112586727 | 112588607 |
| chr1 | 187250918 | 187252902 |
| chr5 | 81190837 | 81192690 |
| chr15 | 99248865 | 99250796 |
| chr20 | 24474831 | 24476123 |
| chr6 | 135421176 | 135422510 |
| chr10 | 74452080 | 74454726 |

| | | |
|---|---|---|
| chrUn_GL000220v1 | 54992 | 57507 |
| chr2 | 213245806 | 213247680 |
| chr16 | 228907 | 230639 |
| chr12 | 42109434 | 42111285 |
| chr15 | 77940281 | 77941612 |
| chr14 | 39979830 | 39981805 |
| chr4 | 15230776 | 15232381 |
| chr22 | 22267472 | 22268695 |
| chr9 | 67775801 | 67777580 |
| chr1 | 2098850 | 2100566 |
| chr14 | 85879630 | 85881786 |
| chr9 | 21703510 | 21705094 |
| chr2 | 102476596 | 102478620 |
| chr3 | 7482030 | 7483342 |
| chr10 | 58264062 | 58265813 |
| chr6 | 60545157 | 60547456 |
| chr14 | 61883767 | 61885473 |
| chr7 | 82374789 | 82378094 |
| chr7 | 102671164 | 102672565 |
| chr2 | 28940209 | 28942420 |
| chr11 | 87859733 | 87861174 |
| chr5 | 123862713 | 123865157 |
| chr2 | 182520450 | 182524051 |
| chr14 | 41476566 | 41477803 |
| chr6 | 97858590 | 97860263 |
| chr12 | 65230828 | 65232665 |
| chr12 | 123329878 | 123331932 |
| chr1 | 195431530 | 195433560 |
| chr7 | 81488591 | 81491578 |
| chr12 | 72383606 | 72385622 |
| chr16 | 30241603 | 30243602 |
| chr6 | 41818157 | 41820035 |
| chr19 | 41263096 | 41264862 |
| chr6 | 82478819 | 82480840 |
| chr10 | 131307796 | 131310496 |
| chr14 | 82876764 | 82878528 |
| chr6 | 74868850 | 74871564 |
| chr2 | 165348724 | 165350670 |
| chr14 | 60825499 | 60828273 |
| chr2 | 50269591 | 50272518 |
| chr11 | 87939912 | 87941003 |
| chr1 | 100082325 | 100084251 |
| chr1 | 70022428 | 70024374 |
| chr2 | 205188747 | 205191458 |
| chr2 | 186640047 | 186642572 |
| chr16 | 69672207 | 69675282 |
| chr12 | 62485937 | 62488351 |
| chr1 | 146695278 | 146696393 |
| chr2 | 51282100 | 51285293 |

| | | |
|---|---|---|
| chr1 | 219148868 | 219151078 |
| chr2 | 139964467 | 139966826 |
| chr10 | 76029214 | 76030943 |
| chr1 | 49885498 | 49886829 |
| chr13 | 52712141 | 52714533 |
| chr4 | 10157428 | 10158878 |
| chr2 | 178446812 | 178448731 |
| chr2 | 85545275 | 85546451 |
| chr1 | 148926195 | 148927444 |
| chr1 | 84184260 | 84185941 |
| chr14 | 41440630 | 41442316 |
| chr7 | 52146666 | 52148968 |
| chr8 | 114450896 | 114453366 |
| chr21 | 9777009 | 9780293 |
| chr10 | 52443410 | 52444974 |
| chr15_KI270905v1_ | 2497659 | 2499509 |
| chr3 | 106685075 | 106686407 |
| chr6 | 25555823 | 25557745 |
| chr10 | 60027023 | 60030227 |
| chr8 | 76584150 | 76586392 |
| chr5 | 132491127 | 132493159 |
| chr14 | 87058230 | 87060727 |
| chr2 | 171744884 | 171746415 |
| chr12 | 70595584 | 70597345 |
| chr14 | 28551985 | 28555644 |
| chr6 | 81065205 | 81068514 |
| chr1 | 46651849 | 46654398 |
| chr1 | 144551048 | 144552853 |
| chr17 | 50079807 | 50081812 |
| chr20 | 10400970 | 10403001 |
| chr4 | 171830105 | 171832529 |
| chr6 | 64838653 | 64839995 |
| chr17_GL383563v3_ | 36836 | 40114 |
| chr22 | 27918687 | 27920202 |
| chr2 | 18499934 | 18501562 |
| chr5 | 81980578 | 81982028 |
| chr7 | 56322895 | 56325006 |
| chr10 | 93050956 | 93053138 |
| chr7 | 61085210 | 61086905 |
| chr13 | 76990698 | 76993041 |
| chr11 | 107513823 | 107515282 |
| chr5 | 119589789 | 119591820 |
| chr9 | 66318696 | 66323139 |
| chr1 | 173867549 | 173871523 |
| chr5 | 22413540 | 22415051 |
| chrX | 123894521 | 123895524 |
| chr2 | 68517150 | 68518162 |
| chr14 | 41608180 | 41610392 |
| chr14 | 33436479 | 33437931 |

| | | |
|---|---|---|
| chr14 | 22085156 | 22087241 |
| chr2 | 196296977 | 196298702 |
| chr6 | 76877929 | 76880332 |
| chr12 | 85060446 | 85062437 |
| chr7 | 81726275 | 81729268 |
| chr1 | 69872359 | 69874530 |
| chr11 | 89863315 | 89867213 |
| chr13 | 64932613 | 64934436 |
| chr16 | 76378122 | 76380600 |
| chr2 | 85319821 | 85322065 |
| chr2 | 142413364 | 142415079 |
| chr20 | 47887691 | 47889994 |
| chr21 | 21293496 | 21295640 |
| chr1 | 104982175 | 104984698 |
| chr2 | 57938859 | 57941106 |
| chr15 | 98864856 | 98866586 |
| chr6 | 81752300 | 81754668 |
| chr4 | 15022685 | 15024200 |
| chr12 | 81412821 | 81416295 |
| chr14 | 62915881 | 62918435 |
| chr2 | 214022801 | 214025238 |
| chr12 | 4320275 | 4322188 |
| chr2 | 177992155 | 177995104 |
| chr5 | 150756654 | 150758318 |
| chr2 | 69828706 | 69831198 |
| chr15 | 20927061 | 20930163 |
| chr9 | 31263536 | 31267550 |
| chr7 | 87018036 | 87019273 |
| chr2 | 216500201 | 216501889 |
| chr4 | 157337783 | 157340245 |
| chr11 | 20467591 | 20469352 |
| chr13 | 106174891 | 106177040 |
| chr1 | 208360556 | 208361957 |
| chr2 | 63038152 | 63040888 |
| chr2 | 5641411 | 5643134 |
| chr2 | 70961082 | 70962649 |
| chr1_KI270765v1_a | 173136 | 175064 |
| chr2 | 184116709 | 184117926 |
| chr1 | 65148703 | 65150695 |
| chr13 | 35039419 | 35040498 |
| chr4 | 157138344 | 157145263 |
| chr5 | 27801677 | 27803211 |
| chr8 | 16273777 | 16276332 |
| chrY | 25382273 | 25383933 |
| chr14 | 52744176 | 52745471 |
| chr11 | 21577210 | 21579965 |
| chr5 | 160422045 | 160424013 |
| chr10 | 104255805 | 104257467 |
| chr18 | 3250274 | 3251331 |

| | | |
|---|---|---|
| chr14 | 106262857 | 106265428 |
| chr7 | 89068623 | 89069811 |
| chr15 | 74372751 | 74375259 |
| chr2 | 186763354 | 186765885 |
| chr3 | 262916 | 265305 |
| chr20 | 33211503 | 33212846 |
| chr10 | 128082263 | 128084230 |
| chr6 | 77241054 | 77243113 |
| chr4 | 94675967 | 94676969 |
| chr14 | 91327821 | 91329822 |
| chr2 | 185125469 | 185126739 |
| chr3 | 80770577 | 80772622 |
| chr9 | 64908473 | 64910498 |
| chr4 | 20518795 | 20522430 |
| chr5 | 94891054 | 94892877 |
| chr16 | 76215788 | 76217773 |
| chr22 | 10774466 | 10777628 |
| chr2 | 146732912 | 146734908 |
| chr6 | 81774886 | 81776571 |
| chr13 | 27369699 | 27372011 |
| chr20 | 29635055 | 29637196 |
| chr8 | 97920775 | 97922068 |
| chr5 | 130694553 | 130696693 |
| chr14 | 89417414 | 89420137 |
| chr7 | 83097126 | 83099244 |
| chr4 | 39061188 | 39062897 |
| chr1 | 96411167 | 96413240 |
| chr5 | 108177984 | 108180309 |
| chr13 | 97344778 | 97346621 |
| chr2 | 162544829 | 162546340 |
| chr6 | 104457150 | 104458854 |
| chr12 | 38334150 | 38336479 |
| chr10 | 16855023 | 16857419 |
| chr15 | 76794201 | 76795698 |
| chr7 | 58090596 | 58091849 |
| chr6 | 99397283 | 99398747 |
| chr22 | 10961755 | 10964929 |
| chr6 | 22322259 | 22325362 |
| chr10 | 61353005 | 61355133 |
| chr12 | 91326018 | 91328529 |
| chr2 | 41011242 | 41013799 |
| chr16 | 69422533 | 69423957 |
| chr2 | 214481382 | 214483769 |
| chr1 | 98258293 | 98261146 |
| chr10 | 61683504 | 61684682 |
| chr10 | 119987968 | 119990718 |
| chr4 | 90948012 | 90949864 |
| chr20 | 9359441 | 9360777 |
| chr1 | 115296959 | 115299527 |

| | | |
|---|---|---|
| chr2 | 146659930 | 146662323 |
| chr6 | 73755781 | 73757436 |
| chr14 | 79460239 | 79462392 |
| chr11 | 26417475 | 26419421 |
| chr19 | 10005167 | 10008234 |
| chr6 | 38989141 | 38990656 |
| chr12 | 38878348 | 38881352 |
| chr14 | 47406319 | 47408137 |
| chr2 | 50154216 | 50157097 |
| chr13 | 64492622 | 64494317 |
| chr3 | 137880334 | 137882662 |
| chr15 | 59689107 | 59690693 |
| chr16 | 72823699 | 72825161 |
| chr19 | 56836078 | 56837771 |
| chr9 | 104646717 | 104647775 |
| chr12 | 68031572 | 68033456 |
| chr10 | 44476096 | 44478143 |
| chr3 | 29425271 | 29426542 |
| chr1 | 218570888 | 218573479 |
| chr12 | 90104197 | 90106041 |
| chr14 | 30500296 | 30502931 |
| chr14 | 31340358 | 31342554 |
| chr12 | 75313281 | 75315226 |
| chr13 | 38767452 | 38770097 |
| chr5 | 27357590 | 27359842 |
| chrY | 56821760 | 56823581 |
| chr10 | 125854353 | 125856769 |
| chr1 | 190470394 | 190472522 |
| chr4 | 85864454 | 85867827 |
| chr14 | 95895950 | 95898254 |
| chr15 | 22291746 | 22293696 |
| chr17_KI270909v1_ | 313659 | 314886 |
| chr4 | 30000381 | 30003271 |
| chr10 | 58398001 | 58400328 |
| chr3 | 158484873 | 158486814 |
| chr14 | 55268466 | 55270598 |
| chr14_KI270846v1_ | 737888 | 739277 |
| chr10 | 115061839 | 115063426 |
| chr10 | 65787256 | 65791936 |
| chr15 | 37882054 | 37886760 |
| chr14 | 31451092 | 31452145 |
| chr13 | 106067959 | 106070233 |
| chr14 | 67241733 | 67243201 |
| chr3 | 81980339 | 81983295 |
| chr1 | 103741166 | 103742255 |
| chr14 | 37403626 | 37406159 |
| chr1 | 222929423 | 222930577 |
| chr1 | 94014835 | 94017029 |
| chr10 | 55279669 | 55281449 |

| | | |
|---|---|---|
| chr4 | 18837412 | 18840326 |
| chr20 | 38827867 | 38830256 |
| chr1 | 101550133 | 101551912 |
| chr10 | 115654091 | 115655852 |
| chr8 | 73008215 | 73009368 |
| chr1 | 211579708 | 211581577 |
| chr12 | 60608098 | 60609936 |
| chr6 | 133050417 | 133053203 |
| chr1 | 110354994 | 110357459 |
| chr5 | 78014999 | 78016737 |
| chr20 | 62894543 | 62896671 |
| chr5 | 101416059 | 101417060 |
| chr10 | 65193872 | 65197349 |
| chr6 | 22087682 | 22089523 |
| chr10 | 109032710 | 109034660 |
| chr2 | 50398149 | 50399920 |
| chr5 | 59006202 | 59008005 |
| chr2 | 79915647 | 79916855 |
| chr1_KI270765v1_a | 98320 | 102360 |
| chr6 | 56513553 | 56515457 |
| chr12 | 32853794 | 32855882 |
| chr13 | 72186300 | 72187983 |
| chr1 | 96506797 | 96509260 |
| chr5 | 112109444 | 112110835 |
| chr20 | 3633556 | 3636175 |
| chr10 | 43773351 | 43774756 |
| chr14 | 102226565 | 102228753 |
| chr14 | 31106009 | 31108125 |
| chr8 | 126866237 | 126867827 |
| chr8 | 15983133 | 15985578 |
| chr9 | 13365823 | 13367786 |
| chr14 | 41453313 | 41455645 |
| chr10 | 47369117 | 47371710 |
| chr1 | 104534911 | 104539018 |
| chr3 | 166104860 | 166106701 |
| chr10 | 58175749 | 58178750 |
| chr6 | 50466765 | 50469059 |
| chr3 | 77556111 | 77557860 |
| chr2 | 194339838 | 194342922 |
| chr11 | 17276548 | 17277939 |
| chr10 | 115930572 | 115932165 |
| chr1 | 187885828 | 187887241 |
| chr5 | 93129437 | 93131446 |
| chr1 | 94060132 | 94062716 |
| chr2 | 208980885 | 208983403 |
| chr10 | 94057721 | 94059783 |
| chr2 | 111306400 | 111307750 |
| chr13 | 66101837 | 66103908 |
| chr14 | 50875458 | 50877735 |

| | | |
|---|---|---|
| chr10 | 73693796 | 73696312 |
| chr1 | 98891791 | 98894688 |
| chr22_KI270734v1_ | 78076 | 80045 |
| chr6 | 65236615 | 65240233 |
| chr1 | 190226588 | 190228335 |
| chr6 | 70686053 | 70687788 |
| chr2 | 97568668 | 97570539 |
| chr2 | 169394175 | 169396401 |
| chr2 | 186939383 | 186941185 |
| chr1 | 218485700 | 218488695 |
| chr2 | 185113991 | 185116328 |
| chr2 | 69558535 | 69560513 |
| chr12 | 43451866 | 43454502 |
| chr2 | 162306387 | 162307676 |
| chr2 | 158947786 | 158950383 |
| chr6 | 91158249 | 91159781 |
| chr1 | 101992361 | 101995461 |
| chr5 | 94240192 | 94242702 |
| chr2 | 84693397 | 84695539 |
| chr1 | 98156507 | 98159342 |
| chr4 | 115769104 | 115770945 |
| chr1 | 80055854 | 80057077 |
| chr12 | 97536310 | 97538875 |
| chr13 | 40858098 | 40860054 |
| chr2 | 97643682 | 97646439 |
| chr12 | 77292558 | 77293976 |
| chr15 | 24379159 | 24380503 |
| chr11 | 85627943 | 85629305 |
| chr14 | 38958381 | 38960257 |
| chr17 | 47648367 | 47649958 |
| chr10 | 73718068 | 73719853 |
| chr10 | 55040994 | 55042704 |
| chr15 | 75275530 | 75277424 |
| chr14 | 85359074 | 85362137 |
| chr1 | 117524841 | 117527658 |
| chr14 | 27869055 | 27870816 |
| chr15 | 48856438 | 48858250 |
| chr22 | 15921218 | 15924663 |
| chr4 | 34143374 | 34144672 |
| chr7 | 79405162 | 79408496 |
| chr7 | 53657804 | 53660085 |
| chr14 | 38896735 | 38898370 |
| chr2 | 51364498 | 51366496 |
| chr7 | 81573342 | 81575935 |
| chr6 | 363316 | 364759 |
| chr13 | 80123854 | 80126460 |
| chr15 | 92881481 | 92882931 |
| chr10 | 112819510 | 112822859 |
| chr2 | 57371230 | 57380533 |

| | | |
|---|---|---|
| chr9 | 62290229 | 62292024 |
| chr17 | 21282539 | 21285620 |
| chr18 | 4461349 | 4463375 |
| chr2 | 186855612 | 186857893 |
| chr5 | 18852725 | 18854151 |
| chr3 | 70516340 | 70518761 |
| chr4 | 93019686 | 93021831 |
| chr11 | 28025035 | 28027168 |
| chr3 | 74791724 | 74793164 |
| chr8 | 112562076 | 112564047 |
| chr8 | 54046072 | 54048663 |
| chr14 | 88020670 | 88022904 |
| chr2 | 168458145 | 168461582 |
| chr22 | 47192414 | 47194278 |
| chr11 | 23204178 | 23206175 |
| chr4 | 156658169 | 156661106 |
| chr8 | 103644300 | 103645973 |
| chr14 | 67527965 | 67530311 |
| chr5 | 59431942 | 59434371 |
| chr1 | 113613306 | 113615643 |
| chr14 | 41220041 | 41221962 |
| chr4 | 156676593 | 156677830 |
| chr6 | 70560424 | 70561903 |
| chr5 | 27482611 | 27484977 |
| chr13 | 62017490 | 62019047 |
| chr3 | 77779967 | 77781809 |
| chr20 | 29665951 | 29668154 |
| chr1 | 94478380 | 94480372 |
| chr8 | 120072728 | 120075364 |
| chr1 | 174999242 | 175001012 |
| chr14 | 88470468 | 88472261 |
| chr20 | 38507425 | 38508982 |
| chr2 | 144432489 | 144434245 |
| chr16 | 76166223 | 76167843 |
| chr10 | 102366956 | 102368744 |
| chr9 | 121760997 | 121763999 |
| chr10 | 81347365 | 81349316 |
| chr2 | 227045726 | 227047703 |
| chr7 | 79546864 | 79548378 |
| chr3 | 57240797 | 57243875 |
| chr7 | 95135168 | 95137033 |
| chr1 | 16062491 | 16064131 |
| chr16 | 76680956 | 76682858 |
| chr14 | 95543728 | 95546011 |
| chr14 | 24822554 | 24824626 |
| chr10 | 80362476 | 80364755 |
| chr14 | 46308215 | 46310161 |
| chr4 | 132996754 | 132998056 |
| chr12 | 65432116 | 65434916 |

| | | |
|---|---|---|
| chr1 | 111713923 | 111715962 |
| chr1 | 103540225 | 103541811 |
| chr3 | 75229802 | 75231996 |
| chr10 | 82986314 | 82987871 |
| chr3 | 106397448 | 106399483 |
| chr18 | 70128676 | 70129704 |
| chr20 | 30917048 | 30918851 |
| chr8 | 97253129 | 97255730 |
| chr14 | 27812784 | 27815456 |
| chr12 | 64373469 | 64374961 |
| chr6 | 123153213 | 123154252 |
| chr11 | 105973709 | 105976635 |
| chr6 | 72530165 | 72531824 |
| chr11 | 56469664 | 56472527 |
| chr1 | 101849433 | 101851623 |
| chr5 | 58989048 | 58991156 |
| chr10 | 38648242 | 38649265 |
| chr16 | 72430031 | 72431617 |
| chr5 | 17526369 | 17528963 |
| chr20 | 49096017 | 49097406 |
| chr7 | 103282664 | 103283685 |
| chr1 | 80446139 | 80448237 |
| chr1 | 191794108 | 191795389 |
| chr10 | 62856654 | 62858682 |
| chr9 | 5339176 | 5341567 |
| chr2 | 203223399 | 203225753 |
| chr12 | 89879315 | 89880492 |
| chr20 | 35741599 | 35744288 |
| chr6 | 102062312 | 102064068 |
| chr7 | 86834228 | 86835342 |
| chr20 | 38947196 | 38949327 |
| chr7 | 81558129 | 81562039 |
| chr3 | 136196055 | 136197722 |
| chr2 | 53805515 | 53807138 |
| chr3 | 178757662 | 178759200 |
| chr6 | 125678537 | 125680926 |
| chr7 | 109293100 | 109294876 |
| chr1 | 16580306 | 16581426 |
| chr2 | 187480148 | 187482471 |
| chr6 | 99897188 | 99899032 |
| chr12 | 100472922 | 100475050 |
| chr14 | 47014968 | 47018691 |
| chr2 | 182005976 | 182007974 |
| chr1 | 80350152 | 80352029 |
| chr4 | 82900144 | 82902109 |
| chr11 | 28546289 | 28549047 |
| chr18 | 67217013 | 67219231 |
| chr15 | 69218180 | 69219735 |
| chr2 | 82992562 | 82994453 |

| | | |
|---|---|---|
| chr6 | 75444420 | 75445609 |
| chr10 | 110939667 | 110941364 |
| chr21 | 20850282 | 20851955 |
| chr6 | 72873346 | 72874965 |
| chr13 | 71637637 | 71639702 |
| chr7 | 79057217 | 79058951 |
| chr1 | 84782082 | 84783744 |
| chr1 | 86204750 | 86206281 |
| chr5 | 34171116 | 34172263 |
| chr7 | 88617603 | 88619856 |
| chr5 | 29749968 | 29752558 |
| chr17_KI270857v1_ | 2171072 | 2173236 |
| chr14 | 40943837 | 40945521 |
| chr6 | 91514852 | 91517422 |
| chr5_KI270897v1_a | 130867 | 132535 |
| chr2 | 187688323 | 187689774 |
| chr6 | 15320089 | 15321786 |
| chr2 | 39995096 | 39996880 |
| chr20 | 55548084 | 55549483 |
| chr2 | 204657133 | 204659308 |
| chr7 | 76903165 | 76904819 |
| chr9 | 64598198 | 64599487 |
| chr19 | 48993338 | 48994711 |
| chr1 | 223200232 | 223203104 |
| chr4 | 156472068 | 156474335 |
| chr12 | 85876603 | 85879560 |
| chr10 | 57997228 | 57999317 |
| chr3 | 81301667 | 81303507 |
| chr10 | 62647175 | 62648888 |
| chr13 | 64064176 | 64065755 |
| chr10 | 51649550 | 51651037 |
| chr15 | 68060583 | 68061967 |
| chr8 | 139513957 | 139515980 |
| chr2 | 42047723 | 42050595 |
| chr2 | 79428155 | 79430396 |
| chr2 | 103133700 | 103136281 |
| chr2 | 34526778 | 34528937 |
| chr1 | 162153414 | 162155120 |
| chr5 | 28205031 | 28208569 |
| chr10 | 43352976 | 43354808 |
| chr10 | 56119274 | 56121491 |
| chr4 | 149774903 | 149776592 |
| chr8 | 93192340 | 93194630 |
| chr4 | 3896723 | 3897739 |
| chr12 | 82063239 | 82064288 |
| chr2 | 68507081 | 68508558 |
| chr10 | 125123841 | 125126878 |
| chr6 | 129718439 | 129719444 |
| chr1 | 102690037 | 102691512 |

| | | |
|---|---|---|
| chr10 | 46181152 | 46183039 |
| chr5 | 39274043 | 39275819 |
| chr2 | 166732702 | 166735431 |
| chr10 | 55342885 | 55344137 |
| chr20 | 5595356 | 5597479 |
| chr13 | 84546483 | 84548469 |
| chr14 | 37427603 | 37429223 |
| chr8_KI270813v1_a | 256628 | 258285 |
| chr1 | 71451436 | 71453226 |
| chr14 | 33243495 | 33244940 |
| chr16 | 61844628 | 61846297 |
| chr12 | 65380838 | 65384052 |
| chr14 | 39737226 | 39738919 |
| chr20 | 9468878 | 9471089 |
| chr6 | 129826578 | 129828425 |
| chr13 | 72150563 | 72151823 |
| chr8 | 17001020 | 17003028 |
| chr1 | 74235928 | 74238380 |
| chr2 | 63919530 | 63922322 |
| chr6 | 70602194 | 70603541 |
| chr13 | 55259220 | 55261732 |
| chr20 | 44218403 | 44219644 |
| chr2 | 197779769 | 197782207 |
| chr1 | 94901401 | 94903818 |
| chr11 | 21470641 | 21473058 |
| chr3 | 84300033 | 84301574 |
| chr2 | 50024730 | 50027293 |
| chr2 | 73784233 | 73786015 |
| chr6 | 97856422 | 97858205 |
| chr20 | 57260495 | 57261657 |
| chr12 | 85467866 | 85469234 |
| chr2 | 41703651 | 41705650 |
| chr4 | 91595945 | 91597499 |
| chr2 | 105642196 | 105644678 |
| chr8 | 16831203 | 16833896 |
| chr1 | 106296199 | 106298643 |
| chr10 | 115738283 | 115740655 |
| chr4 | 136144423 | 136146441 |
| chr9 | 103299788 | 103302024 |
| chr7 | 49870828 | 49872784 |
| chr1 | 192804516 | 192807086 |
| chr8 | 94081212 | 94083745 |
| chr10 | 116422520 | 116424518 |
| chr1 | 90384451 | 90386329 |
| chr4 | 114381350 | 114383025 |
| chr15 | 47597939 | 47599271 |
| chr2 | 181781328 | 181782816 |
| chr14 | 38841207 | 38843217 |
| chr10 | 59625319 | 59627525 |

| | | |
|---|---|---|
| chr14 | 36215661 | 36217442 |
| chr7 | 85308068 | 85309397 |
| chr10 | 55236276 | 55239058 |
| chr5 | 171386489 | 171388982 |
| chr19 | 7057211 | 7059416 |
| chr2 | 67612288 | 67614651 |
| chr1 | 197384066 | 197386326 |
| chr4 | 31734487 | 31736660 |
| chr14 | 29539837 | 29542114 |
| chr14 | 39514852 | 39516347 |
| chr10 | 115160739 | 115162569 |
| chr5 | 111225848 | 111228933 |
| chr14 | 58521289 | 58523294 |
| chr2 | 230442687 | 230444483 |
| chr2 | 95519447 | 95522331 |
| chr10 | 35218180 | 35220155 |
| chr5_KI270791v1_a | 179738 | 181434 |
| chr14 | 43985126 | 43988890 |
| chr4 | 26170208 | 26172833 |
| chr20 | 7916425 | 7918742 |
| chr6 | 79230723 | 79233233 |
| chr6 | 126814689 | 126816549 |
| chr2 | 84860458 | 84862204 |
| chr6 | 147566945 | 147568401 |
| chr14 | 100352383 | 100354974 |
| chr11 | 81932103 | 81933230 |
| chr6 | 86264509 | 86266301 |
| chr15 | 21292958 | 21294209 |
| chr11 | 108287113 | 108290232 |
| chr21 | 42974036 | 42976108 |
| chr17_KI270857v1_ | 548552 | 549764 |
| chr10 | 57809306 | 57811924 |
| chr14 | 87128725 | 87129982 |
| chr4 | 34064200 | 34066450 |
| chr12 | 85043380 | 85048105 |
| chr20 | 14710479 | 14712764 |
| chr5 | 141848723 | 141849894 |
| chr1 | 103102440 | 103103961 |
| chr2 | 231000142 | 231002477 |
| chr11 | 118392157 | 118393162 |
| chr2 | 67489391 | 67491143 |
| chr4 | 69572138 | 69573916 |
| chr9 | 92276928 | 92278701 |
| chr2 | 131642438 | 131644995 |
| chr12 | 85184692 | 85186903 |
| chr6 | 51470984 | 51473242 |
| chr11 | 23827836 | 23830846 |
| chr9 | 63641574 | 63644775 |
| chr2 | 166427979 | 166430222 |

| | | |
|---|---|---|
| chr2 | 192968403 | 192970731 |
| chr21 | 10398665 | 10400451 |
| chr14 | 19351786 | 19353963 |
| chr21 | 10108843 | 10110351 |
| chr10 | 67524257 | 67525431 |
| chr10 | 54617624 | 54619866 |
| chr1 | 117331004 | 117332439 |
| chr2 | 189221873 | 189223392 |
| chr3 | 22758260 | 22760082 |
| chr14 | 102599017 | 102600176 |
| chr8 | 105344344 | 105346008 |
| chr1 | 104994002 | 104995571 |
| chr7 | 95915015 | 95918077 |
| chr12 | 24595865 | 24598319 |
| chr8 | 109292154 | 109293633 |
| chr17 | 70743221 | 70745776 |
| chr10 | 37121408 | 37123219 |
| chr6 | 73075841 | 73077912 |
| chr20 | 50935766 | 50938791 |
| chr2 | 211484631 | 211486661 |
| chr3 | 175710560 | 175713592 |
| chr2 | 222521047 | 222524278 |
| chr14 | 27010559 | 27012522 |
| chrY | 11676184 | 11678709 |
| chr2 | 50917514 | 50920655 |
| chr7 | 94667583 | 94669010 |
| chr8 | 78228960 | 78231559 |
| chr3 | 60859736 | 60860872 |
| chr7 | 81686640 | 81688121 |
| chr3_KI270784v1_a | -421 | 2251 |
| chr2 | 59723949 | 59726328 |
| chr20 | 57357358 | 57359748 |
| chr17 | 60598509 | 60601922 |
| chr1 | 116020578 | 116023104 |
| chr6 | 27250095 | 27252286 |
| chr2 | 88170419 | 88172751 |
| chr1 | 194926720 | 194929128 |
| chr7 | 84649604 | 84652221 |
| chr6 | 22911106 | 22913091 |
| chr10 | 56156846 | 56158284 |
| chr5 | 179975743 | 179977509 |
| chr1 | 104525702 | 104529211 |
| chr1 | 192355067 | 192357158 |
| chrY | 10102436 | 10103686 |
| chr6 | 135667654 | 135670251 |
| chr9 | 73181255 | 73184596 |
| chr20 | 55262282 | 55264601 |
| chr7 | 116590670 | 116593068 |
| chr3 | 167578914 | 167581523 |

| | | |
|---|---|---|
| chr2 | 48419582 | 48421574 |
| chr2 | 58057205 | 58058580 |
| chr3 | 102880058 | 102882691 |
| chr12 | 78196217 | 78197831 |
| chr2 | 179649893 | 179652826 |
| chr1 | 191553099 | 191554899 |
| chr10 | 37114540 | 37116852 |
| chr11 | 24906177 | 24908365 |
| chr11 | 31105285 | 31107715 |
| chr1 | 194432853 | 194434757 |
| chr4 | 34081014 | 34083245 |
| chr14 | 25506547 | 25508242 |
| chr7 | 79462303 | 79464182 |
| chr14 | 32809467 | 32811320 |
| chr21 | 16575654 | 16577260 |
| chr12 | 99329416 | 99331983 |
| chr14 | 27831623 | 27833404 |
| chr2 | 184765447 | 184766765 |
| chr2 | 45567255 | 45569596 |
| chr4 | 26936967 | 26938462 |
| chr6 | 45332874 | 45335460 |
| chr14 | 64033990 | 64035836 |
| chr1 | 99574804 | 99576520 |
| chr4 | 121666718 | 121668889 |
| chr13 | 87192841 | 87193989 |
| chr2 | 91655358 | 91656776 |
| chr16 | 77282629 | 77284279 |
| chr1 | 98070471 | 98072426 |
| chr2 | 182145615 | 182148532 |
| chr2 | 226422618 | 226424455 |
| chr1 | 5671106 | 5673151 |
| chr2 | 91437693 | 91438904 |
| chr2 | 68040101 | 68043048 |
| chr13 | 77663654 | 77666317 |
| chr6 | 8661185 | 8663545 |
| chr6 | 73855364 | 73856749 |
| chr7 | 78940704 | 78942603 |
| chr1 | 83867582 | 83871001 |
| chr2 | 103529203 | 103530732 |
| chr1 | 106143088 | 106144678 |
| chr12 | 41436140 | 41437975 |
| chr2 | 199738789 | 199741331 |
| chr8 | 114104407 | 114106497 |
| chr2 | 67259176 | 67261214 |
| chr15 | 55617780 | 55620521 |
| chr6 | 169196349 | 169197738 |
| chr21 | 6372498 | 6373683 |
| chr16_KI270728v1_ | 223415 | 225825 |
| chr1 | 190282409 | 190284627 |

| | | |
|---|---|---|
| chr14 | 86719264 | 86721937 |
| chr1 | 92244094 | 92246495 |
| chr6 | 123439738 | 123441565 |
| chr15 | 26715852 | 26716905 |
| chr7 | 94151393 | 94154821 |
| chr2 | 172669922 | 172672293 |
| chr5 | 124347491 | 124349362 |
| chr5 | 39453297 | 39454982 |
| chr14 | 19267067 | 19269813 |
| chr1 | 186970548 | 186973086 |
| chr12 | 78891560 | 78893747 |
| chr6 | 149263263 | 149265143 |
| chr10 | 58343371 | 58346217 |
| chr9 | 85787170 | 85789505 |
| chr5_KI270897v1_a | 199282 | 201663 |
| chr12 | 26936958 | 26939838 |
| chr5 | 131643097 | 131645203 |
| chr1_KI270765v1_a | 143416 | 145779 |
| chr17 | 52167152 | 52169117 |
| chr4 | 33917235 | 33920103 |
| chr10 | 37855130 | 37857801 |
| chr4 | 34399244 | 34401116 |
| chr12 | 87986096 | 87987502 |
| chr2 | 154441989 | 154444299 |
| chr6 | 88697858 | 88699402 |
| chr19 | 42925452 | 42927347 |
| chr14 | 27785951 | 27787789 |
| chr14 | 74261361 | 74263138 |
| chr3 | 85250020 | 85251379 |
| chr1 | 167935786 | 167937088 |
| chr1 | 70017240 | 70019335 |
| chr1 | 101847618 | 101849310 |
| chr1 | 145596958 | 145597965 |
| chr8 | 110839544 | 110842316 |
| chr5 | 92749475 | 92751244 |
| chr4 | 25019650 | 25022024 |
| chr8 | 137735322 | 137736751 |
| chr12 | 72305294 | 72306621 |
| chr13 | 103078180 | 103080074 |
| chr2 | 61350301 | 61352189 |
| chr12 | 78774540 | 78775929 |
| chr21 | 16180186 | 16182071 |
| chr1 | 195824515 | 195826553 |
| chr13 | 63174672 | 63176561 |
| chr16_KI270728v1_ | 20139 | 22990 |
| chr14 | 59091232 | 59093410 |
| chr9 | 68282004 | 68283541 |
| chr1 | 94615879 | 94617507 |
| chr7 | 126881419 | 126883647 |

| | | |
|---|---|---|
| chr8 | 96234773 | 96236063 |
| chr13 | 80048502 | 80051398 |
| chr15 | 20209257 | 20211766 |
| chr10 | 97329761 | 97330787 |
| chr6 | 90667851 | 90669717 |
| chr10 | 43122547 | 43124489 |
| chr6 | 64351244 | 64354780 |
| chr2 | 102651572 | 102654370 |
| chr2 | 144797973 | 144799702 |
| chr1 | 237829503 | 237831084 |
| chr2 | 184564102 | 184566532 |
| chr1 | 121569799 | 121572406 |
| chr2 | 204947793 | 204949296 |
| chr7 | 83254968 | 83257025 |
| chr8 | 115577595 | 115578665 |
| chr6 | 97307354 | 97308666 |
| chr10 | 46619753 | 46622240 |
| chr17 | 60729335 | 60730658 |
| chr5 | 177763668 | 177767211 |
| chr14 | 41505778 | 41507249 |
| chr2 | 95448222 | 95450402 |
| chr1 | 143254624 | 143255840 |
| chr2 | 48200149 | 48201363 |
| chr1 | 56517251 | 56520024 |
| chr2 | 225996950 | 226000551 |
| chr8 | 77510574 | 77513282 |
| chr1 | 149343223 | 149345659 |
| chr2 | 187071734 | 187073360 |
| chr2 | 58363477 | 58365400 |
| chr1 | 100705933 | 100708066 |
| chr4 | 24007351 | 24009712 |
| chr9 | 65223525 | 65225376 |
| chr2 | 82464038 | 82466301 |
| chr11 | 101913191 | 101915454 |
| chr2 | 195025792 | 195027397 |
| chr1 | 102324059 | 102325982 |
| chr1 | 71001151 | 71004060 |
| chr12 | 77999805 | 78000815 |
| chr2 | 157309622 | 157312048 |
| chr2 | 95537864 | 95540232 |
| chr14 | 47228938 | 47231130 |
| chr10 | 34621370 | 34623774 |
| chr13 | 65575270 | 65576621 |
| chr2 | 17778140 | 17779959 |
| chr1 | 186949766 | 186951433 |
| chr5 | 30417405 | 30419415 |
| chr20 | 3156901 | 3159600 |
| chr9 | 73245608 | 73247507 |
| chr7 | 84414721 | 84415841 |

| | | |
|---|---|---|
| chr2 | 187579285 | 187581287 |
| chr2 | 67363989 | 67365540 |
| chr14 | 25485448 | 25487589 |
| chr13 | 82747527 | 82749273 |
| chr7 | 83453060 | 83455662 |
| chr4 | 156580509 | 156582794 |
| chr12 | 79719295 | 79721125 |
| chr8 | 136253156 | 136254483 |
| chr2 | 195468658 | 195471024 |
| chr12 | 70666831 | 70668452 |
| chr11 | 55271779 | 55273180 |
| chr1 | 245042219 | 245043463 |
| chr10 | 88985902 | 88987499 |
| chr13 | 106439463 | 106441505 |
| chr6 | 55884121 | 55887132 |
| chr1 | 188494304 | 188496359 |
| chr11 | 106048879 | 106050407 |
| chr2 | 180537604 | 180540701 |
| chr6 | 64429721 | 64432911 |
| chr17 | 68250850 | 68252956 |
| chr10 | 64353244 | 64355459 |
| chr9 | 28791218 | 28793919 |
| chr10 | 46275944 | 46277449 |
| chr2 | 64128561 | 64130475 |
| chr2 | 112365763 | 112367729 |
| chr2 | 34400054 | 34404461 |
| chr7 | 85572691 | 85576369 |
| chr2 | 70225530 | 70227587 |
| chr4 | 97028301 | 97031412 |
| chr7 | 65080396 | 65081699 |
| chr2 | 69825453 | 69827833 |
| chr1 | 107280720 | 107282946 |
| chr2 | 185170528 | 185173594 |
| chr2 | 72395961 | 72398214 |
| chr4 | 178678398 | 178679835 |
| chr1 | 66812706 | 66814636 |
| chr20 | 29430869 | 29434418 |
| chr14 | 30272847 | 30275437 |
| chr15 | 37903427 | 37905345 |
| chr2 | 161963473 | 161966825 |
| chr2 | 231832794 | 231833795 |
| chr7 | 79356344 | 79357818 |
| chr7 | 55591283 | 55594855 |
| chr4 | 33564512 | 33567976 |
| chr2 | 18969763 | 18972600 |
| chr11 | 23122320 | 23125591 |
| chr2 | 95940257 | 95943520 |
| chr22 | 11053204 | 11056705 |
| chr1 | 198887736 | 198890959 |

| | | |
|---|---|---|
| chr11 | 96059903 | 96063194 |
| chr12 | 50985298 | 50988500 |
| chr13 | 80117425 | 80120550 |
| chr2 | 64522470 | 64525735 |
| chr5_KI270897v1_a | 698465 | 701998 |
| chr2 | 178702133 | 178705053 |
| chr13 | 67063382 | 67067333 |
| chr9 | 63827853 | 63830032 |
| chr1 | 236220427 | 236222509 |
| chr1 | 103468380 | 103471224 |
| chr13 | 71705418 | 71709725 |
| chr1 | 74771196 | 74775060 |
| chr9 | 31849518 | 31852244 |
| chr14 | 19322476 | 19325649 |
| chr1 | 100248806 | 100251969 |
| chr12 | 77993728 | 77997101 |
| chr5 | 89566009 | 89568186 |
| chr14 | 100957093 | 100960294 |
| chr21 | 7465301 | 7468966 |
| chr7 | 84161198 | 84165300 |
| chr19 | 42784979 | 42787077 |
| chr10 | 68336438 | 68339074 |
| chr9 | 65800691 | 65803302 |
| chrY | 6253944 | 6257911 |
| chr11 | 81917211 | 81920871 |
| chr7 | 82981156 | 82985164 |
| chr5 | 110568698 | 110570845 |
| chr7 | 87156016 | 87158535 |
| chr4 | 28436819 | 28439482 |
| chr6 | 79515978 | 79518717 |
| chr12 | 25208415 | 25212581 |
| chr10 | 37161341 | 37163604 |
| chr15 | 20367951 | 20372924 |
| chr1 | 90003384 | 90005936 |
| chr14 | 19713376 | 19717038 |
| chr13 | 63911032 | 63914709 |
| chr14 | 19862024 | 19865686 |
| chr7 | 85025845 | 85031680 |
| chr3 | 174438426 | 174442765 |
| chr7 | 100333375 | 100336550 |
| chr10 | 90712217 | 90716133 |
| chr15_KI270850v1_ | 301185 | 304951 |
| chr6 | 26029784 | 26034182 |
| chr10 | 67658472 | 67660794 |
| chr9 | 66328241 | 66331877 |
| chr2 | 86439152 | 86443747 |
| chr1 | 103687396 | 103689834 |
| chr3 | 147613829 | 147618530 |
| chr10 | 63621668 | 63625141 |

| | | |
|---|---|---|
| chr7 | 81107969 | 81110562 |
| chr20 | 30812334 | 30814908 |
| chr2 | 90390077 | 90393191 |
| chr13 | 80344093 | 80346686 |
| chr2 | 57450080 | 57456769 |
| chr4 | 45023659 | 45026545 |
| chr2 | 72544191 | 72546613 |
| chr19 | 101023 | 104452 |
| chr22 | 15874494 | 15879060 |
| chr15 | 20685017 | 20689260 |
| chr6 | 79082044 | 79085571 |
| chr4 | 33352845 | 33356176 |
| chr14 | 41169258 | 41173492 |
| chr12 | 78896150 | 78900440 |
| chr1 | 45686084 | 45688690 |
| chr20 | 45971147 | 45974957 |
| chr2 | 89799501 | 89808284 |
| chr2 | 97163894 | 97168377 |
| chr20 | 30187913 | 30191980 |
| chrY | 56870899 | 56874342 |
| chr2 | 97585754 | 97588119 |
| chr14 | 47136498 | 47140507 |
| chr14 | 42941907 | 42944508 |
| chr14 | 63832061 | 63834513 |
| chr1 | 116910653 | 116913129 |
| chr11 | 23422982 | 23425921 |
| chr7 | 62347286 | 62356662 |
| chr4 | 34430431 | 34433938 |
| chr2 | 44476692 | 44479763 |
| chr6 | 52284534 | 52287620 |
| chr7 | 80678587 | 80682796 |
| chr2 | 162240168 | 162247789 |
| chr15 | 77419933 | 77423297 |
| chr1 | 148668837 | 148671519 |
| chr2 | 225406040 | 225408242 |
| chr10 | 73494363 | 73498645 |
| chr13 | 23569387 | 23572373 |
| chr10 | 88990875 | 88995088 |
| chr2 | 89787659 | 89799207 |
| chr3 | 85216073 | 85218625 |
| chr5 | 121160274 | 121163998 |
| chr2 | 176634454 | 176637304 |
| chr6_GL000256v2_a | 3491875 | 3495582 |
| chr1 | 103291649 | 103295105 |
| chr8 | 76579737 | 76583780 |
| chr4 | 156975337 | 156979054 |
| chr10 | 64774345 | 64778065 |
| chr7 | 80742345 | 80746289 |
| chr7 | 79089728 | 79092715 |

| | | |
|---|---|---|
| chr13 | 57169102 | 57171259 |
| chr14 | 47505298 | 47507535 |
| chr12 | 72628197 | 72632332 |
| chr1 | 103725500 | 103729332 |
| chr6 | 20929881 | 20934887 |
| chr3 | 77034966 | 77039712 |
| chr1 | 16723725 | 16727084 |
| chr14 | 96549941 | 96552871 |
| chr13 | 67236451 | 67239782 |
| chr14 | 42488217 | 42491250 |
| chr2 | 66108032 | 66111798 |
| chr1 | 189634502 | 189637462 |
| chr1 | 149840240 | 149843399 |
| chr10 | 57045829 | 57050582 |
| chr5_KI270897v1_a | 27607 | 30436 |
| chr14 | 40792298 | 40794814 |
| chr11 | 23275460 | 23279123 |
| chr2 | 112375776 | 112378870 |
| chr14 | 38405775 | 38408864 |
| chr14 | 59187684 | 59190644 |
| chr16 | 77095295 | 77097914 |
| chr1 | 102168831 | 102172374 |
| chr6 | 133098149 | 133102183 |
| chr12 | 84527216 | 84530578 |
| chr14 | 88660964 | 88663786 |
| chr7 | 84425909 | 84431694 |
| chr5 | 124654687 | 124656747 |
| chr7 | 79270133 | 79273707 |
| chr6 | 133952361 | 133955242 |
| chr9 | 138301302 | 138304184 |
| chr9 | 66305335 | 66308043 |
| chr1 | 96774470 | 96778244 |
| chr7 | 94472913 | 94474952 |
| chr2 | 169943446 | 169947380 |
| chrUn_GL000218v1 | 59206 | 61853 |
| chr14 | 85846080 | 85850616 |
| chr2 | 186529286 | 186532302 |
| chr1 | 103785070 | 103789414 |
| chr14 | 46728308 | 46733858 |
| chr2 | 183038427 | 183040729 |
| chr2 | 190925088 | 190927281 |
| chr9 | 82911661 | 82913791 |
| chr20 | 9109760 | 9112125 |
| chr7 | 78486125 | 78490531 |
| chr10 | 58367653 | 58370125 |
| chr1 | 244840280 | 244843432 |
| chr14 | 19278870 | 19283804 |
| chr10 | 73675185 | 73677750 |
| chr5 | 27218803 | 27221613 |

| | | |
|---|---|---|
| chr8 | 82665019 | 82668058 |
| chr5 | 134924615 | 134928794 |
| chr1 | 192609430 | 192611677 |
| chr14 | 25802915 | 25805871 |
| chr12 | 28373768 | 28379208 |
| chr1 | 103618332 | 103621999 |
| chr2 | 50839520 | 50842767 |
| chr13 | 113066460 | 113070100 |
| chr10 | 74981886 | 74985318 |
| chr7 | 60910109 | 60919318 |
| chr17 | 22148845 | 22152687 |
| chr20 | 30361833 | 30364821 |
| chr7 | 82436173 | 82442640 |
| chr15_KI270905v1_ | 651047 | 654001 |
| chr7 | 67087279 | 67089727 |
| chr1 | 218525027 | 218528272 |
| chr15 | 20379580 | 20383572 |
| chr16 | 71053551 | 71057202 |
| chr14 | 33989380 | 33991577 |
| chr10 | 87328382 | 87330960 |
| chr1 | 200409555 | 200411745 |
| chr2 | 162310873 | 162314628 |
| chr14 | 46316959 | 46319284 |
| chr6 | 78667281 | 78670825 |
| chr10 | 52516753 | 52519298 |
| chr10 | 90188551 | 90191843 |
| chr2 | 87114752 | 87118973 |
| chr7 | 82725275 | 82729329 |
| chr14 | 48209817 | 48212627 |
| chr22 | 16262248 | 16266778 |
| chr14 | 61750263 | 61752904 |
| chr10 | 42099373 | 42102147 |
| chr8 | 115646134 | 115649022 |
| chr2 | 186678622 | 186681466 |
| chr8 | 115446933 | 115451017 |
| chr6 | 95605818 | 95610081 |
| chr8 | 106735398 | 106737429 |
| chr7 | 81795561 | 81799285 |
| chr21 | 10062311 | 10066677 |
| chr14 | 26436329 | 26441028 |
| chr2 | 186665449 | 186669633 |
| chr12 | 90666912 | 90669853 |
| chr5_KI270897v1_a | 81928 | 86128 |
| chr10 | 58266593 | 58271146 |
| chr2 | 200524952 | 200527752 |
| chr1 | 103198018 | 103201724 |
| chr9 | 41184477 | 41187411 |
| chr2 | 188971377 | 188983034 |
| chr14 | 19930611 | 19933974 |

| | | |
|---|---|---|
| chr15_KI270852v1_ | 27715 | 31261 |
| chr4 | 176780537 | 176783119 |
| chr14 | 64483677 | 64486497 |
| chr1 | 206185793 | 206189855 |
| chr16_KI270728v1_ | 1580613 | 1584352 |
| chr14 | 19717142 | 19720392 |
| chr7 | 84039161 | 84043019 |
| chr14 | 86935305 | 86938310 |
| chr14 | 19950572 | 19955086 |
| chr7 | 95058861 | 95061388 |
| chr8 | 12581233 | 12583486 |
| chr2 | 164835817 | 164839747 |
| chr12 | 84629539 | 84633573 |
| chr15 | 20394250 | 20399848 |
| chr13 | 18172782 | 18180434 |
| chr4 | 114889655 | 114892181 |
| chr1 | 104263193 | 104265399 |
| chr7 | 19093494 | 19098324 |
| chr1 | 218656944 | 218662484 |
| chr1 | 95720180 | 95723273 |
| chr10 | 56542261 | 56546478 |
| chr2 | 211743265 | 211746144 |
| chr1 | 97304967 | 97309369 |
| chr1_KI270765v1_a | 178691 | 182535 |
| chr4 | 126854370 | 126856958 |
| chr12 | 84592939 | 84598230 |
| chr1 | 218654789 | 218656909 |
| chr10 | 4671811 | 4674457 |
| chr12 | 68318209 | 68322616 |
| chr13 | 80549606 | 80552719 |
| chr15_KI270852v1_ | 12003 | 16453 |
| chr1_KI270765v1_a | 170728 | 172853 |
| chr2 | 213313167 | 213315561 |
| chr1 | 119136167 | 119141401 |
| chr1 | 218811474 | 218814247 |
| chr2 | 78866413 | 78869277 |
| chr1 | 102955469 | 102962064 |
| chr2 | 58045291 | 58048625 |
| chr6 | 80333239 | 80336737 |
| chr10 | 74985766 | 74991677 |
| chr4 | 59435323 | 59438055 |
| chr5 | 34158135 | 34160504 |
| chr2 | 91402245 | 91410576 |
| chr14 | 58295883 | 58298804 |
| chr12 | 18303246 | 18305641 |
| chr9 | 66242580 | 66246293 |
| chr3 | 28347398 | 28351422 |
| chr6 | 121437456 | 121440814 |
| chr4 | 35645419 | 35648251 |

| | | |
|---|---|---|
| chr12 | 79079634 | 79082299 |
| chr2 | 168050789 | 168053063 |
| chr20 | 54927396 | 54929840 |
| chr8 | 102651623 | 102654168 |
| chr7 | 81583783 | 81586178 |
| chr1 | 174440434 | 174442696 |
| chr11 | 23077707 | 23080637 |
| chr10 | 47625814 | 47629189 |
| chr7 | 152275702 | 152278367 |
| chr7 | 61038573 | 61040918 |
| chr15 | 20348785 | 20352327 |
| chr1 | 103311639 | 103314642 |
| chr13 | 87679561 | 87684512 |
| chr7 | 82480872 | 82483263 |
| chr14 | 36343211 | 36346580 |
| chr14 | 19775402 | 19783194 |
| chr12 | 84635532 | 84639931 |
| chr7 | 79908295 | 79911063 |
| chr2 | 186842887 | 186846320 |
| chr2 | 95633063 | 95635528 |
| chr10 | 113367900 | 113371059 |
| chr2 | 186837883 | 186842863 |
| chr22 | 10939142 | 10943958 |
| chr2 | 198362997 | 198366345 |
| chr7 | 84073150 | 84079084 |
| chr10 | 94413120 | 94415614 |
| chr21 | 10443535 | 10446465 |
| chr14 | 79374926 | 79378458 |
| chr8 | 95268032 | 95270588 |
| chr1 | 96792412 | 96795663 |
| chr4 | 157125875 | 157129824 |
| chr7 | 61021724 | 61031535 |
| chr1 | 103254236 | 103258649 |
| chr12 | 65825705 | 65831482 |
| chr1 | 206383082 | 206385727 |
| chr12 | 102028487 | 102031130 |
| chr2 | 186483445 | 186488335 |
| chr2 | 188656416 | 188660434 |
| chr4 | 126807565 | 126811055 |
| chr2 | 189042690 | 189046461 |
| chr10 | 47640992 | 47643653 |
| chr8 | 110718738 | 110721482 |
| chr1 | 219178346 | 219182890 |
| chr12 | 65831696 | 65834159 |
| chr1_KI270713v1_r | 18706 | 21944 |
| chr1 | 103215968 | 103221784 |
| chr8 | 106618267 | 106621504 |
| chr5 | 28933219 | 28936358 |
| chr21 | 20821870 | 20824527 |

| | | |
|---|---|---|
| chr7 | 85977371 | 85979913 |
| chrY | 11712117 | 11719405 |
| chr20 | 13782867 | 13785381 |
| chr6 | 82492972 | 82495272 |
| chr14 | 59498392 | 59503778 |
| chr10 | 60559917 | 60563230 |
| chr4 | 34419995 | 34427319 |
| chr10 | 110916459 | 110923114 |
| chr14 | 19432037 | 19434662 |
| chr7 | 79818986 | 79821216 |
| chr14 | 92159936 | 92163359 |
| chr1 | 146409678 | 146412244 |
| chr10 | 131210035 | 131213024 |
| chr2 | 195922766 | 195927478 |
| chr4 | 156749870 | 156753525 |
| chr7 | 83052188 | 83057523 |
| chr11 | 95789557 | 95791963 |
| chr7 | 80981321 | 80984409 |
| chr16 | 72698098 | 72700340 |
| chr4 | 102825590 | 102829175 |
| chrY | 11695598 | 11705727 |
| chr8 | 106630663 | 106633114 |
| chr15 | 20678203 | 20684818 |
| chr2 | 162476037 | 162479064 |
| chr6 | 160615039 | 160618572 |
| chr21 | 10449566 | 10456251 |
| chr22 | 12714358 | 12717269 |
| chr6 | 26922879 | 26924979 |
| chr14 | 25274478 | 25277928 |
| chr2 | 89771969 | 89787623 |
| chr7 | 157932073 | 157934418 |
| chr21 | 16454856 | 16460170 |
| chr13 | 105601576 | 105604343 |
| chr10 | 17230156 | 17233380 |
| chr10 | 68340008 | 68344692 |
| chr1 | 192315568 | 192319427 |
| chr1 | 88285315 | 88288196 |
| chr1 | 219554048 | 219557377 |
| chr10 | 87422371 | 87424895 |
| chr15 | 36497533 | 36501572 |
| chrY | 11686566 | 11690303 |
| chr2 | 57432036 | 57437549 |
| chr16 | 34595789 | 34602006 |
| chr7 | 83596800 | 83600739 |
| chr3 | 174089256 | 174092401 |
| chr12 | 85405485 | 85410270 |
| chr14 | 50879584 | 50882001 |
| chr2 | 97906581 | 97910042 |
| chr6 | 122046719 | 122052224 |

| | | |
|---|---|---|
| chr14 | 37807857 | 37810720 |
| chr7 | 82753518 | 82758508 |
| chr14 | 90381701 | 90384405 |
| chr21 | 21230573 | 21232922 |
| chr14 | 41595645 | 41601170 |
| chr2 | 56897993 | 56900655 |
| chr16 | 34765959 | 34772325 |
| chr9 | 73090855 | 73093677 |
| chr6 | 81170456 | 81175275 |
| chr1 | 244969899 | 244973714 |
| chr7 | 90297860 | 90301001 |
| chr12 | 78587886 | 78591516 |
| chr6 | 57754045 | 57758976 |
| chrUn_GL000218v1 | 143114 | 147040 |
| chr10 | 74951299 | 74954997 |
| chr2 | 144991531 | 144996934 |
| chr14 | 27467944 | 27471193 |
| chr14 | 60482732 | 60487170 |
| chr10 | 91491942 | 91495575 |
| chr7 | 80539662 | 80542974 |
| chr14 | 86288369 | 86292173 |
| chr10 | 17145911 | 17149699 |
| chr12 | 44720202 | 44723175 |
| chr7 | 58104217 | 58113278 |
| chr1 | 239679146 | 239683240 |
| chr5 | 164016883 | 164019091 |
| chr7 | 80276931 | 80280399 |
| chr10 | 66774301 | 66777988 |
| chr14 | 48346681 | 48349389 |
| chr13 | 71680298 | 71682975 |
| chr12 | 65439891 | 65445652 |
| chr11 | 18861125 | 18863678 |
| chr14 | 19242786 | 19248133 |
| chr7 | 84203961 | 84209311 |
| chr10 | 65247980 | 65253003 |
| chr6 | 141313102 | 141315968 |
| chr4 | 156972379 | 156975059 |
| chr16 | 33146497 | 33148761 |
| chr7 | 92559031 | 92563704 |
| chr6 | 74259482 | 74261938 |
| chr1 | 88943573 | 88946544 |
| chr6 | 70534728 | 70538935 |
| chr10 | 65747322 | 65750404 |
| chr7 | 54472996 | 54476837 |
| chr21 | 16417948 | 16425671 |
| chr2 | 61203421 | 61206720 |
| chr7 | 79381476 | 79385448 |
| chr14 | 101945599 | 101949410 |
| chr7 | 89931649 | 89934195 |

| | | |
|---|---|---|
| chr2 | 176942586 | 176945322 |
| chr22 | 16211845 | 16214204 |
| chr13 | 105620183 | 105627557 |
| chr14 | 36111569 | 36114015 |
| chr5_KI270897v1_a | 832324 | 836621 |
| chr16 | 69566559 | 69573904 |
| chr14 | 49304465 | 49306851 |
| chr14 | 69152377 | 69155312 |
| chr6 | 56181139 | 56183974 |
| chr19 | 45766307 | 45770565 |
| chr15 | 20320839 | 20324294 |
| chr5 | 112160619 | 112164184 |
| chr2 | 40510629 | 40513539 |
| chr12 | 57751917 | 57754659 |
| chr2 | 68130232 | 68133376 |
| chr6 | 75242601 | 75245111 |
| chr6 | 70369958 | 70372651 |
| chr14 | 73020986 | 73024028 |
| chr10 | 111852874 | 111855682 |
| chr2 | 95950834 | 95954647 |
| chr20 | 29091326 | 29096688 |
| chr3 | 146607130 | 146610687 |
| chr14 | 28775006 | 28781563 |
| chr21 | 10405111 | 10407758 |
| chr16 | 34625660 | 34630813 |
| chr7 | 79447762 | 79450867 |
| chr2 | 195315286 | 195319335 |
| chr14 | 98820931 | 98823038 |
| chr12 | 79045292 | 79049510 |
| chr11 | 23052839 | 23060267 |
| chr1 | 189891915 | 189895694 |
| chr9 | 138291415 | 138298233 |
| chr7 | 84228754 | 84232380 |
| chr14 | 41895828 | 41899940 |
| chr12 | 84703893 | 84707544 |
| chr7 | 79561685 | 79564924 |
| chrY | 6259683 | 6266644 |
| chr7 | 85698520 | 85702659 |
| chr7 | 82809328 | 82825055 |
| chr15 | 21264719 | 21269379 |
| chr4 | 35701285 | 35704195 |
| chr9 | 67794022 | 67798850 |
| chr7 | 82800474 | 82809227 |
| chr7 | 80145476 | 80149551 |
| chr2 | 187386094 | 187389944 |
| chr1 | 92168278 | 92171719 |
| chr9 | 67784532 | 67786965 |
| chr7 | 84623082 | 84626832 |
| chr15 | 20655717 | 20666253 |

| | | |
|---|---|---|
| chr7 | 84368147 | 84370424 |
| chr20 | 29389634 | 29393520 |
| chr4 | 176734061 | 176737358 |
| chr1 | 89715250 | 89718465 |
| chr10 | 46175098 | 46178107 |
| chr6 | 56176883 | 56181133 |
| chrY | 9527076 | 9533390 |
| chr2 | 45451553 | 45455491 |
| chr10 | 51388081 | 51390497 |
| chr2 | 172070028 | 172075377 |
| chr14 | 28610758 | 28613446 |
| chr20 | 18748004 | 18751083 |
| chr13 | 80560816 | 80563522 |
| chr14 | 32701265 | 32705389 |
| chr1 | 238494852 | 238499014 |
| chr6 | 71380601 | 71384339 |
| chr13 | 71858786 | 71863963 |
| chr13 | 71750036 | 71753663 |
| chr1 | 149886316 | 149890602 |
| chr7 | 87219003 | 87222363 |
| chr17 | 64505946 | 64508472 |
| chr1 | 61214783 | 61217007 |
| chr2 | 57187892 | 57191016 |
| chr7 | 60919694 | 60938517 |
| chr7 | 84700108 | 84704834 |
| chr2 | 44169006 | 44172511 |
| chr4 | 156769844 | 156772453 |
| chr7 | 80622706 | 80627751 |
| chr2 | 174339446 | 174342263 |
| chr7 | 84068081 | 84072679 |
| chr10 | 114966609 | 114969178 |
| chr13 | 106085873 | 106093482 |
| chr1 | 195411829 | 195414697 |
| chr2 | 91993892 | 91997090 |
| chr1 | 86913190 | 86916556 |
| chr12 | 90472492 | 90476055 |
| chr5 | 27922487 | 27929772 |
| chr12 | 85037873 | 85041195 |
| chr13 | 41666971 | 41670876 |
| chr4 | 157039446 | 157044180 |
| chr7 | 81701278 | 81704183 |
| chr12 | 85575386 | 85577736 |
| chr2 | 189215620 | 189218446 |
| chr2 | 172081490 | 172085125 |
| chr2 | 55381666 | 55384441 |
| chr4 | 89900725 | 89903935 |
| chr1 | 97730632 | 97733257 |
| chr1 | 108691945 | 108695889 |
| chr1 | 104117029 | 104123665 |

| | | |
|---|---|---|
| chr17 | 65556817 | 65559753 |
| chr15_KI270852v1_ | 20757 | 26582 |
| chr20 | 49915084 | 49917337 |
| chrY | 11261892 | 11265847 |
| chr2 | 186846761 | 186852493 |
| chr4 | 49307976 | 49310058 |
| chr1 | 103109521 | 103113739 |
| chr1 | 197603078 | 197608623 |
| chr22_KI270734v1_ | 96902 | 99587 |
| chr7 | 62370552 | 62374728 |
| chr2 | 55614367 | 55618157 |
| chr7 | 87415645 | 87418460 |
| chr15_KI270905v1_ | 2684091 | 2687651 |
| chr1 | 103690316 | 103696892 |
| chr14 | 41081452 | 41085349 |
| chr14 | 76786574 | 76789411 |
| chr1 | 107166466 | 107169531 |
| chr1 | 102890821 | 102896504 |
| chr19 | 18635270 | 18639038 |
| chr15 | 20621703 | 20625060 |
| chr1 | 219275495 | 219277707 |
| chr1 | 92295532 | 92297887 |
| chr16_KI270728v1_ | 1806800 | 1810478 |
| chr2 | 169285332 | 169288980 |
| chr9 | 25968213 | 25970676 |
| chr20 | 56367423 | 56370583 |
| chr6 | 72748890 | 72752236 |
| chr11 | 22936611 | 22940396 |
| chr6 | 122061256 | 122066385 |
| chr2 | 188987804 | 188992891 |
| chr12 | 75644919 | 75649009 |
| chr7 | 83100538 | 83105423 |
| chr10 | 17026928 | 17030331 |
| chr16 | 80603384 | 80606057 |
| chr8 | 12434941 | 12439133 |
| chr1 | 103039374 | 103044838 |
| chr1 | 93549056 | 93551663 |
| chr17 | 68247362 | 68250655 |
| chr7 | 84145653 | 84150295 |
| chr1 | 111349197 | 111351507 |
| chr12 | 84655573 | 84661252 |
| chr5 | 143963999 | 143966243 |
| chr10 | 65228695 | 65235447 |
| chr1 | 103179741 | 103185006 |
| chr6 | 80126636 | 80128956 |
| chr9 | 6943465 | 6946587 |
| chr7 | 9936438 | 9940551 |
| chr14 | 27180141 | 27185565 |
| chr2 | 57077526 | 57080920 |

| | | |
|---|---|---|
| chr2 | 176669917 | 176672690 |
| chr9 | 64613009 | 64616907 |
| chr2 | 199908806 | 199912860 |
| chr6 | 101572730 | 101575895 |
| chr10 | 74976263 | 74979872 |
| chr4 | 32477046 | 32479505 |
| chr2 | 194266589 | 194268779 |
| chr3 | 85478794 | 85481836 |
| chr11 | 90707041 | 90710477 |
| chr13 | 67144410 | 67149168 |
| chr1 | 100049419 | 100052700 |
| chr4 | 32112339 | 32115305 |
| chr12 | 77937639 | 77944088 |
| chr2 | 50318551 | 50321535 |
| chr8 | 108114248 | 108117415 |
| chr21 | 15888827 | 15891550 |
| chr5 | 27033895 | 27036137 |
| chr7 | 82739401 | 82742326 |
| chr14 | 48425160 | 48428813 |
| chr10 | 103887841 | 103890936 |
| chr12 | 25322210 | 25328019 |
| chr12_GL877875v1_ | 5249 | 8639 |
| chr1 | 105970844 | 105973922 |
| chr7 | 79505039 | 79510509 |
| chr4 | 115020807 | 115022991 |
| chr16 | 76538145 | 76541692 |
| chr1 | 70090668 | 70093671 |
| chr22 | 16330234 | 16332973 |
| chrUn_GL000195v1 | 177118 | 180105 |
| chr4 | 115779930 | 115786471 |
| chr14 | 21266847 | 21270113 |
| chr10 | 52215262 | 52218228 |
| chr10 | 46817295 | 46822879 |
| chr14 | 36317488 | 36323254 |
| chr10 | 38485963 | 38495364 |
| chr12 | 39754190 | 39760113 |
| chr10 | 81716225 | 81718882 |
| chr15 | 37872197 | 37881923 |
| chr12 | 81776978 | 81780398 |
| chr14 | 28380537 | 28383723 |
| chr14 | 25919233 | 25922054 |
| chr4 | 116089560 | 116092984 |
| chr8 | 115583122 | 115588268 |
| chr10 | 46556593 | 46561074 |
| chr7 | 83978127 | 83982359 |
| chr10 | 66929813 | 66932432 |
| chr3 | 76985636 | 76989669 |
| chr9 | 10063803 | 10067530 |
| chr7 | 84031881 | 84036725 |

| | | |
|---|---|---|
| chr2 | 35167003 | 35170129 |
| chr21 | 16577286 | 16582170 |
| chr22 | 15774349 | 15779112 |
| chr8 | 113077381 | 113081469 |
| chr7 | 82358687 | 82363127 |
| chr2 | 165573252 | 165578900 |
| chr10 | 67117029 | 67120443 |
| chr1 | 98469394 | 98473760 |
| chr9 | 138315633 | 138321300 |
| chr14 | 19793781 | 19798474 |
| chr1 | 101902159 | 101908847 |
| chr17 | 22123157 | 22131504 |
| chr1 | 97917587 | 97921448 |
| chr21 | 9739153 | 9743329 |
| chr14 | 27634755 | 27638700 |
| chr12 | 43670992 | 43674200 |
| chr16 | 34087324 | 34093418 |
| chr17 | 65721079 | 65725581 |
| chr16 | 36191595 | 36200324 |
| chr10 | 56276830 | 56280010 |
| chr12 | 77828333 | 77832733 |
| chr1 | 104031461 | 104035310 |
| chr12 | 45730065 | 45733970 |
| chr7 | 80885252 | 80890859 |
| chr7 | 80698728 | 80701725 |
| chr8 | 112553901 | 112557540 |
| chr21 | 16585265 | 16589987 |
| chr14 | 19934544 | 19937899 |
| chr2 | 199459536 | 199463944 |
| chr11 | 16451104 | 16454978 |
| chr1 | 50194906 | 50197758 |
| chr10 | 54763721 | 54767024 |
| chr14 | 41978137 | 41982383 |
| chr6 | 76633898 | 76636869 |
| chr14 | 19905031 | 19908508 |
| chr12 | 84640638 | 84655086 |
| chr14 | 26545440 | 26548727 |
| chr8 | 77332399 | 77335434 |
| chr8 | 113310686 | 113315883 |
| chr14 | 41835758 | 41841360 |
| chr4 | 107030597 | 107033997 |
| chr6 | 63195416 | 63199466 |
| chr7 | 78313332 | 78315979 |
| chr8 | 110701788 | 110704919 |
| chr15_KI270905v1_ | 828737 | 831863 |
| chr14 | 87674502 | 87677535 |
| chr1 | 103014713 | 103018574 |
| chr12 | 84984111 | 84986993 |
| chr14 | 23745582 | 23750021 |

| | | |
|---|---|---|
| chr8 | 113372943 | 113378668 |
| chr14 | 19484510 | 19490098 |
| chr17 | 22188643 | 22192862 |
| chr1 | 113394767 | 113398726 |
| chr14 | 88821629 | 88825950 |
| chr2 | 185572871 | 185575959 |
| chr2 | 162286351 | 162299684 |
| chr6 | 50683637 | 50686758 |
| chr7 | 124229359 | 124233883 |
| chr12 | 25216242 | 25219346 |
| chr7 | 77936742 | 77941662 |
| chr8 | 76669917 | 76673590 |
| chr2 | 226206800 | 226209972 |
| chr16 | 36122741 | 36135683 |
| chr8 | 95993032 | 95996415 |
| chr2 | 189139719 | 189145363 |
| chr12 | 77721249 | 77725448 |
| chr1 | 143234225 | 143243665 |
| chr8 | 113399929 | 113405921 |
| chr7 | 62302669 | 62310534 |
| chr7 | 62400247 | 62412393 |
| chr5 | 88726903 | 88730920 |
| chr9 | 67787222 | 67792404 |
| chr1 | 83859860 | 83864058 |
| chr17 | 26783117 | 26789230 |
| chr2 | 50234269 | 50238190 |
| chr14 | 37268789 | 37273787 |
| chr10 | 106153493 | 106156316 |
| chr14 | 54474613 | 54478430 |
| chr14 | 39428065 | 39432958 |
| chr6 | 72228674 | 72234310 |
| chr14 | 27568739 | 27571479 |
| chr8 | 112139708 | 112144275 |
| chr6 | 121880101 | 121885117 |
| chr1 | 172135344 | 172148294 |
| chr4 | 107168340 | 107173988 |
| chr1 | 197526120 | 197530007 |
| chr14 | 26448992 | 26453627 |
| chr2 | 176674888 | 176680164 |
| chr2 | 51935273 | 51938823 |
| chr13 | 81210103 | 81213373 |
| chr1 | 188602144 | 188606302 |
| chr7 | 81099597 | 81102383 |
| chr14 | 19425953 | 19432024 |
| chr7 | 82144165 | 82149635 |
| chr1 | 101951293 | 101955074 |
| chr2 | 97509066 | 97512910 |
| chr14 | 67379937 | 67383679 |
| chr1 | 97323547 | 97328381 |

| | | |
|---|---|---|
| chr10 | 60570258 | 60574629 |
| chr14 | 40110603 | 40114541 |
| chr12 | 90944995 | 90948634 |
| chr10 | 68070994 | 68076108 |
| chr1 | 191639013 | 191644521 |
| chr8 | 113426392 | 113431930 |
| chr1 | 198919561 | 198924194 |
| chr2 | 193938988 | 193943541 |
| chr8 | 114048387 | 114052547 |
| chr12 | 45723873 | 45727114 |
| chr12 | 63219667 | 63222607 |
| chr2 | 77671247 | 77677161 |
| chr8 | 97457993 | 97461271 |
| chr20 | 28578626 | 28587558 |
| chr17 | 22134845 | 22140435 |
| chr14 | 27598411 | 27601465 |
| chr3 | 95675131 | 95685638 |
| chr1 | 98431401 | 98435398 |
| chr14 | 47057841 | 47064671 |
| chr6 | 56098561 | 56105294 |
| chr8 | 113433622 | 113442037 |
| chr7 | 97010948 | 97025912 |
| chr5_KI270897v1_a | 820066 | 824277 |
| chr14 | 60432498 | 60437007 |
| chr15_KI270905v1_ | 4818194 | 4821388 |
| chr14 | 28752000 | 28760918 |
| chr14 | 26469477 | 26475906 |
| chr1 | 187976295 | 187979905 |
| chr1 | 86725459 | 86728769 |
| chr5 | 27937802 | 27944366 |
| chr9 | 41007804 | 41010970 |
| chr2 | 196166935 | 196171625 |
| chr14 | 84015467 | 84019922 |
| chr15 | 38078393 | 38081989 |
| chr18 | 3453300 | 3457276 |
| chr12 | 88138831 | 88143432 |
| chr14 | 39037484 | 39041526 |
| chr15 | 37780112 | 37784828 |
| chr2 | 164973903 | 164977243 |
| chr2 | 58662519 | 58666989 |
| chr14 | 43588581 | 43592567 |
| chr14 | 26441207 | 26444115 |
| chr1 | 125177622 | 125185060 |
| chr14 | 47541386 | 47545602 |
| chr1 | 125079418 | 125085874 |
| chr10 | 67268609 | 67272332 |
| chr6 | 93392596 | 93397081 |
| chr7 | 81877054 | 81880114 |
| chr8 | 90763908 | 90767785 |

| | | |
|---|---|---|
| chr1 | 191316968 | 191321083 |
| chr5 | 59094343 | 59098201 |
| chr22 | 15697590 | 15702394 |
| chr7 | 62440824 | 62450508 |
| chr2 | 195445225 | 195451784 |
| chr14 | 68110405 | 68114597 |
| chr1 | 100214867 | 100217680 |
| chr14 | 26735777 | 26739603 |
| chr12 | 73063291 | 73066220 |
| chr14 | 39671100 | 39676472 |
| chr7 | 82910233 | 82913017 |
| chr2 | 103958792 | 103964926 |
| chr15 | 20284361 | 20287800 |
| chr1 | 100350035 | 100354652 |
| chr12 | 57902498 | 57906231 |
| chr17 | 22142184 | 22147014 |
| chr2 | 40198386 | 40201969 |
| chr7 | 81982164 | 81988997 |
| chr3 | 77000480 | 77005075 |
| chr1 | 104411681 | 104416052 |
| chr7 | 80456929 | 80460277 |
| chr10 | 66074233 | 66076955 |
| chr6 | 63688349 | 63691271 |
| chr9 | 5435568 | 5439116 |
| chr8 | 96265829 | 96269393 |
| chr8 | 112976411 | 112983074 |
| chr2 | 103669119 | 103671606 |
| chr8 | 115662827 | 115669076 |
| chr8 | 110356372 | 110361702 |
| chr9 | 62471569 | 62474969 |
| chr1 | 97978018 | 97981250 |
| chr14 | 27946575 | 27949340 |
| chr1 | 125068791 | 125076292 |
| chr1 | 93180839 | 93185476 |
| chr20 | 8657620 | 8664867 |
| chr1 | 69574831 | 69577314 |
| chr14 | 36552376 | 36557741 |
| chr11 | 105910520 | 105916192 |
| chr14 | 36461582 | 36465836 |
| chr11 | 4354079 | 4357435 |
| chr22 | 15852493 | 15859131 |
| chr4 | 93056349 | 93061868 |
| chr2 | 209333416 | 209336529 |
| chr11 | 107927619 | 107930247 |
| chr2 | 242144853 | 242147050 |
| chr11 | 23592364 | 23596906 |
| chr9 | 66003619 | 66007650 |
| chr1 | 167913426 | 167915719 |
| chr2 | 187467324 | 187470387 |

| | | |
|---|---|---|
| chr1 | 155061027 | 155063833 |
| chr15 | 20639532 | 20642090 |
| chr14 | 41120670 | 41125856 |
| chr5 | 27986971 | 27989641 |
| chr4 | 79602388 | 79605490 |
| chr2 | 195481591 | 195487952 |
| chr11 | 87316545 | 87319120 |
| chrX | 154378316 | 154380483 |
| chr15_KI270905v1_ | 607853 | 610003 |
| chr2 | 83271331 | 83274277 |
| chr14 | 87379028 | 87381533 |
| chr14 | 100426425 | 100429849 |
| chr1 | 74730821 | 74733458 |
| chr2 | 57901795 | 57906400 |
| chr13 | 110711776 | 110713908 |
| chr12 | 84618515 | 84621226 |
| chr10 | 95674301 | 95676559 |
| chr2 | 48730734 | 48733827 |
| chr8 | 127176383 | 127179180 |
| chr14 | 90299623 | 90302138 |
| chr2 | 189176113 | 189180570 |
| chr1 | 143184535 | 143189479 |
| chr10 | 58741892 | 58745319 |
| chr14 | 28318128 | 28321268 |
| chr2 | 51245411 | 51248212 |
| chr2 | 187575679 | 187579258 |
| chr1 | 196890810 | 196893111 |
| chr10 | 127231860 | 127234468 |
| chr14 | 28735776 | 28740545 |
| chr2 | 49448678 | 49451669 |
| chr2 | 39225396 | 39229734 |
| chr8 | 110766721 | 110770059 |
| chr7 | 89330244 | 89333750 |
| chr15 | 94378637 | 94380894 |
| chr2 | 50591133 | 50594187 |
| chr6 | 45092443 | 45095327 |
| chr2 | 56826348 | 56828850 |
| chr16 | 73670347 | 73672652 |
| chr1 | 102147862 | 102150852 |
| chr2 | 195006950 | 195010346 |
| chr6 | 74492076 | 74494358 |
| chr11 | 97903319 | 97905370 |
| chr10 | 69405934 | 69408615 |
| chr7 | 11116633 | 11120113 |
| chr2 | 184782782 | 184785504 |
| chr2 | 211583484 | 211586660 |
| chr14 | 41313031 | 41316076 |
| chr1 | 219564561 | 219567107 |
| chr10 | 87874225 | 87879535 |

| | | |
|---|---|---|
| chr7 | 62337081 | 62341523 |
| chr14 | 25555471 | 25558480 |
| chr2 | 189879025 | 189881359 |
| chr9 | 66597298 | 66600522 |
| chr3 | 77806771 | 77809757 |
| chr1 | 61925532 | 61929625 |
| chr7 | 83512619 | 83515162 |
| chr2 | 169140604 | 169142997 |
| chr2 | 15681199 | 15683639 |
| chr2 | 193393015 | 193396241 |
| chr12 | 78151472 | 78155291 |
| chr10 | 64483064 | 64485493 |
| chr1 | 98215790 | 98219221 |
| chr1 | 185371592 | 185374893 |
| chr12 | 19157562 | 19161015 |
| chr12 | 17504241 | 17506912 |
| chr14 | 58744619 | 58746917 |
| chr14 | 30618204 | 30622594 |
| chr2 | 50561635 | 50564329 |
| chr16 | 76081102 | 76085260 |
| chr13 | 94804462 | 94807577 |
| chr18 | 76495356 | 76498641 |
| chr1 | 35760408 | 35762888 |
| chr10 | 111985369 | 111988572 |
| chr10 | 55795797 | 55798776 |
| chr8 | 65491451 | 65494535 |
| chr3 | 144028768 | 144031205 |
| chr7 | 25854707 | 25858008 |
| chr7 | 103911567 | 103914866 |
| chr10 | 54552961 | 54555454 |
| chr1 | 104311890 | 104315337 |
| chr1 | 90857564 | 90859664 |
| chr1 | 218166769 | 218172784 |
| chr4 | 157717744 | 157720244 |
| chr5 | 161632989 | 161635660 |
| chr6 | 70594628 | 70598502 |
| chr7 | 152228771 | 152231094 |
| chr12 | 78086184 | 78088752 |
| chr12 | 85075758 | 85078115 |
| chr10 | 105797352 | 105800427 |
| chr7 | 79242199 | 79244611 |
| chr7 | 62293763 | 62302577 |
| chr2 | 61871069 | 61873811 |
| chr7 | 83853453 | 83857311 |
| chr7 | 83719915 | 83722824 |
| chr14 | 46171707 | 46176145 |
| chr18 | 2931412 | 2933630 |
| chr1 | 32213949 | 32216193 |
| chr1 | 62258693 | 62260942 |

| | | |
|---|---|---|
| chr6 | 61969558 | 61972884 |
| chr13 | 67022491 | 67026437 |
| chr8 | 39916079 | 39920105 |
| chr7 | 82018999 | 82021767 |
| chr10 | 51815864 | 51818401 |
| chr10 | 66010103 | 66014911 |
| chr16 | 78240262 | 78242328 |
| chr9 | 73157492 | 73162903 |
| chr7 | 7878125 | 7880632 |
| chr7 | 86444073 | 86446513 |
| chr5_KI270897v1_a | 770185 | 774267 |
| chr3 | 81002234 | 81004553 |
| chr12 | 38735907 | 38738125 |
| chr8 | 99633914 | 99637327 |
| chr20 | 54214317 | 54216826 |
| chr17 | 60831056 | 60835248 |
| chr1 | 238646533 | 238649232 |
| chr2 | 179951656 | 179954473 |
| chr1 | 148487424 | 148491146 |
| chr11 | 25574013 | 25576355 |
| chr16 | 34713439 | 34716029 |
| chr12 | 82356901 | 82360702 |
| chr3 | 174458356 | 174460944 |
| chr6 | 90546105 | 90548710 |
| chr14 | 26597368 | 26600400 |
| chr2 | 197109803 | 197112755 |
| chr2 | 85540943 | 85544471 |
| chr9 | 62434042 | 62436497 |
| chr7 | 81519513 | 81523151 |
| chr14 | 26963963 | 26967437 |
| chr14 | 27414432 | 27416858 |
| chr10 | 55538795 | 55542429 |
| chr2 | 40813018 | 40817524 |
| chr1 | 169986589 | 169991347 |
| chr7 | 94393618 | 94396231 |
| chr7 | 84264779 | 84267440 |
| chr2 | 104705997 | 104709661 |
| chr8 | 57441329 | 57443461 |
| chr15 | 53978706 | 53981726 |
| chr5 | 55158222 | 55161064 |
| chr11 | 130124930 | 130127397 |
| chr14 | 32928976 | 32932547 |
| chr1 | 240112127 | 240114205 |
| chr1 | 72207266 | 72209598 |
| chr3 | 1273585 | 1275682 |
| chr15 | 47908900 | 47911361 |
| chr2 | 56370292 | 56373475 |
| chr10 | 91863814 | 91866396 |
| chr2 | 21844188 | 21846229 |

| | | |
|---|---|---|
| chr6 | 52419323 | 52423005 |
| chr6 | 155915632 | 155917855 |
| chr16 | 76723019 | 76725932 |
| chr1 | 187266519 | 187268889 |
| chr2 | 241353969 | 241358367 |
| chr12 | 65844015 | 65846452 |
| chr15 | 43062261 | 43064616 |
| chr7 | 79627597 | 79629934 |
| chr4 | 31471969 | 31474081 |
| chr2 | 181130249 | 181132977 |
| chr1 | 145096423 | 145098995 |
| chr7 | 104566108 | 104568394 |
| chr5 | 121835825 | 121839261 |
| chr11 | 90201243 | 90205451 |
| chr1 | 87914520 | 87917953 |
| chr16 | 34659201 | 34664967 |
| chr15 | 35541257 | 35546083 |
| chr2 | 200418380 | 200420560 |
| chr14 | 80054740 | 80058353 |
| chr2 | 187483364 | 187490189 |
| chr12 | 65821887 | 65825330 |
| chr12 | 22058565 | 22060571 |
| chr1 | 105170335 | 105173917 |
| chr5 | 129377307 | 129380552 |
| chr1 | 70807536 | 70809726 |
| chr15 | 71558732 | 71561941 |
| chr21 | 7575111 | 7577350 |
| chr20 | 16998746 | 17001205 |
| chr3 | 76806238 | 76808421 |
| chr10 | 105953935 | 105956450 |
| chr5 | 105901481 | 105903610 |
| chr2 | 188882163 | 188885681 |
| chr17 | 79779809 | 79783112 |
| chr6 | 123389431 | 123394112 |
| chr20 | 40398364 | 40400526 |
| chr2 | 91573626 | 91576468 |
| chr11 | 65425182 | 65428777 |
| chr14 | 22492630 | 22495132 |
| chr14 | 28174442 | 28176754 |
| chr5 | 88245989 | 88248352 |
| chr12 | 87060138 | 87063620 |
| chr6 | 45430311 | 45440621 |
| chr13 | 86563925 | 86566793 |
| chr2 | 50208315 | 50212921 |
| chr2 | 204758998 | 204763902 |
| chr1 | 107340812 | 107344114 |
| chr1 | 31690483 | 31692698 |
| chr20 | 21508132 | 21510154 |
| chr14 | 33619676 | 33623031 |

| | | |
|---|---|---|
| chr11 | 20604055 | 20607898 |
| chr20 | 53486479 | 53490192 |
| chr1 | 105336082 | 105339996 |
| chr2 | 188700307 | 188702715 |
| chr20 | 61389959 | 61393230 |
| chr7 | 84290055 | 84294372 |
| chr2 | 35146960 | 35151840 |
| chr2 | 184897710 | 184900170 |
| chr9 | 66372075 | 66375799 |
| chr10 | 57842826 | 57845809 |
| chr11 | 84509971 | 84511993 |
| chr6 | 72302215 | 72305649 |
| chr7 | 82973217 | 82975844 |
| chr19 | 33175130 | 33177993 |
| chr16 | 83307331 | 83310676 |
| chr2 | 66806571 | 66810047 |
| chr12 | 85928053 | 85932515 |
| chr5 | 92550615 | 92555632 |
| chr3 | 76863525 | 76866348 |
| chr10 | 6964378 | 6966841 |
| chr10 | 99057334 | 99059797 |
| chr8 | 10881089 | 10883467 |
| chr2 | 35007544 | 35012160 |
| chr15 | 36293686 | 36295721 |
| chr13 | 81615367 | 81619000 |
| chr1 | 102022276 | 102025954 |
| chr1 | 69762133 | 69764607 |
| chr3 | 121647133 | 121649358 |
| chr14 | 44218537 | 44220554 |
| chr2 | 55230912 | 55233340 |
| chr12 | 65879476 | 65882726 |
| chr6 | 160556254 | 160558967 |
| chr21 | 7592705 | 7594899 |
| chr14 | 47384805 | 47387011 |
| chr11 | 16342090 | 16346845 |
| chr2 | 59802006 | 59805992 |
| chr14 | 61255449 | 61259799 |
| chr6 | 2513129 | 2515689 |
| chr6 | 132670389 | 132672666 |
| chr20 | 10177574 | 10180865 |
| chr10 | 90723199 | 90726218 |
| chr19 | 35738853 | 35741085 |
| chr10 | 110006829 | 110010461 |
| chr20 | 61872427 | 61874606 |
| chr12 | 14773287 | 14778358 |
| chr10 | 62977661 | 62980337 |
| chr1 | 6612121 | 6614147 |
| chr1 | 198770708 | 198773926 |
| chr14 | 31692648 | 31695690 |

| | | |
|---|---|---|
| chr14 | 19668241 | 19671762 |
| chr21 | 10394380 | 10397263 |
| chr1 | 94752835 | 94755530 |
| chr3 | 105683873 | 105687702 |
| chr10 | 61750129 | 61753679 |
| chr4 | 172591332 | 172593435 |
| chr5 | 114268478 | 114270886 |
| chr10 | 17084710 | 17087211 |
| chr14 | 28590942 | 28594404 |
| chr4 | 28025741 | 28028457 |
| chr21 | 9764490 | 9770483 |
| chr6 | 125232147 | 125235095 |
| chr16 | 34895338 | 34906892 |
| chr8 | 65152448 | 65154612 |
| chr13 | 35287744 | 35295735 |
| chr15_KI270852v1_ | -306 | 3036 |
| chr8 | 119745141 | 119749258 |
| chr10 | 116705020 | 116708002 |
| chr5_KI270897v1_a | 147215 | 153094 |
| chr14 | 22487081 | 22490013 |
| chr6 | 142111823 | 142114746 |
| chr12 | 10212054 | 10214283 |
| chr14 | 35772904 | 35777480 |
| chr2 | 68262068 | 68264373 |
| chr7 | 105109264 | 105112271 |
| chr12 | 85098402 | 85102490 |
| chr12 | 75177572 | 75180429 |
| chr14 | 57021639 | 57023979 |
| chr3 | 76990981 | 76994495 |
| chr1 | 97824278 | 97827135 |
| chr10 | 46708037 | 46710863 |
| chr2 | 193314011 | 193317067 |
| chr14 | 19261001 | 19264117 |
| chr10 | 55240138 | 55242440 |
| chr12 | 90757681 | 90760800 |
| chr5 | 168765055 | 168767788 |
| chr11 | 111925695 | 111927989 |
| chr10 | 86732952 | 86735482 |
| chr1 | 98748882 | 98751053 |
| chr12 | 114391461 | 114394368 |
| chr13 | 49003796 | 49006627 |
| chr3 | 95711995 | 95714846 |
| chr2 | 47798719 | 47801480 |
| chr7 | 80273993 | 80276073 |
| chr15 | 20416967 | 20422041 |
| chr14 | 54562251 | 54565077 |
| chr7 | 11062359 | 11064821 |
| chr1 | 97869772 | 97874984 |
| chr14 | 21906064 | 21908949 |

| | | |
|---|---|---|
| chr12 | 96580635 | 96582913 |
| chr5 | 127796378 | 127799679 |
| chr14 | 19955569 | 19959182 |
| chr1 | 195852359 | 195855455 |
| chr12 | 16715923 | 16720258 |
| chr7 | 92366047 | 92368267 |
| chr4 | 30462956 | 30465454 |
| chr14 | 27318627 | 27322031 |
| chr2 | 50670645 | 50673889 |
| chr1 | 89150317 | 89153040 |
| chr2 | 27637798 | 27640635 |
| chr12 | 76420616 | 76423384 |
| chr7 | 9513996 | 9516640 |
| chr6 | 97979406 | 97981983 |
| chr1 | 87963721 | 87966520 |
| chr1 | 106816612 | 106819353 |
| chr15 | 62431508 | 62434144 |
| chr1 | 143198555 | 143203831 |
| chr10 | 115991240 | 115993766 |
| chr2 | 50732435 | 50736072 |
| chr15 | 35286363 | 35288702 |
| chr15_KI270850v1_ | 250088 | 258450 |
| chr11 | 133452576 | 133455753 |
| chr14 | 42544551 | 42547586 |
| chr2 | 44578414 | 44581482 |
| chr6 | 152299931 | 152302982 |
| chr14 | 28727787 | 28730585 |
| chr6 | 132816968 | 132819123 |
| chr1 | 172246503 | 172248678 |
| chr6 | 156963801 | 156966160 |
| chr2 | 60150782 | 60153030 |
| chr9 | 31067854 | 31070060 |
| chr8 | 113974284 | 113978485 |
| chr8 | 90063224 | 90066634 |
| chr3 | 70064520 | 70066651 |
| chr7 | 79150191 | 79152528 |
| chr12 | 44206799 | 44208871 |
| chr1 | 107410039 | 107413107 |
| chr1 | 99179978 | 99183374 |
| chr1 | 215184750 | 215187200 |
| chr2 | 77866871 | 77869016 |
| chr15_KI270852v1_ | 96414 | 103647 |
| chr14 | 95534325 | 95538746 |
| chr2 | 76821853 | 76825224 |
| chr2 | 51551042 | 51555281 |
| chr15_KI270905v1_ | 4814132 | 4817801 |
| chr11 | 38950232 | 38952816 |
| chr12 | 83210566 | 83212908 |
| chr17 | 15404336 | 15406504 |

| | | |
|---|---|---|
| chr11 | 89932413 | 89936148 |
| chr10 | 56019507 | 56022809 |
| chr1 | 93758525 | 93760793 |
| chr13 | 23172126 | 23174272 |
| chr14 | 99415455 | 99419857 |
| chr2 | 211451593 | 211456981 |
| chr10 | 57115909 | 57119034 |
| chr2 | 73069244 | 73072322 |
| chr3 | 104766853 | 104769902 |
| chr14 | 100380475 | 100384713 |
| chr7 | 114957627 | 114960391 |
| chr10 | 78968324 | 78970492 |
| chr1 | 148892799 | 148895209 |
| chr1 | 103360334 | 103363244 |
| chr7 | 93303650 | 93306707 |
| chr2 | 226799366 | 226804592 |
| chr14 | 29227700 | 29233538 |
| chr10 | 67426936 | 67430591 |
| chr3 | 29927094 | 29929520 |
| chr6 | 133751609 | 133754494 |
| chr20 | 13550192 | 13552325 |
| chr13 | 83760333 | 83762695 |
| chr14 | 41159820 | 41165595 |
| chr1 | 103246325 | 103252002 |
| chr2 | 36026276 | 36028326 |
| chr2 | 187836211 | 187839246 |
| chr1 | 188295315 | 188297987 |
| chr11 | 85662028 | 85665696 |
| chr1 | 66629876 | 66632004 |
| chr2 | 166948603 | 166951226 |
| chr20 | 28601324 | 28605724 |
| chr7 | 54752854 | 54755493 |
| chr7 | 62451533 | 62456819 |
| chr10 | 49954131 | 49956835 |
| chr6 | 26053864 | 26059710 |
| chr1 | 103533502 | 103535934 |
| chr1 | 188656658 | 188658900 |
| chr14 | 86784438 | 86788957 |
| chr12 | 90939171 | 90941668 |
| chr6 | 133013858 | 133016957 |
| chr14 | 19180020 | 19182131 |
| chr8 | 91629649 | 91633597 |
| chr12 | 118185241 | 118190214 |
| chr2 | 118065172 | 118067238 |
| chr4 | 126306648 | 126308650 |
| chr2 | 226113902 | 226122859 |
| chr1 | 237439564 | 237442598 |
| chr2 | 59497773 | 59500283 |
| chr2 | 59779824 | 59781952 |

| | | |
|---|---|---|
| chr2 | 224585204 | 224588470 |
| chr13 | 107442400 | 107444409 |
| chr14 | 26505971 | 26512571 |
| chr7 | 55190215 | 55192233 |
| chr10 | 95374607 | 95378134 |
| chr14 | 85999614 | 86001760 |
| chr16 | 52602082 | 52604318 |
| chr15 | 49420851 | 49423668 |
| chr1 | 116077332 | 116079740 |
| chr14 | 105145324 | 105148294 |
| chr6 | 133291259 | 133293577 |
| chr2 | 39539451 | 39542318 |
| chr10 | 54031792 | 54034036 |
| chr10 | 100868116 | 100870352 |
| chr14 | 89290922 | 89294608 |
| chr2 | 221153626 | 221156927 |
| chr1 | 149319488 | 149321492 |
| chr1 | 51438832 | 51441151 |
| chr10 | 46541593 | 46544719 |
| chr11 | 28187802 | 28192550 |
| chr1 | 191165374 | 191168973 |
| chr10 | 67762976 | 67765391 |
| chr14 | 32254532 | 32259179 |
| chr1 | 143538241 | 143541177 |
| chr1 | 103945635 | 103948598 |
| chr11 | 26410754 | 26413948 |
| chr16 | 35185824 | 35188236 |
| chr12 | 82320790 | 82323060 |
| chr14 | 98839796 | 98842359 |
| chr4 | 17809711 | 17812565 |
| chr15 | 101751715 | 101753889 |
| chr6 | 160643977 | 160647213 |
| chr4 | 175637452 | 175640160 |
| chr10 | 78214446 | 78216446 |
| chr2 | 185799275 | 185801319 |
| chr2 | 219689547 | 219692721 |
| chr12 | 81227850 | 81231420 |
| chr3 | 88058567 | 88060731 |
| chr7 | 84211728 | 84215032 |
| chr22 | 15839269 | 15842533 |
| chr3 | 77047778 | 77049931 |
| chr2 | 210309728 | 210314581 |
| chr12 | 77799651 | 77802794 |
| chr1 | 102897197 | 102901032 |
| chr1 | 71271101 | 71273970 |
| chr1 | 96174560 | 96178434 |
| chr8 | 59966448 | 59969689 |
| chr14 | 71800450 | 71802970 |
| chr2 | 202566026 | 202568245 |

| | | |
|---|---|---|
| chr3 | 182632281 | 182634572 |
| chr14 | 100249903 | 100253032 |
| chr1 | 188714957 | 188717069 |
| chr2 | 88772016 | 88777129 |
| chr1 | 200172882 | 200175821 |
| chr14 | 96537056 | 96540519 |
| chr7 | 88743789 | 88746177 |
| chr15 | 77219634 | 77222762 |
| chr10 | 62057650 | 62061183 |
| chr10 | 101441709 | 101443809 |
| chr1 | 81977849 | 81981155 |
| chr10 | 84189902 | 84192300 |
| chr1 | 125151405 | 125154926 |
| chr7 | 11138940 | 11143606 |
| chr10 | 79710928 | 79713265 |
| chr10 | 43199202 | 43202184 |
| chr1 | 102912937 | 102917141 |
| chr1 | 199242233 | 199244751 |
| chr14 | 19255193 | 19259984 |
| chr16 | 81093763 | 81095810 |
| chr16 | 77146640 | 77150950 |
| chr7 | 79710223 | 79716692 |
| chr1 | 185864091 | 185866391 |
| chr4 | 116939430 | 116942490 |
| chr7 | 58113736 | 58119496 |
| chr2 | 218451694 | 218453989 |
| chr8 | 16824168 | 16827960 |
| chr1 | 59846577 | 59849074 |
| chr1 | 195315321 | 195318564 |
| chr6 | 19835375 | 19838643 |
| chr2 | 57889549 | 57897988 |
| chr20 | 30897030 | 30899717 |
| chr14 | 33608892 | 33611875 |
| chr12 | 84869437 | 84872316 |
| chr10 | 74079897 | 74082003 |
| chr11 | 23862030 | 23864365 |
| chr1 | 188413866 | 188416609 |
| chr10 | 54761492 | 54763510 |
| chr18 | 4290307 | 4292328 |
| chr6 | 135039421 | 135042041 |
| chr4 | 13471444 | 13474050 |
| chr7 | 64689343 | 64692171 |
| chr14 | 40596443 | 40600347 |
| chr10 | 103240977 | 103243009 |
| chr7 | 51860587 | 51862721 |
| chr14 | 43035020 | 43038379 |
| chr1 | 98038089 | 98041677 |
| chr7 | 80412922 | 80416627 |
| chr2 | 86623134 | 86627101 |

| | | |
|---|---|---|
| chr14 | 44896485 | 44900024 |
| chr8 | 121498210 | 121500676 |
| chr13 | 41688937 | 41696853 |
| chr12 | 56723653 | 56725989 |
| chr4 | 31878909 | 31881061 |
| chr10 | 66237326 | 66239564 |
| chr10 | 62543065 | 62545153 |
| chr15 | 38068717 | 38071580 |
| chr5 | 127240475 | 127244334 |
| chr6 | 91251858 | 91254469 |
| chr14 | 22454531 | 22458003 |
| chr8 | 115431241 | 115436381 |
| chr8 | 9174627 | 9178261 |
| chr19 | 41858719 | 41861953 |
| chr7 | 82164542 | 82168199 |
| chr20 | 29056197 | 29060445 |
| chr2 | 72624986 | 72627491 |
| chr4 | 35581968 | 35584848 |
| chr14 | 40577321 | 40580168 |
| chr3 | 148081676 | 148085258 |
| chr12 | 78946303 | 78953314 |
| chr10 | 46545061 | 46550241 |
| chr12 | 31451650 | 31453685 |
| chr1 | 217768484 | 217774379 |
| chr14 | 52860461 | 52864445 |
| chr4 | 120567497 | 120570073 |
| chr7 | 78353680 | 78358504 |
| chr14 | 28872843 | 28875171 |
| chr15 | 53105230 | 53108246 |
| chr1 | 78628840 | 78631296 |
| chr15_KI270852v1_ | 250635 | 254088 |
| chr10 | 59170876 | 59173651 |
| chr15 | 38252936 | 38257995 |
| chr14 | 26486787 | 26490071 |
| chr12 | 98501979 | 98504899 |
| chr10 | 38224872 | 38227453 |
| chr6 | 75294844 | 75298888 |
| chr12 | 78528148 | 78530882 |
| chr14 | 26476090 | 26481443 |
| chr2 | 189113683 | 189121302 |
| chr15 | 55740290 | 55742766 |
| chr5 | 50397727 | 50399733 |
| chr10 | 37950821 | 37954616 |
| chr14 | 19338935 | 19342011 |
| chr6 | 55824397 | 55828936 |
| chr14 | 27054620 | 27058037 |
| chr2 | 175395724 | 175399114 |
| chr2 | 38561953 | 38564340 |
| chr10 | 115478549 | 115483975 |

| | | |
|---|---|---|
| chr16 | 54284794 | 54288482 |
| chr1 | 97967893 | 97970825 |
| chr2 | 50222320 | 50224408 |
| chr2 | 165133701 | 165135917 |
| chr20 | 53098120 | 53102370 |
| chr2 | 235495205 | 235497215 |
| chr2 | 195307918 | 195312343 |
| chr14 | 101116455 | 101120942 |
| chr6 | 124095102 | 124097468 |
| chr1 | 211759563 | 211763199 |
| chr1 | 185960543 | 185963302 |
| chr5 | 129026118 | 129030499 |
| chr1 | 92816228 | 92818857 |
| chr7 | 96346084 | 96348300 |
| chr2 | 197514550 | 197516643 |
| chr9 | 66323206 | 66326838 |
| chr15 | 49371084 | 49375602 |
| chr7 | 79429546 | 79431634 |
| chr7 | 90959865 | 90962599 |
| chr14 | 38904916 | 38908068 |
| chr20 | 28588026 | 28590260 |
| chr2 | 50309572 | 50317006 |
| chr5 | 121573490 | 121576676 |
| chr21 | 27243194 | 27245586 |
| chr15 | 73866097 | 73868487 |
| chr12 | 79795837 | 79799207 |
| chr14 | 24162830 | 24165859 |
| chr12 | 17588647 | 17591874 |
| chr15 | 21249832 | 21253586 |
| chr16_KI270728v1_ | 1502863 | 1506670 |
| chr2 | 201376682 | 201378871 |
| chr1 | 143270222 | 143274597 |
| chr14 | 50785182 | 50788523 |
| chr15 | 48152307 | 48154934 |
| chr9 | 64572394 | 64575189 |
| chr14 | 40268100 | 40271074 |
| chr2 | 161354615 | 161356961 |
| chr15 | 58058784 | 58060817 |
| chr10 | 44278866 | 44281924 |
| chr2 | 204886343 | 204888435 |
| chr2 | 51156790 | 51159565 |
| chr7 | 62311284 | 62315551 |
| chr4 | 105101011 | 105103992 |
| chr6 | 71114926 | 71116932 |
| chr7 | 127940089 | 127942714 |
| chr2 | 183284846 | 183287975 |
| chr1 | 90479078 | 90483091 |
| chr2 | 144128027 | 144130646 |
| chr7 | 92054183 | 92060688 |

| | | |
|---|---|---|
| chr20 | 29414567 | 29419021 |
| chr14 | 31171743 | 31174392 |
| chr1 | 215045071 | 215047673 |
| chr10 | 62998135 | 63001110 |
| chr8 | 39891434 | 39894214 |
| chr1 | 97131112 | 97134671 |
| chr13 | 70957954 | 70961615 |
| chr1 | 117046641 | 117049956 |
| chr13 | 57852861 | 57856702 |
| chr14 | 49463068 | 49466064 |
| chr11 | 24575507 | 24578768 |
| chr14 | 19681495 | 19686047 |
| chr1 | 189117367 | 189120028 |
| chr10 | 65154153 | 65157273 |
| chr2 | 78905134 | 78908744 |
| chr12 | 53005792 | 53008076 |
| chr2 | 189099310 | 189105019 |
| chr14 | 28761488 | 28765835 |
| chr12 | 44928136 | 44932142 |
| chr2 | 176117084 | 176119114 |
| chr11 | 95776160 | 95779191 |
| chr1 | 103763542 | 103768368 |
| chr10 | 91906673 | 91910508 |
| chr12 | 57454068 | 57457737 |
| chr1 | 196167947 | 196170171 |
| chr13 | 109790045 | 109793433 |
| chr2 | 177220217 | 177223339 |
| chr12 | 89451337 | 89453786 |
| chr21 | 10474715 | 10480273 |
| chr7 | 82477794 | 82480151 |
| chr12 | 85551558 | 85553674 |
| chr7 | 77022834 | 77026951 |
| chr6 | 72211035 | 72215162 |
| chr5 | 71762526 | 71764579 |
| chr10 | 129430580 | 129433460 |
| chr2 | 164701204 | 164704742 |
| chr2 | 40997041 | 40999484 |
| chr11 | 84058464 | 84060939 |
| chr2 | 63013610 | 63016513 |
| chr8 | 111186324 | 111188416 |
| chr2 | 67424904 | 67427294 |
| chr7 | 95204816 | 95207624 |
| chr14 | 68707304 | 68709355 |
| chr14 | 25237433 | 25240038 |
| chr2 | 97152550 | 97156211 |
| chr10 | 38126122 | 38128968 |
| chr7 | 16855671 | 16858224 |
| chr5 | 11054573 | 11057250 |
| chr7 | 11055857 | 11058295 |

| | | |
|---|---|---|
| chr14 | 36251905 | 36255887 |
| chr12 | 74537096 | 74541711 |
| chr12 | 84759028 | 84761600 |
| chr6 | 133078665 | 133082453 |
| chr5 | 36449042 | 36451125 |
| chr4 | 34789124 | 34791767 |
| chr9 | 65212881 | 65218022 |
| chr2 | 186590690 | 186593425 |
| chr5 | 91635212 | 91639989 |
| chr13 | 86782498 | 86785376 |
| chr5 | 27245970 | 27249608 |
| chr1 | 61227483 | 61231586 |
| chr2 | 186600625 | 186603651 |
| chr5 | 18559394 | 18561548 |
| chr1 | 216251316 | 216254835 |
| chr5 | 27341957 | 27344554 |
| chr2 | 64200962 | 64204039 |
| chr1 | 193181917 | 193185493 |
| chr13 | 105230801 | 105233669 |
| chr7 | 82335710 | 82339366 |
| chr20 | 13399855 | 13403348 |
| chr14 | 29660682 | 29663766 |
| chr6 | 72099613 | 72103973 |
| chr10 | 88506354 | 88509090 |
| chr12 | 92844175 | 92846639 |
| chr11 | 100105806 | 100108291 |
| chr15 | 99868387 | 99870498 |
| chr14 | 19867339 | 19872106 |
| chr1 | 71078988 | 71081961 |
| chr6 | 78526985 | 78529928 |
| chr6 | 64150790 | 64153816 |
| chr5 | 157795616 | 157799220 |
| chr10 | 112947577 | 112951450 |
| chr14 | 19844498 | 19852027 |
| chr11 | 99111726 | 99113740 |
| chr4 | 42278200 | 42280378 |
| chr7 | 48279405 | 48281698 |
| chr2 | 195080398 | 195082620 |
| chr4 | 125614969 | 125617402 |
| chr11 | 25077466 | 25081127 |
| chr7 | 82664496 | 82668305 |
| chr2 | 188270998 | 188274451 |
| chr4 | 45708978 | 45711120 |
| chr14 | 85823005 | 85825224 |
| chr2 | 45609674 | 45613355 |
| chr16 | 78096469 | 78100586 |
| chr9 | 40937279 | 40940073 |
| chr7 | 83575140 | 83581514 |
| chr17 | 61155346 | 61158289 |

| | | |
|---|---|---|
| chr13 | 107141716 | 107146151 |
| chr15_KI270852v1_ | 33406 | 38058 |
| chr1 | 194893814 | 194896457 |
| chr2 | 103244464 | 103246564 |
| chr4 | 35409526 | 35414637 |
| chr1 | 52053795 | 52056697 |
| chr5 | 123833926 | 123836392 |
| chr14 | 41773533 | 41776267 |
| chr11 | 60604944 | 60607245 |
| chr3 | 394145 | 397941 |
| chr4_KI270896v1_a | 289285 | 291991 |
| chr6 | 71957093 | 71960298 |
| chr12 | 53842572 | 53844693 |
| chr12 | 83447634 | 83450592 |
| chr1 | 198274272 | 198277109 |
| chr14 | 48476112 | 48479304 |
| chr6 | 50053945 | 50056540 |
| chr6 | 160973641 | 160976198 |
| chr2 | 194930080 | 194932919 |
| chr1 | 102474242 | 102478912 |
| chr2 | 195254360 | 195257488 |
| chr1 | 191189039 | 191191865 |
| chr7 | 62342184 | 62346371 |
| chr9 | 29847717 | 29850289 |
| chr2 | 187443729 | 187451854 |
| chr20 | 37065060 | 37069143 |
| chr11 | 27360302 | 27365565 |
| chr1 | 96769110 | 96771216 |
| chr10 | 62038439 | 62040781 |
| chr5 | 62237494 | 62240246 |
| chr2 | 45029102 | 45032050 |
| chr6 | 65184589 | 65187650 |
| chr5 | 93068765 | 93071768 |
| chr12 | 65891673 | 65897464 |
| chr1 | 104996349 | 104998565 |
| chr14 | 50668180 | 50670635 |
| chr8 | 14544875 | 14547293 |
| chr1 | 76540900 | 76543188 |
| chr12 | 83755515 | 83758434 |
| chr14 | 88789639 | 88795334 |
| chr14 | 19413741 | 19419444 |
| chr2 | 181286703 | 181290100 |
| chr4 | 89910938 | 89918827 |
| chr14 | 36056504 | 36060698 |
| chr1 | 83865081 | 83867328 |
| chr10 | 63463640 | 63469044 |
| chr5 | 171404921 | 171409763 |
| chr2 | 190934752 | 190937497 |
| chr3 | 80130305 | 80132672 |

| | | |
|---|---|---|
| chr10 | 4426098 | 4432223 |
| chr7 | 81501174 | 81505590 |
| chr8 | 17915616 | 17918576 |
| chr8 | 16537286 | 16539937 |
| chr6 | 94147533 | 94150415 |
| chr1 | 76389637 | 76391967 |
| chr7 | 83287953 | 83291036 |
| chr14 | 35545614 | 35550019 |
| chr2 | 205137933 | 205139950 |
| chr12 | 63431507 | 63433799 |
| chr15 | 54656746 | 54660958 |
| chr2 | 194490052 | 194494679 |
| chr2 | 163625015 | 163629046 |
| chr2 | 187000790 | 187003722 |
| chr1 | 192012639 | 192016737 |
| chr12 | 33715949 | 33718177 |
| chr7 | 83154719 | 83158606 |
| chr13 | 104952769 | 104955832 |
| chr7 | 7092789 | 7094881 |
| chr11 | 55278164 | 55282101 |
| chr7 | 152431257 | 152434216 |
| chr5_GL339449v2_a | 792003 | 795732 |
| chr1 | 104571462 | 104575188 |
| chr10 | 105880522 | 105882908 |
| chr6 | 133068013 | 133074225 |
| chr7 | 79656117 | 79658565 |
| chr14 | 33574542 | 33577738 |
| chr4 | 117513754 | 117516281 |
| chr5 | 28765753 | 28770582 |
| chr10 | 93683447 | 93685733 |
| chr5 | 21026868 | 21029671 |
| chr12 | 22469659 | 22472698 |
| chr7 | 93845003 | 93848447 |
| chr8 | 115408283 | 115414061 |
| chr2 | 145004350 | 145008273 |
| chr10 | 107589452 | 107591929 |
| chr6 | 140085689 | 140088233 |
| chr1 | 81989279 | 81991352 |
| chr3 | 84450298 | 84452410 |
| chr1 | 163319111 | 163323333 |
| chr8 | 123414901 | 123418158 |
| chr1 | 108510993 | 108513243 |
| chr2 | 184153920 | 184156275 |
| chr7 | 77796108 | 77799199 |
| chr14 | 64201005 | 64203041 |
| chr4 | 163920928 | 163923388 |
| chr14 | 19326585 | 19330975 |
| chr14 | 64908111 | 64910715 |
| chr8 | 90644535 | 90646563 |

| | | |
|---|---|---|
| chr21 | 21124817 | 21127205 |
| chr18 | 66535058 | 66537213 |
| chr10 | 95693043 | 95697382 |
| chr10 | 42066064 | 42094548 |
| chr2 | 187390134 | 187392994 |
| chr2 | 209876778 | 209881281 |
| chr1 | 172320185 | 172327358 |
| chr20 | 29077086 | 29081635 |
| chr14 | 86184324 | 86187007 |
| chr1 | 187569155 | 187572789 |
| chr1 | 189278766 | 189283638 |
| chr1 | 76943083 | 76947102 |
| chr4 | 32789432 | 32792419 |
| chr6 | 64615265 | 64617638 |
| chr7 | 89022517 | 89026437 |
| chr7 | 85131907 | 85134804 |
| chr12 | 39618543 | 39620929 |
| chr4 | 62048885 | 62051222 |
| chr14 | 32365946 | 32368027 |
| chr7 | 85888149 | 85893663 |
| chr2 | 55930395 | 55933347 |
| chr2 | 189853942 | 189856448 |
| chr2 | 184699835 | 184702370 |
| chr6 | 94208168 | 94211394 |
| chr8 | 132812251 | 132814748 |
| chr21 | 6638219 | 6641095 |
| chr2 | 100232437 | 100234936 |
| chr18 | 3602271 | 3606909 |
| chr8 | 111253397 | 111256960 |
| chr12 | 84786220 | 84788747 |
| chr7 | 103774888 | 103778465 |
| chr4 | 157353466 | 157355665 |
| chr10 | 101829654 | 101834928 |
| chr15 | 57221579 | 57225619 |
| chr1 | 153668794 | 153672023 |
| chr14 | 88157407 | 88159473 |
| chr11 | 14298675 | 14301417 |
| chr9 | 100493224 | 100495898 |
| chr14 | 85778504 | 85780517 |
| chr14 | 28129082 | 28132011 |
| chr1 | 190169987 | 190172095 |
| chr2 | 180903096 | 180907743 |
| chr14 | 41062805 | 41065279 |
| chr14 | 97623483 | 97625649 |
| chr7 | 91279443 | 91281758 |
| chr4 | 76146186 | 76149179 |
| chr7 | 8583415 | 8586582 |
| chr4 | 125738936 | 125741260 |
| chr20 | 8017485 | 8020513 |

| | | |
|---|---|---|
| chr2 | 83501869 | 83505300 |
| chr5 | 52839801 | 52841963 |
| chr1 | 62081743 | 62084273 |
| chr15 | 92897489 | 92902637 |
| chr9 | 62850573 | 62854375 |
| chr1 | 217681460 | 217684594 |
| chr16 | 76420394 | 76425362 |
| chr20 | 51796920 | 51800978 |
| chr14 | 100952140 | 100956831 |
| chr2 | 58651169 | 58655136 |
| chr14 | 68155700 | 68158762 |
| chr3 | 39149798 | 39151845 |
| chr10 | 50559149 | 50561336 |
| chr5 | 60230372 | 60232648 |
| chr12 | 73919051 | 73923573 |
| chr1 | 195779632 | 195782456 |
| chr10 | 67089911 | 67096210 |
| chr2 | 52157020 | 52159700 |
| chr14 | 100674313 | 100676627 |
| chr1 | 218078852 | 218081445 |
| chr20 | 58902072 | 58905509 |
| chr2 | 182110764 | 182113022 |
| chr14 | 39916674 | 39920881 |
| chr12 | 72575656 | 72578141 |
| chr12 | 57460984 | 57463083 |
| chr2 | 24727069 | 24729172 |
| chr16 | 36139386 | 36143901 |
| chr2 | 166006020 | 166008977 |
| chr11 | 89474393 | 89476541 |
| chr6 | 38024573 | 38026638 |
| chr8 | 136871309 | 136874597 |
| chr6 | 120237375 | 120240816 |
| chr10 | 108154046 | 108156738 |
| chr17 | 65204097 | 65206274 |
| chr1 | 219163470 | 219167733 |
| chr9 | 62362932 | 62368146 |
| chr2 | 50095004 | 50097189 |
| chr2 | 40158190 | 40161040 |
| chr1 | 103204957 | 103213720 |
| chr11 | 26830189 | 26832634 |
| chr1 | 213571861 | 213574748 |
| chr7 | 112449229 | 112452192 |
| chr12 | 65265718 | 65269181 |
| chr10 | 71294782 | 71296975 |
| chr1 | 100894414 | 100899498 |
| chr10 | 56832937 | 56836146 |
| chr12 | 80199911 | 80204094 |
| chr6 | 78659986 | 78663406 |
| chr5 | 173917378 | 173920920 |

| | | |
|---|---|---|
| chr13 | 63027934 | 63032305 |
| chr14 | 28595163 | 28599930 |
| chr14 | 41025834 | 41028882 |
| chr11 | 30017810 | 30020727 |
| chr4 | 90917777 | 90922377 |
| chr2 | 57437981 | 57444133 |
| chr2 | 99965345 | 99967422 |
| chr7 | 88831597 | 88836755 |
| chr20 | 28556575 | 28563785 |
| chr21 | 9906049 | 9910852 |
| chr12 | 89709830 | 89713831 |
| chr17 | 22154731 | 22159297 |
| chr15 | 35371610 | 35374982 |
| chr6 | 124900828 | 124903347 |
| chr2 | 60067340 | 60069812 |
| chr1 | 112702213 | 112704840 |
| chr1 | 67676342 | 67678937 |
| chr12 | 62241172 | 62243444 |
| chr4 | 156806030 | 156810535 |
| chr14 | 101823802 | 101826429 |
| chr2 | 188509894 | 188514127 |
| chr4 | 98285161 | 98287497 |
| chr13 | 67074449 | 67078345 |
| chr14 | 29060504 | 29063688 |
| chr21 | 21540315 | 21543903 |
| chr10 | 51902682 | 51904923 |
| chr6 | 135865771 | 135868256 |
| chr11 | 123579978 | 123583420 |
| chr11 | 91113517 | 91116093 |
| chr17 | 21665683 | 21669299 |
| chr15 | 34496976 | 34499946 |
| chr10 | 112952069 | 112954527 |
| chr9 | 62843320 | 62848285 |
| chr10 | 54888545 | 54893714 |
| chr20 | 29486502 | 29488555 |
| chr16 | 36135699 | 36139059 |
| chr14 | 88535219 | 88537308 |
| chr12 | 62375449 | 62377876 |
| chr2 | 177504440 | 177506535 |
| chr10 | 60070704 | 60073069 |
| chr2 | 72989817 | 72992350 |
| chr9 | 62355140 | 62360830 |
| chr2 | 65811661 | 65813885 |
| chr8 | 115463507 | 115468511 |
| chr1 | 96810139 | 96812369 |
| chr11 | 85365767 | 85368039 |
| chr4 | 49560667 | 49562750 |
| chr3 | 33865301 | 33870702 |
| chr1 | 97298016 | 97300593 |

| | | |
|---|---|---|
| chr10 | 79298309 | 79302430 |
| chr12 | 78599347 | 78601746 |
| chr2 | 161122416 | 161125433 |
| chr12 | 99769906 | 99773833 |
| chr1 | 246367528 | 246369873 |
| chr6 | 74355772 | 74358838 |
| chr6 | 63759069 | 63761929 |
| chr14 | 49422955 | 49425827 |
| chr5 | 161413340 | 161415832 |
| chr8 | 117182336 | 117186869 |
| chr14 | 28138415 | 28140614 |
| chr2 | 91577432 | 91581571 |
| chr4 | 176742288 | 176745615 |
| chr4 | 2959222 | 2963321 |
| chr1 | 161864387 | 161866435 |
| chr14 | 103393040 | 103395972 |
| chr14 | 100694329 | 100698046 |
| chr2 | 66665749 | 66670034 |
| chr2 | 58198621 | 58201240 |
| chr16 | 46406208 | 46411622 |
| chr6 | 114120681 | 114123207 |
| chr2 | 175743273 | 175745773 |
| chr9 | 63891119 | 63894475 |
| chr2 | 58162388 | 58165423 |
| chr11 | 26946470 | 26948631 |
| chr7 | 85122784 | 85126099 |
| chr14 | 46752265 | 46756163 |
| chr2 | 76784443 | 76786635 |
| chr6 | 49966047 | 49969917 |
| chr4 | 175832266 | 175836635 |
| chr1 | 77944950 | 77947886 |
| chr14 | 28366010 | 28372654 |
| chr2 | 231713278 | 231716778 |
| chr10 | 43699788 | 43703971 |
| chr12 | 81086625 | 81089297 |
| chr2 | 57957193 | 57961261 |
| chr2 | 181301712 | 181304719 |
| chr7 | 58049749 | 58067792 |
| chr10 | 65766401 | 65769598 |
| chr4 | 34238756 | 34241275 |
| chr1 | 101912561 | 101917758 |
| chr1 | 179044819 | 179046840 |
| chr12 | 77906451 | 77908821 |
| chr4 | 156632912 | 156637612 |
| chr1 | 189658895 | 189662549 |
| chr2 | 1664011 | 1666352 |
| chr11 | 111836315 | 111839301 |
| chr5 | 151756985 | 151760216 |
| chr17 | 22105501 | 22109516 |

| | | |
|---|---|---|
| chr8 | 112359618 | 112362613 |
| chr12 | 26191746 | 26194520 |
| chr6 | 69390026 | 69392657 |
| chr5 | 92393920 | 92396052 |
| chr5 | 107369021 | 107372168 |
| chr7 | 10797988 | 10800118 |
| chr12 | 45645197 | 45647914 |
| chr1 | 171089645 | 171091862 |
| chr14 | 34433836 | 34437640 |
| chr7 | 58032131 | 58041324 |
| chr2 | 45576545 | 45579743 |
| chr7 | 81619020 | 81622345 |
| chr10 | 65296955 | 65300361 |
| chr9 | 67867666 | 67871828 |
| chr17 | 65947444 | 65949983 |
| chr14 | 101133804 | 101136635 |
| chr1 | 106608100 | 106610148 |
| chr7 | 86641247 | 86645621 |
| chr1 | 99889630 | 99893648 |
| chr11 | 123722654 | 123725066 |
| chr10 | 102916746 | 102920048 |
| chr1 | 218864608 | 218868958 |
| chr3 | 169771086 | 169775686 |
| chr8 | 84409290 | 84411717 |
| chr13 | 77642699 | 77645821 |
| chr2 | 162703624 | 162706453 |
| chr10 | 57730044 | 57732370 |
| chr3 | 81848935 | 81854683 |
| chr16 | 81938428 | 81941223 |
| chr1 | 190476354 | 190479423 |
| chr2 | 179986482 | 179989320 |
| chr4 | 35127609 | 35129732 |
| chr12 | 85151882 | 85156343 |
| chr1 | 95935658 | 95938824 |
| chr6 | 62360569 | 62362956 |
| chr7 | 80473007 | 80478910 |
| chr20 | 30388003 | 30392422 |
| chr7 | 93223192 | 93227521 |
| chr14 | 46821760 | 46826350 |
| chr4 | 73178544 | 73180585 |
| chr5_KI270897v1_a | 783057 | 787269 |
| chr11 | 34925912 | 34929174 |
| chr1 | 109188178 | 109190999 |
| chr6 | 2099932 | 2103044 |
| chr10 | 74432958 | 74435528 |
| chr7 | 152380534 | 152384067 |
| chr3 | 117727345 | 117729650 |
| chr3 | 78910075 | 78912907 |
| chr15 | 57339723 | 57342381 |

| | | |
|---|---|---|
| chrUn_GL000218v1 | 156077 | 160300 |
| chr10 | 48534424 | 48537059 |
| chr11 | 107432826 | 107435958 |
| chr10 | 89938599 | 89941207 |
| chr1 | 106565966 | 106569731 |
| chr8 | 72212792 | 72215474 |
| chr2 | 102422356 | 102426054 |
| chr2 | 58181378 | 58185961 |
| chr8 | 60613712 | 60617131 |
| chr16 | 56846692 | 56849113 |
| chr13 | 67193246 | 67195997 |
| chr2 | 58644122 | 58646205 |
| chr2 | 41655622 | 41658301 |
| chr5 | 24713188 | 24715222 |
| chr7 | 62390450 | 62392600 |
| chr14 | 28604763 | 28609903 |
| chr2 | 77057417 | 77060665 |
| chr2 | 186748505 | 186751245 |
| chr7 | 78468134 | 78471423 |
| chr1 | 195961419 | 195964007 |
| chr6 | 49199415 | 49201887 |
| chr5 | 89687916 | 89692287 |
| chr2 | 159871370 | 159876546 |
| chr10 | 84521051 | 84524325 |
| chr15 | 54011632 | 54013922 |
| chr2 | 159164875 | 159167667 |
| chr10 | 90274716 | 90277063 |
| chr2 | 179402225 | 179405161 |
| chr2 | 44641810 | 44644901 |
| chr20 | 15989198 | 15991634 |
| chr6 | 54090938 | 54094766 |
| chr6 | 20954464 | 20957370 |
| chr11 | 25355867 | 25358027 |
| chr7 | 40406773 | 40409201 |
| chr10 | 131869715 | 131872733 |
| chr10 | 50859927 | 50863543 |
| chr15 | 37820033 | 37822293 |
| chr12 | 39614393 | 39618087 |
| chr14 | 35955225 | 35958076 |
| chr12 | 59962422 | 59965604 |
| chr9 | 63998399 | 64001620 |
| chr9 | 62296292 | 62300178 |
| chr7 | 54060179 | 54063785 |
| chr1_KI270713v1_r | 2174 | 5606 |
| chr1 | 94225240 | 94228686 |
| chr2 | 195713045 | 195717026 |
| chr2 | 69576217 | 69578928 |
| chr20 | 34475578 | 34478670 |
| chr10 | 35643640 | 35646058 |

| | | |
|---|---|---|
| chr2 | 162142965 | 162145935 |
| chr7 | 82271852 | 82276310 |
| chr1 | 103021823 | 103030240 |
| chr7 | 79831214 | 79833750 |
| chr5 | 118362685 | 118364905 |
| chr12 | 42393212 | 42396120 |
| chr2 | 212352527 | 212357501 |
| chr8 | 89270822 | 89274788 |
| chr10 | 4723359 | 4726960 |
| chr1 | 97929748 | 97933888 |
| chr2 | 189167255 | 189170862 |
| chr2 | 187978889 | 187981276 |
| chr22 | 11016746 | 11021768 |
| chr2 | 63541224 | 63544996 |
| chr14 | 45135188 | 45138161 |
| chr21 | 16569055 | 16573522 |
| chr12 | 83289798 | 83292216 |
| chr2 | 184185040 | 184188708 |
| chr21 | 20968101 | 20973662 |
| chr1 | 5649792 | 5651838 |
| chr2 | 41782273 | 41784789 |
| chr14 | 66398832 | 66401723 |
| chr10 | 57878536 | 57881652 |
| chr12 | 84547828 | 84553740 |
| chr5 | 122154817 | 122157510 |
| chr14 | 36944781 | 36949692 |
| chr14 | 19791476 | 19793569 |
| chr2 | 209478526 | 209480929 |
| chr5 | 117682108 | 117685748 |
| chr10 | 115813838 | 115817285 |
| chr10 | 117271754 | 117273895 |
| chr7 | 94971925 | 94974609 |
| chr1 | 172913513 | 172915556 |
| chr12 | 121793625 | 121796257 |
| chr14 | 70058944 | 70061152 |
| chr2 | 191018058 | 191020782 |
| chr3 | 25425707 | 25434018 |
| chr10 | 58191166 | 58197136 |
| chr7 | 89159235 | 89162498 |
| chr14 | 85563881 | 85567753 |
| chr2 | 172387042 | 172389244 |
| chr9 | 65967805 | 65972357 |
| chr7 | 94067135 | 94071021 |
| chr13 | 71787177 | 71791267 |
| chr1 | 186285841 | 186288281 |
| chrUn_GL000220v1 | 4184 | 9829 |
| chr7 | 126569902 | 126571931 |
| chr14 | 60288307 | 60292931 |
| chr17 | 742194 | 745529 |

| | | |
|---|---|---|
| chr10 | 61475091 | 61477792 |
| chr6 | 120761243 | 120763279 |
| chr5 | 41123369 | 41125868 |
| chr6 | 5465083 | 5467514 |
| chr7 | 82406370 | 82408422 |
| chr1 | 144905544 | 144908818 |
| chr1 | 66168628 | 66171054 |
| chr8 | 76202259 | 76205733 |
| chr2 | 192682591 | 192685492 |
| chr20 | 25920554 | 25926282 |
| chr4 | 130978748 | 130981075 |
| chr16 | 46401274 | 46406172 |
| chr8 | 113835435 | 113838777 |
| chr13 | 80331527 | 80334025 |
| chr1 | 103623236 | 103626274 |
| chr15 | 20356343 | 20362287 |
| chr2 | 80992628 | 80996071 |
| chr10 | 87324132 | 87327051 |
| chr2 | 178614998 | 178620285 |
| chr9 | 20385866 | 20388332 |
| chr2 | 50680174 | 50683224 |
| chr7 | 53861944 | 53865756 |
| chrY | 11680658 | 11686363 |
| chr11 | 49549035 | 49551596 |
| chr3 | 77028370 | 77031818 |
| chr10 | 52817744 | 52821772 |
| chr17 | 56834344 | 56836595 |
| chr14 | 59722734 | 59725316 |
| chr10 | 17045911 | 17049880 |
| chr7 | 58043322 | 58049593 |
| chr14 | 47207928 | 47212839 |
| chr17_KI270908v1_ | 251063 | 253324 |
| chr2 | 192330329 | 192332932 |
| chr2 | 63216953 | 63220558 |
| chr4 | 47218957 | 47221352 |
| chr12 | 23761537 | 23764169 |
| chr11 | 105280911 | 105284249 |
| chr2 | 76603437 | 76606380 |
| chr3 | 29345120 | 29348153 |
| chr6 | 64139025 | 64142519 |
| chr10 | 25243401 | 25246447 |
| chr1 | 148627487 | 148631762 |
| chr14 | 77551652 | 77553754 |
| chr7 | 84012345 | 84014925 |
| chr5 | 166330628 | 166333402 |
| chr10 | 93478956 | 93482651 |
| chr18 | 71510336 | 71512551 |
| chr1 | 198924435 | 198930883 |
| chr2 | 50183931 | 50187509 |

| | | |
|---|---|---|
| chr2 | 66511850 | 66514430 |
| chr1 | 198357424 | 198361584 |
| chr1 | 96035905 | 96038872 |
| chr8 | 101539960 | 101542269 |
| chr14 | 41445917 | 41448131 |
| chr10 | 55395548 | 55399591 |
| chr7 | 80483966 | 80490048 |
| chr7 | 88349879 | 88351977 |
| chr1 | 87082156 | 87087654 |
| chr7 | 82885729 | 82888106 |
| chr14 | 100972651 | 100976994 |
| chr10 | 116682677 | 116685067 |
| chr15 | 37769806 | 37776020 |
| chr16 | 36167013 | 36191199 |
| chr2 | 39066295 | 39069271 |
| chr1 | 103572211 | 103575753 |
| chr8 | 76532045 | 76534288 |
| chr3 | 80183053 | 80187036 |
| chr5 | 19778782 | 19781774 |
| chr3 | 22938987 | 22941493 |
| chr16 | 79593635 | 79596833 |
| chr2 | 189080433 | 189084271 |
| chr7 | 80508466 | 80514659 |
| chr7 | 81095606 | 81099464 |
| chr14 | 65411079 | 65413869 |
| chr16_KI270728v1_ | 934009 | 938009 |
| chr7 | 81992856 | 81997325 |
| chr9 | 68972412 | 68975053 |
| chr12 | 84027049 | 84030750 |
| chr8 | 106725235 | 106730120 |
| chr19 | 45642588 | 45646024 |
| chr5 | 30581236 | 30584746 |
| chr6 | 133043695 | 133048312 |
| chr6 | 124959488 | 124962116 |
| chr12 | 103288769 | 103290938 |
| chr1 | 195703497 | 195705660 |
| chr2 | 76273822 | 76277255 |
| chr7 | 84185684 | 84197553 |
| chr10 | 50188237 | 50191807 |
| chr6 | 120444855 | 120446970 |
| chr5 | 27017981 | 27022079 |
| chr16 | 61639256 | 61641332 |
| chr10 | 115468530 | 115471202 |
| chr12 | 18317768 | 18320895 |
| chr17 | 79385163 | 79387195 |
| chr2 | 173400019 | 173402282 |
| chr11 | 104186430 | 104192614 |
| chr14 | 27333973 | 27338556 |
| chr11 | 56461685 | 56464395 |

| | | |
|---|---|---|
| chr6 | 49422664 | 49425235 |
| chr6 | 108419411 | 108423728 |
| chr15 | 35920017 | 35922631 |
| chr6 | 85460197 | 85462601 |
| chr14 | 45201973 | 45205234 |
| chr14 | 38721059 | 38724313 |
| chr10 | 91818477 | 91821099 |
| chr2 | 213004807 | 213010556 |
| chr2 | 185539174 | 185542599 |
| chr2 | 177153431 | 177156176 |
| chr14 | 89198703 | 89200708 |
| chr11 | 103119579 | 103124038 |
| chr7 | 77908204 | 77912958 |
| chr14 | 28768457 | 28772907 |
| chr7 | 86913515 | 86917288 |
| chr1 | 103714452 | 103717049 |
| chr3 | 101723945 | 101727081 |
| chr12 | 750435 | 753956 |
| chr22 | 10761669 | 10763775 |
| chr2 | 74505041 | 74509329 |
| chr10 | 46737259 | 46741914 |
| chr3 | 3845285 | 3848942 |
| chr6 | 56753432 | 56755858 |
| chr6 | 98018333 | 98021342 |
| chr11 | 91382618 | 91385322 |
| chr1 | 102996969 | 103005321 |
| chr10 | 110284117 | 110286762 |
| chr10 | 65129563 | 65131763 |
| chr6 | 133372137 | 133377207 |
| chr8 | 108560070 | 108562961 |
| chr2 | 83924515 | 83927775 |
| chr7 | 79418477 | 79424519 |
| chr2 | 105522307 | 105524760 |
| chr2 | 184687046 | 184690355 |
| chr8 | 106181657 | 106185262 |
| chr11 | 25126521 | 25128689 |
| chr1 | 99727862 | 99730849 |
| chr10 | 88282642 | 88285915 |
| chr9 | 63543800 | 63546742 |
| chr4 | 31508927 | 31510970 |
| chr2 | 56036797 | 56039138 |
| chr11 | 23852760 | 23855598 |
| chr1 | 64834285 | 64837150 |
| chr1 | 148453779 | 148459863 |
| chr12 | 100046506 | 100051031 |
| chr4 | 23133326 | 23137114 |
| chr9 | 66254944 | 66258052 |
| chr14 | 96342409 | 96345967 |
| chr2 | 94960495 | 94964636 |

| | | |
|---|---|---|
| chr7 | 80628022 | 80633636 |
| chr2 | 140705093 | 140707268 |
| chr12 | 85251954 | 85257474 |
| chr5 | 170916782 | 170919994 |
| chr10 | 46719982 | 46724301 |
| chr5 | 180258951 | 180261754 |
| chr1 | 102774880 | 102778602 |
| chr5 | 108300048 | 108303008 |
| chr2 | 188937077 | 188942503 |
| chr1 | 103226948 | 103230460 |
| chr4 | 33246806 | 33250236 |
| chr4 | 115642115 | 115645748 |
| chr4 | 157345753 | 157349141 |
| chr2 | 62850010 | 62853317 |
| chr14 | 40622061 | 40627257 |
| chr2 | 185506796 | 185508972 |
| chr5 | 27013379 | 27016653 |
| chr7 | 61045155 | 61058093 |
| chr1 | 88263688 | 88266545 |
| chr12 | 97757393 | 97759967 |
| chr1 | 93331074 | 93333532 |
| chr22 | 12625523 | 12628813 |
| chr10 | 131270922 | 131273364 |
| chr12 | 33439670 | 33442187 |
| chr11 | 26122236 | 26124313 |
| chr7 | 84081979 | 84087603 |
| chr1 | 97742935 | 97745595 |
| chr3 | 175961045 | 175963471 |
| chr7 | 87624895 | 87633079 |
| chr14 | 38228539 | 38231333 |
| chr9 | 73149281 | 73154200 |
| chr2 | 214411664 | 214413692 |
| chr11 | 786949 | 789415 |
| chr14 | 23783026 | 23785370 |
| chr2 | 188479819 | 188485480 |
| chr9 | 66112486 | 66115641 |
| chr10 | 63519409 | 63522925 |
| chr20 | 13490689 | 13493480 |
| chr2 | 184887757 | 184892201 |
| chr14 | 23305791 | 23308295 |
| chr7 | 61041066 | 61045121 |
| chr6 | 19668678 | 19671861 |
| chr13 | 107162672 | 107165966 |
| chr5 | 28213834 | 28216127 |
| chr14 | 62401303 | 62403958 |
| chr2 | 203684224 | 203686620 |
| chr1 | 2173598 | 2175831 |
| chr10 | 109069409 | 109072669 |
| chr10 | 49850763 | 49853950 |

| | | |
|---|---|---|
| chr7 | 87738123 | 87741265 |
| chr7 | 86014199 | 86018199 |
| chr14 | 42372827 | 42375056 |
| chr4 | 158019031 | 158021305 |
| chr14 | 81527168 | 81530284 |
| chrUn_GL000219v1 | 86225 | 88735 |
| chr1 | 89297950 | 89300197 |
| chr12 | 85970016 | 85973534 |
| chr2 | 68450972 | 68453316 |
| chr2 | 197704663 | 197708486 |
| chr7 | 85126114 | 85130134 |
| chr7 | 61058404 | 61069155 |
| chr11 | 105779329 | 105782290 |
| chr15 | 40463965 | 40466625 |
| chr11 | 23681135 | 23684964 |
| chr11 | 24525879 | 24530188 |
| chr5 | 39861219 | 39863436 |
| chr8 | 115593006 | 115598492 |
| chr16 | 83014851 | 83018300 |
| chr11 | 22670729 | 22675230 |
| chr6 | 101908719 | 101911532 |
| chr2 | 74528257 | 74531114 |
| chr6 | 137871314 | 137874598 |
| chr2 | 189163484 | 189167100 |
| chr8 | 115567900 | 115571773 |
| chr6 | 79947246 | 79949850 |
| chr10 | 74955957 | 74964258 |
| chr1 | 192453139 | 192457190 |
| chr10 | 44985009 | 44987373 |
| chr7 | 80452040 | 80454360 |
| chr14 | 88421988 | 88424540 |
| chr6 | 94188330 | 94191150 |
| chr5 | 94473261 | 94475857 |
| chr8 | 75063307 | 75065554 |
| chrY | 11691969 | 11695521 |
| chr12 | 100218513 | 100222013 |
| chr4 | 135401600 | 135404409 |
| chr7 | 89144790 | 89147009 |
| chr1 | 61101519 | 61103724 |
| chr10 | 49165104 | 49168638 |
| chr18 | 28704087 | 28706215 |
| chr4 | 115119213 | 115122654 |
| chr1 | 114392700 | 114396015 |
| chr5 | 148418186 | 148420537 |
| chr15 | 20667367 | 20675495 |
| chr10 | 59232114 | 59237848 |
| chr3 | 149966103 | 149970605 |
| chr7 | 85672594 | 85675320 |
| chr2 | 163714979 | 163722625 |

| | | |
|---|---|---|
| chr2 | 103261801 | 103263880 |
| chr14 | 35406268 | 35408803 |
| chr2 | 60336285 | 60339157 |
| chr14 | 58826346 | 58828612 |
| chr2 | 162188052 | 162192197 |
| chr2 | 182824961 | 182829066 |
| chr14 | 100240873 | 100243876 |
| chr7 | 85019512 | 85025774 |
| chr17 | 58005071 | 58008177 |
| chr6 | 45448470 | 45454129 |
| chr14 | 37137170 | 37141783 |
| chr1 | 84995045 | 84997960 |
| chr12 | 127068873 | 127071225 |
| chr10 | 67778161 | 67781413 |
| chr2 | 154089761 | 154092284 |
| chr1 | 97829741 | 97832942 |
| chr20 | 16049847 | 16054003 |
| chr1 | 86112089 | 86115353 |
| chr14_KI270847v1_ | 298883 | 301687 |
| chr14 | 19390472 | 19394232 |
| chr5 | 108135492 | 108137903 |
| chr7 | 81953212 | 81959006 |
| chr10 | 42312482 | 42317629 |
| chr5 | 93596638 | 93604399 |
| chr2 | 80669901 | 80671985 |
| chr6 | 80282346 | 80286304 |
| chr12 | 88637714 | 88642055 |
| chr8 | 76402154 | 76410342 |
| chr11 | 27053340 | 27055890 |
| chr1 | 143229597 | 143233903 |
| chr4 | 126782168 | 126785841 |
| chr4 | 32467591 | 32469983 |
| chr12 | 83030490 | 83032940 |
| chr2 | 40619718 | 40621781 |
| chr14 | 63283191 | 63285906 |
| chr12 | 43422511 | 43429163 |
| chr21 | 10355611 | 10359341 |
| chr6 | 93044454 | 93048671 |
| chr13 | 86384437 | 86386609 |
| chr16 | 73490751 | 73493312 |
| chr13 | 106098139 | 106101943 |
| chr1 | 90572350 | 90574661 |
| chr7 | 79385756 | 79391748 |
| chr10 | 55148965 | 55154755 |
| chr4 | 156517708 | 156519896 |
| chr16 | 34948714 | 34962738 |
| chr22 | 15901810 | 15905403 |
| chr1 | 68195985 | 68198714 |
| chr5 | 89384517 | 89386712 |

| | | |
|---|---|---|
| chr5 | 116663840 | 116666772 |
| chr14 | 47096726 | 47099060 |
| chr13 | 87274875 | 87277550 |
| chr16 | 56449485 | 56451964 |
| chr1 | 197694511 | 197697148 |
| chr21 | 10012505 | 10017124 |
| chr6 | 45736920 | 45740699 |
| chr7 | 78760688 | 78765551 |
| chr6 | 27890882 | 27893826 |
| chr2 | 238105779 | 238108023 |
| chr2 | 40644351 | 40647754 |
| chr10 | 68230192 | 68233350 |
| chr7 | 82294315 | 82299706 |
| chr10 | 56160397 | 56162527 |
| chr1 | 143189510 | 143198389 |
| chrUn_GL000218v1 | 140173 | 142312 |
| chr11 | 31818201 | 31821277 |
| chr10 | 47633065 | 47640951 |
| chr14 | 51092838 | 51097249 |
| chr2 | 99253392 | 99256778 |
| chr12 | 59068309 | 59073286 |
| chr2 | 113586497 | 113589286 |
| chr12 | 99569714 | 99573281 |
| chr7 | 82846345 | 82850287 |
| chr14 | 40521923 | 40525504 |
| chr14 | 85416268 | 85420007 |
| chr8 | 113929731 | 113932922 |
| chr7 | 83862691 | 83867288 |
| chr2 | 194742124 | 194745011 |
| chr14 | 37560144 | 37563417 |
| chr11 | 59556830 | 59559271 |
| chr1 | 148443220 | 148447344 |
| chr11 | 49331134 | 49334695 |
| chr5 | 92376870 | 92379666 |
| chr10 | 89081830 | 89085799 |
| chr8 | 115612863 | 115618902 |
| chr7 | 79514981 | 79517222 |
| chr1 | 192228050 | 192231217 |
| chr7 | 93259190 | 93262530 |
| chr7 | 85341103 | 85345088 |
| chr7 | 83648523 | 83651608 |
| chr12 | 79933725 | 79937347 |
| chr12 | 76353069 | 76355653 |
| chr2 | 34342893 | 34346924 |
| chr5 | 27701633 | 27706144 |
| chr3 | 35779643 | 35782023 |
| chr10 | 42216408 | 42218522 |
| chr10 | 45115495 | 45117994 |
| chr1 | 179880721 | 179884177 |

| | | |
|---|---|---|
| chr14 | 29188415 | 29191363 |
| chr22 | 10704613 | 10707429 |
| chr1 | 170054725 | 170057970 |
| chr10 | 73447100 | 73449243 |
| chr4 | 130008245 | 130011189 |
| chr3 | 110575006 | 110577347 |
| chr1 | 222311935 | 222316436 |
| chr10 | 110963814 | 110967254 |
| chr6 | 76467641 | 76469846 |
| chr15 | 56642073 | 56645302 |
| chr11 | 126652571 | 126655086 |
| chr11 | 76834262 | 76836746 |
| chr14 | 30128472 | 30130973 |
| chr2 | 183405194 | 183408259 |
| chr6 | 106052930 | 106055307 |
| chr15 | 48134485 | 48140785 |
| chr11 | 18527294 | 18530271 |
| chr22 | 15759976 | 15764927 |
| chr14 | 100967707 | 100971415 |
| chr8 | 58889627 | 58891968 |
| chr20 | 53212872 | 53215800 |
| chr20 | 57342206 | 57345658 |
| chr3 | 177108748 | 177110926 |
| chr2 | 83473056 | 83476753 |
| chr14 | 62602657 | 62605725 |
| chr3 | 174244411 | 174246843 |
| chr2 | 169637932 | 169642062 |
| chr2 | 183310242 | 183312283 |
| chr14 | 19873836 | 19882510 |
| chr8 | 90495489 | 90497992 |
| chr12 | 98880734 | 98883705 |
| chr7 | 95920261 | 95924269 |
| chr2 | 164891853 | 164896888 |
| chr14 | 26841240 | 26846487 |
| chr10 | 88756715 | 88758780 |
| chr2 | 144833768 | 144838202 |
| chr12 | 85241414 | 85246635 |
| chr14 | 49451797 | 49454271 |
| chr1 | 189618454 | 189621203 |
| chr12 | 84092728 | 84096243 |
| chr20 | 14489642 | 14492772 |
| chr10 | 128691995 | 128694635 |
| chr21 | 10613693 | 10617042 |
| chr1 | 145091764 | 145095936 |
| chr7 | 85217568 | 85221292 |
| chr3 | 85499594 | 85504366 |
| chr2 | 66583043 | 66587348 |
| chr12 | 81440840 | 81448422 |
| chr7 | 85771947 | 85776684 |

| | | |
|---|---|---|
| chr20 | 24406994 | 24409921 |
| chr14 | 28218331 | 28220779 |
| chr2 | 71148482 | 71151093 |
| chr2 | 39661492 | 39664485 |
| chr7 | 80803354 | 80809210 |
| chr3 | 73041276 | 73043320 |
| chr4 | 20505688 | 20510464 |
| chr5 | 19591037 | 19593089 |
| chr4 | 107348296 | 107351094 |
| chr7 | 85093495 | 85098882 |
| chr14 | 31386434 | 31389949 |
| chr5 | 91382756 | 91387401 |
| chr2 | 161316180 | 161321078 |
| chr8 | 90120788 | 90123372 |
| chr11 | 65504842 | 65509221 |
| chr10 | 81316712 | 81320647 |
| chr17 | 21836353 | 21838775 |
| chr13 | 104652565 | 104655528 |
| chr12 | 73389869 | 73392836 |
| chr4 | 93062093 | 93066273 |
| chr14 | 49283465 | 49287267 |
| chr3 | 153321030 | 153323155 |
| chr4 | 175270293 | 175273265 |
| chr2 | 193803771 | 193806621 |
| chr20 | 51957445 | 51964153 |
| chr1 | 215130162 | 215134505 |
| chr10 | 66141273 | 66144855 |
| chr11 | 56447837 | 56449881 |
| chr7 | 85320826 | 85327211 |
| chr2 | 176128766 | 176130994 |
| chr14 | 41399791 | 41403324 |
| chr14 | 41601817 | 41605715 |
| chr2 | 180981143 | 180984529 |
| chr12 | 76084076 | 76086830 |
| chr1 | 102917660 | 102928209 |
| chr4 | 175808090 | 175814552 |
| chr9 | 63874303 | 63877495 |
| chr2 | 95532325 | 95536268 |
| chr1 | 154661020 | 154663334 |
| chr22 | 15862353 | 15867398 |
| chr2 | 97180213 | 97183255 |
| chr11 | 38648027 | 38651954 |
| chr2 | 162510509 | 162514205 |
| chr12 | 15323146 | 15327552 |
| chr7 | 82621591 | 82626629 |
| chr11 | 105278097 | 105280824 |
| chr9 | 62326468 | 62330001 |
| chr12 | 91000398 | 91004405 |
| chr16 | 70242562 | 70248141 |

| | | |
|---|---|---|
| chr10 | 79287759 | 79290350 |
| chr2 | 102388329 | 102390957 |
| chr14 | 100949602 | 100952116 |
| chr7 | 82977441 | 82979949 |
| chr10 | 60171201 | 60174675 |
| chr1 | 199706462 | 199708744 |
| chr10 | 65306110 | 65309925 |
| chr10 | 2788704 | 2791193 |
| chr2 | 193378389 | 193380820 |
| chr21 | 10009244 | 10012481 |
| chr16_KI270728v1_ | 1830925 | 1836088 |
| chr1 | 217507754 | 217510280 |
| chr20 | 64084041 | 64086537 |
| chr10 | 90679388 | 90682441 |
| chr17 | 26789464 | 26800590 |
| chr2 | 194562322 | 194564720 |
| chr10 | 106966227 | 106968918 |
| chr10 | 45166988 | 45171671 |
| chr1 | 98393218 | 98399873 |
| chr2 | 184580876 | 184583533 |
| chr6 | 92916192 | 92918812 |
| chr14 | 86652631 | 86657500 |
| chr13 | 40612537 | 40614555 |
| chr1 | 105256195 | 105259482 |
| chr1 | 197145244 | 197148403 |
| chr12 | 72312426 | 72315991 |
| chr13 | 35163653 | 35167167 |
| chr13 | 62363489 | 62366051 |
| chr11 | 26186904 | 26191040 |
| chr3 | 82984912 | 82987838 |
| chr10 | 101798611 | 101800811 |
| chr17 | 64858342 | 64860543 |
| chr21 | 15869621 | 15871791 |
| chr3 | 147415531 | 147420059 |
| chr7 | 66996258 | 66999277 |
| chr2 | 226045482 | 226053412 |
| chr16 | 77189515 | 77192935 |
| chr2 | 56299619 | 56303239 |
| chr14 | 29464551 | 29467959 |
| chr7 | 88696695 | 88700582 |
| chr14 | 26590025 | 26597336 |
| chr14 | 18654678 | 18658639 |
| chr14 | 31962146 | 31965806 |
| chr14 | 40092191 | 40095450 |
| chr1 | 95115965 | 95121247 |
| chr2 | 226183388 | 226193170 |
| chr14 | 36739789 | 36743587 |
| chr16 | 64785013 | 64788080 |
| chr20 | 30889473 | 30896549 |

| | | |
|---|---|---|
| chr1 | 96963053 | 96969079 |
| chr8 | 98640178 | 98642493 |
| chr2 | 72179058 | 72181146 |
| chr11 | 83551006 | 83553040 |
| chr15 | 81474434 | 81477288 |
| chr2 | 80351831 | 80353890 |
| chr6 | 96520801 | 96525007 |
| chr6 | 129646710 | 129649256 |
| chr11 | 73659955 | 73662313 |
| chr2 | 178852047 | 178854604 |
| chr2 | 184619436 | 184625516 |
| chr3 | 130110104 | 130112498 |
| chr16 | 34708749 | 34713378 |
| chr4 | 126699529 | 126703333 |
| chr6 | 78982130 | 78988082 |
| chr10 | 55549518 | 55553756 |
| chr12 | 55217521 | 55220614 |
| chr5_KI270897v1_a | 7095 | 9849 |
| chr8 | 99527667 | 99530610 |
| chr2 | 199310276 | 199312925 |
| chr10 | 117167030 | 117170746 |
| chr8 | 106344833 | 106348093 |
| chr13 | 67227786 | 67235146 |
| chr14 | 29684214 | 29688214 |
| chr2 | 175945337 | 175949146 |
| chr7 | 80015861 | 80019902 |
| chr1 | 98335148 | 98337585 |
| chr10 | 94985109 | 94989644 |
| chr6 | 122178591 | 122182506 |
| chr2 | 161278570 | 161282319 |
| chr10 | 51801149 | 51805761 |
| chr9 | 61720312 | 61722834 |
| chr15 | 96328051 | 96331180 |
| chr12 | 78404120 | 78406878 |
| chr14 | 20629744 | 20634191 |
| chr6 | 64478008 | 64482547 |
| chr16_KI270853v1_ | 1019947 | 1026310 |
| chr8 | 116184406 | 116187099 |
| chr10 | 61316020 | 61319213 |
| chr1 | 218691220 | 218695369 |
| chr1 | 96493311 | 96495801 |
| chr16 | 34736311 | 34744682 |
| chr10 | 67098240 | 67102180 |
| chr4 | 32974417 | 32978189 |
| chr7 | 84198757 | 84202111 |
| chr5 | 125610540 | 125615042 |
| chr9 | 10108851 | 10111290 |
| chr16 | 53203094 | 53205215 |
| chr14 | 46905112 | 46908082 |

| | | |
|---|---|---|
| chr1 | 91519795 | 91522066 |
| chr12 | 90959172 | 90962011 |
| chr14 | 33536981 | 33541183 |
| chr15_KI270905v1_ | 797297 | 802397 |
| chr8 | 107994117 | 107996349 |
| chr14 | 32736322 | 32738787 |
| chr10 | 65368735 | 65372531 |
| chr20 | 57714692 | 57717218 |
| chr1 | 185871187 | 185874504 |
| chr2 | 213036114 | 213039806 |
| chr1 | 99467663 | 99470640 |
| chr2 | 42163434 | 42166131 |
| chr1 | 100975899 | 100979385 |
| chr14 | 79910742 | 79913058 |
| chr7 | 83804289 | 83807157 |
| chr1 | 103007051 | 103014260 |
| chr13 | 84070750 | 84073200 |
| chr1 | 98367533 | 98370986 |
| chr2 | 20347747 | 20351338 |
| chr5 | 27029565 | 27033842 |
| chr2 | 59002962 | 59008035 |
| chr6 | 77934166 | 77936784 |
| chr7 | 79548891 | 79553899 |
| chr11 | 107044659 | 107047596 |
| chr7 | 100952356 | 100959541 |
| chr7 | 85052594 | 85057951 |
| chr8 | 115415663 | 115420350 |
| chr7 | 11101583 | 11107849 |
| chr22 | 12306206 | 12310493 |
| chr6 | 133183467 | 133187303 |
| chr10 | 87001305 | 87004926 |
| chr2 | 227490641 | 227492673 |
| chr2 | 161407602 | 161414372 |
| chr6 | 85674720 | 85678216 |
| chr14 | 45728353 | 45731108 |
| chr10 | 62980976 | 62983133 |
| chr11 | 22733543 | 22737839 |
| chr6 | 12577415 | 12579632 |
| chr4 | 146915394 | 146917981 |
| chr10 | 60067152 | 60070615 |
| chr18 | 68837388 | 68840560 |
| chr14 | 59333099 | 59335512 |
| chr2 | 163933046 | 163935618 |
| chr6 | 84754893 | 84759387 |
| chr5 | 43481748 | 43485448 |
| chr11 | 107137722 | 107141849 |
| chr4 | 176926927 | 176929117 |
| chr2 | 161867288 | 161870872 |
| chr11 | 23819391 | 23821618 |

| | | |
|---|---|---|
| chr10 | 75410869 | 75413932 |
| chr7 | 79288115 | 79291162 |
| chr14 | 19744734 | 19749051 |
| chr1 | 196353485 | 196356934 |
| chr20 | 54270964 | 54274266 |
| chr1 | 187932696 | 187936272 |
| chr1 | 234604553 | 234608421 |
| chr10 | 79516227 | 79518814 |
| chr10 | 37703456 | 37706651 |
| chr14 | 28962967 | 28967441 |
| chr6 | 27753156 | 27755890 |
| chr14 | 32383719 | 32385796 |
| chr1 | 93843897 | 93846867 |
| chr12 | 24172590 | 24174951 |
| chr2 | 69327985 | 69332376 |
| chr8_KI270813v1_a | 261842 | 269115 |
| chr3 | 168837340 | 168839881 |
| chr4 | 107148971 | 107151438 |
| chr1 | 148632263 | 148637914 |
| chr4 | 102303092 | 102305717 |
| chr14 | 50827838 | 50831548 |
| chr14 | 30154219 | 30159392 |
| chr4 | 18192595 | 18194983 |
| chr9 | 11408084 | 11411222 |
| chr1 | 215874772 | 215876908 |
| chr2 | 194344365 | 194346478 |
| chr12 | 65302110 | 65310107 |
| chr6 | 103022360 | 103024506 |
| chr3 | 608710 | 610766 |
| chr10 | 108197989 | 108200749 |
| chr12 | 7194648 | 7199680 |
| chr10 | 41851982 | 41857227 |
| chr7 | 85285856 | 85289945 |
| chr12 | 26032396 | 26037034 |
| chr14 | 102208398 | 102211513 |
| chr1 | 102451912 | 102454728 |
| chr5 | 27672620 | 27674651 |
| chr5 | 171201987 | 171204301 |
| chr2 | 66676814 | 66679860 |
| chr15 | 20191308 | 20195192 |
| chr1 | 102876335 | 102882129 |
| chr14 | 85408629 | 85413528 |
| chr22 | 20274326 | 20276783 |
| chr22 | 15628687 | 15634003 |
| chr12 | 74918606 | 74921567 |
| chr21 | 10644415 | 10647895 |
| chr1 | 148679412 | 148681804 |
| chr7 | 49267232 | 49269530 |
| chr4 | 90308034 | 90310041 |

| | | |
|---|---|---|
| chr2 | 67275387 | 67277976 |
| chr10 | 110730953 | 110734768 |
| chr1 | 103552626 | 103555911 |
| chr1 | 99685694 | 99690397 |
| chr7 | 118417082 | 118419757 |
| chr10 | 65447774 | 65452985 |
| chr1 | 194819591 | 194822484 |
| chr6 | 80038663 | 80043201 |
| chr22 | 10751294 | 10756438 |
| chr6 | 75822707 | 75824832 |
| chr6 | 56055974 | 56058520 |
| chr5 | 178749411 | 178751498 |
| chr4 | 27939242 | 27941745 |
| chr22 | 11050051 | 11053114 |
| chr10 | 66925149 | 66928687 |
| chr2 | 182979723 | 182984691 |
| chr8 | 100163343 | 100166210 |
| chr2 | 165292700 | 165294764 |
| chr1 | 109094619 | 109097383 |
| chr8 | 112287234 | 112292045 |
| chr1 | 215668117 | 215672571 |
| chr2 | 83827195 | 83832283 |
| chr6 | 77024000 | 77027376 |
| chr14 | 37368042 | 37373416 |
| chr12 | 127490238 | 127492656 |
| chr18 | 60463154 | 60465190 |
| chr14 | 27549988 | 27554515 |
| chr1 | 198897394 | 198910956 |
| chr14 | 30914569 | 30918490 |
| chr7 | 77878537 | 77881579 |
| chr6 | 72817897 | 72822333 |
| chr4 | 174157595 | 174160490 |
| chr5_GL949742v1_a | 22066 | 24863 |
| chr15 | 55407053 | 55410502 |
| chr1 | 186091651 | 186095774 |
| chr8 | 64295244 | 64298188 |
| chr15 | 57225981 | 57228621 |
| chr4 | 177614705 | 177617524 |
| chr7 | 83160940 | 83164809 |
| chr7 | 90596416 | 90600526 |
| chr2 | 36544513 | 36547870 |
| chr3 | 23803157 | 23806028 |
| chr10 | 51619323 | 51622339 |
| chr8 | 76118267 | 76120749 |
| chr1 | 186173385 | 186175719 |
| chr6 | 95230718 | 95234958 |
| chr14 | 33510781 | 33515231 |
| chr6 | 26170367 | 26172501 |
| chr2 | 188538608 | 188541509 |

| | | |
|---|---|---|
| chr15 | 94619521 | 94622014 |
| chr7 | 90617882 | 90620547 |
| chr7 | 78363721 | 78370324 |
| chr7 | 90708930 | 90711387 |
| chr16_KI270853v1_ | 2050161 | 2056370 |
| chr8 | 39424611 | 39427211 |
| chr16 | 77417086 | 77419253 |
| chr2 | 159383711 | 159386701 |
| chr2 | 48510884 | 48514519 |
| chr1 | 198894149 | 198897166 |
| chr21 | 16069354 | 16076151 |
| chr1 | 172275048 | 172279937 |
| chr12 | 30668699 | 30671278 |
| chr13 | 71474785 | 71479807 |
| chr2 | 215420693 | 215424007 |
| chr14 | 25811099 | 25814362 |
| chr7 | 6254477 | 6257682 |
| chr11 | 103452979 | 103456287 |
| chr18 | 3446525 | 3450244 |
| chr8 | 112950743 | 112954152 |
| chr12 | 84717091 | 84721545 |
| chr2 | 226785968 | 226790068 |
| chr17 | 18603208 | 18608356 |
| chr2 | 57472136 | 57474766 |
| chr8 | 82460462 | 82464300 |
| chr14 | 37253182 | 37258488 |
| chr2 | 91719317 | 91722299 |
| chr7 | 7806276 | 7811645 |
| chr7 | 82233257 | 82237985 |
| chr11 | 31966753 | 31969234 |
| chr2 | 40628112 | 40632990 |
| chr5 | 93821477 | 93824781 |
| chr12 | 85031985 | 85037162 |
| chr7 | 97026316 | 97030823 |
| chr6 | 73806660 | 73809594 |
| chr14 | 38468842 | 38473740 |
| chr1 | 218346667 | 218351894 |
| chr10 | 82139830 | 82142898 |
| chr11 | 32891744 | 32895253 |
| chr11 | 89978518 | 89983327 |
| chr21 | 9002003 | 9006902 |
| chr1 | 192903156 | 192905957 |
| chr6 | 55977905 | 55984067 |
| chr13 | 71524764 | 71529625 |
| chr7 | 82833224 | 82842323 |
| chr2 | 186629653 | 186636626 |
| chr3 | 109946904 | 109950068 |
| chr14 | 35805705 | 35810377 |
| chr8 | 90890325 | 90892777 |

| | | |
|---|---|---|
| chr10 | 87006792 | 87010635 |
| chr13 | 71828109 | 71833223 |
| chr7 | 83749192 | 83752999 |
| chr7 | 84057727 | 84062825 |
| chr14 | 41920270 | 41928343 |
| chr14 | 36937783 | 36942739 |
| chr14 | 60748718 | 60752743 |
| chr7 | 79333423 | 79339530 |
| chr10 | 101006295 | 101009380 |
| chr7 | 80585436 | 80590123 |
| chr2 | 66769376 | 66777031 |
| chr13 | 71620827 | 71626928 |
| chr20 | 53655445 | 53661934 |
| chr1 | 98004621 | 98009024 |
| chr4 | 115231648 | 115242385 |
| chr17 | 76732213 | 76736601 |
| chr2 | 73255094 | 73258294 |
| chr4 | 175803309 | 175807308 |
| chr14 | 33098317 | 33101921 |
| chr1 | 143209041 | 143228724 |
| chr2 | 51655286 | 51659621 |
| chr1 | 112393037 | 112397816 |
| chr2 | 68806916 | 68810361 |
| chr2 | 59201186 | 59204834 |
| chr3 | 89110053 | 89112916 |
| chr14 | 29722798 | 29726950 |
| chr1 | 189954836 | 189960975 |
| chr2 | 62703034 | 62708532 |
| chr19 | 44888182 | 44891596 |
| chr2 | 226476552 | 226481072 |
| chr14 | 25884680 | 25890129 |
| chr10 | 56038065 | 56042200 |
| chr12 | 78989863 | 78994608 |
| chr6 | 72310133 | 72313236 |
| chr5 | 141039970 | 141043726 |
| chr14 | 103122470 | 103125062 |
| chr2 | 88764812 | 88768861 |
| chr11 | 23604420 | 23606995 |
| chr1 | 218871399 | 218875994 |
| chr2 | 189201911 | 189207752 |
| chr7 | 62333582 | 62337067 |
| chr22 | 12598774 | 12603993 |
| chr11 | 83905288 | 83908453 |
| chr7 | 83007684 | 83014471 |
| chr2 | 53198388 | 53201314 |
| chr7 | 10041617 | 10044748 |
| chr15 | 50351290 | 50355473 |
| chr14 | 40398490 | 40401779 |
| chr14 | 48810119 | 48813419 |

| | | |
|---|---|---|
| chr14 | 19370572 | 19374139 |
| chr5 | 39739980 | 39743921 |
| chr12 | 74613366 | 74619348 |
| chr16 | 70232129 | 70235593 |
| chr5 | 27400208 | 27403577 |
| chr14 | 88424936 | 88428454 |
| chr2 | 63840642 | 63844650 |
| chr6 | 55841981 | 55848211 |
| chr10 | 120939169 | 120943502 |
| chr1 | 214876385 | 214878983 |
| chr2 | 63048801 | 63052580 |
| chr16 | 76234085 | 76236945 |
| chr5 | 28105410 | 28109031 |
| chrUn_GL000195v1 | 88656 | 94751 |
| chr2 | 36848587 | 36851198 |
| chr6 | 74393849 | 74397148 |
| chr2 | 177223382 | 177229761 |
| chr7 | 84236213 | 84240340 |
| chr12 | 86386619 | 86388904 |
| chr2 | 188585932 | 188589840 |
| chr2 | 181673622 | 181676742 |
| chr14 | 86256151 | 86258935 |
| chr1 | 191458344 | 191462761 |
| chr1 | 219173114 | 219176614 |
| chr7 | 102467305 | 102469936 |
| chr1 | 98403521 | 98407743 |
| chr10 | 42317782 | 42322078 |
| chr4 | 157221669 | 157228734 |
| chr2 | 189125480 | 189132629 |
| chr7 | 83072708 | 83078196 |
| chr12 | 84908280 | 84911872 |
| chr2 | 195420679 | 195424112 |
| chr2 | 209043747 | 209046845 |
| chr12 | 78144054 | 78150909 |
| chr3 | 111069582 | 111072048 |
| chr17 | 14611964 | 14614652 |
| chr12 | 21646646 | 21649778 |
| chr2 | 36882423 | 36885602 |
| chr6 | 98053779 | 98058581 |
| chr2 | 166251639 | 166254805 |
| chr7 | 79468757 | 79472880 |
| chr11 | 23585313 | 23591984 |
| chr14 | 52768438 | 52773126 |
| chr4 | 34407045 | 34409546 |
| chr12 | 65325905 | 65329277 |
| chr10 | 116383990 | 116388395 |
| chr2 | 102515960 | 102520922 |
| chr20 | 31222987 | 31245863 |
| chr5_KI270897v1_a | 836829 | 841733 |

| | | |
|---|---|---|
| chr14 | 40884856 | 40889998 |
| chr14 | 38513716 | 38516418 |
| chr16_KI270728v1_ | 989007 | 993310 |
| chr14 | 26578265 | 26587988 |
| chr7 | 82684547 | 82688887 |
| chr20 | 38524354 | 38527579 |
| chr14 | 100583760 | 100589744 |
| chr2 | 51626175 | 51630164 |
| chr16 | 76446963 | 76450598 |
| chr16 | 33506477 | 33510323 |
| chr12 | 70275725 | 70278753 |
| chr14 | 44659767 | 44662876 |
| chr14 | 100978163 | 100985090 |
| chr7 | 79394208 | 79400710 |
| chr3 | 97435851 | 97438957 |
| chr6 | 56167553 | 56172157 |
| chr1 | 197539298 | 197542620 |
| chr13 | 63090556 | 63093684 |
| chr4 | 175868329 | 175877221 |
| chr6 | 78459408 | 78462885 |
| chr2 | 56012169 | 56016369 |
| chr12 | 24469959 | 24474641 |
| chr4 | 115107556 | 115117010 |
| chr9 | 73177518 | 73180874 |
| chr5 | 135755138 | 135757853 |
| chr11 | 31235602 | 31240983 |
| chr7 | 93376216 | 93380515 |
| chr5 | 27563875 | 27568736 |
| chr8 | 39374211 | 39382155 |
| chr11 | 90220486 | 90223759 |
| chr7 | 84091070 | 84097295 |
| chr1 | 103080645 | 103084223 |
| chr12 | 53479771 | 53483944 |
| chr1 | 86403332 | 86406706 |
| chr1 | 86497902 | 86501232 |
| chr13 | 71756406 | 71761788 |
| chr7 | 84283100 | 84287973 |
| chr14 | 32198883 | 32203464 |
| chr4 | 32464269 | 32467472 |
| chr6 | 13120328 | 13124762 |
| chr11 | 31710637 | 31714698 |
| chr3 | 95715047 | 95722674 |
| chr12 | 85029148 | 85031456 |
| chr1 | 75881794 | 75885388 |
| chr7 | 84329106 | 84335457 |
| chr6 | 84245356 | 84251219 |
| chr7 | 82795058 | 82799533 |
| chr6 | 80268807 | 80273328 |
| chr1 | 107944865 | 107948511 |

| | | |
|---|---|---|
| chr2 | 172045719 | 172051381 |
| chr14 | 21381497 | 21385621 |
| chr15_KI270851v1_ | 51763 | 55442 |
| chr14 | 19284186 | 19289508 |
| chr3 | 174461448 | 174467495 |
| chr2 | 189091457 | 189097589 |
| chr6 | 48713499 | 48717264 |
| chrUn_GL000195v1 | 113579 | 118058 |
| chr5 | 41939266 | 41942699 |
| chr14 | 32895766 | 32899062 |
| chr1 | 85018130 | 85022285 |
| chr2 | 58568352 | 58571278 |
| chr8 | 124480941 | 124486086 |
| chrX | 56776923 | 56781396 |
| chr1 | 78001513 | 78007031 |
| chr2 | 209678727 | 209682129 |
| chr1 | 97846411 | 97854457 |
| chr21 | 16535489 | 16542407 |
| chr1 | 148622718 | 148626595 |
| chr6 | 121891971 | 121898812 |
| chr6 | 125375225 | 125378630 |
| chr12 | 79775952 | 79779095 |
| chr1 | 103749397 | 103755838 |
| chr2 | 77305240 | 77308715 |
| chr12 | 77840591 | 77850344 |
| chr7 | 78830343 | 78832732 |
| chr4 | 29978215 | 29981322 |
| chr7 | 10254342 | 10260217 |
| chr7 | 62277442 | 62282242 |
| chr11 | 104246707 | 104250421 |
| chr5 | 108020911 | 108025956 |
| chr2 | 41128790 | 41134197 |
| chr5_KI270897v1_a | 870734 | 873934 |
| chr1 | 103070861 | 103076027 |
| chr10 | 62811863 | 62817538 |
| chr14 | 88611657 | 88615757 |
| chr22 | 22269767 | 22275619 |
| chr12 | 62389378 | 62394610 |
| chr2 | 188368921 | 188371563 |
| chr14 | 53806099 | 53809808 |
| chr14 | 42701664 | 42705151 |
| chr12 | 78860516 | 78871259 |
| chr13 | 105337153 | 105341217 |
| chr15 | 38352333 | 38356452 |
| chr20 | 30823299 | 30826559 |
| chr12 | 96486971 | 96490870 |
| chr10 | 47646573 | 47649829 |
| chr10 | 38495905 | 38527046 |
| chr10 | 83537307 | 83540731 |

| | | |
|---|---|---|
| chr17_GL383563v3_ | 30978 | 34473 |
| chr2 | 102945022 | 102949339 |
| chr4 | 35147326 | 35152273 |
| chr6 | 74317725 | 74322008 |
| chr11 | 114602577 | 114606348 |
| chr10 | 76541990 | 76545370 |
| chr9 | 11651800 | 11654757 |
| chr14 | 40487283 | 40491920 |
| chr1 | 149608817 | 149612260 |
| chr10 | 101063654 | 101068132 |
| chr7 | 84365246 | 84368118 |
| chr2 | 194568336 | 194572168 |
| chr7 | 82102950 | 82107743 |
| chr20 | 10013215 | 10016479 |
| chr1 | 103192472 | 103197175 |
| chr10 | 101393762 | 101398019 |
| chr12 | 39599486 | 39606972 |
| chr7 | 84043399 | 84048811 |
| chr12 | 82324826 | 82330063 |
| chr7 | 83476808 | 83481231 |
| chr14 | 91969546 | 91974071 |
| chr10 | 58286922 | 58292902 |
| chr10 | 42296224 | 42311989 |
| chr7 | 91379037 | 91384844 |
| chr7 | 79637056 | 79641605 |
| chr14 | 62846073 | 62852460 |
| chr14 | 36857456 | 36861276 |
| chr14 | 47929189 | 47933575 |
| chr13 | 66988784 | 66997299 |
| chr9 | 64497439 | 64501963 |
| chr2 | 178450783 | 178453756 |
| chr2 | 59343770 | 59348246 |
| chr2 | 66565658 | 66575068 |
| chr12 | 79015196 | 79021231 |
| chr1 | 189106152 | 189110335 |
| chr17 | 74775334 | 74778783 |
| chr10 | 65224028 | 65228441 |
| chr13 | 71700947 | 71705279 |
| chr22 | 20298291 | 20301280 |
| chr16 | 69642807 | 69647539 |
| chr1 | 102968690 | 102980474 |
| chr6 | 72645790 | 72648531 |
| chr11 | 22160452 | 22163532 |
| chr9 | 63388279 | 63392538 |
| chr14 | 36313168 | 36317369 |
| chr7 | 83683125 | 83685990 |
| chr2 | 212387459 | 212392970 |
| chr2 | 189108652 | 189113573 |
| chr1 | 103412657 | 103421065 |

| | | |
|---|---|---|
| chr13 | 63059854 | 63068715 |
| chr1 | 86720671 | 86724149 |
| chr7 | 8438621 | 8442076 |
| chr8 | 79365806 | 79368171 |
| chr14 | 105470135 | 105472700 |
| chr1 | 103035094 | 103037991 |
| chr20 | 43458322 | 43463447 |
| chr3 | 89164359 | 89169235 |
| chr15_KI270850v1_ | 226514 | 231416 |
| chr10 | 65490713 | 65498449 |
| chr7 | 80602969 | 80608049 |
| chr2 | 189148700 | 189154751 |
| chr14 | 84534474 | 84537283 |
| chr4 | 40054535 | 40059748 |
| chr17 | 26800674 | 26814077 |
| chr11 | 79420757 | 79423557 |
| chr14 | 99469626 | 99477860 |
| chr2 | 187495478 | 187500960 |
| chr2 | 210586444 | 210591947 |
| chr9 | 13277699 | 13283162 |
| chr14 | 39060875 | 39064567 |
| chr7 | 57676662 | 57682250 |
| chr10 | 37234572 | 37241057 |
| chr15_KI270851v1_ | 32688 | 35577 |
| chr7 | 94655289 | 94664232 |
| chr2 | 74533275 | 74537176 |
| chr10 | 55833792 | 55837252 |
| chr20 | 30744370 | 30747235 |
| chr4 | 166003576 | 166007393 |
| chr1 | 98448728 | 98455218 |
| chr7 | 88954229 | 88958790 |
| chr7 | 94092940 | 94096744 |
| chr7 | 94180151 | 94182804 |
| chr11 | 23579632 | 23585048 |
| chr9 | 62810827 | 62814341 |
| chr2 | 63483811 | 63489558 |
| chr4 | 33365976 | 33368869 |
